# The Genetic Origin of the Indo-Europeans

**DOI:** 10.1101/2024.04.17.589597

**Authors:** Iosif Lazaridis, Nick Patterson, David Anthony, Leonid Vyazov, Romain Fournier, Harald Ringbauer, Iñigo Olalde, Alexander A. Khokhlov, Egor P. Kitov, Natalia I. Shishlina, Sorin C. Ailincăi, Danila S. Agapov, Sergey A. Agapov, Elena Batieva, Baitanayev Bauyrzhan, Zsolt Bereczki, Alexandra Buzhilova, Piya Changmai, Andrey A. Chizhevsky, Ion Ciobanu, Mihai Constantinescu, Marietta Csányi, János Dani, Peter K. Dashkovskiy, Sándor Évinger, Anatoly Faifert, Pavel N. Flegontov, Alin Frînculeasa, Mădălina N. Frînculeasa, Tamás Hajdu, Tom Higham, Paweł Jarosz, Pavol Jelínek, Valeri I. Khartanovich, Eduard N. Kirginekov, Viktória Kiss, Alexandera Kitova, Alexeiy V. Kiyashko, Jovan Koledin, Arkady Korolev, Pavel Kosintsev, Gabriella Kulcsár, Pavel Kuznetsov, Rabadan Magomedov, Mamedov Aslan Malikovich, Eszter Melis, Vyacheslav Moiseyev, Erika Molnár, Janet Monge, Octav Negrea, Nadezhda A. Nikolaeva, Mario Novak, Maria Ochir-Goryaeva, György Pálfi, Sergiu Popovici, Marina P. Rykun, Tatyana M. Savenkova, Vladimir P. Semibratov, Nikolai N. Seregin, Alena Šefčáková, Mussayeva Raikhan Serikovna, Irina Shingiray, Vladimir N. Shirokov, Angela Simalcsik, Kendra Sirak, Konstantin N. Solodovnikov, Judit Tárnoki, Alexey A. Tishkin, Viktov Trifonov, Sergey Vasilyev, Ali Akbari, Esther S. Brielle, Kim Callan, Francesca Candilio, Olivia Cheronet, Elizabeth Curtis, Olga Flegontova, Lora Iliev, Aisling Kearns, Denise Keating, Ann Marie Lawson, Matthew Mah, Adam Micco, Megan Michel, Jonas Oppenheimer, Lijun Qiu, J. Noah Workman, Fatma Zalzala, Anna Szécsényi-Nagy, Pier Francesco Palamara, Swapan Mallick, Nadin Rohland, Ron Pinhasi, David Reich

## Abstract

The Yamnaya archaeological complex appeared around 3300BCE across the steppes north of the Black and Caspian Seas, and by 3000BCE reached its maximal extent from Hungary in the west to Kazakhstan in the east. To localize the ancestral and geographical origins of the Yamnaya among the diverse Eneolithic people that preceded them, we studied ancient DNA data from 428 individuals of which 299 are reported for the first time, demonstrating three previously unknown Eneolithic genetic clines. First, a “Caucasus-Lower Volga” (CLV) Cline suffused with Caucasus hunter-gatherer (CHG) ancestry extended between a Caucasus Neolithic southern end in Neolithic Armenia, and a steppe northern end in Berezhnovka in the Lower Volga. Bidirectional gene flow across the CLV cline created admixed intermediate populations in both the north Caucasus, such as the Maikop people, and on the steppe, such as those at the site of Remontnoye north of the Manych depression. CLV people also helped form two major riverine clines by admixing with distinct groups of European hunter-gatherers. A “Volga Cline” was formed as Lower Volga people mixed with upriver populations that had more Eastern hunter-gatherer (EHG) ancestry, creating genetically hyper-variable populations as at Khvalynsk in the Middle Volga. A “Dnipro Cline” was formed as CLV people bearing both Caucasus Neolithic and Lower Volga ancestry moved west and acquired Ukraine Neolithic hunter-gatherer (UNHG) ancestry to establish the population of the Serednii Stih culture from which the direct ancestors of the Yamnaya themselves were formed around 4000BCE. This population grew rapidly after 3750-3350BCE, precipitating the expansion of people of the Yamnaya culture who totally displaced previous groups on the Volga and further east, while admixing with more sedentary groups in the west. CLV cline people with Lower Volga ancestry contributed four fifths of the ancestry of the Yamnaya, but also, entering Anatolia from the east, contributed at least a tenth of the ancestry of Bronze Age Central Anatolians, where the Hittite language, related to the Indo-European languages spread by the Yamnaya, was spoken. We thus propose that the final unity of the speakers of the “Proto-Indo-Anatolian” ancestral language of both Anatolian and Indo-European languages can be traced to CLV cline people sometime between 4400-4000 BCE.

Summary Figure:
The origin of Indo-Anatolian and Indo-European languages.
Genetic reconstruction of the ancestry of Pontic-Caspian steppe and West Asian populations points to the North Caucasus-Lower Volga area as the homeland of Indo-Anatolian languages and to the Serednii Stih archaeological culture of the Dnipro-Don area as the homeland of Indo-European languages. The Caucasus-Lower Volga people had diverse distal roots, estimated using the *qpAdm* software on the left barplot, as Caucasus hunter-gatherer (purple), Central Asian (red), Eastern hunter-gatherer (pink), and West Asian Neolithic (green). Caucasus-Lower Volga expansions, estimated using *qpAdm* on the right barplot as disseminated Caucasus Neolithic (blue)-Lower Volga Eneolithic (orange) proximal ancestries, mixing with the inhabitants of the North Pontic region (yellow), Volga region (yellow), and West Asia (green).

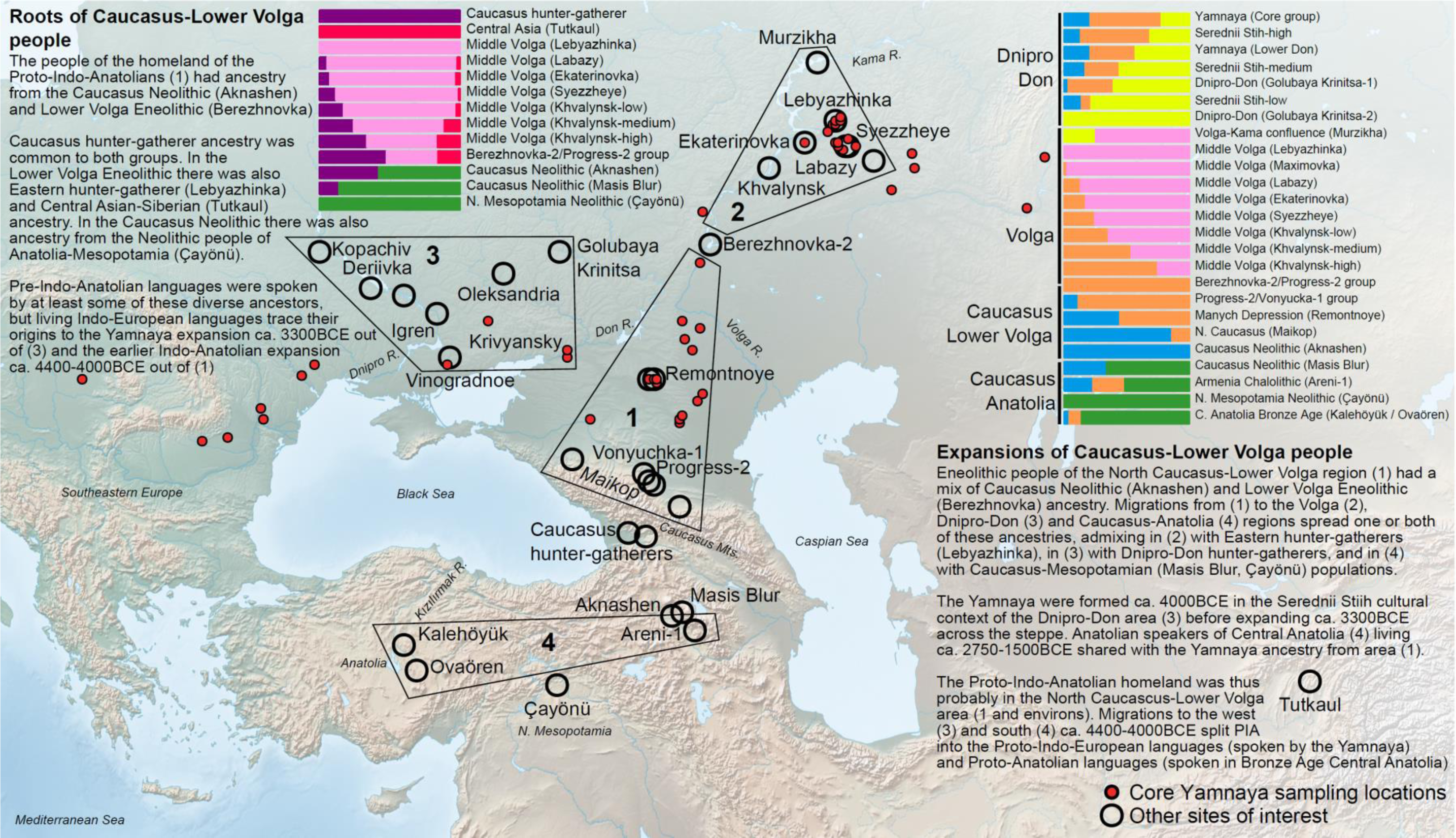

## Introduction

Between 3300-1500 BCE, people of the Yamnaya archaeological complex and their descendants, in subsequent waves of migration, spread over large parts of Eurasia, contributing to the ancestry of people of Europe, Central and South Asia, Siberia, and the Caucasus. The spread of Indo-European language and culture^1–7^ transformed all these regions. Despite the centrality of the Yamnaya expansion to the human story of Bronze Age Eurasia, their ancestral origins are poorly understood. A first challenge has been the sparse sampling of the Yamnaya themselves across their enormous geographic distribution. The remarkable long-range mobility of the Yamnaya, quickly spreading over a vast region, adds further difficulties to tracing, from radiocarbon dating, the origins of their material culture and associated genetic profile. Nor can these origins be traced to the numerous earlier Eneolithic cultures that preceded the Yamnaya, and among whom their ancestors must be sought, as these have been sampled even more poorly and unsystematically.

The first formal study of the origins of the Yamnaya identified two disparate sources of ancestry: a northern, “Eastern Hunter-Gatherer” (EHG) source from far eastern Europe, and a southern, West Asian source related to present-day Armenians.^2^ The latter source was revealed, by ancient DNA, to be related to some of the region’s earliest inhabitants: Paleolithic-Mesolithic “Caucasus Hunter-Gatherers” (CHG) of Georgia,^8^ and Neolithic people of the Zagros^9^ and South Caucasus.^6,10,11^ Additional discoveries further complicated the stories of both the northern and southern ancestors of the Yamnaya. First, it was noted that both CHG and EHG were part of an interaction sphere across the boundary between West Asia and eastern Europe,^9^ suggesting the existence of intermediate populations and raising the question of when and where these came together to form the Eneolithic antecessors of the Yamnaya. Second, it was recognized that the steppe itself was an admixture zone of EHG with “Western Hunter-Gatherers” (WHG^12^).

Mesolithic hunter-gatherers from Ukraine were succeeded by more WHG-admixed Neolithic hunter-gatherers in the Dnipro valley,^13^ representing a local reshuffling within the European portion of a ∼7,000km-long trans-Eurasian cline of boreal hunter-gatherers.^14^ What was the relative contribution of the EHG (who were present in the Volga River at Lebyazhinka^2^ ca. 5660-5535 BCE) and these more western Ukraine Neolithic hunter-gatherers (UNHG) of the Dnipro to later populations? Third, it was discovered that the Yamnaya had not only CHG-related, but also Anatolian Neolithic ancestry, absent in the early known steppe inhabitants, and derived from European farmer neighbors west of the steppe^5^. This ancestry was later shown to be of rather Anatolian-Levantine-Mesopotamian origin, and to be mediated not from Europe but from the Caucasus neighbors south of the steppe.^6^ Such ancestry must have been added following the expansion of Neolithic farmers into the Caucasus, introduced thence into the steppe as a later exogenous element, distinct from the earlier CHG-related one. Finally, it was recognized that European steppe populations were formed not only by northern-southern admixture, but included, in at least some Eneolithic and Bronze Age people of the North Caucasus, contributions related to Siberians from further east.^5^ What was the extent of the spread of this eastern ancestry and did the Yamnaya themselves possess it?

Here we present a unified population genetic analysis of 372 newly reported individuals dating from 6400-2000 BCE, as well as increased quality data for 61 individuals. The present study serves as the formal technical report for 299 of the newly reported individuals and 55 of the individuals with increased quality data; more than 80% of the individuals are from Russia, but the dataset is also significant in including dozens of individuals from westward expansion of Steppe cultures along the Danube (Supplementary Information, section 1, Online Table 1). Technical details of the 803 ancient DNA libraries that are the basis for the newly reported data (and an additional 195 libraries that failed our screening) are presented in Online Table 2, while details of 198 newly generated radiocarbon dates on these individuals are presented in Online Table 3. A parallel study^15^ presents a combined archaeological and genetic analysis of population transformations in the North Pontic Region (Ukraine and Moldova) and serves as the formal report for the data from the other 73 of the newly analyzed individuals and the other 5 individuals with increased quality data, with both studies co-analyzing the full dataset. We grouped individuals into analysis labels based on geographical and temporal information, archaeological context, and genetic clustering (Online Table 4 lists all individuals used for analyses, with their labels). The potential of the combined dataset for shedding light on this period can be appreciated from the fact that it adds 79 analyzed Eneolithic people from the steppe and its environs (from Russia or Ukraine, west of 60E longitude and south of 60N latitude, between 5000-3500BCE) to 82 published^5,7,13,15–20^ and a total of 286 Yamnaya/Afanasievo individuals compared to 75 in the literature.^2,4–6,13,21–29^

### Discovery of three pre-Bronze Age genetic clines that collapsed after Yamnaya expansion

Principal Component Analysis (PCA) of ancient individuals from the Pontic-Caspian steppe and adjacent areas of Southeastern Europe, the Caucasus and West Asia reveals that most of the Eneolithic people of the steppe as well as the later Bronze Age Yamnaya fall on non-overlapping genetic gradients (Figure 1). Visual impressions from a two-dimensional PCA must be evaluated cautiously, as populations at intermediate PCA positions, may not, in fact, be mixtures of more extreme ones, and the plot may suggest alternative ways of modeling each population of interest. For example, PC1 correlates (from left to right) to the differentiation between inland West Asians (Caucasus and Iran) to East Mediterranean (Anatolian-European) populations^10^, but also to the differentiation between Siberians and European hunter-gatherers^14^. On the other hand, PC2 differentiates between Neolithic and earlier populations from northern Eurasia (top; including Europe and Siberia) and West Asia (bottom: Anatolia-Mesopotamia-Caucasus-Iran). The Eneolithic and Bronze Age populations occupy the middle of the PCA: how did the earlier groups surrounding them across these orthogonal directions combine to form them?

**Figure 1:**
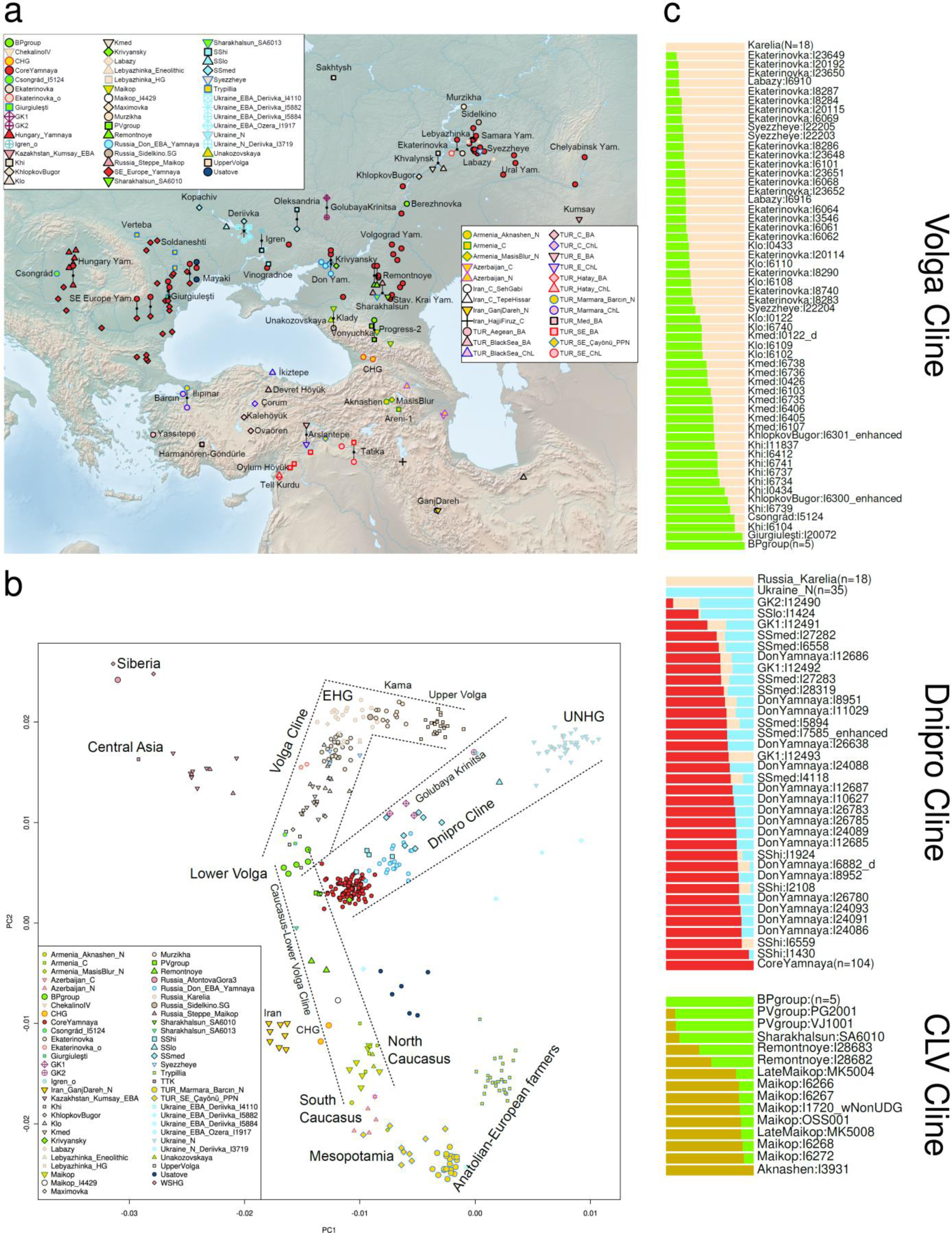
Three Eneolithic clines and their neighbors in space and time. (a) Map with analyzed sites. (b) PCA analysis using axes formed by a set of ancient West European hunter-gatherer (WHG), Siberian, West Asian, and European farmer populations. Selected individuals relevant to this study are projected^37^ (Methods). (c) qpAdm models fitted on individuals of the populations of the three clines. The Volga Cline is generated by admixture between Lower Volga (BPgroup) people with upriver Eastern hunter-gatherers (EHG). Populations of the Dnipro Cline have UNHG or UNHG+EHG admixture relative to the Core Yamnaya (the hunter-gatherer source along this cline is significantly variable). The Caucasus-Lower Volga Cline is generated by admixture between lower Volga people with those from the Neolithic Caucasus (Aknashen-related).

To answer these questions in a statistically rigorous way, we implemented a new model competition tournament framework around qpWave/qpAdm methods^2,30^ to fit and distinguish among alternative models (Methods; Supplementary Information, section 2). Briefly summarized, the idea of this methodology is that an admixture model *X* that includes a set of sources describes the admixture history of a target population *T* well if it: (i) reconstructs the shared genetic drift of *T* with both distant outgroup populations and the sources of alternative competing models, but also (ii) renders these competing models infeasible by showing that they cannot model this shared drift with the sources of *X*. In our framework, models are evaluated against a conservative set of distant outgroups as an initial filtering step; if they fit poorly, they are rejected; if not, they are further evaluated by comparing them against each other in symmetrical fashion (all-against-all) to identify a smaller set of promising models.

With this note of caution, we observe that in the PCA in the Eneolithic-Bronze Age steppe there are three clines (geographically denoted as “Volga”, “Dnipro”, and “Caucasus-Lower Volga”), which diverge, in PCA space, from an area that includes populations enclosed by the Lower Don (at the site of Krivyansky), Lower Volga (at Berezhnovka-2), and north Caucasus mountains (at Progress-2, Vonyuchka-1, and Sharakhalsun^5^). From these similar beginnings the three clines extend outward into distinct directions corresponding to their geographical neighbors: both towards the EHG and UNHG representing the pre-Eneolithic people that lived in the Volga-Don-Dnipro area of eastern Europe, and towards the CHG and Caucasus Neolithic representing the pre-Eneolithic people that lived in the Caucasus and West Asia. In what follows, we introduce the key populations of each of the three clines and show how these can be modeled in terms of proximate sources. We also infer the ancestry origins of the people of the three clines to discover what is shared among all of them and unique to each of them.

#### Volga Cline

The “Volga Cline” consists of sites on waterways that drain into the Caspian Sea and is suggestive of a zone of ongoing human contact within its region. The Eneolithic individuals fall at positions that correlate extraordinarily well to their position on the Volga River as one moves downstream: the Volosovo-attributed Sakhtysh (in the Upper Volga) and Murzikha (near the Kama-Volga confluence)^14^ constitute the upriver portion of the cline, situated in PCA space between EHG and UNHG. The Volga Cline then distinctly “bends” in PCA space and the knee of the cline is occupied by EHG groups, including those sampled in the northwest of Russia in Karelia^2,19^ and those of the Middle Volga, suggesting that this widely dispersed set of hunter-gatherers, which has also been called the Sidelkino Cluster based on its oldest representative^19,22^ were the major population of much of eastern Europe. Past the knee, in the downriver portion of the cline, the hunter-gatherer affinity decreases starting at the Middle Volga: Labazy, Lebyazhinka, Ekaterinovka, Syezzheye, then Khvalynsk (4500-4350 BCE) and Khlopkov Bugor, finally reaching the Lower Volga at Berezhnovka (4450-3960 BCE) (Fig. 1a). The decrease of hunter-gatherer affinity is counterbalanced by increased affinity towards populations of the Caucasus, suggesting that it is generated by an unsampled CHG-related source—that existed somewhere between Georgia (where the known CHG individuals were sampled^8^) and the Lower Volga— interacting with the northern EHG natives. Archaeological correlates for such south-north interactions do exist, and begin with the expansion of the Seroglazovo forager culture around the Lower Volga estuary ∼6200 BCE, with some ceramic and lithic typological parallels with Caucasus cultures, and continue to the unsampled North Caucasus Neolithic cemetery dated ∼4800 BCE near Nalchik.^31,32^

At the end of the cline, the four individuals from the newly reported Lower Volga site of Berezhnovka-2 can be grouped with the PG2004 individual of the Progress-2^5^ site in the north Caucasus into a “Berezhnovka-2-Progress-2 cluster” (abbreviated to “BPgroup”). This proves that the CHG-related ancestry found at Progress-2 extended well into the steppe in the Lower Volga. The second individual from Progress-2 (PG2001) is grouped with another north Caucasus individual from Vonyuchka-1^5^ into a related “Progress-2-Vonyuchka-1 cluster” (abbreviated as “PVgroup”). PVgroup and BPgroup are distinct (p=0.0006), but their genetic differentiation was small in magnitude (F_ST_=-0.002±0.002; Extended Data Table 1) suggesting movement between the north Caucasus piedmont and Lower Volga sites. The two locations also shared a distinctive burial pose on the back with raised knees, later typical of Yamnaya and currently dated earliest in the Samara region at Lebyazhinka-5 and in a few graves at Ekaterinovka dated before 4500 BCE. It is clear from the PCA (Fig. 1b) that BPgroup differs from PVgroup in that the former is shifted towards the Afontova Gora-3 Upper Paleolithic individual from Siberia,^33^ West Siberian hunter-gatherers,^4^ and Central Asians such as a 7,500-year old Neolithic individual from Tutkaul (TTK) in Tajikistan.^19^ We will see below that Siberian/Central Asian ancestry was one of the constitutive elements of the Lower Volga-North Caucasus Eneolithic population represented by the two groups.

A natural interpretation of the Volga cline is that upriver EHG-related ancestors and downriver Berezhnovka-related ones came together to form communities along the length of the river, resulting in a highly variable set of sampled individuals along the genetic gradient. While the origin of the upriver EHG ancestry is clear, as it has antecedents in eastern Europe for thousands of years,^19^ that of the downriver Berezhnovka group is less so, as (i) no earlier individuals from the Lower Volga have been sequenced, (ii) the genetic position of the Berezhnovka people is distinct from that of all preceding groups, and (iii) the BPgroup cannot be modeled as a clade with any contemporary or earlier groups (p<0.001). Whatever the origin of BPgroup, a point to which we will return below, we can use it as a proximate source and test Volga cline populations and individuals for consistency with a history of mixture of people related to the BPgroup and EHG (using Karelia^2,19^ as an EHG source well outside the Volga area and unlikely to be part of the riverine mating network), as suggested by the PCA. Seven Volga cline populations fit this model (p-values of 0.04 to 0.72) with the only consistently poor fits for Upper Volga, Murzikha, Maximovka, and “Klo” (the Khvalynsk individuals with low Berezhnovka relatedness) (p-values of 1e-66 to 0.006). Three of these populations (other than Klo which we discuss below) are arrayed in the upriver portion of the Volga cline, before its PCA “bend” (from EHG towards the UNHG). Individuals along the downriver portion of the cline can be well-modeled with only the two sources (BPgroup and EHG) (Fig. 1c).

People on the Volga Cline buried at the Ekaterinovka cemetery likely died between 5050-4450 BCE (based on radiocarbon dates on three herbivore bones including a domesticated sheep in the graves of individuals we analyzed that are not expected to be affected by marine reservoir effects; Online Table 1). The Ekaterinovka people were already in the process of mixing with BPgroup-related people from the Lower Volga (24.3±1.3% on average). This contrasts to the earlier hunter-gatherer from Lebyazhinka, who had the lowest estimate of Berezhnovka ancestry on the Volga Cline of only 7.9±3.6%, providing a baseline of this component prior to the Eneolithic and which can also be modeled with only EHG-related ancestry (p=0.21) while Ekaterinovka cannot (p=2e-4). Mixing intensified over time so that 100-200 years later at the site of Khvalynsk^34^ which is ∼120km from Ekaterinovka (date range of 4500-4350 BCE based on two herbivore bones in the graves of individuals we analyzed), we observe a continuous gradient of admixture which we divide for convenience into three groups: “Khavlynsk high (Khi)” (76.8±1.9% BPgroup), “Khvalynsk medium (Kmed)” (57.3±1.7% BPgroup), and “Khalynsk low (Klo)” (41.2±1.6% BPgroup). Individuals on the downriver portion of the Volga cline exhibited a range of Berezhnovka ancestry from ∼14-89% (Fig. 1c) and thus were not clearly dominated by either the old EHG ancestors of the region or the Lower Volga newcomers. Genetic differentiation between Lower Volga (BPgroup) and Ekaterinovka was strong (F_ST_=0.030±0.001; Extended Data Table 1) and quite probably reflected at least two different linguistic-cultural communities interacting with each other.

A genetically Volga Cline individual not from the Volga Basin is from Csongrád-Kettőshalom in Hungary, whose direct date is 4331-4073 cal BCE. This individual is estimated to have 87.9±3.5% of its ancestry from the BPgroup (Fig. 1c) comparable to the most extreme “Khvalynsk high” individuals. The Csongrád individual is one among a group of steppe-like graves that appeared in Southeastern Europe in the late 5^th^ millennium BCE including a cemetery at Giurgiuleşti,^35^ Moldova, from which one individual (I20072; 4330-4058BCE) is consistent with being a clade (p=0.90) with BPgroup, and another cemetery at Mayaky, Ukraine.^36^ Archaeological analysis has documented long-distance movement of Balkan copper to the Volga-Cline site of Khvalynsk,^34^ and the Csongrád and Mayaky individuals were plausibly part of the cultural exchange that mediated this process—a process our results show has no evidence of being contributed to genetically by people with ancestry typical of the Dnipro and Don basins. As we will now see, migrants with ancestry from the Lower Volga Eneolithic populations at the southern extreme of the Volga Cline did settle in the Dnipro area and generated the second major cline of the steppe.

#### (2) Dnipro Cline

The Dnipro Cline is formed at one end by Neolithic individuals living along the Dnipro River rapids whose union of calibrated radiocarbon dates is 6242-4542 BCE (UNHG), and at the other end by the Serednii Stih population represented by 13 individuals with good quality data whose union of radiocarbon date ranges uncorrected for freshwater reservoir effects are 4996-3372 BCE. The Dnipro Cline also includes the great majority of later Yamnaya individuals who expanded widely, most of whom are from a genetically homogeneous subset, and we used a large group of these individuals that have high quality data (n=104) to represent “Core Yamnaya” (Supplementary Information, section 2). Close to the Core Yamnaya in PCA are two Eneolithic groups: the Serednii Stih individual from Krivyansky in the Lower Don (4359-4251 BCE), and the PVgroup from the north Caucasus we discussed above as related to the Berezhnovka Lower Volga population. Nonetheless, the Core Yamnaya cannot be modeled as derived from either of these two earlier sources or indeed any other single source (p<1e-4). Their ancestry must have involved some admixture as their position along the highly variable Dnipro/Serednii Stih-associated cline also suggests. People from the Dnipro Cline as a whole are also fully distinct from those of the Volga Cline in PCA, and no pair of populations from the Volga and Dnipro clines form a genetic clade (p<1e-7). This distinctiveness spans a period of three millennia, beginning with earlier groups from Ukraine (UNHG), continuing with those of the Eneolithic Serednii Stih culture, and ending with the Yamnaya at the beginning of the Bronze Age, documenting the distinctiveness of the communities of these two great eastern European rivers and the relative lack of migration between them. A more geographically localized Yamnaya population of the Lower Don (n=23), many (n=17) of which are from the site of Krivyansky, bear no affinity to the Eneolithic individual from the area (Fig. 1). The Yamnaya can thus be traced neither to the north Caucasus (PVgroup), nor to the Lower Don (Krivyansky), nor to the Volga (BPgroup and the rest of the Volga cline). Yet, their position on the Dnipro cline, generated by populations of UNHG ancestry suggests that they emerged there, as a descendant community of people of the Serednii Stih culture.

The genetic heterogeneity of the Serednii Stih contrasts with the homogeneity of the Core Yamnaya (Fig. 1) which occupies one end of the Dnipro cline. The Core Yamnaya homogeneity is remarkable given that this cluster includes individuals sampled across 5,000 km from Hungary to southern Siberia, a vast slice of Eurasia across which the Yamnaya expanded but, for whatever reason, hardly admixed, at least initially, and at least for the elite subset of people afforded burial in kurgans, with any of the people that previously occupied it. Individuals of the Serednii Stih culture are arrayed along the Dnipro Cline with individuals of high or low Yamnaya affinity found at different sites. Closest to the Core Yamnaya genetically is a Serednii Stih individual from Vinogradnoe from the coast of the Azov Sea which we group with two other individuals from Oleksandria and one from Igren into an “SShi” cluster of greatest Yamnaya affinity. The sampled SShi group does not form a clade with the Core Yamnaya (p=2×10^-7^). A female from Kopachiv (I7585)^38^, represented by a long bone found loose in a Trypillia phase BI-II settlement, is part of a second “SSmed” cluster that is further along the Dnipro Cline; this group also includes three individuals from Oleksandria and three from Deriivka. The SShi and SSmed subsets are largely contiguous with each other, but individual I1424 from Moliukhiv Bugor (“SSlo”) is much further apart and close to the UNHG. The true variation within the Serednii Stih plausibly included individuals that fill gaps along the cline, e.g., between SSlo and SSmed, and even extended beyond the sampled variation, occupying the position of the Core Yamnaya itself. The Don Yamnaya largely overlap the Serednii Stih individuals, and the Don Yamnaya are discontinuous with the earlier Eneolithic individual from that location (p=7e-15). An interesting material correlate is seen in settlement continuity at stratified sites of the Konstantinovka culture on the Lower Don where the Don Yamnaya continued to settle in the same place as the earlier Serednii Stih, a continuity not seen in the Volga-Ural steppes, where most Eneolithic settlement sites exhibited no re-use by the Yamnaya.

*qpAdm* analysis reveals that all groups visually on the Dnipro Cline in the PCA can be well modeled with either UNHG or GK2 (individual I12490 from Golubaya Krinitsa in the Middle Don dated 5610-5390 BCE) at one extreme, and Core Yamnaya on the other (p-values between 0.07 and 0.85). Some populations of the cline (SSmed) can be modeled as Core Yamnaya and either GK2 (p=0.43) or UNHG (p=0.27); others, like the Don Yamnaya, can be modeled only as Core Yamnaya and UNHG (p=0.08) but not GK2 (p=0.0001); and others, like SShi, as Core Yamnaya and GK2 (p=0.08) but not UNHG (p=0.003). Thus, the hunter-gatherer end of the Dnipro Cline is not clearly UNHG or GK2. We therefore model individuals of the Cline with ancestry from any population from the UNHG-EHG cline (Fig. 1c), observing that individuals can be modeled as a mix in which UNHG ancestry predominates but EHG ancestry is also present in individuals (similar to GK2). This reflects the admixture of Caucasus-Lower Volga ancestry with hunter-gatherers of the Dnipro-Don (or UNHG-GK2) area, rather than other areas of eastern Europe (such as the Volga area) in which the hunter-gatherer population was EHG. Using Core Yamnaya as a source for the Serednii Stih is, of course, ahistorical, as they postdate the Serednii Stih, and so the model of Core Yamnaya + UNHG/GK2 admixture must be interpreted as admixture between local Neolithic residents of the Dnipro-Don area with a second, unsampled, Eneolithic source, which together account for the ancestry of the Core Yamnaya and—with even more UNHG/GK2 ancestry—of the Dnipro cline as a whole.

The Don, situated geographically between the Dnipro and Volga, is represented in our data by individuals from Golubaya Krinitsa (in the Middle Don) and Krivyansky (in the Lower Don). Golubaya Krinitsa contained two archaeologically contrasting styles of graves, one compared to Dnipro Neolithic graves and the other like Serednii Stih.^39^ The GK2 individual can be modeled as 66.6±4.7% UNHG and 33.4±4.7% EHG (p=0.39), suggesting that intermediate populations between the Dnipro hunter-gatherers (represented by UNHG) and the EHG existed not only in the Upper Volga (the upriver portion of the Volga cline), but also in the Middle Don. When we examine populations using the most ancient sources (Karelia, UNHG, and CHG) of the steppe and Caucasus, we see that the Eneolithic population of the Lower Don at Krivyansky and Neolithic individuals from Golubaya Krinitsa can all be well modeled with variable proportions of CHG-related ancestry (Fig. 2a). The most CHG-related ancestry is seen at Krivyansky (58.9±2.4%); there is less (25.3±2.1%) in three individuals which (Fig. 1) we group as GK1; and individual GK2 is consistent with having none or very little (4.0±2.2%), fitting the simpler EHG+UNHG model mentioned above. Thus, the Neolithic and Eneolithic individuals of the Don were a mixture of European hunter-gatherer ancestries (intermediate between the Dnipro-sampled UNHG and the Volga-sampled EHG, paralleling the intermediate geographic position of the Don) and southern CHG-related ancestry (Fig. 2a). When did the CHG-related ancestry reach the Don area? Its presence in a ^14^C-dated individual of the GK1 group (I12491/5557-5381 BCE) and others from the region^7^ suggest it was present there as early as the Neolithic. However, its absence from GK2 of similar ^14^C age proves that it was not a general feature of the Neolithic population. Both GK1/GK2 dates may be too early given that archaeologists of Golubaya Krinitsa interpreted people of the site as in contact with people of the much later Eneolithic Serednii Stih Culture.^40^ Moreover, an outlier Serednii Stih individual from Igren (I27930; 4337-4063 cal BCE) is consistent with all its ancestry coming from GK2; this could be an example of long-distance migration from the Don to the Dnipro, but also casts some doubt on the much older date of the GK2 individual, as genetic identity across more than a millennium in two different locations seems implausible given the diverse admixtures taking place throughout the steppe during the Eneolithic. The interpretation of the Golubaya Krinitsa population is further complicated by uncertainties as to their date due to freshwater reservoir effects in individuals who have a diet heavily reliant on freshwater fish. This can make nominal dates up to a millennium too old in this region.^41^ Further sampling along the Don would shed light on the distinctive processes and temporality of the ancestry change along this major river and place both the Golubaya Krinitsa individuals and those of Krivyansky on the Don mouth in their proper context.

**Figure 2.**
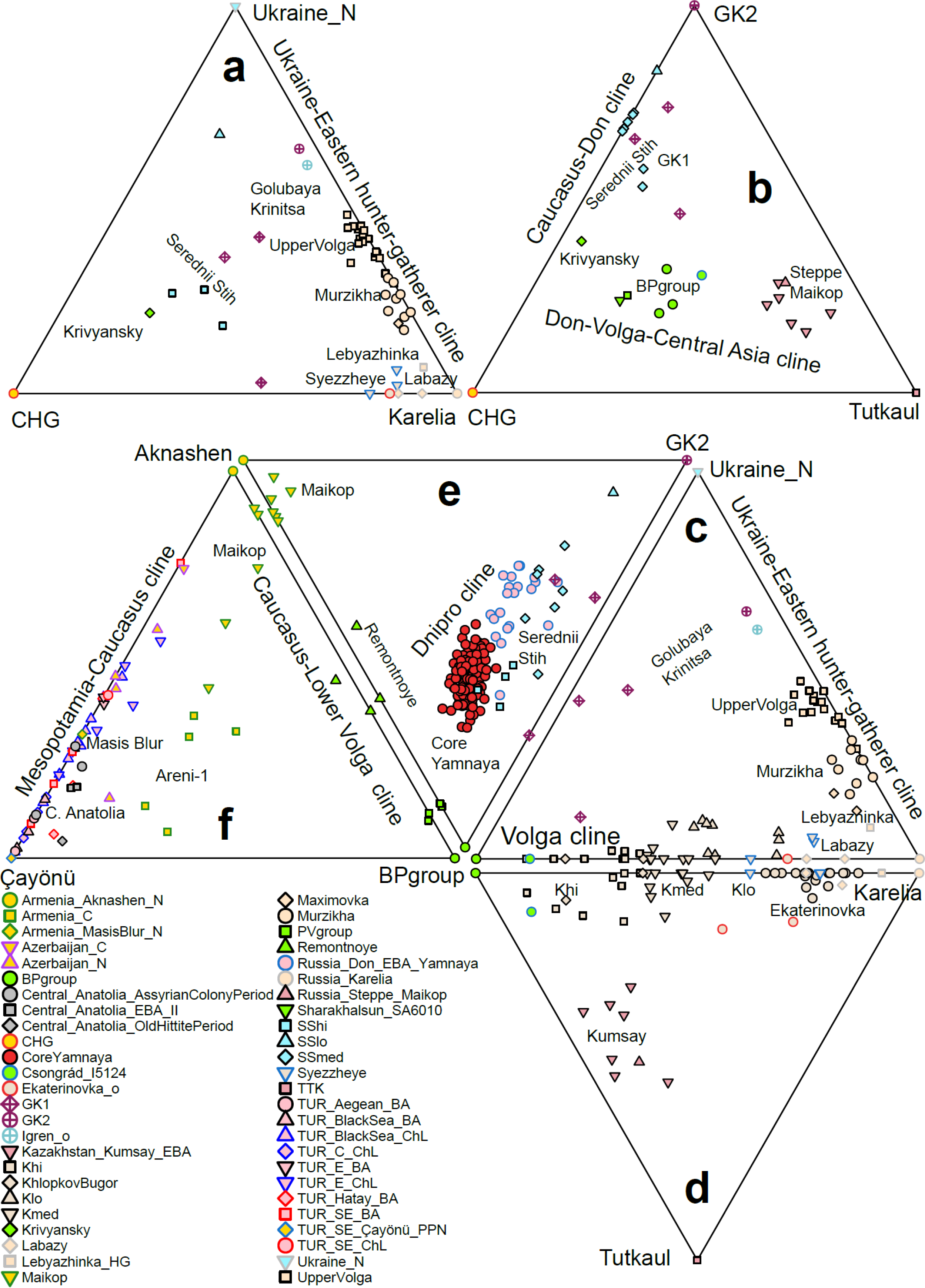
The three Eneolithic clines in the context of Eneolithic and Bronze Age admixture. Six 3-source models elucidate a complex history of admixture. Individuals plotted at the triangle edge fit (p>0.05); the simpler 2-source model is plotted for individuals with a negative coefficient from one of the three sources. The corners of each triangle represent the sources. Unplotted individual all give fits at P<0.05 and so should be viewed as poorly described by the model. (a) Caucasus and European hunter-gatherer admixtures in the “Old Steppe”: Krivyansky on the Lower Don received much more CHG-related admixture than upriver people of the Middle Don at Golubaya Krinitsa. In the Middle and Upper Volga and the Kama River, populations belonged to the old EHG cline with negligible CHG-related influence. (b) The “Don-Volga” difference. On the Lower Volga and North Caucasus piedmont, the BPgroup did receive CHG-related ancestry like its western Lower Don counterpart at Krivyansky; but, unlike Krivyansky, it also received ancestry from Central Asia; this eastern influence was higher still in the Bronze Age Steppe Maikop. (c) The “Volga Cline” vis-à-vis the Don: populations at Khvalynsk, Klopkov Bugor, and Ekaterinovka are clinal between the Berezhnvoka cluster on the Lower Volga and the upriver EHG-like populations of the Middle Volga (Labazy and Lebyazhinka). (d) the “Volga Cline” vis-à-vis Central Asia: a slight excess of Central Asian ancestry in the Khi subset of Khvalynsk. (e) the “Dnipro” cline: the Core Yamnaya are on one end of a cline that also includes the Don Yamnaya and Serednii Stih populations. The cline is formed by admixture from the “Caucasus-Lower Volga” (CLV) cline that is formed by differential admixture of Neolithic Caucasus and BPgroup people. The CLV Cline includes diverse people buried in kurgans at Berezhnovka, Progress-2, Remontnoye, and Maikop sites Klady and Dlinnaya-Polyana ∼5000-3000 BCE. (f) “West Asian”: CLV ancestry first appears in the Chalcolithic population at Areni-1 in Armenia and is also present in the Bronze Age at Maikop. The majority of the ancestry in both populations is from West Asian sources from the Mesopotamia-Caucasus (or Çayönü-Masis Blur-Aknashen) cline. Chalcolithic and Bronze Age Anatolians lack CLV ancestry but traces of it can be found in Bronze Age Central Anatolians.

It has been suggested^7^ that the Yamnaya were formed by a substantial contribution of ∼65% Golubaya Krinitsa people from the Middle Don, that already had ∼20-30% CHG-related ancestry, with an additional ∼35% CHG-related ancestry. This scenario implies that they were formed in the Don area as the result of the CHG-related admixture observed there. Our results contradict this as the Core Yamnaya do not fit models with CHG-related and either GK1/GK2 sources (p<1e-6), suggesting that they have ancestry not accounted for by the model of ref.^7^ To understand the source of this ancestry, we fit the model of Fig. 2a (with the most ancient sources: Karelia, UNHG, and CHG) and observed that its failure (p=2×10^-20^) is explained by the fact that it severely underestimates their shared genetic drift with both Afontova Gora-3 from Upper Paleolithic Siberia (Z=-5.2) and Anatolian Neolithic (Z=-6.8).^6^ Thus, the Yamnaya must have Siberian- and Anatolian-related ancestry and cannot be a simple mixture of Caucasus- and Middle Don hunter-gatherers. A Volga source of the Siberian-related ancestry is strongly suggested by the fact that the Volga cline is shifted away from the Dnipro cline precisely in the direction of Siberian populations (Fig. 1b). That the Volga cline populations had such ancestry is proven by the fact that the model of Fig. 2a fails them precisely for the same reason as it does the Core Yamnaya as it also underestimates shared drift with Afontova Gora-3, e.g., for BPgroup (p=1×10^-8^ and Z=-4.5). This extra ancestry in BPgroup is also affirmed positively by the fact that it can be modeled as a mixture of Krivyansky and ∼24% Central Asian (Siberian-related) Tutkaul^19^ ancestry (p=0.13). When we fit both Krivyansky and the BPgroup with the same model that includes all relevant ancestries (Fig. 2b)—CHG, GK2, and Tutkaul—we see that indeed Krivyansky has little to no Central Asian ancestry (5.1±3.6%) but it can be fitted as 56.7±2.6% CHG-related and 43.3±2.6% GK2 alone (p=0.37), while BPgroup does have 29.3±2.2% Tutkaul ancestry. The model of Fig. 2b corrects for the missing Siberian-related ancestry in the Yamnaya, predicting shared genetic drift with Afontova Gora-3 reasonably accurately (Z=-1.7), but still fails (p=1e-9) as it does not predict shared drift with Anatolian Neolithic (Z=-6.1). Thus, while ancestry from the Volga can explain the Siberian relatedness of the Core Yamnaya it cannot explain the Anatolian Neolithic relatedness as this was not a component of Volga cline populations.

Our new data resolve the extent of the spread of eastern “Central Asian” or “Siberian” ancestry into the Pontic-Caspian steppe. It was present, during the Eneolithic, on the Volga and in the North Caucasus Steppe, but further west on the Don there still existed populations without much or any of it like those at Krivyansky and Golubaya Krinitsa. When we repeat our modeling of the Volga Cline as a mixture of BPgroup and EHG sources but add either a western (UNHG) or eastern (Tutkaul) source (Fig. 2c,d) we see that individuals on the cline remain largely well-modeled as linear combinations of the two groups: Fig. 2c shows the characteristic “bend” of the Volga Cline with a portion showing variable Berezhnovka ancestry and the other (including many individuals from the Upper Volga and Murzikha) showing variable UNHG ancestry which increases further still in the GK2 individual from the Don. Fig. 2d shows that individuals of the Volga Cline have more Tutkaul ancestry than is explained by the simpler Berezhnovka-Karelia model; however, the deviations are small (4.4±2.6% Tutkaul ancestry for “Khi”). The Eneolithic Volga was an admixture zone between downriver BPgroup people with upriver EHG ones that included Central Asian ancestry mainly via BPgroup. Crucially, the Core Yamnaya fail all models of Fig. 2a-d (p<1e-8), and thus its origins must include a different blend of ancestry than the CHG-EHG-UNHG-Tutkaul ancestries involved in these models. As we will now see, this ancestry came from a third cline formed between the Caucasus Neolithic populations and those of the Lower Volga.

#### (3) Caucasus-Lower Volga Cline (CLV)

The Yamnaya are on the edge of the Dnipro cline, having less UNHG/GK2-related ancestry than other cline populations; thus, they cannot be modeled in terms of them alone (Fig. 1), but must have possessed more of a second source of ancestry. We found that the only consistently fitting (p=0.67) two-way model for the Core Yamnaya involved 73.7±3.4% of the SShi subset of the Serednii Stih population and 26.3±3.4% from a population represented by a sample of two individuals from Eneolithic burial sites at Sukhaya Termista I (I28682) and Ulan IV (I28683), dated 4152-3637 BCE near the village of Remontnoye, north of the Manych Depression on the watershed between the Lower Don and Caspian. The Remontnoye population is on neither the Volga nor Dnipro clines and is neither genetically close (Fig. 1) nor forms a clade (p<1e-10) to any other single sampled population. We determined that it had at least two sources: a southern one from the Caucasus—either descendants of the Aknashen Neolithic in Armenia^6^, or ancestors of people of the Bronze Age Maikop^5^ culture—and a northern one from a population from the low-EHG end of the Volga Cline such as the BPgroup. The Caucasus component is about half when using either Aknashen (44.6±2.7%; p=0.66) or Maikop (48.1±2.9%; p=0.44) as the proxy for the southern source. We also observed that the main cluster of Maikop individuals, including those buried in kurgans in Klady and Dlinnaya-Polyana, can be modeled as having 86.2±2.9% (p=0.50) Aknashen ancestry. Thus, there exists a Caucasus-Lower Volga (CLV) cline: Aknashen-Maikop-Remontnoye-Berezhnovka. These four populations are arrayed in order of decreasing Caucasus Neolithic component, concordant with their south-to-north geographical location. However, there were also populations of the CLV cline that bucked this latitudinal trend, such as the people of the North Caucasus at Progress-2 and Vonyuchka-1 that, unlike their Maikop neighbors, had little Caucasus Neolithic ancestry and were most like the people of Berezhnovka-1 in the Lower Volga. These violations of the genetic-geographic pattern prove long-range connectivity across the CLV area; they also caution us not to easily interpret genetic position along the CLV cline as predictive of position within the CLV geography.

What was the proximal source for the southern ancestry of the intermediate populations of the CLV cline? Aknashen makes a poor choice, as it is both geographically remote from the steppe and earlier by two millennia (5985-5836 BCE) than Remontnoye. Neither is Maikop a good proximal source; it is geographically closer, but postdates (3932-2934 BCE) Remontnoye.

Settlements at Meshoko and Svobodnoe, dated 4466-3810 BCE,^42^ provide a temporally, geographically, and archaeologically plausible source, as they exhibit exchanges of exotic stone, copper, and stone mace heads with Volga Cline sites, setting the context for the expansion of Aknashen-like ancestry northward and Berezhnovka-like ancestry southward. These settlements are temporally earlier than Maikop and later than two individuals from Eneolithic Unakozovskaya (ref.^5^ 4607-4450 BCE, and this study) in the North Caucasus; however, unlike Aknashen and Maikop, the Unakozovskaya population is not a good genetic source for Remontnoye, as the model BPgroup+Unakozovskaya fails (p<0.001) by overestimating (Z=3.8) shared genetic drift with the CHG. The Unakozovskaya was not exactly the same genetically as the Maikop who succeeded them (p=2e-11) but were genetically similar (Fig. 1) and can be modeled as 95.3±6.3% Maikop and 4.7±6.3% CHG (p=0.46). Thus, there were three elements of ancestry in the North Caucasus in the Eneolithic: (i) Aknashen-related ancestry was dominant, representing the spread of the Neolithic from the south across the Caucasus mountains; (ii) there was some variation in CHG-related ancestry as suggested by the Maikop-Unakozovskaya contrast; and (iii) there was also a small component of northern Lower Volga ancestry of about one seventh in the Maikop on average. Thus, in the north Caucasus there lived, side by side, both “high steppe” ancestry people genetically close to the Lower Volga Berezhnovka population (individuals at Progress-2 and Vonyuchka-1), as well as “low steppe” ancestry people in which the Lower Volga ancestry had been diluted by the greater contribution of the (Aknashen-related) Caucasus Neolithic.

The Remontnoye and Berezhnovka people, like the Maikop people, were buried in kurgans. Thus, the kurgan burial rite was widespread 5000-3000 BCE among people of diverse ancestry from both the edges and middle of the CLV Cline, suggesting that—regardless of its ultimate origin and whether it was culturally adopted or spread by migration—it was common among the people of the CLV region.^22^ In contrast, a distinctive position of the body on the back with knees raised and the floor of the burial pit covered with red ochre was shared by all the steppe groups including Serednii Stih, groups on the Volga Cline, and Remontnoye, while the Maikop burial position was contracted on one side. Thus, some funeral customs united Maikop with the steppes and others separated them.

The discovery of the CLV Cline suggests a solution to the question of the origin of the Dnipro Cline and thus the genetic origins of the Yamnaya. Most of their ancestors were people of the CLV Cline, similar to the sampled Remontnoye individuals. These CLV ancestors were drawn into the Dnipro-Don region and mixed with local groups to form Serednii Stih people and eventually the Yamnaya. It must be emphasized that the CLV and Dnipro-Don sources need not have been identical to the sampled Remontnoye and SShi populations or have lived close to the sampling locations of these two populations. The Dnipro Cline can be fit (Fig. 2e) by a 3-way model in which the GK2 admixed with groups of mixed Aknashen and Berezhnovka ancestry. We note the aforementioned caveat that either of GK2 or UNHG could be contributing to the Dnipro Cline, but chose GK2 in Fig. 2e as this model has a higher p-value (p=0.93) for the Core Yamnaya than the alternative with UNHG as the source (p=0.04); however, we do not take this as evidence that the GK2 population was a better source than the UNHG as we have far better data for UNHG (n=35 individuals) than GK2 (n=1), which provides more power to detect slight but qualitatively unimportant oversimplifications in models. Note also, that GK2 is itself ∼2/3 UNHG in ancestry, and that the proportion of either GK2 (22.5±1.8%) or UNHG (17.7±1.3%) is similar, and about one fifth. A full exploration of 3-way models (Supplementary Information section 2) reveals that the Yamnaya could have been formed from diverse (but similar) distal sources which include populations of (i) Neolithic or Chalcolithic age from Armenia^6,9^ and Azerbaijan^43,44^ representing the “Caucasus Neolithic”, (ii) GK2, UNHG, or Serednii Stih representing the Dnipro-Don area, and (iii) BPgroup or PVgroup representing the Lower Volga-north Caucasus Eneolithic. What is invariant among the class of 2- and 3-way models for the Core Yamnaya is that they posit their descent from people of the CLV Cline (the remaining four fifths of their ancestry) who admixed with Dnipro-Don people of substantial UNHG ancestry.

Our results show that movement of people and culture we document as having occurred along the CLV Cline was the vector by which Caucasus-derived ancestry like that present in the Aknashen Neolithic population flowed into the steppe and into the ancestors of the Yamnaya^45^. Crucially, the successful Remontnoye+SShi model predicts shared genetic drift with the Anatolian Neolithic outgroup well (Z=-0.8). CLV cline populations can account for both Siberian-related (via the Lower Volga component) and Anatolian Neolithic-related (via the Caucasus Neolithic component) affinities of the Yamnaya. Archaeological evidence shows that Balkan copper was traded during the late 5^th^ millennium BCE across the steppes to North Caucasus farmer sites (Svobodnoe) and to the Volga (Khvalynsk), while Neolithic pots like those from Svobodnoe appeared in Dnipro-Don steppe sites connected with the Seredni Stih culture (Novodanilovka), documenting an active period of cultural exchange that was the context for the movement of groups of mixed BPgroup/Aknashen-related ancestry into the Dnipro-Don steppes.

##### CLV impact in the Caucasus and Anatolia

CLV Cline people also had an impact further south, in Armenia and Anatolia (Fig. 2f). The earliest evidence of steppe ancestry south of the Caucasus is at Areni-1 in Chalcolithic Armenia around 4000 BCE^9^, documenting its southward penetration which parallels the incursion of Caucasus ancestry generating the Volga/Dnipro clines on the steppe. Our analysis (Supplementary Information section 2) clarifies that in Areni-1 the Lower Volga ancestry (26.9±2.3% BPgroup) admixed with a local “Masis Blur”-related Neolithic substratum, in contrast to the North Caucasus (at Maikop) where it combined with an “Aknashen”-related Neolithic substratum. The Aknashen/Masis Blur distinction of the Neolithic population of Armenia reflected the dilution of the native CHG ancestry that was higher in Aknashen than in Masis Blur.^6^ We can model Masis Blur as 33.9±8.6% Aknashen and 66.1±8.6% Çayönü ancestry (p=0.47) associated with the Pre-Pottery Neolithic of the Tigris Basin of Mesopotamia^46^, thus documenting the spread of early Neolithic ancestry into the Caucasus that formed a cline of diminishing Mesopotamian-related and increasing CHG-related ancestry: Çayönü-Masis Blur-Aknashen. Using CHG as the source, we see that the two populations from Armenina differed indeed in their retention of CHG ancestry, with more (42.0±3.8%) in Aknashen than in Masis Blur (13.7±4.0%). Some Anatolian Chalcolithic and Bronze Age groups can be derived entirely from this north-south Caucasus-Mesopotamian cline (Fig. 2f), while others also have ancestry from the east-west Mesopotamian-Anatolian cline, lacking any steppe ancestry.^22,43,45,47,48^

The discovery of the Mesopotamian-Caucasus cline allows us to study the ancestry of the population of Bronze Age Central Anatolia^22^ from the Early Bronze Age (2750-2500 BCE), Assyrian Colony (2000-1750 BCE), and Old Hittite (1750-1500 BCE) periods. We cannot be certain of what languages were spoken by these individuals in what may well have been multilingual societies, but we document for the first time that they had a small amount of CLV cline ancestry combined with Mesopotamian (Çayönü) ancestry (Supplementary Information, section 2; Fig. 2f; Extended Data Fig. 1). The inferred amount of ancestry from the CLV or CLV-influenced source depends on the amount of “dilution” of this ancestry in the source: more such ancestry is required from populations of higher dilution. For example, it is estimated as 10.8±1.7% ancestry (p=0.14) from the BPgroup, or about double 19.0±2.4% from Remontnoye (p=0.19)—whose own ancestry is about half from the BPgroup—or 33.5±4.8% of Armenia_C ancestry (p=0.10)—where the BPgroup ancestry is lower.

The exact source of the steppe ancestry in Anatolia cannot be precisely determined, but it is noted that all fitting models involve some of it (Extended Data Fig. 1a). Some of the steppe-related sources can be rejected on chronological grounds; for example, the Core Yamnaya itself (12.2±2.0%; p=0.10) as well as western Yamnaya-derived populations from Southeastern Europe such as from Boyanovo or Mayaky Early Bronze Age^36^ (Extended Data Fig. 1b). Moreover, when we consider pairs of steppe sources (and can thus place the steppe ancestry at varying points along the Volga, Dnipro, and CLV clines), we observe a negative hunter-gatherer contribution (−3.4±2.6% EHG) on the Volga cline, and also on the Dnipro cline (−2.3±2.7% UNHG or −3.9±3.5% GK2); thus, there is no evidence that the admixing population had more EHG/UNHG/GK2 ancestry than the BPgroup/Core Yamnaya endpoints of these two clines (Supplementary Information section 2). The admixing population in this analysis contributed a significant amount of BPgroup ancestry (8.8±2.7%) from the CLV cline and was consistent with being on that cline (p=0.129). Thus, a model in which the steppe ancestry is derived from the Caucasus-Lower Volga Eneolithic is not only geographically and chronologically plausible but also genetically so. The steppe+Mesopotamian class of models fit the Central Anatolian Bronze Age but do not fit any of the Chalcolithic/Bronze Age Anatol0ian regional subsets (p<0.001; the BPgroup+Çayönü model is shown in Extended Data Fig. 1c), indicating that their success is not due to their general applicability. Moreover, the steppe ancestry in the Central Anatolian Bronze Age is observed in all individuals of the three periods (Extended Data Fig. 2d) and is thus not driven by any outlier individuals within the population. Its presence in both Early Bronze Age individuals from Ovaören south of the Kızılırmak river and in Middle Late Bronze Age individuals from Kalehöyük just within the bend of the river is consistent with the idea that the Kızılırmak formed an Anatolian-Hattic linguistic boundary that was crossed some time before the ca. 1730 BCE conquest of Hattusa by the Hittites.^49^ Regardless of the linguistic identity of the sampled individuals, the truly unique blend of CLV and Mesopotamian ancestries found in the Central Anatolia Bronze Age calls for an explanation.

How and when did this blend reach Central Anatolia? We note that populations along the path from the steppe to Central Anatolia can all be modeled with BPgroup ancestry and distinctive substratum ancestries along the north-south / Caucasus-Mesopotamia cline: Aknashen-related in the North Caucasus Maikop; Masis Blur-related in the South Caucasus Chalcolithic population of Armenia at Areni-1; and Mesopotamian Neolithic for the Central Anatolian Bronze Age (Extended Data Fig. 1e, f). This series of admixtures had certainly begun by ca. 4300-4000BCE (the date range of the Armenia_C population^9^) and can be dated using DATES to 4382±63BCE (Extended Data Fig. 2f). The Pre-Pottery Neolithic population of Çayönü was itself genetically halfway between that of Mardin^10^, 200km to the east, and the Central Anatolian pottery Neolithic at Çatalhöyük along the east-west / Mesopotamian-Anatolian cline. Chalcolithic/Bronze Age people from Southeastern and Central Anatolia all had ancestry from the same Çatalhöyük-Mardin continuum and such populations may have been proximal sources for the Çayönü-related ancestry of the Central Anatolian Bronze Age population (Supplementary Information section 2). If the Proto-Anatolian population was formed in this region by the admixture of CLV cline people with Mesopotamian ones then their descendants may have been present there at the unknown site of Armi whose Anatolian personal names are recorded by their neighbors in the kingdom of Ebla in Syria.^50^ We thus propose the following hypothesis: that CLV cline people migrated southwards ca. 4400BCE, or about a millennium before the appearance of the Yamnaya, (admixing with different substratum populations along the way) and then westwards before finally reaching Central Anatolia.

We in fact find Y-chromosome evidence that is consistent with the autosomal evidence. Sporadic instances of the steppe-associated Y-chromosome haplogroup R-V1636 in West Asia occurred at Arslantepe^43^ in Eastern Anatolia and Kalavan^9^ in Armenia in the Early Bronze Age (∼3300-2500 BCE) among individuals without detectible steppe ancestry^45^ and these could be remnants of the dilution process. This haplogroup was found in the male individual from Remontnoye, both individuals from Progress-2^5^ and two of three males from Berezhnovka, in addition to its occurrence in eleven individuals of the Volga Cline and thus was a prominent lineage of the pre-Yamnaya steppe. Isolated instances have also been found beyond the steppe in Corded Ware individuals from Esperstedt in Germany^17^ and Gjerrild in Denmark.^51^ The expansive distribution of R-V1636 on the steppe and beyond contrasts with its disappearance on the steppe after the Yamnaya arrived on the scene: a single individual (SA6010; 2886-2671 BCE) from Sharakhalsun^5^ has it, with a genetic profile consistent with CLV ancestry (Fig. 2), the last detected holdout of this once pervasive population (Fig. 3).

**Figure 3.**
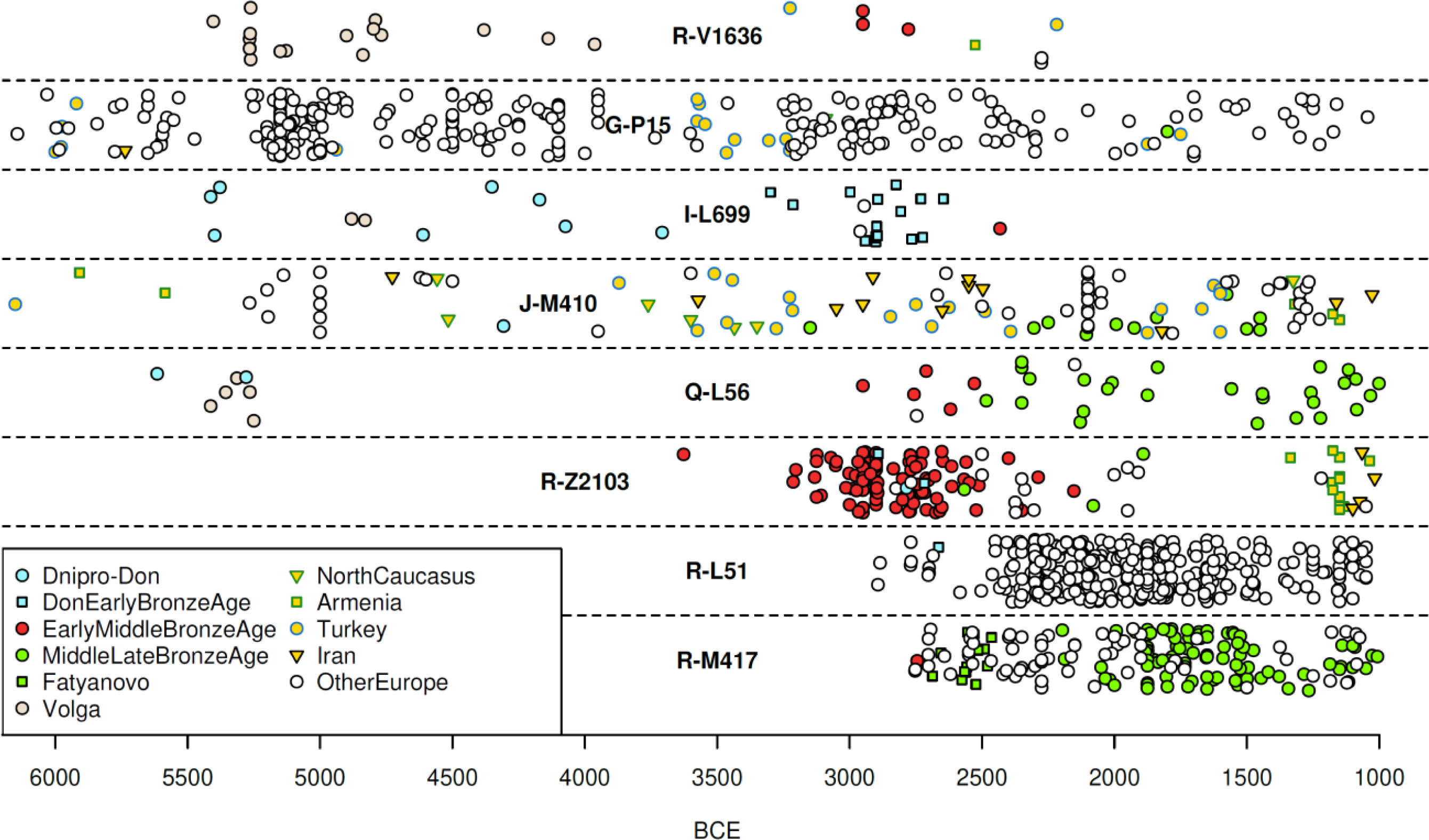
Patrilineal succession. Temporal distribution of key Y-chromosome haplogroups from Kazakhstan, Kyrgyzstan, Mongolia, Russia, Turkmenistan, Ukraine, Uzbekistan, and comparative regions of Europe and West Asia 6000-1000 BCE. The Early and Middle Bronze Age group includes the Yamnaya, Afanasievo, Poltavka, Catacomb, Chemurchek, and North Caucasus cultures; the Middle and Late Bronze Age group individuals of diverse cultures down to 1000 BCE including those of the Sintashta, Andronovo, Potapovka, and Srubnaya cultures.

### The Yamnaya expansion broke correlations between geography and genetics

We have traced the origins of the Yamnaya to the Dnipro Cline and the populations of the Serednii Stih culture: the Yamnaya were formed as people of the CLV cline admixed with people of the Dnipro-Don area having UNHG ancestry. Deeper in time, the CLV cline was formed by the admixture of Aknashen-related and BPgroup-related people who, in turn, were formed by earlier mixtures still: the Caucasus Neolithic represented at Aknashen by the admixture of CHG people with Neolithic farmers of the Fertile Crescent^6,10^ and the lower Volga Eneolithic people represented by BPgroup had ancestries that were related to CHG, EHG, and people from Siberia or Central Asia. Dating this complex sequence of admixtures could be done by generating time transects of fine resolution in all relevant areas from which the ancestors of the Yamnaya were drawn across the millennia until they finally combined to form the Yamnaya genetic profile somewhere in the territory of the Serednii Stih culture: seeing the admixture “as it happened” through the lens of ancient DNA. Our study has revealed the outlines of this millennia-long process and future studies may fill in the details.

A different way is to date the admixture itself in the genomes of the Yamnaya using methods like DATES^52^ to measure the average sizes of stretches of ancestry related to UNHG/EHG hunter-gatherer populations on the one hand, and West Asian/Caucasus-related populations on the other, as this reflects the number of generations elapsed since mixture began and stretches of ancestry broke down. This population contrast aligns to the differentiation along PC2 (Fig. 1). We would also like to model the Core Yamnaya in terms of ancestry along the Dnipro cline itself (their last and most proximal admixture event), but unfortunately this is challenging given that the Yamnaya themselves are the end of the Dnipro cline (Fig. 1). The inferred date of 4038±48 BCE (Extended Data Fig. 2a) should thus be viewed with caution given the complex history of the ancestors of the Yamnaya, and admixture may have taken place both before and after this date.

Nonetheless, an Eneolithic time frame (with a small standard error of <2 generations) proves that the admixture derived using qpAdm and observed visually in PCA did not occur in the remote past, but corresponds, at least in part, to the efflorescence of the Serednii Stih culture that our reconstruction points to as ancestral to the Yamnaya.

Uncertainty about where, exactly, within the territory of the Serednii Stih culture the ancestors of the Core Yamnaya lived contrasts with their expansive distribution after the formation of the Yamnaya archaeological horizon: individuals we identified as “Core Yamnaya” (Extended Data Table 2) cluster in a small portion of the PCA (Fig. 1) and are from several countries: China, Hungary, Kazakhstan, Moldova, Romania, Ukraine (Extended Data Table 2), and 15 different locations in Russia (Fig. 4a). The homogeneity is also evident in a mean F_ST_ of 0.005, comparable to that between modern northern Europeans (Extended Data Table 3). This remarkable homogeneity across vast geographical distances of the “eastern” expansion of the Yamnaya shows that many of them mixed very little if at all with any of the people that inhabited the Eurasian steppe before them. The Don Yamnaya (Fig. 4a) are distinctive and can be modeled with 79.4±1.1% Core Yamnaya and 20.6±1.1% UNHG ancestry; the actual proportion of Core Yamnaya ancestry may be lower if, as is plausible, the Core Yamnaya admixed with a Serednii Stih population of partial UNHG ancestry (e.g., 40.0±4.7% with SSmed as the Serednii Stih source). The Don Yamnaya were formed in the late 4^th^ millennium BCE (Extended Data Fig. 2b), a time during which unmixed UNHG, after a millennium or more of the Serednii Stih culture, would be rare if they existed at all.

**Figure 4:**
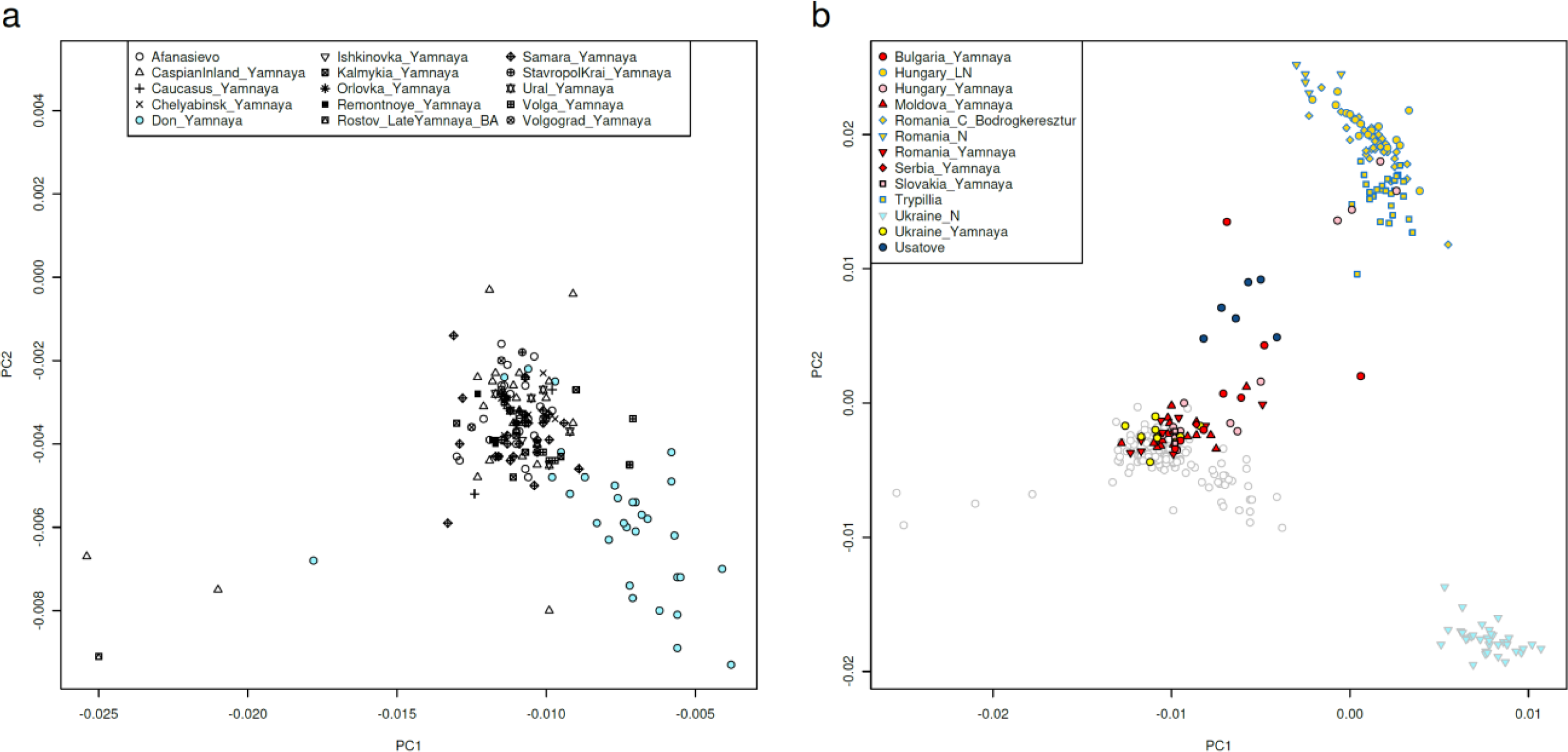
Population structure in people with a Yamnaya cultural affiliation. Individuals are projected in the same space as in Fig. 1. (a) showing that the Core Yamnaya cluster (black symbols) from diverse sites is differentiated from the Don Yamnaya (blue) who tend towards the UNHG. (b) Yamnaya individuals in the West (Ukraine, Hungary, Slovakia, and Southeastern Europe) include a tight cluster of individuals as well as others that tend towards the direction of European Neolithic and Chalcolithic groups from Romania and Hungary. Individuals from Russia are shown in grey circles in panel (b).

The western expansion of the Core Yamnaya also brought them into southeastern Europe; Yamnaya there or other individuals of “high steppe ancestry” can be found as far west and south as Albania and Bulgaria.^6^ Many western Yamnaya cluster with the Core Yamnaya, but many also deviate in the direction of Neolithic and Chalcolithic populations of southeastern and central Europe (Fig. 4b) and can be modeled with admixture from such populations (Extended Data Table 4). This admixture also took place in the late 4^th^ millennium BCE (Extended Data Fig. 2c), after the sporadic early Chalcolithic migrations into southeastern Europe from the steppe.^36^ It is interesting that after the Don Yamnaya formed they participated little or not at all in the Core Yamnaya expansion to either the Altai or SE Europe, and thus the Lower Don represented a cul-de-sac for the Yamnaya expansion.

The late 4^th^ millennium BCE admixtures with European farmers and UNHG-admixed populations frame the Dnipro-Don region from west and east, providing another line of evidence for the formation of the Yamnaya within this region. Y chromosome haplogroup sharing—which traces the entirely male line and is of particular interest in societies that have patrilineal traditions—(Fig. 3) is less informative for tracing the origins of the Core Yamnaya, but proves continuity of the Don Yamnaya with their Serednii Stih ancestors. Haplogroup I-L699 was an important lineage in the Dnipro area since the Neolithic hunter-gatherer period, continued to be prevalent among the Serdenii Stih, and in the Don Yamnaya was dominant (17/20 instances).

The Core Yamnaya belonged primarily to haplogroup R-M269 (49/51 instances) most of which could be determined as belonging to the Z2103 sub-lineage (41/51). This lineage is unprecedented in our sampling of the steppe before the Yamnaya period; its closest relative is the L51 lineage which dominated the Beaker group^3^ and mainland Europe outside the steppe (Fig. 3), with a slightly more distant relative in the R-PF7563 lineage found in Pylos in Mycenaean Greece.^45^ With an estimated time of formation of ∼4450 BCE (https://www.yfull.com/tree/R-L23/; v11.04.00), the R-L23 lineage unifies Beaker, Yamnaya, and Mycenaean Y-chromosomes within an Eneolithic timeframe, which is consistent with the ancestors of these three groups being part of a single population in the Yamnaya period itself since population divergences are always lower than the genetic divergences of specific haplotypes. It is a challenge for future ancient DNA studies to find the population in which the Eneolithic R-L23 founder lived and to trace his R-Z2103 descendants. Their absence from the Eneolithic record, together with the evidence (discussed below) for isolation in the formative period of the Yamnaya suggest that he might have been part of a small group not yet sampled.

That the Core Yamnaya are part of the Dnipro cline may suggest an origin in the Dnipro basin itself, but (a) the Dnipro cline is generated by admixture with Dnipro-Don people (UNHG/GK2-related), and (b) the Yamnaya on the Don are also part of this cline, so an alternative origin in the Don area cannot be excluded. An origin of the Core Yamnaya further east, in the Caucasus-Volga region is unlikely given that they are not part of the Volga or CLV Clines. Conversely, placing Yamnaya origins west of the Dnipro is implausible as the Core Yamnaya are the population of the Dnipro Cline that is maximally derived from the eastern CLV Cline and they also do not have the European farmer-derived ancestry of western populations such as the Usatove (Fig. 1b).^15^ The Core Yamnaya share ancestry with people of the whole Dnipro-Don-Volga-Caucasus region, but their ancestral mix includes all components also found in the Serednii Stih, while these are lacking elsewhere (Extended Data Fig. 3). A more western origin of the Core Yamnaya would also bring their latest ancestors in proximity to the place of origin of the Corded Ware complex whose origin is itself in question but must have certainly been in the area of central-eastern Europe occupied by the Globular Amphora culture west of the Core Yamnaya. The Corded Ware population, which could trace a large part of its ancestry to the Yamnaya,^2^ was formed by admixture concurrent with the Yamnaya expansion^52^ (Extended Data Fig. 2d), shared segments of IBD proving connections within a shallow genealogical timeframe, and had a balance of ancestral components from the Caucasus and eastern Europe indistinguishable from the Yamnaya.^6^ In combination, these lines of evidence suggests that it was formed indeed by early 3^rd^ millennium BCE admixture with Yamnaya, or, at the very least, genetically Yamnaya ancestors that need not have been Yamnaya in the archaeological sense. The geographical homelands of the Corded Ware and Yamnaya would then conceivably be in geographical proximity to allow for their synchronous emergence and shared ancestry. The Dnipro-Don area of the Serednii Stih culture fits the genetic data, as it explains the ancestry of the nascent Core Yamnaya and places them in precisely the area from which both Corded Ware, and Southeastern European Yamnaya (in the west) and the Don Yamnaya (in the east) could have emerged by admixture of the Core Yamnaya with European farmers and UNHG respectively.

### From Serednii Stih to Yamnaya: the 4^th^ millennium BCE

We estimated the population growth trajectory of Core Yamnaya using HapNe-LD, a methodology that can infer effective population size fluctuations in low-coverage ancient DNA data.^53^ Figure 5 shows the results separately analyzed for Core Yamnaya dating to the first three hundred years of our sampling (n=25) who produce a 95% confidence interval of 3829-3374 BCE for the time before growth, and 3642-3145 BCE for Core Yamnaya groups from the later three hundred years (n=26). In both cases, these correspond to growth from an effective number of reproducing individuals of a few thousand people. These intervals overlap at 3642-3374 BCE, corresponding to the late Serednii Stih period. Taken together with the admixture dating, these findings point to a scenario where the Serednii Stih were largely formed by admixture before 4000 BCE likely somewhere within the geographic span of the Dnipro-Don Cline. Half a millennium later, a subgroup of them developed cultural innovations that allowed them to expand dramatically, manifesting in a way that can be detected in the archaeological record around 3300 BCE in both the Pontic and Caspian Steppes.

**Figure 5:**
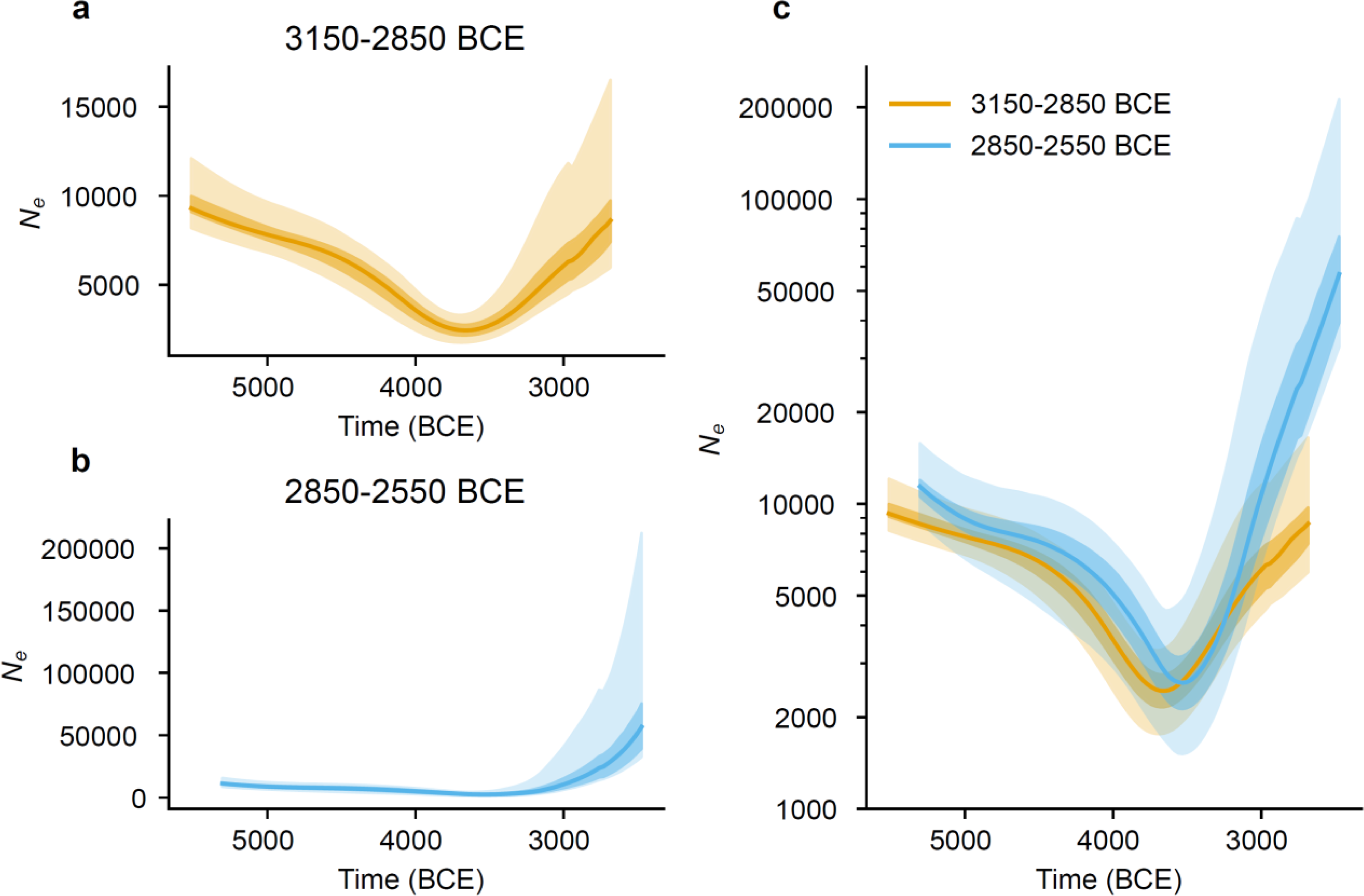
Trajectory of the Yamnaya expansion. We use HapNe-LD to estimate the changes in effective population size over time of Yamnaya ancestors, performing the computation separately for the individuals from the earlier three hundred years (a) of our sampling, and the later three hundred years (b); shading shows uncertainty intervals. We infer an extraordinary population expansion (c) after 3642-3374 BCE (intersection of 95% confidence intervals for the two analyses for the minimum), from a time when the effective size is a few thousand to an order of magnitude larger.

We tested for segments of the genome Identical-By-Descent (IBD) between pairs of individuals^54^, and found that the Yamnaya expansion transformed the interconnectedness of steppe populations. Before the Yamnaya, IBD links of ≥20cM did exist between regional populations (Fig. 6a), but this network of connections expanded dramatically in the Yamnaya period (Fig. 6b). Prior to the Yamnaya period, the rate of IBD links for individuals separated by more than 500km was vanishingly low (Fig. 6c), but in Yamnaya times, it was measurably non-zero (at a few percent) for distance separations between 500-5000km (Fig. 6d). We also studied close genetic relatives, defined as sharing at least three ≥20cM segments or a total sum of IBD ≥100cM. Both before and during the Yamnaya period, close relatives are only detected living within 500km, with a greatly elevated rate in the same cemetery (Fig. 6e, f). We examined Yamnaya-Afanasievo individuals in kurgans or kurgan cemeteries represented by at least two individuals (Fig. 6g), and found that around 14.4% of individual pairs were close relatives within kurgans and 7.4% of individual pairs were close relatives across kurgans of the same cemetery. These patterns are general across Yamnaya kurgan cemeteries (they are not dominated by one or a few sites with large numbers of samples). The observed rate of close relatives is much less than the 29.0% rate among pairs of individuals in Hazleton North chambered tomb in Neolithic Britain ∼3700BCE^55^ (p=0.00075; Fisher’s exact test), where 27 of 35 sequenced individuals were all found to be part of the same genetically tightly connected pedigree. These findings disprove theories that kurgans were “family tombs”^56^ of biological relatives. Instead, kurgan cemeteries largely included individuals that were biological kin only in the sense of sharing common descent for a population that lived many centuries in the past; if there were kinship links within the same kurgan, they were largely non-biological ones.

**Figure 6:**
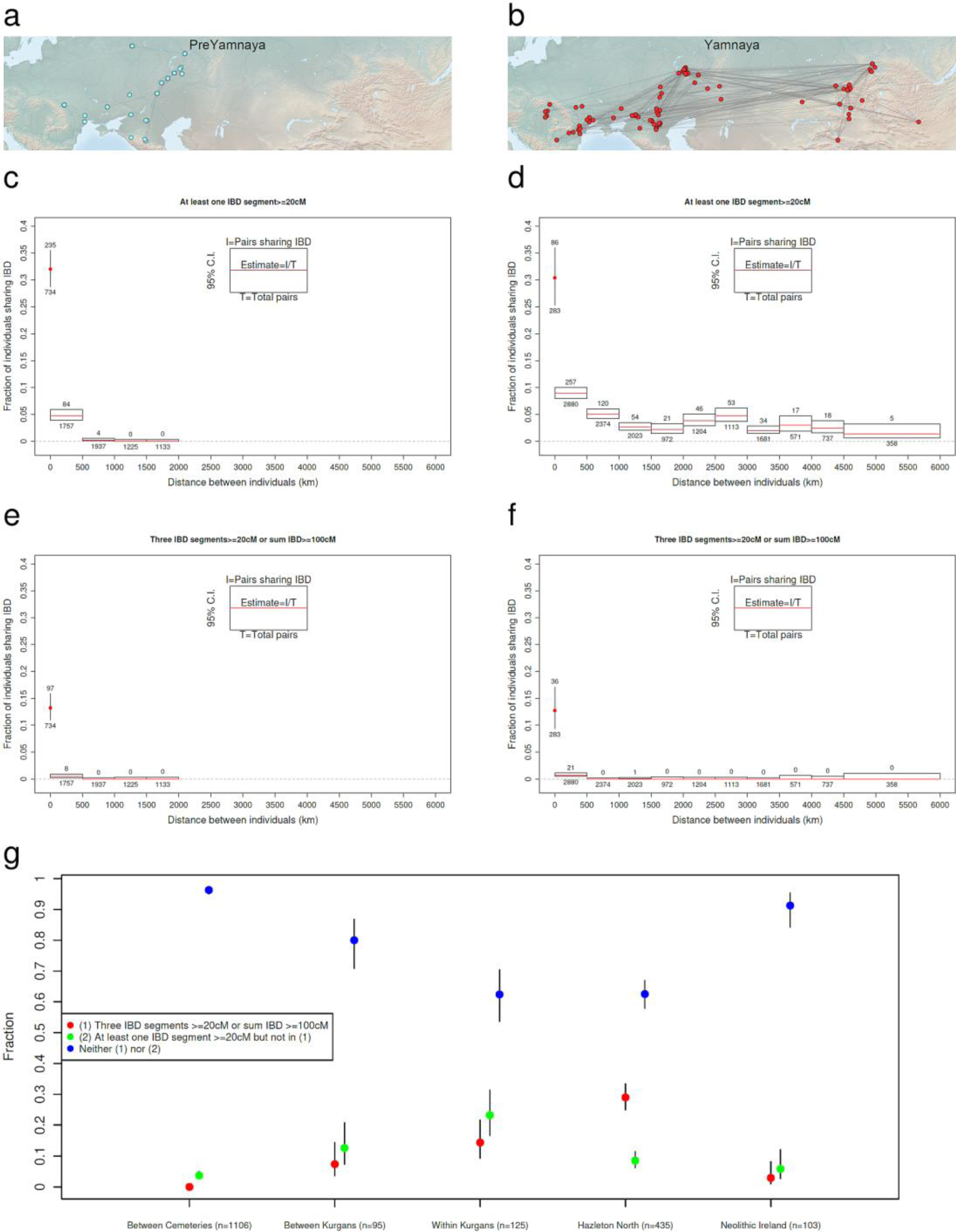
IBD analysis of the Yamnaya and their predecessors. Pairs of individuals linked by at least one IBD segment ≥20cM in length reveal a sparse and highly connected network in the Pre-Yamnaya (a) and Yamnaya (b) groups. No detectible IBD is found in the Pre-Yamnaya period beyond the scale of 1000km (c); Yamnaya share more IBD with each other at short distance scales but IBD sharing extends all the way to the ∼6000km scale of their geographical distribution. However, closely related individuals only occur at short distance scales in both Pre-Yamnaya (e) and Yamnaya (f) groups, indicating that the IBD sharing in the Yamnaya was a legacy of their common origin. (g) In a set of 9 Yamnaya cemeteries, and a total of 25 kurgans closely or distantly related individuals are virtually absent in inter-cemetery comparisons, more are found in inter-kurgan/within-cemetery comparisons, and more still in intra-kurgan comparisons; nonetheless, most Yamnaya individuals in all comparisons were unrelated. Kurgan burial of close kin was less common than in the case of a local patrilineal dynasty as at a Neolithic long cairn at Neolithic Hazleton North,^55^ but more common than in Neolithic monuments of Neolithic Ireland.^57^

### The origin and spread of the first speakers of Indo-Anatolian languages

Different terminologies exist to designate the linguistic relationship of Anatolian and Indo-European languages. The traditional view includes both within an “Indo-European” (IE) group in which Anatolian languages usually represent the first split^58,59^. An alternative terminology, which we use here, names the entire linguistic group “Indo-Anatolian” (IA) and uses IE to refer to the set of related non-Anatolian languages such as Tocharian, Greek, Celtic, and Sanskrit.^6,49^ Dates between 4300-3500 BCE have been proposed for the time of IA split^49,59–61^ predating both the first attestation of the Hittite language in Central Anatolia (post-2000 BCE^49^) and the expansion of the Yamnaya archaeological culture (post-3300 BCE). We identify the Yamnaya population as Proto-IE for several reasons. First, the Yamnaya were formed by admixture ∼4000 BCE and began their expansion during the middle of the 4^th^ millennium BCE, corresponding to this linguistic split date between IE and Anatolian. Second, the Yamnaya were the source of the Afanasievo migration to the east^62^ a leading candidate for the split of the ancestral form of Tocharian, widely recognized as the second split after that of Anatolian.^63^ Third, the Yamnaya can be linked to the languages of Armenia^45^ via both autosomal and Y-chromosome ancestry after ∼2500 BCE, and to the languages of the Balkans^13^ such as Greek.^45,47^ Fourth, the Yamnaya can be linked indirectly to other IE speakers via the demographically and culturally transformative Corded Ware and Beaker archaeological cultures of the 3^rd^ millennium BCE that postdate it by centuries. Most people of the Corded Ware culture of central-northern Europe had about three quarters of Yamnaya ancestry,^2^ a close connection within a few generations that can be traced to the late 4^th^ millennium BCE. The Beaker archaeological culture of central-western Europe also shared a substantial amount of autosomal ancestry with the Yamnaya and were also linked to them by their possession of R-M269 Y-chromosomes.^3^ The impact of these derivative cultures in Europe leaves no doubt that they were linguistically Indo-European as most later Europeans were; the Corded Ware culture itself can also be tentatively linked via both autosomal ancestry and R-M417 Y-chromosomes with Indo-Iranian speakers via a long migratory route that included Fatyanovo^20^ and Sintashta^4,22^ intermediaries. A recent study proposed a much deeper origin of IA/IE languages^64^ to ∼6000 BCE or about two millennia older than our reconstruction and the consensus of other linguistic studies. The technical reasons for these older dates will doubtlessly be debated by linguists. From the point of view of archaeogenetics, we point out that the post-3000 BCE genetic transformation of Europe by Corded Ware and Beaker cultures on the heels of the Yamnaya expansion is hard to reconcile with linguistic split times of European languages consistently >4000 BCE as no major pan-European archaeological or migratory phenomena that are tied to the postulated South Caucasus IA homeland ∼6000 BCE can be discerned.

The Yamnaya culture stands as the unifying factor of all attested Indo-European languages. Yet, the homogeneity of the Yamnaya patrilineal community was formed out of the admixture of diverse ancestors, via proximal ancestors from the Dnipro and CLV clines (Fig. 2e). Yamnaya and Anatolians share ancestry from the CLV Cline (Fig. 2e,f), and thus, if the earliest IA language speakers shared any genetic ancestry at all—the possibility of an early transfer of language without admixture must not be discounted—then the CLV Cline is where this ancestry must have come from. On the Anatolian side, we see that ancestry from the southern Caucasus Neolithic end of the CLV Cline was impactful during the Chalcolithic and Bronze Ages^45^ and Bronze Age Central Anatolians over the time span of Hittite presence there also had traces of Lower Volga-related ancestry which implies an origin north of the Caucasus (Fig. 2f; Extended Data Fig. 1). On the steppe side, we see that mixed Lower Volga/Caucasus Neolithic ancestry was present in the Dnipro Cline and maximized in the Yamnaya population along that cline (Fig. 2e). IBD analysis identifies long (≥30cM) segments shared by Eneolithic individuals from Berezhnovka-2 in the Lower Volga with Khvalynsk, Igren-8 Serednii Stih, and Areni-1 Armenian Chalcolithic populations, providing strong direct evidence for the impact of Lower Volga ancestry on the Middle Volga, Dnipro, and South Caucasus regions, and active gene flow among these regions around the time the sampled individuals lived (Extended Data Table 5). The individual from Vonyucka-1 in the North Caucasus, in fact, has an IBD link (15.2cM) with an early Bronze Age Anatolian from Ovaören. Indo-Anatolian languages must have been spread widely by people carrying CLV cline ancestry (Fig. 2) >4000BCE. However, only two descendant groups transmitted their languages to later groups: the Yamnaya in the Dnipro-Don area, aided by the mobility of their horse-wagon technology, and the Proto-Anatolians in the south, surviving in the diverse linguistic landscape of ancient Western Asia long enough for their languages to be recorded in writing after 2000BCE. Whatever their deeper origins in time out of the diverse constituents of CLV cline populations, the Indo-Anatolians must have been part of that cline. Genetics has little to say whether within this cline the IA languages were first spoken in the Caucasus end of the cline and spread into the steppe along with the spread of Caucasus ancestry, or vice versa, or even if a linguistic unity uncoupled with ancestry existed within the CLV continuum. DNA has traced back the ancestors of both Anatolian and IE speakers to the part of the CLV Cline that was north of the Caucasus mountains, bringing them into proximity with each other and uncovering their common CLV ancestry. However, it cannot adjudicate, on its own, who among the proximate and diverse distal ancestors of the CLV people were Pre-IA speaking. Future studies of the dynamics and temporality of intra-CLV contacts (to which genetics may add its information) and of the cultures of CLV people (as reconstructed by archaeology and linguistics) may decide who among them were most likely to have been the “original” Indo-Anatolians.

Linguistic evidence has been advanced in favor of different solutions of the Proto-IE origins problem for more than two centuries and we review some recent proposals relevant to our reconstruction of early IA/IE history.

First, the presence of some cereal terminology in IA languages and even more in IE was suggested to reflect a subsistence strategy that relied in part on agriculture; this was interpreted as providing evidence against a geographic origin of the populations that spread Indo-European languages east of the Dnipro valley, the easternmost point in which agriculture was used (along with foraging and herding) during the Eneolithic.^65^ Our genetic findings are consistent with this constraint. If a Caucasus Neolithic population like that at Aknashen spread IA languages to the north (via the CLV cline to the Dnipro-Don area) it would almost certainly have had a cereal vocabulary, and then this vocabulary would have been retained during the Serednii Stih culture of the Eneolithic down to the time of the Yamnaya as agriculture continued to be used there.^65^

Second, the fact that Anatolian languages are attested largely in western Anatolia has been interpreted as evidence for entry into Anatolia from the west (via the Balkans),^49^ and thus we need compelling genetic evidence to provide a strong synthetic case for an eastern route. In fact, however, our genetic data does provide such a strong case, greatly increasing the plausibility of scenarios of an eastern entry of Proto-Anatolian speaking ancestors into Anatolia.^66^ This is because we find that Central Anatolian Early Bronze Age people who were plausibly speakers of Anatolian languages based on their archaeological contexts, were striking genetic outliers from their neighbors due to having a minority component of their ancestry from the CLV (plausibly from the people who brought the ancestral form of Anatolian languages to Anatolia), the majority of their ancestry from Mesopotamian Neolithic farmers, and little or no ancestry from the Neolithic and Chalcolithic Anatolians who were overwhelming the source populations of other Early Bronze Age Anatolians. Mesopotamian Neolithic ancestry almost certainly had an eastern geographic distribution, while the Central Anatolian Bronze Age people had no evidence of the European farmer or European hunter-gatherer ancestry that CLV have encountered if they had migrated to Anatolia from the west, so the genetic data favor an eastern route. How then could it be that there is no linguistic evidence of Anatolian speakers in eastern Anatolia? We propose that the archaeologically momentous expansion of the Kura-Araxes archaeological culture in the Caucasus and eastern Anatolia after around 3000BCE may have driven a wedge between steppe and West Asian speakers of IA languages, isolating them from each other and perhaps explaining their survival in western Anatolia into recorded history. That the expansion of the Kura-Araxes archaeological culture could have had a profound enough demographic impact to have pushed out Anatolian-speakers, is attested by genetic evidence showing that in Armenia, the spread of the Kura-Araxes culture was accompanied by the complete disappearance of CLV ancestry that had appeared there in the Chalcolithic (Fig. 2f).^9,45,67^

The Kura-Araxes culture may not be the only reason for the IA split. The ancestors of the Yamnaya did not only become separated from their Anatolian linguistic relatives but from other steppe populations as well. The homogenization of the Yamnaya ancestral population during the 4^th^ millennium BCE, both in terms of its autosomal ancestry, and in terms of its Y-chromosome lineage, attest to a period of relative isolation and the cessation of admixture. Such isolation would foster linguistic divergence of the languages spoken in the pre-Yamnaya community with those of their linguistic relatives on the steppe. This isolation must have persisted even after the sudden appearance of the Yamnaya archaeological horizon. Mobility and geographical dispersal provided ample opportunities for the resumption of admixture, yet the genetic homogeneity of the “Core Yamnaya” across much of the steppe leaves little room for the absorption of any pre-existing steppe communities: they all seem to disappear in the face of the Yamnaya juggernaut. Did mixing occur between the segment of the Yamnaya population not buried in kurgans and locals they encountered while the kurgan-buried elite largely avoided it with some exceptions?^15^ The rise of the Yamnaya in the Steppe at the expense of their predecessors was followed by their demise after a thousand years (Fig. 3), displaced by descendants of people of the Corded Ware culture. Was this the demise of the kurgan elites of the Yamnaya or of the population as a whole? The steppe was dominated by many and diverse groups later still, such as the Scythians and Sarmatian nomads of the Iron Age. These groups are certainly very diverse genetically, but their kurgans scattered across the steppe attest to the persistence of at least some elements of culture that began in the Caucasus-Volga area seven thousand years ago before blooming, in the Dnipro-Don area, into the Yamnaya culture that first united the steppe and impacted most of Eurasia. To what symbolic purpose did the Yamnaya and their precursors erect these mounds we may not ever fully know. If they aimed to preserve the memory of those buried under them, they did achieve their goal, as the kurgans, dotting the landscape of the Eurasian steppe, drew generations of archaeologists and anthropologists to their study, and enabled the genetic reconstruction of their makers’ origins presented here.

## Methods

### Terminology for archaeological cultures and geographic locations

For archaeological cultures and geographic locations that span more than one modern country, we used the prevalent term in the archaeological and genetic literature, for example using “Yamnaya” which is the common term used in Russia and most of Eastern Europe instead of the Ukrainian “Yamna”. For archaeological cultures and locations that are confined to a single country, we generally use the local terminology, for example we refer to the archaeological cultures of “Usatove” and “Trypillia” and “Serednii Stih” and the river “Dnipro” with the Ukrainian terms rather than the corresponding Russian terms “Usatovo”, “Tripolye,” “Sredni Stog” and “Dniepr”.

### Sampling ancient individuals

The skeletal remains analyzed here were almost all sampled in ancient DNA clean rooms either at Harvard Medical School, the University of Vienna or the Institute for Archaeogenomics in Budapest. If available and accessible, we prioritized sampling petrous bones, taking bone powder from the cochlea by sandblasting and milling^68^, or directly drilling into the cochlea after physical surface cleaning, or drilling through the cranial base to minimize damage to intact skulls^69^. If we could not sample from the cochlea, we sought to sample a tooth, prioritizing the cementum layer after physical surface cleaning^70^. If neither a cochlea nor a tooth was available, we sought to sample a dense cortical bone, which we analyzed by drilling and collecting powder after physical surface cleaning. For some samples that could not leave the museum, we sampled on site, either drilling directly into the cochlea, the tooth root, or bone after physical surface removal. We sometimes dislodged auditory ossicles during sandblasting or drilling into the cochlea. When this happened during the cleaning procedure, we generally stopped the destructive sampling and collected the ossicle(s)^71^. As suggested in the study that recognized the high preservation of DNA in ossicles, we cleaned the ossicle with 10% Bleach and radiated it ultraviolet light for 10 minutes before submerging it in extraction buffer without attempting to produce powder.

### Ancient DNA data generation

The samples we studied were processed in our laboratories between 2013 and 2023 and therefore were analyzed with changing protocols. Details and protocols used for each library can be found in Online Table 2. At Harvard Medical School, where the majority of wet laboratory work was done, we initially carried out all DNA extractions and Illumina library preparations manually, using small batches of samples and silica columns for DNA cleanup^72–74^. Since 2018, we used automated liquid handlers (Agilent Bravo Workstations) for both DNA extraction^75^ and library preparation with magnetic beads (see supplementary material in ^76^ for automated double-stranded library preparation, and ref. ^77^ for automated single-stranded library preparation). We treated DNA extracts with USER (NEB) during library preparation to cut DNA at uracils; this treatment is inefficient at terminal uracils and leaves a damage pattern expected for ancient DNA at the terminal bases that can be filtered out for downstream analysis while allowing a library to be authenticated as old. All libraries were either dual barcoded through double-stranded ligation or dual indexed through indexing PCR at the end of single-stranded library preparation to allow pooling before sequencing.

Before 2015, we screened libraries for mitochondrial DNA before attempting to capture nuclear loci^78^. In the next couple of years, we added an increasing number of nuclear SNPs (between 10 and 4000) as targets into the screening capture since mitochondrial DNA quality does not always correlate well with nuclear DNA quality and quantity. We later increased the number of targeted SNPs in our nuclear capture from about 390,000 (390k) ^2,79^ to about 1.24 million (1240k)^80^ for libraries passing the mitochondrial capture with nuclear spike-in. Later, we dropped the screening capture altogether and added the mitochondrial probes to the 1240k probes (1240k+). In 2022, we switched from the 1240k homebrew capture to a kitted capture product available from Twist Biosciences^81^.

For ancient DNA data generated in the Budapest at the Institute of Archaeogenomics, HUN-REN Research Centre for the Humanities, we followed the protocol described in ^82^.

### Bioinformatic processing

All ancient DNA libraries were sequenced with paired-end reads. We then performed the following steps: preprocessing, alignment and post-alignment filtering for variant calling. The goal of preprocessing is to take raw sequenced products and create merged sequences for alignment. We demultiplex reads, binning these to whichever library each read belongs to using the identifying barcodes and indices, trim these identifying markers as well as any residual adapter sequences, and merge each paired-end read into a single molecule using the overlap of the paired-end reads as a guide, employing a modified version of *SeqPrep* (https://github.com/jstjohn/SeqPrep). The resulting single-ended reads are aligned to both the *hg19* human genome reference (https://www.internationalgenome.org/category/grch37/) and the inferred ancestral Reconstructed Sapiens Reference Sequence (RSRS) mitochondrial sequence^83^ using the *samse* aligner of *bwa*^84^. Duplicate molecules are marked by barcode bin, based on the same start/stop positions and orientation. The computational pipelines with specific parameters used are publicly available on GitHub at https://github.com/dReichLab/ADNA-Tools and https://github.com/dReichLab/adna-workflow.

We used a ‘pseudohaploid genotyping’ approach to determine a randomly selected allele at SNP sets of interest. To represent the allele at each SNP, we randomly selected sequences from a pool of all sequences covering that position with a minimum data quality; our criteria were a minimum mapping quality of at least 10, and a base quality of at least 20, after trimming sequences by 2 base pairs at both 5’ and 3’ ends to remove damage artifacts. We assessed ancient DNA authenticity by using *contamMix-1.0.1051*^85^ to search for heterogeneity in mitochondrial DNA sequences which are expected to be non-variable in uncontaminated individuals, and also ANGSD to teset for heterogeneity in X chromosome sequences which are expected to be homozygous in male individuals.^86^ We also evaluated authenticity of the ancient samples by using *pmdtools*^87^ to measure the rate of cytosine-to-thymine mutations in the first and last nucleotides (in untrimmed sequences) which is expected for genuine ancient DNA^73^, and by computing the ratio of Y chromosome to the sum of X and Y chromosome sequences which is expected to be very low for females and to have a very much higher value for males. We determined a consensus for mitochondrial DNA using *bcftools* (https://github.com/samtools/bcftools) and *SAMTools*^88^ requiring a minimum of 2-fold coverage to call the nucleotide and a majority rule to determine its value. We used *HaploGrep2* to determine mitochondrial haplogroups based on the phylotree database (mtDNA tree build 17).^89,90^

### Principal Components Analysis

Individuals in Fig. 1b are projected analysis in *smartpca*^37^ using parameters newshrink: YES and lsqporject: YES: on a PCA space whose axes are formed by the following set of populations: OberkasselCluster (set of trans-Alpine WHG individuals identified in^19^), Russia_Firsovo_N, Iran_HajjiFiruz_C^4^, Iran_C_SehGabi^9^, Iran_C_TepeHissar^91^, Israel_C^92^, Germany_EN_LBK^2,17,82,93^

### F_ST_ estimation

F_ST_ was computed in *smartpca*^37^ with parameters inbreed: YES and fstonly: YES.^94^

### Visualizing the three Eneolithic Clines

Three models are fitted for Eneolithic cline populations using qpAdm^2^ and with OldAfrica, Russia_AfontovaGora3, CHG, Iran_GanjDareh_N, Italy_Villabruna, Russia_Sidelkino.SG, Turkey_N set of Right populations (Fig. 1c).

### Model competition with qpAdm/qpWave

We use qpWave/qpAdm methods^2,30^ on diverse target and source populations from the steppe and adjacent areas (Supplementary Information section 2). We use OldAfrica, Russia_AfontovaGora3, CHG, Iran_GanjDareh_N, Italy_Villabruna, Russia_Sidelkino.SG, Turkey_N as the set of Right populations for most analyses. For the analysis of Anatolian populations, we expanded this set to OldAfrica, CHG, Iran_GanjDareh_N, Italy_Villabruna, Russia_AfontovaGora3, Russia_Sidelkino.SG, TUR_Marmara_Barcın_N, TUR_C_Boncuklu_PPN, TUR_C_Çatalhöyük_N, Natufian to gain leverage for differentiating between different West Asian sources. For faster computation, we ran qpWave/qpAdm on precomputed output from qpfstats runs (https://github.com/DReichLab/AdmixTools/blob/master/qpfs.pdf) with poplistname that includes Han.DG, and all target, source, and Right populations, and parameters allsnps: YES, inbreed: NO. Separate qpWave/qpAdm runs directly on genotype files were performed as needed when the target or source populations were not present in the qpfstats output with parameter basepop: Han.DG. Feasible models are identified as having p>0.05, all standard errors ≤0.1, and admixture proportions within ≤2 standard errors from 0 and 1. Target or source populations are removed from the Right set. Competition of models A and B involves two qpWave/qpAdm runs in which all sources of A \ B and B \ A (\ denotes set difference) are placed on the Right set. Details of all analyses can be found in Supplementary Information section 2.

### Y-chromosome haplogroup inference

We used the methodology described in ref. ^6^ which used the YFull YTree v. 8.09 phylogeny (https://github.com/YFullTeam/YTree/blob/master/ytree/tree_8.09.0.json) to denote Y-chromosome haplogroups in terminal notation.^95^

### Estimates of dates of admixture

We used DATES^4,52^ to estimate a date of admixture for the Core Yamnaya, Don Yamnaya, Eastern European Yamnaya, Corded Ware, and Caucasus-Anatolian populations (Extended Data Fig. 2). For the Core Yamnaya and Caucasus-Anatolian populations, we used sets of diverse West Asian and European hunter-gatherer populations as the two sources. For the Don Yamnaya we used the Core Yamnaya and UNHG as the two sources.

For the Eastern European Yamnaya we used the Core Yamnaya and a diverse set of Neolithic/Chalcolithic “European farmers” from Fig. 4b. For the Corded Ware we used the Core Yamnaya and Globular Amphora as the two sources. It is more important to use many source samples even if they are not identical to the true ones; picking the wrong sources does not bias the date estimate^52^.

### Identity-by-Descent (IBD) segment detection

We used ancIBD^54^ to detect IBD segments of length ≥8cM.

### Geographical distance estimation

To study the decay of IBD with geographical distance, we estimate distance between sites based on their latitude and longitude (Online Table 2) using the Haversine distance as implemented in distHaversine^96^ of the package *geosphere* in R.

### Estimates of effective population sizes

We ran HapNe-LD (version 1.20230726 ^18^) using default parameters and providing pseudo-haploid genotypes as input. Briefly, HapNe-LD uses a summary statistic measuring long-range correlations between markers to infer fluctuations in the effective population size (defined as the inverse of the coalescence rate) through time. We studied two distinct sets of unrelated individuals all of which had a coverage of at least 0.7x on the target autosomal SNPs and with a standard deviation on their estimated date smaller than 180 years (∼6 generations). The first group consists of 25 Core Yamnaya individuals with estimated dates ranging between 4500 and 4800 BP. The second group contains 26 Core Yamnaya individuals ranging from 4800 to 5100 BP.

If no evidence of effective population size fluctuations can be detected in the data, HapNe-LD produces a flat line. An output containing fluctuations should thus be interpreted as the detection of changes in historical effective population sizes. Recent admixture between highly differentiated populations (Fst > 0.1) might lead to biases in LD-based analyses that induce fluctuations similar to a population bottleneck. However, HapNe implements a test to flag the presence of recent structure in the data, which was not detected in both sample sets (approximate p>=0.1), suggesting that the observed signal instead reflects variation in the effective population size of these groups.

In our analyses, the effective population size is defined as the inverse of the instantaneous coalescence rate. This quantity corresponds to twice the number of breeding individuals in an idealized population. We note that, in addition to changes in the number of individuals in the population (census size), several factors, such as changes in population structure, selection, and cultural practices,^97^ can have an influence on the effective population size. These additional factors may in part be responsible for the effective size fluctuations observed in the Core Yamnaya.

Approximate confidence intervals were obtained using bootstrap with different chromosome arms as resampling units. The beginning of the expansion was determined by using the location of the minimum of each bootstrapped trajectory. We converted the results into years by assuming 28.6 years per generation for the median minimum location and 25.6 and 31.5 years per generation for the lower and upper bounds, respectively.^98^ We used these values, corresponding to the estimated number of years per generation for males (31.5) and females (25.6) to account for uncertainty in the conversion factor.

## Data Access

Genotype data for individuals included in this study can be obtained from the Harvard Dataverse repository through the following link (XXX).The DNA sequences reported in this paper are deposited in the European Nucleotide Archive under the accession number XXX. Other newly reported data such as radiocarbon dates and archaeological context information are included in the manuscript and supplementary files.

## Acknowledgments

We thank Alexey G. Nikitin for valuable advice and critical feedback. We thank Nicole Adamski, Rebecca Bernardos, Nasreen Broomandkhoshbacht, Daniel Fernandes, Matthew Ferry, Eadaoin Harney, Kirsten Mandl, Susanne Nordenfelt, Kristin Stewardson, Balázs G. Mende, and Zhao Zhang for laboratory or bioinformatics work, and L’ubov Bembeeva, Bianca Preda-Bălănică, István Ecsedy, Andrey I. Gotlib, Volker M. Heyd, Skorobogatov Andrey Mikhailovich, Nina Morgunova, Andrei Soficaru, Svetlana S. Tur, and Piotr Włodarczak for anthropological work and critical comments. TH’s research was supported by a grant from the Hungarian Research, Development and Innovation Office (FK128013), the Bolyai Scholarship of the Hungarian Academy of Sciences, and by the ÚNKP-23-5 New National Excellence Program of the Ministry for Culture and Innovation from the source of the National Research, Development and Innovation Fund. Pavel Flegontov and Leonid Vyazov were supported by the Czech Ministry of Education, Youth and Sports (program ERC CZ, project no. LL2103). Pavel Flegontov was supported by the Czech Science Foundation (project no. 21-27624S); the European Union Operational Programme "Just Transition" (LERCO project no. CZ.10.03.01/00/22_003/0000003). We acknowledge support from the Polish scientific project grant NCN OPUS 2015/17/B/HS3/01327, as well as Russian Science Foundatino grant #22-18-00470 to Alexey A. Tishkin. We acknowledge support from the Museum of the Institute of Plant and Animal Ecology (UB RAS, Ekaterinburg) to Pavel Kosintsev. The ancient DNA data generation and analysis was supported by the National Institutes of Health (R01-HG012287), the John Templeton Foundation (grant 61220), by a private gift from Jean-Francois Clin, by the Allen Discovery Center program, a Paul G. Allen Frontiers Group advised program of the Paul G. Allen Family Foundation and by the Howard Hughes Medical Institute (DR). The author-accepted version of this article, that is, the version not reflecting proofreading and editing and formatting changes following the article’s acceptance, is subject to the Howard Hughes Medical Institute (HHMI) Open Access to Publications policy, as HHMI lab heads have previously granted a nonexclusive CC BY 4.0 license to the public and a sublicensable license to HHMI in their research articles. Pursuant to those licenses, the author-accepted manuscript can be made freely available under a CC BY 4.0 license immediately upon publication.,

## Author Contributions

I.L., N.P., D.A., L.V., and D.R. wrote the manuscript and supplementary materials with input from all co-authors. A.S.-N., P.F.P., S.M., N.R., R.P. and D.R. supervised different aspects of the study. I.L. and N.P. carried out the main genetic analyses. I.L., R.F., H.R., I.O., P.F.P. analyzed genetic data. D.A. and L.V. edited archaeological information. D.A., A.Kh., E.K., N.Sh., S.C.A., S.V.A., E.Ba., Z.B., A.B., P.C., A.C., I.C., M.Co., M.Cs, J.D., N.E., S. É, A.Fa., P.F., A.Fr., M.F., A.G., T.Ha., T.Hi., P.Je., V.Kh., V.Ki., A.Kit., A.Ko., G.K., N.N., B.B., P.D., P.Ja., E.Ki., A.Kiy., J.K., P.K., R.M., M.M., E.Me., V.M., E.Mo., J.M., N.M., M.N., O.N., M.R., M.O-G., G.P., S.P., T.S., N.Se., A. Š, M.R.S., I.S., V.Sh., A.Si., K.S., K.N.S., J.T., A.A.T., V.T., S.V., O.F., A.Sz-N., and R.P. sampled anthropological remains and/or contributed to the creation of the archaeological supplement. A.A., E.S.B., M.Ma., A.Mi., and S.M. carried out bioinformatic data processing. K.C., F.C., O.Ch., E.C., L.I., A.Ke., D.K., A.M.L., M.Mi., J.O., L.Q., J.N.W., F.Z., N.R., and carried out wet laboratory work.

## Ethics Statement

The individuals studied here were all analyzed with the goal of minimizing damage to their skeletal remains, with permission from local authorities in each location from which they came. Every sample is represented by stewards such as archaeologists or museum curators, who are either authors or thanked in the Acknowledgments. Open science principles require making all data used to support the conclusions of a study maximally available, and we support these principles here by making fully publicly available not only the digital copies of molecules (the uploaded sequences) but also the molecular copies (the ancient DNA libraries themselves, which constitute molecular data storage). Those researchers who wish to carry out deeper sequencing of libraries published in this study should make a request to corresponding author D.R. We commit to granting reasonable requests as long as the libraries remain preserved in our laboratories, with no requirement that we be included as collaborators or co-authors on any resulting publications.

---

**Extended Data Table 1:** F_ST_ values among select populations of the Dnipro, Don, Volga, and Caucasus areas. F_ST_ values are shown below the diagonal and their standard errors above it.

|  | BPgroup | CoreYamnaya | Ekaterinovka | GK1 | Khi | KhlopkovBugor | Klo | Kmed | Labazy | Maikop | Maximovka | Murzikha | PVgroup | Remontnoye | Russia_Caucasus_LateMaikop | Russia_Don_EBA_Yamnaya | SShi | SSmed | Syezzheye | Ukraine_N | Unakozovskaya | UpperVolga |
| --- | --- | --- | --- | --- | --- | --- | --- | --- | --- | --- | --- | --- | --- | --- | --- | --- | --- | --- | --- | --- | --- | --- |
| BPgroup |  | 0.001 | 0.001 | 0.003 | 0.001 | 0.002 | 0.001 | 0.001 | 0.002 | 0.001 | 0.002 | 0.001 | 0.002 | 0.002 | 0.005 | 0.001 | 0.001 | 0.001 | 0.003 | 0.001 | 0.003 | 0.001 |
| CoreYamnaya | 0.011 |  | 0.000 | 0.003 | 0.000 | 0.002 | 0.000 | 0.001 | 0.002 | 0.001 | 0.002 | 0.001 | 0.002 | 0.001 | 0.004 | 0.000 | 0.001 | 0.001 | 0.002 | 0.001 | 0.002 | 0.000 |
| Ekaterinovka | 0.030 | 0.032 |  | 0.003 | 0.001 | 0.002 | 0.000 | 0.001 | 0.002 | 0.001 | 0.002 | 0.001 | 0.002 | 0.002 | 0.004 | 0.000 | 0.001 | 0.001 | 0.003 | 0.001 | 0.003 | 0.000 |
| GK1 | 0.042 | 0.041 | 0.045 |  | 0.003 | 0.007 | 0.003 | 0.003 | 0.005 | 0.004 | 0.006 | 0.003 | 0.005 | 0.005 | 0.018 | 0.003 | 0.004 | 0.005 | 0.009 | 0.003 | 0.006 | 0.003 |
| Khi | 0.007 | 0.014 | 0.019 | 0.039 |  | 0.002 | 0.001 | 0.001 | 0.002 | 0.001 | 0.002 | 0.001 | 0.002 | 0.002 | 0.004 | 0.001 | 0.001 | 0.001 | 0.003 | 0.001 | 0.003 | 0.001 |
| KhlopkovBugor | 0.010 | 0.017 | 0.022 | 0.037 | 0.008 |  | 0.002 | 0.002 | 0.003 | 0.003 | 0.003 | 0.002 | 0.003 | 0.003 | 0.009 | 0.002 | 0.003 | 0.003 | 0.005 | 0.002 | 0.004 | 0.002 |
| Klo | 0.018 | 0.022 | 0.008 | 0.041 | 0.009 | 0.013 |  | 0.001 | 0.002 | 0.001 | 0.002 | 0.001 | 0.002 | 0.002 | 0.004 | 0.001 | 0.001 | 0.001 | 0.003 | 0.001 | 0.003 | 0.001 |
| Kmed | 0.014 | 0.018 | 0.015 | 0.042 | 0.006 | -0.002 | 0.002 |  | 0.002 | 0.001 | 0.002 | 0.001 | 0.002 | 0.002 | 0.005 | 0.001 | 0.001 | 0.001 | 0.003 | 0.001 | 0.003 | 0.001 |
| Labazy | 0.032 | 0.034 | 0.009 | 0.048 | 0.021 | 0.027 | 0.010 | 0.016 |  | 0.002 | 0.003 | 0.002 | 0.003 | 0.003 | 0.007 | 0.002 | 0.002 | 0.003 | 0.004 | 0.002 | 0.004 | 0.002 |
| Maikop | 0.031 | 0.025 | 0.064 | 0.064 | 0.037 | 0.043 | 0.052 | 0.045 | 0.067 |  | 0.003 | 0.001 | 0.002 | 0.002 | 0.006 | 0.001 | 0.002 | 0.002 | 0.003 | 0.001 | 0.003 | 0.001 |
| Maximovka | 0.044 | 0.041 | 0.021 | 0.048 | 0.033 | 0.033 | 0.021 | 0.028 | 0.021 | 0.076 |  | 0.002 | 0.003 | 0.003 | 0.007 | 0.002 | 0.003 | 0.003 | 0.004 | 0.002 | 0.003 | 0.002 |
| Murzikha | 0.056 | 0.053 | 0.034 | 0.065 | 0.044 | 0.047 | 0.034 | 0.039 | 0.034 | 0.088 | 0.018 |  | 0.002 | 0.002 | 0.004 | 0.001 | 0.001 | 0.001 | 0.003 | 0.001 | 0.003 | 0.001 |
| PVgroup | -0.002 | 0.012 | 0.035 | 0.046 | 0.010 | 0.012 | 0.024 | 0.018 | 0.038 | 0.025 | 0.048 | 0.061 |  | 0.003 | 0.006 | 0.002 | 0.002 | 0.003 | 0.004 | 0.002 | 0.003 | 0.002 |
| Remontnoye | 0.012 | 0.011 | 0.040 | 0.041 | 0.015 | 0.020 | 0.028 | 0.024 | 0.046 | 0.012 | 0.052 | 0.065 | 0.011 |  | 0.006 | 0.002 | 0.002 | 0.002 | 0.004 | 0.002 | 0.003 | 0.002 |
| Russia_Caucasus_LateMaikop | 0.025 | 0.020 | 0.058 | 0.065 | 0.033 | 0.037 | 0.048 | 0.041 | 0.059 | -0.001 | 0.063 | 0.081 | 0.026 | 0.002 |  | 0.004 | 0.006 | 0.007 | 0.011 | 0.004 | 0.007 | 0.004 |
| Russia_Don_EBA_Yamnaya | 0.014 | 0.005 | 0.029 | 0.040 | 0.014 | 0.019 | 0.019 | 0.018 | 0.030 | 0.030 | 0.037 | 0.048 | 0.016 | 0.015 | 0.025 |  | 0.001 | 0.001 | 0.003 | 0.001 | 0.003 | 0.001 |
| SShi | 0.009 | 0.011 | 0.027 | 0.034 | 0.013 | 0.014 | 0.017 | 0.017 | 0.029 | 0.030 | 0.036 | 0.048 | 0.010 | 0.016 | 0.034 | 0.011 |  | 0.002 | 0.004 | 0.001 | 0.003 | 0.001 |
| SSmed | 0.011 | 0.010 | 0.021 | 0.034 | 0.011 | 0.012 | 0.015 | 0.015 | 0.021 | 0.030 | 0.030 | 0.041 | 0.013 | 0.014 | 0.019 | 0.008 | 0.004 |  | 0.004 | 0.001 | 0.003 | 0.001 |
| Syezzheye | 0.045 | 0.047 | 0.022 | 0.059 | 0.034 | 0.035 | 0.026 | 0.033 | 0.029 | 0.082 | 0.043 | 0.050 | 0.049 | 0.056 | 0.077 | 0.042 | 0.040 | 0.034 |  | 0.003 | 0.004 | 0.003 |
| Ukraine_N | 0.046 | 0.039 | 0.036 | 0.047 | 0.040 | 0.042 | 0.032 | 0.037 | 0.036 | 0.063 | 0.038 | 0.048 | 0.049 | 0.049 | 0.055 | 0.029 | 0.031 | 0.017 | 0.055 |  | 0.003 | 0.001 |
| Unakozovskaya | 0.059 | 0.057 | 0.094 | 0.090 | 0.068 | 0.069 | 0.083 | 0.076 | 0.096 | 0.034 | 0.107 | 0.117 | 0.058 | 0.039 | 0.030 | 0.060 | 0.062 | 0.061 | 0.107 | 0.092 |  | 0.003 |
| UpperVolga | 0.044 | 0.040 | 0.021 | 0.048 | 0.033 | 0.035 | 0.019 | 0.028 | 0.019 | 0.073 | 0.015 | 0.027 | 0.049 | 0.051 | 0.067 | 0.033 | 0.035 | 0.026 | 0.038 | 0.029 | 0.103 |  |

**Extended Data Table 2:**
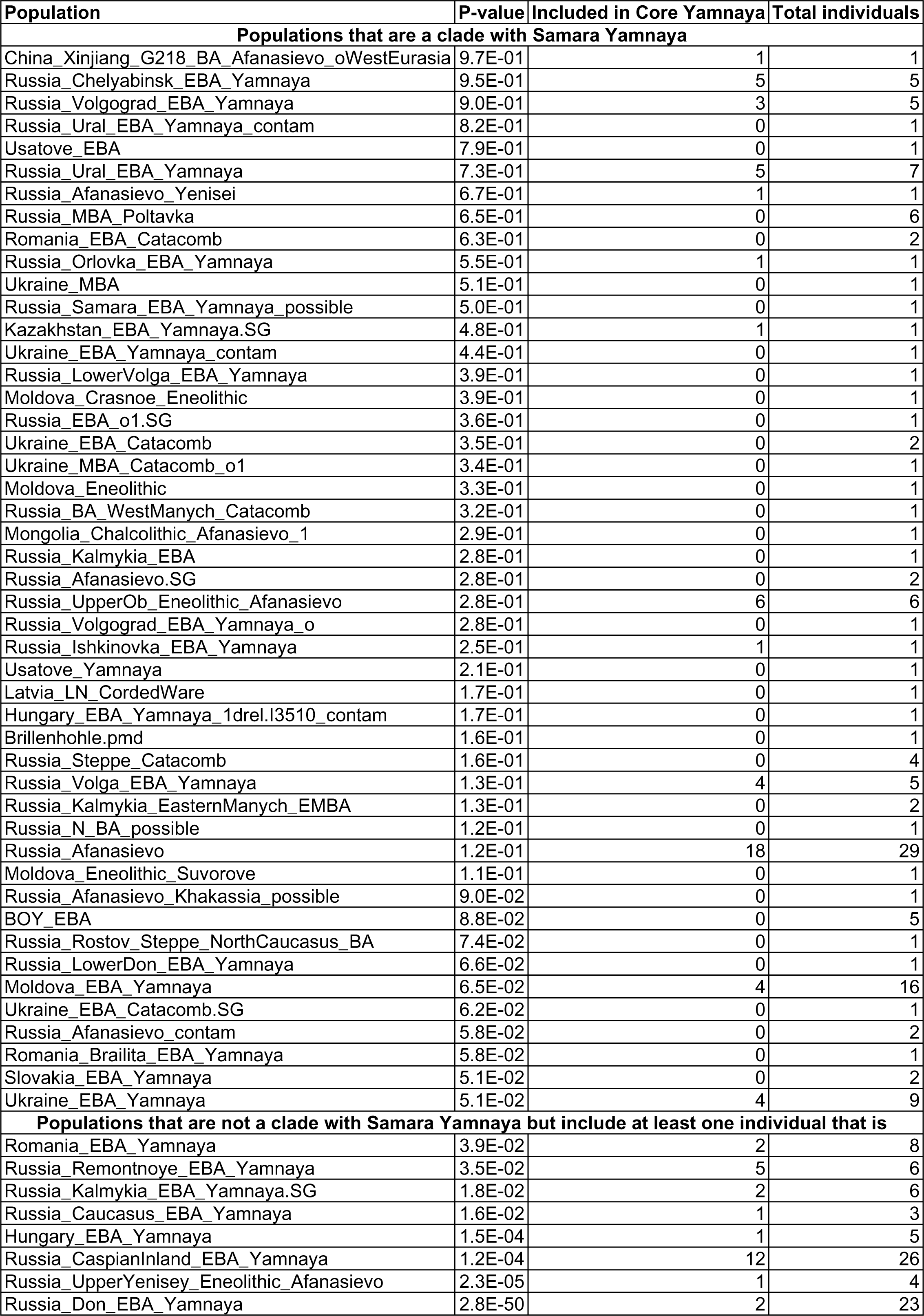
Extraordinary Genetic Homogeneity in the Core Yamnaya. We tested all populations and individuals for cladality with Samara Yamnaya. We list populations for which this is not rejected (p>0.05) and populations that include individuals that fit Core Yamnaya selection criteria (p>0.2, at least 300k SNPs, and Yamnaya or Afanasievo culture).

| Population | P-value | Included in Core Yamnaya | Total individuals |
| --- | --- | --- | --- |
| **Populations that are a clade with Samara Yamnaya** |  |  |  |
| China_Xinjiang_G218_BA_Afanasievo_oWestEurasia | 9.7E-01 | 1 | 1 |
| Russia_Chelyabinsk_EBA_Yamnaya | 9.5E-01 | 5 | 5 |
| Russia_Volgograd_EBA_Yamnaya | 9.0E-01 | 3 | 5 |
| Russia_Ural_EBA_Yamnaya_contam | 8.2E-01 | 0 | 1 |
| Usatove_EBA | 7.9E-01 | 0 | 1 |
| Russia_Ural_EBA_Yamnaya | 7.3E-01 | 5 | 7 |
| Russia_Afanasievo_Yenisei | 6.7E-01 | 1 | 1 |
| Russia_MBA_Poltavka | 6.5E-01 | 0 | 6 |
| Romania_EBA_Catacomb | 6.3E-01 | 0 | 2 |
| Russia_Orlovka_EBA_Yamnaya | 5.5E-01 | 1 | 1 |
| Ukraine_MBA | 5.1E-01 | 0 | 1 |
| Russia_Samara_EBA_Yamnaya_possible | 5.0E-01 | 0 | 1 |
| Kazakhstan_EBA_Yamnaya.SG | 4.8E-01 | 1 | 1 |
| Ukraine_EBA_Yamnaya_contam | 4.4E-01 | 0 | 1 |
| Russia_LowerVolga_EBA_Yamnaya | 3.9E-01 | 0 | 1 |
| Moldova_Crasnoe_Eneolithic | 3.9E-01 | 0 | 1 |
| Russia_EBA_o1.SG | 3.6E-01 | 0 | 1 |
| Ukraine_EBA_Catacomb | 3.5E-01 | 0 | 2 |
| Ukraine_MBA_Catacomb_o1 | 3.4E-01 | 0 | 1 |
| Moldova_Eneolithic | 3.3E-01 | 0 | 1 |
| Russia_BA_WestManych_Catacomb | 3.2E-01 | 0 | 1 |
| Mongolia_Chalcolithic_Afanasievo_1 | 2.9E-01 | 0 | 1 |
| Russia_Kalmykia_EBA | 2.8E-01 | 0 | 1 |
| Russia_Afanasievo.SG | 2.8E-01 | 0 | 2 |
| Russia_UpperOb_Eneolithic_Afanasievo | 2.8E-01 | 6 | 6 |
| Russia_Volgograd_EBA_Yamnaya_o | 2.8E-01 | 0 | 1 |
| Russia_Ishkinovka_EBA_Yamnaya | 2.5E-01 | 1 | 1 |
| Usatove_Yamnaya | 2.1E-01 | 0 | 1 |
| Latvia_LN_CordedWare | 1.7E-01 | 0 | 1 |
| Hungary_EBA_Yamnaya_1drel.I3510_contam | 1.7E-01 | 0 | 1 |
| Brillenhohle.pmd | 1.6E-01 | 0 | 1 |
| Russia_Steppe_Catacomb | 1.6E-01 | 0 | 4 |
| Russia_Volga_EBA_Yamnaya | 1.3E-01 | 4 | 5 |
| Russia_Kalmykia_EasternManych_EMBA | 1.3E-01 | 0 | 2 |
| Russia_N_BA_possible | 1.2E-01 | 0 | 1 |
| Russia_Afanasievo | 1.2E-01 | 18 | 29 |
| Moldova_Eneolithic_Suvorove | 1.1E-01 | 0 | 1 |
| Russia_Afanasievo_Khakassia_possible | 9.0E-02 | 0 | 1 |
| BOY_EBA | 8.8E-02 | 0 | 5 |
| Russia_Rostov_Steppe_NorthCaucasus_BA | 7.4E-02 | 0 | 1 |
| Russia_LowerDon_EBA_Yamnaya | 6.6E-02 | 0 | 1 |
| Moldova_EBA_Yamnaya | 6.5E-02 | 4 | 16 |
| Ukraine_EBA_Catacomb.SG | 6.2E-02 | 0 | 1 |
| Russia_Afanasievo_contam | 5.8E-02 | 0 | 2 |
| Romania_Brailita_EBA_Yamnaya | 5.8E-02 | 0 | 1 |
| Slovakia_EBA_Yamnaya | 5.1E-02 | 0 | 2 |
| Ukraine_EBA_Yamnaya | 5.1E-02 | 4 | 9 |
| **Populations that are not a clade with Samara Yamnaya but include at least one individual that is** |  |  |  |
| Romania_EBA_Yamnaya | 3.9E-02 | 2 | 8 |
| Russia_Remontnoye_EBA_Yamnaya | 3.5E-02 | 5 | 6 |
| Russia_Kalmykia_EBA_Yamnaya.SG | 1.8E-02 | 2 | 6 |
| Russia_Caucasus_EBA_Yamnaya | 1.6E-02 | 1 | 3 |
| Hungary_EBA_Yamnaya | 1.5E-04 | 1 | 5 |
| Russia_CaspianInland_EBA_Yamnaya | 1.2E-04 | 12 | 26 |
| Russia_UpperYenisey_Eneolithic_Afanasievo | 2.3E-05 | 1 | 4 |
| Russia_Don_EBA_Yamnaya | 2.8E-50 | 2 | 23 |

**Extended Data Table 3:** F_ST_ values among populations that include Core Yamnaya individuals. F_ST_ values are shown below the diagonal and their standard errors above it.

|  | Hungary_EBA_Yamnaya | Moldova_EBA_Yamnaya | Romania_EBA_Yamnaya | Russia_Afnasievo | Russia_CaspianInland_EBA_Yamnaya | Russia_Caucasus_EBA_Yamnaya | Russia_Chelyabinsk_EBA_Yamnaya | Russia_Don_EBA_Yamnaya | Russia_Kalmykia_EBA_Yamnaya.SG | Russia_Remontnoye_EBA_Yamnaya | Russia_Samara_EBA_Yamnaya | Russia_UpperOb_Eneolithic_Afnasievo | Russia_UpperYenisey_Eneolithic_Afnasievo | Russia_Ural_EBA_Yamnaya | Russia_Volga_EBA_Yamnaya | Russia_Volgograd_EBA_Yamnaya | Ukraine_EBA_Yamnaya |
| --- | --- | --- | --- | --- | --- | --- | --- | --- | --- | --- | --- | --- | --- | --- | --- | --- | --- |
| Hungary_EBA_Yamnaya |  | 0.001 | 0.001 | 0.001 | 0.001 | 0.002 | 0.001 | 0.001 | 0.001 | 0.001 | 0.001 | 0.001 | 0.001 | 0.001 | 0.001 | 0.001 | 0.001 |
| Moldova_EBA_Yamnaya | 0.001 |  | 0.001 | 0.000 | 0.000 | 0.001 | 0.001 | 0.000 | 0.001 | 0.001 | 0.000 | 0.001 | 0.001 | 0.001 | 0.001 | 0.001 | 0.001 |
| Romania_EBA_Yamnaya | 0.001 | 0.001 |  | 0.001 | 0.001 | 0.002 | 0.001 | 0.001 | 0.001 | 0.001 | 0.001 | 0.001 | 0.001 | 0.001 | 0.001 | 0.001 | 0.001 |
| Russia_Afnasievo | 0.006 | 0.004 | 0.005 |  | 0.000 | 0.001 | 0.001 | 0.000 | 0.001 | 0.001 | 0.000 | 0.001 | 0.001 | 0.001 | 0.001 | 0.001 | 0.001 |
| Russia_CaspianInland_EBA_Yamnaya | 0.004 | 0.003 | 0.002 | 0.006 |  | 0.001 | 0.001 | 0.000 | 0.001 | 0.001 | 0.000 | 0.001 | 0.001 | 0.001 | 0.001 | 0.001 | 0.001 |
| Russia_Caucasus_EBA_Yamnaya | 0.001 | 0.002 | 0.001 | 0.003 | 0.003 |  | 0.002 | 0.001 | 0.002 | 0.002 | 0.001 | 0.002 | 0.002 | 0.002 | 0.002 | 0.002 | 0.002 |
| Russia_Chelyabinsk_EBA_Yamnaya | 0.008 | 0.009 | 0.009 | 0.010 | 0.009 | 0.009 |  | 0.001 | 0.001 | 0.001 | 0.001 | 0.001 | 0.001 | 0.001 | 0.001 | 0.001 | 0.001 |
| Russia_Don_EBA_Yamnaya | 0.006 | 0.005 | 0.006 | 0.008 | 0.006 | 0.005 | 0.012 |  | 0.001 | 0.001 | 0.000 | 0.001 | 0.001 | 0.001 | 0.001 | 0.001 | 0.001 |
| Russia_Kalmykia_EBA_Yamnaya.SG | 0.007 | 0.005 | 0.004 | 0.005 | 0.001 | 0.004 | 0.011 | 0.007 |  | 0.001 | 0.001 | 0.001 | 0.001 | 0.001 | 0.001 | 0.001 | 0.001 |
| Russia_Remontnoye_EBA_Yamnaya | 0.004 | 0.004 | 0.003 | 0.004 | 0.000 | 0.003 | 0.010 | 0.006 | -0.049 |  | 0.001 | 0.001 | 0.001 | 0.001 | 0.001 | 0.001 | 0.001 |
| Russia_Samara_EBA_Yamnaya | 0.003 | 0.002 | 0.003 | 0.005 | 0.003 | 0.003 | 0.008 | 0.005 | 0.005 | 0.004 |  | 0.001 | 0.001 | 0.001 | 0.001 | 0.001 | 0.001 |
| Russia_UpperOb_Eneolithic_Afnasievo | 0.006 | 0.005 | 0.004 | 0.002 | 0.003 | 0.006 | 0.010 | 0.008 | 0.001 | 0.003 | 0.004 |  | 0.001 | 0.001 | 0.001 | 0.001 | 0.001 |
| Russia_UpperYenisey_Eneolithic_Afnasievo | 0.011 | 0.010 | 0.008 | 0.009 | 0.009 | 0.009 | 0.015 | 0.012 | 0.009 | 0.007 | 0.010 | 0.006 |  | 0.001 | 0.001 | 0.001 | 0.001 |
| Russia_Ural_EBA_Yamnaya | 0.002 | 0.002 | 0.001 | 0.004 | 0.003 | 0.003 | 0.006 | 0.005 | 0.004 | 0.003 | 0.001 | 0.003 | 0.008 |  | 0.001 | 0.001 | 0.001 |
| Russia_Volga_EBA_Yamnaya | 0.004 | 0.004 | 0.003 | 0.005 | 0.005 | 0.005 | 0.007 | 0.007 | 0.008 | 0.007 | 0.003 | 0.007 | 0.011 | 0.003 |  | 0.001 | 0.001 |
| Russia_Volgograd_EBA_Yamnaya | 0.005 | 0.003 | 0.004 | 0.007 | 0.005 | 0.004 | 0.009 | 0.007 | 0.007 | 0.005 | 0.004 | 0.007 | 0.009 | 0.003 | 0.006 |  | 0.001 |
| Ukraine_EBA_Yamnaya | 0.003 | 0.001 | 0.001 | 0.004 | 0.002 | 0.002 | 0.008 | 0.005 | 0.003 | 0.003 | 0.002 | 0.004 | 0.009 | 0.001 | 0.004 | 0.004 |  |

**Extended Data Table 4:**
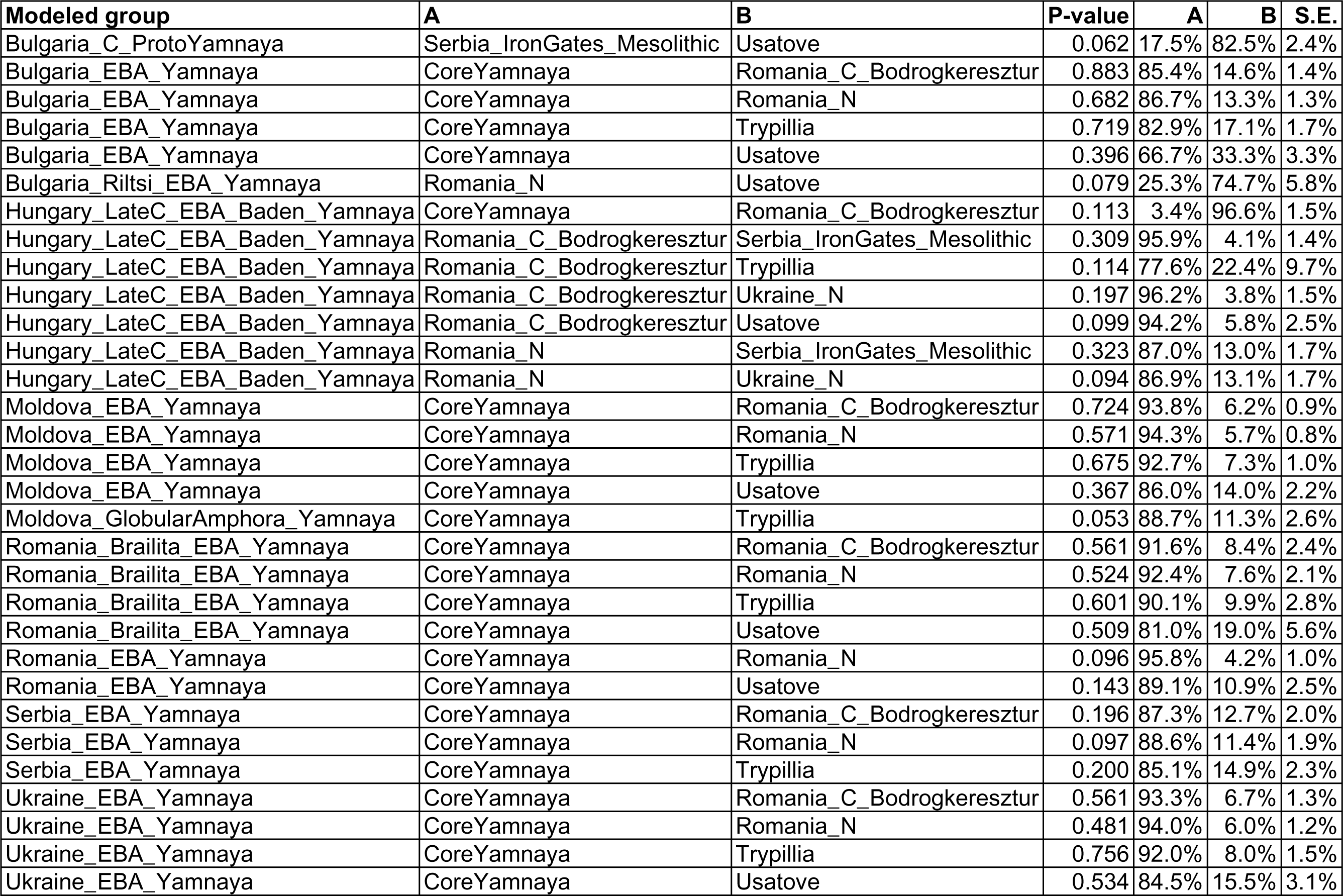
*qpAdm* models that fit non-Core Yamnaya. We use the following sources to model Yamnaya-related populations other than the Core and Don Yamnaya: CoreYamnaya, Romania_C_Bodrogkeresztur, Romania_N, Serbia_IronGates_Mesolithic, Trypillia, Ukraine_N, Usatove. The Baden individuals from Hungary represent a reburial into a kurgan^82^ and are predominantly of European farmer, not Yamnaya, ancestry. The Riltsi individual is shown with Usatove ancestry here and can also be modeled with about half Remontnoye ancestry, as the Usatove have ancestry from the CLV cline.^15^

**Extended Data Table 5:** Cross-regional shared Identity-by-Descent (IBD) segments. We list all segments≥12cM shared between individuals from two different regions defined as follows: “Dnipro cline”: CoreYamnaya, GK1, GK2, Russia_Don_EBA_Yamnaya, SShi, SSlo, SSmed, Ukraine_N; “Volga cline”: Ekaterinovka, Khi, KhlopkovBugor, Klo, Kmed, Labazy, Lebyazhinka_HG, Maximovka, Murzikha, Syezzheye, UpperVolga; “Caucasus-Lower Volga Eneolithic”: BPgroup, PVgroup; “CLV-South”: Remontnoye, Maikop, Unakozovskaya, Armenia_C, TUR_C_Kalehöyük_MLBA, TUR_C_Ovaören_EBA

| Individual 1 | Individual 2 | Group 1 | Group 2 | Segment length (cM) |
| --- | --- | --- | --- | --- |
| I22201 | I1924 | BPgroup | SShi | 35.8 |
| I22202 | I6734 | BPgroup | Khi | 32.1 |
| I1634 | I22199 | Armenia_C | BPgroup | 31.4 |
| I6300_enhanced | I22202 | KhlopkovBugor | BPgroup | 22.0 |
| I6406 | I22200 | Kmed | BPgroup | 20.1 |
| PG2004 | I11837 | BPgroup | Khi | 18.4 |
| I6301_enhanced | I22199 | KhlopkovBugor | BPgroup | 18.2 |
| I6301_enhanced | PG2001 | KhlopkovBugor | PVgroup | 17.6 |
| I28683 | PG2004 | Remontnoye | BPgroup | 16.6 |
| I10567 | I28682 | Russia_CaspianInland_EBA_Yamnaya | Remontnoye | 16.2 |
| PG2001 | I3950 | PVgroup | Russia_Afanasievo | 15.9 |
| PG2001 | I6062 | PVgroup | Ekaterinovka | 15.9 |
| I22199 | I8282 | BPgroup | Ekaterinovka | 15.8 |
| I22201 | I10208 | BPgroup | Moldova_EBA_Yamnaya | 15.5 |
| I1924 | I20188 | SShi | Klo | 15.4 |
| I32501 | I8448 | Russia_UpperYenisey_Eneolithic_Afanasievo | Murzikha | 15.4 |
| I12637 | I8457 | Moldova_EBA_Yamnaya | Murzikha | 15.4 |
| I32821 | I8449 | Russia_UpperOb_Eneolithic_Afanasievo | Murzikha | 15.4 |
| MA2213_wNonUDG.SG | VJ1001 | TUR_C_Ovaören_EBA | PVgroup | 15.2 |
| I32501 | I8455 | Russia_UpperYenisey_Eneolithic_Afanasievo | Murzikha | 15.2 |
| I6301_enhanced | I22199 | KhlopkovBugor | BPgroup | 14.9 |
| I8411_enhanced | I26785 | UpperVolga | Russia_Don_EBA_Yamnaya | 14.9 |
| I22201 | I1924 | BPgroup | SShi | 14.8 |
| I22199 | I28682 | BPgroup | Remontnoye | 14.8 |
| I0122 | I22202 | Klo | BPgroup | 14.6 |
| I32501 | I8454 | Russia_UpperYenisey_Eneolithic_Afanasievo | Murzikha | 14.5 |
| I22199 | I6734 | BPgroup | Khi | 14.5 |
| I22201 | I11752 | BPgroup | Russia_Afanasievo | 14.3 |
| I6064 | I22199 | Ekaterinovka | BPgroup | 14.2 |
| I0122 | I22199 | Klo | BPgroup | 14.2 |
| I1634 | I1924 | Armenia_C | SShi | 13.9 |
| I6301_enhanced | I22201 | KhlopkovBugor | BPgroup | 13.9 |
| I6918 | I8446 | Russia_Volgograd_EBA_Yamnaya | Maximovka | 13.9 |
| I22202 | I6739 | BPgroup | Khi | 13.9 |
| PG2004 | I23651 | BPgroup | Ekaterinovka | 13.7 |
| I0357 | I11842 | Russia_Samara_EBA_Yamnaya | Murzikha | 13.7 |
| I22202 | I3952 | BPgroup | Russia_Afanasievo | 13.7 |
| I0122 | I20190 | Klo | Russia_Samara_EBA_Yamnaya | 13.6 |
| I8951 | I11842 | Russia_Don_EBA_Yamnaya | Murzikha | 13.5 |
| PG2004 | I8290 | BPgroup | Ekaterinovka | 13.4 |
| I0231 | I8456 | Russia_Samara_EBA_Yamnaya | Murzikha | 13.4 |
| I25159 | I22199 | Russia_Afanasievo | BPgroup | 13.3 |
| I4111 | I6109 | Ukraine_N | Klo | 13.3 |
| I22199 | I26787 | BPgroup | Russia_Don_EBA_Yamnaya | 13.3 |
| I6301_enhanced | PG2004 | KhlopkovBugor | BPgroup | 12.9 |
| I8449 | I2105 | Murzikha | Ukraine_EBA_Yamnaya | 12.9 |
| I20189 | I22200 | Ekaterinovka | BPgroup | 12.8 |
| I6297 | I22201 | Russia_Orlovka_EBA_Yamnaya | BPgroup | 12.8 |
| I6705 | I28682 | Russia_Samara_EBA_Yamnaya | Remontnoye | 12.8 |
| I32821 | I22200 | Russia_UpperOb_Eneolithic_Afanasievo | BPgroup | 12.7 |
| I32501 | I8449 | Russia_UpperYenisey_Eneolithic_Afanasievo | Murzikha | 12.6 |
| I22201 | I6739 | BPgroup | Khi | 12.4 |
| I0231 | I28682 | Russia_Samara_EBA_Yamnaya | Remontnoye | 12.3 |
| PG2004 | I6739 | BPgroup | Khi | 12.3 |
| I6918 | I22200 | Russia_Volgograd_EBA_Yamnaya | BPgroup | 12.3 |
| I22201 | I3952 | BPgroup | Russia_Afanasievo | 12.2 |
| I6406 | I1450 | Kmed | Russia_Samara_EBA_Yamnaya | 12.2 |
| I22199 | I5273 | BPgroup | Russia_Afanasievo | 12.1 |
| I4114 | I12964 | Ukraine_N | UpperVolga | 12.1 |
| I11838 | I23651 | Russia_Volga_EBA_Yamnaya | Ekaterinovka | 12.0 |
| I6907 | I11841 | Russia_Samara_EBA_Yamnaya | Murzikha | 12.0 |
| I22201 | I1924 | BPgroup | SShi | 12.0 |

**Extended Data Figure 1:**
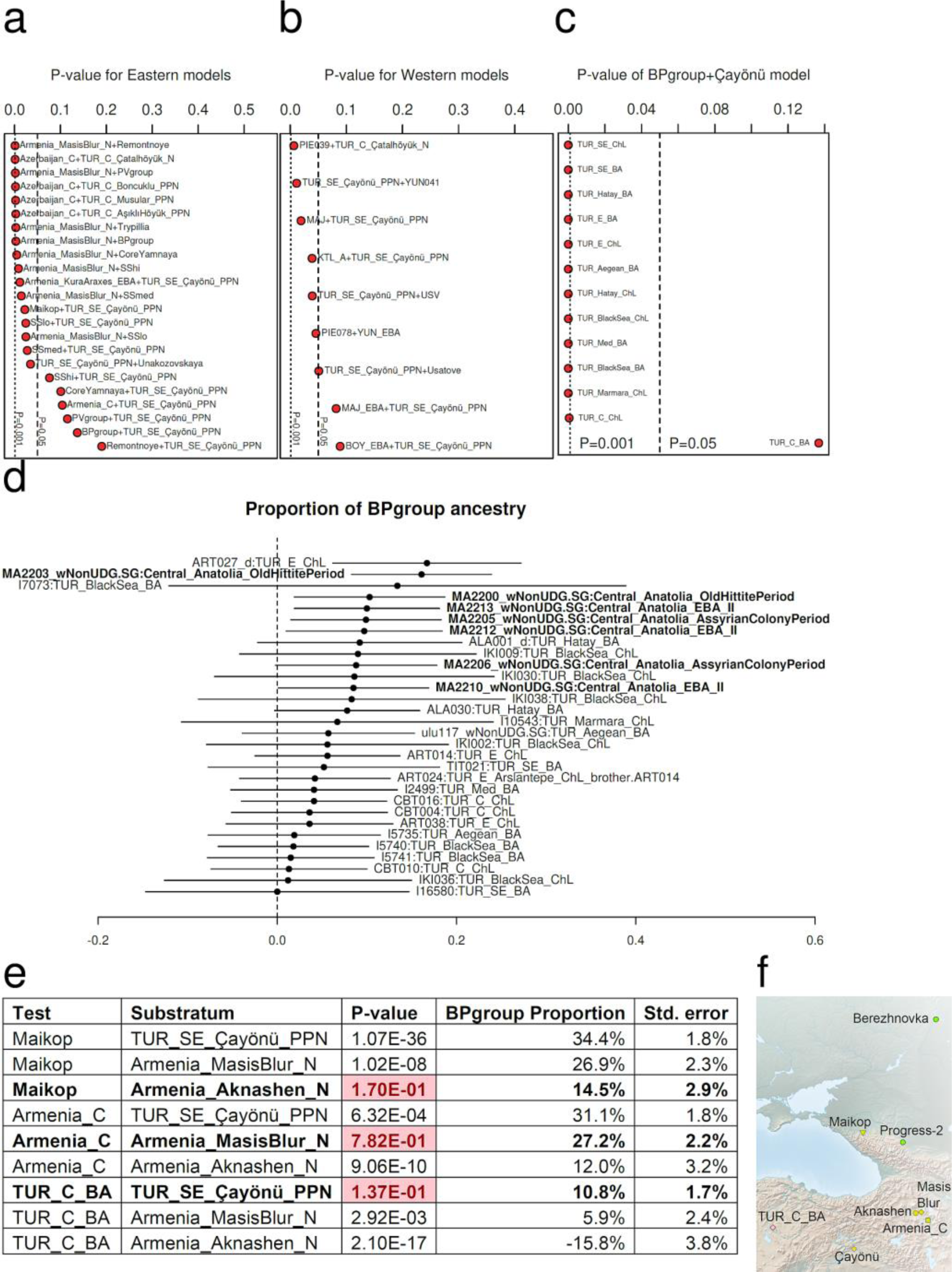
The origin of Central Anatolian Bronze Age. (a) Fitting models include Mesopotamian (Çayönü) and steppe ancestry. (b) Models with western sources from Southeastern Europe fail except those with Mayaky or Boyanovo EBA sources both of which are Yamnaya-derived. (c) The steppe (BPgroup)+Çayönü model fails all Chalcolithic/Bronze Anatolians except Central Anatolian Bronze Age. (d) Steppe (BPgroup) ancestry observed in all individuals of the Central Anatolian Bronze Age (±3s.e. shown). (e) BPgroup-related ancestry admixed with different substrata: Aknashen-related in the North Caucasus Maikop, Masis Blur-related in Chalcolithic Armenia, and Mesopotamian-related (Çayönü) in the ancestors of the Central Asian Bronze Age, following the route (f) from the North Caucasus to Anatolia.

**Extended Data Figure 2:**
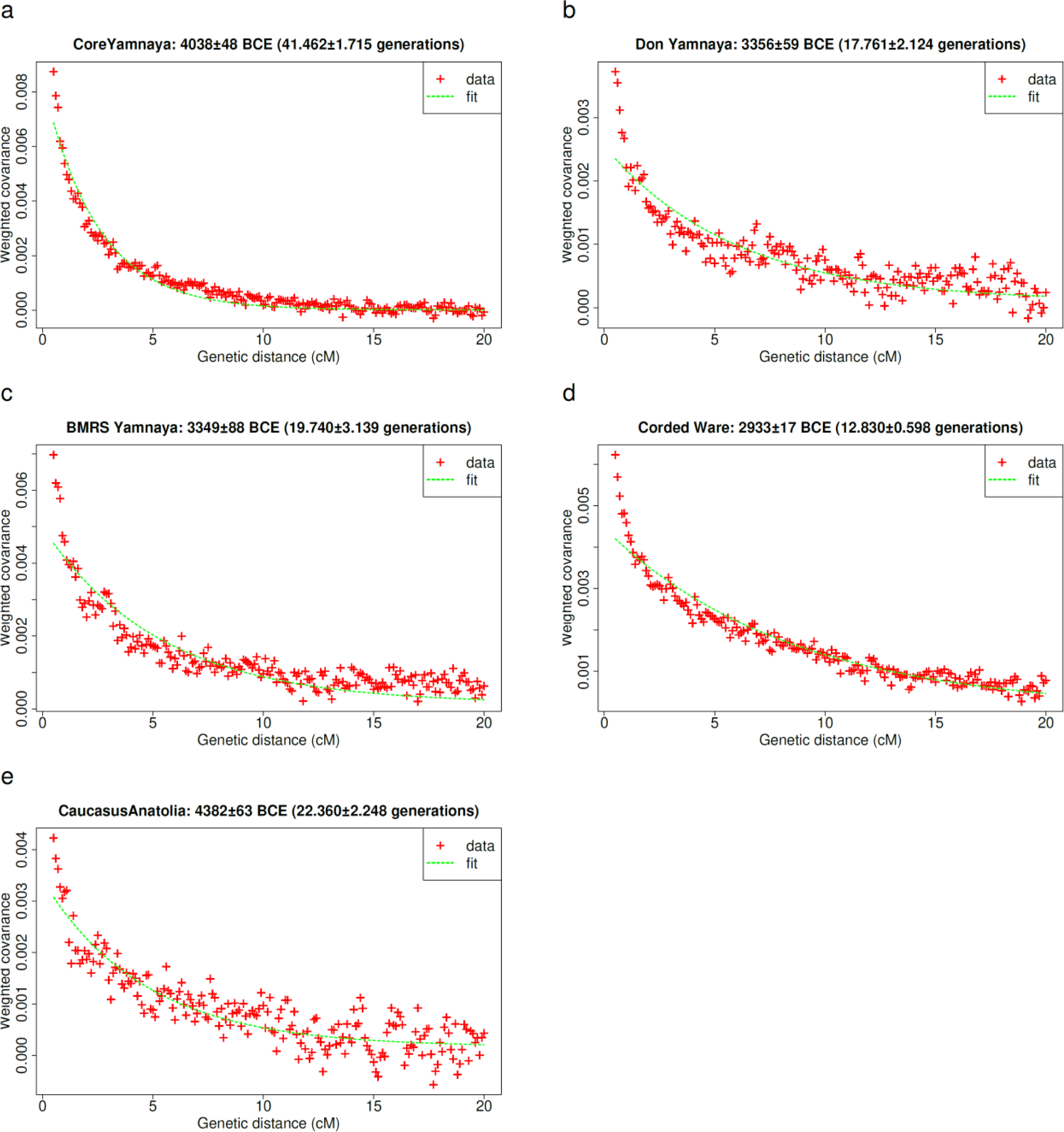
Admixture time estimates. We estimate admixture times for the Core Yamnaya as a mixture of European hunter-gatherer and West Asian populations (a), for the Don Yamnaya as a mixture of Core Yamnaya and UNHG (b), for the Bulgaria-Moldova-Romania-Serbia (BMRS) Yamnaya as a mixture of Core Yamnaya and European Neolithic/Chalcolithic farmers (c), for the Corded Ware as a mixture of Core Yamnaya and Globula Amphora (d), and for Caucasus-Anatolia populations (Maikop-Armenia_C-TUR_C_BA) as a mixture of European hunter-gatherer and West Asian populations which occurred ca. 4400BCE (e). The Core Yamnaya were formed ca. 4000BCE, followed by admixture ca. 3350 BCE with UNHG and European farmers in the east and west of the Dnipro-Don region and <3000BCE in central-eastern Europe.

**Extended Data Figure 3:**
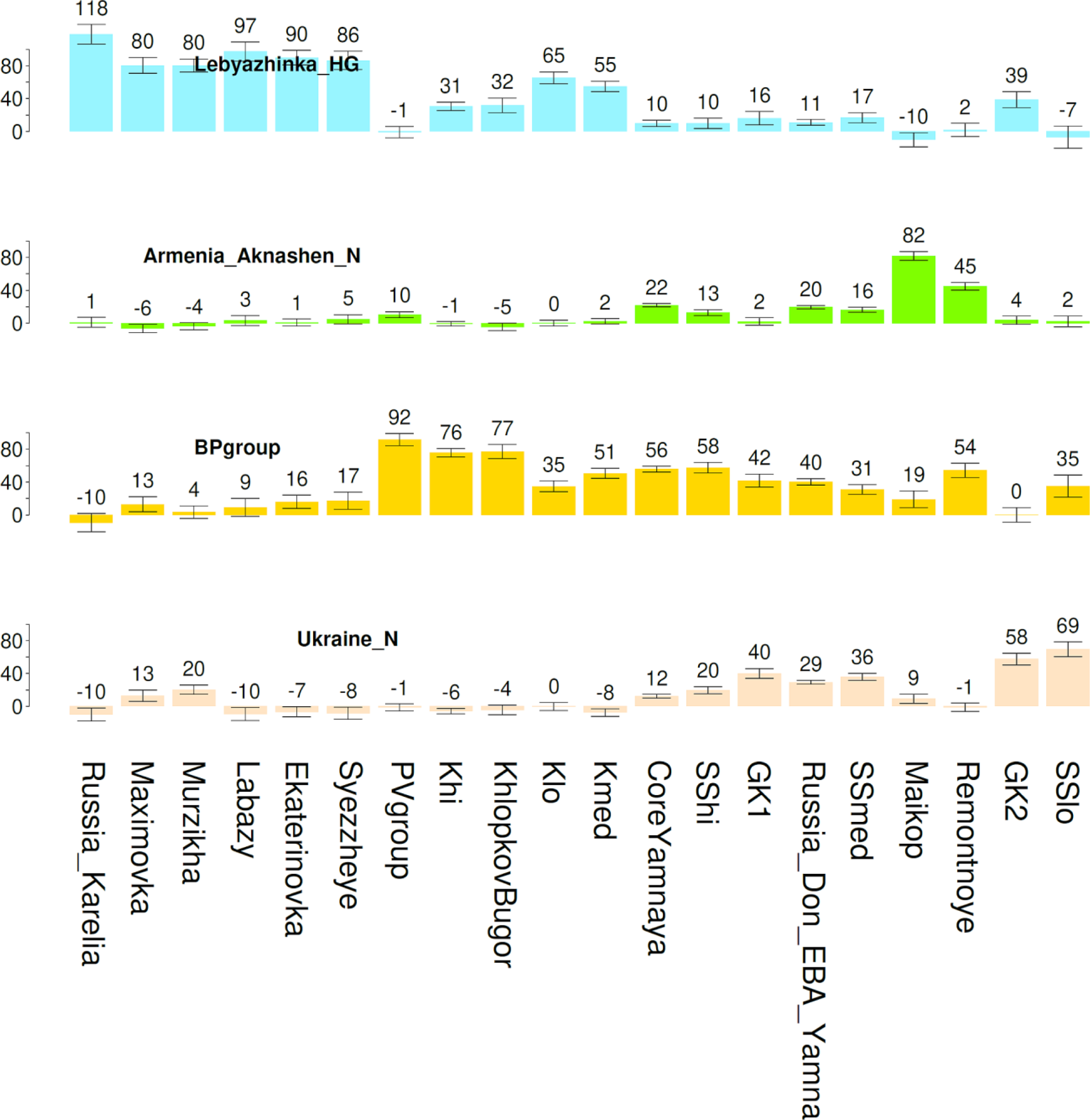
A 4-way model for the entire Dnipro-Don-Volga-Caucasus region. Error bars show ±1 standard error.

## Online Tables

**Online Table 1: Ancient individuals with newly reported genome-wide data.**

**Online Table 2: Technical details of newly reported ancient DNA libraries.**

**Online Table 3: Newly reported direct radiocarbon dates.**

**Online Table 4: All ancient individuals including in genome-wide analysis.**

