## Supplementary Information for "The Genetic Origin of the Indo-Europeans"

**Table of Contents**

**SI 1** – Archaeological Supplement 1-147

**SI 2** – The proximate and distal origins of the Yamnaya and the origins and dispersals of Indo-Anatolian and Indo-European languages 148-217

#### Supplementary Information 1

### Archaeological Supplement

David Anthony and Leonid Vyazov, editors

### 1. NEOLITHIC AND ENEOLITHIC OF THE FOREST ZONE OF THE EAST EUROPEAN PLAIN

In Northern Eurasian archaeology, the term "Neolithic" denotes a developmental phase in material culture characterized by the utilization of polished and drilled stone tools alongside pottery by hunter-fisher-gatherer communities. In chronological terms, the Neolithic was introduced to the forested zone of East Europe around ~7000-6500 BCE. This period gradually transitions into the Eneolithic phase, initiating in the steppes approximately from ~5200/5000 BCE. This transition is marked by an increase in the presence of metal artifacts, initially acquired through imports and subsequently manufactured across diverse regions within this expansive area.
The genome data for the following two sites, Sakhtysh-2 and Murzikha-2, was recently presented by Zeng et al. (2023), supplemented by detailed archaeological descriptions. Here we include the summarized observation on their chronology and genetic results.

#### 1.1 Sakhtysh-2

*Summary by D. Anthony*

Sakhtysh-2 is located 200km NE of Moscow in the upper Volga region. It is a multi-period riverside cemetery of the Middle Neolithic Lyalovo culture (conventionally mid 6th to early 4th millennium BCE) and the Late Neolithic Volosovo culture (conventionally late 5th to mid 4th millennium BCE). The Lyalovo graves (15 reported in 2013) were in extended supine position with red ochre on the grave floor & heads oriented to the SE; Volosovo (57 reported) were in extended supine position without red ochre & heads to the SW. All had carbon and nitrogen isotopic values most like otters analyzed from a nearby site, Minnino, documenting the long-term primary role of fish in the human diet. Lyalovo graves contained ornaments of elk (Alces alces) and bear teeth. 36 of 57 Volosovo graves contained beads made of stone, bone, animal teeth, and amber. 90% of 367 amber beads came from two Volosovo graves.

| Sample ID | Burial code | Radiocarbon date | Attribution | Y-chromosome | mtDNA |
|---|---|---|---|---|---|
| I12494 | raskop 1, grave 12 | 3621-3373 calBCE (4705±25 BP, PSUAMS-9400) | Volosovo (Lyalovo in 2013 publ.) | I2a-S21825 | U5a1d |
| I12496 | grave 13 (layer II), | 5500-3600 BCE | | n/a (female) | U4c1 |
| I12498 | grave 56 (level IIa), | 5500-3600 BCE | | R1b-R-L754 | U5a2b2 |
| I12500 | grave 13 (layer IIa), | 4400-3600 BCE | Volosovo | n/a (sex unknown) | U5a2d |
| I12962 | grave 54 (level IIa) | 3768-3649 calBCE (4940±19 BP) [R_Combine: (4900±30 BP, PSUAMS-9401); (4964±23 BP, AAR-15051)] | Volosovo | n/a (female) | U5a1d |
| I12964 | grave 34 (layer IIa) | 3630-2904 calBCE (4540±150 BP, GIN-7276) | Volosovo | R1b-R-Y13200 | U4b1b1 |
| I8404 | grave 32 (layer IIa) | 3952-3792 calBCE (5060±25 BP, PSUAMS-9086) | Volosovo | R1b-R-Y13200 | U2e1h |
| I8407 | raskop 3, KV 16 layer I | 5500-3600 BCE | | n/a (female) | U4a1 |
| I8408 | grave 33 (layer IIa) | 3800-3600 BCE | Volosovo | R1b-R-Y13200 | R |
| I8409 | grave ? (layer IIa 1988) | 4047-3961 calBCE (5195±25 BP, PSUAMS-9087) | Volosovo | R1a-R-YP4141 | U5a1d |
| I8410 | grave 40 (layer IIa) | 5617-5477 calBCE (6585±35 BP, PSUAMS-9405) | Lyalovo | Q1b-Q-M930 | U4a1 |

| Sample ID | Burial code | Radiocarbon date | Attribution | Y-chromosome | mtDNA |
|---|---|---|---|---|---|
| I8411 | grave 42 (layer IIa) | 5323-4610 calBCE (6060±150 BP, GIN-6586) | Lyalovo | R1b-R-Y13200 | U5a1d |
| I8413 | grave 29 (layer II) | 5500-3600 BCE | | n/a (female) | K1b2 |
| I8414 | grave 35 (layer IIa) | 3283-2049 calBCE (4080±180 BP, GIN-7273) | Volosovo | I2–I-P37 | U5a2d |
| I8415 | shurf N1 (layer 8) | 5500-3600 BCE | | n/a (female) | U5a2d |
| I8416 | grave 61 (layer IIa) | 5633-5483 calBCE (6650±35 BP, PSUAMS-9406) | Lyalovo | n/a (female) | U5a2d |
| I8417 | grave 58 (layer IIa) | 4342-4248 calBCE (5430±25 BP, PSUAMS-9088) | Volosovo | Q1b-Q-Y6802 | U2e1e |
| I8418 | grave 39 (layer IIa) | 5500-3600 BCE | | R1a-R-M198 | U5a2d |
| I8419 | grave 11 (layer IIa) | 4400-3600 BCE | Volosovo | n/a (female) | U2e1e |
| I8437 | grave 10 (layer II) | 3635-2896 calBCE (4540±160 BP, GIN-6234) | Volosovo | n/a (female) | U2e1e |
| I8438 | grave 43 (layer IIa) | 5500-5200 BCE | Lyalovo | R1b-R-P297 | U4b1a |

#### 1.2 Murzikha-2

*Summary by D. Anthony & A. Chizhevsky*

Murzikha-2 was an Eneolithic cemetery in the northern broadleaf forest zone near the Volga-Kama river junction, below which the Volga doubles in width. The site was on the south side of the Kama, originally sited on a low peninsula today used as an approach to a bridge. Dated to about 4400 BCE by dates on human bone, the cemetery had 18 Eneolithic graves, 6 single and 12 collective. The standard burial pose apparently was sitting forward, with the head and arms resting on raised knees (probably tied in position). Three individuals were sitting side by side in each of four collective graves. The cemetery contained three or four generations of a single extended family; all 14 analyzed were relatives.

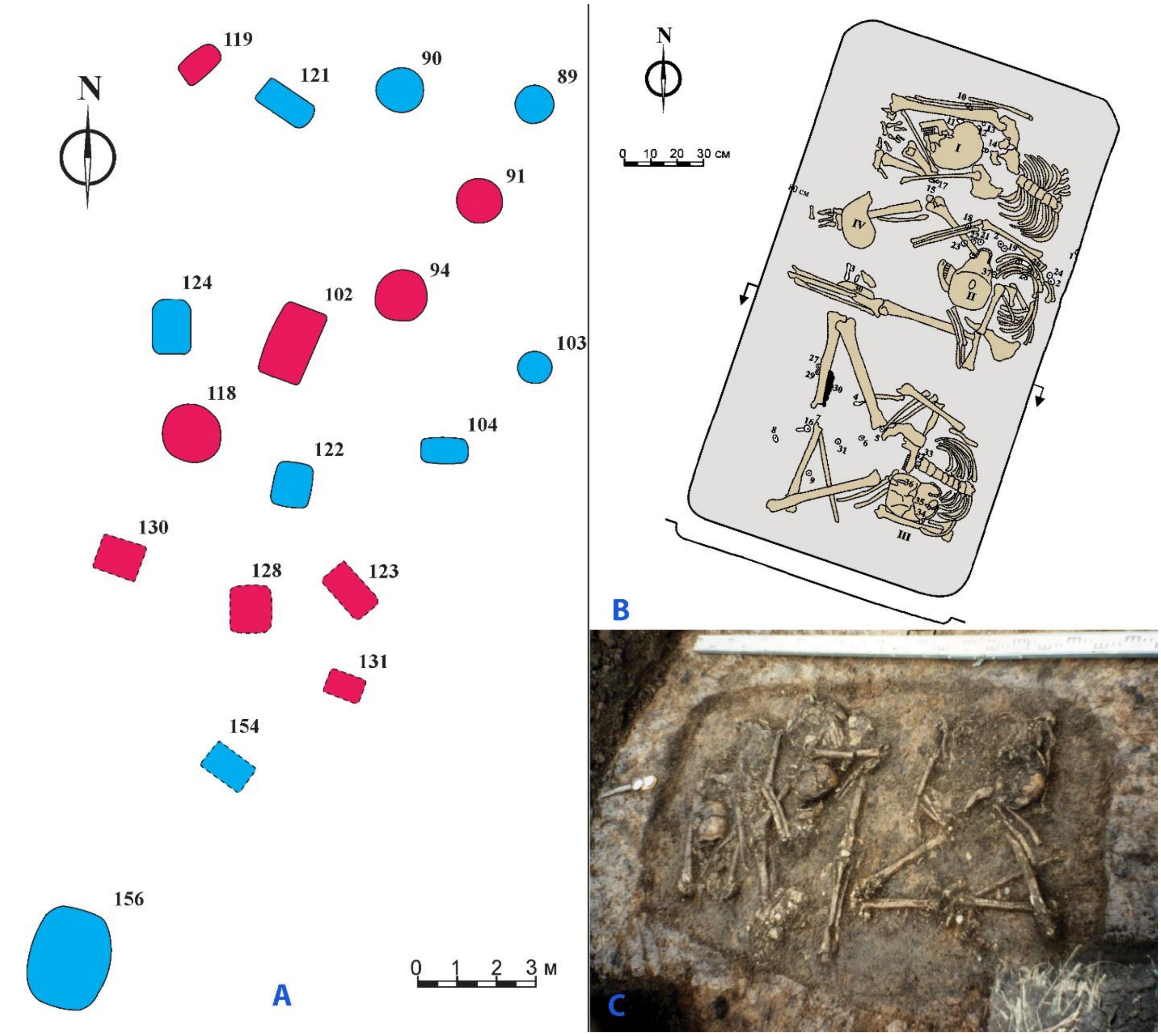

**Fig. 1.1. Eneolithic burials at Murzikha-2. A – general plan of the cemetery, sequenced burials are highlighted in red, B and C – plan and photo of burial 102 (images by Andrey Chizhevskiy).**

Two polished stone pendants of serpentine and one copper wire hoop used as ornaments were obtained from the steppes to the south. One grave contained a spatula-like tool made of a domesticated cattle scapula, but the economy retained a large role for hunting and fishing.

The genetics of the Murzikha II family are on the northern end of a cline of Volga populations. Within the cline Murzikha II is aligned with the Upper Volga Lyalovo & Volosovo hunter-fisher populations at Sakhtysh-2, located 600km upriver but sharing the forest zone, and not with the middle Volga Ekaterinovka Mys and Syezzheye populations, located less than half that distance downriver, but in a different ecological zone, the steppes.

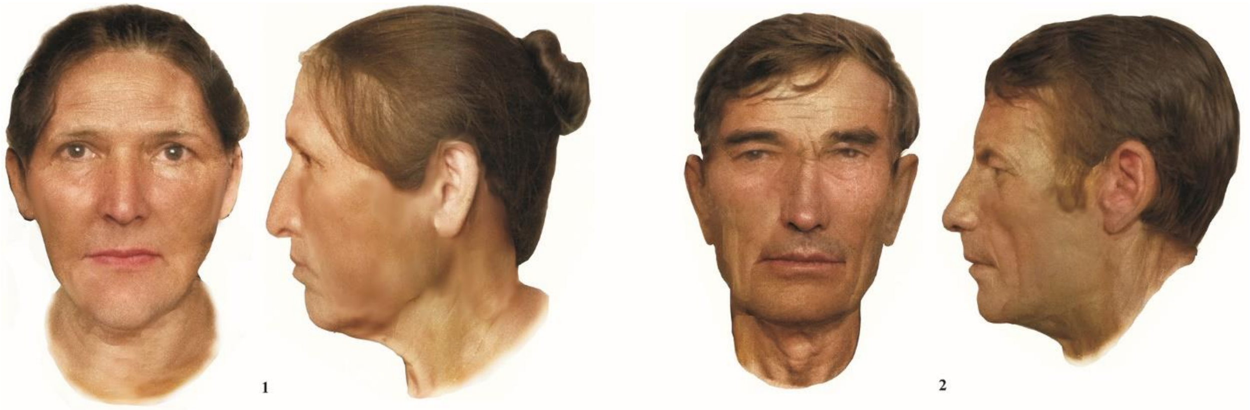

**Fig. 1.2. People buried in Murzikha.** A reconstruction by Aleksey Nechvaloda (image contributed by Andrey Chizhevskiy).

According to family analysis by Iñigo Olalde: Murzikha Family A (14 Members) (I8455-I8744-I8454 are grandfather-father-daughter. All three individuals are 2nd/3rd degree relatives of I8448, I8451, and I11841 (who are all 2nd/3rd degree relatives of each other); all three are also 2nd/3rd degree relatives of I8456 (who is also 2nd/3rd degree relative of I8451 and I11841). The grandfather (I8455) is also a 2nd/3rd degree relative of I8449 (who is a 2nd/3rd degree relative of I8448 and I11841), I8453 (who is a 2nd/3rd degree relative of I8449, I8450, and I11841), and I8450 (who is the brother of I11030 and is a 2nd/3rd degree relative of I8448, I8452, I8453, and I11841); the father (I8744) is also a 2nd/3rd degree relative of I8449 and I8452 [who is a 1st degree relative of I8451 and a 2nd/3rd degree relative of I11841 and I11842 (who is also a 2nd/3rd degree relative of I8457)]).

| Sample ID | Burial code | Radiocarbon date | Sex | Y-chromosome | mtDNA |
|---|---|---|---|---|---|
| I11030 | Murzikha II B91 skeleton 1 | 4447-4351 calBCE (5560±25 BP, PSUAMS-8296) | M | Q1a-Q-F1096 | U2e2a |
| I11841 | Murzikha II B131 | 4599–4347 calBCE (5630±60 BP, Ki-10038) | M | Q1a-Q-F1096 | U5a1d2b |
| I11842 | Murzikha II B94 skeleton 2 | 4538–4365 calBCE (5630±40 BP, Ki-9428) | F | | U4a1 |
| I8448 | Murzikha II B102 skeleton 1 A | 4358-4256 calBCE (5470±25 BP, PSUAMS-8281) | M | Q1a-Q-F1096 | U2e2a |
| I8449 | Murzikha II B102 skeleton 1g | 4547-4373 calBCE (5655±25 BP, PSUAMS-8334) | M | Q1a-Q-F1096 | U5a2b1 |
| I8450 | Murzikha II B104 | 4681-4463 calBCE (5730±30 BP, PSUAMS-8282) | M | Q1a-Q-F1096 | U2e2a |
| I8452 | Murzikha II B118 skeleton 3 | 4532-4358 calBCE (5615±30 BP, PSUAMS-8283) | F | | U5a1d2 |
| I8453 | Murzikha II B119 | 4653-4452 calBCE (5700±30 BP, PSUAMS-8335) | F | | U4a |
| I8454 | Murzikha II B123 skeleton 1 | 4446-4346 calBCE (5550±25 BP, PSUAMS-8292) | F | | R1b |
| I8455 | Murzikha II B128 skeleton 2 | 4446-4274 calBCE (5515±30 BP, PSUAMS-8293) | M | Q1a-Q-F1096 | U5a2b1 |
| I8456 | Murzikha II B128 skull I(?) | 4678-4494 calBCE (5730±25 BP, PSUAMS-8336) | M | Q1a-Q-F1096 | U2e2a |
| I8451 | Murzikha II B118 skeleton 2 | 4602-4403 calBCE (5670±30 BP, PSUAMS-8393) | M | Q – Q-L472 | U5a1d2 |
| I8457 | Murzikha II B130 skeleton 1 | 4700-4200 BCE | M | Q1a-Q-YP1669 | U5a1d2b |

| Sample ID | Burial code | Radiocarbon date | Sex | Y-chromosome | mtDNA |
|---|---|---|---|---|---|
| I8744 | Murzikha II B118 skeleton 1 | 4700-4200 BCE | M | Q –<br>Q-L472 | U4d |

<u>*Bibliography to section 1:*</u>

*Chizhevskii, A.A. 2008. Pogrebeniya epokhi Eneolita Murzikhinskogo II mogil'nika. Trudy II (XVIII) Vserossiiskogo Arkheologicheskogo C'ezda v Suzdale 2008 g. Tom I: 367-371. Moskva: Institut Arkheologii RAN.*

*Piezonka, H. E. Kostyleva, M. G. Zhilin, M. Dobrovolskaya, and T. Terberger. 2013. Flesh or fish? First results of archaeometric research of prehistoric burials from Sakhtysh-2a, Upper Volga region, Russia. Documenta Praehistorica XL (2013): 57-73. DOI: 10.4312/dp.40.6*

*Zeng TC, Vyazov LA, Kim A, et al. Postglacial genomes from foragers across Northern Eurasia reveal prehistoric mobility associated with the spread of the Uralic and Yeniseian languages. bioRxiv; 2023. DOI: 10.1101/2023.10.01.560332.*

#### 2. ENEOLITHIC OF THE FOREST-STEPPE AND STEPPE ZONES

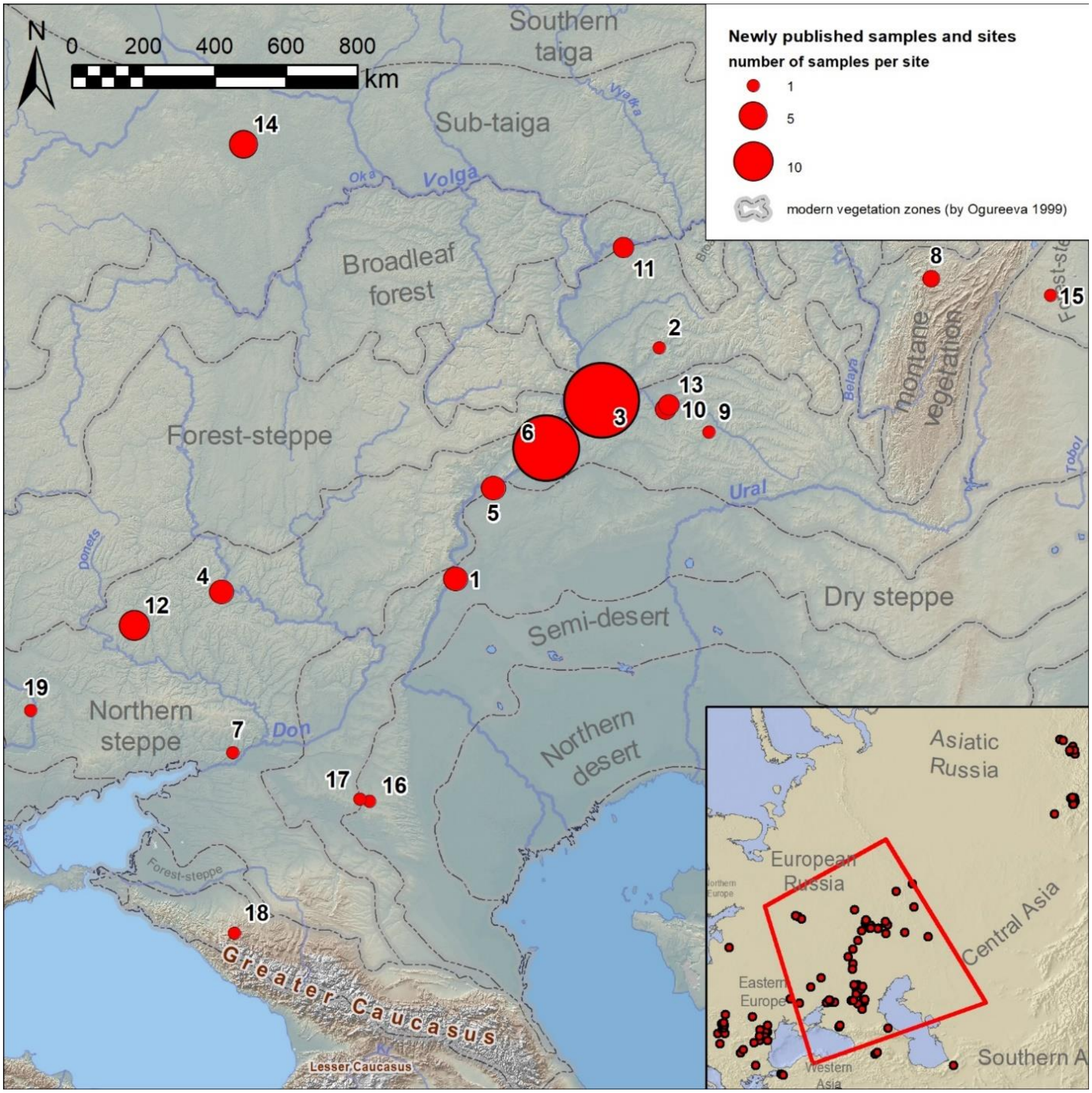

**Fig. 2.1. Newly published samples from Neolithic and Eneolithic sites on the East European Plain.** 1 - Berezhnovka-2 (Volgograd Oblast, Nikolayevsky District); 2 - Chekalino-4 (Samara Oblast, Sergiyevsky District); 3 – Ekaterinovka Mys (Samara Oblast, Bezenchuksky District, Ekaterinovka); 4 - Golubaya-Krinitsa (Voronezh Oblast, Rossoshansky District); 5 - Khlopkov-Bugor (Saratov Oblast, Voskresensky District); 6 - Khvalynsk-1 and -2 (Saratov Oblast, Khvalynsky District); 7 - Krivyansky-9 (Rostov Oblast, Oktyabrsky District); 8 - Kulmetovskiy Grot (Chelyabinsk Oblast, Katav-Ivanovsky District); 9 - Labazy (Orenburg Oblast, Kurmanayevsky District); 10 - Maksimovka-1 (Samara Oblast, Neftegorsky District); 11 - Murzikha-2 (Tatarstan, Alexeyevsky District, Mokrye Kurnali); 12 - Olexandria (Ukraine, Kharkiv Oblast, Kupiansk District, Kurilivka municipality); 13 - S'yezzheye (Samara Oblast, Bogatovsky District); 14 - Sakhtysh-2a and 8 (Ivanovo Oblast, Teykovsky District); 15 - Shatrovo-1 (Chelyabinsk Oblast, Chebarkulsky District); 16 - Sukhaya-Termista-1 and 17 - Ulan-4 (Rostov Oblast, Remontnensky District); 18 - Unakozovskaya (Republic of Adygea, Maykopsky District); 19 - Vovnihy-2 (Ukraine, Dnipropetrovsk Oblast, Dnipro District, Solone municipality).

#### 2.1 Berezhnovka-2

*Summary by D. Anthony & N. Shishlina*

Two groups of kurgans (1 and 2) were excavated near the village of Berezhnovka on the east (left) side of the Volga 1950-1957 before the construction of the Volgograd dam and reservoir in 1958-61 flooded the site. Here the west (right) bank of the Volga was a series of white limestone bluffs that stood 100m over the floodplain, cut by steep ravines, while the left bank was a flat, arid steppe. Yamnaya kurgan cemeteries were strung along the left bank at intervals of about 15km, according to Merpert, and Berezhnovka-1 and 2 together formed one such cluster NW of the village of Berezhnovka. 1 and 2 were located about 3km apart on a landform that in the early 20th century was an island surrounded by phragmites marshes. The location could have been a peninsula in the Eneolithic. In the steppes phragmites reed marshes were historically an important source of winter fodder for cattle and construction materials for floors, beds, roofs, and wagon covers.

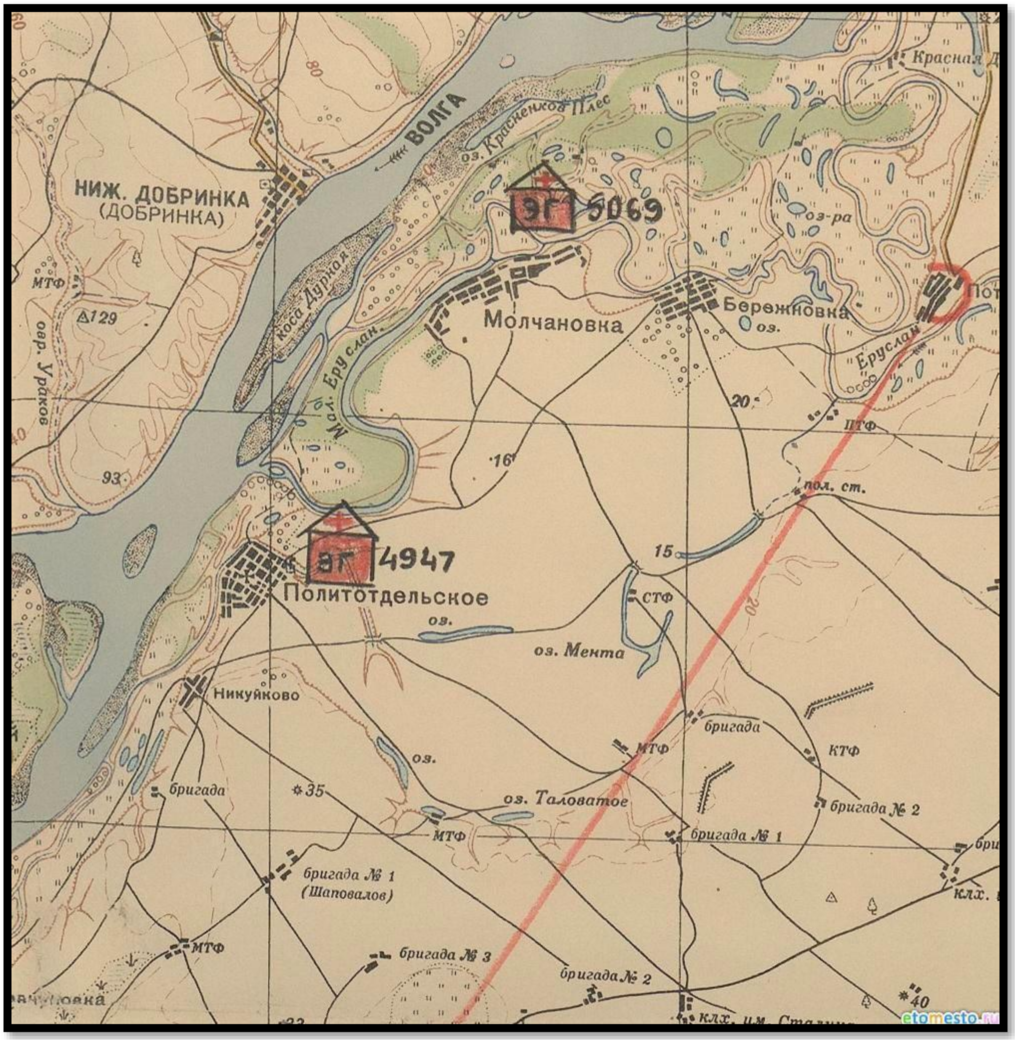

**Fig. 2.2. Berezhnovka-1 (south) and 2 (north) before dams flooded the region.**

Kurgan 9 at Berezhnovka-2 covered graves that seemed archaic compared to the other Yamnaya kurgans such as Berezhnovka-1 kurgan 5, grave 20 (I22197). In Merpert's foundational 1974 synthesis of the Yamnaya culture he designated the k.9 graves as representing the earliest

Yamnaya phase, extending back into the Eneolithic and found only on the Volga. Marija Gimbutas's theory that the 'Kurgan culture' originated on the lower Volga was based partly on Merpert's interpretation of Berezhnovka-2 k.9 (Mallory 1977). After the discovery of the Eneolithic Khvalynsk cemetery in 1977 with graves like Berezhnovka k.9, these graves are regarded as probably Eneolithic, dated before the Yamnaya period.

Here we confirm that the graves under k.9 date to the late 5th-millennium BCE, contemporary with the late Khvalynsk culture. The raised-knee posture, use of red ochre, bird-bone tubes, flint lamellar unifacial blades, bone beads, and two small copper rings in grave 9 make an assemblage that would fit easily in the graves at Khvalynsk. New radiocarbon dates were obtained with the help of Natasha Shishlina in the State Historical Museum, Moscow, and Konstantin Marzhorin in the Saratov Museum.

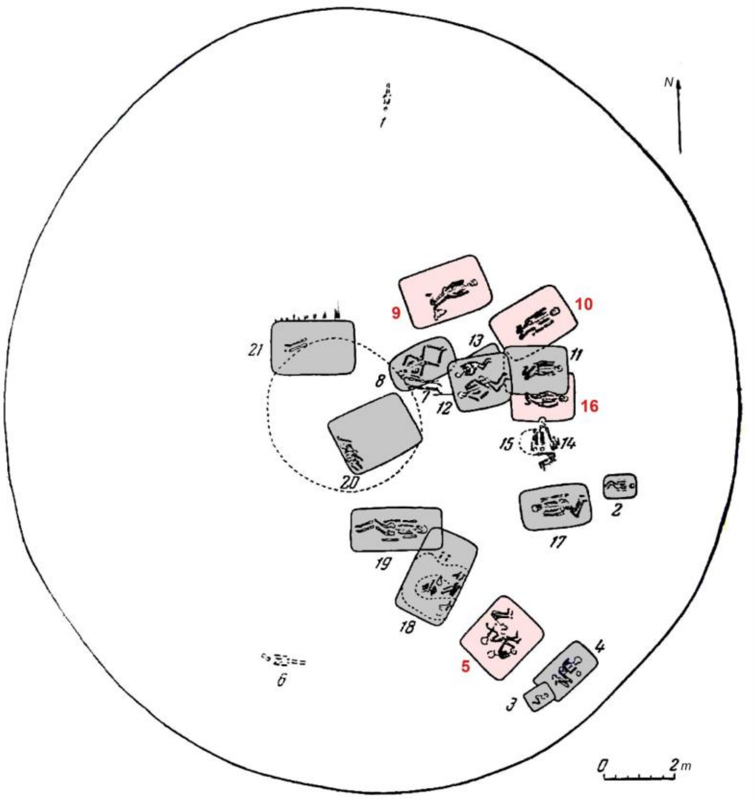

**Fig. 2.3. Berezhnovka-2, kurgan 9. The sequenced burials are marked in red.**

#### *Dates, FRE, and diet*

The oldest two radiocarbon results, one on a human from k.9 grave 5 (I22199) dated 4929-4730 calBCE, and the other on a bird (swan-size) bone from k.9 grave 9 dated 4841-4711 calBCE, fall in the early 5th millennium BCE and probably are affected by a Freshwater Reservoir Effect (FRE) making them too old. Swans have a riverine diet subject to FRE, and the human in grave 9 (I22200) with the bird bone gave a radiocarbon date 700 years later. The other Eneolithic individuals (I22202, I22201, and I22200) have dates on human teeth that fall in the late 5th millennium BCE contemporary with and partly after Khvalynsk, supporting the artifactual similarities with Khvalynsk.

This difference in FRE in between grave 5 and the others implies that one Eneolithic individual at Berezhnovka-2 (I22199, with Y-haplogroup Ia2 like Serednii Stih) had a more riverine diet that created a FRE, while others buried beside him (I22202, I22201, and I22200 all with Y-haplogroup R1b) had a more terrestrial diet and therefore yielded accurate radiocarbon dates.

Variation in diet within the Eneolithic Volga Cline is documented also by the variation in the consumption of dairy foods between Progress-2 and other Eneolithic sites sampled for dairy proteins in human dental calculus. Dairy peptides were absent from seven individuals analyzed from Khvalynsk and Khlopkov Bugor by Wilkin et al. (2021). Berezhnovka was not analyzed for dairy peptides. But dairy peptides were present in one of three Eneolithic individuals sampled from Progress-2 (PG2001) in the North Caucasus steppes at the southern end of the Volga cline (Scott et al 2022). Probably the concept of obtaining and preparing dairy foods was learned from Caucasus farmers who consumed milk products earlier, but dairy was not widely adopted in the steppes until the Yamnaya period (Wilkin et al. 2021).

##### *Genetics*

The genetics of Berezhnovka-2 on the lower Volga were so similar to some individuals at Vonjucka and Progress-2 in the North Caucasus steppes that either a recent migration or an ongoing system of mate exchanges occurred between these two regions, separated by 800km and in quite different ecological settings. The variation in FRE suggested at Berezhnovka-2 could have been caused by the movement of people with different diets from distinct ecologies between Progress-2 (terrestrial, so accurate radiocarbon dates) and Berezhnovka (riverine, so possibly a large FRE).

At Berezhnovka-2 two Y-haplotypes in the R1b R-V1636 clade are typical in males in the Eneolithic Volga Cline and are found also at Progress-2. The single Y I2a (with a high FRE) is atypical; it was typical of Dnipro-Don cline males. Three of the four mt-haplogroups are common in the Volga Cline.

However, mt-H2a1 was found in Neolithic Anatolia, the Caucasus, and European Neolithic farmers including LBK. It is attested in the late 5th millennium BCE both in the Volga Cline (Berezhnovka-2, Khvalynsk, Khlopkov Bugor) and in the Caucasus (Areni-1, where R1b R-V1636 also appears). It is not attested on the Dnipro prior to Serednii Stih. H2a could indicate mate exchanges between the Caucasus and the Volga Cline accompanying the spread of CHG-related ancestry into the Volga Cline. The Khvalynsk male buried with the largest number of Balkan copper beads (I0122) was mt-H2a. H2a persisted in the 4th millennium BCE in Trypillia (Verteba cave), Maikop, Steppe Maikop, Usatove, and Yamnaya populations.

| Sample ID | Burial code | Radiocarbon dates | Y-chromosome | mtDNA |
|---|---|---|---|---|
| I22199 | kurgan 9, burial 5 | 4929-4730 calBCE (5955±20 BP, PSUAMS-8815) | I2a-I-S12195 | H13a1 |
| I22201 | kurgan 9, burial 10 | 4446-4333 calBCE (5525±30 BP, PSUAMS-9809) | R1b-R-V1636 | U2e1'2'3 |
| I22200 | kurgan 9, burial 9 | 4049-3961 calBCE (5200±25 BP, PSUAMS-8840) | R1b-R-Y106006 | H2a1 |
| I22202 | kurgan 9, burial 16 | 4235-4173 calBCE (5308±18 BP) [R_combine: (5295±25 BP, PSUAMS-9810), (5310±25 BP, PSUAMS-8841)] | (F) | U5a1f |

*Mallory, J. P. 1977. The chronology of the early kurgan tradition (part two). Journal of Indo-European Studies 5/4 (1977) 339–368.*

*Merpert, N. I. 1974. Drevneishie Skotovody Volzhsko-Uralskogo Mezhdurechya. Moskva 1974.*

*Scott, Ashley, S. Reinhold, T. Hermes, A. A. Kalmykov, A. Belinskiy, A. Buzhilova, N. Berezina, A. R. Kantorovich, V.E. Maslov, F. Guliyev, B. Lyonnet, P. Gasimov, B. Jalilov, J. Eminli, E. Iskandarov, E. Hammer, S. E. Nugent, R. Hagan, K. Majander, P. Onkamo, K. Nordqvist, N. Shishlina, E. Kaverzneva, A. I. Korolev, A. A. Khokhlov, R.V. Smolyaninov, S.V. Sharapova, R. Krause, M. Karapetian, E. Stolarczyk, J. Krause, S. Hansen, W. Haak & Christina Warinner. 2022. Emergence and intensification of dairying in the Caucasus and Eurasian steppes. Nature Ecology and Evolution 6, 813–822. https://doi.org/10.1038/s41559-022-01701-6*

*Wilkin, Shevan, A. Ventresca Miller, R. Fernandes, R. Spengler, W. T. Taylor, D. R. Brown, D. Reich, D. Kennett, B.J. Culleton, L. Kunz, C. Fortes, A. Kitova, P. Kuznetsov, A. Epimakhov, A.K. Outram, E. Kitov, A. Khokhlov, D. Anthony, and Nicole Boivin. 2021. Dairying enabled Early Bronze Age Yamnaya steppe expansions. Nature 598: 629–633. https://doi.org/10.1038/s41586-021-03798-4*

#### 2.2 Chekalino-4

*Summary by D. Anthony & L. Vyazov*

Chekalino-4 is a Neolithic-Eneolithic settlement site on the east (left) bank of the Sok River, 3.7km south of Chekalino village. The Sok flows into the Volga north of Samara through the steppe/forest-steppe ecotone. The site yielded a rich inventory of Early Neolithic and Eneolithic lithics and ceramics, and 5 hearths, 8 pits, and middens of freshwater Unio shells. A single grave of a female aged 50-55 posed crouched on her side was found in the settlement area and was assumed to be Early Neolithic. But radiocarbon dates show that this grave was deposited long after most other artifacts, in the mid-4th millennium BCE.

The Chekalino-4 female belonged to a northern, EHG-rich population like late-5th-millennium Murzikha on the Kama-Volga confluence 180km to the north. Elements of this population moved south into the Samara region in the early and middle 4th millennium BCE at Chekalino and Maximovka, apparently displacing Volga-Cline populations genetically like S'yezzheye and Ekaterinovka Mys. It is surprising that at both Chekalino-4 and Maximovka on the Samara River these EHG-rich 4th-millennium BC graves are found inserted into older Neolithic settlements.

The Yamnaya arrival in the Samara region about 3300 BCE introduced another new population that displaced Chekalino-4-type genetics. The Yamnaya migrants built a wall of kurgan cemeteries consisting of at least six archaeologically attested kurgan groups distributed over 40 km along the southeast bank of the Sok from Chekalino-4 to Grachevka. No Yamnaya kurgans were built on the northwest side of the Sok.

| Sample ID | Burial code | Radiocarbon dates | Y-chromosome | mtDNA |
|---|---|---|---|---|
| I6303 | Chekalino-4 grave | 3645-3528 calBCE (4815±20 BP, PSUAMS-2909) | (F) | U5a2+16294 |

*Mamonov, A.E. 2000. Neolit. In Istoriya Samarskogo Povolzh'ya S Drevneishikh Vremen do Nashikh Dnei: Kammennyi Vek, pp141-176. Samara: Integratsiia.*

#### 2.3 Ekaterinovka Mys

*Summary by D. Anthony & A. Korolev*

Ekaterinovka Mys, discovered and excavated 2013-2018 by A.A. Korolev, A.F. Kochkina, and D.A. Stashenkov is an Eneolithic cemetery of 100 graves located on a peninsula surrounded on three sides by a minor Volga tributary, the Bezenchuk River. Before the Saratov Reservoir was filled the Eneolithic cemetery was about 6km above the Bezenchuk-Volga confluence; today it is on the edge of the reservoir.

'Ekaterinovskiy Cape' is the English translation used by the excavators themselves, but English 'cape' usually refers to a seacoast landform (Cape Horn, Cape Cod). A better English word might be 'point', used colloquially for landforms that jut into lakes, rivers, or bays. But 'mys' has no exact English equivalent, so we use the Russian here. Also, 'Ekaterinovka' is the nominative case of the village name, as on an English-language map, while 'Ekaterinovskiy Mys' is 'the mys of Ekaterinovka', using a case ending that has no English equivalent. The name here, 'Ekaterinovka Mys', uses undeclined (as in English) Russian words for this place-name. But we make no claim to accuracy in these murky waters.

The Volga River in this region makes a loop 90km east and then 90km west around a white limestone ridge standing 250m above the river, a natural landmark called the Samara Luka that coincides with the ecotone between the forest-steppe on its north and the steppe on its south. The Samara River flows into the Volga at the eastern apex of the Luka. Ekaterinovka Mys was located on the southern or steppe side of the Samara Luka, 50km west (downstream) of the modern city of Samara.

The normal burial pose at Ekaterinovka Mys was supine with legs straight like S'yezzh'e and most Neolithic/Early Eneolithic cemeteries. But at least five graves (23, 85, 86, 81, 90, and perhaps 52) were posed supine with raised knees, early examples of this distinctive pose, later standard at Khvalynsk and in Yamnaya graves.

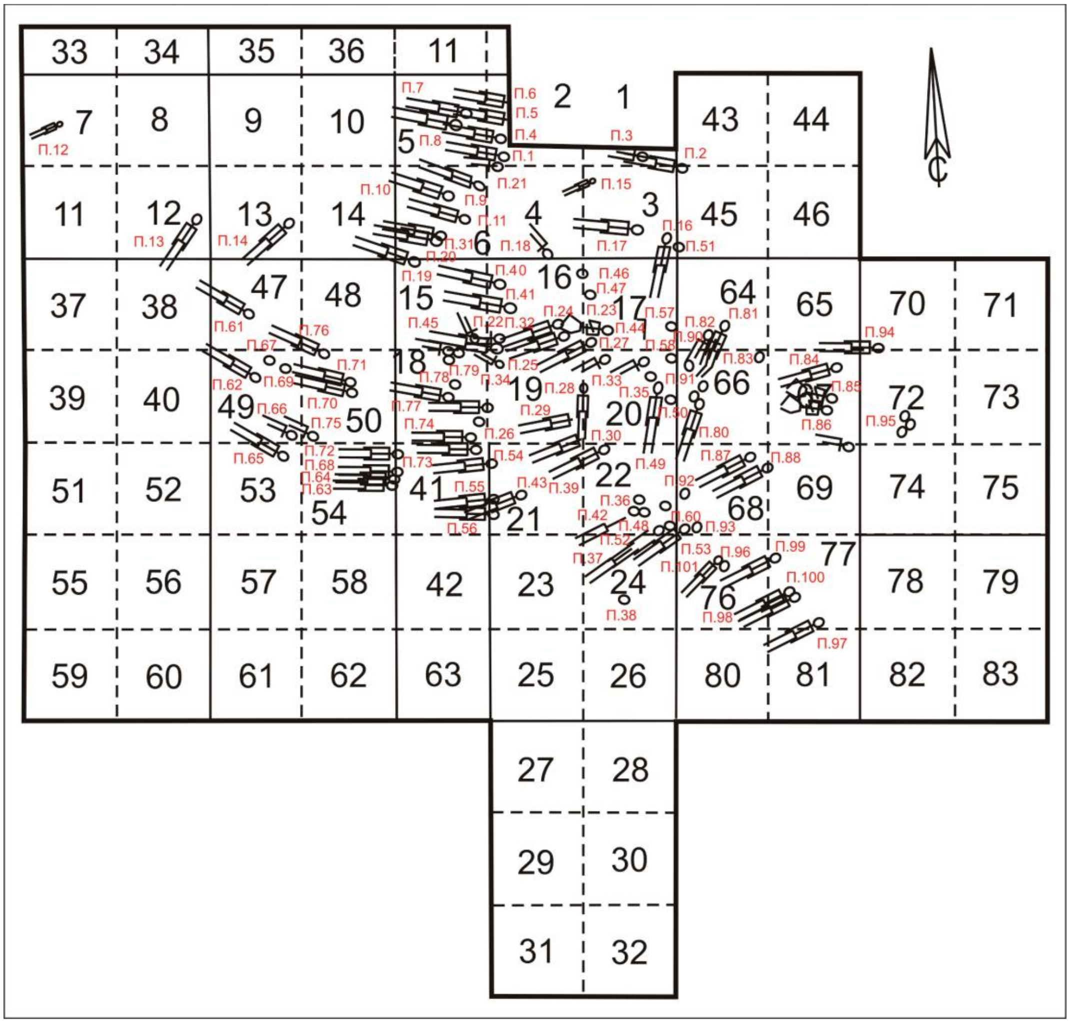

**Fig. 2.4. Preliminary Schematic Plan of Ekaterinovka Mys cemetery.** Formal plan will be published with the final report and could change. Grave 45 was in squares 15 & 16. The five graves with raised knees were 81 in and 90 in sq.64, 85 & 86 in sq.67, and 23 in sq.17.

Graves were arranged beside each other in rows. One large row was on a line NW to SE with heads oriented to the N/NE; other overlapping rows were oriented N to S or NE to SW with heads oriented E/SE. A few graves did not conform to any row and were oriented heads to the N/NE.

Graves without any artifacts (about half) had little red ochre or none. Graves with artifacts had intense red ochre on the floor. Horse, sheep, and goat bones were found in the ochre-stained soil above the graves and in some graves; a 6-week-old domesticated goat kid was on the left arm of grave 45 (image). A sheep tooth was examined in the Reich lab for comparison of its DNA with other ancient sheep but it did not yield sufficient data. The sacrifices of domesticated mammals were like those at the contemporary cemetery at S'yezzh'e (two horse head-and-hoof deposits in red ochre above the human graves, cattle images carved in antler) and later at Khvalynsk. No copper was found; the cemetery is thought to pre-date the appearance of Balkan-derived copper on the Volga about 4500 BCE.

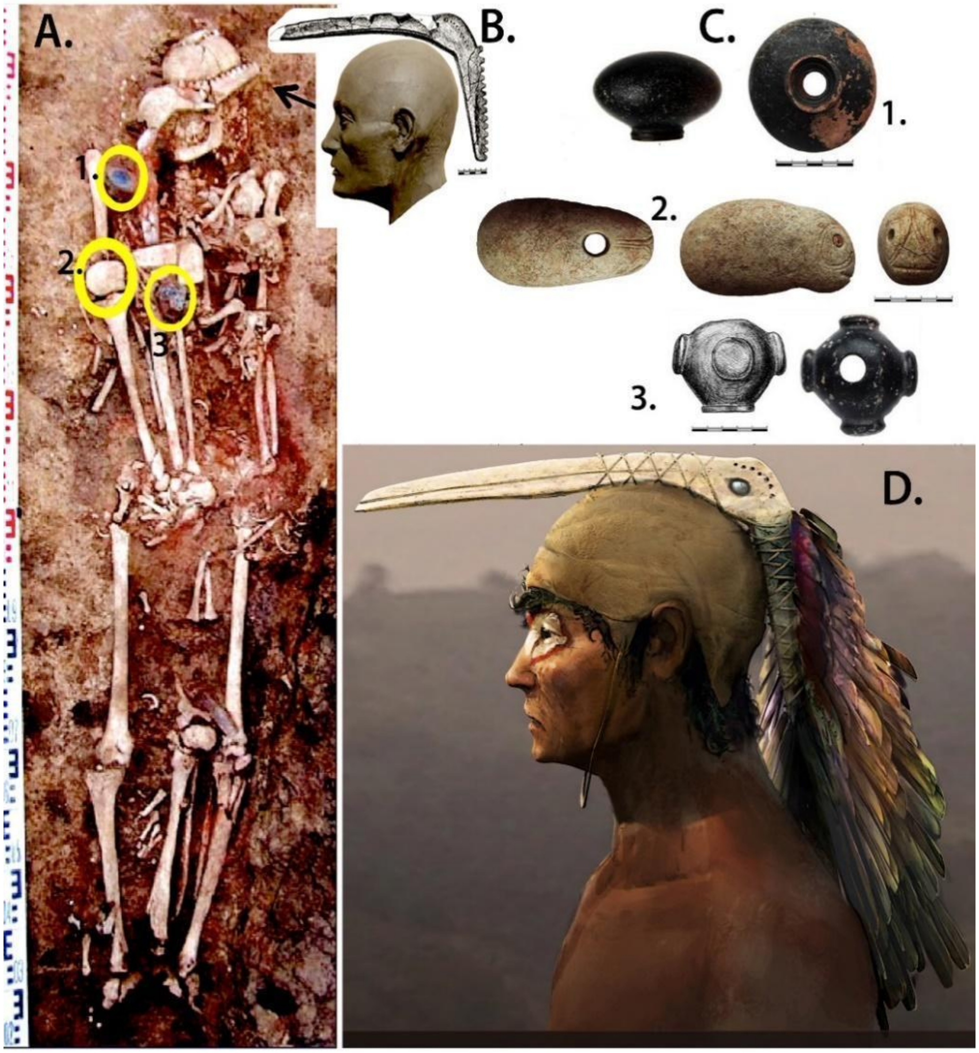

**Fig. 2.5. Ekaterinovka Mys grave 45. A.** Grave 45 with three mace heads circled. A tibia from another individual rests against his right arm and another between his tibias. Two severed hands of two individuals rest on his left hip, perhaps from the same two that yielded the tibias and two of the mace heads. A domesticated goat kid rests on his left arm. A long-beaked bird carved from antler rests across his head and face. **B**. Reconstruction of the head and face of grave 45, with a line drawing of the carved antler bird head arranged over his head. **C**. Three mace-heads from grave 45, found on his right arm, one a zoomorphic representation of a fish (?). **D**. Artist's rendition of the grave 45 male with the antler bird arranged as the crest of a hat, with eye paint and feather colors from the feather and eye patch colors of the Volga glossy ibis. A,B, and C after Korolev et al. 2018, Figures 2, 4, 7, 9, 10, & 12, used with permission. D is by Russel Story, used with permission. From Anthony et al. 2022 Figure 14.

Human bone from Grave 45 gave a radiocarbon date 700-800 years older (5311-5218 calBCE/6280±25BP/ PSUAMS-2882) than the goat tooth in the same grave (4550-4450 BCE, see table below), showing a strong FRE on dates on human bone. Four dates were obtained from terrestrial animal (marmot) incisors used as ornaments and these align with a date obtained previously on an organic residue from a pottery sherd. These 5 dates, shown below, suggest that the cemetery was in use primarily during the period 4800-4500 BCE; the midpoint of five means is 4628 BCE.

The ceramics are typical of the Samara culture like the site of S'yezzh'e on the Samara River. They were shell-tempered, comb-stamped and collared. Fifteen polished stone mace heads of various shapes were found, the largest and most varied collection of stone mace heads from any steppe cemetery. They occurred in six certainly male graves (40, 45, 69, 77, 76, 90) and in other graves of uncertain sex. A female in grave 31 had more than 1000 Unio shell beads arranged in multiple belts and sewn on the edges of her clothing. Grave 45 (image) had three stone maces and a carved antler long-beaked bird head that might have been a crest on an elaborate hat, found lying across his face; the severed hands and tibias of two other individuals were arranged on his body; and a domesticated goat kid was on his left arm.

Ekaterinovka Mys belongs to the Early Eneolithic period when a few standard items including long flint blades, boar's tusk ornaments, long belts made of shell, bone, stone, and animal teeth beads, and collared ceramic pots appeared in graves across the steppes, from the Dnipro to the Don and the Volga, before the spread of copper. The most elaborate dress ornaments were at the Mariupol cemetery in the coastal Azov steppes, to which other sites are compared.

##### *Genetics*

Ekaterinovka Mys represents the northern end of the Volga Cline, averaging 24% ancestry from CHG-related populations whose source population in the south is unknown. A millennium earlier the Lebyazhinka IV hunter-gatherer, also from the Samara region, had 7% CHG-related ancestry from the same CHG-related source. It is possible that the source population was related to hunter-gatherers of the Seroglazivka culture who appeared around the lower Volga around 6200 BCE at sites such as Kair Shak III and Dzhangar. After about 5000 BCE the flow of this southern CHG-related ancestry increased significantly in the Samara region, perhaps with the spread of domesticated animals.

Cattle and sheep first appeared in the mid-to-late 6th millennium BCE in the Dnipro-Azov steppes. At Rozdol'noe on the Kalmius River 60km northeast of Mariupol a domesticated calf mandible was dated 5559 ± 45 calBCE (Ua-42031/6609 ± 49 BP) by Kotova and Anthony in 2017, suggesting that domesticated animals spread to the Volga from the west. A spread from the Caucasus is possible, but the earliest radiocarbon dated domesticates in the North Caucasus piedmont were associated with the Meshoko culture dated earliest at Unakozovskaya Cave (I1722, I2055, I2056) between 4676-4367 BCE, a millennium after Rozdol'noe.

Grave 97 (I20116) had the mutation marking the branching event at the root of the Yamnaya R1b Y-haplogroup subclade, (**R-P297**). This mutation also was present in some Latvian hunter-gatherers dated 5500-4500 BC (I4626-28, I434, I4636, I4439), Lyalovo hunter-gatherers on the upper Volga dated 5200 BC (MOS244, MOS225), and in the Samara hunter-gatherer dated 5500 BC (I0124), an upper Volga-Baltic-Samara distribution. It is surprising that the subsequent mutations in this important subclade (R-P297 -> R-M269 -> R-L23 -> R-Z2103) are not documented in our steppe sample set until their appearance in Yamnaya individuals more than 1000 years later.

Dates on terrestrial animals and organic residue

| Ekaterinovka Mys Grave # | Lab | Sampled material | Age BP | calBC (95%) | δ13C | δ15N |
|---|---|---|---|---|---|---|
| grave 40 | PSUAMS 8194 | beaver incisor | 5750 ± 25 | 4686-4505 | -20.7 | 6.7 |
| grave 45 | PSUAMS 4568 | goat tooth | 5680±20 | 4550-4450 | Nd | |

| Ekaterinovka Mys Grave # | Lab | Sampled material | Age BP | calBC (95%) | δ13C | δ15N |
|---|---|---|---|---|---|---|
| grave 101 | PSUAMS 8195 | sheep tooth | 6025 ± 40 | 5028-4798 | Nd | |
| Grave 60 | PSUAMS 8218 | marmot tooth | 5745±30 | 4689-4517 | Nd | |
| potsherd | SPb-2251 | organic residue | 5673±120 | 4795-4267 | Nd | |

#### Human dates and haplogroups

| Sample ID | Burial code | Ekaterinovka Mys 14C dates | Y-group | mt-group |
|---|---|---|---|---|
| I20114 | grave 94 | 5100-4400 BCE | P1-<br>P-P337 | U4a1 |
| I20116 | grave 97 | 5100-4400 BCE | R1b- R-P297 | n/a (<2x) |
| I20118 | grave 99 | 5100-4400 BCE | (female) | U5a1+@16192 |
| I20189 | grave 101/2 | 5311-5073 calBCE (6255±30 BP, PSUAMS-9699) | (female) | U4a2 |
| I20192 | grave 100 | 5297-5050 calBCE (6210±30 BP, PSUAMS-9700) | (female) | U4a2 |
| I23648 | burial80, Skeleton 1 | 5100-4400 BCE | (female) | U5a1+@16192 |
| I23649 | burial80, Skeleton 2 | 5100-4400 BCE | (female) | U5a2 |
| I23650 | burial80a, Skeleton 3 | 5100-4400 BCE | (female) | U5a1+@16192 |
| I23651 | burial 82 | 5100-4400 BCE | Q1b- Q-M930 | U4a |
| I23652 | burial 86 | 5100-4400 BCE | (female) | U4a2 |
| I3546 | LP17.1 | 5477-5331 calBCE (6442±34 BP, DeA-8214) | Q1b- Q-L53 | U5a1+@16192 |
| I6057 | grave 1 | 5372-5222 calBCE (6350±20 BP, PSUAMS-2871) | (female) | U4a2c |
| I6059 | Grave 17 | 5315-5215 calBCE (6295±25 BP, PSUAMS-2873) | (female) | U2e1f1 |
| I6060 | Grave 19 | 5461-5221 calBCE (6355±25 BP, PSUAMS-2900) | (female) | U4a1 |
| I6061 | Grave 20 | 5470-5226 calBCE (6375±25 BP, PSUAMS-2901) | (female) | U4a3 |
| I6062 | Grave 24 | 5324-5214 calBCE (6305±25 BP, PSUAMS-2874) | R1b- R-V1636 | U5a1i |
| I6063 | Grave 31 | 5468-5225 calBCE (6370±25 BP, PSUAMS-2875) | (female) | U5a1 |
| I6064 | Grave 39 | 5474-5321 calBCE (6415±30 BP, PSUAMS-2876) | R1b- R-L754 | U4a3 |
| I6066 | Grave 41 | 5313-5214 calBCE (6290±25 BP, PSUAMS-2878) | R1b- R-V1636 | U5a1+@16192 |
| I6068 | Grave 45 | 4578-4451 calBCE (5680±20 BP, PSUAMS-4568) | R1b- R-L754 | U5a1+@16192 |
| I6069 | Grave 49 | 5467-5223 calBCE (6365±25 BP, PSUAMS-2881) | (female) | U5a1+@16192 |
| I6101 | Grave 53 | 5312-5212 calBCE (6280±25 BP, PSUAMS-2882) | (female) | U4a1 |
| I8282 | B.2 | 5318-5214 calBCE (6300±25 BP, PSUAMS-4307) | Q1b- Q-M930 | U5a1 |
| I8283 | B.36 skull 2 | 5312-5212 calBCE (6280±25 BP, PSUAMS-4280) | R1b- R-L754 | U4d |
| I8284 | B.36 skull 3 | 5301-5054 calBCE (6220±30 BP, PSUAMS-4308) | (female) | U5a1+@16192 |
| I8285 | B.57 | 5324-5214 calBCE (6305±25 BP, PSUAMS-4309) | R1b- R-V1636 | U2e1b |
| I8286 | B.70 | 5459-5219 calBCE (6350±25 BP, PSUAMS-4310) | Q1b- Q-M930 | U4a1 |
| I8287 | B.71 | 5471-5228 calBCE (6380±25 BP, PSUAMS-4311) | Q1b- Q-M930 | U5a1 |
| I8289 | B.74 | 5368-5217 calBCE (6330±25 BP, PSUAMS-4313) | (female) | U5a1+@16192 |
| I8290 | B.77 | 5316-5134 calBCE (6270±25 BP, PSUAMS-4314) | (female) | U5a1b |

| Sample ID | Burial code | Ekaterinovka Mys 14C dates | Y-group | mt-group |
|---|---|---|---|---|
| I8740 | B.79 skeleton 2 | 5474-5326 calBCE (6420±30 BP, PSUAMS-9551) | R1b - ? | U5a1+@16192 |
| I20114 | grave 94 | 5100-4400 BCE | P1- P-P337 | U4a1 |
| I8738 | B.55 (34d) | 5100-4400 BCE | .. | n/a (<2x) |

*Korolev, A., A. Kochkina, and D. Stashenkov 2019. The Early Eneolithic burial ground at Ekaterinovsky Cape in the forest-steppe Volga region. Documenta Praehistorica XLVI: 388-397. DOI: 10.4312/dp.46.24*

*Korolev, A. I., A.F. Kochkina, D.A. Stashenkov, Aleksandr Khokhlov, & N.V. Roslyakova. 2018. Unikal'noe pogrbenie mogil'nika epokhi rannego Eneolita Ekaterinovskii Mys na Srednei Volge. Stratum plus (2): 285-302.*

*Kotova, Nadezhda. 2018. Revisiting the Neolithic chronology of the Dnieper steppe region with consideration of a reservoir effect for human skeletal material. Sprawozdania Archeologiczne 70: 47-66.*

#### 2.4 Golubaya Krinitsa

*Summary by D. Anthony*

The Golubaya Krinitsa site consists of three places with mixed occupation and graves on a sandy ridge west of the middle Don River (un-named in the map below but visible in the image on the right), on a tributary, the Chernaya Kalitva River. The sites were occupied in the Upper Paleolithic, Neolithic, Eneolithic, Bronze, & Iron Ages.

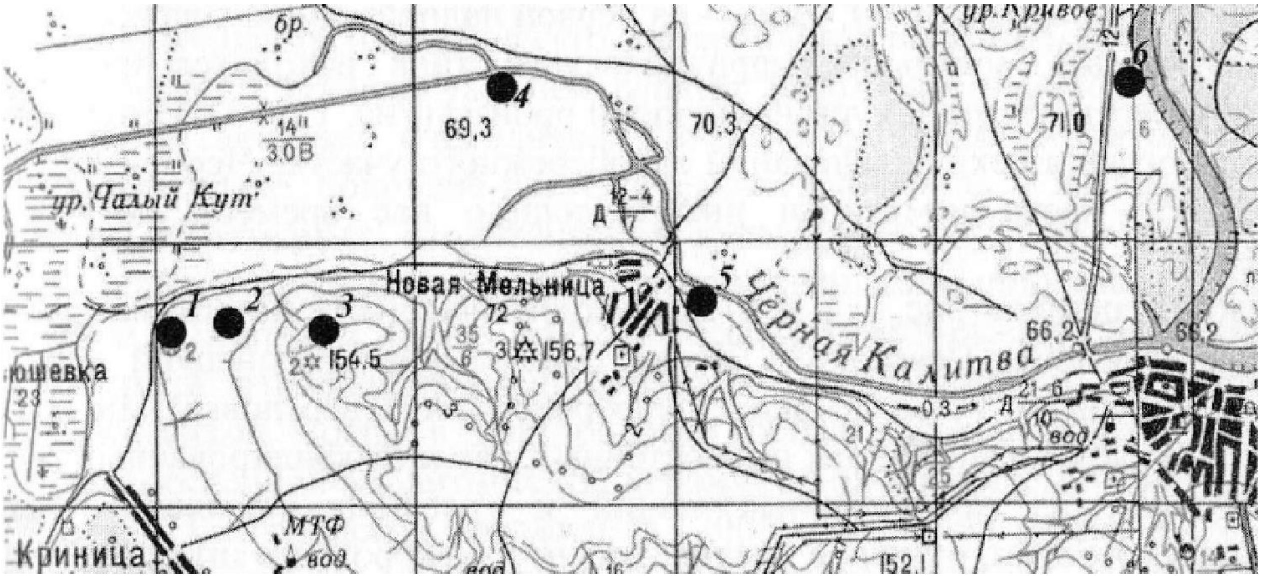

**Fig. 2.6. Three sites compose Golubaya Krinitsa (Blue Spring):** 1. Settlement with Paleolithic to Bronze Age artifacts and scattered graves of both Early (Mariupol type) and Late Eneolithic (Serednii Stih type) (1st publication of Golubaya Krinitsa was just this site); 2. Neolithic settlement; 3. Early Eneolithic cemetery (Mariupol type, Dnepr-Donets culture, sampled by Willerslev).

Two chronological-cultural components have been sampled for aDNA. Dnieper-Donets Mariupol-type objects occurred in graves at GK #3—boars tusk pendants (map) and long flint blades were deposited with burials in supine extended position. The excavators ascribed these graves to influence from the Dnipro Neolithic of Dereivka I type.

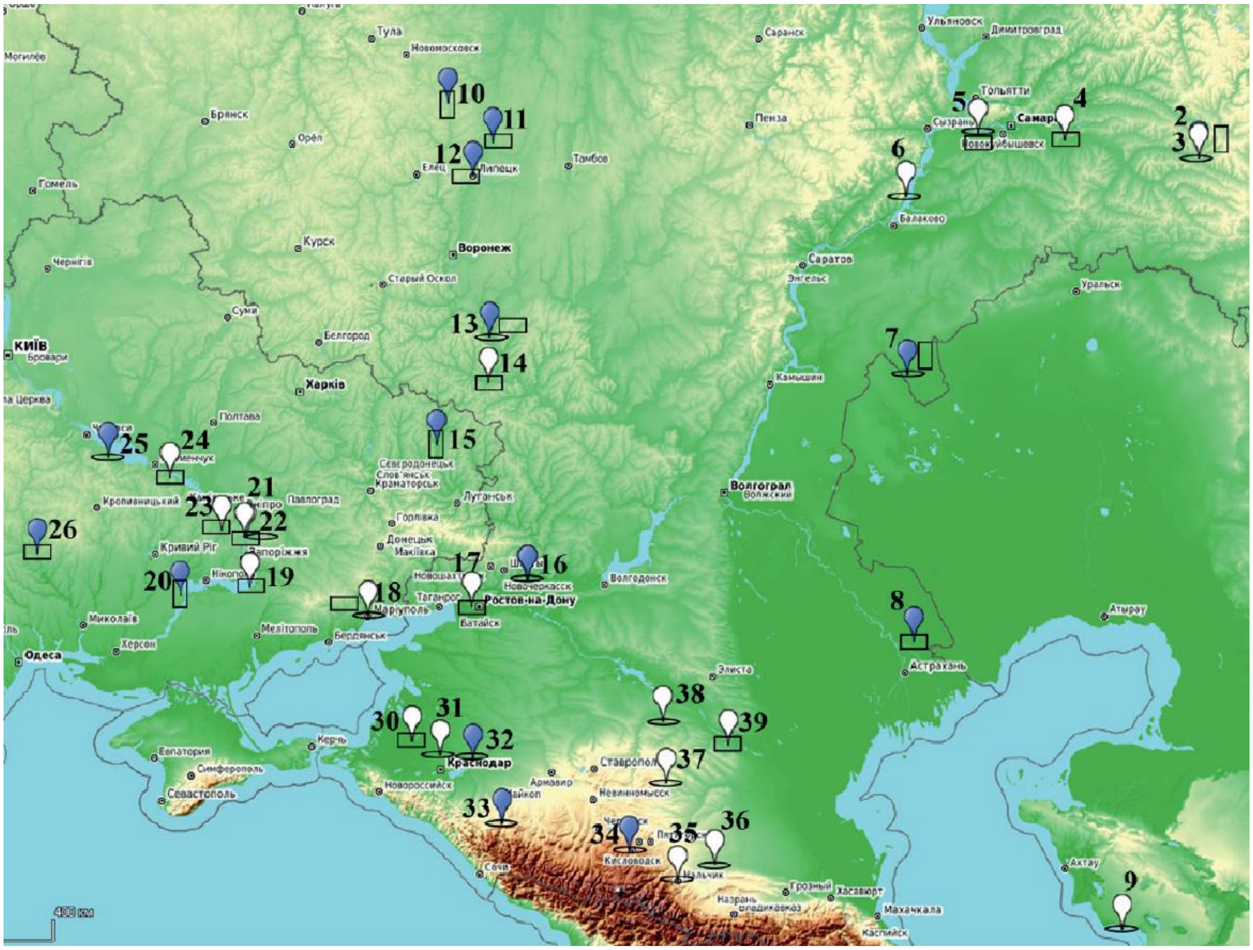

**Fig. 2.7. Boars tusk ornaments 4700-4000 BCE.** 14-Golubaya Krinitsa; White, cemeteries; Blue, settlements. Flat Rectangle, cut rectangular tusk plaque; Cigar-shape, lunate whole tusk; Upright Rectangle, weapon: From Skorobogatov 2020.

1 – Mellyatamak V; 2 – Ivanovskaya; 3 – Krasnoyarka; 4 – S"ezzhee; 5 – Ekaterinovskii Mys; 6 – Khvalynskie I and II; 7 – Varfolomeevka; 8 – Kair-Shak III; 9 – Tokmak; 10 – Dolgoe; 11 - Vasil'evskii Kordon 17; 12 – Lipetskoe Ozero; 13 – Cherkasskaya; 14 – Golubaya Krinitsa; 15 – Podgorovka I; 16 – Razdorskoe I; 17 – Liventsovskii; 18 – Mariupol'skii; 19 – Lysogorskii; 20 – Mikhailovka; 21 – Yasinovatskii; 22 – Petro-Svistunovo; 23 – Nikol'skii; 24 – Dereivskii; 25 – Molyukhov Bugor; 26 – Pugach-2; 27 – Vykhvatinskii; 28 – Bernashevka; 29 – Luka-Vrublevetskaya; 30 – Staronizhesteblievskaya; 31 – Novotitarovskaya; 32 – Svobodnoe; 33 – Meshoko; 34 – Zamok; 35 – Nal'chikskii; 36 – Chernoyarskaya; 37 – Vesyolaya roshcha; 38 – Aigurskii; 39 – Chograi.

GK also had later, Serednii Stih-type graves supine with raised knees, with Serednii Stih-type pottery and early-to-late 4th-millennium BCE dates. Settlements and cemeteries with similar Serednii Stih materials spread up the Don River ca. 4200-4000 BCE. GK was located between the steppe Serednii Stih sites on the lower Don, where Serednii Stih might have first evolved according to Kotova; and the Serednii Stih sites that spread up the middle and upper Don, replacing the Pit & Comb ware Middle Don Neolithic culture (not present at GK).

People displaying local variants of Serednii Stih ceramics, tools, ornaments, & funeral culture lived as hunter-gatherers without domesticated animals on the middle and upper Don into the early Yamnaya period, as at Vasilievskii Kordon (11 on map) near Lipetsk dated 3700-3300 BCE, 300km north of GK. Repin style pottery on the lower Don retained elements of middle Don Serednii Stih pottery and indeed this could be the origin of the Repin style in the steppes. In the steppes the Repin pottery style was the only kind of pottery at Repin on the lower Don and was 10% of the whole vessels at Early Yamnaya Mikhailovka stratum II on the lower Dnipro, both sites dated 3400-3100 BCE.

##### *Genetics*

Two GK samples (I12490 and I12491) are reported here, and Allentoft et al 2022 reported six more (NEO113, NEO204, NEO207, NEO209, NEO210, NEO212).

One of our samples (GK2) has affinity with Ukraine_N and the other (GK1) with Serednii Stih, so it seems we sampled one of each of the Neolithic/Eneolithic components present at GK.

*Allentoft et al. 2022 found two Y I2a males at GK, and one male R1a (R-M459). They commented that this R1a was ancestral to Corded Ware R1a. Both males from GK that we sampled had Y R1a (R-M459). GK contained both Y I2a, like Ukraine _N and Serednii Stih, and pre-CWC R1a, as did one Khvalynsk and one Steppe Maikop male.*

| Sample ID | Burial code | Golubaya Krinitsa | Y-group | mtDNA |
|---|---|---|---|---|
| I12491 | GK grave 8 juvenis M | 5557-5381 calBCE (6520±30 BP, PSUAMS-8936) | R1a-M459 | U5a2b |
| I12490 | GK16-gr 15 | 5610-5390 calBCE (6535±30 BP, PSUAMS-8934) | R1a-M459 | U4b1 |

*Allentoft, Morten E. et a. 2024. Population genomics of post-Glacial western Eurasia.*

*https://doi.org/10.1038/s41586-023-06865-0*

*Skorobogatov A. M. 2020. Neolithic – Eneolithic wild boar tusk artifacts found in the East European steppe and forest-steppe (in Russian). In: Bulletin of the Moscow Regional State University. Series: History and Political Sciences, 2020, vol. 5 Circumpontica II: 10–27. DOI: 10.18384/2310-676X-2020-5-10-27 (source of map)*

*Skorobogatov, A.M. & R.V. Stol'ianinov 2013. Srednestogovske materialy v basseine verkhnego i srednego Dona. Rossiiskaya Arkheologiya 2: 126-136.*

#### 2.5 Khlopkov Bugor

*Summary by D. Anthony & L. Vyazov*

The Khlopkov Bugor Eneolithic cemetery was located on a high bluff overlooking the Volga on its west bank 130km south of the Khvalynsk cemetery. Riverine erosion partly destroyed it before rescue excavations recovered 15 graves. The bodies were posed supine with raised knees, like all Volga Cline graves dated after 4500 BCE. Initial analysis identified 6 males, 7 females, and two adults too fragmentary to determine age. No subadults and no aged individuals were recovered—only young and mature adults.

Khokhlov (2010) was able to analyze 5 males (4a, 5, 6, 10, 14) and 6 females (4b, 7, 8, 12, 13, &

15) by osteological criteria. Experts disagreed on the indicators of sex in Grave 6 (I6302), designated a male in 1986, revised to a female in 1988, returned to a male two decades later (Khokhlov 2010:439), and now confirmed to have been genetically male. The Grave 6 male was 40-45 years old at death and genetically was the father of the female in Grave 15 (I6300), aged 25-30. Two other individuals sampled for DNA were females unrelated within 3 degrees; neither could have been 15's mother, but 8 (I6905) could have been a maternal relative of 6's mother. The square-jawed female in Grave 7 (I6301) was a second-degree relative, probably the paternal grandmother, of the male I6107 in grave 4 at Khvalynsk II.

The KB graves contained typical Khvalynsk ceramic vessels, beads made of shell, bone, and stone, and two polished stone maces, one vaguely zoomorphic, the other an eared oblong type with D-shaped ridges carved into its sides interpreted by some as symbolic harness markings. Similar maces with D ridges are found in the Samara region, in several Eneolithic graves in the North Caucasus steppes (Dzhangar, Ulan Tolga), and in settlements of the Trypillia B1 culture. No copper was found at Khlopkov Bugor, an absence interpreted to mean that the Balkan copper trade had not yet reached the Volga.

During the brief interval between Khlopkov Bugor and Khvalynsk, between a grandmother (KB7) and her grandson (Khvalynsk II:4), perhaps around 4500 BCE, Balkan copper began to flow through exchange relationships in the Volga steppes and was concentrated at Khvalynsk.

Three of the four individuals that passed aDNA screening were females, and the only male had too low a SNP count to determine the Y-haplogroup.

| Sample ID | Burial code | Radiocarbon date | Y-group | mtDNA |
|---|---|---|---|---|
| I6301 | Grave 7 | 5213-5035 calBCE (6170±25 BP, PSUAMS-2908) | female | U5a1a1 |
| I6300 | Grave 15 Da of I6302 | 5215-5045 calBCE (6185±25 BP, PSUAMS-2907) | female | U5a1f1 |
| I6302 | Grave 6 Fa of I6300 | 5292-5032 calBCE (6190±35 BP, PSUAMS-2949) | M too few SNP's | H13a1a |
| I6905 | Grave 8 | 5209-4995 calBCE (6135±25 BP, PSUAMS-4256) | female | H13a1a |

*Khokhlov, A. A. 2010. Naslenie Khvalynskoi Eneoliticheskoi kul'tury po antropologicheskim materialam gruntovykh mogil'nikov Khvalynsk I, Khvalynsk II, Khlopkov Bugor. In: S. A. Agapov (ed.) Khvalynskie eneoliticheskie mogil'niki i Khvalynskaya Eneoliticheskaya kul'tura: Issledovanija materialov. SROO IEKA "Povolzh'e" (Samara 2010) 407–583.*

#### 2.6 Khvalynsk

*Summary by D. Anthony with help from D. Agapov and N. Shishlina*

Khvalynsk is the largest excavated Eneolithic cemetery in the Don-Volga-Ural steppes (201 recorded graves). More copper artifacts were found (373 objects) than at other late fifth millennium BC steppe cemetery and more sacrificed animals were found (at least 106 sheep-goat, 29 cattle, and 16 horses). Before flooding by the Saratov reservoir, it was located on the right (west) bank of the Volga and was divided into two burial plots (1 and 2). Khvalynsk-1 was excavated in 1977-79 and contained 158 excavated individuals. Erosion from the reservoir claimed an unknown number of additional graves. Khvalynsk-2 was located about 120m SW of 1 and was salvaged in 1987-89 as the Volga bank continued to erode westward. It yielded 43 recovered individuals. Both were in use simultaneously by people with the same rituals, pottery, stone tools, and copper artifacts—the same 'culture'. It was separated intentionally into two burial plots. Much of the Khvalynsk-1 collection was lost after a flood in a storage facility, so aDNA was obtained from only 5/158 individuals, while 2 is largely preserved and yielded 26/43 individuals useful for aDNA analysis.

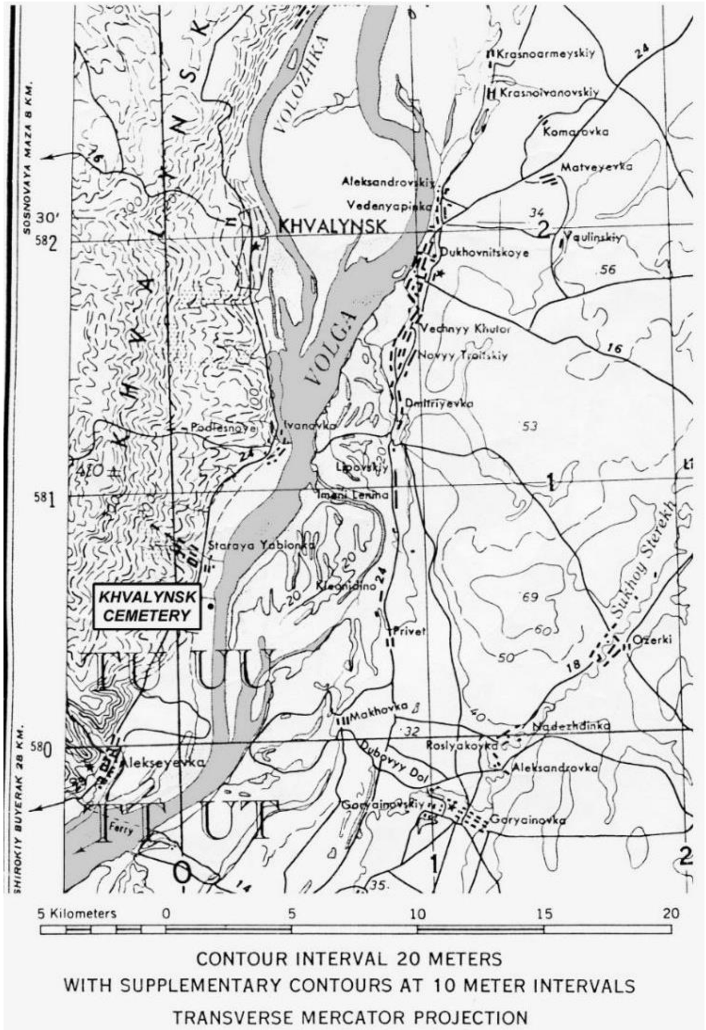

**Fig. 2.8. Pre-dam topographic location of Khvalynsk.**

Khvalynsk-1 was four times larger than 2 and had males and females in about equal proportions according to skeletal analyses. But only one male at Khvalynsk-1 was in the 18-30 age range, prime military age. Three older males at Khvalynsk I were buried with four polished stone maces. All three also had copper ornaments and sacrificed domesticated mammals, two distinctions accorded to less than 15% of individuals at 1.

At Khvalynsk-2, 20 of 26 (80%) adults assigned a skeletal sex by Khokhlov were males, a very different sex profile. Seven of the 20 were in the 18-30 age range, suggesting that males in this age range were intentionally buried at Khvalynsk-2. If 1 and 2 came from the same population then 87.5% of males in this age range were buried at 2. One third of the 43 individuals at 2 were subadults, most of them under age 7. Perhaps males of prime military age were buried in II with selected wives and children. Males and females in 2 had 10x more Balkan copper items than was found in 4x more graves in 1, and the richest grave at 2 also was furnished with a polished stone mace, and cattle, sheep, and goat sacrifices. Khvalynsk-2 could have been a burial plot for a male sodality with enhanced access to copper, perhaps warriors, buried with selected wives.

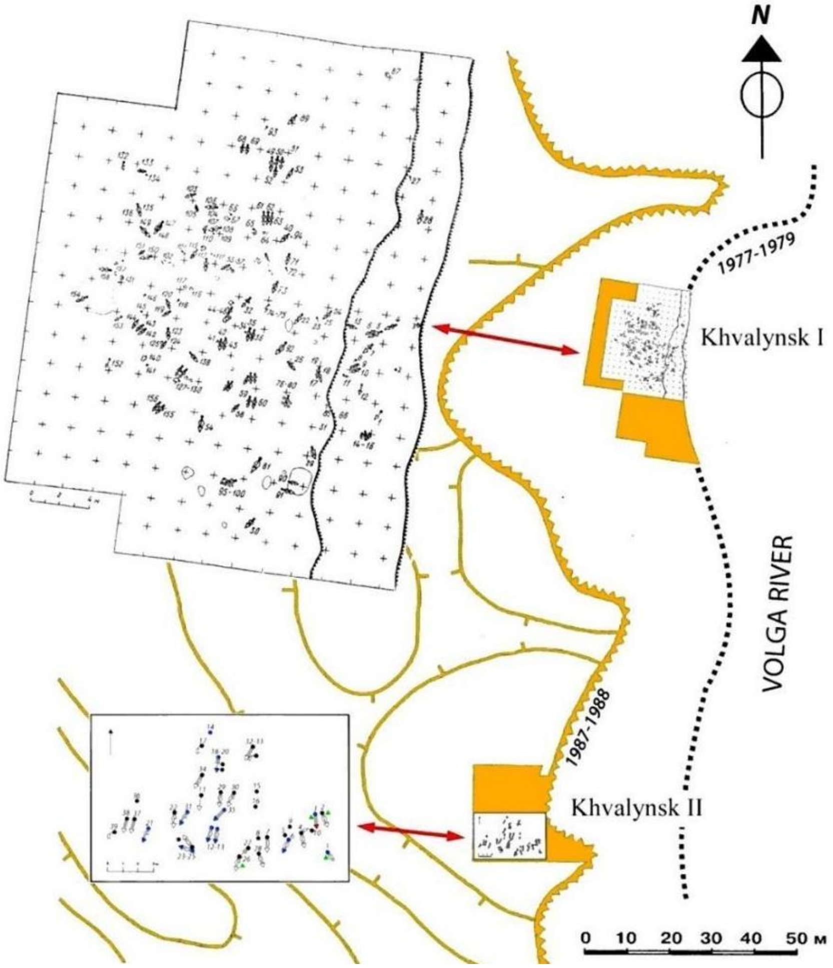

**Fig. 2.9. Khvalynsk 1 and 2. Situation plan.**

Radiocarbon dates from terrestrial animal bones from I and II were tightly grouped between 4450-4250 BCE (Shishlina et al., Radiocarbon, 2009). A sheep-goat bone made into a ring was dated 4450-4355 BCE (GrA-29178, 5565±40 BP) from grave 147 at Khvalynsk-1; and a cattle bone was dated 4448-4362 BCE (GrA-34100, 5570± 40 BP) from grave 10 at Khvalynsk II. These dates are now confirmed by four more dates obtained on other ungulate bone artifacts from Khvalynsk-1 by N. Shishlina and colleagues. They are 4340-4251 BCE (DeA 38095), 4338-4248 BCE (IGANams 9188), 4440-4332 BCE (IGANams 9189), and 4341-4372 BCE (Poz 137344) (Shishlina 2023).

The standard burial pose was supine with tightly raised knees, oriented NW, N, or or NE, with some exceptions. Red ochre was deposited by the head and feet. This orientation, pose, and use of red ochre were also standard in Sredenii Stih graves and were retained later in Yamnaya culture graves.

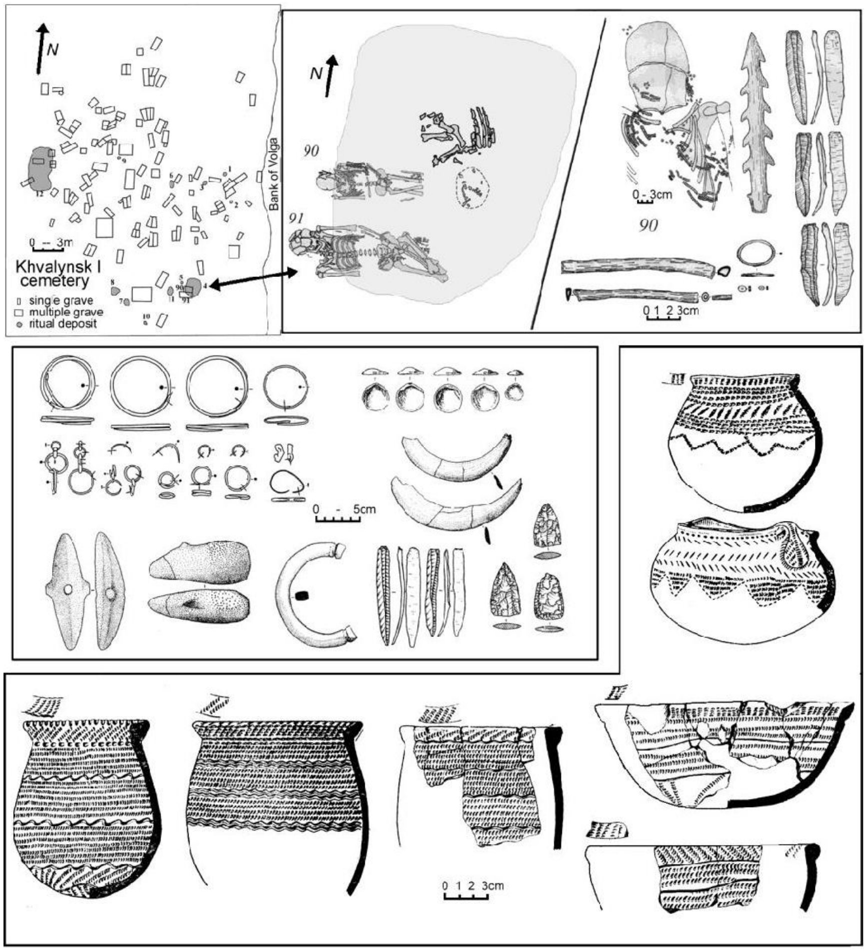

**Fig. 2.10. Khvalynsk I plan and objects. Top:** cemetery plan and Sacrificial Deposit 4 containing bones of 2 cattle, 1 sheep-goat, & 1 horse above Graves 90 & 91 with bird-bone tube, antler harpoon, flint blades, and copper ring. **Middle:** grave artifacts including the broken mace and whole mace from grave I:108, a polished stone bracelet probably from Svobodnoe in the North Caucasus, & fossil *Glycemeris* shell ornaments; **Bottom:** ceramic pots and bowls from Khvalynsk I. From Anthony 2007: Figure 9.7.

Copper was used for ornamental rings, beads, and pendants. It had trace elements most like Balkan copper from Ai Bunar, and is thought to have been imported from the Varna and Trypillia B1 cultures in the form of both finished objects and lumps of smelted copper, the latter found in two graves as well as in the cemetery cultural level outside the graves. It then was worked and welded into beads and other ornaments by local steppe craft workers, who copied Trypillia welding methods but displayed less control over temperature (imperfect welds).

##### *Patricentered families*

aDNA analysis of 5 individuals from 1 revealed none related within 3 degrees. Of the 26 analyzed at 2, 18 (69%) were related to at least one other individual. 17 of 18 relatives at Khvalynsk-2 were males; the lone related female was the sister of the male with whom she was buried in the richest mace-grave (2:24 & 25). None of the other 5 females in 2 was related to anyone, while 17 (89%) of the 19 DNA-assigned males had at least one relative in 2. Among these relatives were three father-son pairs, and probable brother’s sons (2:22,27) were buried near their probable paternal uncles (2:12, 13).

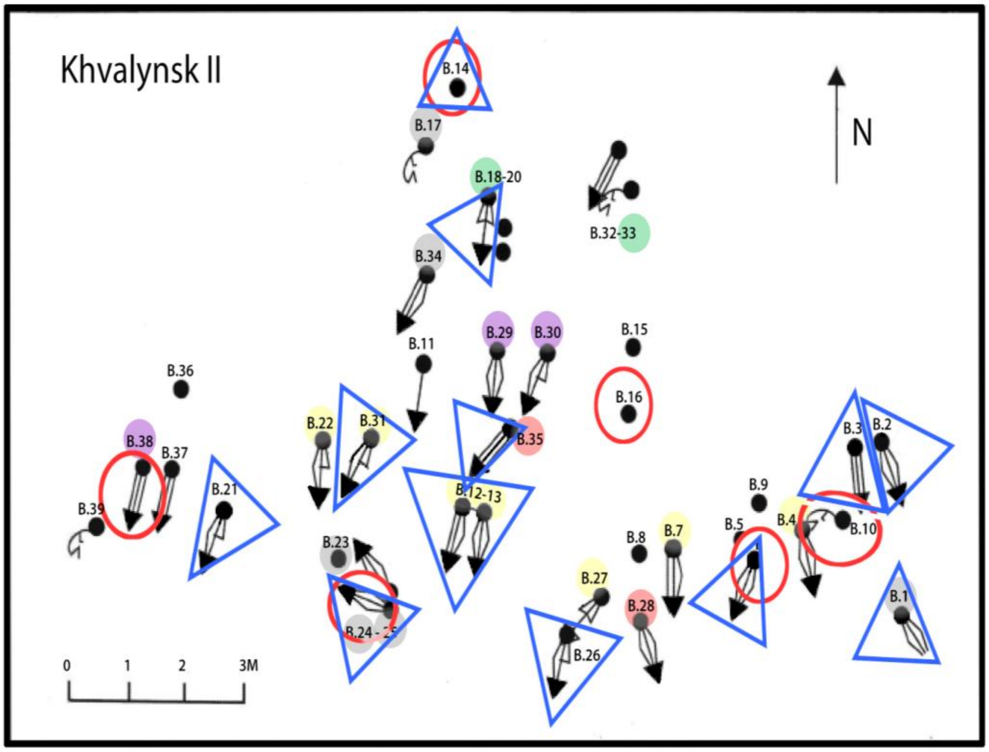

**Fig. 2.11. Khvalynsk II with family relationships indicated by color superimposed on copper finds (blue triangle) and animal sacrifices (red circle).** Icons with straight lines were supine with raised knees; icons with curved lines were half-sitting with raised knees; black circles were isolated skulls. Related individuals are color coded with five colors: yellow (dominant), green, grey, orange, & purple. 24 (grey, I6407) buried with his sister (25, I6734) and the skull of a 3rd-degree paternal male relative 23 (I20188) at their feet, was the richest grave at Khvalynsk, and the only one oriented SE. From D. Agapov 2010: figure 5 and Anthony et al. 2022.

No mothers or daughters, and only one sister of the Khvalynsk II males were present, supporting its interpretation as a multi-generational burial plot for a male sodality, drawn from a limited set of families mostly affiliated with the R1b-V1636 Y-haplogroup, but including individuals with R1a-M459 (like later Corded Ware), Q1a-L472 (like Murzikha to the north), 12a-L699 (a widespread Serednii Stih haplogroup), and J1-CTS1026 (unrelated to anyone).

A male from grave 4 at Khvalynsk-2 (I6107) was a 2nd-degree relative of an adult female (I6301) buried in grave 7 at Khlopkov Bugor, 130km to the south on the Volga. The absence of copper in 15 graves at Khlopkov Bugor suggests that it was earlier than Khvalynsk. If so, then the I6107 male from Khvalynsk was the descendant of the female in KB7, who would in this case have been his paternal (because they had different mt-haplogroups) grandmother, testifying to upriver movement.

*Multi-regional population*

Khvalynsk was extraordinarily diverse genetically, containing individuals with affinity to Ekaterinovka Mys at the northern end of the Volga Cline and to Progress-2 and Vonjuchka at the southern end, 1000km apart. Cranio-facial types also included broader ‘northern’ faces and narrower ‘southern’ ones. Variation in δ13C also seems to identify female mt-haplogroups from different Volga riverine catchments that differed in their depletion in carbon. δ13C was correlated with some mtDNA haplogroups according to Schulting in Anthony et al. 2022, suggesting that some females at Khvalynsk came from distinct riverine catchments with differing concentrations of δ13C. Khvalynsk was a meeting-place and cemetery used by groups from many places in the Volga River valley and beyond.

##### *Early horse domestication*

An early stage of horse management in the Khvalynsk period was indicated by horse DNA from Dnipro-Don-Volga steppe sites published in Librado et al. (2021) showing a shift away from wild horses designated NEO-NCAS, from hunter-gatherer archaeological sites with all wild fauna dated 5600-5200 BCE, toward DOM2 in horses in Eneolithic sites with some domesticated cattle & sheep dated 4700-4000 BCE. Horses from Eneolithic Semenovka (Librado's Ukr11_Ukr_m4185) in the lower Dnipro steppes and Oroshaemoe I (RN96_Rus_m4586) in the lower Volga steppes were shifted towards DOM2, perhaps showing early selection for domesticated genetic traits.

In addition, horses were sacrificed and buried with cattle, sheep-goats, and humans at Khvalynsk where no obviously wild mammals were included; polished stone mace-heads shaped liked horse heads proliferated across the steppes and spread into the lower Danube valley between 4400-4000 BCE; and a migrant from the Volga Cline who was buried at Csongrad, Hungary (I5124) had the lower trunk and pelvic musculature of a habitual rider dated 4442–4243 calBCE (5470 ± 40 BP/ Poz-41865) (Trautmann et al. 2023). Eneolithic horses, even if they were more skittish than modern horses, might have been ridden in quiet settings such as herding, where they permitted a mounted shepherd to oversee three times more sheep than a pedestrian shepherd, producing a surplus useful in hosting feasts.

*Agapov, S.A. (ed.) 2010. Khvalynskie eneoliticheskie mogil'niki i Khvalynskaya Eneoliticheskaya kul'tura: Issledovanija materialov. SROO IEKA "Povolzh'e" (Samara).*

*Anthony, David W., A. A. Khokhlov, S. A. Agapov, D. S. Agapov, R. Schulting, and D. Reich. 2022. The Eneolithic cemetery at Khvalynsk on the Volga River. Praehistorische Zeitschrift 97(1): 22-67. https://doi.org/10.1515/pz-2022-2034.*

*Shishlina, N.I., J. van der Plicht, & M.A. Turetsky 2018. The Lebyazhinka burial ground (middle Volga region, Russia): new 14C dates and the reservoir effect. Radiocarbon 60 Special Issue 5(2): 681–690. DOI:10.1017/RDC.2017.94*

*Shishlina, N.I. 2023. Khvalynskii i Eneoliticheskii mogil'nik: Novye radiouglorodnye daty. Voprosy Arkheologii Povolzh'ya (Samara) v. 10.*

*Trautmann et al. 2023. First Bioanthropological Evidence for Yamnaya Horsemanship. Science Advances 9.1–13*

| Sample ID | Burial code | Radiocarbon dates on humans | Y-group | maDNA |
|---|---|---|---|---|
| I0122 | Khvalynsk II Grave 12 | 4936-4730 calBCE (5960±25 BP, PSUAMS-4031) | R1b-<br>R-V1636 | H2a1 |
| I0426 | Khvalynsk II, Grave 32 | 4550-4300 | (female) | U4a |
| I0433 | Khvalynsk II Grave 1 | 4697-4539 calBCE (5760±25 BP, PSUAMS-4032) | R1a-<br>R-M459 | U5a1i |
| I0434 | Khvalynsk II Grave 17 | 5198-4853 calBCE (6070±25 BP, PSUAMS-4033) | Q –<br>Q-L472 | U4 |
| I11837 | Khvalynsk I Grave 40 | 4550-4300 | R1b-<br>R-L754 | U4a1c |
| I6102 | Khvalynsk I Grave 17 | 4703-4547 calBCE (5775±25 BP, PSUAMS-2883) | (female) | T2a1b |
| I6103 | Khvalynsk I Grave 30 | 4983-4795 calBCE (5995±25 BP, PSUAMS-2884) | I2a- | U5a2d |

| Sample ID | Burial code | Radiocarbon dates on humans | Y-group | maDNA |
|---|---|---|---|---|
| | | | I-L699 | |
| I6104 | Khvalynsk I Grave 127 | 4537-4362 calBCE (5625±25 BP, PSUAMS-2885) | R1b-<br>R-L389 | U2e1a1 |
| I6105 | Khvalynsk I Grave 147 | 4449-4352 calBCE (5565±25 BP, GrA-29178) | (female) | U2e1b |
| I6106 | Khvalynsk II Grave 2 | 4940-4790 calBCE (5975±25 BP, PSUAMS-2902) | (female) | T2a1b |
| I6107 | Khvalynsk II Grave 4 | 4935-4786 calBCE (5965±20 BP, PSUAMS-2903) | R1b-<br>R-L389 | U5a2d |
| I6108 | Khvalynsk II Grave 6 | 5204-4905 calBCE (6085±25 BP, PSUAMS-4250) | (female) | U4a |
| I6109 | Khvalynsk II Grave 7 | 4836-4715 calBCE (5900±25 BP, PSUAMS-4148) | R1b-<br>R-L389 | U5a1+@16192 |
| I6110 | Khvalynsk II Grave 10 | 4491-4342 calBCE (5570±40 BP, GrA-34100) | (female) | U4 |
| I6299 | Khvalynsk II Grave 18 | 5209-4958 calBCE (6125±20 BP, PSUAMS-2906) | Q1a-<br>Q-YP1669 | U2e2a1 |
| I6402 | Khvalynsk II Grave 29 | 4789-4613 calBCE (5840±25 BP, PSUAMS-4150) | R1b-<br>R-L754 | H2a1 |
| I6403 | Khvalynsk II Grave 13 | 4945-4792 calBCE (5985±25 BP, PSUAMS-4200) | R1b-<br>R-L754 | H2a |
| I6404 | Khvalynsk II Grave 19 | 5311-5084 calBCE (6260±25 BP, PSUAMS-4151) | (female) | U5a1 |
| I6405 | Khvalynsk II Grave 21 | 4934-4728 calBCE (5955±25 BP, PSUAMS-4152) | R1b-<br>R-L754 | U5a2d |
| I6406 | Khvalynsk II Grave 22 | 4929-4726 calBCE (5950±25 BP, PSUAMS-4153) | R1b-<br>R-L754 | U4d |
| I6407 | Khvalynsk II Grave 24 | 4983-4795 calBCE (5995±25 BP, PSUAMS-4154) | Q1a-<br>Q-YP1669 | U2e1b |
| I6408 | Khvalynsk II Grave 35 | 5209-5006 calBCE (6150±25 BP, PSUAMS-4155) | R1b-<br>R-L754 | U4a1 |
| I6412 | Khvalynsk II, Grave 38 | 4695-4508 calBCE (5755±25 BP, PSUAMS-4156) | R1b-<br>R-L754 | H13a2a |
| I6734 | Khvalynsk II Grave 25 | 4678-4494 calBCE (5730±25 BP, PSUAMS-4162) | (female) | U2e1b |
| I6735 | Khvalynsk II Grave 26 | 5206-4935 calBCE (6100±25 BP, PSUAMS-4163) | J1-<br>J-CTS1026 | U4a |
| I6736 | Khvalynsk II Grave 27 | 4987-4797 calBCE (6000±25 BP, PSUAMS-4304) | R1b-<br>R-L754 | U5a1a1 |
| I6737 | Khvalynsk II Grave 28 | 4783-4555 calBCE (5820±25 BP, PSUAMS-4545) | R1b-<br>R-L754 | U5a1a2 |
| I6738 | Khvalynsk II Grave 31 | 4889-4722 calBCE (5930±25 BP, PSUAMS-4305) | R1b-<br>R-V1636 | R1b1 |
| I6739 | Khvalynsk II Grave 33 | 4540-4369 calBCE (5640±25 BP, PSUAMS-4164) | Q1a-<br>Q-YP1669 | U2e2a1 |
| I6740 | Khvalynsk II Grave 34 | 5206-4909 calBCE (6095±25 BP, PSUAMS-4306) | Q1a-<br>Q-YP1669 | U5a1i |
| I6741 | Khvalynsk II Grave 30 | 4945-4792 calBCE (5985±25 BP, PSUAMS-4223) | R1b-<br>R-L389 | U4b1+293+138<br>34 |

#### 2.7 Krivyanskiy-9, kurgan 1:19

*Summary by David Anthony & Anatoly Faifert*

An Eneolithic grave was located on a low promontory overlooking the northern edge of the lower Don floodplain meadows. Similar burials reportedly were found further along the promontory. The northern edge of Kurgan 1, built in the Early Bronze Age (EBA), covered the Eneolithic grave. Kurgan 1 was 46m in diameter, and kurgans 2 and 5 were half that size, built close to 1. All three contained graves dated to the EBA and of the MBA Catacomb culture. The Eneolithic individual found in grave 19 under kurgan 1 was submitted for DNA analysis .

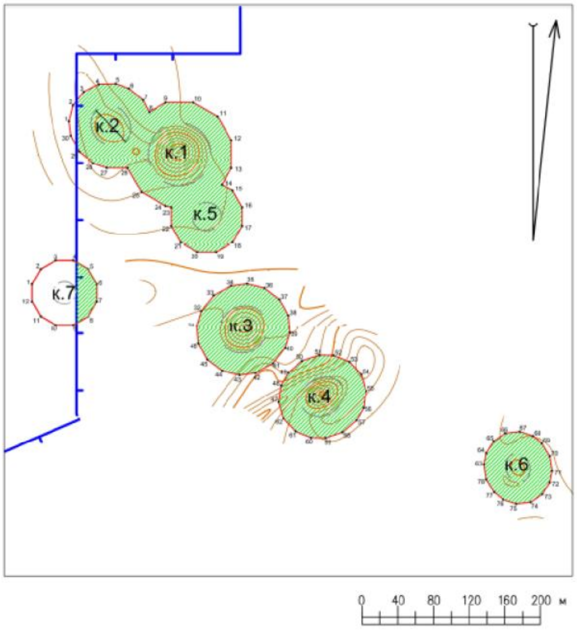

**Fig. 2.12. Schematic preliminary plan of kurgans 1-7 at Krivyanskiy-9. Final plan may be different.**

##### *2.7.1 Burial 19 (individual ID I11828): 4359-4251 calBCE*

When cleaning the west side of the central N-S baulk of Kurgan 1, 16 m north of its center, under the northern margin of the kurgan, grave 19 was discovered containing a burial of the Eneolithic era. The burial pit was round in plan with a concave bottom; many red ochre grains were on the floor. The grave was partially destroyed by a bulldozer.

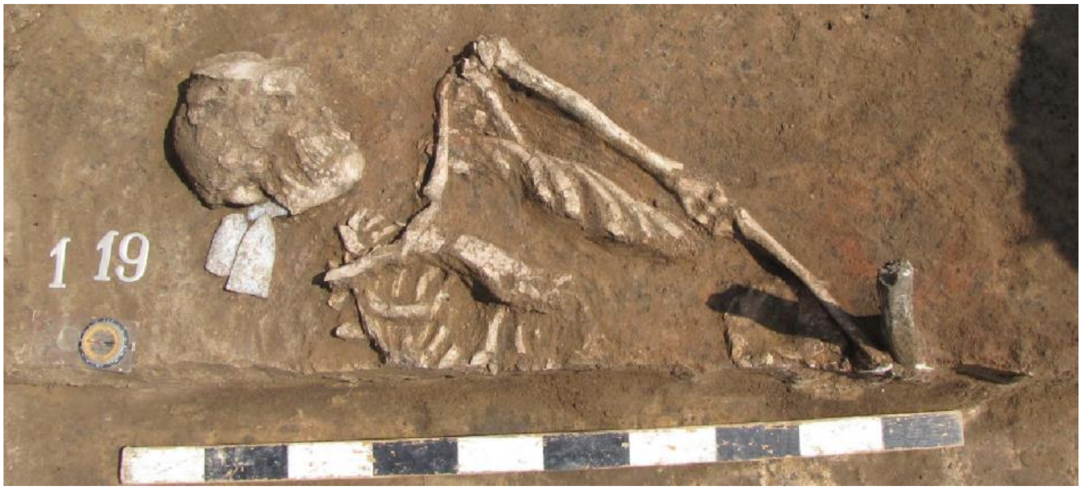

**Fig. 2.13. Grave 19 preserved in the baulk; lower part was cut by bulldozer.**

At the bottom of the pit lay the skeleton of an adult man, on his back, knees raised, with the skull towards the NNE. The left arm was extended, the left thigh was raised (the top of the femur was cut off in ancient times by a later grave pit). The burial pose, artifacts, and date are typical of the

Serednii Stih culture, broadly defined.

The burial is dated to 4359-4251 calBCE (5465±30 BP, PSUAMS-7866).

Inventory:

1. A retouched flint blade displaced by burrowing rodents was found in the fill 10 cm above the floor of the grave. It retained a small bulb of percussion. Its tip was broken and missing, but was found in a deeper level (see #3). One edge exhibits small semi-circular flake scars from retouching, with a noticeable gloss. The flint is grayish-brown in color, translucent but patinated. Dimensions: 125x25x6 mm.
2. On the right side of the skull were two large bifacial flint projectile points and an axe-shaped blank. They were elongated triangles in shape with edges tapering to a point in the upper third. The base was straight and thinned for hafting. Large, shallow flake scars covered both surfaces. The edges were sharpened with deep, shallow flake scars followed by fine edge retouch, forming a finely ribbed surface at the edges. The flint is brown, translucent, with inclusions of small organic residues, but patinated in places. Dimensions were 96x48x11 mm. and 78x34x6 mm.
3. The axe-shaped flint blank is bifacial, sub-trapezoidal in shape. The cortex is preserved on one face. The edges are sharpened with a large, flat, long flakes followed by fine retouch. The flint is brown, translucent, with inclusions of small organic residues. According to its technical and morphological traits, it could be made into either a chopping tool or a projectile point. Dimensions: 77x49x13 mm.

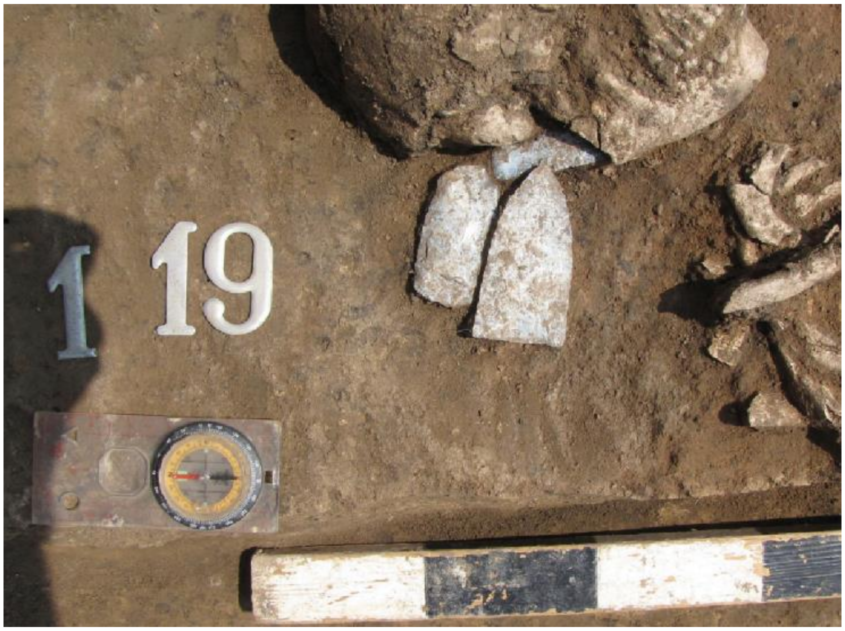

**Fig. 2.14. Grave 19, detail.**

4. Displaced into soils beneath the grave floor were another lamellar unifacial flint blade and the broken tip of the first lamellar flint blade. The flint blade displayed varying color ranging from light gray to white, with small white inclusions. There is a mottled patina on the ventral surface. Dimensions: 70x34x8 mm.

The I31755 male had a Caucasus-derived Y-haplogroup, J2a J-M319, variants of which were shared with Aknashen and Maikop, but he lacked the Aknashen-type Neolithic CHG and instead exhibited only the older CHG variant related to Mesolithic CHG, like the Berzhnovka/Progress-2 population. His paternal ancestry was rare in the sampled steppe populations. His mt-haplogroup, T2a1b, was widespread among steppe women, found in Ukraine Neolithic, Serednii Stih and Volga Cline groups. In PCA the Krivyanskiy-9 male was very close to the Yamnaya cluster although not in the Yamnaya clade.

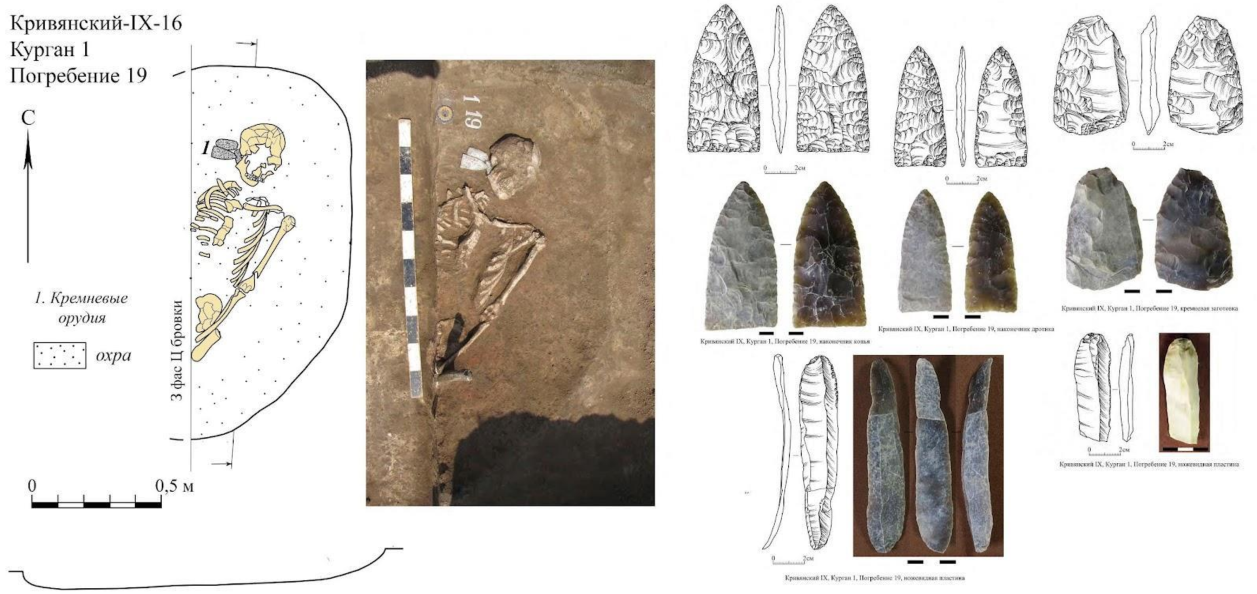

**Fig. 2.15. Burial 19 at Krivyanskiy-9** (image by Anatoly Faifert).

| Sample ID | Burial code | Radiocarbon date | Y-group | mtDNA |
|---|---|---|---|---|
| I31755 | Kurgan 19 Grave 1 | 4359-4251 calBCE (5465±30 BP, PSUAMS-7866) | J2a1a1a2b1b (J-M319) | T2a1b |

#### 2.8 Lebyazhinka-5

*Summary by D. Anthony & L. Vyazov*

The Sok River is a left-bank tributary of the Volga north of Samara that marks the NW margin of the elevated plateau known as the Obshchiy Syrt, a steppe tableland that projects from the southwestern Urals and separates the Volga and Ural River drainages. On the Sok left or south bank at least three Eneolithic settlement sites and several burial places were found near Lebyazhinka village. At Lebyazhinka the foot of a 150m-high ridge on the opposite bank projects into the wide marshes around the lower Sok, making this a convenient place to cross, as the modern bridge testifies.

**Lebyazhinka-1** is a multi-phase occupation site beginning in the Early Neolithic (early Elshanka-type ceramics). **Lebyazhinka-3** was a single-period settlement dated 4700-4500 BC and occupied by Early Eneolithic hunter-fishers using Samara Neolithic-style pottery (like that at S'yezzh'e, see this SI) who left only wild animal bones. Late Eneolithic **Lebyazhinka-6** was a single-period settlement dated 4100-3800 BC with Khvalynsk-style pottery, and about 33% of the fauna were bones of domesticated cattle and sheep-goat.

At **Lebyazhinka-4** a male known in the literature as the Samara Hunter-Gatherer (I0124) was buried in a multi-phase Neolithic (hunter-gatherer with ceramics) and Eneolithic (Khvalynsk-culture) settlement site that contained one extended supine burial dated 5660-5535 calBCE (6680±30 BP, Beta-392490), with a whole ceramic pot of the Early Neolithic Elshanka type, dated at other sites between 6200-5000 BCE. His Y-haplogroup R1b R-P297 showed the basal mutation at the root of the Yamnaya R1b Y-haplogroup subclade. His mt-haplogroup, U5a1d, was among the most common steppe mt-haplogroups.

A millennium later in the same area another male (I20116) showed the same R1b R-P297 in grave 97 at Ekaterinovka Mys dated ca. 4700-4500 BCE. After that this Y-lineage disappeared from sampled populations until reappearing in Yamnaya, but we can observe that its early distribution included the northern end of the Volga Cline.

The Eneolithic settlement at Lebyazhinka-4 produced a polished stone mace head with a zoomorphic eared shape compared to a horse head.

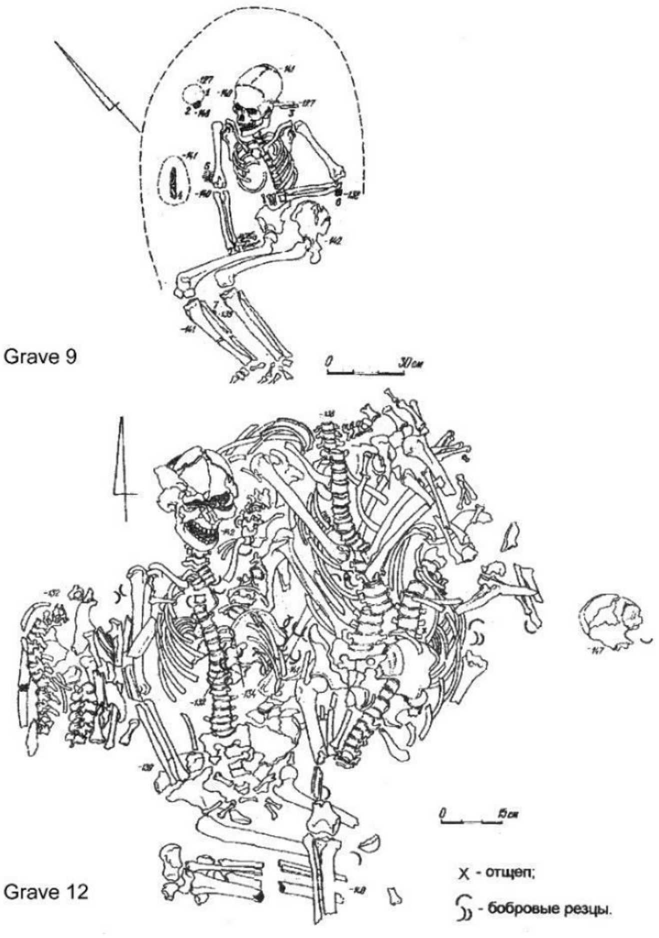

**Fig. 2.16. Burials 9 and 12 at Lebyazhinka-5.**

At **Lebyazhinka-5** several grave pits were partly disturbed by a LBA Srubnaya settlement. This project sampled graves 8, 9, and 12 at Lebyazhinka-5.

Graves 8 (I6907, a female) and 9 (I6906, female) were thought to be Eneolithic, largely because the pose was supine with raised knees like Khvalynsk, and the area near Lebyazhinka farm witnessed intense Eneolithic activity. But both graves were dated by radiocarbon to the Yamnaya period between 3336-2907 calBCE. It is interesting that they seem to have lacked kurgans, making them rare examples of Yamnaya-era graves with Yamnaya genetics but without kurgans.

Grave 12 at Lebyazhinka V was Eneolithic and contained parts of five individuals: two adult females, two adult males, and a juvenile. The most complete was skeleton #2 (I6908), an adult female posed supine with tightly raised knees, probably resulting from the lower legs being bound against the thighs with the feet flat on the ground. This distinctive burial pose was not found in Siberia, Kazakhstan, or the northern Russian/Baltic forests, where an extended position on the back was typical.

A radiocarbon sample from a terrestrial animal, a marmot incisor ornament (one of many found in clusters throughout the deposit) was dated 4838-4612 calBCE (5865±40 BP, GrA-64051) by Shishlina et al (2017). The date is applied to the entire grave based on the intermixed nature of the bones. An articulated human knee (proximal tibia/fibula and distal femur) from another individual was found overlying (see image) the raised knees of individual #2. If the 4800-4600 BCE date applies to her, then she is among the earliest dated individuals buried in this distinctive pose, which later (4500-3800 BCE) was typical across the steppes: in the Volga Cline (Khvalynsk), Serednii Stih (Krivyanskiy-9), and in the Berezhnovka-Progress-2 graves. It is also known as ‘the Yamnaya position’ because of its frequency in graves contemporary with 8 and 9.

Individual #2 in Grave 12 was almost complete and in anatomical order. She was placed over a grave pit that contained four partial individuals, two adult men (#3 & #5), one adult woman (#1), and an adolescent (#4). All heads were missing with most lower limbs, but partly articulated torsos included vertebrae and pelvises. The loss of heads and most limbs could have resulted from exposure before burial. They were buried in two compact layers with #4 and #5 at the bottom. Similar deposits of individuals overlaid in compact layers were continued at Khvalynsk, with whole and partial remains. The dated marmot tooth was located near #4 and 5.

Grave 12 at Lebyazhinka V was contemporary with Ekaterinovka Mys where 90% of the graves were posed supine with legs extended but a few were supine with raised knees like #2; and with S’yezzh’e (all supine extended). This distinctive burial position began to occur here and in a few graves at Ekaterinovka Mys before 4500 BCE in sites lacking copper artifacts, arguably dated before Balkan copper spread to the Volga.

| Sample ID | Burial code | Radiocarbon date | Y-group | mtDNA |
|---|---|---|---|---|
| I0124 | Leb. IV, grave 1 Samara H-G | 5660-5535 calBCE (6680±30 BP/ Beta-392490) | R1b-<br>R-P297 | U5a1d |
| I6908 | Leb. V, grave 12 | 4838-4612 calBCE (5865±40 BP/ GrA-64051) | (female) | U4a |
| I6906 | Leb. V, grave 9 | 3336-3033 calBCE (4475±20 BP/<br>PSUAMS-4257) | female | U4a |
| I6907 | Leb. V, grave 8 | 3022-2907 calBCE (4355±20 BP/ PSUAMS-4258) | female | H13a1a2 |
| marmot | Leb. V, grave 12 | 4838-4612 calBCE (5865±40 BP/ GrA-64051) | Terrestrial mammal | |

*Korolev, A., M. Kulkova, V. Platonov, N. Roslyakova, A. Shalapinin and Y.E. Yanish. 2018. Archaeological Materials of Eneolithic Settlements in Forest-Steppe Zone of the Volga Region: A Source for Diet and Chronology. Radiocarbon 60 Special Issue 5(2): pp. 1587 – 1596. DOI:*

*https://doi.org/10.1017/RDC.2018.114*

*Shishlina, N.I., J. van der Plicht, & M.A. Turetsky 2018. The Lebyazhinka burial ground (middle Volga region, Russia): new 14C dates and the reservoir effect. Radiocarbon 60 Special Issue 5(2): 681–690. DOI:10.1017/RDC.2017.94*

#### 2.9 Maximovka

*Summary by D. Anthony*

Flint microliths typical of Mesolithic & Neolithic hunter-fishers were found at Maximovka with Early Neolithic pottery, but the graves inserted into the site were dated later, 4153-3633 BCE. Together with a similar EHG-rich individual found at Chekalino-4 (3600-3500 BC), also inserted into a Neolithic settlement site on the Sok River not far north of Samara (I6303), these graves perhaps indicate a southward shift of a Murzikha-type population into the Samara valley in the early 4th millennium BCE. The Yamnaya population replaced this EHG-rich population in the Samara valley in the earliest phase of the Yamnaya culture, beginning 3300-3100 BCE.

The Maximovka graves include a pair of 1st-degree relatives, either father-son or brothers, and their identical mt-DNA haplotypes suggest brothers. Their radiocarbon dates are offset for an unknown reason; perhaps one brother had a diet with more fish (?) and his dates reflect a slight FRE.

| Sample ID | Burial code | Radiocarbon date | Y-group | mtDNA | Relatives |
|---|---|---|---|---|---|
| I6904 | Maximovka | 4153-3962 calBCE (5205±25 BP, PSUAMS-4255) | I2a-I-P37 | U5a1d2b | 1st-degree relative of I8742 |
| I8742 | Maximovka S.1 B.1 K.1 | 3710-3633 calBCE (4885±25 BP, PSUAMS-8284) | I2a-I-P37 | U5a1d2b | 1st-degree relative of I6904 |
| I8446 | Maximovka S.1 B.1 K.2 | 3946-3711 calBCE (5025±25 BP, PSUAMS-5923) | R1a-R-YP4141 | U4a1 | |

#### 2.10 Progress-2 and Vonyuchka

*Summary by D. Anthony*

Progress-2 was excavated 2009-2010 under the direction of S.Y. Berzina. The multi-period kurgan cemetery was located on the left bank of the Malka River, itself a left tributary of the Terek River in the central Caucasus steppes east of Piatigorsk. The central Caucasus steppes consist of high grass-covered ridges that form a watershed, cut by streams flowing northeast into the Caspian (through the Terek River), north into the Manych Depression, or northwest into the Sea of Azov/Black Sea (through the Kuban), with the glaciated peaks of the Caucasus visible 100km to the south. Progress-2 is one of many Eneolithic grave sites including Vonyuchka concentrated in the central Caucasus steppes, the upper Terek tributaries, and the Manych Depression. Very few Eneolithic graves are found in the NW Caucasus Kuban drainage.

The oldest and largest of these Eneolithic cemeteries was excavated in 1929-30 in the city of Nalchik and yielded one date of 4840–4820 BCE (GrA-24442, 5910 ± 45 BP). Nalchik has not been sampled for aDNA. Nalchik differed from other Eneolithic sites in its size (121 burials, while the later Eneolithic cemeteries such as Progress-2 usually have 2-4 individuals) and in the fact that 75% of the Nalchik burials were posed contracted on the left (mainly females) or right (mainly males) sides, while Progress-2 and almost all later Eneolithic graves were posed supine with raised knees, like most Khvalynsk and Serednii Stih graves (although contracted-on-the-side graves continued at Khvalynsk as a small minority). Burials contracted on one side were later also typical of the Maikop culture.

About 25% of the graves at Nalchik in which a specific pose was clear were posed supine with raised knees. They can be interpreted as later graves in a multi-component cemetery, showing a shift in funeral ritual, a hypothesis supported by the discovery in one of the supine-with-raised knees graves (#83) of the only copper artifact found at Nalchik, a ring, and a serpentine stone bracelet like one found at Khvalynsk, dated 4500-4300 BCE there. Or the raised-knee graves at Nalchik could be older than Khvalynsk, contemporary with the raised-knee graves at Lebyazhinka V:12 and Ekaterinovka Mys in the Samara region, dated 4700-4500 BCE. More radiocarbon dates are needed to identify the oldest raised-knee graves, but Nalchik shows that the raised-knee posture was an Eneolithic innovation that replaced an older ritual (contracted on the side) in the North Caucasus.

Progress-2, Vonyuchka, and the other late Eneolithic graves in the central Caucasus steppes were in either simple earthen pits or in niches (“catacombs”) dug into one wall of a pit, with the grave floor intensely stained by red ochre. The shallow grave pit was covered by a small mound, perhaps 10-15m in diameter and 1 to 1.5m high. These were among the oldest kurgans in the steppes, dated firmly to the late 5th millennium BCE. Small kurgans were built also over Serednii Stih pit graves in the lower Don-Azov steppes (Novodanilovka) and in the steppes bordering the northern Danube delta (Suvorovo) at this time, but not over Volga Cline graves on the Volga. Small kurgans were a regional funeral practice in the Eneolithic.

The image shows PG 2004: Progress-2, kurgan 4, grave 9. PG2003 was grave 12. 9 & 12 were contemporary. 12 shown on right with intense red ochre.

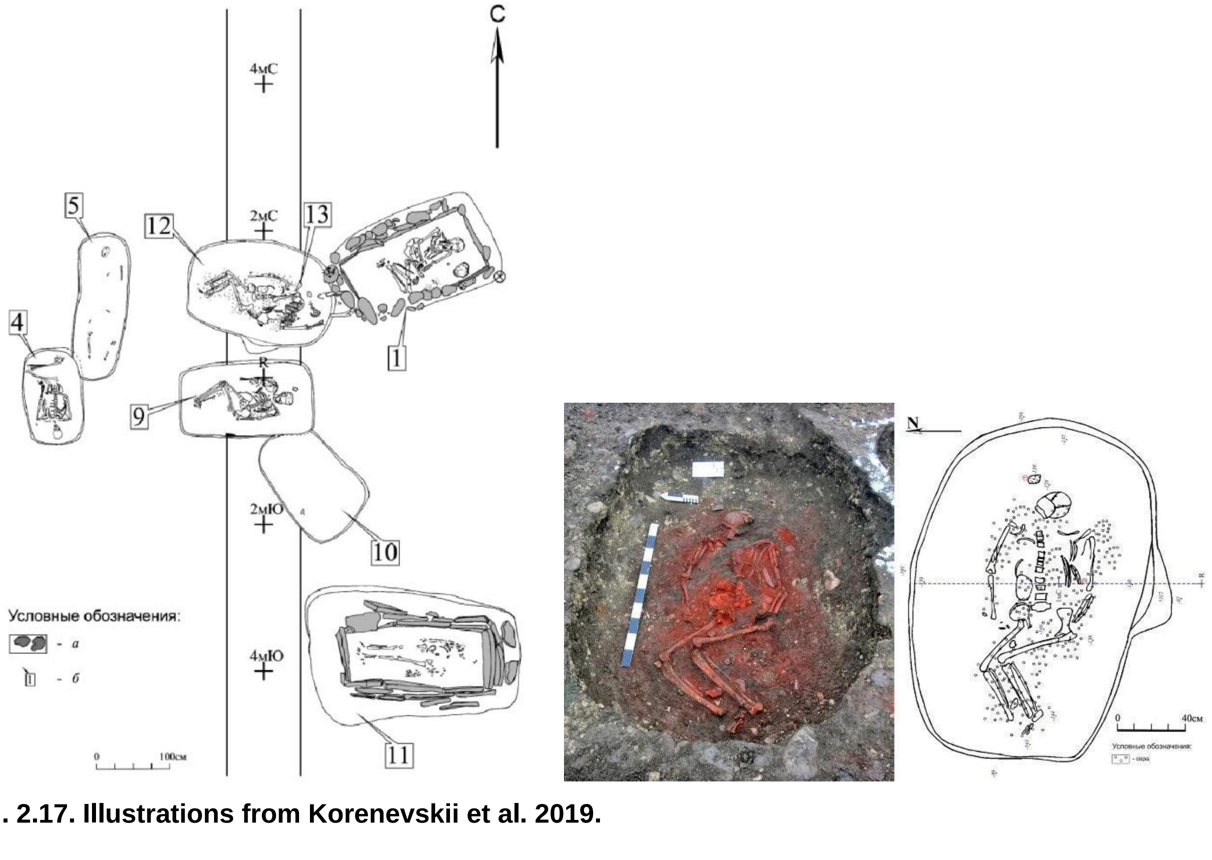

**Fig. 2.17. Illustrations from Korenevskii et al. 2019.**

At Vonjucka-1 (VJ 1001) kurgan 1 covered an Eneolithic grave 8, discovered in 2010 in the eastern suburbs of Pyatigorsk. Local archaeologists named it after an adjacent slow stream called ‘Stinker’. Archaeologists who published with Wang et al. (2019) later tried to change the site name to Konstantinovskii-1, after a town northeast of the site. But the whole genome had already been published under the name Vonjucka-1 (Wang et al. (2019: 3).

The female in k.1 grave 8 was positioned supine with raised knees on a floor intensely colored with red ochre in a chamber dug into the side of the grave shaft, or a ‘catacomb’. The grave contained a small undecorated ceramic pot, a flint blade and scraper, and a pestle.

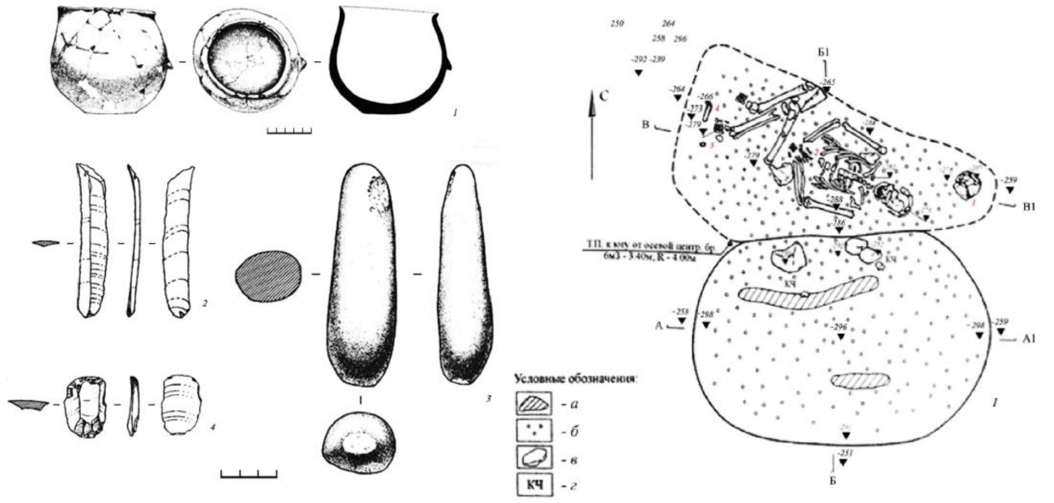

**Fig. 2.18. Illustrations from Korenevskii et al. 2019.**

| Sample ID | Burial code | Radiocarbon date | Y-group | mtDNA |
|---|---|---|---|---|
| VJ1001 | Vonjucka-1 kurgan 1, grave 8 | 4337-4177 calBCE (5409±24 BP, MAMS-29823) | (F) | T2a1b |
| PG2001 | Progress-2 kurgan 1, grave 37 | 4994-4802 BCE (6012±28 BP, MAMS-110564) on human bone // MAMS-13010 on charcoal in grave 5397 +- 28 BP, 4338-4074 BCE | R1b1a2–R-V1636 | I3a |
| PG2004 | Progress-2 kurgan 4, grave 9 | 4240-4047 calBCE (5304±25 BP, MAMS-11210) | R1b1a2-R-V1636 | H2 |

*Korenevskii, S.N., N. Y. Berezina, Y. B. Berezin, U. G. Gresky 2019. Novy pogrebeniya protoYamnoi kul’tury na Stavropol’e (new burials of the protoYamna culture in Stavropol). In: Kh. A. Amirkhanov (ed.), Gory Kavkaza i Mesopotamskaya Step’na Zare Bronzovogo Veka. Institut Arkheologii RAN (Moskva 2019) 155–178. (source of illustrations)*

*Korenevskii, S. N. 2012. Rozhdenie Kurgana: Pogrebal’nye pamyaniki Eneoliticheskogo vremeni PreKavkaz’ya I Volgo-Donskogo mezhdurech’ya (The birth of kurgans: grave rituals of the Eneolithic period in the Caucasus piedmont and between the Volga and Don Rivers). Moskva: Institut Arkheologii RAN.*

#### 2.11 Remontnoye Late Steppe Eneolithic

*Summary by D. Anthony & N. Shishlina*

Two Eneolithic graves dated between 4152-3637 BCE (I28682, I28683) were discovered near the modern village Remontnoye on the watershed between the Black and Caspian Seas north of the Manych Depression. The Manych Depression was formed as a spillway that flowed from the Caspian into the Black Sea when glacial meltwater filled the North Caspian Depression between about 11,000 BC and 9,000 BC. Until about 9,000 BC the Manych held a river 600km long flowing west into what is now the lower Don and Sea of Azov. After 9,000 BC the Manych river gradually became a series of winding lakes containing phragmites and sedge marshes important for animal fodder.

Graves from the steppe Eneolithic are usually were placed under small mounds. These mounds, usually solitary, were the earliest burials in the steppes north of the North Caucasus, and were distributed from the lower Kuban to the Caspian Depression. Yamnaya culture burials are either incorporated into the mounds of the Eneolithic period or serve as primary mounds themselves or are incorporated into mounds from the Yamnaya period. Yamnaya mounds form separate clusters within steppe burial grounds, ranging from 2-3 to 10-12 mounds.

Steppe Eneolithic burials were placed in both simple pits and pits with a side chamber (catacombs). Bodies were usually arranged supine with raised knees, like Khvalynsk and Serednii Stih, or sometimes in a contracted position on their sides. Inventory items typically include saiga astragali, bone rods, and pottery vessels similar to Serednii Stih.

Eneolithic camps and graves were found at Sukhaya Termista I, Ulan IV, and Peschany V on the north side of the Manych. Poorly preserved Peschany V (I28681) gave a radiocarbon date older than the other two (4329-4046 calBCE /5340±50 BP/ Gr-54939), but only Sukhaya Termista and Ulan IV provided sufficient data for DNA analysis.

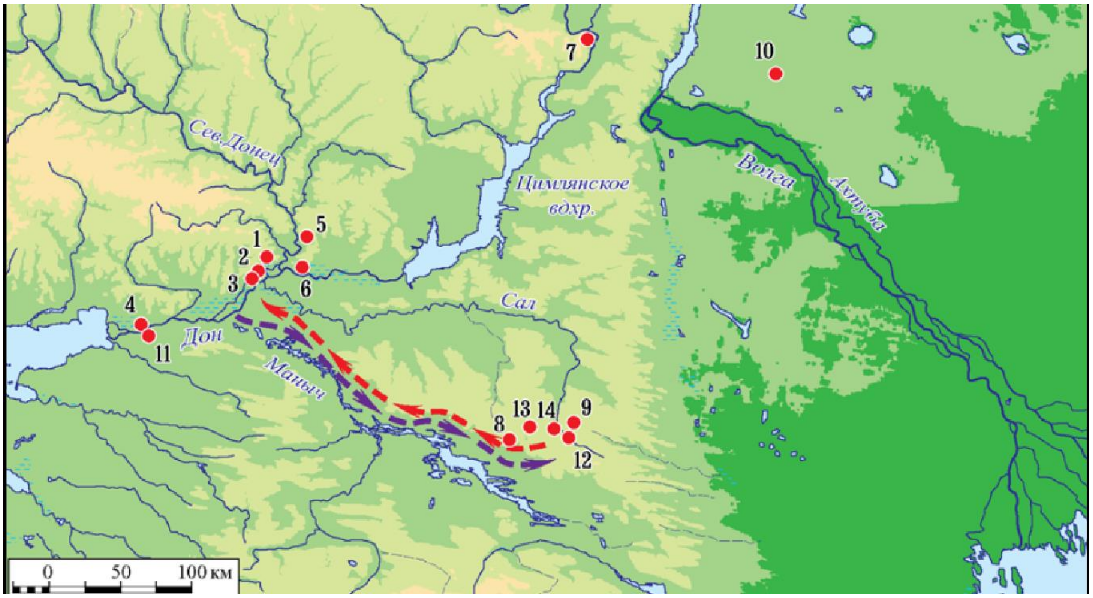

**Fig. 2.19. The Remontnoye sites are: 12, Peschany-5; 13, Ulan-4; and 14, Sukhaya Termista-1.** Shishlina postulated that the same population moved seasonally from camp sites on the lower Don. Krivyansky-9 was near #2 & 3. From Shishlina et al. 2018.

The Eneolithic graves around Remontnoye were interpreted by Shishlina as representing a population that moved seasonally with its cattle and sheep between the Manych steppes (summer) and the lower Don (winter). Seasonal (summer) settlements from the Eneolithic and Yamnaya culture have been discovered along the Sal and Manych river valleys. Eneolithic radiocarbon dates in the late 5$^{th}$/early 4$^{th}$ millennium BCE indicate a period when domesticated animals were used as a ritual currency in funeral sacrifices at Khvalynsk but were milked only rarely, according to studies of dairy peptides in human dental calculus.

##### *Genetics*

A Remontnoe male at Ulan IV k3:12 had a V-1636 Y-haplogroup like the Volga Cline males, but otherwise both he and the female at Sukhaya Termista I were enriched in Aknashen-related CHG ancestry. They occupy a PCA position between Berzhnovka-Progress-2, which did not attest Aknashen-related ancestry, and a Caucasus population like Neolithic Aknashen in Armenia, the latter perhaps acquired through mate exchanges with the Meshoko-culture farmers who occupied the Kuban region 4600-3800 BCE but are unsampled because they did not use cemeteries (for Meshoko see map in Golubaya Krinitsa section). One female at Sukhaya Termista had mt-haplogroup H2a, also found in the Caucasus at Areni-1 and in Maikop.

Steppe Maikop people occupied this same region, 200km north of the Caucasus peaks, about 3600-3300 BCE, showing more West Siberian and Maikop ancestry than the Remontnoye population.

| Sample ID | Burial code | Radiocarbon date | Y-group | mtDNA |
|---|---|---|---|---|
| I28682 | Sukhaya Termista I k1:14 | 3766-3637 calBCE (4903±30 BP, OxA-29798) | 18-25 F | mt/H2a |
| I28683 | Ulan IV k3:12 | 4152-3804 calBCE (5160±40 BP, Gr49047) | 40-45 M R1b R-V1636 | mt/R0 |
| I28681 | Peschany V | 4329-4046 calBCE (5340±50 BP, Gr-54939) | juven M | Too few SNP's |

##### *2.11.1.Kurgan 1: grave 4 (ID I28682): 3766-3637 calBCE*

In 2009, Natalia Shishlina excavated a Steppe Eneolithic burial under kurgan 1 at the Sukhaya Termista-1 site. The burial was unearthed in the center of a multi-phase kurgan (see the description in SI 5.3.1).

The burial was made on the ancient ground surface and covered by a small poorly preserved mound.

A shallow oval-shaped pit was oriented in east-west direction, with a slight deviation to the north. Its dimensions are 190 by 107 centimeters, with wall heights ranging from 30 to 40 centimeters. Within the pit lay the skeleton of a female individual aged 18-25 y.o. was positioned gently contracted on her side, with the skull oriented west. The cranial bones are coated with ochre.

A bronze (or possibly silver) temple ring, measuring 2.2 by 1.7 centimeters and completing 1.5 rotations, was found beneath the right temporal bone. Additionally, a sheep vertebra was located 40 centimeters south of the skull.

A.A. Kazarnitskiy characterizes the cranial features as an exceptionally elongated and narrow dolichocephalic cranial vault (with damage to the basilar part), a wide forehead, and a face that is notably tall with average width, relatively narrow, and orthognathic. The horizontal profiling of the face is moderate at the upper level and highly pronounced at the zygomatic points. The orbits are notably wide and elevated, while the nose is very tall, narrow, and sharply projecting, with a high nasal bridge. According to the formula developed by M. Trotter and G. Gleser, the body length measures 154 cm.

The radiocarbon dating of the burial yielded an age of 3766-3637 calBCE (4903±30 BP, OxA-29798).

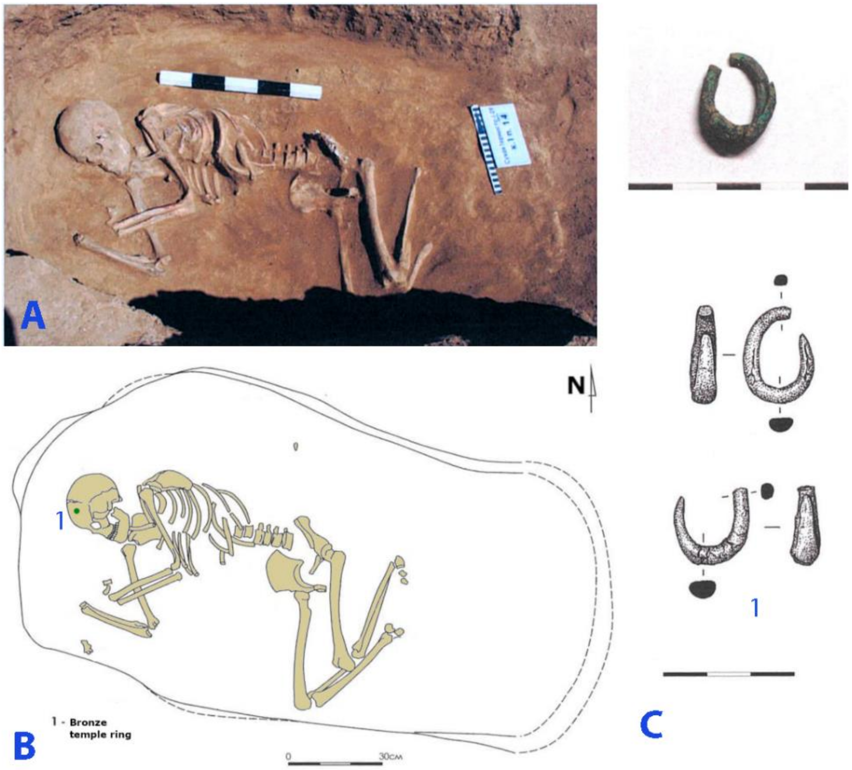

**Fig. 2.20. Sukhaya Termista-1, kurgan 1, burial 14.** A, B – plan and photo of the burial, C – grave goods: bronze temple ring (image by Natalia Shishlina).

##### *2.11.2 Ulan-4, kurgan 3, burial 12 (ID I28683): 4152-3804 calBCE*

In 2010, Natalia Shishlina excavated another Steppe Eneolithic burial 12 under kurgan 3 at the Ulan-4 site. It was located in the northwest sector of the kurgan and was the oldest grave in the burial site, primary for the stratigraphically first mound constructed at this site. Burial 12 was partially disrupted by a later grave 13 attributed to the Early Catacomb culture.

The grave pit is a shallow, oval-shaped feature, with an approximate depth of 40 cm from the ancient ground surface. The pit is oriented along a northeast-southwest axis, with a length along the northeast-southwest axis of 174 cm and a width along the northwest-southeast axis estimated to be around 110 cm.

At the bottom of the grave, at a depth of -335 cm from the top of the kurgan, the skeleton of a male aged 45-55 years was found positioned on his back, the skull oriented to the southwest. The original posture of the legs is difficult to determine because of the later disturbance of grave 13. The skull was found in situ likely on a raised platform, slightly tilting leftward, with the facial bones facing northeastward. In the left frontal bone near the coronal suture, there was evidence of a pre-mortem open cranial injury, completely healed. All skeleton bones were heavily coated with red ochre. Fragments of organic bedding, with approximate dimensions of 140 × 60 cm, were preserved around the skeleton. No artifacts were found in the burial.

The skeleton was C14-dated to 4152-3804 calBCE (5160±40 BP, Gr49047).

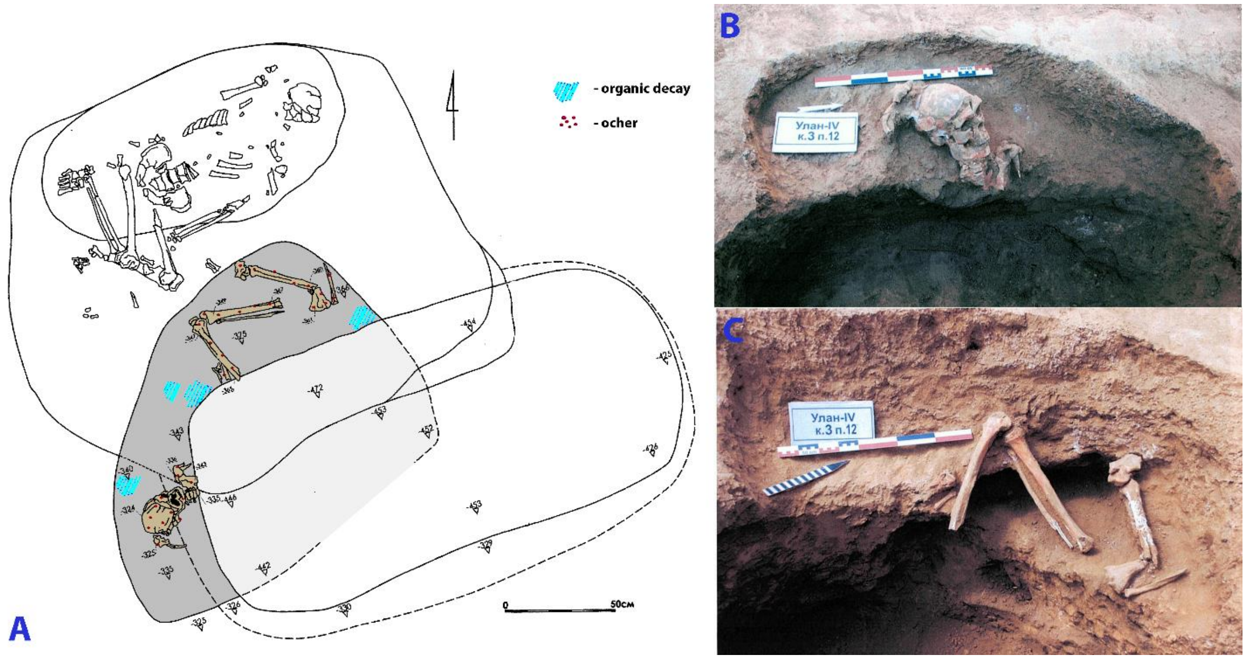

**Fig. 2.21. Ulan-4, kurgan 3, burial 12.** A – plan of burial 12, B-C – photos of the western and the northern parts of the burial (image by Natalia Shishlina).

#### 2.12 Syezzheye

*Summary by D. Anthony*

Syezzheye (pronounced SYEHZ-ZHYE-YEH) is an Early Eneolithic cemetery near the south bank of the Samara River, which flows into the Volga from the southwestern Urals and marks the northern steppe border. Nine Eneolithic graves held single individuals in extended supine position, heads to the east/northeast, stained with red ochre. Radiocarbon dates on human bone are too old because of FRE (Freshwater Reservoir Effects).

Comb-impressed, collared, shell-tempered pottery representing 27 vessels was found in the graves and the ochre-stained sacrificial surface above the graves, the type collection for the Eneolithic Samara culture (see image). Since similar Samara-culture pottery was found also in the graves at Ekaterinovka Mys with a similar burial rite, the radiocarbon dates on terrestrial animal bone from that site, 4800-4500 BC, probably broadly indicate the age of Syezzheye.

Copper did not occur at Syezzheye or Ekaterinovka Mys, probably dated before Balkan copper appeared on the Volga after 4500 BCE. Graves 6 and 7 at Syezzheye were richly decorated with shell and bone beads and boar’s tusk plaques like those at Yasinuvatka on the Dnieper and Mariupol on the Azov coast, also dated before 4500 BCE. Concentrated in these two graves but also in the ochre-stained sacrificial deposits above the graves, were found, in addition to the pottery, four polished stone adzes 10-12cm long, six short (ca.5cm) polished stone axes, one retouched flint blade 12 cm long, a bone harpoon, bone plaques carved in the shapes of bulls, ducks, and horses, and the head-and-hoof remains of two horses.

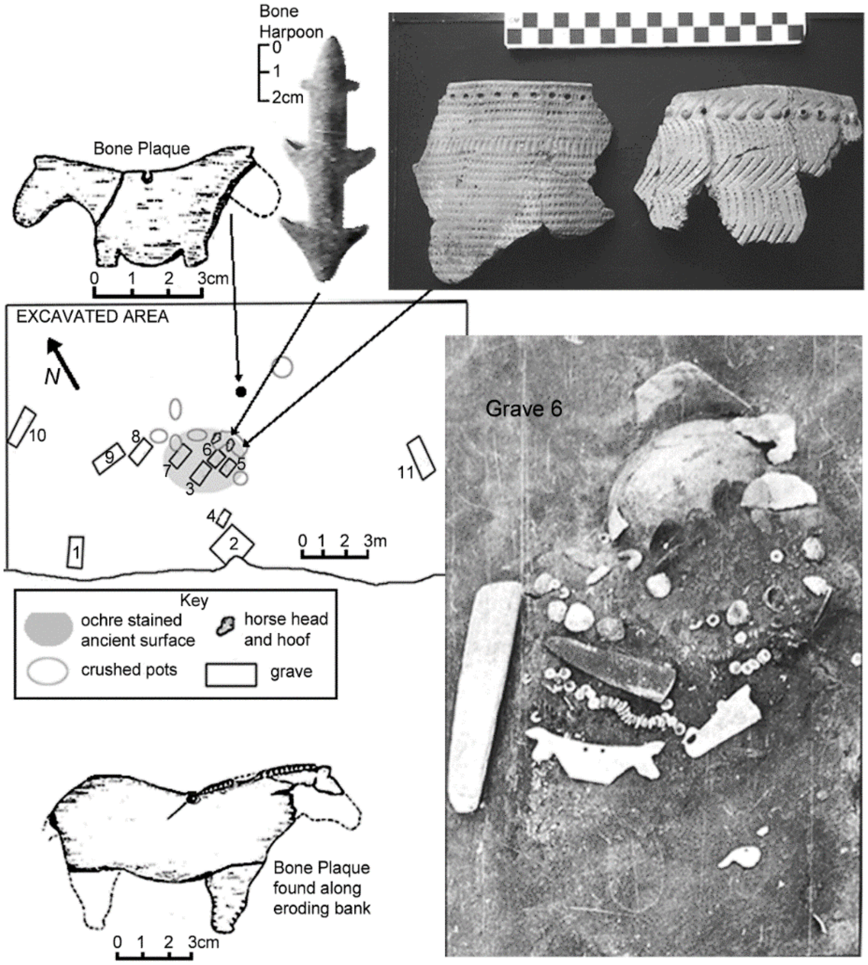

**Fig. 2.22. Selected artifacts from Syezzheye.**

On a PCA the Syezzheye individuals show affinity to Ekaterinovka Mys. Both were admixed with a CHG-related ancestry the source of which is not identified, but it was distinct from the Aknashen-related farmers of the Meshoko culture in the NW Caucasus 4600-3800 BCE. It is possible that the unsampled Seroglazovka foragers (with ceramics) who appeared at sites such as Kair-Shak-3 and Dzhangar on the lower Volga after about 6200 BCE (Vybornov et al. 2018) introduced this Mesolithic-like CHG-related ancestry into the steppes, accounting for its presence in a small amount (ca. 7%) even in the Samara Hunter-Gatherer from Lebyazhinka-4 at 5500 BCE. The flow of this archaic CHG-related ancestry into the Volga Cline increased at Ekaterinovka Mys and after 4500 BCE at Khvalynsk and Khlopkov Bugor, possibly the result of increased movement up and down the Volga encouraging wide-ranging mate exchanges with the CHG-rich Berezhnovka-Progress-2 population.

A population with northern EHG-dominant genetics appeared less than 5km from Syezzheye at Maksimovka (I6904, I8742, I8446), dated 4100-3700 BCE, centuries after Syezzheye. These graves perhaps indicate a southward shift of a Murzikha-type population also seen at Chekalino-4 above, into the Samara valley in the early 4th millennium BCE, perhaps pushing the CHG-admixed Syezzheye population south into the steppes.

| Sample ID | Burial code | Syezzheye radiocarbon dates | Y chromosome | mtDNA |
|---|---|---|---|---|
| I22203 | grave 2, skeleton 2 | 4881-4720 calBCE (5925±25 BP, PSUAMS-8842) | R1b-R-V1274 | U5a1d |
| I22204 | grave 2, skeleton 3 | 5600-4700 BCE | R1b-R-V1636 | R1b1 |
| I22205 | grave 3 | 5557-5381 calBCE (6520±30 BP, PSUAMS-8843) | female | U2e1 |

*Vasiliev, I.B., and G.I. Matveeva, 1979. Mogil'nik u s. S'yezhee na R. Samare. Sovietskaya Arkheologiia (4): 147-166.*

*Vybornov, A. A.,M. Kulkova, P. Kosintsev, V. Platonov, S. Platonova, B. Philippsen, and L. Nesterova. 2018. Diet and chronology of the Neolithic-Eneolithic cultures (from 6500 to 4700 cal BC) in the lower Volga basin. Radiocarbon 60(5): 1–14.*

*Anthony, David W. 2007. The horse, the wheel, and language: How Bronze Age riders from the Eurasian Steppes shaped the modern world. Princeton: Princeton University Press, pp. 189-192.*

#### 3. ENEOLITHIC OF THE ALTAI AND MIDDLE YENISEI: AFANASIEVO CULTURE SITES

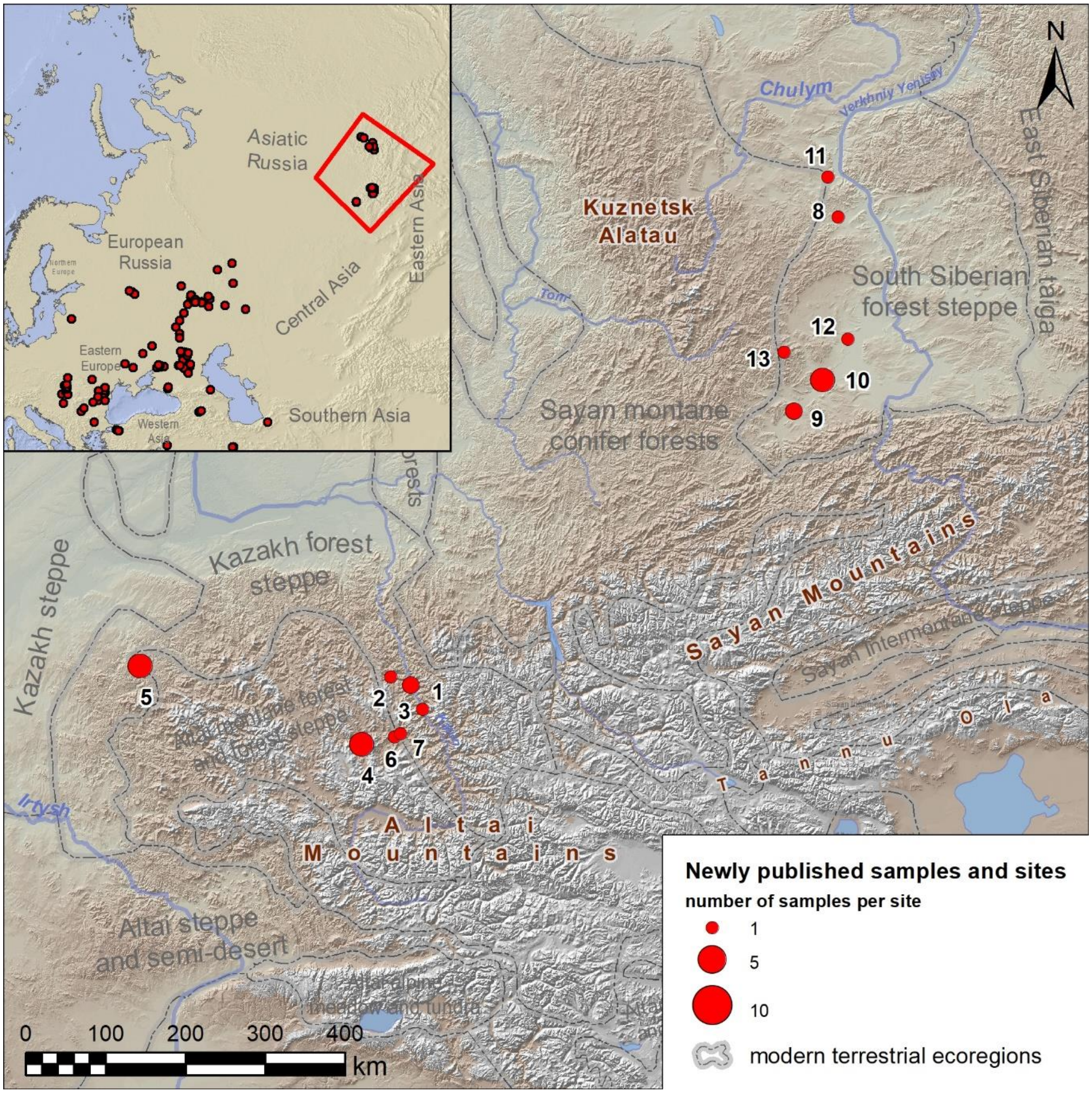

**Fig. 3.1. Newly published samples from the Eneolithic and Bronze Age sites in the Altai region and the Minusinsk Basin, Russia.** 1 - Ayrydash-1 (Altai Republic, Chemalsky District, Kuyus Village); 2 - Bolshoy-Tolgoyok (Altai Republic, Shebalinsky District); 3 - Choburak-1 (Altai Republic, Chemalsky District, Elanda Village); 4 - Elo-1, 2 and Nizhny-Tyumechin-1 (Altai Republic, Ongudaysky District, Elo Village); Inskoy-Dol and Khankarinsky-Dol (Altai Krai, Krasnoshchyokovsky District); 6 - Karakol-2 (Altai Republic, Ongudaysky District); 7 - Kurota-2 (Altai Republic, Ongudaysky District); 8 - Afanasieva-Gora (Bateni) (Republic of Khakassia, Bogradsky District, Pervomayskoye Village); 9 - Beltyry (Republic of Khakassia, Askizsky District); 10 - Kamyshta-1 and 2 (Republic of Khakassia, Askizsky District, Katanov Village); 11 - Karasuk-3 (Republic of Khakassia, Bogradsky District, Pervomayskoye Village); 12 - Okunev-Ulus (Republic of Khakassia, Ust-Abakansky District); 13 - Uzunchul (Republic of Khakassia, Askizsky District).

#### 3.1 Choburak-1 site

*Summary by A. Tishkin*

The archaeological site Choburak-1 is located on the right bank of the Katun' River, a bit south from the Elanda village, in the Chemalsky district of the Altai Republic, Russia. It includes more than 60 features dated to various chronological periods from the Eneolithic to the Early Middle Ages.

##### *3.1.1 Choburak-1, kurgan 36*

Kurgan 36 was explored by the Chemalsk archaeological expedition of Altai State University, in the eastern part of the Choburak-1. The kurgan was a rounded structure made of medium and small stones. The mound was 8.8 m in diameter and up to 0.3 m high. In the center of the burial structure, the stones were almost completely absent, which is characteristic of the burial mounds of the Afanasievo culture.

###### *3.1.1.1 Burial 36:2 (Individual ID I25159)*

Under the central part of the mound was a grave measuring 2.36x1.26 m and up to 0.97 m deep. The skeleton was placed in the supine position, with his legs bent at the knees and his head oriented to the east. The skeleton was poorly preserved, the thoracic part and ribs were destroyed by rodents, whose burrows were in the filling of the grave pit. A small hole in the skull has been found. The skeleton and the bottom of the grave were covered with traces of ochre. The individual was a female, mt-haplogroup W3a1.

The grave goods are represented by an ornamented pointed-base pottery vessel with a clearly visible crust on the inner surface of the walls. The grave is attributed to the Afanasievo culture of the Eneolithic.

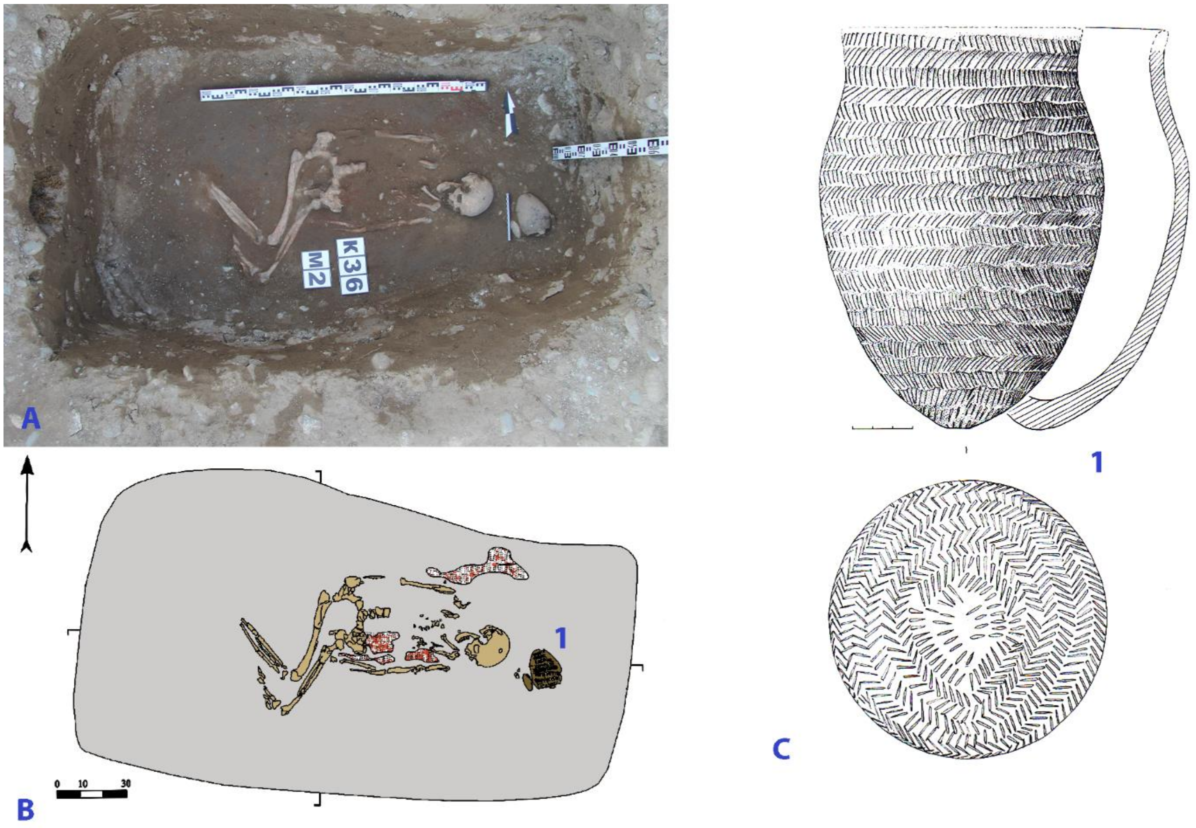

**Fig. 3.2. Choburak-1, kurgan 36, burial 2. A and B – photo and plan of the burial, C – pottery vessel.**

### 4. EARLY BRONZE AGE OF THE VOLGA-URAL STEPPE

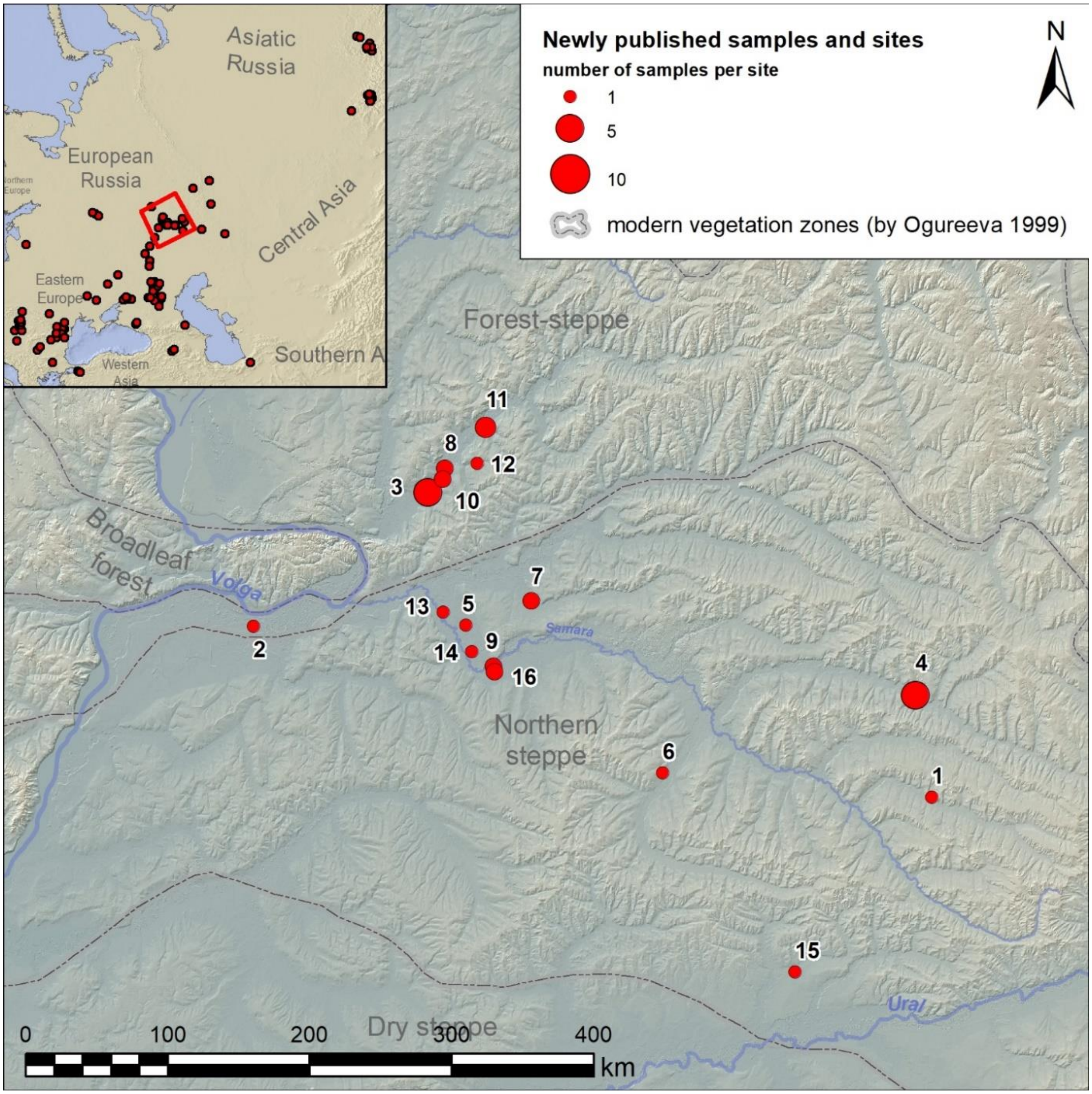

**Fig. 4.1. Newly published samples from the Bronze age sites in the Middle Volga and the Volga-Ural regions, Russia.** 1 - Bogolubovka (Orenburg Oblast, Novosergiyevsky District); 2 – Ekaterinovka Mys (Samara Oblast, Bezenchuksky District); 3 - Grachevka-1 and 2 (Samara Oblast, Krasnoyarsky District); 4 - Krasikovo (Orenburg Oblast, Krasnogvardeysky District); 5 - Krasnosamarskoe-4 (Samara Oblast, Kinel District); 6 - Kurmanaevka-3 (Orenburg Oblast, Kurmanayevsky District); 7 - Kutuluk-1 (Samara Oblast, Kinel District); 8 – Lebyazhinka-5 (Samara Oblast, Krasnoyarsky District); 9 - Leschevo-1 and 3 (Samara Oblast, Neftegorsky District); 10 - Lopatino-1 and 2 (Samara Oblast, Krasnoyarsky District); 11 - Nizhnaya-Orlyanka-1 (Samara Oblast, Sergiyevsky District); 12 - Orlovka (Samara Oblast, Sergiyevsky District); 13 - Podlesny-1 (Samara Oblast, Kinel District); 14 - Poplavskoe-1 (Samara Oblast, Kinel District); 15 - Shumayevo-1 (Orenburg Oblast, Tashlinsky District); 16 - Utyovka-11 (Samara Oblast, Neftegorsky District).

#### 4.1 Bogolubovka site (Volga-Ural steppe)

*Summary by P. Kuznetsov*

The site is situated on the edge of the first river terrace on the Bolshoy Uran River's left bank (the Samara River's right tributary). It is part of a group of five kurgan burial sites and three settlements dated to the Bronze Age, which stretches the left bank of the river for 30 km. This area also has six left tributaries of the Bolshoy Uran River. The steppe landscape, network of tributaries, and relatively low left bank contributed to the productive development of animal husbandry during the Bronze Age.

##### *4.1.1 Bogolubovka, kurgan 11*

The diameter of the burial mound is 26 meters, with a height of 0.67 meters.

*Burial 11:6 (individual ID I11831): 3074-2904 calBCE*

Burial 6 is located under the center of the mound. The pit is rectangular in shape, measuring 220x135 cm, with a depth of up to -165 cm from the modern surface of the mound's center. The burial was disrupted by a burrowing rodent. The skull, ribs, and bones of the left leg are preserved. All bones are stained with ochre, and organic bedding is identified at the bottom of the pit. The individual was a male aged 17-25, Y-haplogroup R1b (R-KMS67), mt-haplogroup U4a1e.

The skull is anthropologically classified as mesomorphic, Europoid. The burial is attributed to the late stage of the Yamnaya culture of the Urals based on the orientation of the burial pit and the presumed position of the deceased (Morgunova, ed., 2014).

The burial is dated to 3074-2904 calBCE (4355±25 BP, PSUAMS-10736).

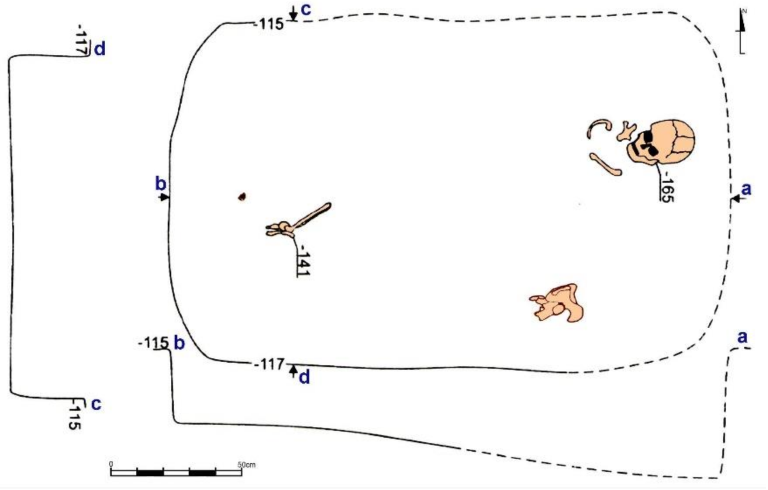

**Fig. 4.2. Bogolubovka, kurgan 11, burial 6** (image by Pavel Kuznetsov)**.**

#### 4.2 Grachevka-1 site (Middle Volga forest-steppe)

*Summary by P. Kuznetsov*

The site is located approximately 15 km from the mouth of the Sok River, a right tributary of the Volga River. It is positioned on the edge of the floodplain terrace on the left bank of the Sok River, where multiple clusters of kurgans are dispersed along a 40 km stretch, spaced roughly 3-5 km apart from each other.

###### *4.2.1 Grachevka-1, kurgan 2*

The diameter of the mound is 18 m, height 0.3 m.

###### *4.2.1.1 Burial 2:1 (individual ID I7479): 2899-2703 calBCE*

Burial 1 is the only one found in the kurgan, it is located 4.3 m to the west of the centre of the mound. The dimensions of the pit are 1.52×1.34 m. The profile of the burial pit is shallow, with steep walls, and a depth of -152 cm from the modern top of the mound. The skeleton is positioned on their back with bent legs and outstretched arms. The head is oriented to the east-northeast. Near the eastern wall of the pit, a clay vessel was discovered, inverted with its bottom up.

The buried individual is a female child, aged 7-8 years. According to Aleksandr Khokhlov's determination, the skull of the deceased is Europoid. Her mt-haplogroup is H13a1a1.

Between femur and tibia of the deceased's left leg, bones of two limbs of a sheep or a goat were found. Another sheep or goat long bone was uncovered under the elbow of the left arm of the skeleton. This is an unusual type of sacrificial offering.

Between the right arm and the knee of the right leg of the skeleton, two pairs of thin-walled hollow bones, potentially avian, with neatly trimmed ends, were discovered. The length of the bones in one pair measured 8.3 cm, while in the second pair, it was 6.1 cm. The diameter of the bones ranged from 1.5 to 2.0 cm. These findings could possibly be attributed to parts of a Pan flute (Kuznetsov et al., 2018).

Based on burial inventory, specifically the flat-bottomed vessel, the grave is attributed to Early stage of the Poltavka culture. The radiocarbon date of the burial is 2899-2703 calBCE (4220±20 BP, PSUAMS-4264).

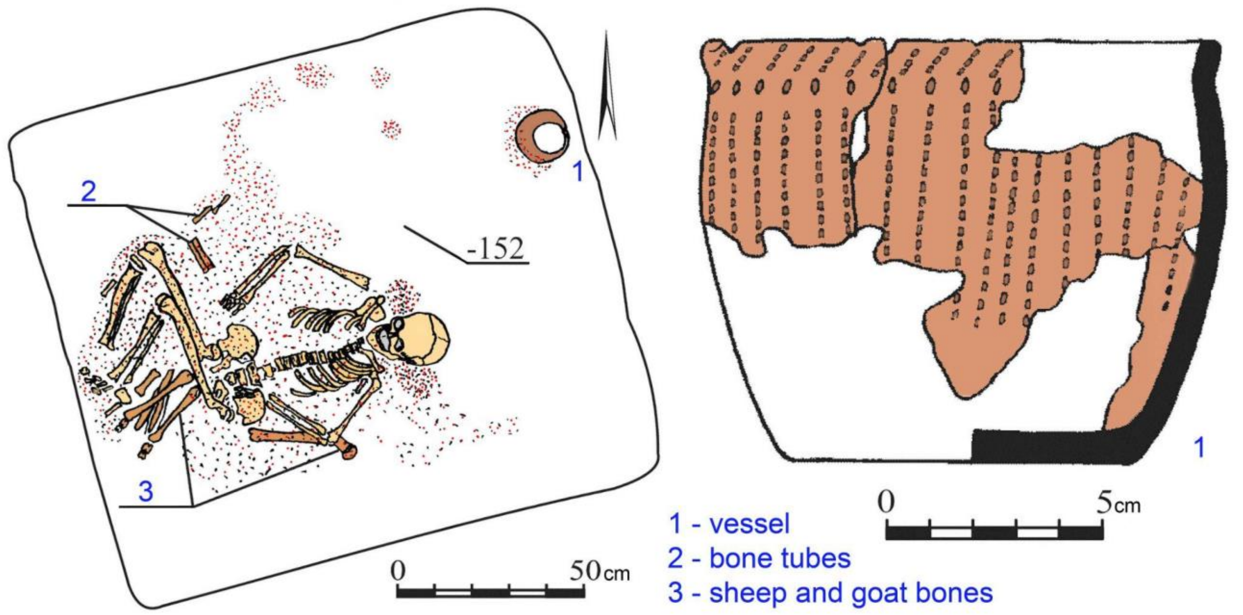

**Fig. 4.3. Grachevka-1, kurgan 2, burial 1.** Plan of the burial and a flat-bottomed pottery vessel from it (image by Pavel Kuznetsov).

#### 4.3. Grachevka-2 site (Middle Volga forest-steppe)

*Summary by P. Kuznetsov*

The site is located nearby the Grachevka-1 kurgan group.

##### *4.3.1 Grachevka-2, kurgan 5*

The diameter of the kurgan mound is 20 m, height is 0.24 m.

###### *4.3.1.1 Burial 5:1 (individual ID I0245): 2896-2589 calBCE*

Burial 1 is located 4 m to the south of the centre of the mound. The dimensions of the pit are 1.42-1.52 m × 0.75-1.2 m. The profile of the burial pit is shallow, with steep walls, a depth of -70 cm from the modern top of the mound.

The skeleton of a male aged 35-45 was positioned on his back with bent legs and outstretched arms with his head oriented to the east-northeast. The skull shows flattening of the occipital bone, likely a result of cradle deformation. According to Aleksandr Khokhlov, the skull belongs to a Europoid individual.

Pieces of ochre measuring: 1×1 cm, 1×2 cm, and 2×3 cm were discovered in the scull area. Additionally, some stains of ochre powder were found in the southern half of the burial pit. Between the knees of the deceased lay the vertebrae and ribs of an animal, identified by Natalia Roslyakova as belonging to a large ram. No other grave goods were found (Kuznetsov et al., 2018).

The burial is dated to 2896-2589 calBCE (4179 ± 55 AA-53804) and attributed to the late stage of Yamnaya culture of the Volga, based on the orientation of the skeleton, ochre presence, and C14 dating.

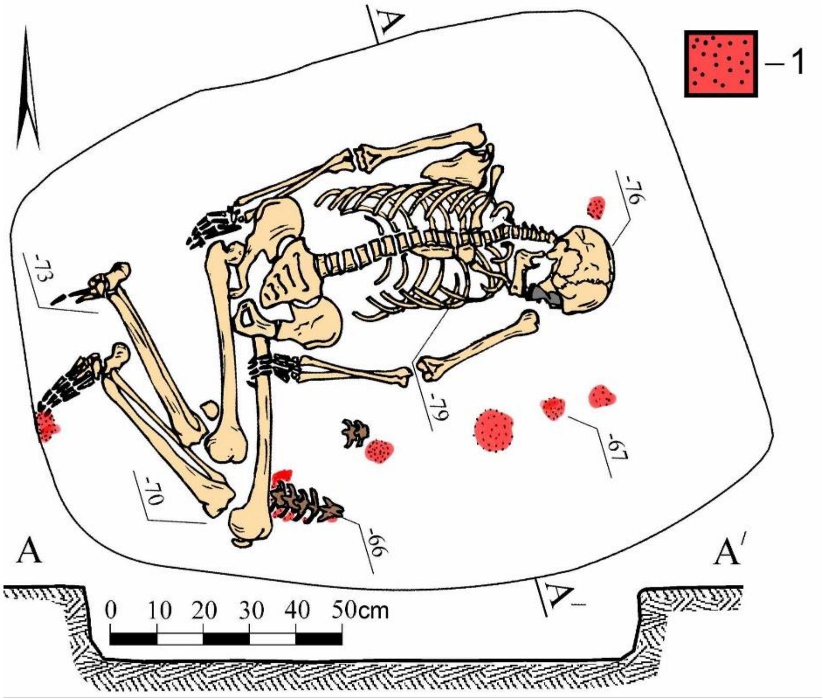

**Fig. 4.4. Grachevka-2, kurgan 5, burial 1** (image by Pavel Kuznetsov)**.**

*4.3.1.2 Burial 5:2 (individual ID I7490): 3092-2917 calBCE*

Burial 2 was excavated 1.6 meters southeast from the center of the mound, and it served as the primary burial. The dimensions of the pit were 1.74×0.8-0.94 meters. The profile of the grave was shallow, with steep walls, reaching a depth of -123 centimeters from the surface of the mound's peak. The deceased, a female older than 45 years, was laid on her back with her legs bent, arms stretched out, and head facing east. Fragments of organic bedding or a shroud were found around the bones. A clay pot stood near the occipital region of the skull. To the right of the skull, three human teeth (1) and four small flint chips measuring 0.3–0.5 cm (2) were placed. A cluster of ochre was observed around and beneath them, forming a circular patch with a diameter of up to 20 cm. According to Aleksandr Khokhlov's determination, the skull belongs to a Europoid individual.

The burial is dated to 3092-2917 calBCE [R-Combine 3110-2870 calBCE (4342±56 AA-53804); 3150-2750 calBCE (4330±60 IGAN-2875); 3282-2918 calBCE (4410±25 BP, PSUAMS-4272)] and attributed to the late stage of Yamnaya culture of the Volga, based on the orientation of the skeleton, partial ochre presence, and C14 dating.

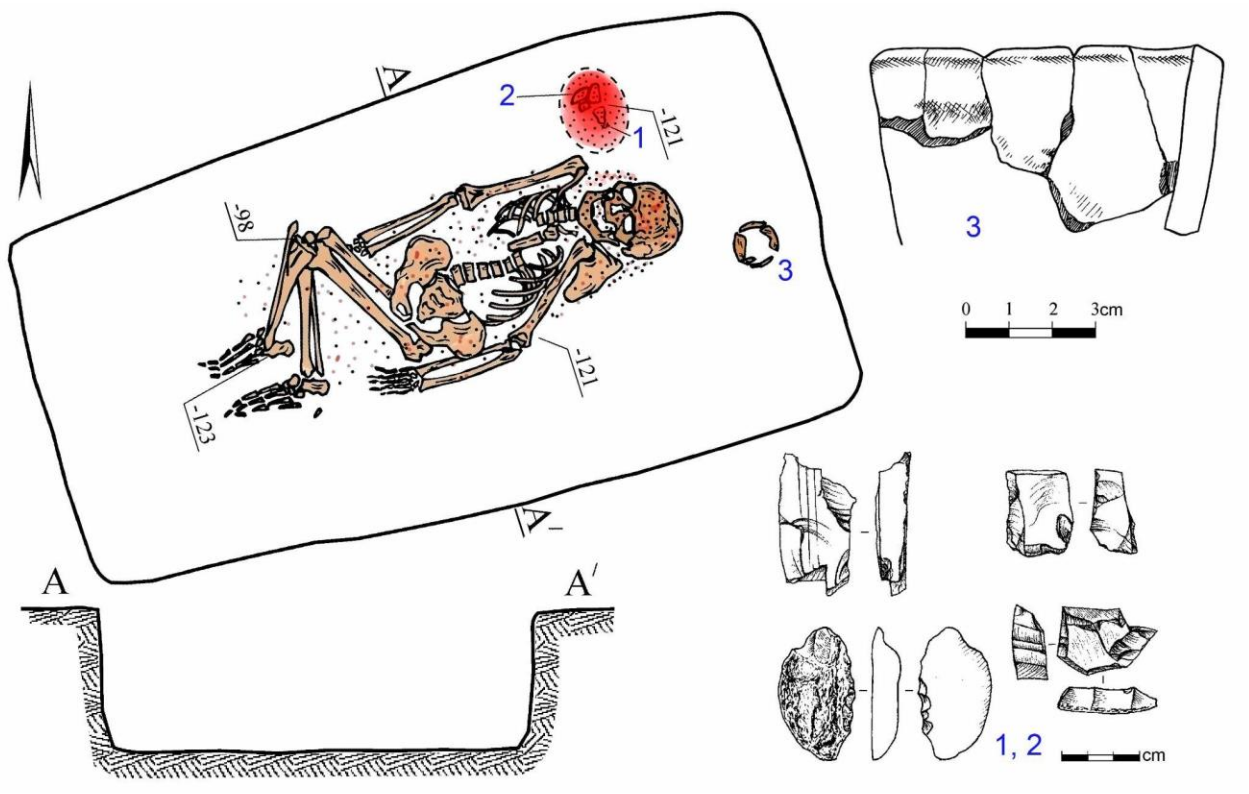

**Fig. 4.5. Grachevka-2, kurgan 5, burial 2.** 1 – human teeth, 2 – flakes, 3 – pottery vessel (image by Pavel Kuznetsov).

##### *4.3.2 Grachevka-2, kurgan 7*

The diameter of the mound is 19.5 m, height 0.32 m.

*4.3.2.1 Burial 7:1 (individual ID I0242): 3320-2885 calBCE*

Burial 1 was 9.1 m to the south-southeast of the centre and is considered peripheral. The burial pit had the shape of an elongated irregular oval and was oriented with its long sides along the northeast-southwest line, the walls were steep. The dimensions of the pit at ground level were 0.95 m X 0.25-0.58 m, depth of -104 cm from the modern top of the mound.

At the bottom of the burial pit was the poorly preserved skeleton of a female infant six-month-old. Only parts of the skull and leg bones preserved. The occipital bone of the skull lies on the ground above the bottom of the pit by 1.5-2 cm. The localization of the preserved skeleton bones suggests the position of the buried child on her back with a slight tilt to the right side. The head was placed on a "pillow" of dark soil with clay inclusions. Near the leg bones, an ochre stain with a diameter of 8 cm was recorded.

Behind the buried child's skull, at 0.08 m to the north-northeast, a destructed pottery vessel was found, with an ocher spot measuring 3.5 cm X 1.5 cm on its neck.

At the bottom of the pit, between the ochre stain and the southwest wall of the grave, a ragment of a light brown sandstone slab of elongated shape was found (Kuznetsov et al., 2018).

The burial is dated to 3320-2885 calBCE (4361±55 BP, AA-53807) and attributed to the late stage of the Yamnaya culture of the Volga region based on the orientation of the skeleton, partial ochre staining, and C14 dating.

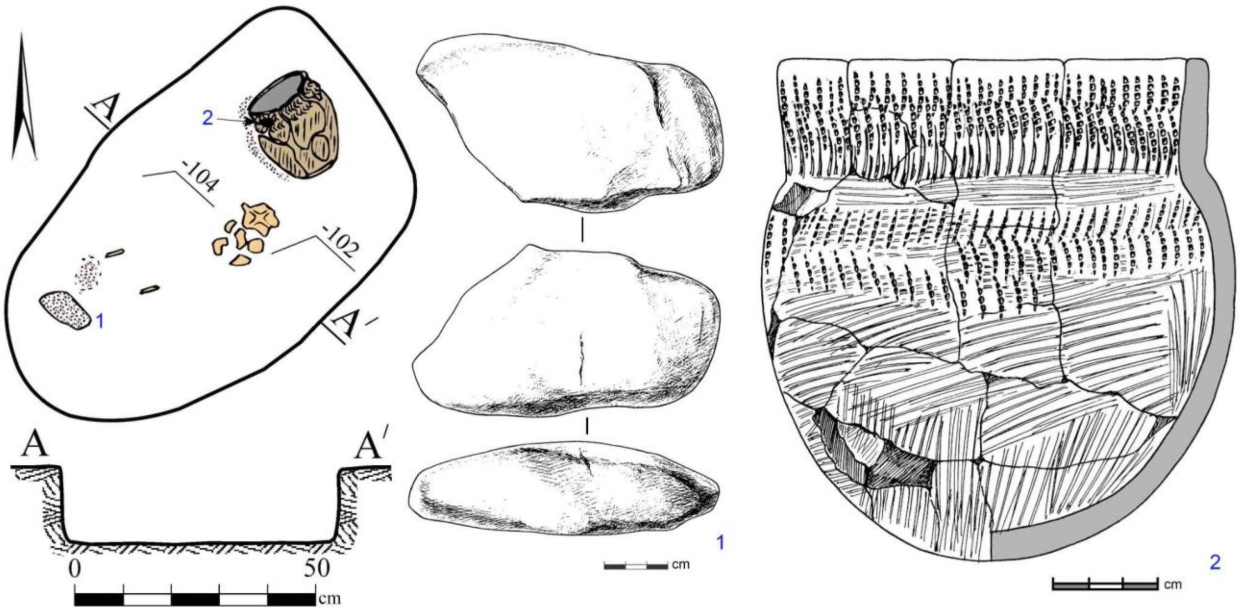

**Fig. 4.6. Grachevka-2, kurgan 7, burial 1.** 1 – stone slab piece, 2 – pottery vessel (image by Pavel Kuznetsov).

*4.3.2.2 Burial 2 (individual ID I0243): 3335-2912 calBCE*

Burial 2 was unearthed at a distance of 1.2 meters south of the mound's center and is considered as the primary burial in the kurgan. The dimensions of the pit of a rectangular shape with rounded corners were 1.76×0.8-0.94 meters. The grave was oriented along the northeast-southwest axis.

At the bottom of the grave, the skeleton of a six-month-old male infant was discovered, lying on its back with bent legs, oriented with the head facing northeast. The left arm was bent at the elbow, while the bones of the right arm were absent. Initially, the legs were positioned with the knees up but later fell to different sides. A layer of dark material, possibly a covering, was observed on and around the buried remains, with a thickness of up to 3–4 centimeters. Fragments of black-colored bedding, resembling leather, with a metallic sheen, were found at the bottom of the pit. Ochre was scattered throughout the grave, with the highest concentration observed on the skull, pelvis, and feet of the deceased. Pieces of ochre were also found near the left femur bone.

The grave goods are represented by two potrttery vessels, one discovered behind the skull of the buried individual, lying on its side, and another, a large one, found against the western wall. Adjacent to the left leg's shinbone were fragments of a Unio shell and two small flint chips without traces of processing.

The burial is dated to 3335-2912 calBCE (4419±56, AA-53808) and attributed to the late stage of the Yamnaya culture of the Volga region based on the orientation of the skeleton, partial ochre staining, and C14 dating.

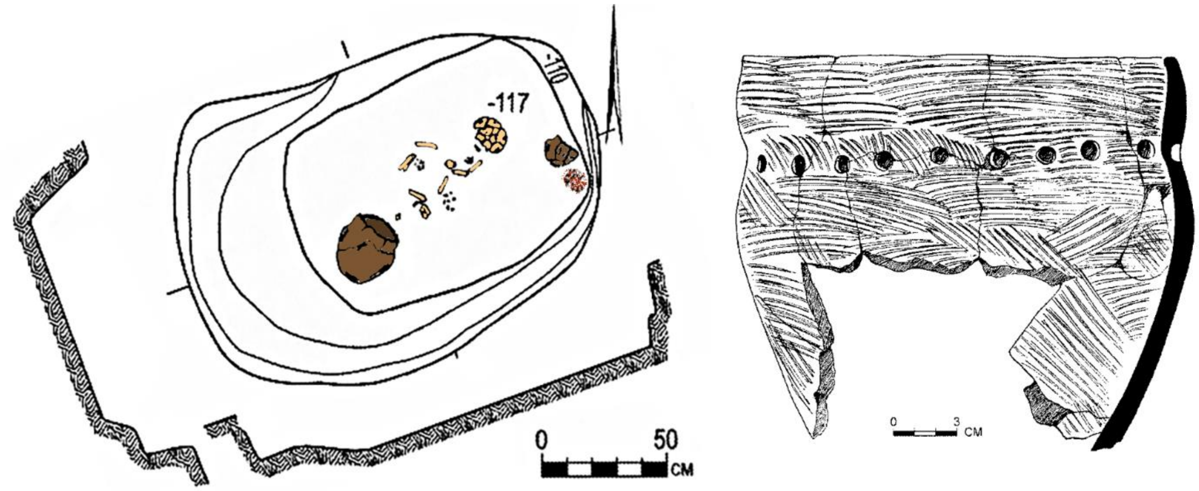

**Fig. 4.7. Grachevka-2, kurgan 7, burial 2.** Plan of the burial and one of the pottery vessels found in the grave (image by Pavel Kuznetsov).

#### 4.4 Krasikovo-1 site (Volga-Ural steppe)

*Summary by P. Kuznetsov*

The site occupies an elevated area in the Obschy Syrt Upland on the left bank of the Tok River, a right tributary of the Samara River. It includes six rounded earthen mounds ranging from 13 to 40 meters in diameter and from 0.25 to 2 meters in height. Topographically, the mounds are divided into three groups:

1) elongated in a chain along the southwest-northeast line, mounds 1, 2, and 3;
2) located in a chain along the northwest-southeast line, with mounds 4 and 5 situated 40 meters to the north-northwest of mound 3;
3) mound 6, which is the largest in the complex.

The total length of the burial mound group from north to south is approximately 170 meters, with an estimated width of the area of about 150 meters.
The site was investigated by the Orenburg archaeological expedition.

###### ***4.4.1 Krasikovo-1, kurgan 1***

The diameter of the mound is 28 meters, with a height of 1.12 meters. According to Olga Khokhlova, the paleosols beneath mound 1 contain phytoliths of arid flora.

###### *4.4.1.1 Grave 1:1 (individual ID I12241): 2858-2577 calBCE*

Burial 1 was located 2 meters south of the center of the mound and was the main burial in the kurgan. The dimensions of the pit at ground level were 190×150 cm, and at the bottom of the pit were 165×100 cm. The depth was up to -295 cm from ground level. The pit was likely covered with planks.

The skeleton of an adult was found at the bottom of the pit, positioned on the right side with some inclination onto the back, with the legs bent to the right. A large part of the skeleton was preserved in situ: the legs, pelvis, partially the spine and ribs, and also the left arm, which lay over the chest, with the hand resting on the pelvic and thigh bones. The individual was oriented with the head to the east-southeast. The body lay with a turn to the right side. The legs of the buried individual were bent.

As defined by Aleksandr Khokhlov and A.A. Grigoryev, the skull is Europoid, dolichocephalic, and mesomorphic. The skeleton bones are large. The estimated height is approximately 177.6-178.2 cm. He had Y-haplogroup R1b (R-M269) and mt-haplogroup H2a1.

At the bottom of the burial pit, beneath the skeleton, there was a layer of ochre. Ochre covered the upper part of the body and legs.

The skeleton lay on a plant mat, which was best preserved in the western part of the mound, under the leg bones. Only white decay remained from the mat, and the bones of the feet were heavily stained with ochre. Along the southern wall and bottom of the burial pit, where the pelvic bones of the buried individual were located, a large fragment of red ochre was found (Morgunova et al., 2014; Morgunova and Kul'kova, 2019).

The burial is attributed to the late stage of the Yamnaya-Poltavka culture of the Volga-Ural region. The basis for this conclusion includes the position on the right side, orientation, partial ochre covering, and radiocarbon dating, 2858-2577 calBCE (4110±20 BP, PSUAMS-10776).

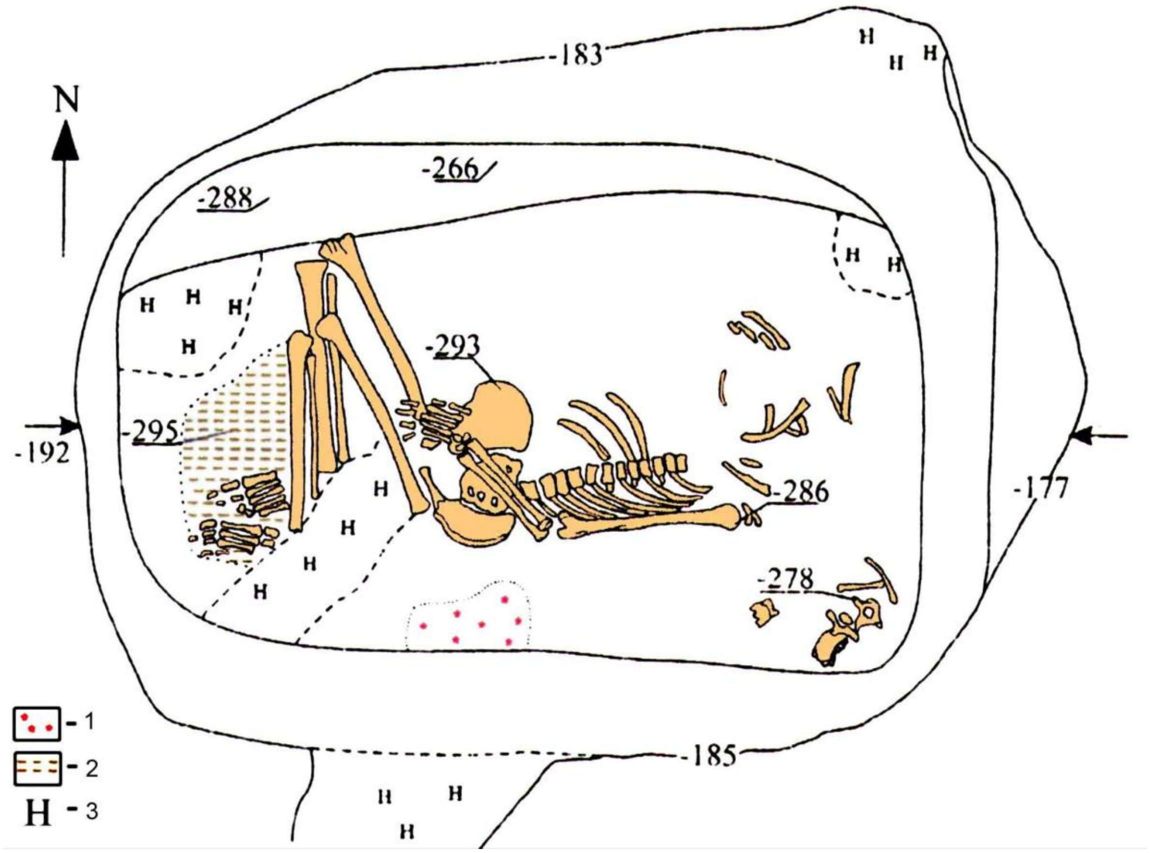

**Fig. 4.8. Krasikovo-1 site, kurgan 1, burial 1. 1 – ocher, 2 – white decay (plant mat), 3 – krotovinas (image by Pavel Kuznetsov).**

##### *4.4.2 Krasikovo-1, kurgan 2*

The diameter of mound 2 reached 24 meters, with a height of 61 centimeters on the southern slope and 89 centimeters on the northern slope. The mound was constructed using soil from the circular ditch surrounding the submound platform. The diameter of the submound platform, surrounded by the ditch, was 16-17 meters. According to Olga Khokhlova, the paleosols beneath mound 1 contain phytoliths of arid flora.

###### *4.4.2.1 Grave 2:1 (individual ID I12242)*

Burial 2:1 was located 2 meters south of the mound's center and is considered the primary burial within the mound.

The grave pit dimensions at the surface level were 280x221 centimeters, and at the pit bottom level, they were 200x120 centimeters, with a depth of -309 centimeters from the surface. Human bones and fragments of wood from the covering were found at different levels within the pit. In the northeast corner, at a depth of -122 to -155 centimeters from the surface (or -262 to -295 centimeters from the top), compactly arranged human bones were discovered, including pelvic bones, leg and arm bones, and ribs, which were partially disturbed by a groundhog burrow. Some bones were displaced to the southeast corner of the pit. Upon the bones, a pectoral or gorget made of boar tusk was found.

A man's skull aged 30-40 years was found at a depth of -292 centimeters from the surface. It was positioned vertically, facing east, on a clay pedestal reinforced with densely packed clay around the edges. The clay layer had cracked significantly, presumably due to drying. The mandibula was displaced northeast of the skull. A large quartzite arrowhead was placed in front of the facial bones, above the clay layer.

Therefore, the burial consisted of a separate interment of the skull on a clay pedestal and a cluster of compactly arranged bones in the eastern part of the mound, partially disturbed by two krotovinas (ground hogs). The burial may have remained uncovered for some time, as the clay layer around the skull developed deep cracks.

The skull belongs to a male, 30-40 years old. According to Aleksandr Khokhlov and A.P. Grigoriev, the skull was of a hypermorphous Europoid variant. The skeleton had the largest bones in the burial ground. The body length was 181.8-183.3 centimeters. His Y-haplogroup was R1b (R-Y87909), mt-haplogroup U5a1a1.

Grave goods included a processed boar tusk and a quartzite arrowhead.

A bedding consisting of three layers was found. The first layer (the lowest) was a 5-centimeter-thick coating of white clay. The second layer was a covering of brown organic matter, no more than 0.5-1 centimeters thick, containing small pieces and ochre powder. The third layer, no more than 1 centimeter thick, consisted of a bedding of reeds or rushes. The thickest layer of bedding was concentrated in the western part of the mound, where the skull rested. A patch of ochre was found on the bedding in the northwest part of the pit. In the eastern part the bedding was disrupted by a krotovina (Morgunova et al., 2014; Morgunova and Kul'kova, 2019).

*Grave 2:2 (individual ID I10033): 2900-2697 calBCE*

Grave 2 was located 4.5-5 m to the south of the kurgan's center and was interpreted as the secondary burial in the mound.

The grave pit was recorded at the buried soil level. In the grave, an adult genetically determined female was found in a flexed position on the back, with her head to the west. The skull (highly fragmented) and the leg bones were oriented to the left side. The legs were in a strongly flexed position. The ribs and bones of the left arm were fragmented and found in a non-anatomical arrangement. The left hand was positioned on the pelvic bones, while the right hand partially overlapped the bones of the right foot. Red ochre stains were observed around the skull, ribs, and right foot (Morgunova et al., 2014; Morgunova and Kul'kova, 2019).

According to the Aleksandr Khokhlov and A.A. Grigoryev, the skull closely resembles ancient North Ural types. The muscular relief is well developed. Her estimated height is approximately 152.6 cm. Her mt-haplogroup was T1a1, a haplotype found earlier among European Neolithic farmers such as LBK.

This burial was dated to 2900-2697 calBCE (4215±25 BP, PSUAMS-6152) and attributed to the late stage of the Yamnaya culture in the Volga-Ural region.

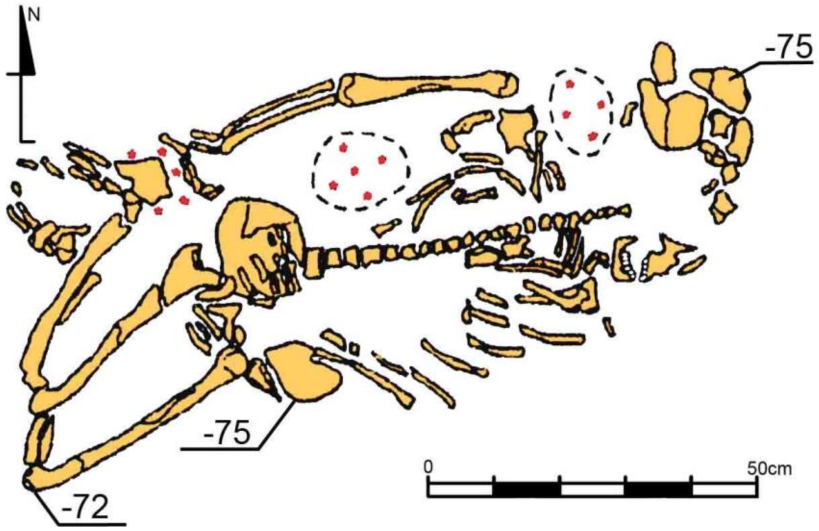

**Fig. 4.9. Krasikovo-1 site, kurgan 1, burial 2 (image by Pavel Kuznetsov).**

##### *4.4.3 Krasikovo-1, kurgan 3*

The diameter of kurgan 3 on the modern surface was 18 meters, with a height ranging from 0.25 to 0.3 meters, however, the excavations revealed that the initial platform, on which the kurgan was constructed, reached only 17 meters. The highest point of the kurgan was in its center.

###### *4.4.3.1 Grave 3:3 (individual ID I10034): 2881-2633 calBCE*

Burial 3, secondary in the kurgan, was located 1.4 m south-southwest of the kurgan center. It was recorded at a depth of -118 to -125 cm from the surface. The boundaries of the pit were not traced. The child's skeleton lay slightly flexed on the back, with the head to the northwest. The tibiae and feet of the deceased were absent (it is impossible to determine whether they were removed by rodents or deliberately severed during burial). The arms of the deceased were positioned behind the back. No inventory was found (Morgunova et al., 2014; Morgunova and Kul'kova, 2019; Khokhlov and Grigoryev 2019). Cultural attribution is unknown, however, the radiocarbon date 2881-2633 calBCE (4165±25 BP, PSUAMS-10771) points at the late Yamnaya period.

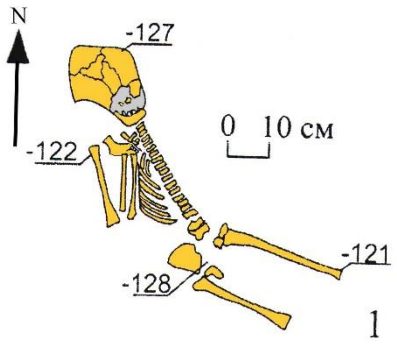

**Fig. 4.10. Krasikovo-1 site, kurgan 3, burial 3 (image by Pavel Kuznetsov).**

###### *4.4.3.2 Grave 3:4 (individual ID I12243): 2851-2506 calBCE*

Burial 4 was the primary one in the kurgan. The pit was covered with planks placed along the long sides of the pit. Its dimensions were 180×120 cm, and at the bottom, they reached 170×90 cm. The depth was up to -249 cm from the highest point of the mound.

The deceased was put in the pit on their back with legs strongly bent to the right. The orientation of the body was with the head to the northwest, facing south. The hands of the deceased were placed on the pelvic and femur bones. The skeleton lay on straw, with traces of its weave in some places. The bones of the feet were heavily stained with ochre.

According to the classification by Aleksandr Khokhlov and A.P. Grigoriev, the skull is Europoid, gracile, and leptomorphic. The skeleton bones are large. Physical development was strong. The body length was 176 cm. His Y-haplogroup was R1b (R-KMS67), mt-haplogroup H6a1a.

At the bottom of the pit, a bedding was laid. It consisted of a dense brown-colored mass with a well-preserved plant structure. Scanning electron microscopy showed that this mass included decomposed stems of steppe grasses and amorphous heavily decomposed animal organic matter. Based on this, we reconstruct the bedding as a two-layered item.

In the southeast corner of the mound, to the southeast of the skull, the reains of a ceramic vessel were found. The vessel had an egg-shaped form, gray in color, with the upper part decorated with imprints of cord stamps (Morgunova et al., 2014, Morgunova and Kul'kova, 2019).

This burial was dated to 2851-2506 calBCE (4095±20 BP, PSUAMS-10737) and attributed to the late stage of the Yamnaya culture in the Volga-Ural region.

#### 4.5 Krasnosamarskoe-4 site (Middle Volga steppe)

*Summary by P. Kuznetsov & D. Anthony*

The kurgan group is situated 4 kilometers southwest of the village of Krasnosamarskoye in the Kinelsky District, along the shoreline of one of the fishery ponds. The burial ground was discovered in 1995 during a survey by Pavel Kuznetsov, Oleg Mochalov, and David Anthony. In 1999, barrows 1, 2, and 3 were excavated by Pavel Kuznetsov and David Anthony, kurgan 5 was excavated in 2005, and kurgan 7 in 2008, by Pavel Kuznetsov and Oleg Mochalov.

Geomorphologically, the site is located on the second terrace of the Volga River valley, contiguous with the first terrace of the Samara River, a left tributary of the Volga. The terrace is composed of alluvial clays and loams, with a thickness of approximately 3-5 meters, underlain by sands and a layered alluvial deposit. Until 1966, an oxbow of the Samara River was located in close proximity to the site. The terrain is predominantly flat, with absolute elevations around 50 meters and a height amplitude associated with micro-relief of 2 meters. The area belongs to the southern part of the forest-steppe zone, bordering the steppe zone with chernozem soils.

##### *4.5.1 Krasnosamarskoe-4, kurgan 7*

###### *4.5.1.1 Burial 7:5 (individual ID I11838): 2851-2498 calBCE*

The burial was made at a depth of 1.99 meters below ground level, or 0.73 meters below the subsoil surface. The pit had a rectangular shape with rounded corners, measuring 2.02 meters by 1.33 meters, oriented in the west-east direction.

At the bottom, the skeleton of a 55-65-year-old male was found lying on his back with a slight tilt to the right, oriented with the head to the west-southwest. Initially, the buried individual's skull was positioned on a raised surface, possibly an organic pillow, but later it tilted with the facial part to the left. The arms were stretched along the body, with the left hand extended and the right hand bent at a right angle. The legs were strongly bent at the knees and initially positioned with the knees upward, but under the pressure of the soil, they tilted to the right.

Beneath the skeleton, there was a layer decayed organics of black color (likely remains of bedding) that nearly followed the contours of the pit, with a thickness of up to 2 millimeters.

In the area of the right lower ribs of the buried individual, an oval-shaped pebble with a diameter of 2.3 centimeters was discovered. Traceological analysis conducted by Igor Gorashchuk indicated that the pebble could have been used as a projectile for a sling.

According to Aleksandr Khokhlov's classification, the skull is mesomorphic, mesocranial, and Europoid. It fits well into the existing small series of skulls from previously excavated barrows in the same burial ground.

This burial was dated to 2851-2498 calBCE (4085±25 BP, PSUAMS-10774) and attributed to the late stage of the Yamnaya culture in the Volga-Ural region.

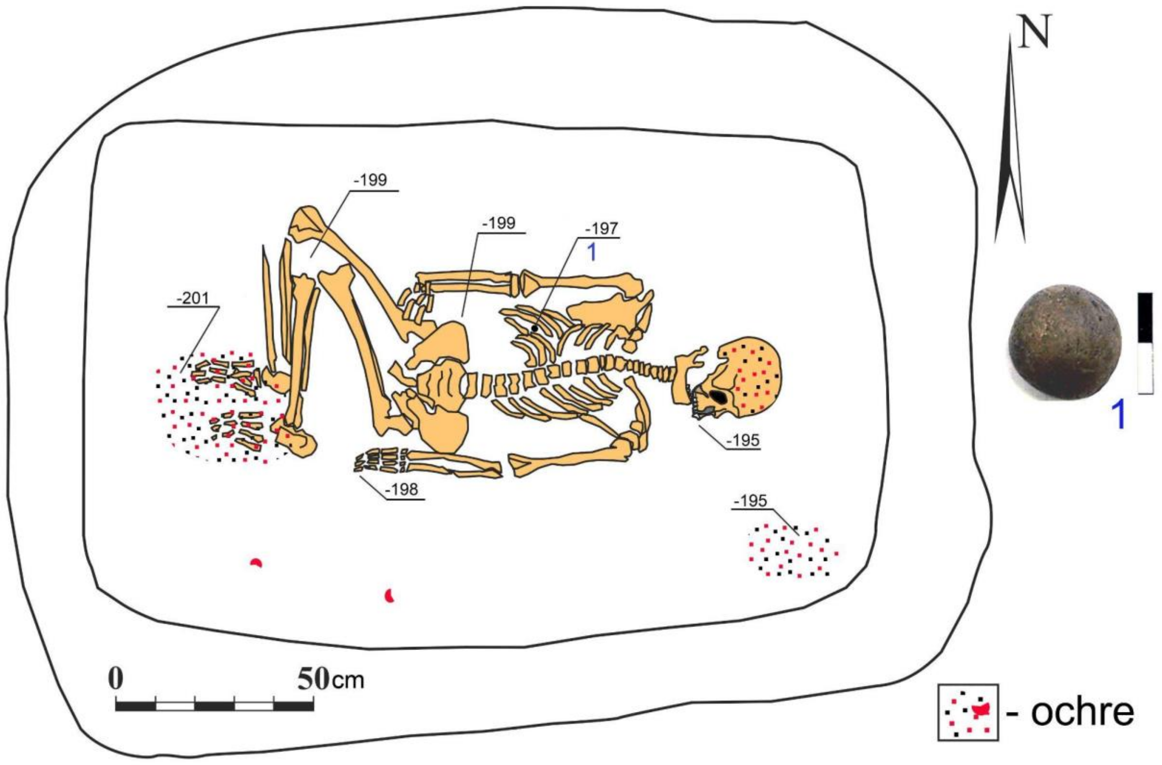

**Fig. 4.11. Krasnosamarskoe-4, kurgan 7, burial 5. 1 – pebble (image by Pavel Kuznetsov).**

#### 4.6 Kurmanaevka-3 site (Volga-Ural steppe)

*Summary by P. Kuznetsov*

Kurmanayevka-3 was excavated under the direction of S.V. Bogdanov and A.Yu. Kravtsov. The site is located in the western part of the Orenburg Trans-Urals, on the right bank of the Buzuluk River (a left tributary of the Samara River, left tributary of the Volga), on the floodplain terrace, the rear part of which is bounded by the ancient lake Chernoe. The kurgan cemetery consists of 3 mounds, all of which belong to the late Yamnaya period. The mounds are earthen and rounded in plan.

##### *4.6.1 Kurmanaevka-3, kurgan 3*

The diameter of the mound is 20 m, height is 0.23 m. (I25159

###### *4.6.1.1 Burial 3:1 (individual ID I0441): 2876-2635 calBCE*

Burial 1 was located under the center of the mound, the only grave in the mound. The pit is uneven and trapezoidal, and the bottom is uneven, with organic bedding at the bottom. Dimensions of the pit at the ground level: 260×150 cm, at the bottom level of the pit: 150×110 cm. Depth to -193 cm from the modern top of the mound.

The skeleton of an adult woman was located at the bottom of the pit in a position on the right side, with some tilt onto the back, with legs bent to the right. In situ, a large part of the skeleton was preserved: legs, pelvis, partial spine, and ribs. The left arm, lying over the chest, its hand is placed under the right arm. The right arm is semi-bent and directed towards the right thigh. The buried woman was oriented with her head to the Northeast. The legs of the buried woman were bent. At the bottom of the burial pit, near the right pelvis and bones of the right leg, ochre was found scattered. Ochre was sprinkled over the right hand.

Here, on top of the hand, beneath the layer of ochre, lay a bone-polished pin and a ring-shaped cut of a hollow bone (Bogdanov 1999).

The woman's age was upper mature. Her mt-haplogroup was H2b, a haplogroup found among European farmers such as LBK and Lengyel.

C14 data 2876-2635 calBCE (4159±19 BP) [R_Combine: (4234±60 BP, AA-47805), (4150±20 BP, PSUAMS-8721)]

Cultural attribution - Late Yamnaya-Poltavka stage of the Yamnaya culture of the Volga-Ural region based on the position on the right side, the orientation, partial ochre sprinkling, and C14 dating.

#### 4.7 Kutuluk-1 site (Middle Volga steppe)

*Summary by P. Kuznetsov*

The site is located at the edge of the first terrace on the left bank of the Kutuluk River, a right tributary of the Bolshoi Kinel River, left tributary of the Volga, and occupies the left bank of the small stream Trostinka. The kugran group includes 7 mounds.

##### 4.7.1 Kutuluk-1, kurgan 3

###### 4.7.1.1 Burial 3:4 (individual ID I11840): 2921-2878 calBCE

Burial 4 is the primary burial in kurgan 3 and was located in the central part of the mound. The burial pit had a rectangular shape with oval corners, with vertical walls, oriented with the long sides in the west-east direction. The dimensions of the pit at the level of the subsoil were 1.9 meters by 1.32 meters, with a depth of -157 centimeters from ground level.

At the bottom of the pit, there were the bones of a 40-45-year-old male, which had been displaced by borrowing rodents. It is likely that the buried individual was lying on their back, with the head oriented to the west. The bones of the skull and part of the left leg were partially covered with ochre.

This burial was dated to 2921-2878 calBCE (4280±25 BP, PSUAMS-7868) and attributed to the late stage of the Yamnaya culture in the Volga-Ural region.

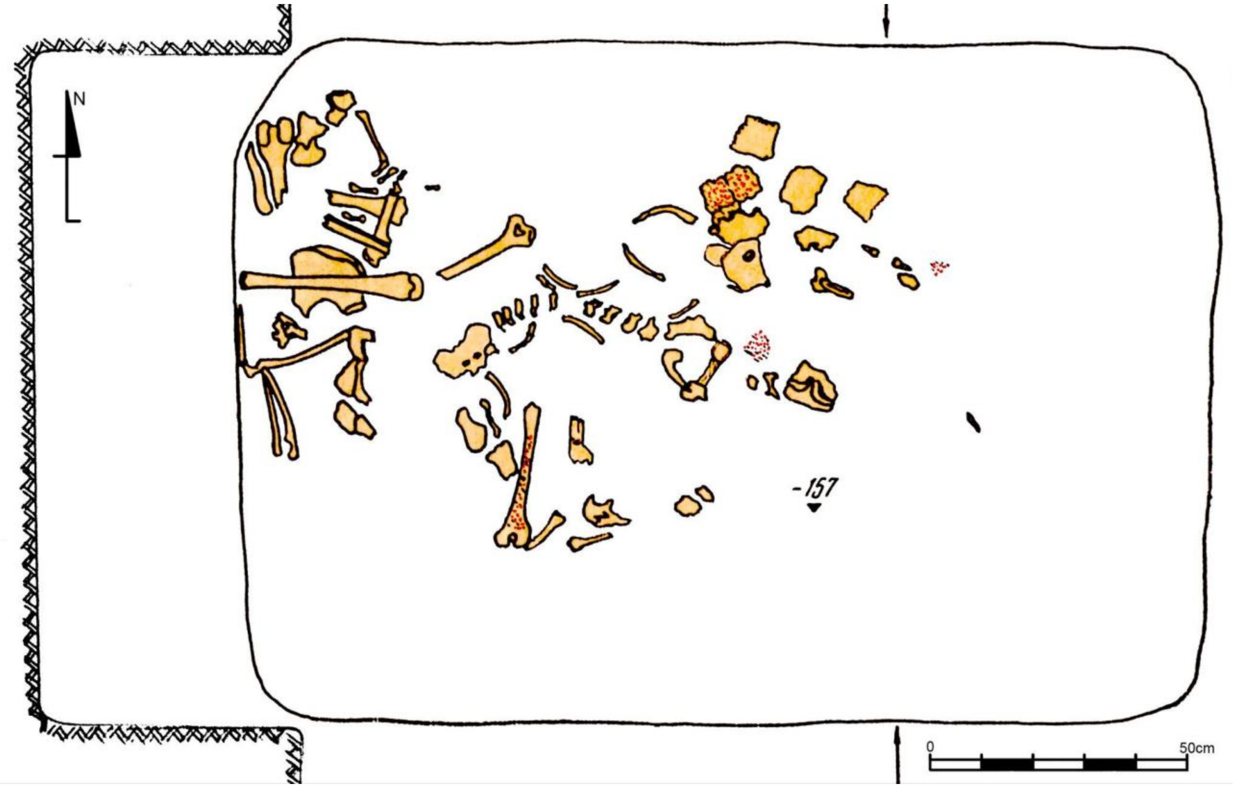

**Fig. 4.12. Kutuluk-1, kurgan 3, burial 4** (image by Pavel Kuznetsov)**.**

##### 4.7.2 Kutuluk-1, kurgan 4

The kurgan was excavated by Pavel Kuznetsov in 1990.

###### 4.7.2.1 Burial 4:1 (individual ID I20158): 2925-2880 calBCE

Burial 1 was the primary grave in the mound . It was located 1.6 m to the south of its center. The grave pit, oriented along the northeast-southwest line, had a rectangular shape, 2.8 to 1.7 m, the corners of the pit were rounded and the walls were steep. The bottom of the grave was at 2.04 m from the top of the kurgan.

At the bottom, a skeleton of a male person aged 30-39 years old was recorded in the supine position. The skull was slightly displaced onto the chest. The right arm was extended along the body, the left was half bent at the elbow. The legs were bent and stood upright, after the burial they fell to the right. The front part of the skull, pelvis, and legs were intensively painted with ocher. The skeleton was underlaid by a layer of organic decay 0.5-0.7 cm thick reflecting a mat made from bark or other plant material.

On the elbow of the left arm of the buried man lay a solid copper sword-scepter 48.7 cm long, a unique object (Kuznetsov 1990).

According to the definition of Alexander Khokhlov, the skull belongs to the massive, hypermorphic-mesocranial type, widespread in the territory from the Volga region to the Dnipro. The height of the buried man was about 176 cm. During his life, he engaged in intense physical activity, possibly moving a lot on foot.

On the leg bones of the buried person, there are no curvatures diagnostic for horse riding, characteristic of later nomads.

The buried person was dated by 14C several times, which yielded ages of 3340-2890 calBCE (4400±70, OxA-4306), 3340-2880 calBCE (4370±75, AA-12570), 2919-2875 calBCE (4270±25 BP, PSUAMS-10741). The combination of the dates gives 2925-2880 calBCE as the most probable date of the burial.

The burial provides a reference assemblage for the late phase of the Yamnaya culture in the Middle Volga region.

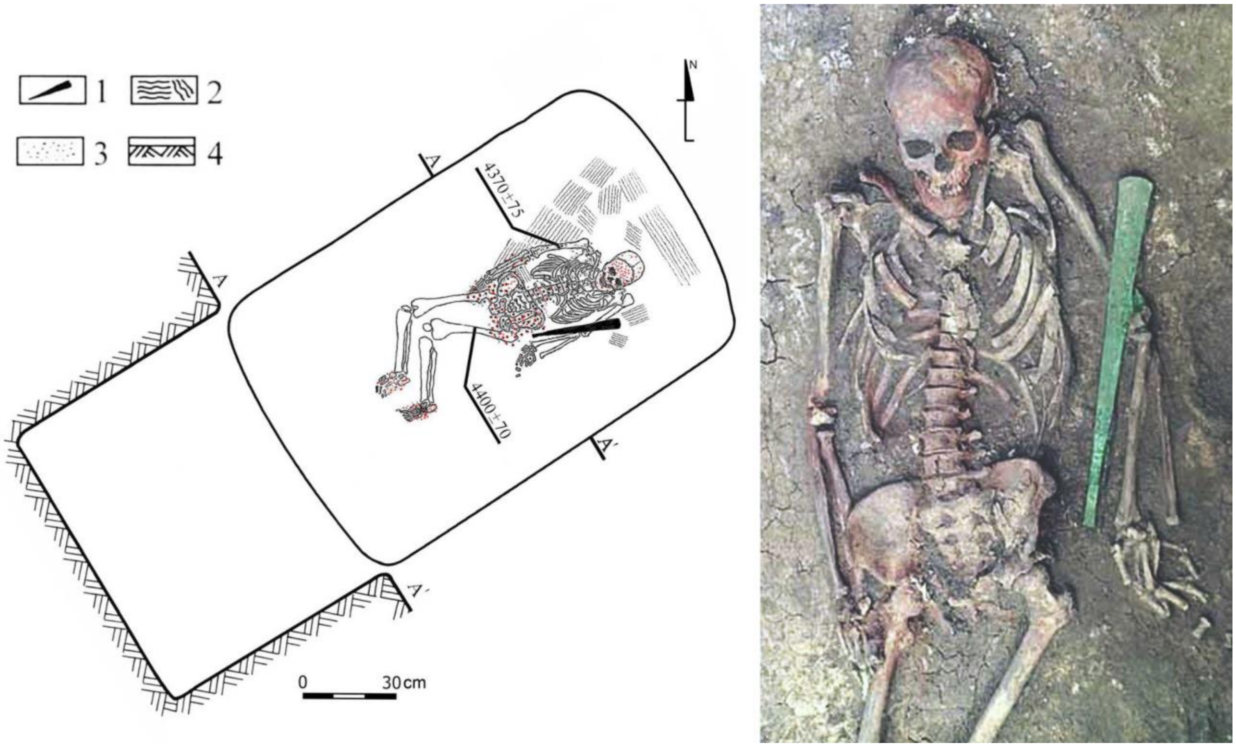

**Fig. 4.13. Kituluk-1 site, burial 1 under kurgan 4.** 1 - sword-sceptre, 2 - decayed organics, 3 - ocher, 4 - subsoil level on the profile (image and photo by Pavel Kuznetsov).

#### 4.8 Lopatino-1 site (Middle Volga forest-steppe)

*Summary by P. Kuznetsov*

The site is located on the edge of a floodplain terrace of the left bank of the Sok River, a left tributary of the Volga River. The ensamble comprises 35 burial mounds, making it the largest kurgan site complex in the Krasnoyarsk archaeological microdistrict, in terms of the number of mounds.

##### *4.8.1 Lopatino-1, kurgan 22*

Excavations by I.B. Vasilyev, Kuznetsov, and A.P. Semenova in 1987.

###### *4.8.1.1 Burial 22:1 (individual ID I10714)*

Burial 1 was excavated in the central part of the mound. The burial pit was damaged by borrowing rodents and later digging. Presumably, it was oriented along the northeast-southwest axis. The dimensions and depth of the pit are undetermined.

Various bones, fragments of a skull, and parts of the mandibula of a male aged 7-14 years were found at different depths within the grave pit, partially covered with ochre. The burial is attributed to the late stage of the Yamnaya culture of the Volga region, based on the pit orientation and partial ochre staining.

#### 4.9 Lopatino-2 site (Middle Volga forest-steppe)

*Summary by P. Kuznetsov*

See Lopatino-1 site.

##### *4.9.1 Lopatino-2, kurgan 2*

###### *4.9.1.1 Burial 2:3 (individual ID I25026): 3011-2702 calBCE*

Burial 3 is the primary grave in the kurgan, excavated in the central part of the mound.

The dimensions of the grave pit are 1.82×1.38 meters, it had shallow, vertical walls, and a depth of up to -133 cm from ground level. A male individual aged 50-55 years old was buried in the supine position with his legs bent at knees and extended arms, with his head to the northwest. According to Aleksandr Khokhlov's assessment, the skull of the buried individual is of medium robustness, with a long, low, dolichocephalic cranial vault, a wide and sloping forehead, a wide and low face, and a relatively well-profiled face with a somewhat low nasal bridge, characteristic of the Europoid type.

Abundant ochre staining was observed around the skull, arms, ribs, and thigh bones. It appears that the ochre was sprinkled after the individual was placed in the pit. Remnants of a covering were found on the skeleton - a thin organic layer of dark brown color with a thickness of up to 0.5 cm. Organic bedding remnants of dark brown color with a thickness of up to 1 cm were found at the bottom of the pit.

A stone arrowhead made of dark red flint was found 10 cm south of the skull. It has a triangular shape with a notch at the base. Another arrowhead, made of light green flint, was found between the ribs of the deceased under the sternum. It also has a triangular shape with a marked notch at the base. The arrowhead points towards the head of the buried individual, suggesting he may have been killed by an arrow (Kuznetsov et al., 2023).

The burial is attributed to the late stage of Yamnaya culture, dating from 3010-2700 calBCE, based on orientation, partial ochre staining, and radiocarbon dating 3011-2702 calBCE (4270±40, GIN 11456).

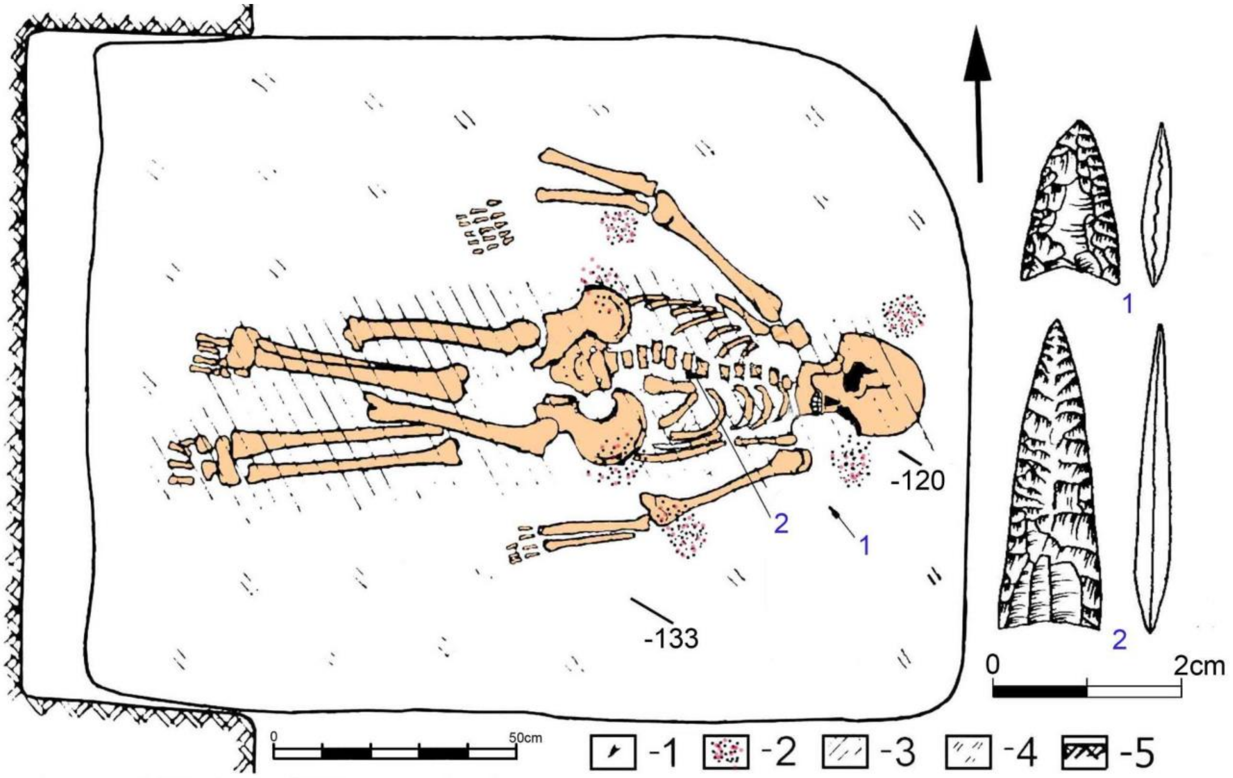

**Fig. 4.14. Lopatino-2, kurgan 2, burial 3** (image by Pavel Kuznetsov)**.**

#### 4.10 Leschevo-1 site (Middle Volga steppe)

*Summary by P. Kuznetsov*

The site is located 750 meters southwest of Lake Leschevo (an old oxbow on the ancient channel of the Samara River) and 2 kilometers west of the northeast outskirts of the village of Utevka. It is situated on the edge of the floodplain terrace and was discovered by Oleg Mochalov.

##### *4.10.1 Leschevo-1, kurgan 1*

The diameter of the mound is 20 meters, and the height is 0.3–0.35 meters. Beneath the mound, burial 1 and one completely empty round pit of unclear purpose were discovered.

###### *4.10.1.1 Burial 1:1 (individual ID I6730): 3091-2918 calBCE*

Burial 1 is located 3.4 meters southeast of the center of the mound and is the primary and only burial in this mound.

The dimensions of the pit are 2.22 meters by 1.35 meters, with a deep profile and vertical walls, reaching a depth of up to -152 cm from ground level.

Two skeletons were found at the bottom of the pit:

Skeleton 1 (ID I6730): A woman aged 20-30 years, placed on her back with bent legs and arms semi-bent at the elbows. The skull and lower jaw are lying on the spine, with the head facing to the northwest.

Skeleton 2: An infant's skeleton found near the left pelvic bone of Skeleton 1. It is completely covered with ochre (Kuznetsov and Mochalov 2014a, b).

Between the southwest corner of the pit and the left foot of Skeleton 1, two objects were found lying close to each other:

1. A flint knife measuring 7.2 cm in length and 3.8 cm in width, with sharp cutting edges. It likely served as a cutting tool.
2. A cylindrical sandstone disk measuring 4.3 cm in diameter and 4.3 cm in height. It could have been used as a whetstone or for grinding mineral compounds such as ochre.

The burial is dated to 3091-2918 calBCE (4390±20 BP, PSUAMS-4159)and attributed to the late stage of Yamnaya culture.

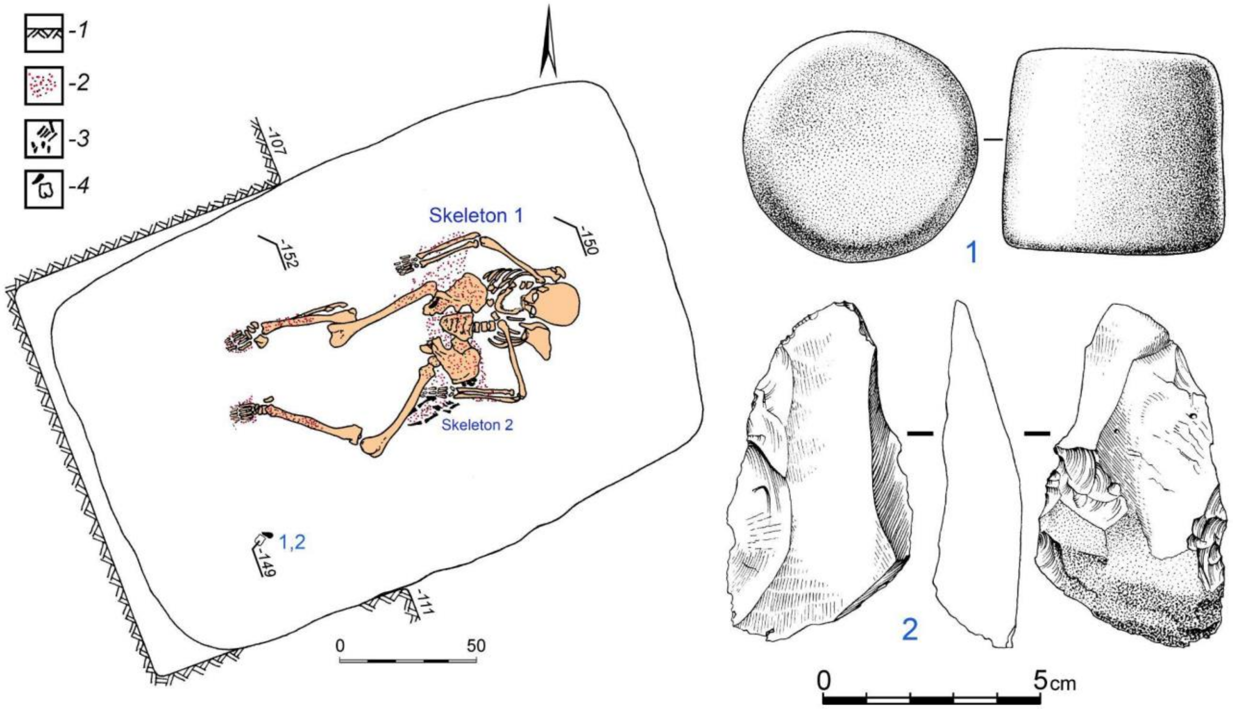

**Fig. 4.15. Leschevo-1, kurgan 1, burial 1** (image by Pavel Kuznetsov). 1-2 – grave goods.

##### *4.10.2 Leschevo-1, kurgan 2*

Kurgan 2 is located in close proximity to the larger kurgan 1. Initially, it appeared to be the largest in the cemetery. However, in coarse of the total station measuring, it was determined that the mound sits on a natural elevation platform. The diameter of the mound is 16 meters, and the height is up to 0.35 meters.

*4.10.2.1 Burial 2:2 (individual ID I6731): 3083-2916 calBCE*

Burial 2 is located 0.6 meters to the south-southwest from the center of the mound, and it is the primary burial. The burial pit 2.38 meters by 1.83 meters was covered with planks, with maximum dimensions of 2.5 meters by 0.2 meters by 0.1 meters. The walls of the burial pit gradually narrowed towards the bottom. The dimensions of the pit at the bottom level are 2.02 meters by 1.35 meters, with a depth of up to -163 centimeters from the ground level.

At the bottom of the burial pit, the skeleton of a male aged 50-60 years was found, lying on his back with a tilt to the left side, with his head to the east-southeast. The skull shifted towards the left shoulder blade. The skull lay on an elevation made of subsoil clay. The spinal column, rib cage, and arm bones lay predominantly on the posterior (dorsal) surface. The arms of the buried individual were stretched alongside the body. The bones of the forearms and the right hand were found on the right thigh. The left hand lay beneath the left femur. The pelvic bones lay on the left side. The legs of the buried individual were bent to the left at the knees. Presumably, the buried individual was placed in a fetal position on their back. Subsequently, under the pressure of the soil, the head of the buried individual shifted towards the long southern wall of the burial pit.

At the bottom of the burial pit, beneath the skeleton, a layer of ochre was found. However, there was no ochre sprinkling on the buried individual itself.

Near the right shoulder of the skeleton, a copper dagger with a bone handle was discovered. Its total length is 22.05 centimeters, including blade of 14.6 x 4.9 centimeters. The tip has an oval ending. The handle of the dagger is possibly made of antler. The length of the handle is 7.45 centimeters, and its width is 3.9 centimeters (Kuznetsov and Mochalov 2014a, b).

The burial is dated to 3083-2916 calBCE (4380±20 BP, PSUAMS-4160) and attributed to the late stage of Yamnaya culture.

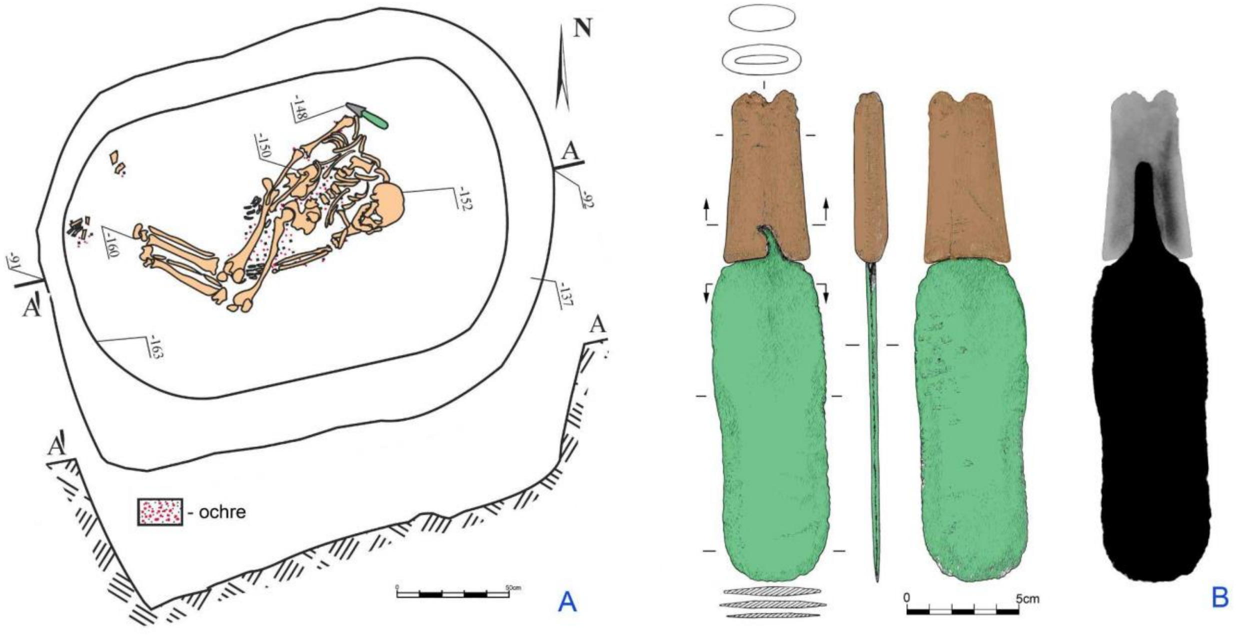

**Fig. 4.16. Leschevo-1, kurgan 1, burial 1 . A – plan of the burial, B – bronze knife.**

#### 4.11 Nizhnaya Orlyanka-1 site (Middle Volga forest-steppe)

*Summary by P. Kuznetsov*

The site is located at the edge of the floodplain terrace of the left bank of the Orlyanka River (a left tributary of the Volga River).

##### *4.11.1 Nizhnaya-Orlyanka-1, kurgan 1*

The diameter of the mound is 16 meters, with a height of 0.83 meters. The kurgan was excavated by Pavel Kuznetsov in 1992.

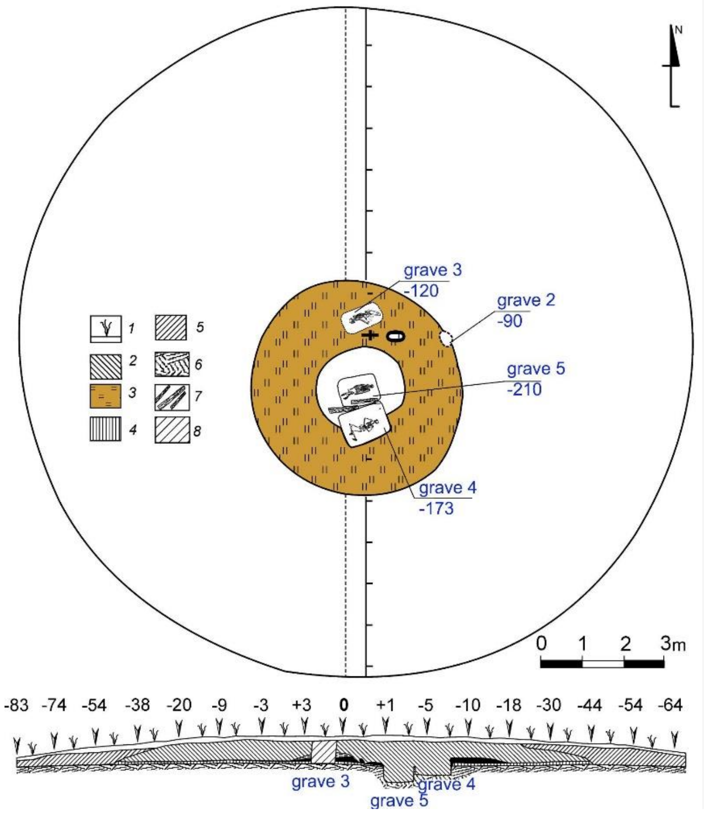

**Fig. 4.17. Nizhnaya-Orlyanka-1, kurgan 1. General plan of the mound.** 1 – plow layer, 2 – mound, 3 – ejecta from burial 5, 4 – buried soil, 5 – redeposited humic layer, 6 – subsoil, 7 – wooden desks covering burial 5, 8 – stratigraphic section (image by Pavel Kuznetsov).

###### *4.11.1.1 Burial 1:4 (individual ID I6727): 2831-2476 calBCE*

Burial 4 was located 1.9 m to the south of the centre of the mound. This is a secondary burial that cuts the ejecta from the grave pit of burial 5. Dimensions of the pit: 2.40 m x 1.78 m. The profile of the burial pit is shallow, with steep walls, and a depth of -173 cm from the modern top of the mound.

The buried male 35-45 years old lay on his right side, turned onto his back with his legs bent to the right, arms bent at the elbows, and fingers interlocked, with his head to the east-northeast. The forehead parts of the skull and feet are covered with ochre stains. At the knee of the right leg, there was a copper dagger with a pronounced, long handle.

Anthropologically, the skeleton is characterized as massive, sharply profiled, Europoid type.

There are two radiocarbon dates from this burial, 3022-2576 calBCE (4230±80, OxA 4255) and 2831-2476 calBCE (4055±20 BP, PSUAMS-4157). R-Combine fails, and we use the last one as the more precise one. Based on stratigraphic position, position of the skeleton on its right side, and the radiocarbon dating, the burial is attributed to Poltavka culture.

*4.11.1.2 Burial 1:5 (individual ID I6728): 3083-2916 calBCE*

Burial 5 was a primary burial in the kurgan and was located 0.83 meters south of its center. A circular ejecta of subsoil clay from the burial pit was discovered around the burial. Burial construction of the grave included robust wooden longitudinal covering of the burial pit. The dimensions of the pit were 2.00 × 1.55 meters, with vertical walls and a depth of up to 210 centimeters from the surface of the kurgan's center. The skeleton of a 40-45-year-old male was found lying on his back with bent legs, arms extended, and head facing east. The bones of the buried individual were densely covered with ochre, and plant bedding was traced beneath them. Alexander Khokhlov identified the skull as belonging to a hypermorphic, massive, paleo-European/proto-European type.

The is attributed to the early stage of the Yamnaya culture. Two radiocarbon dates were obtained from the human bones of the buried person, 3500-2900 calBCE (4510±75, OxA-4254) and 3076-2912 calBCE (4370±20 BP, PSUAMS-4544). A combination of them gives 3083-2916 calBCE as the most probable age of the burial.

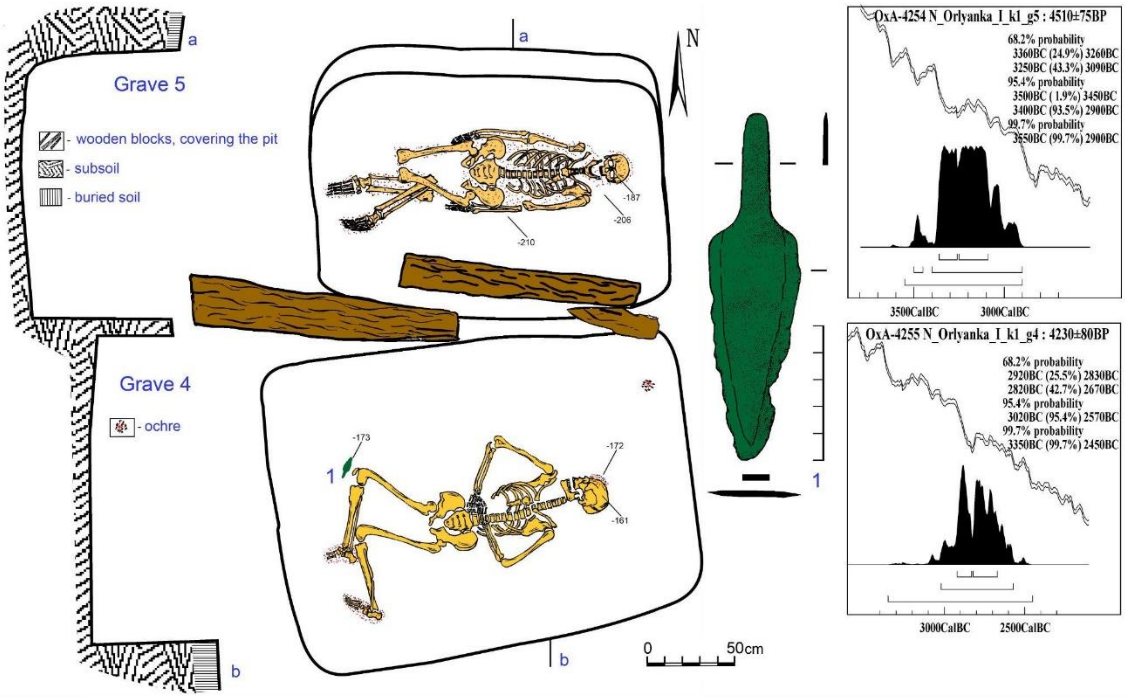

**Fig. 4.18. Nizhnaya-Orlyanka-1, kurgan 1, burials 4 and 5** (image by Pavel Kuznetsov).

###### ***4.11.2 Nizhnaya-Orlyanka-1, kurgan 4***

The diameter of the mound is 10 m, height is 0.15 m. The kurgan was excavated by Pavel Kuznetsov in 1992.

###### *4.11.2.1 Burial 4:2 (individual ID I6729): 3321-2931 calBCE*

Burial 2 was located in the center and was the primary one in the mound. Dimensions of the pit: 2.88 m x 1.56 m, with vertical walls., a depth of -210 cm from the modern top of the mound.

The buried male 18-25 years old was laid on his back with bent legs, and arms extended. Initially, the legs were bent at knees and stood vertically, with his head to the east-northeast. The forehead parts of the skull, pelvis, and feet are covered with ochre stains. A fragment of organic bedding was present at the bottom of the pit. No grave goods were found.

Alexander Khokhlov identified the buried person as Hypermorphous, massive, paleo-Europeanoid/proto-Europeanoid type.

Based on burial rite, specifically orientation and partial ochre coverage, the burial was attributed to the late stage of Yamnaya culture. However, two radiocarbon dates, 3495-2930 calBCE (4520±75, OxA-12573) and 3270-2920 calBCE (4425±20 BP, PSUAMS-4158), demonstrate the chronological position more typical for the early stage of the Yamnaya assemblage. The combination of the dates gives 3321-2931 calBCE [R-Combine: (4520±75, OxA-12573); (4425±20 BP, PSUAMS-4158)] as the most probable dating of burial 5.

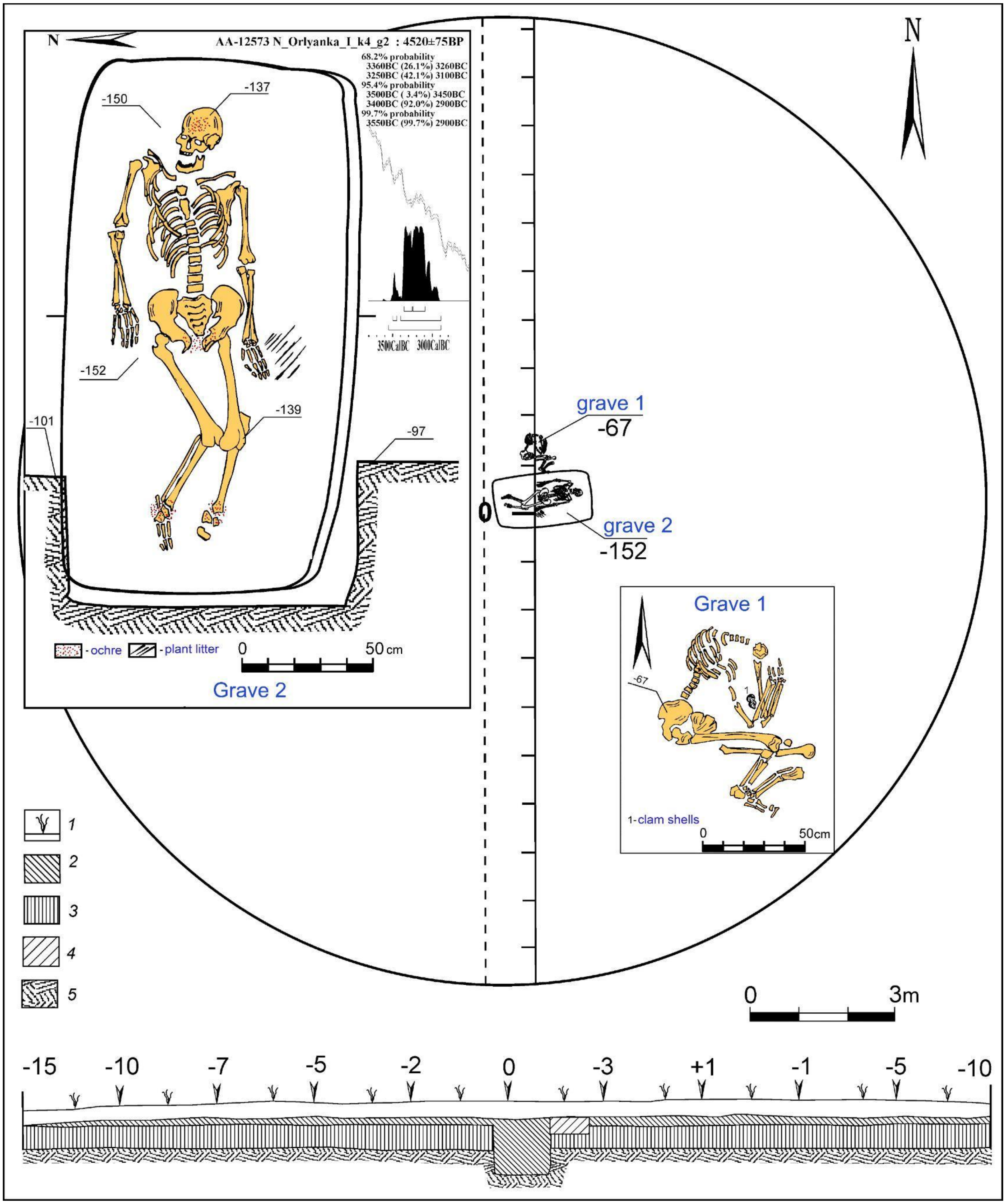

**Fig. 4.19. Nizhnaya-Orlyanka-1, kurgan 4, burial 2** (image by Pavel Kuznetsov).

#### 4.12 Orlovka-1 site (Middle Volga forest-steppe)

*Summary by P. Kuznetsov*

The site is located at the edge of the first floodplain terrace of the left bank of the Chagra River, a left tributary of the Volga River.

##### *4.12.1 Orlovka-1, kurgan 2*

The diameter of the mound is 59 m, height is 1.09 m.

###### *4.12.1.1 Burial 2:2 (individual ID I6297): 3022-2907 calBCE*

Burial 2 is located in the centre of the mound and interpreted as the primary grave in the kurgan. The dimensions of the grave pit are 2.25 m x 1.25 m. The profile of the burial pit is shallow, with steep walls, and a depth of 124 cm from the modern top of the mound.

At the bottom of the grave pit, a female skeleton of unidentified age was discovered. It was laid on her back, with the legs slightly bent at knees, and arms extended along the body, with her head to the northeast. The skull and mandibula were resting on the vertebra. Ochre covers the skull, upper body, arms, legs, and feet.

Between the long northern wall of the pit and the skull of the buried individual, a bronze knife, an awl (a stilet-like item), and flakes of flintified limestone were found (Ovchinnikova and Fadeev 2007).

The burial is attributed to the late stage of the Yamnaya culture of the Volga-Ural region, based on orientation, and partial ochre staining. The radiocarbon dates the burial to 3022-2907 calBCE (4355±20 BP, PSUAMS-2904).

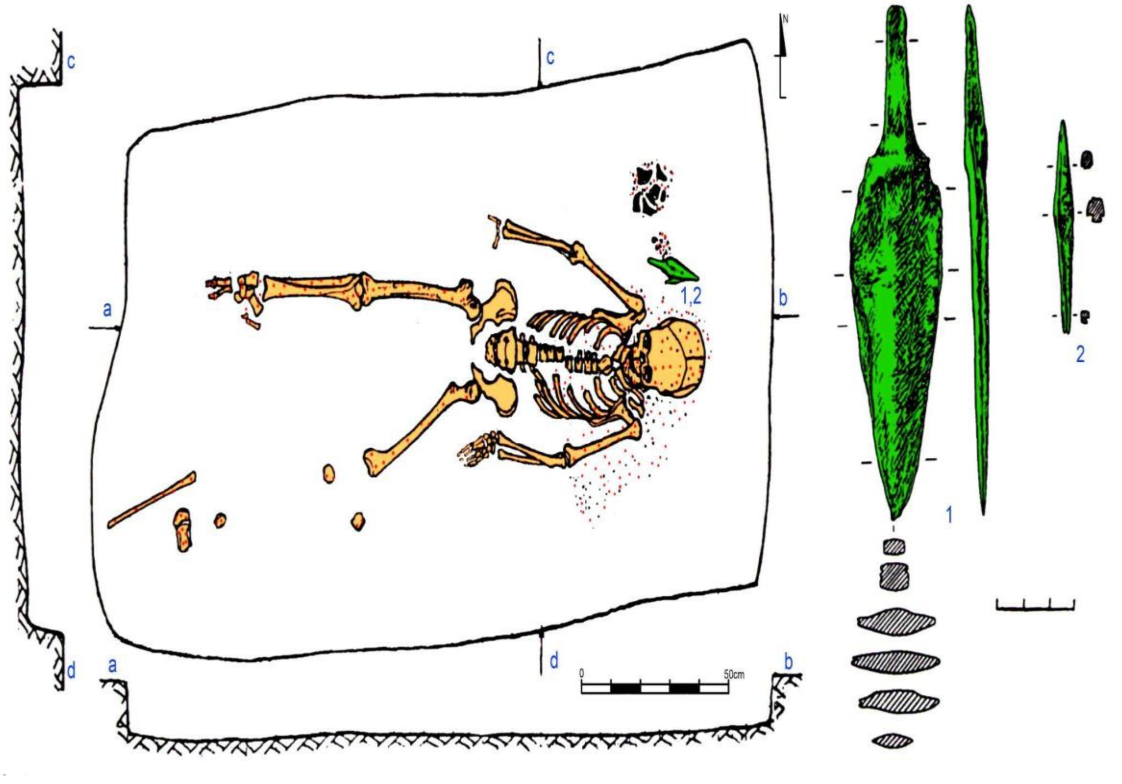

**Fig. 4.20. Orlovka-1, kurgan 2, burial 2** (image by Pavel Kuznetsov).

#### 4.13 Podlesny-1 site (Middle Volga steppe)

*Summary by P. Kuznetsov*

The site is located on the edge of the floodplain terrace of the left bank of the Samara River, a left tributary of the Volga River, on the northwest outskirts of the village of Podlesny. Podlesny-1 is one of eight Bronze Age kurgan groups investigated along a 12 km stretch of the floodplain terrace between the villages of Podlesny and Spiridonovka.

##### *4.13.1 Podlesny-1, kurgan 3*

The diameter of the mound is 24 m, height is 0.35 m.

###### *4.13.1.1 Burial 3:3 (individual ID I1450): 3333-3028 calBCE*

Burial 3 is located in the centre of the mound and is the primary one. The pit was covered with planks placed along the long walls. Dimensions of the pit: 1.9 m x 1.2 m. The walls of the burial pit are vertical, depth of -156 cm from the modern top of the mound.

At the bottom of the burial pit, the skeleton of a 45-55-year-old male was discovered lying on his back and oriented with his head to the east. The legs of the buried individual were stretched out, and at the bottom of the burial pit, beneath the skeleton, ochre was found scattered. Ochre covered the upper part of the body and legs of the buried individual. No artefacts were found.

According to the determination of Aleksandr Khokhlov, the skull is Europoid, dolichocephalic. The skeleton bones are large and robust. The physical development was strong, and height of the buried individual reached approximately 175.1 cm (Barynkin et al., 2006).

The burial is identified as belonging to the early phase of the Yamnaya culture in the Volga-Ural region, characterized by its orientation, extensive ochre staining, and extended posture. Radiocarbon dating places this burial among the oldest dated Yamnaya graves, dating back to approximately 3333-3028 calBCE (4465±20 BP, PSUAMS-4412).

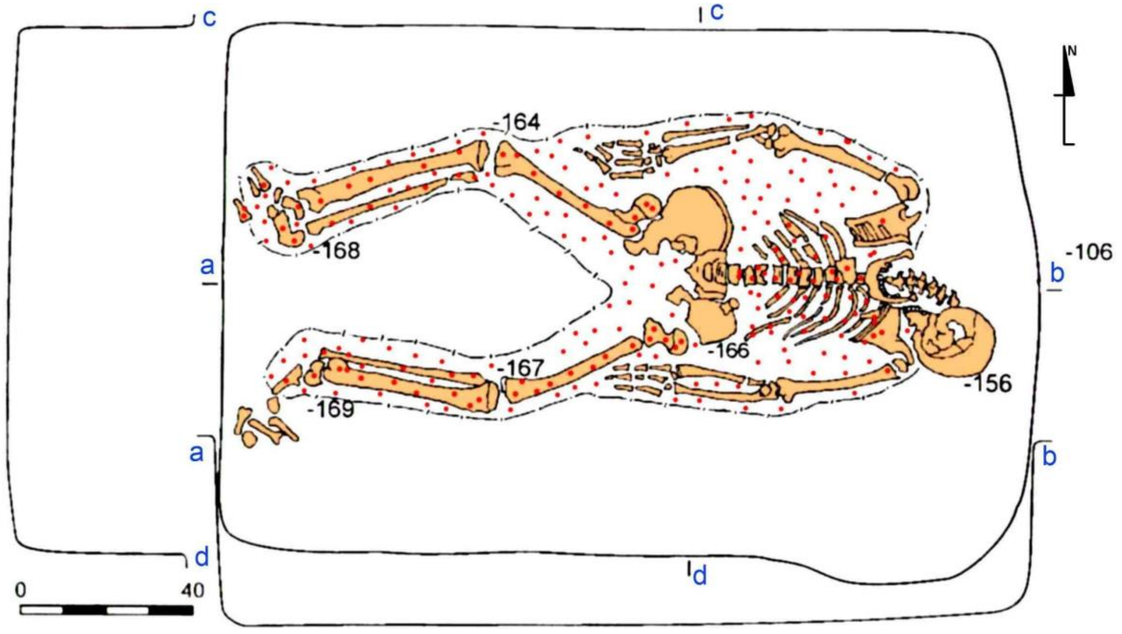

**Fig. 4.21. Podlesny-1, kurgan 3, burial 3** (image by Pavel Kuznetsov).

#### 4.13 Poplavskoe-1 site (Middle Volga steppe)

The settlement of Poplavskoye is situated on the southern shore of Lake Lebyazhye, approximately 2.5 kilometers south of the village of Krasnosamarskoye in the Kinel district. During summer seasons of 1992-1994, Yuri Kolev excavated the site and unearthed a late Bronze Age structure as well as a burial complex dated to the Early Bronze Age.

##### *4.13.1 Poplavskoe-1, burial 1, skeleton 3 (individual ID I7677): 2911-2881 calBCE*

Burial 1 situated at the presumed northwest periphery of a Bronze Age settlement, where the cultural layer was partially disturbed in course of construction of a drilling machine. It was identified on the subsoil level, at a depth of 1.03 meter from the ground surface.  The grave pit had a rectangular shape with rounded corners, its dimensions at the upper lever reached 250x185 cm. The long sides of the burial pit are oriented along the northwest-southeast axis. The burial contained three skeletons, from which skeleton 3 (individual ID I7677) was the primary one and two other were placed into the grave pit later.

<u>Burial 1, skeleton 3 (individual ID I7677)</u>. The skeleton was discovered lying on the bottom of the pit, on its back with the head oriented to the north-northeast. The bones of the deceased were preserved in anatomical order. The legs, bent at the knees, were laid on the right side. The arms, bent at the elbows, were laid with the hands on the clavicles. There were no traces of ocher found in the grave.

The grave is attributed to the late stage of Yamnaya culture and dated to 2911-2881 calBCE (4270±20 BP, PSUAMS-4276).

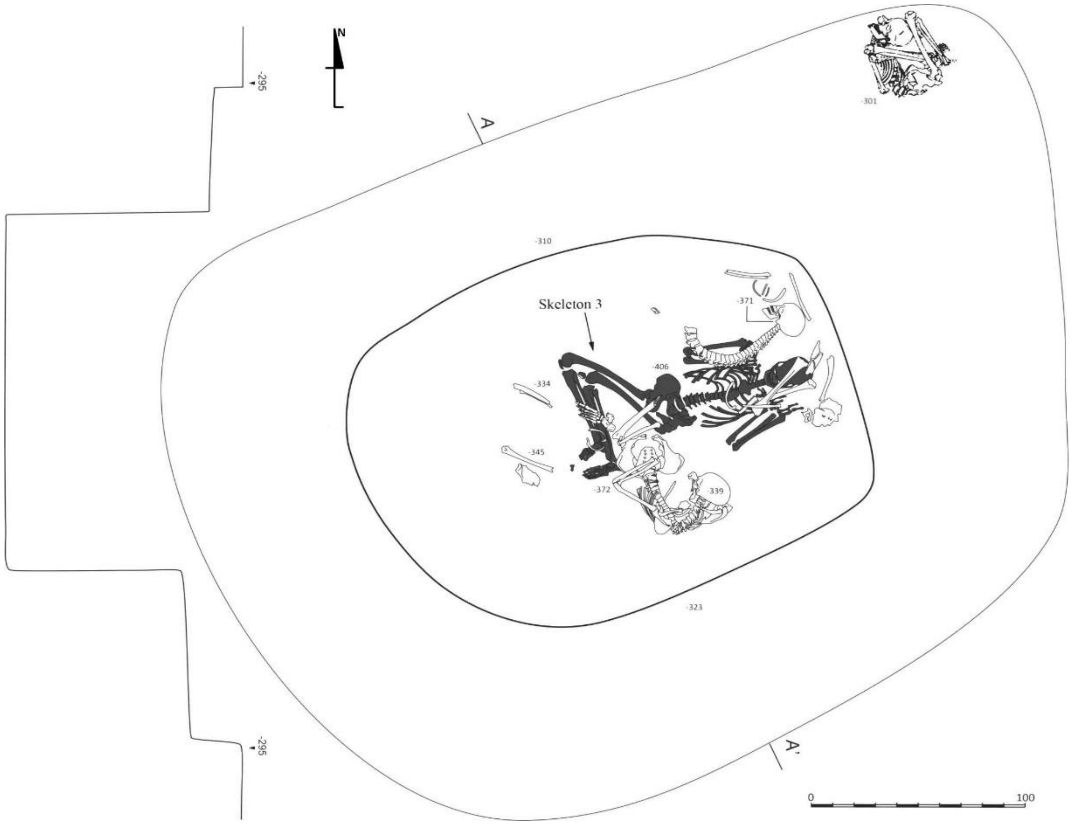

**Fig. 4.22. Poplavskoe-1 occupation site, burial 1** (image by Pavel Kuznetsov).

#### 4.14 Utevka-9 site (Middle Volga steppe)

*Summary by P. Kuznetsov*

The site is located at the edge of the floodplain terrace of the left bank of the Samara River, a left tributary of the Volga River, on the southeast outskirts of the village of Utevka, and it is crossed by the Utevka stream (a left tributary of the Samara River). Within the Utevka valley (10 km x 5 km), at least 11 burial mounds have been identified. The total number of mounds within the Utevka Valley reaches one hundred. This is the highest concentration of mounds in the entire Samara River basin, including the largest mounds in the Middle Volga (Utevka-1 kurgan group includes mounds from 80 m to 110 m in diameter).

The site was discovered in 1998 by Oleg Mochalov and A.V. Plaksin, and in 2008 S.V. Bogdanov (Orenburg) and N.V. Lebedeva (Samara) excavated kurgan 4 of Utevka-9.

##### *4.14.1 Utevka-9, kurgan 4*

The diameter of the mound is 40 m, height 0.5 m.

###### *4.14.1.1 Burial 4:1 (individual IDs I10362 and I10363): 2877-2630 calBCE*

Burial 1 was located in the centre of the mound. The dimensions of the deep (the exact depth is not specified) pit reach 2.4 m x 1.75 m, the walls are steep. The burial contained two skeletons.

Skeleton 1 (individual ID I10362), anthropologically and genetically identified as female, was laid in the supine position with her head oriented to the east. The spine was slightly curved. The right arm was extended along the torso, with the radius bone displaced and its end broken. The hand was under the right thigh bone, which was not lying straight but was shifted to the side. The left arm was lying on the pelvic bones, with the hand and finger phalanges shifted in different directions. A fragment of the upper part of the femur bone was found near the left hip bone. To the west, in a spot of a large burrow, a fragment of the tibia bone was found. The soil around the skull and skeleton was stained with dark red ochre.

Twenty centimetres north of the right hand, a flint flake was discovered, and fifteen centimetres north of the right elbow, a flint blade was found.

Aleksandr Khokhlov determined the cranial as Europoid. Some teeth were lost during life; strong dental calculus development is evident in individual teeth; the nose was broken during life, with the nasal bones damaged by a direct blow from an opponent, slightly from the side and to the right (a common occurrence for this time period), suggestive of a hook-like blow. The mortal age of the deceased was ≥55 yrs.

Skeleton 2 (individual ID I10363) anthropologically and genetically identified as female, was unearthed in the area disturbed by two large burrows. The skull, without the mandibula, was lying on the left side. The legs were bent at the knees. The bones were arranged as if the buried individual was standing on their knees in a pose of adoration. Near the skull, knees, and ankles of the buried man, there were patches of ochre. The legs of the buried individual were directly above the skull of skeleton 1. Thus, burial of skeleton 2 was positioned above skeleton 1, and must me relatively later than skeleton 1. Presumably, the buried individual was oriented with their head to the northeast (Kuznetsov et al., 2011, 2015).

Aleksandr Khokhlov determined the cranial as Europoid and noticed the presence of a metopic suture. The postcranial part mainly consists of bones from the lower extremities. They appear small, moderately long, and proportional (left femur length: 79.2). There is flattening of the upper third of the femoral shaft (hyperplatymeria - 67.2 and platymeria - 75.0). The femurs exhibit mesokneeism (68.7 and 67.2). Height ranges from 157 to 162 cm – average to below average. Shoulder width (left clavicle 132.0?) – approximately 31.5 cm (Razhev, 2003) – small. No pathological changes were observed.

Burial 1, kurgan 4 at Utevka-9 is attributed to Yamnaya culture, based on position, orientation, and ochre staining. The burial yields two radiocarbon dates.

The C14 dating of skeleton 1 is 2877-2630 calBCE (4155±30 BP, PSUAMS-10732), and the C14 dating of skeleton 2 is 3352-3101 calBCE (4510±25 BP, PSUAMS-10733). Stratigraphic relation between the skeletons makes the earlier date 3352-3101 calBCE (4510±25 BP, PSUAMS-10733) for skeleton 2 irrelevant and supposes presence of FRE in this case.

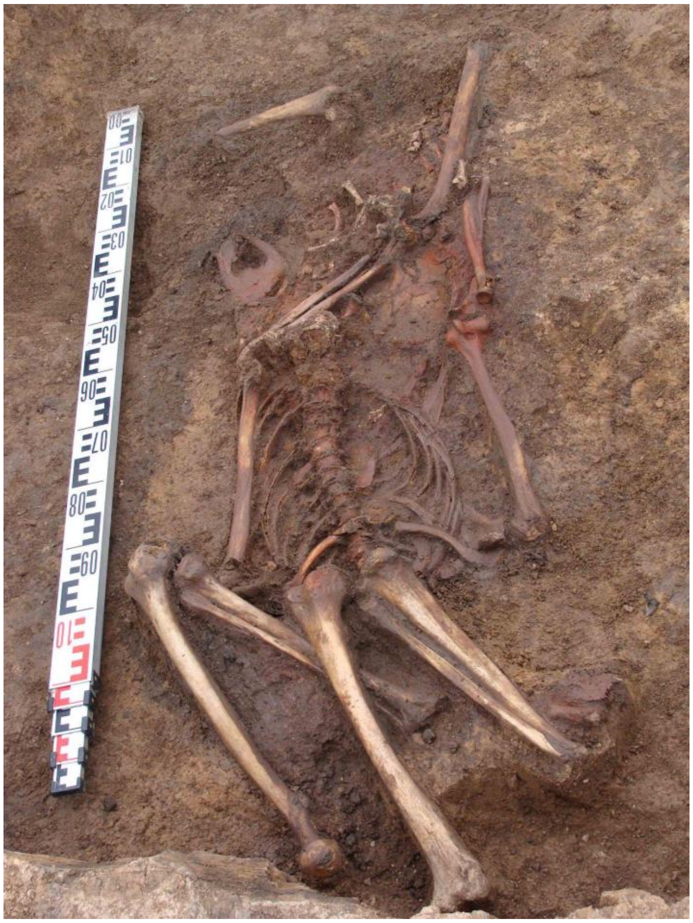

**Fig. 4.23. Utevka-9, kurgan 4, burial 1, skeletons 1 and 2 (image by Pavel Kuznetsov).**

*Bibliography to section 4:*

*Barynkin, P.P., Zudina, V.N., Kramarev, A.I., Salugina, N.P., Tsibin, V.A., & Khokhlov, A.A. (2006). Study of the Bronze Age barrows near the village of Podlesny on the Samara River. In Archaeological Issues of the Volga Region (in Memory of I.B. Vasilyev) (Vol. 4, pp. 293-313). Scientific and Technical Center Publishing, Samara.*

*Bogdanov, S.V. (1999). Kurgans of the Early Bronze Age in the Vicinity of the Village of Kurmanayevka. In Archaeological Sites of the Orenburg Region, Issue 3: 12-19. Orenburg: OGPU Publishing House.*

*Khokhlov, A.A., & Grigoryev, A.P. (2019). Morphological characteristics of the anthropological sample from the Krasikovsky I burial mound. In Archaeological Sites of the Orenburg Region, Issue 14: 69-81). Orenburg: OGPU Publishing House.*

*Kuznetsov P.F., Mochalov O.D., Khokhlov A.A., Anthony D. (2018). Grachevka Kurgans. Archaeology, Anthropology, Genome Analysis. Samara: SGSPU.*

*Kuznetsov, P.F. (1990). Unique Burial of the Early Bronze Age on the Kutuluk River. In Antiquities of the East European Forest-Steppe. Kuibyshev: KGPI Publishing House: 135-137.*

*Kuznetsov, P.F., & Mochalov, O.D. (2014a). Features of Late Yamnaya population kurgans in the Samara Valley during the Early Bronze Age. In Archaeological Sites of the Orenburg Region, Issue 11: 90-95. Orenburg: OGPU Publishing House.*

*Kuznetsov, P.F., & Mochalov, O.D. (2014b). A special type of metal artifacts of the Yamnaya culture (based on the results of work in the Samara Valley). Proceedings of the Samara Scientific Center of the Russian Academy of Sciences, 16(3), 291–297.*

*Kuznetsov, P.F., Mochalov, O.D., Myshkin, V.N., & Khokhlov, A.A. (2015). Study of the Utevka-9 burial ground in the Samara Zavolzhye. Archaeological Issues of the Volga Region (Vol. 5, pp. 52-60). Samara.*

*Kuznetsov, P.F., Myshkin, V.N., & Khokhlov, A.A. (2023). Lopatino-2 kurgan group. In Questions of Archaeology of the Volga Region (Vol. 10, pp. 156-171). Samara: Samara State Pedagogical University.*

*Kuznetsov, P.F., Popova, L.M., Mochalov, O.D., Myshkin, V.N., & Pastukhova, O.S. (2011). Investigation of barrows and settlements in the Samara region. Archaeological Discoveries of 2008. Moscow, pp. 371-373.*

*Morgunova, N.L. (ed.). Bogolyubovskiy Kurgan Cemetery of the Timber Grave Culture in the Orenburg Region. Orenburg : OGPU Publishing House, 2014.*

*Morgunova, N.L., Evgenyev, A.A., Kryukova, E.A., Kharlamov, P.V., Faizullin, A.A., Golieva, A.A. (2019). Krasikovo-1 Bronze Age Barrow Cemetery in the Orenburg Region. In Archaeological Sites of the Orenburg Region, Issue 14: 04-39. Orenburg: OGPU Publishing House.*

*Morgunova, N.L., Kul'kova, M.A. (2019). Results of Radiocarbon Dating of the Krasikovo-1 Barrow Cemetery. In Archaeological Sites of the Orenburg Region, Issue 14: 39-45. Orenburg: OGPU Publishing House.*

*Ovchinnikova, N.V., & Fadeev, V.G. (2007). Results of the study of the Yamnaya culture burial mound Orlovka I. In Samara Region in the History of Russia (Vol. 3, pp. 24-34). Samara: SOIKM named after P.V. Alabin.*

### 5. EARLY BRONZE AGE OF THE VOLGA-DON STEPPES

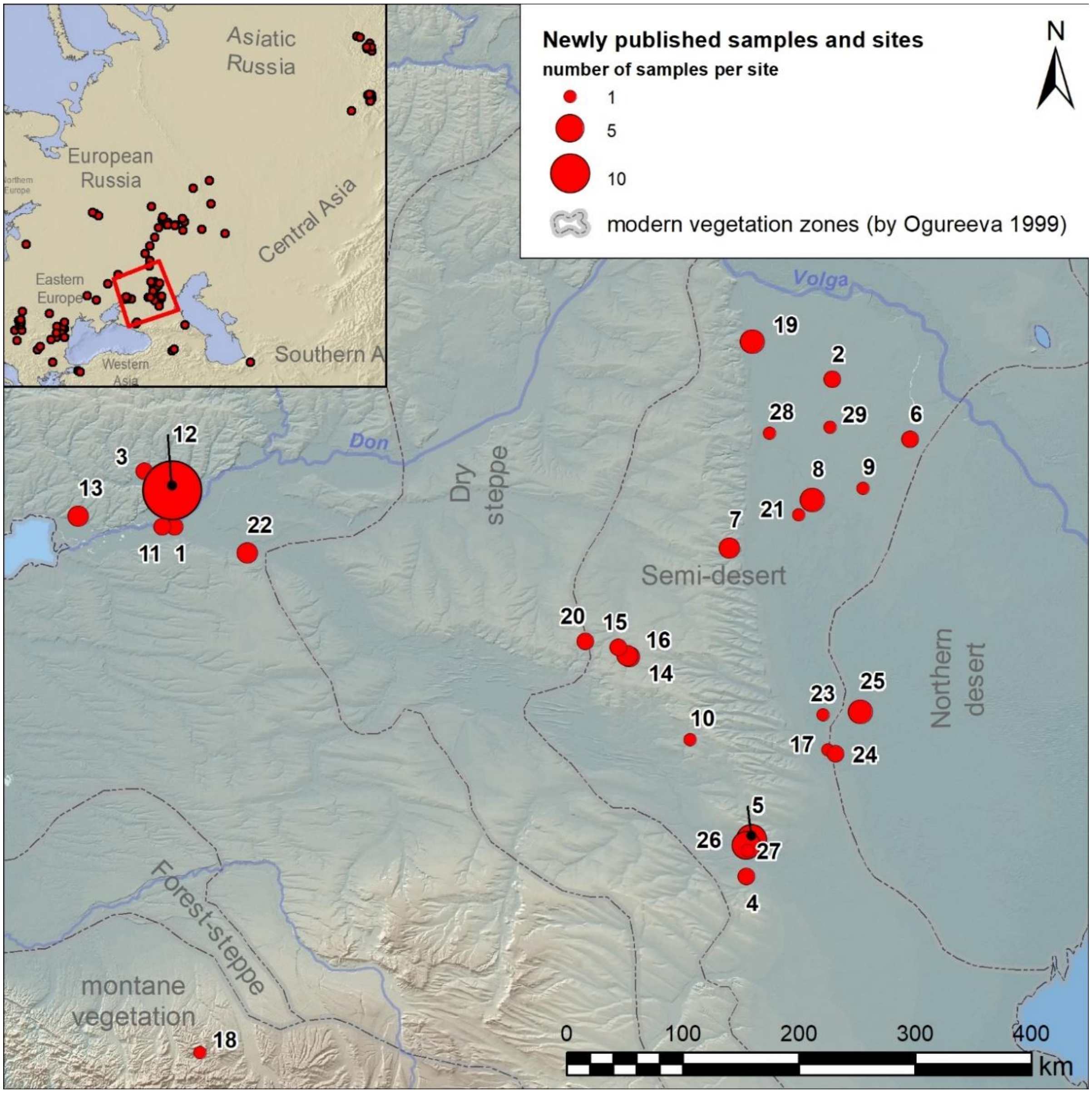

**Fig. 5.1. Newly published samples from the Lower Don region, the Volga-Don interfluves and the Caucasus Piedmont (Russia).** 1 - Arpachin (Rostov Oblast, Bagayevsky District); 2 - Balkin (Republic of Kalmykia, Maloderbetovsky District); 3 - Cherkasov-4 (Rostov Oblast, Aksaysky District); 4 - Chograi-1 and 2 (Stavropol Krai, Arzgirsky District); 5 - Chograi-4 and 5 (Republic of Kalmykia, Iki-Burulsky District); 6 - Dzhangar-Kermen-Tolga (Republic of Kalmykia, Yustinsky District); 7 - Ergeninskiy-1 (Republic of Kalmykia, Ketchenerovsky District); 8 - Evdyk-1 (Republic of Kalmykia, Ketchenerovsky District); 9 - Idzhil-2 (Republic of Kalmykia, Oktyabrsky District); 10 - Khar-Zukha-2 (Republic of Kalmykia, Priyutnensky District); 11 - Krestovy (Rostov Oblast, Aksaysky District); 12 - Krivyansky-7 and 9 (Rostov Oblast, Oktyabrsky District); 13 - Mokro-Chaltyrsky-1 and Chaltyrsky-11 (Rostov Oblast, Myasnikovsky District); 14 - Peschany-4 and 5, 15 - Sukhaya-Termista-1 and 2, 16 - Temrta-4 (Rostov Oblast, Remontnensky District); 17 - Tsagan-Usn-5 (Republic of Kalmykia, Yashkulsky District); 18 - Tsarskaya (Republic of Adygea, Maykopsky District); 19 - Tsatsa (Volgograd Oblast, Svetloyarsky District); 20 - Ulan-4 (Rostov Oblast, Remontnensky District); 21 - Ulan-Tolga (Republic of Kalmykia, Ketchenerovsky District); 22 - Vesely (Rostov Oblast, Vesyolovsky District); 23 - Volga-Chogray-Channel-37, 24 – 53, 25 – 56 (Republic of Kalmykia, Yashkulsky District); 26 - Vostochny-Manych-2 and 3; 27 - Vostochny-Manych-right-bank-1 (Republic of Kalmykia, Iki-Burulsky District); 28 - Zakhanata (Republic of Kalmykia, Sarpinsky District); 29 - Zergenta-2 (Republic of Kalmykia, Oktyabrsky District).

#### 5.1 Peschany-4 site

*Summary by N. Shishlina*

The Kurgan burial site Peschany-IV, excavated by Natalia Shishlina in 2015 (kurgan 13) and N.V. Leonova in 2019 (kurgan 17), is situated 6.1 km southeast of the village of Remontnoye in the Remontnensky District of the Rostov Region, Russia. It is located on a small elevated plateau between the Dzhurak-sal River and the Peschanyaya Balka, at the northern border of the Kumo-Manych Depression. Geographically, the area represents the western extremity of the Yergeni, characterized by a developed network of ravines and gullies. The steppes are heavily drained by the valleys of small rivers. During dry years, the valleys of the intermittent rivers can be quite extensive, featuring terraces and wide floodplain areas, sometimes reaching up to a kilometre in width. The predominant vegetation type occupying much of the Yergeni upland consists of complex Artemisia-Stipoid steppes. On relatively elevated microrelief areas, there are clusters of grasses, while areas with solonchaks feature Artemisia and saline vegetation.

The burial site comprises 26 kurgans grouped into several clusters, extending in a chain along a northwest-southeast axis.

The phases of the site, their chronology, and main features are as follows: (1) Eneolithic (3500-3300 BCE); (2) Yamnaya culture (2900-2600 BCE); (3) Early Catacomb culture (2800-2600 BCE); (4) East/West Manych Catacomb culture (2550-2200 BCE); (5) Early Iron Age (800 BCE to 100 CE); and (6) Medieval period (1170-1260 CE).

Radiocarbon dating of Yamnaya burials at Peschany-4 determined on human bone, sheep bone, horse bone, willow wood, plant litter, and charcoal site yielded 27 dates, indicating a timeframe from 2900 to 2600 BCE.

##### *5.1.1 Peschany-4, kurgan 13*

The height of the mound ranges from 150 to 184 cm, with a diameter of 28-31 m. It contains 6 burials. Burial 6 serves as the primary one for mound 1, and it was overlaid by burial 5, attributed to the Yamnaya culture.

###### *5.1.1.1 Burial 13:6 (individual ID I29558)*

The burial construction consisted of a rectangular pit with rounded corners, oriented along a northwest-southeast axis, with sloping walls. The dimensions of the pit at the top were 136 × 93 cm, and at the bottom were 148 × 107 cm. Initially, the walls of the pit were up to 150-160 cm high. A wooden structure was erected over the grave, possibly supported by rounded wooden posts.

At the bottom of the burial pit, at a depth of -310 cm, lay the skeleton of a child, positioned on its back, with the skull oriented to the south. The position of the head suggests that a pillow lay beneath it. Initially, the legs were bent at the knees, later spreading in a diamond shape. The foot bones also fell to the side.

Nine centimeters from the left elbow at the bottom of the burial pit lay four sheep astragali without traces of processing.

The skeleton rested on a plant bedding measuring 102 × 74 cm. On the skeleton, covering the leg bones, lay decayed material in which twigs or fibers could be discerned, arranged perpendicular to the spinal column. This construction may have been supported by wooden posts but later collapsed onto the skeleton. Rounded patches of red pigment were observed near the foot bones, and the elbow of the right arm was stained with pigment.

According to A.A. Kazarnitsky, a fragmented skull and postcranial skeleton of a child were found in the burial. The level of dental development corresponds to an age of around 6-7 years. There is a small through-hole (outer diameter 2-3 mm, inner diameter about 1 mm) with porosity on the external surface in the center of the left temporal bone. Also present are all long tubular bones of the upper and lower limbs (femur length - 243 mm, tibia - 205 mm, humerus - 174 mm), scapulae, clavicles, ribs, pelvic bones, fragments of several vertebrae from all sections, hand bones, and foot bone (Shishlina et al., 2023).

Dietary data that may influence radiocarbon dating results or reflect migration processes include:

$\delta 13C = -16.5 \pm 0.25‰$

$\delta 15N = 16.6 \pm 0.3‰$

Radiocarbon dates:

Wood: GrA64624, 4145 ± 35, 2878–2585 calBC (δ13C –26.1‰)

Human bone: GrA64622, 4130 ± 35, 2871–2579 calBC

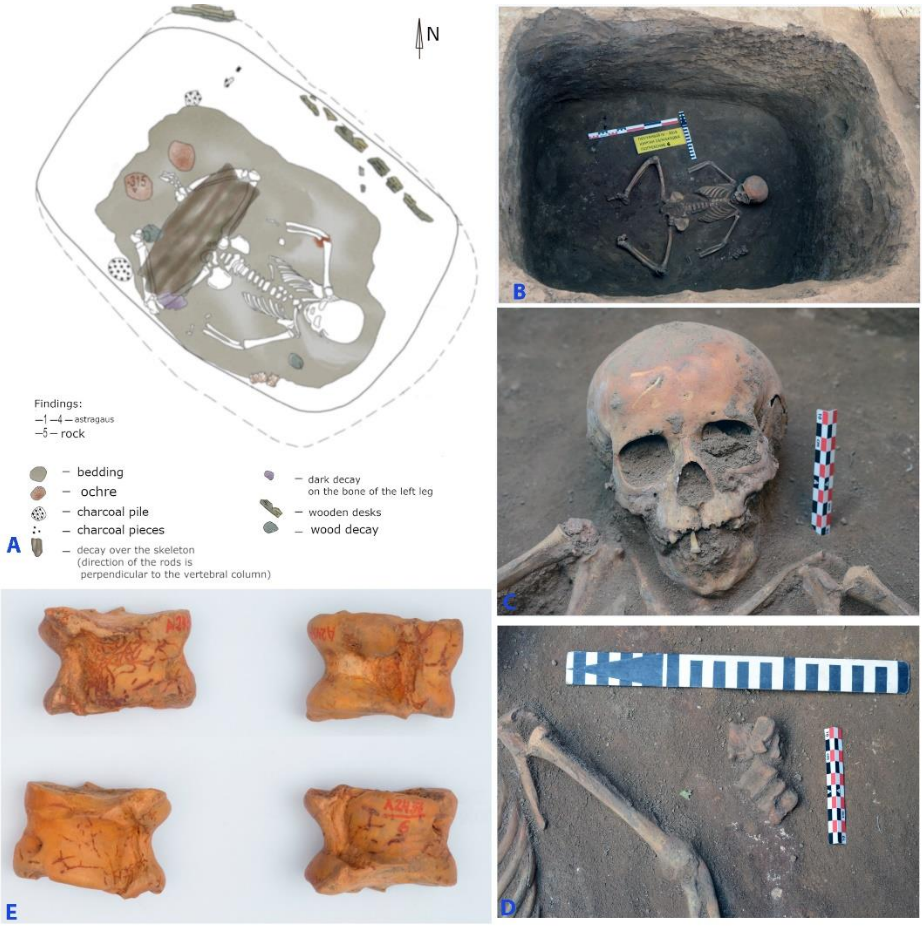

**Fig. 5.2. Peschany-4, kurgan 13, burial 6.** A and B – plan and photo of the burial, C and D – details, E – astragali from the grave.

#### 5.2 Peschany-5 site

*Summary by N. Shishlina*

The kurgan group Peschany-5 is located in the Remontnensky district of the Rostov region, Russia, on a small elevated plateau between the Jurak-Sal River and the Peschanya Balka gully. It consisted of a chain of 65 mounds constructed along the northwest-southeast line on the watershed summit.

##### *5.2.1 Peschany-5, kurgan 5*

The kurgan was excavated by Natalia Shishlina in 2007. In total, six burials have been discovered in the mound, spanning three phases of the monument's existence. The earliest phase, Phase 1, is attributed to the Early Catacomb culture; Phase 2 corresponds to the final stage of the Early Catacomb culture; Phase 3 is associated with the Eastern Manych Catacomb culture.

The main mound of mound 5 was constructed over a burial from the early Catacomb culture (Phase 1). Burials 3 and 4, which belong to Phase 2, were accompanied by infilling of the mound. Burials 5 and 6 are attributed to Phase 3. The most recent burial in the mound is burial 1, the cultural affiliation of which remains undetermined.

###### *5.2.1.1 Burial 5:5 (individual ID I29573)*

Burial 5 is a secondary burial incised into the mound of phase 2, was located in the northwest part of the mound.

The burial construction is a T-shaped catacomb (the axes of the entrance pit and the chamber are perpendicular). The oval entrance pit had dimensions of 128 x 116 cm at the top edge, narrowing towards the bottom. The bottom of the entrance pit descended into the entrance to the chamber with two segmental steps. The entrance was located in the northern wall, had an arched shape, with a width of 90 cm at the base and a height of 75 cm. To the north of the entrance pit was the chamber, the dimensions of which at the bottom were 180 x 120 cm.

At the bottom of the chamber lay the skeletons of two adult individuals, arranged in a non-anatomical order, "package" style, along the east-west axis. At the eastern edge of the package lay the skull of a 50-60-year-old man (individual ID I29573) on the left parietal and temporal bones with a collapse on the facial part, the lower jaw was located next to the skull, disjointed. Postcranial skeleton bones belong to two individuals, some of them with characteristic signs of age-related growths. The long bones were laid almost parallel to each other along the southwest-northeast line. A fragment of a bone artifact in the form of a rod with a sharpened end, 10.4 cm long, was found near the femur bone.

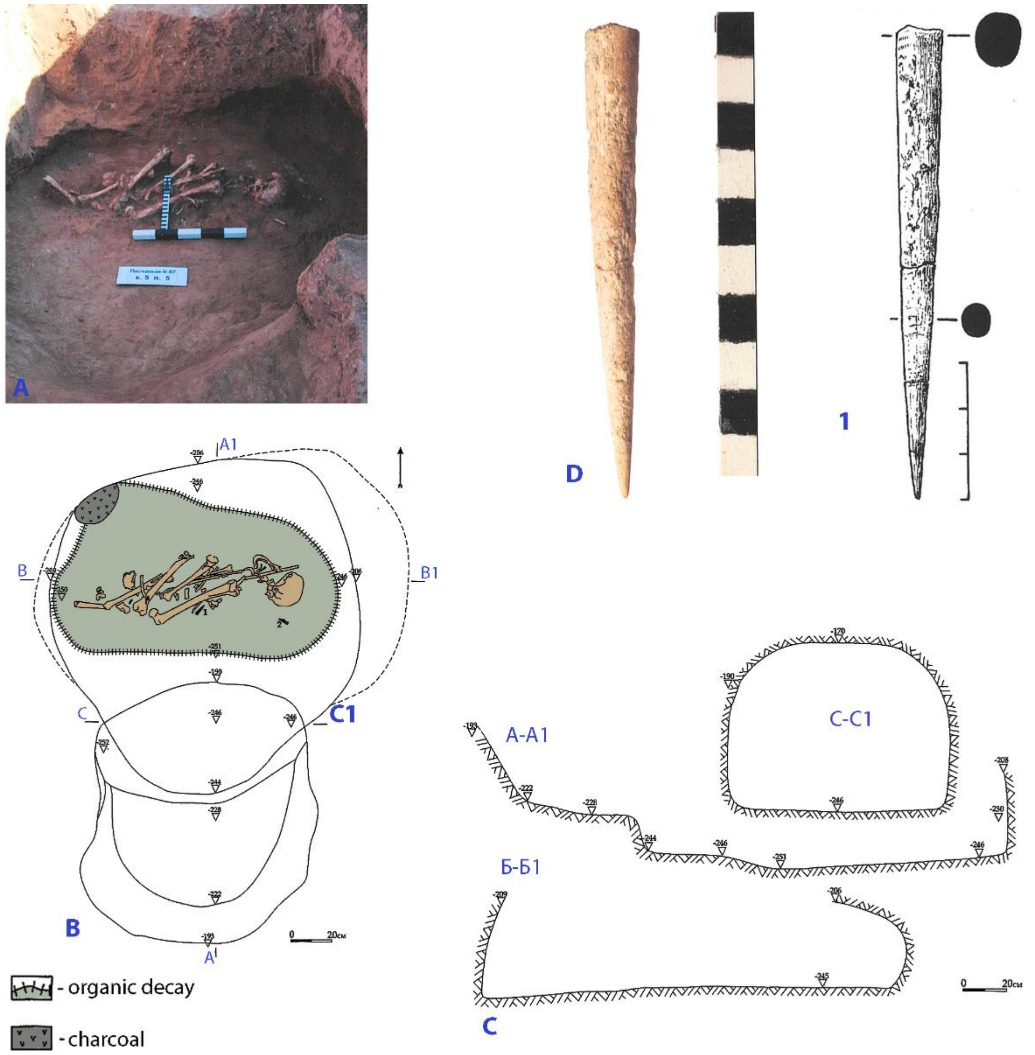

**Fig. 5.3. Peschany-5, kurgan 5, burial 5.** A – photo of the burial, B and C – plan and profile of the burial, D – bone item (image by Natalia Shishlina).

#### 5.3 Sukhaya Termista-1

*Summary by N. Shishlina*

The kurgan group of Sukhaya Termista-I is located on the northeastern outskirts of the village of Remontnoye in the Remontnensky District of the Rostov Region, on the left bank of one of the marshy channels of the Djurak-Sal River, on the second terrace of the Sukhaya Termista ravine. The burial mound group consisted of two mounds, stretched along the northwest-southeast line.

##### *5.3.1 Sukhaya Termista-1, kurgan 1*

In the burial mound, 16 burials were discovered. The main mound was constructed over the late Eneolithic burial 14, the mound of which was likely extremely small and not clearly visible in the mound profiles. However, adjacent to this burial, the main mound was built over the Yamnaya burial 13, into which the Yamnaya burial 11 was soon inserted. The height of the main Yamnaya burial mound was approximately 60-70 cm, with a diameter of 22 m. A ditch was constructed around the ancient mound. A substantial western backfill, associated with two child burials of the Eastern Manych Catacomb culture, covered the ancient mound and the already filled-in ditch. The third stratigraphic horizon is associated with burials from the Late Middle Bronze Age to the Late Bronze Age. In the Early Iron Age, three burials were inserted, and one more burial was added during the Polovtsian period.

###### *5.3.1.1 Burial 1:11 (individual ID I29571|RISE240)*

Burial 11 was incised into mound phase 1 and covered by mound 3, being a primary burial for the last one.

It was made in a rectangular pit with straight walls measuring 160 x 132 cm at the top edge, 172 x 115 cm at the bottom, initial wall height of 240 cm. At the bottom of the pit along the northern wall lay a female skeleton in a contracted position, with the skull oriented to the east. The position of the skull indicates that a headrest was placed under the head, the legs were raised upwards, preserved in their original position, leaning against the northern wall, with the soles of the feet on the ground.

A hammer-headed bone pin was found next to the left hand.

A two-layered vegetal litter was traced under the skeleton; a layer of ochre up to 5 mm thick was fixed on the pelvic bones, in the abdominal area, near the foot bones, and on the skull.

A.A. Kazarnitskiy observed traces of weak temporal deformation on the skull and reported the following craniological characteristics. The cranial vault is very high and wide with medium length, hyper-brachycranial according to the cranial index and gypsycranial (relatively high) according to the height-length index; broad forehead; high and wide face, orthognathic; horizontal facial profiling is moderate at the upper level and very sharp at the zygomatic points level; orbits are moderately wide and low; nose of average height, wide, sharply projecting; nasal bridge high; canine fossa of medium depth. Body length according to the formula of M. Trotter and G. Gleser is 162 cm, with the consideration of the length of the forearm bones, 165 cm.

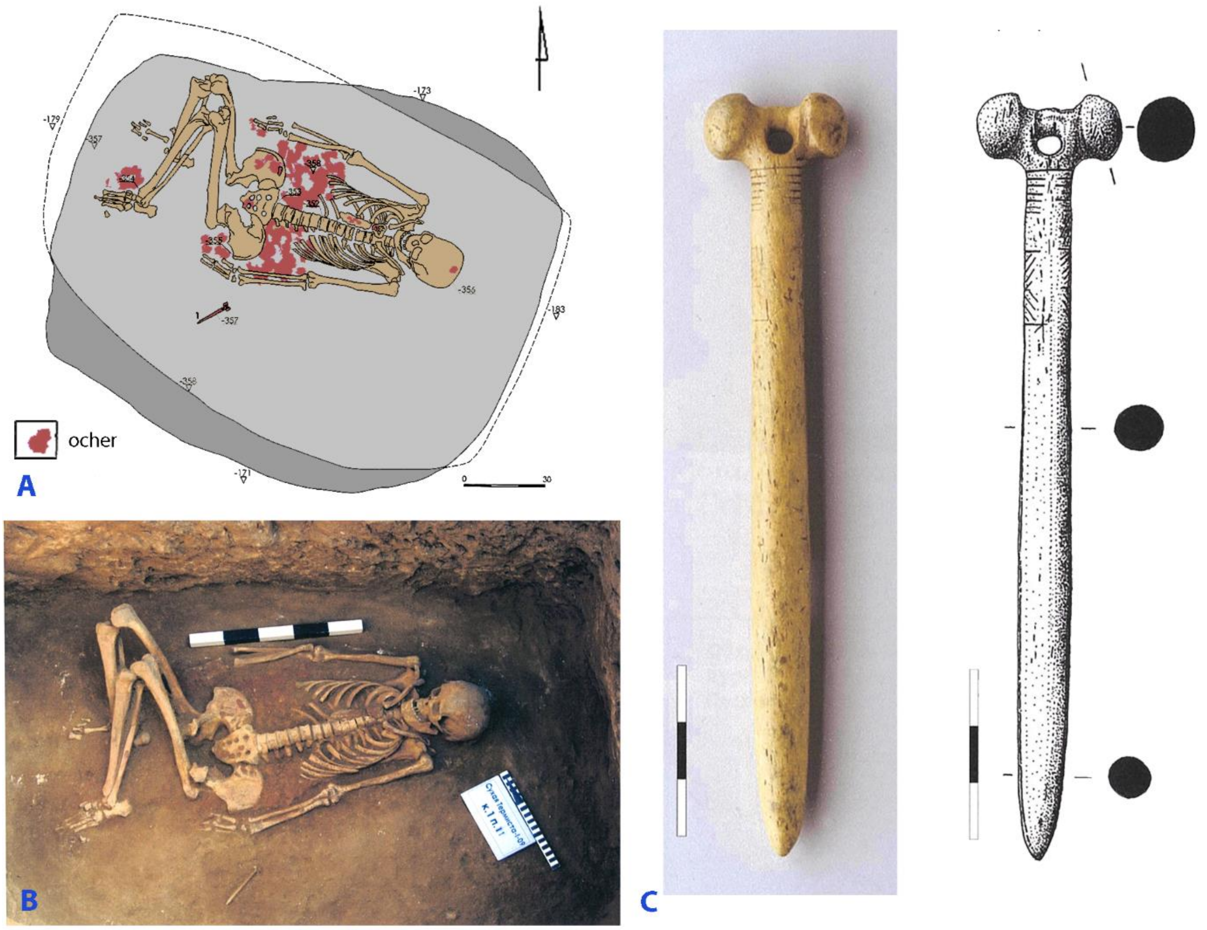

**Fig. 5.4. Sukhaya Termista-1, kurgan 1, burial 11. A and B – plan and photo of the burial, C - hummer-headed bone pin from the grave.**

*Bibliography to section 5:*

*Shishlina, N., Kazarnitsky, A., Leonova, N., Kuznetsova, O., & Sevastyanov, V. (2023). Yamnaya group from the Peschany IV burial ground in the Lower Don Eurasian steppes: local or non-local? Archaeolingua, 45, 29-43. Budapest.*

### 6. EARLY AND MIDDLE BRONZE AGE OF THE LOWER DON

The Don River, the Tanais of Greek colonists, flowed into the Sea of Azov, a northeastern branch of the Black Sea, through a wide marshland that began around its confluence with the Northern Donets on its right bank, widened after its left-bank confluence with the Manych River, and ended at the Gulf of Taganrog about 100km downstream. The estuarine marshes of the lower Don were a productive fishery, its broad floodplain supported dense gallery forests with wild game including red deer and horses, and the large stands of Phragmites reeds were a rich source of winter fodder and building materials. The strategic location of the lower Don at the junction of east-west and north-south trade routes made this region central for the Don Cossacks and for Medieval and Classical commercial traders. The steppes around the lower Don contained many Scythian kurgans and are regarded by historians as part of the Wild Field, occupied by nomadic pastoralists, who retreated into the lower Don estuary for their winter quarters.

Archaeology, anthropology, and genetics agree that during the Eneolithic and Bronze Age the populations of the lower Don occupied a zone of contact between the materially distinct cultures of the North Caucasus (south), the Eurasian steppes (east), the northern forest zone (north), the Pontic steppes (west), and beyond them the agricultural communities of southeastern Europe (far west). Stratified settlements in the lower Don go back to the earliest Neolithic about 6500 BCE. Eneolithic settlements such as Liventsovka and Razdor'ske are distinctive in that they exhibit continuous occupation from the late Eneolithic into the Yamnaya period, although the Yamnaya occupations have a lower density of both artifacts and features than the Eneolithic occupations. In the Volga-Caspian-Ural steppes, Eneolithic settlements such as Varfolomievka were abandoned in the Yamnnaya period and Yamnaya settlement sites are almost unknown. In the lower Don there was more apparent continuity from the Eneolithic into early Yamnaya both archaeologically (in settlement patterns) and genetically (in their larger retention of Seredni Stih genetics than is seen in Core Yamnaya).

#### 6.1 Cherkasov-4 site

*Summary by A. Faifert & D. Anthony*

The Cherkasov-4 site is located on a high watershed north of the Tuzlov River 37 km NE of Rostov-on-Don. 24 mounds have been excavated forming a southwest-northeast line of kurgans 1110 m. in length on the crest of the watershed east of the newly constructed Platov airport.

##### *6.1.1 Cherkasov-4, kurgan 2*

The kurgan diameter was 28 (E-W) x 29 (N-S)m, height 0.75 m. 12 burials were recorded in the mound. The main burial was burial 11, of the Early Bronze Age. Burial 1, in a grave with a side chamber or 'catacomb' dug into one side wall of the main grave pit, is close to it in time, also from the Early Bronze Age. Burial 8 was in the Middle Bronze Age, probably with construction of a larger mound over it. After this a group of later MBA 'Catacomb' burials were made: 2, 3, 4, 5, 6, 7, 9, & 10. The last burial was 12, in the Late Bronze Age.

###### *6.1.1.1 Burial 2:6 (individual ID I8949): 2464-2290 calBCE*

The burial belongs to the late stage of the lower Don Catacomb Culture of the Middle Bronze Age. It is dated 2464-2290 calBCE.

The grave was in a 'catacomb' or chamber dug into one wall of the main grave shaft. The fill of the main shaft contained layers of humus mixed with clay subsoil or 'materik'. In plan it had an irregular oval shape, 1.3x1.4 m, elongated N-S. In the northern part, at the bottom of the entrance shaft, was a brazier made of a broken ceramic sherd including a rim section. The rim is decorated with transverse nail indentations. A fragment of a vertical lug protruded from the shoulder. The interior clay paste was black with no visible impurities. The surface was brown, and lightly stroked with intersecting comb impressions. The reconstructed height of the vessel would be about 24 cm, with a diameter of about 30 cm.

A compact pile of charcoal was found in the southern part of the shaft floor. The entrance to the side chamber was a little above the level of the floor. The width of the entrance hole is 0.55 m. The chamber measured 1.75 x 2.05 m, irregularly rectangular in shape. In the central part of the chamber floor was a faint whitish trace of decayed plant litter.

On top of the litter lay the skeleton of an adult male 35-45 years old, on his back, with his skull on the south. The skull is turned to the right side, the right arm is extended beside the spine, the left arm is also parallel to the spine. The legs are bent to the right, the tibia are bent parallel to the femurs, with the heels to the pelvis. A small spot of bright red ochre was recorded 0.10 m east of the right knee.

Inventory:

1) 10cm south of the skull was a large flat-based ceramic vessel near the wall of the chamber. Its upper part was smooth, its lower part was covered with lightly swiped comb impressions. Its body is squatly rounded with a high sloping shoulder and a narrow rim, low and thick, with its inner edge cut flat. A herringbone belt of three lines of oblique indentations was applied with a narrow pointed tool along the outer edge of the raised rim. Two annular lugs with transverse stamped indentations were applied to the shoulders. The internal bottom of the vessel displays a raised "snowflake" pattern made of applied strips of clay twisted, pinched, and pressed into the bottom before firing. Sherd interiors are black with calcareous particles. The vessel surface is black to light brown. Vessel height is 25 cm, bottom diameter 18 cm, body diameter 40 cm, rim diameter, inside, 14 cm.
2) East of this pot, and partially under it, was a wooden bowl with yellow organic remains inside. The upper edge is beveled inwards. The diameter is about 20 cm, the height is 5.5 cm, the width of the rim is 0.9 cm.
3) The wooden bowl was partially placed on top of a large flat rectangular wooden dish (preserved in the form of a stain) measuring 40x75 cm. The long side of the dish was parallel to the buried male.
4) On top of the dish were the chopped bones of the spine of a domestic *Bos*.
5) The entrance to the chamber was closed by a wooden tripartite wheel made of oak. The three parts were united with wooden dowels (the type of wood has not been preserved) through two holes. One side board is 17x60 cm, the thickness along the edge is 7 cm, in the center there is an oval hole - 4.5 x 7.0 cm. The middle board had a thickened nave, its length is 60 cm, the maximum thickness of the nave was 15 cm, the diameter of the nave at the base is 18 cm, there are also two holes along the edge, the diameter of the hole is 10x13 cm. The other side board was 45 cm long, thickness 6 cm, groove spacing 36 cm, groove hole diameter 4 cm.

The burial is dated to 2464-2290 calBCE (3885±25 BP, PSUAMS-10864).

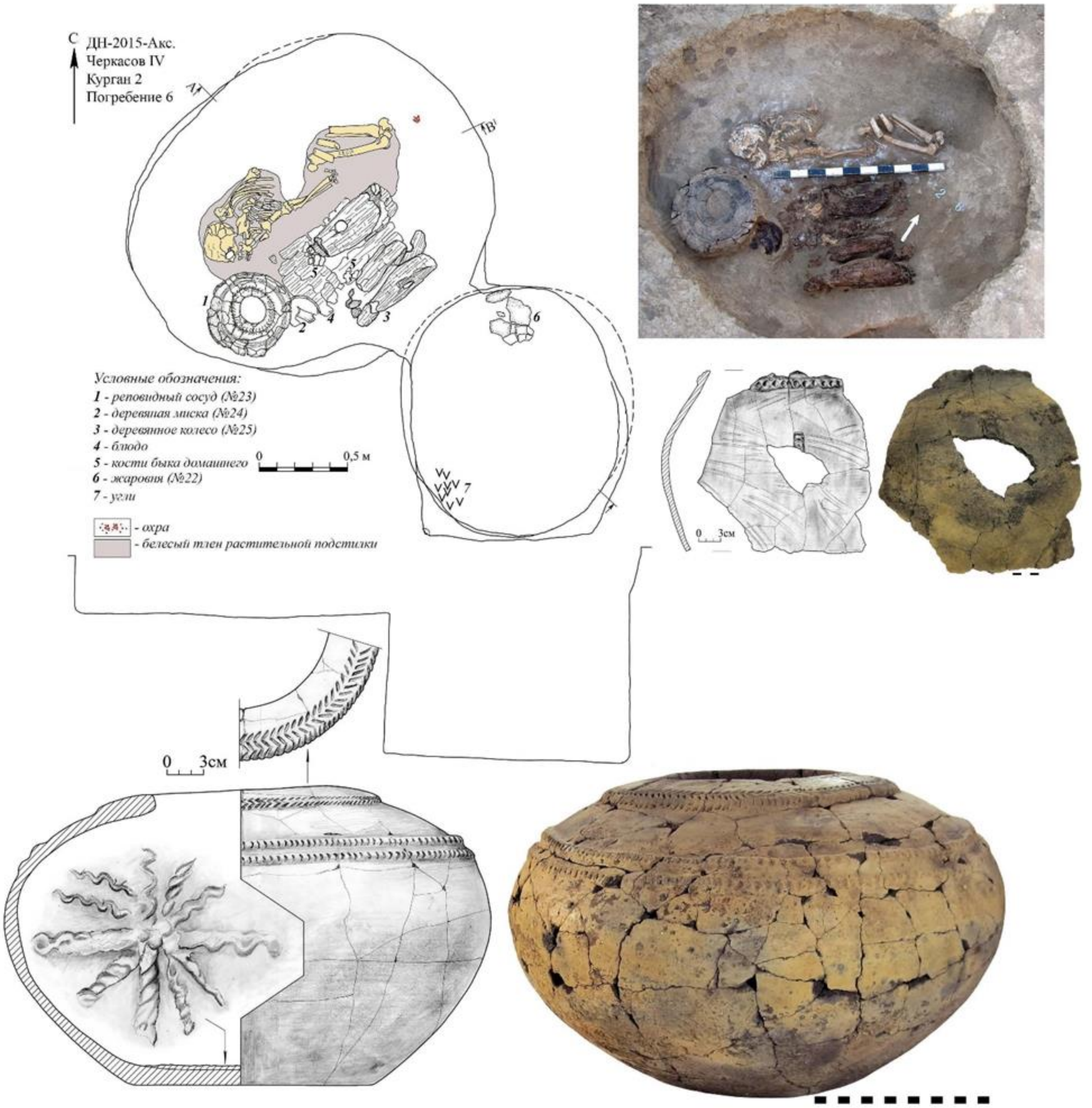

**Fig. 6.1. Cherkasov-4, kurgan 2, burial 6** (image by Anatoly Faifert).

##### *6.1.2 Cherkasov-4, kurgan 7*

3 burials were found in the mound. The mound embankment was almost completely destroyed by plowing. The main burial for the mound is burial 3, of the MBA. Burial 1 was LBA and burial 2 was disturbed. The main burial 3 was assigned archaeologically to the late stage of the Catacomb Culture of the MBA.

###### *6.1.2.1 Burial 7:3 (individual ID I26200)*

Burial 3 was the earliest in the mound. It was in a side chamber connected to the main grave pit. In plan the oval pit measured 1.35 x 1.95 m. A sloping step was built along its south-eastern wall.

The entrance to the side chamber was an oval hole .5 m wide and 0.4 m high, located in the northern pit wall. The entrance hole was covered with wooden timbers preserved in fragments. A low rim up to 4cm high was raised in front of the entrance to the chamber. A step up to 35cm high led to it. The oval chamber measured 1.65 x 2.45 m and the arched roof was 50-60cm high.

At the bottom of the chamber lay the skeleton of an adult male 25-35 years old, on his back, with his skull to the west. The front part of the skull is turned to the south, the right arm is extended beside the spine, the left is bent at the elbow at an obtuse angle, the hand is on top of the right hip joint and next to the right hand. The legs are bent parallel to the right, the tibia are strongly pressed with the heels to the pelvis.

Inventory:

1) At the bottom, next to the skull to the southwest stood a large ceramic vessel. On the inside of its flat bottom is a cross made of two twisted strips of clay with surfaces bearing transverse stamps. The body is wide and squat, with the largest diameter in the middle. On its shoulder are two parallel applied clay strips decorated with vertical indentations. Between them is a zigzag line of applied clay strips with the same decoration. The interior clay paste is black with calcareous particles. Its surface is covered with stroked comb marks and is black to light brown. Streaks of organic residue stain the shoulder. Height 25 cm, bottom diameter 22 cm, body diameter 48 cm, rim diameter inside 14.5 cm.
2) In the soil near the skull was a cylindrical bead carved from shell or soft calcareous stone. The dimensions are 0.7x0.5-0.6x0.2 cm.

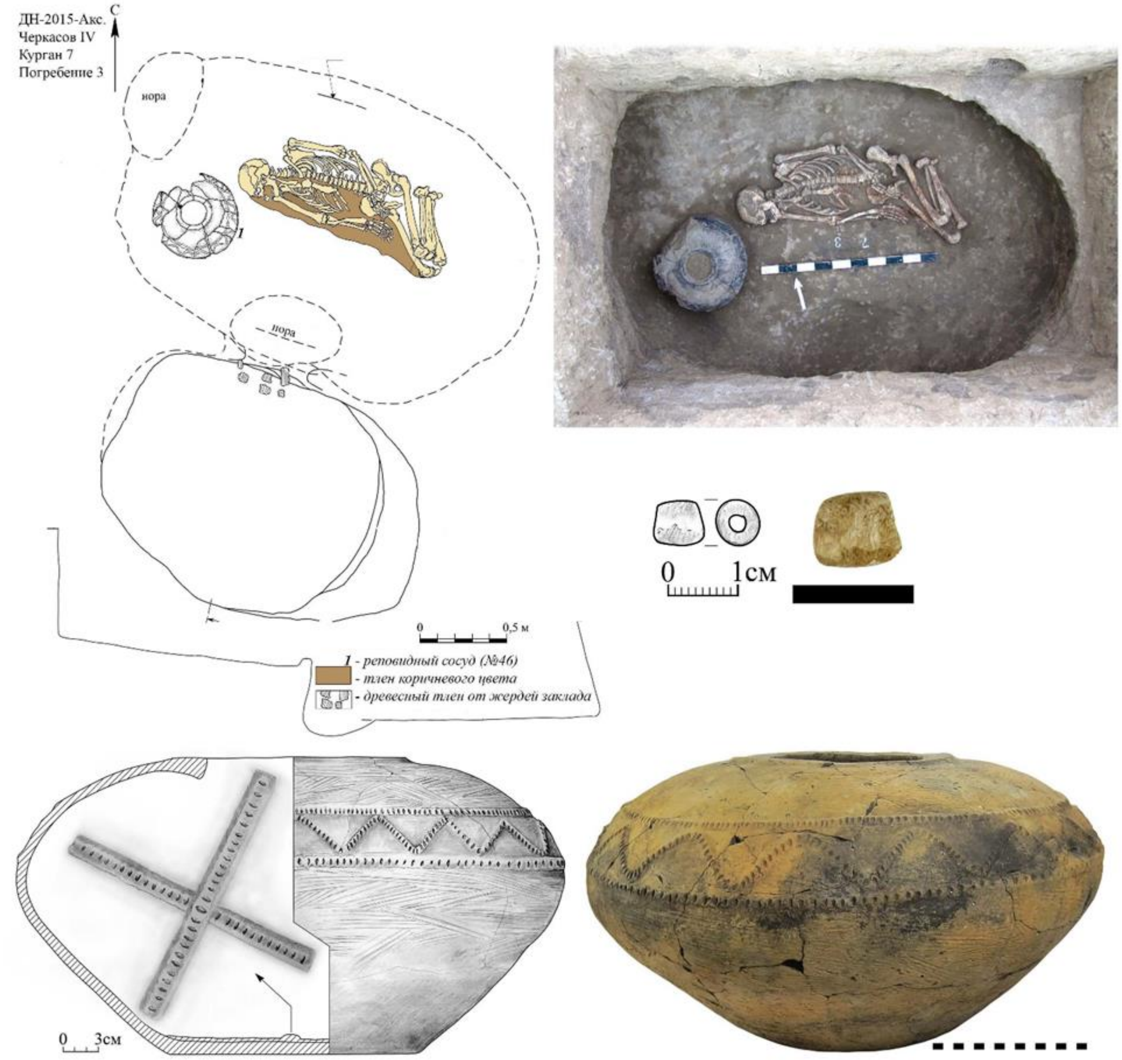

**Fig. 6.2. Cherkasov-4, kurgan 7, burial 3** (image by Anatoly Faifert).

#### 6.2 Krivyansky-7 site

*Summary by A. Faifert & D. Anthony*

The Krivyansky-7 and Krivyansky-9 kurgan cemeteries are part of a group of kurgans northeast of the village of Krivyanskaya on the Kadamovka River floodplain near a small lake, 41 km NE of Rostov-on-Don. The kurgan cemeteries were situated on elevated ridges extending out into the floodplain meadows. The excavations were carried out in 2016 by Olga Bespalaya.

##### *6.2.1 Krivyansky-7, kurgan 2*

Kurgan 2 is located on a low ridge trending west-east to the west of Kurgan 1. The height of the mound is 1.25 m, its diameter 31x34 meters. Its shape was slightly oval extending NW-SE. A concrete power line support was placed in the northern part of the mound. The mound was explored by 6 trenches oriented along the NW-SE line (deviation to the west by 65 degrees), leaving 3 baulks that had both facades cleaned. 17 burials and 12 utility pits were recorded in the mound, mostly from a settllement of the $8^{th}$-$10^{th}$ century CE. The first mound was built in the Early Bronze Age over grave 8. The materials have not been published before.

*6.2.1.1 Burial 2:8 (individual ID I8950): 2907-2704 calBCE*

The first mound was raised over a single grave of the Early Bronze Age in the center. Clay removed from the grave pit was spread as a thin lens on the original humus up to 4 m out from the grave. The top of the pit is rectangular, 1.15x1.85m, and the bottom widened to 1.25x1.95 m. The floor is 2.60-2.65m deep, almost level. It was covered with a stain of organic matter varying from dark brown to ochre brown.

On this organic material lay the skeleton of an adult male, on his back, with his skull to the side, face turned south. The right arm was parallel to the spine, the left was bent across the body, with the left hand touching the right forearm. The legs are bent at the knees to the right. The feet and shins of the buried are painted with red ochre. No funeral inventory was found.

The burial is dated to 2464-2290 calBCE (3885±25 BP, PSUAMS-10864).

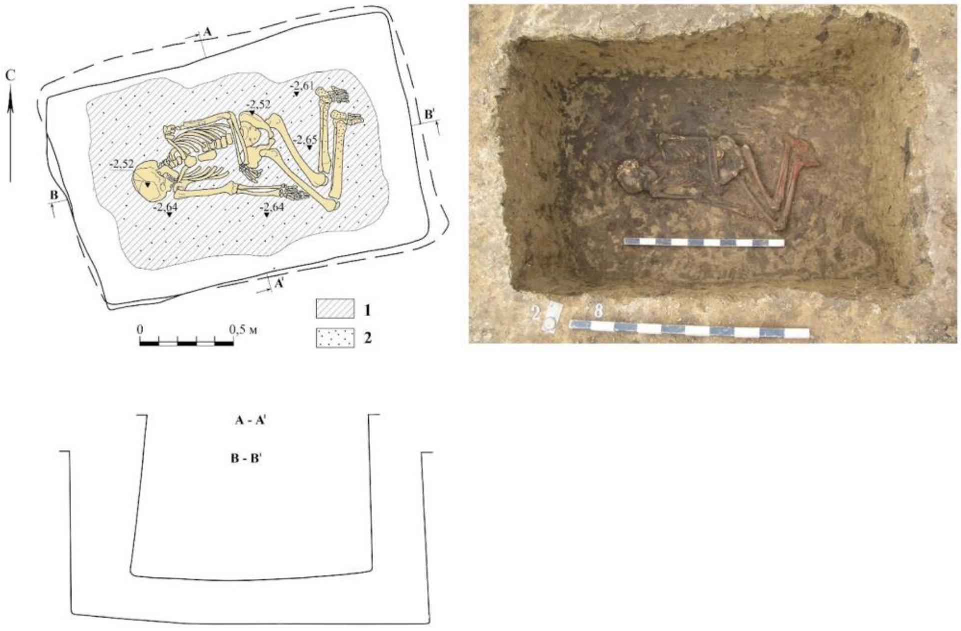

**Fig. 6.3. Burial 8 at kurgan 2 Krivyansky-7.** ***1*** *- brown leftovers;* ***2*** *- ocher spot (Bespalaya 2016, modified).*

#### 6.3 Kryviansky-9 site

*Summary by A. Faifert & D. Anthony*

As was noted above, Krivyanskiy-9 was a cluster of 7 kurgans located northeast of the village of Krivyanskaya on the Kadamovka River floodplain, 41 km NE of Rostov-on-Don. Like the village of Kryvianksii itself, the kurgan cemetery was situated on a low ridge or peninsula extending into the floodplain meadows, rising 2-3m above them. The Krivyanskiy-9 cluster contained 7 kurgans arranged in an uneven line trending NW-SE on the crest of a low ridge. The excavations were carried out in 2016 by Olga Bespalaya.

A GIS-based study of published Yamnaya kurgan cemeteries by Palalidis (2023) found that, among 223 published Yamnaya kurgans in the lower Don region, the median and mean number of Yamnaya graves in each kurgan was 1; the maximum was 3 (Palalidis 2023).

Krivyanskiy-9 is largely unpublished, and it deviates from this lower Don EBA pattern significantly in three ways. First, there are unusual EBA group graves like Kurgan 4 grave 20 (4:20) with four unrelated individuals (3 males and one female), and 4:21 containing 5 unrelated males, three of them children. Group graves are unusual in the EBA. 2nd, kurgans such as K1 and K4 have many EBA/Yamnaya and MBA/Catacomb single individuals in separate graves, adding to the unusually high number of individuals from Krivyanskiy-9.

Finally, while most EBA and MBA kurgans in the lower Don and elsewhere contain few or no genetically related individuals, at Krivyanskiy-9, 12 of the excavated EBA/MBA individuals were genetically related within 3 degrees (1st cousins or closer). These 12 related individuals included 7 males and 5 females whose Catacomb graves were scattered across Kurgans 1, 3, and 4 after they were built and before they were later enlarged. The Catacomb family seems to represent a shift to a more family-based use of these kurgans in the early MBA.

Six of the seven related males had R1b Yamnaya-like Y-haplogroups. Only one was I2a, which was the most common EBA Y-haplogroup on the lower Don. He was a 3rd-degree relative, perhaps a maternal cousin, of several R1b males among the 7, and he had the only polished stone mace head in the graves reported here. Related individuals were scattered across the three oldest kurgans:1, 4, and 3.

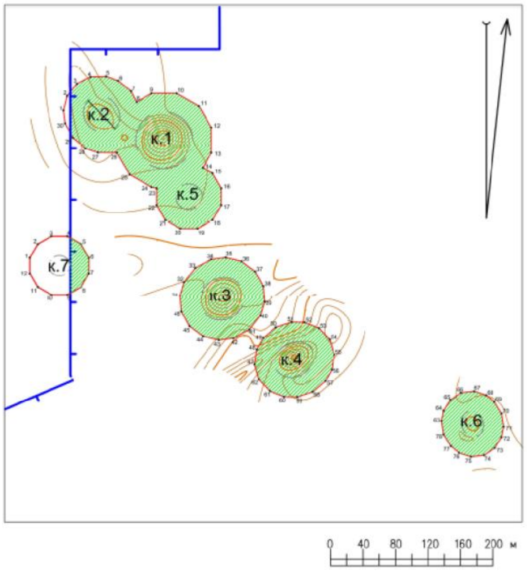

**Fig. 6.4. Preliminary schematic plan of kurgans 1-7 at Krivyanskiy-9** (image by Egor Kitov)**.**

##### *6.3.1 Krivyanskiy-9, kurgan 1*

Kurgan 1 is located at the eastern end of a low ridge or peninsula extending into floodplain meadows. The mound was 1.9 m high, diameter 46 meters. Excavations revealed 30 burials, a ditch, and 20 utility pits in or under the mound. Eneolithic grave 19 was created long before the north edge of the EBA kurgan covered it. Similar Eneolithic burials reportedly were found along the crest of the linear ridge. The base of the Kurgan 1 mound later overlapped with two other kurgans (2 & 5) built beside it, making a cluster of mounds associated with graves of the Eneolithic, EBA and MBA.

*6.3.1.1 Burial 1:18 (individual ID I8951): 2868-2584 calBCE*

Grave 1:18 was the main one for the second, enlarged mound. It contained a pair of subadults and was mostly destroyed by intentional robberies of the center of the mound committed in the 19th and 20th centuries. The robber's trench approached the center from the north. The original grave 18 pit was a rectangular pit, 1.2x1.6 m. In the fill of the northern half the scattered bones of 2 children, and the femur of an adult were found. On the pit floor was a dark brown/ochre brown organic stain. By the southern wall lay bones of the right arm and part of the right ribs, and judging by them, the position was supine, head to the west, arms parallel to the body, legs slightly tucked to the right at the knees, feet pulled to the east. The feet were painted with bright red ochre. A tibia of the older child lay in the northern half of the pit at the bottom, probably in situ.

Inventory:

1) Fragments of a small ornamented molded egg-shaped pot were found in the robbers' trench and grave fill. The surface of the vessel is covered with impressions made with a narrow pointed tool and a thin 3-5 toothed stamp: in the upper and lower parts there is a belt of three lines of an oblique stamp, the space between them in the middle part is filled with rectangles of several strips of a toothed stamp (one element is made with a narrow point) arranged vertically or horizontally. The sherd interior is dark gray with an admixture of crushed shells. The surface is from light brown to dark gray. H. -10.5; D. -13.0 cm.
2) At the right hand from the south lay a limestone disk. Thickness -3.9; D. -7.6x7.9 cm.
3) At 7 cm south of the right knee is a stone disk made of yellow porous shell rock. Thickness -4.1; d. -8.0x8.3 cm.
4) There is another stone disk of dark gray dense shale limestone with ferruginous layers close to it. The base is sanded, the edges of the lower platform are beaten, rounded to the top. Thickness -3.8; d. -6.6x6.9 cm.
5) There was a bone awl under the right shin made from a rib fragment of a small ungulate. Length -10.5; Cross section -1.1x1.9 cm.
6) At the left heel there was a bone tube made of tubular bird bone, with the epiphyses cut off. The diameter is not complete, glued from fragments. Length -7.0; Cross section -0.5x0.7 cm.
7) To the north of it there is also a bone tube made of the tubular bone of a large bird, with evenly trimmed epiphyses. Glued from fragments. Length -10.2; Cross section -0.9x1.3 cm.
8) A flint scraper was found under the right foot. According to the technical and morphological characteristics, the product can be interpreted as a planing tool. The flint is grey, spotted, with whitish inclusions. Dimensions: 65x35x13 mm.
9) A miniature flint chip was found in the filling of the litter near the western wall. Dimensions: 11x13x2 mm.
10) A fragment of the top of the profile of a jar pot of the Late Bronze Age was found in the soil of robbery. The shoulder is rounded, the mouth is narrowed, the corolla is highlighted by a thickening bent outwards. The clay is dense black in color with rare fine sand. The surface is dark brown, smoothed. The diameter is 19.0 cm.

The burial is dated to 2868-2584 calBCE (4130±20 BP, PSUAMS-8550).

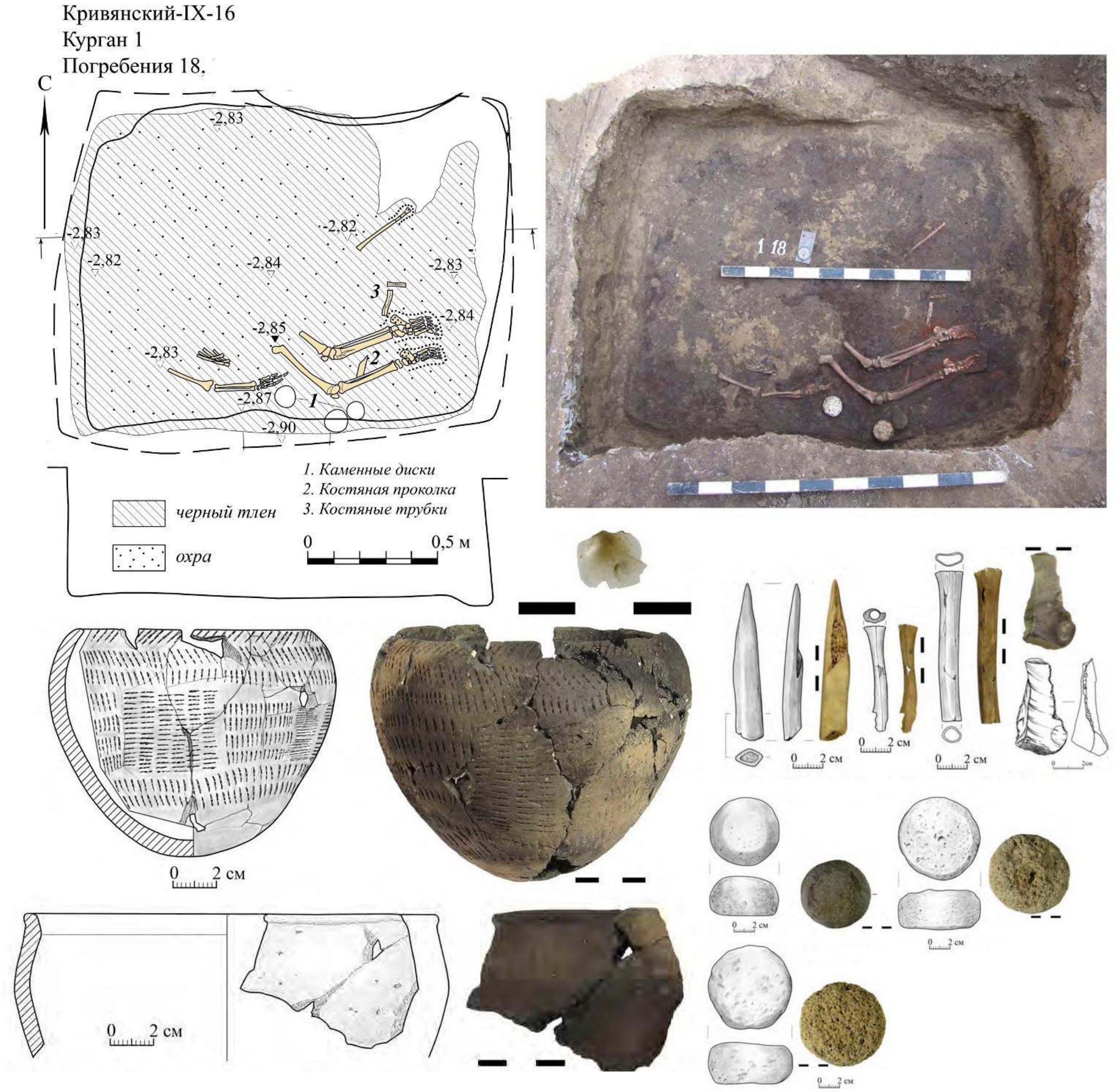

**Fig. 6.5. Krivyansky-9, kurgan 1, burial 18** (image by Anatoly Faifert).

##### *6.3.1.2 Burial 1:27 (individual ID I11829): 3330-2933 calBCE*

Grave 1:27 was recorded just to the north of Grave 1:18, but was much older, the oldest grave associated with kurgan 1 except for Eneolithic grave 19, described in a previous section. The original EBA burial mound 1 is associated with it. The grave pit is small, oval in shape, measuring 0.85 x 1.25 m, the walls are slightly expanded to the bottom. On the floor, sections of the transverse decay of the collapsed ceiling were found, probably caused by the robbers' trench into burial 18.

On the floor of grave 1:27 lay the skeleton of an adult female, mt-haplogroup U3a2, a haplogroup associated with agricultural populations aand unique to this individual in the steppes. Autosomally she is deviated toward the Caucasus populations, an outlier in the lower Don. She was heavily crouched, on her left side. She was robust, tall, and displayed unusual craniofacial traits said to appear Negroid, combined with with a severe occipital osseous deformity. Her arms were folded in front of the face, the knees are tucked into the elbows, the heels to the pelvis. The dark brown decay of the bedspread or clothing was noted in places. There is a red ochre stain on the right side of the lower jaw.

On the floor was a flint tool made of a retouched blade, trapezoidal in cross-section. The edges bear semi-circular flake scars from retouching. The flint is brown, translucent. Dimensions: 35x14x5 mm.

The burial is dated to 3330-2933 calBCE (4440±25 BP, PSUAMS-7867).

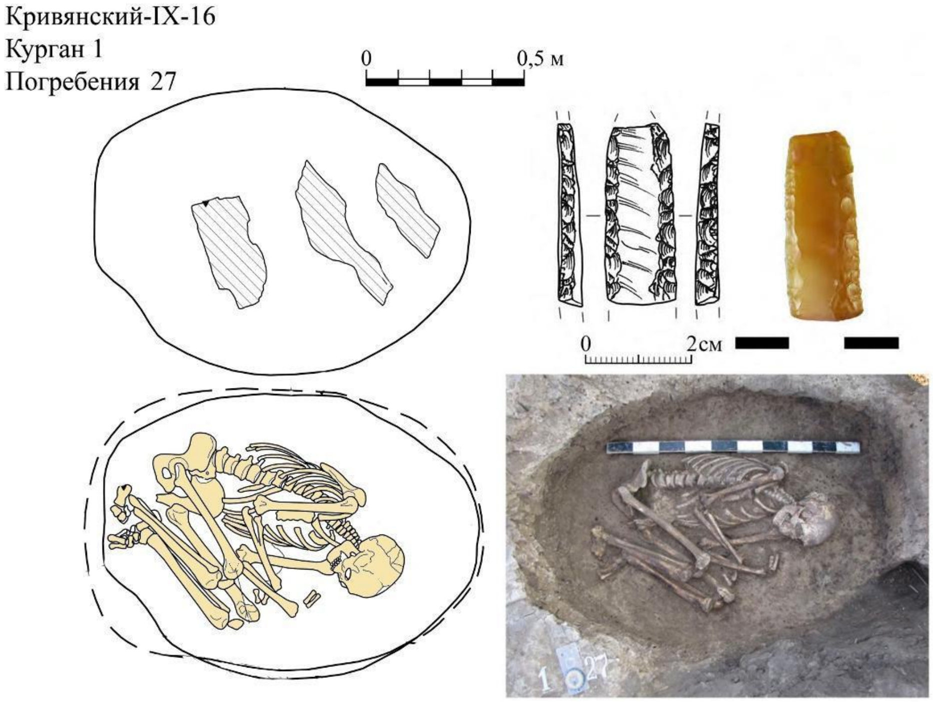

**Fig. 6.6. Krivyansky-9, kurgan 1, burial 27** (image by Anatoly Faifert).

##### *6.3.2 Krivyanskiy-9, Kurgan 2*

Kurgan 2 is located close to kurgan 1 in the middle part of a low peninsula elongated west-east, probably an old levee that formed between the ancient channels of the Kadamovka river. The height of kurgan 2 is 0.3 m, the diameter is 24x25 meters. 5 burials and 54 household pits were recorded in or under the mound. The earliest complex under the mound was designated Site 2 (stone tools of the Late Neolithic-Eneolithic period, perhaps associated with Eneolithic grave 1:19.). Kurgan 2 was erected in the EBA over Grave 2 after kurgan 1 was built. The mound also contains LBA and probably Medieval (#1) graves.

*6.3.2.1 Burial 2:2 (individual ID I8477): 2881-2633 calBCE*

Grave 2:2 was dug into the ancient ground surface. The pit was rectangular, measuring 1.05x1.70 m. The walls of the pit expand towards the bottom, the floor is flat. Charcoal was found in the fill and on the floor. In the center of the floor was an organic litter that left a stain colored dark brown/ochre brown.

On the litter lay the poorly preserved skeleton of an adult female (genetic determination), lying on her right side, skull to the west. The head was previously probably located on a pillow and was displaced to the right along with the cervical spine after decomposition. The right arm is stretched out lengthwise, the left bent at the elbow at 90 degrees, the left hand on top of the right forearm. The legs are bent to the right: the right hip is 135 degrees, the shin is 45; the left hip is 90 degrees, the shin is 40, the feet are brought together. There is a spot of bright red ochre along the right arm, spots of the same ochre near the knees.

A small molded round-bottomed vessel stood upside down 10 cm south of the skull. The body is spherical in shape, with a short rounded shoulder and a short thinly bent rim. There is a horizontal belt of vertical finger pinches along the shoulder. The clay is black with finely ground grog or chamotte temper. The surfaces are lightly brushed with combs, and are from dark gray to light brown in color. H. 7.8; D. body 9.7; D. rim 8.3 cm.

The burial is dated to 2881-2633 calBCE (4165±25 BP, PSUAMS-7978).

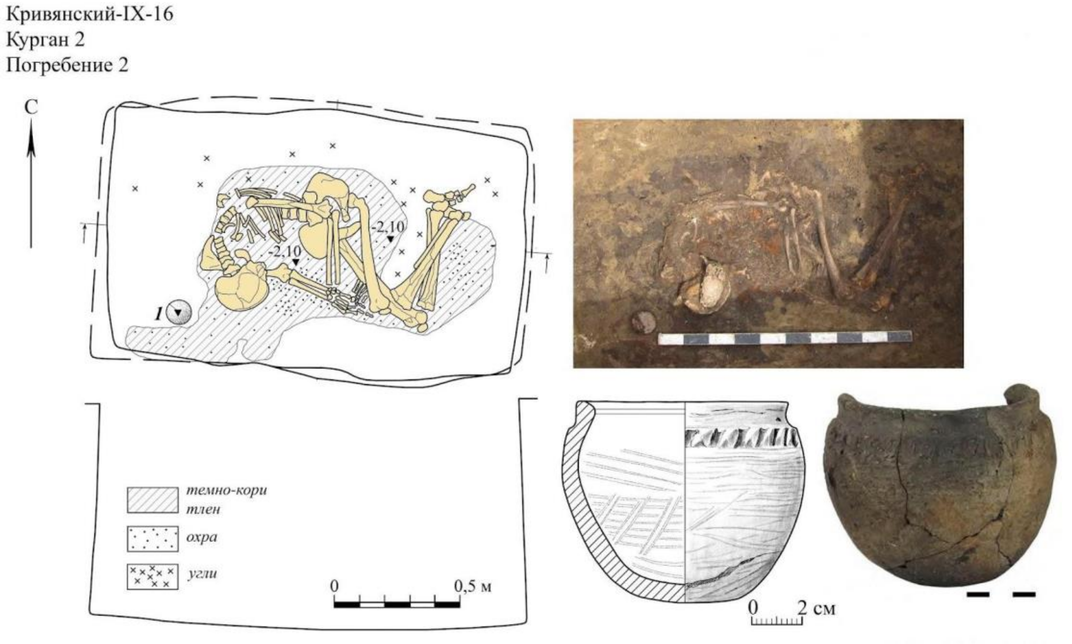

**Fig. 6.7. Krivyansky-9, kurgan 2, burial 2** (image by Anatoly Faifert).

##### *6.3.4 Krivyanskiy-9, Kurgan 3*

Kurgan 3 is located in the southern part of the Krivyanskiy-9 kurgan group, on a low ridge north of a washed-out old channel of the Kadamovka River. The diameter of the mound is 30 m, the maximum modern height was 1.4 m, the height of the from the horizon of the original ground surface was 2.1 m.

There are 19 burials of all ages in kurgan 3. The first mound was built in the EBA over grave 3:15. In the MBA, grave 3:18 was placed in the eastern part of the kurgan, as well as graves 3:11, 3:16, and 3:17 in other parts of kurgan 3.

*6.3.4.1 Burial 3:16 (individual ID I26435): 2916-2781 calBCE*

The entrance of the grave pit was sub-rectangular, measuring 1.5 x 1.8 m. The side chamber was wide-oval in shape, measuring 2.0 x 3.0 m. There was a thin litter in the form of dark brown organic stain along the floor of the side chamber, the bottom of which was spread with sand.

On the chamber floor lay the skeleton of an adult man, on his back, twisted to his right side, with his skull to the west (displaced by the collapse of the vault). The arms are extended with the hands to the thigh. The legs are bent to the right: the femurs at an angle of 160 degrees, the tibia at 45-80 degrees, the feet are brought together. There was no inventory.

His Y-haplogroup was R1b R-M12149, mt-haplogroup U2e. He was one member of the Catacomb culture family of 12 individuals that was buried across kurgans 3, 4, and 1. As a 2$^{nd}$-degree relative he could have been an uncle or nephew or half-sibling of the R1b male in K4:27 and the girls in K3:18a and 3:18b, and was a 3$^{rd}$-degree relative (1$^{st}$ cousin?) of three others in 4:25 and 4:26, and 1:30a.

The burial is dated to 2916-2781 calBCE (4260±25 BP, PSUAMS-10818).

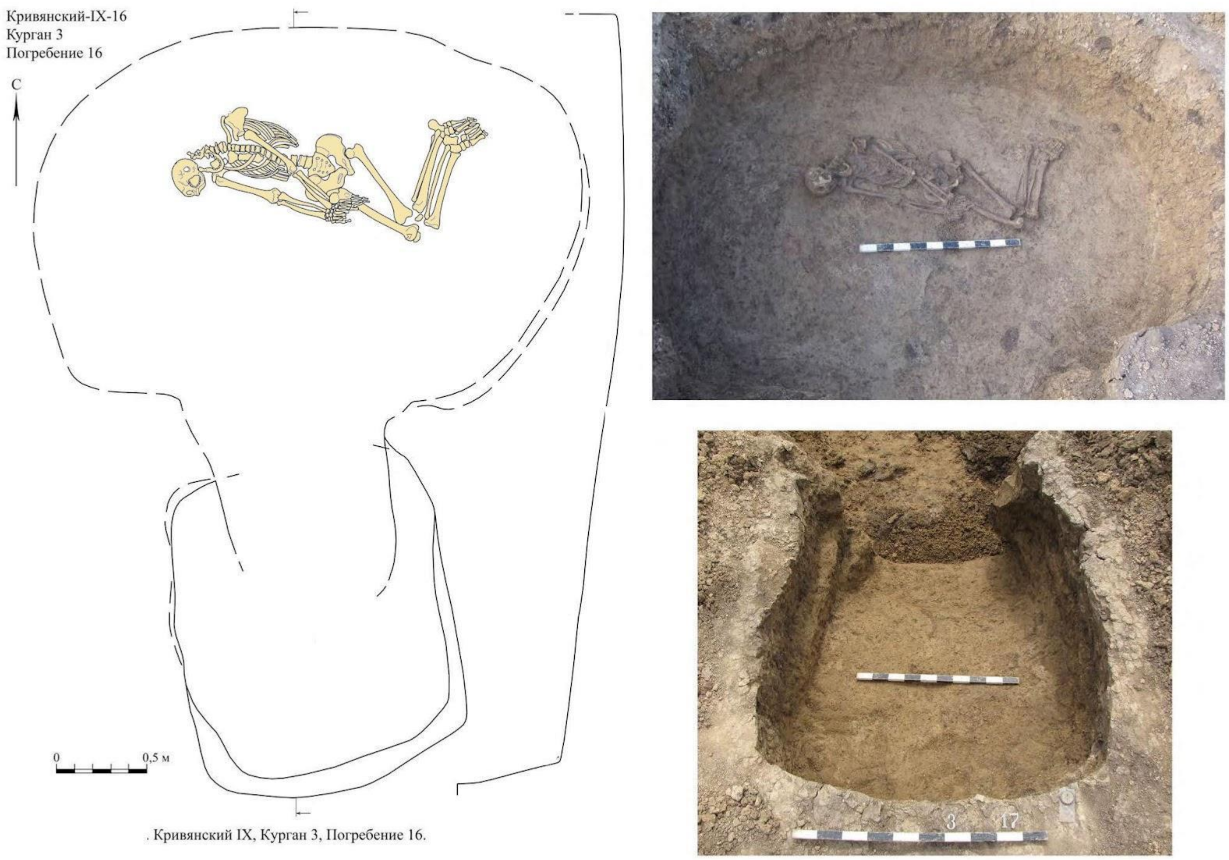

**Fig. 6.8. Krivyansky-9, kurgan 3, burial 16** (image by Anatoly Faifert).

##### *6.3.5 Krivyanskiy-9, kurgan 4*

Kurgan 4 is located in the southern part of the Kurgan group. The height of the mound from the buried soil is 2.0 m (the maximum height above modern surface was 1.45 m), the diameter is 40 m. The visually definable boundaries of the mound are increased due to erosion and plowing. 27 burials were recorded in kurgan 4.

The earliest is grave 4: 11 (not sampled), dug in the EBA in the original ground surface before the construction of the kurgan embankment. The original mound was enlarged twice, associated with burials of the Early and Middle Bronze Age.

The earliest phase of Kurgan 4 was built in the EBA over four graves: 18 (1 male),19 (2 males), 20 (3 males & 1 female), & 21 (5 males), placed on a small hill with a diameter of 10 m. These four graves contained 12 individuals: 11 males, all but one belonging to the I2a Y-haplogroup; and 1 female belonging to mt-haplogroup U5a2b. All were genetically unrelated within 3 degrees. The reason why males of various ages died and were buried together in group graves is not obvious. Perhaps an epidemic disease killed them. This small EBA cemetery was covered by the original kurgan 4 mound.

The mound was enlarged by the addition of a second layer of the embankment in the MBA over the Catacomb-culture grave 27. After the completion of the second embankment, strong erosion of the Kadamovka tributaries occurred on the site, marked by clay-sand layers in parts of the mound. The third expansion of the Kurgan 4 mound was connected with a group of MBA Catacomb burials: 16,23,24,25.

##### *6.3.5.1 Grave 4:18 (individual ID I26785): 2919-2875 calBCE*

Grave 18 in Kurgan 4 was part of an EBA group including graves 19, 20, & 21. The grave pit is rectangular measuring 1.10x1.75 m. A thin dark brown decay was observed in parts of the pit floor.

On the floor lay the skeleton of an adult man, on his back, with his skull north. The skull was displaced to the west. The right arm is extended, away from the backbone, the left is bent at the elbow toward the left side of the pelvis. The legs are bent to the right: the femurs at an angle of 120 degrees, the tibia parallel at an angle of 60 degrees. The feet and the lower part of the shins of the buried are stained with brown ochre.

His Y-haplogroup was R1b (R-M12149), unlike the other males in this group of graves; and his was the only grave containing a single male. His mt-haplogroup was U5a1j.

Funeral inventory was not found. The burial is dated to 2919-2875 calBCE (4270±25 BP, PSUAMS-10748).

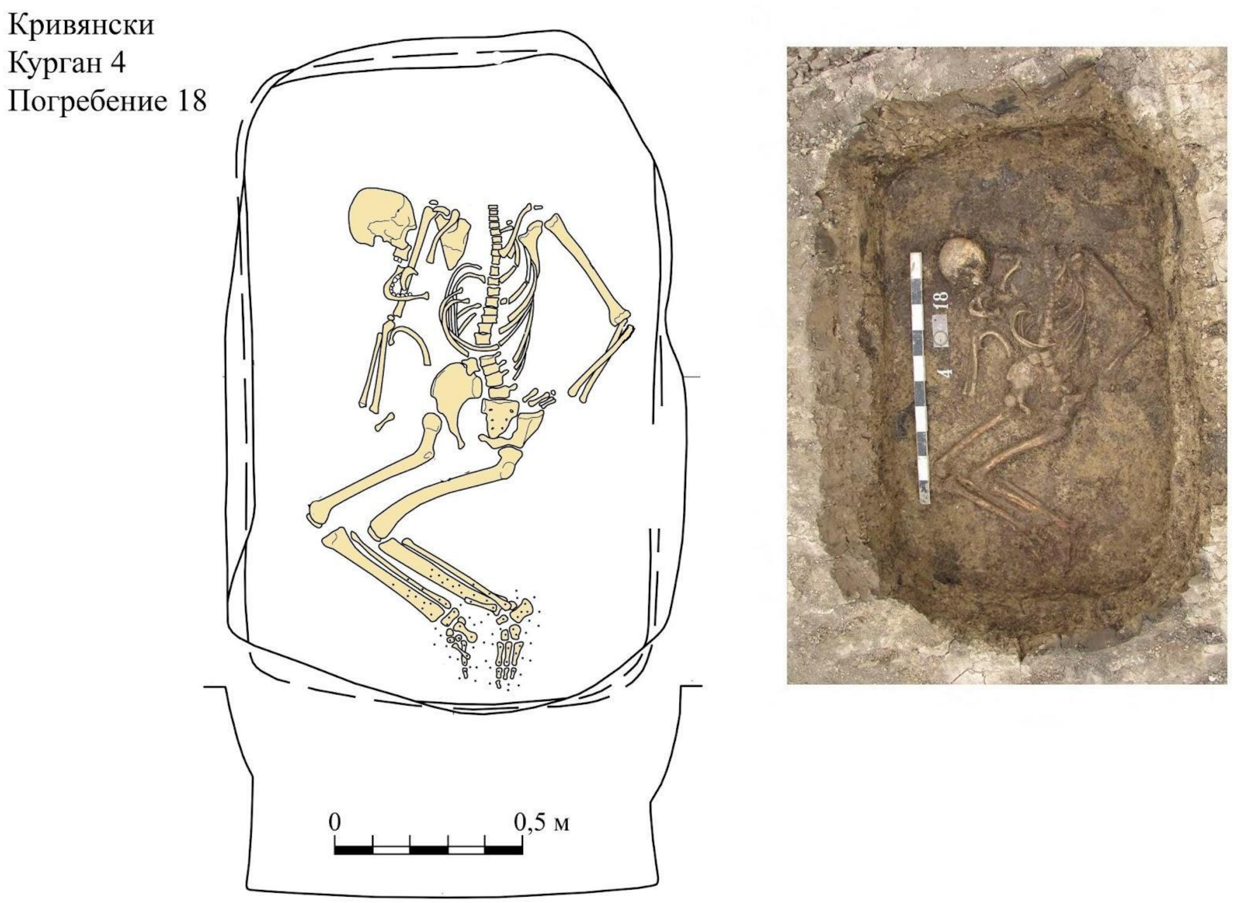

**Fig. 6.9. Krivyansky-9, kurgan 4, burial 18** (image by Anatoly Faifert).

##### *6.3.5.2 Grave 4:19 (IDs I24085 and I24086): 2882-2636 calBCE*

*Iindividual ID I24085 2882-2636 calBCE (4170±25 BP, PSUAMS-10742)*
*I24086 3011-2887 calBCE (4315±25 BP, PSUAMS-10810)*

Grave 4:19 contained two genetically unrelated males. The radiocarbon date ranges do not quite overlap for the two individuals; they are separated by 5 years, 2887-2882 calBCE. The grave pit was rectangular measuring 1.35x1.70 m, the walls are obliquely expanded to the bottom. Grave 19 is thought to have been in a group of 4 graves with 18, 20, & 21, probably covered by the earliest mound. On the floor of grave 19 was a thin decay of dark brown organic litter. The bones of two men, both Y-haplogroup I2a, lay on the litter, their skulls to the west.

Individual "A" (southern): an adult male, lying on his back, skull tilted to the right, arms extended along the trunk, legs bent parallel to the right, bent at the knees: femoral at an angle of 130 degrees, tibial at 20-30 degrees, feet brought together. His Y-haplogroup was I2a (I-L699), mt-haplogroup U5a1a1.

Individual "B" (north): a male teenager lying on his right side, his skull turned to the right, his arms extended parallel with the spine, his legs bent parallel to the right, bent at the knees. The feet of both males are stained with red ochre. Y-haplogroup I2a (I-L699), mt-haplogroup R1a1a.

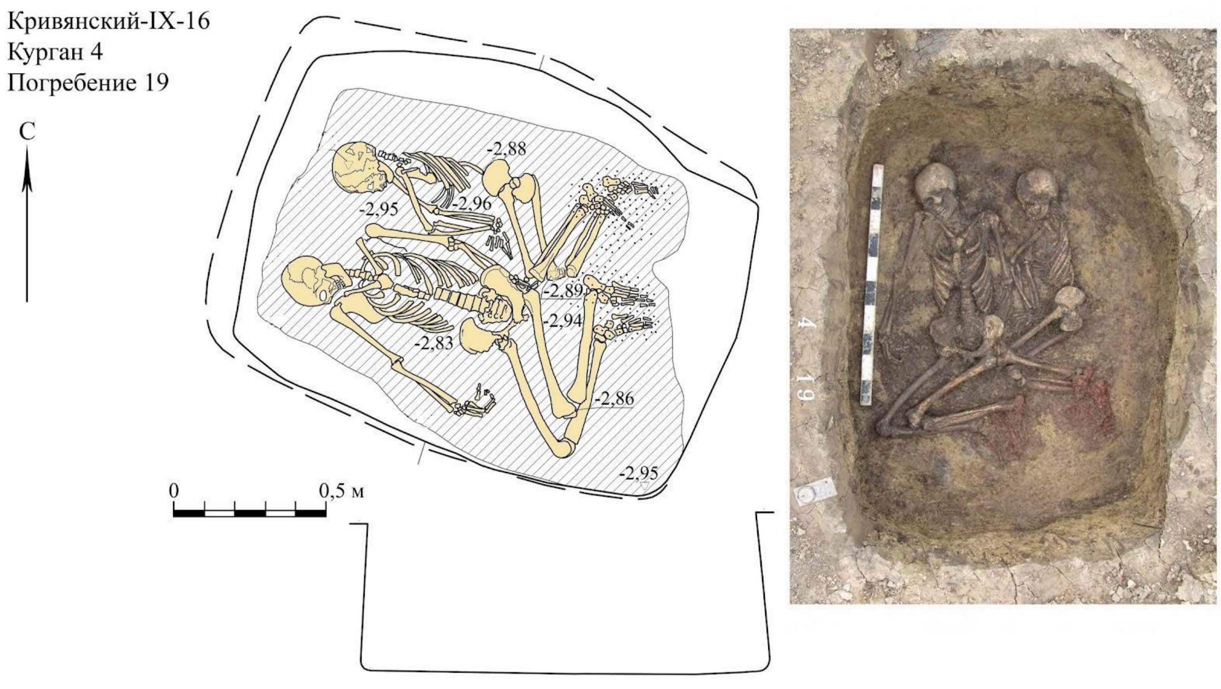

**Fig. 6.10. Krivyansky-9, kurgan 4, grave 19** (image by Anatoly Faifert).

##### *6.3.5.3 Grave 4:20 (IDs I12685, I24088, and I24087): 2917-2887 calBCE*

| | | |
|---|---|---|
| *Individual ID* | *I12685 A* | *2926-2877 calBCE (4285±25 BP, PSUAMS-8589)* |
| | *I24088 B* | *2921-2878 calBCE (4280±25 BP, PSUAMS-10811)* |
| | *I24087 C* | *3013-2889 calBCE (4320±30 BP, PSUAMS-10743)* |

Grave 4:20 contained 4 unrelated individuals including the only female in this group of EBA graves. The pit was rectangular measuring 1.15x1.85 m, the walls slightly expanded at the bottom, the bottom dimension was 1.35x2.0 m. There was a thin dark brown decay of organic bedding on the floor.

The 4 individuals were an adult man and three children of different ages. The adult and two children passed screening for aDNA: individuals A, B, and C. Genetic analysis showed that none were related to each other or to anyone else in the lower Don sample set within 3 degrees. However, A and B shared a T2a mt-haplogroup. All seem to have died at about the same time, but no skeletal indicators of violence were described.

Individual "A": a male child aged 5-7 years, Y-haplogroup I2a, mt-haplogroup T2a, was lying in the south, crouched on his right side, with his skull to the west. His hands are folded in front, his knees are tucked to the right.

Individual "B": an adult male, Y-haplogroup I2a, mt-haplogroup T2a, on his back, with his skull to the west. The right arm is extended touching the upper part of the right thigh, the left is bent at the elbow, touching the left side of the pelvis. The legs are bent at the knees to the right, the feet lie side by side, painted with red ochre with the lower parts of the shins.

Individual "C": an adult teenage female, mt-haplogroup U5a2b, lying against the wall of the pit, on her back, with her skull to the east. Her arms are stretched along her spine, hands on the pelvic bones. The legs are bent at the knees, bent to the left.

Individual "D": (not sampled for aDNA) a child about 2 years old, lying in the eastern part between the buried "B" and "C", stretched out on his back, with his skull to the east. Sprinkles of brown ochre were placed on the bones.

Inventory:

1. To the west of the skull of the buried "B" lay two ribs of a large ungulate. Length -36.0; Cross-section -1.0 x 2.7 cm; and Length -33.5; Cross section -1.0x2.5 cm.

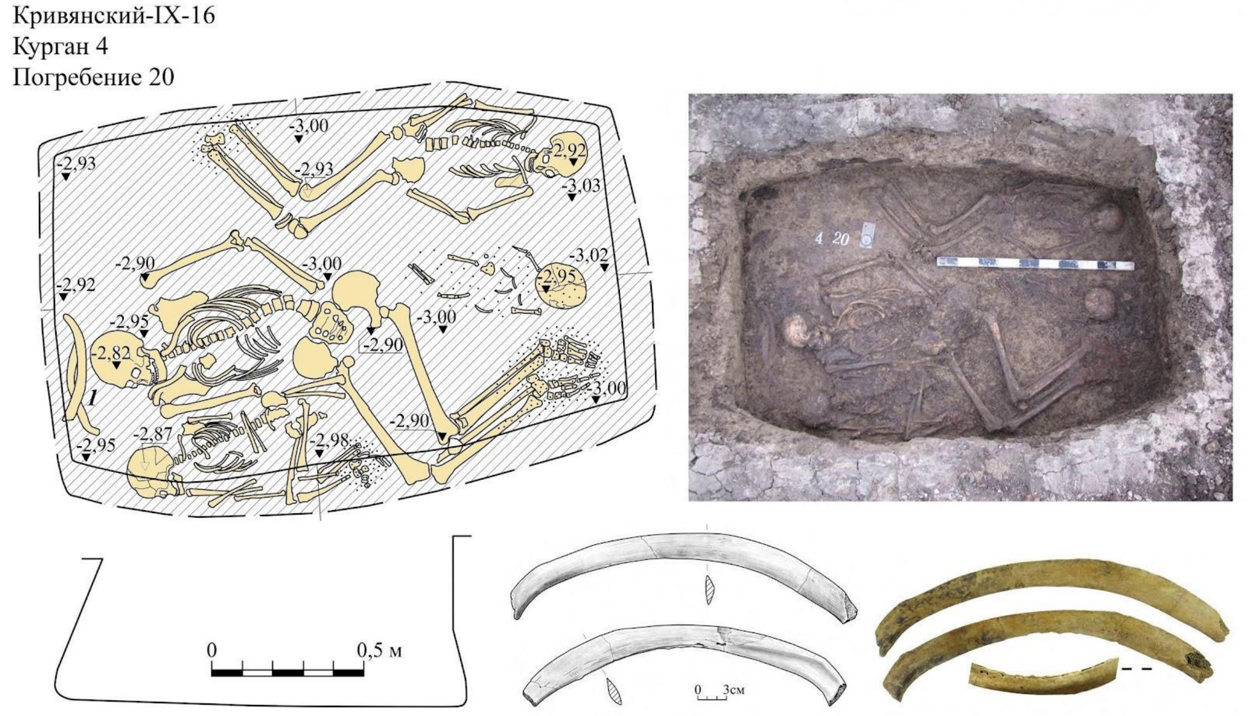

**Fig. 6.11. Krivyansky-9, kurgan 4, grave 20** (image by Anatoly Faifert).

##### *6.3.5.4 Grave 4: 21:*

| | | |
|---|---|---|
| *Individual ID* | *I12686 ?* | *3345-3096 calBCE (4533±18 BP, R-Combine)* |
| | *I12687 C* | *2904-2701 calBCE (4225±25 BP, PSUAMS-7980)* |
| | *I24089 E* | *2926-2877 calBCE (4285±25 BP, PSUAMS-8589)* |
| | *I24090 A* | *2926-2877 calBCE (4285±25 BP, PSUAMS-10813)* |
| | *I24091 B* | *3013-2889 calBCE (4320±30 BP, PSUAMS-10814)* |

Grave 4:21 contained five unrelated males of various ages. It was a sub-rectangular pit measuring 1.15x1.8 m at its opening, the walls were obliquely expanded to the bottom, the bottom dimensions were 1.45x2.05 m. A dark brown stain from organic bedding covered the floor.

Grave 21 contained two adult males, one of whom was ‘young adult’; and three male sub-adults, aged 3. 4, and 10. Genetically, none were related within 3 degrees to any individuals sampled from the lower Don. All seem to have died about the same time, but no skeletal signs of violence were reported. All belonging to Y-haplogroup I2a, but they exhibited 5 different mt-haplogroups. It is not obvious why they were buried together.

Individual A was studied for dairy peptides in his dental calculus. He had dairy peptides from drinking cow, sheep, and horse milk in his dental calculus (Wilkin et al. 2021).

Individual "A": a young adult male with Y-haplogroup I2a, mt-haplogroup U4a1, lay in the southern part of the pit on his back with his skull to the west. The skull is shifted to the south, the bones of the arms are stretched along the backbone, the left one is slightly bent at the elbow. The legs are bent at the knees and bent to the right. The feet are stained with ochre.

Individual "B": a male child over 10 years old, Y-haplogroup I2a, mt-haplogroup U5a1g1, was in the middle part of the pit, with his skull to the west, later displaced to the east. The arms are extended straight, the legs are bent to the right, slightly bent at the knees. The feet are stained with ochre.

Individual "C": an adult male, Y-haplogroup I2a, mt-haplogroup T2a1a, was lying against the northern wall, on his back, with his skull to the west. The skull is turned to the right, the right arm is extended along the spine, the left is bent at the elbow. The legs are bent at the knees and bent to the right. The feet of the buried are stained with ochre.

Individual "D": a male child about 3 years old, Y-haplogroup I2a, mt-haplogroup H15a1, lying in a corner of the pit, on his back, with his skull to the east. The skull is turned to the left, arms outstretched, legs bent at the knees and bent to the left. The skeleton of the child was sprinkled with 17 fragments of a ferruginous fine-grained mineral of dark lilac color.

Individual "E": a male child aged about 4 years old, Y-haplogroup I2a, mt-haplogroup U4a2, was in the eastern part of the pitl, on his left side, with his skull to the east. The arms are stretched out in front of the backbone, the legs are bent at the knees and bent to the left. On top of his pelvis lies the right knee of the buried "C".

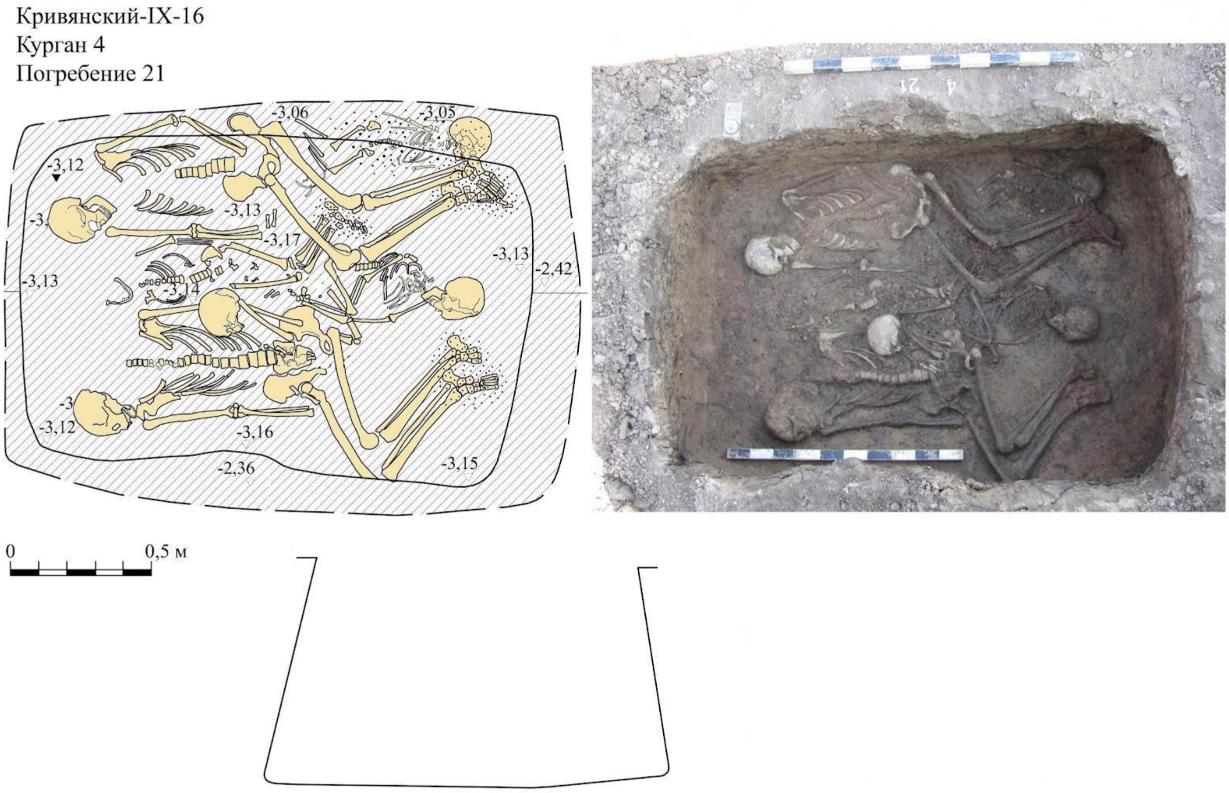

**Fig. 6.12. Krivyansky-9, kurgan 4, grave 21** (image by Anatoly Faifert).

##### *6.3.5.4 Grave 4:22 (individual ID I26781): 2627-2473 calBCE*

Grave 22 in Kurgan 4 was a Catacomb culture burial of the MBA. The vertical grave shaft was disturbed by graves 26 and 27. The side chamber containing the buried individual was an oval shape measuring 1.70x2.45 m. In the north-eastern part of the chamber, a faint decay of organic bedding was recorded, and ochre in the northern part.

The skeleton of an adult female lay on the organic mat. The right arm is bent at the elbow and pointed to the northeast, the left arm is also bent at the elbow and pointed to the southwest. The legs are bent at the knees and bent parallel to the right, the feet are brought together.

Inventory:

1. To the southwest of the spine were three lumps of dark red ochre. 1.5x1.8x2.6; 1.5x2.0x3.3; 1.7x2.8x3.4cm.
2. In the western part there was a brazier made from a rim sherd of a large ceramic pot. The shoulder is high and rounded, and a thin wavy roller is formed in the upper part by finger pinches. The rim is high, smoothly bent outward, horizontally beveled. The clay is dark gray with fine sand. The surface is light brown to black in color, sooty. Size -24.0x2.8; Diameter -24.0-25.0 cm.
3. In the southeastern part of the chamber there was a large pot with a base with a small diameter. The vessel has high rounded shoulders, and a thin wavy roller is formed around the upper shoulder by finger pinches. The neck is short, rim smoothly bent outward, flattened at the top. The clay is black in color without visible impurities. The surfaces are covered with smoothed combs. The surface colors vary from orange to dark gray. H -24.5; d. body -31.0; neck diameter -21.0 cm.
4. 30 cm to the west of the pot was a small molded cup. The bottom is flat, the body is conically expanded to the top, the upper edge is rounded. A small loop-shaped handle, flattened in cross section, is attached to the rim. The clay is black with fine sand. The surface is from bright orange to black, smoothed. The diameter is 5.8; The diameter of the bottom is 6.8; The diameter of the rim is 12.2.

The burial is dated to 2627-2473 calBCE (4040±25 BP, PSUAMS-10820).

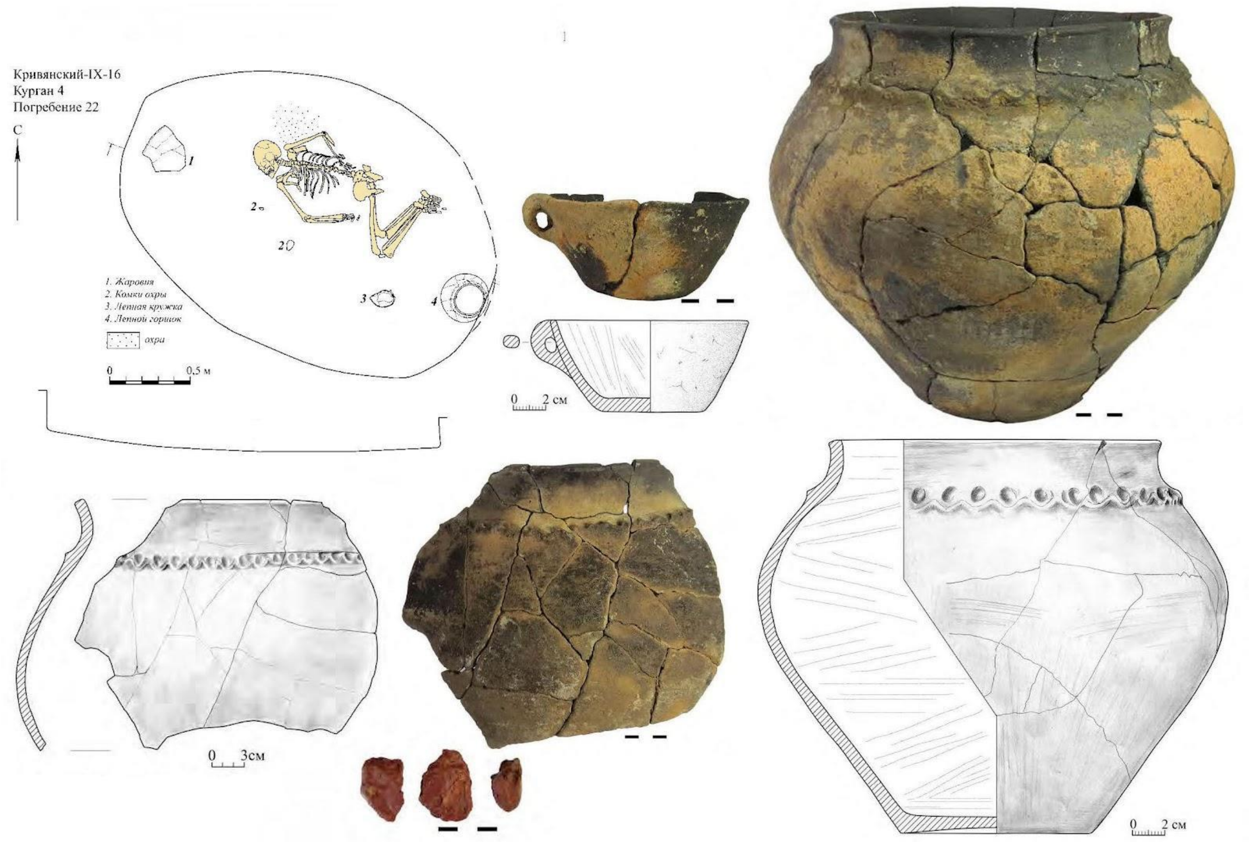

**Fig. 6.13. Krivyansky-9, kurgan 4, burial 22** (image by Anatoly Faifert).

##### *6.3.5.5 Grave 4:25 (individual ID I26782): 2862-2573 calBCE*

Grave 25 in Kurgan 4 was a Catacomb-culture grave of a young girl of the Middle Bronze Age. She was a member of an MBA family numbering 12 sampled individuals scattered across kurgans 1, 3, and 4 at the Krivyanskiy-9 kurgan cemetery in the MBA. This was a significant shift in the relatedness of the people using the Krivyanskiy-9 kurgan cemetery, from no relatives among the EBA people to a family cemetery in the MBA.

The girl in 4:25 was a sibling of I26634, buried in 1:30; and of I26639, buried in 1:30A; and I266340, 1:30B. She probably was a niece of I26158, buried in 3:14.

The rectangular grave opening measured 1.10 x 1.65 m. The side chamber was on the west side of the pit. It was oval shape, 1.85 m wide.

At the bottom of the chamber lay the skeleton of a female child aged about 3 to 4 years, mt-haplogroup U4b3, on her back, skull to the west. The arms are stretched out parallel, the legs are bent to the right at the knees.

On the left side of her skull was a bronze temporal ring, rolled up from a rounded wire in 2.3 turns, the ends were suspended. A thin wire is spirally wound on top, preserved in small sections. Diameter -1.5; Cross-section -0.25 cm.

The burial is dated to 2862-2573 calBCE (4105±25 BP, PSUAMS-10821).

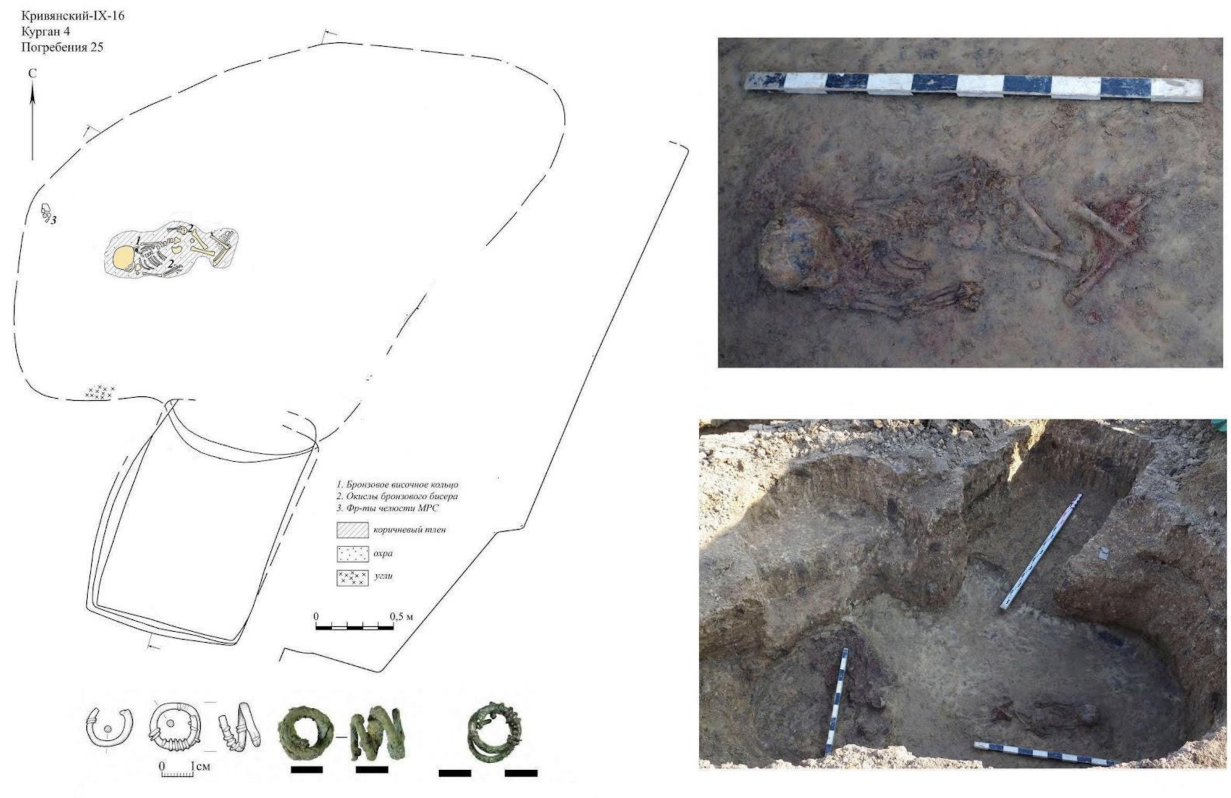

**Fig. 6.14. Krivyansky-9, kurgan 4, burial 25** (image by Anatoly Faifert).

##### *6.3.5.6 Grave 4:26 (individual ID I26783): 2847-2499 calBCE*

Grave 4:26 was a Catacomb Culture grave of the MBA with the richest grave inventory in the sampled individuals from this cemetery.

The pit entrance hole was sub-rectangular measuring 1-3 x 1.7 m. The side chamber disturbed the earlier Burial 4:27. On the floor lay the skeleton of a genetically male boy aged about 10 years, on his back, with his skull south. The right arm is extended beside the spine, the left is bent at the elbow, the left hand is on the right hip joint. The legs are bent at the knees and bent parallel to the right. There is an ochre stain in front of the knees. In the northeast corner of the chamber floor was a patch of light organic staining from bedding materials.

His Y-haplogroup was I2a, but he was a 3$^{rd}$-degree relative, perhaps a maternal cousin, of the adult R1b male buried beside him in grave 4:27; and also of both males in 3:18a and 3:18b; and of the R1b male in 3:16.
Inventory:

1. The boy wore a belt embroidered with 10 centrally pierced flat disc beads made of shell. The diameter is 1.3 cm; diameter of the central hole is 0.2 cm.
2. On the right at the entrance to the chamber there was a brazier for coals from a potsherd. The shoulder is short, rounded with a smooth transition into a high neck narrowed to the top, the upper edge is thinned and tilted inward. On the rim there is a horizontal

herringbone indentation with a toothed stamp. The clay is black in color without visible impurities. The surfaces are covered with smoothed combs. They are light brown in color. The size is -20.0x25.0; The diameter of the rim is about 28.0 cm.

3. A clay pot stood 40 cm northeast of the boy's skull. The bottom is flat, the body smoothly expands to the upper third, with a smooth transition to a high rounded shoulder. The upper neck is tilted outward, rounded at the top. On the rim and the upper part of the shoulder there are 4 horizontal belts stamped with a thin string, with rows of oblique notches between them. Triangles stamped with a cord-wrapped stamp descend from the lower belt along the body. The interior clay is black with calcareous and ferruginous particles. The surface is from light orange to dark gray. c. -17.7; d. bottom -10.5; D. body -22.0; d. corolla -19.5 cm.
4. Under the pot there was a polished stone mace/pommel made of dark gray dense limestone with layers of quartz, with a drilled central perforation. Rounded, flattened in cross section, one-sided drilling. Height -4.7; Diameter -6.5; Hole diameter -1.7-1.9 cm.

The burial is dated to 2847-2499 calBCE (4085±20 BP, PSUAMS-10822).

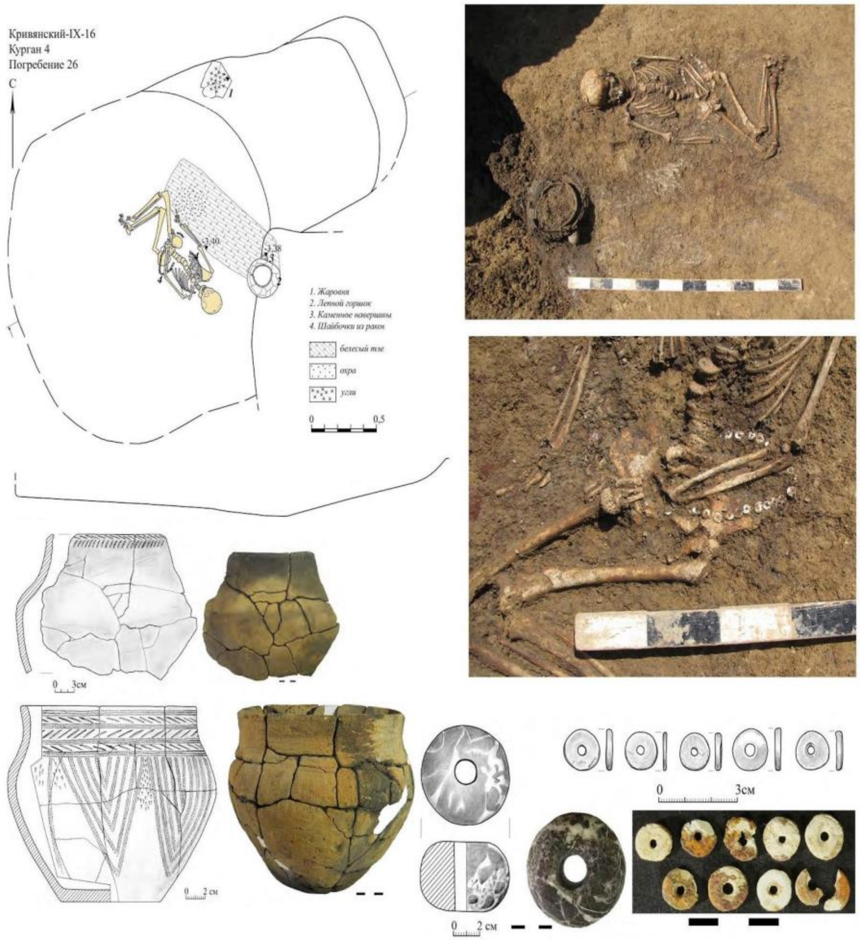

**Fig. 6.15. Krivyansky-9, kurgan 4, burial 26** (image by Anatoly Faifert).

*6.3.5.7 Burial 27 (individual ID I26638): 2889-2672 calBCE*

Grave 4:27 was a Catacomb grave of the MBA, a member of the MBA Catacomb family described above. The pit was rectangular measuring 1.25 x 1.95 m. In the western half, horizontal steps up to 0.25 m wide ran along the long walls. The entrance to the side chamber was narrow, a hole 0.72 m wide. The vault of the chamber collapsed in ancient times. An inclined step led to the chamber. The chamber was ovoid measuring 1.7 x 2.15 m. In the central part, there was a dark stain of decayed organic bedding.

On the floor of the chamber lay the skeleton of a buried adult, on his back, slightly to the right, with his skull to the south. The right arm is stretched beside the spine, the left hand on the backbone, slightly bent at the elbow, the hand in the groin area. The legs are bent at the knees and bent parallel to the right.

This male in 4:27 had Y-haplogroup R1b (M-12149) and mt-haplogroup W3a1. He was a 1$^{st}$-degree relative of the male in k3:18b; and a 2$^{nd}$-degree relative of the male buried beside him in 4:27, as well as the male in 3:16 and the male in 3:18a, and the male in 3:14a.

No inventory was found. The burial is dated to 2889-2672 calBCE (4190±25 BP, PSUAMS-10746).

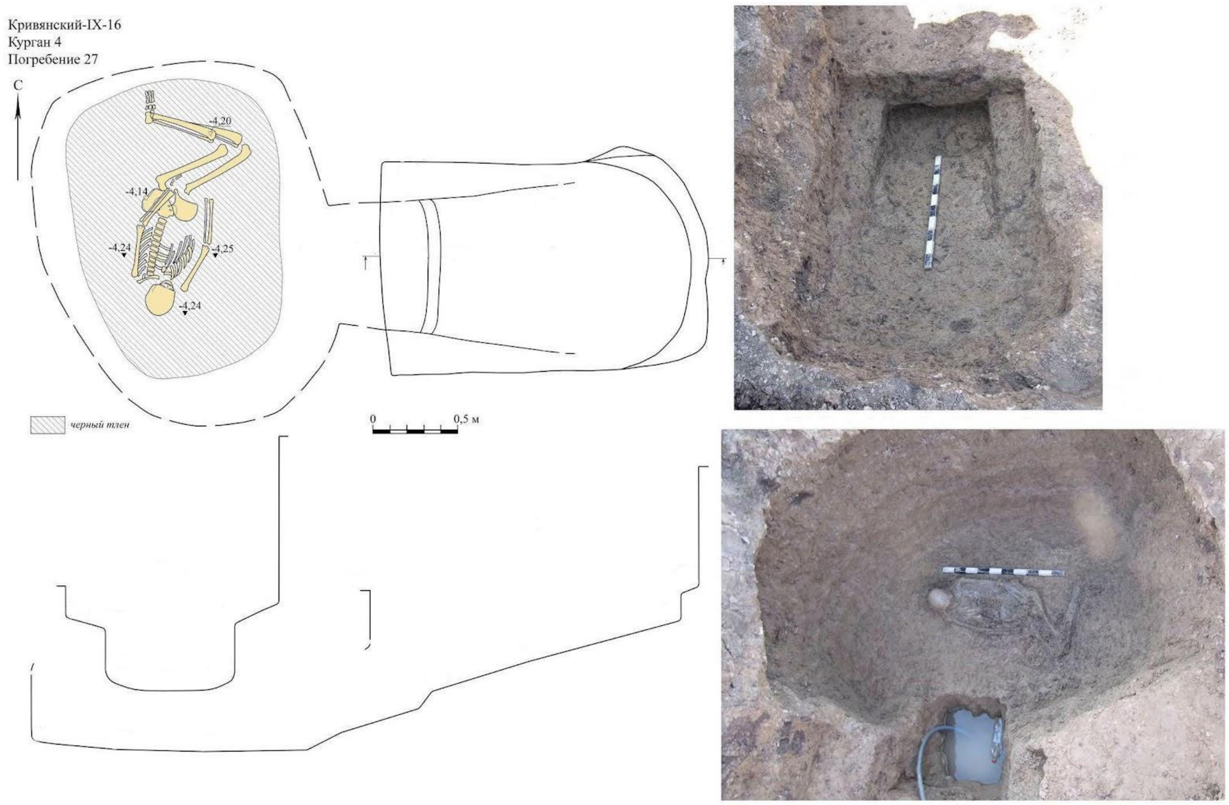

**Fig. 6.16. Krivyansky-9, kurgan 4, burial 27** (image by Anatoly Faifert).

#### 6.4 Chaltyrsky-11 site

The Chaltyrsky-11 kurgan cemetery is a local component within the group of kurgans, situated near the village of Chaltyr 15km NW of Rostov-on-Don overlooking a small river valley that opens into the Don estuary today, but perhaps was on the coast of the the Gulf of Taganrog in ancient times. The excavations were carried out in 2016 by Vadim Yatsenko.

##### *6.4.1. Chaltyrsky-11, kurgan 15*

Kurgan 15 is located in the southern part of the burial ground on a plowed field, on a low natural hill. Diameter 32 x 24 m, height 0.4 m. The northern part of the mound has been destroyed by an existing main gas pipeline. 8 burials were found in the mound, 1 of them from the Early Bronze Age.

###### *6.4.1.1 Grave 7 (individual ID I11029): 2896-2677 calBCE*

Grave 15:7 was the earliest in the mound. The northwestern part is damaged by a Scythian burial. The grave pit had rounded corners. Pit dimensions: length 1.52 m, width probably about 0.90 m.

On the pit floor was the skeleton of a 35-45-year-old genetically determined female, originally identified as male based on her robust skeleton. She was on her back with a half-turn to the right, oriented west-southwest. The skull is turned to the right, to the east. The left part of the skeleton and the pelvis were "cut off" by the burial of the 4th century BC. The right arm is bent at an angle, the right hand lies in front of the stomach, palm down. The right femur forms a right angle with the axis of the spine, bent at the knee at an acute angle.

Her mt-haplogroup was U2e1h. She had no relatives within 3 degrees among the examined samples.

Inventory:

1. Under her right hand was an unfinished polished stone hammer-axe head made of red quartzite. The working edge is not sharpened but is rounded, the butt is flattened, the surface is without traces of grinding. The hole for the axe handle is not finished. It is drilled to a depth of 1.5 cm. Hammer-axe dimensions: 16 × 6.5 × 6.1 cm, hole diameter: 3 × 2.8 cm.

The burial is dated to 2896-2677 calBCE (4205±25 BP, PSUAMS-10735).

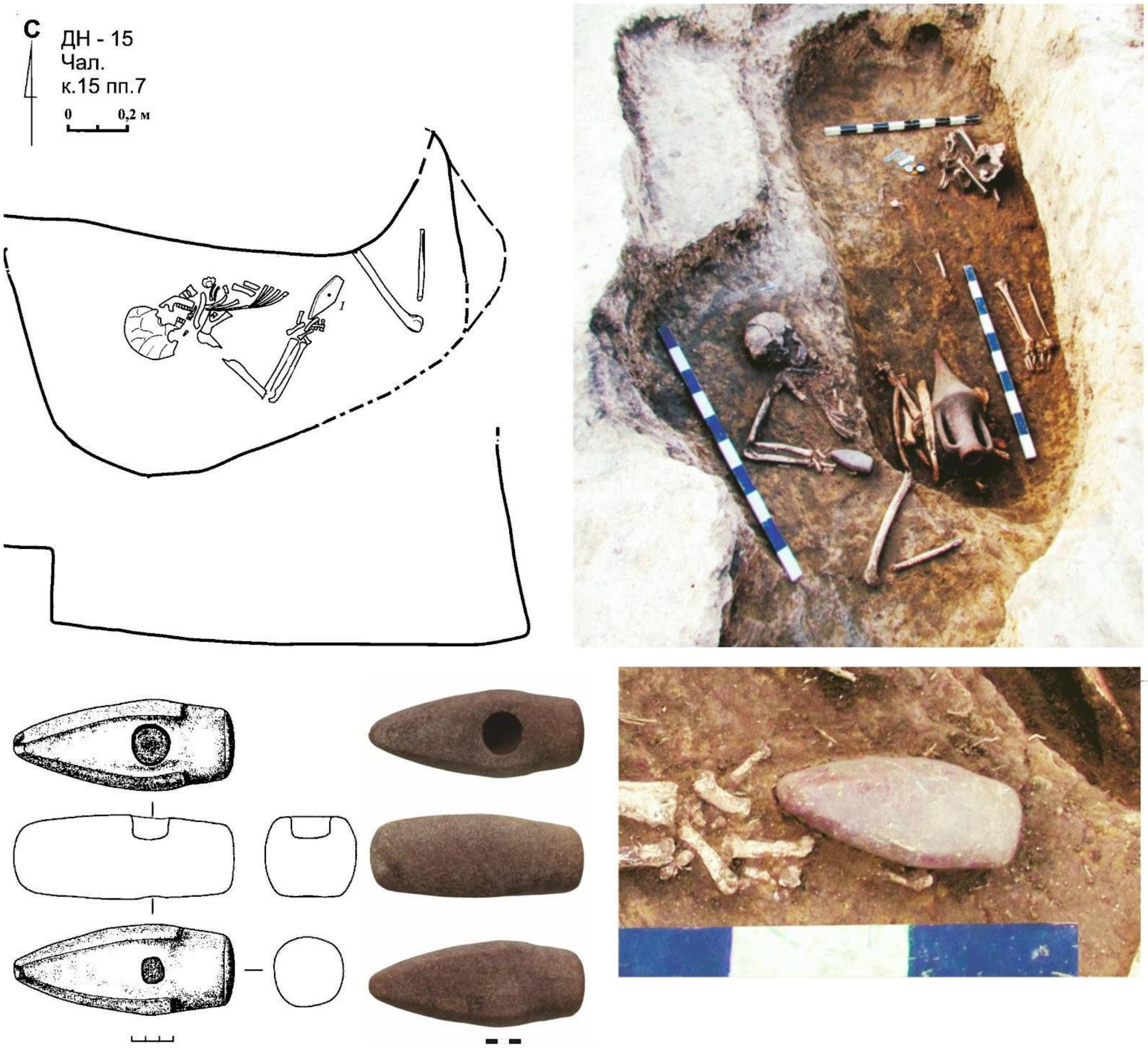

**Fig. 6.17. Chaltyrsky-11, kurgan 15, burial 7. 1 - stone axe** (image by Anatoly Faifert).

#### 6.5 Mokro-Chaltyrsky-1 site

*Summary by A. Faifert & D. Anthony*

The kurgan group "Mokro-Chaltyrsky-1" is located on the southwestern outskirts of the Chaltyr village 18km NW of Rostov-on-Don, on an elevated watershed between the rivers Safyannaya and Mokry Chaltyr overlooking the Don estuary, probably open sea in ancient times. The ancient Greek port of Tanais was on this coast just 10km to the west.

##### *6.5.1 Mokro-Chaltyrsky-1, kurgan 3*

The kurgan is located in the western part of the kurgan group on a small natural hill. The low mound is 0.7 m in height. The northern part has been destroyed by a trench of a gas pipeline.

###### *6.5.1.1 Grave 3:9 (IDs I8952 and I24093): 2907-2877 calBCE*

| | | |
|---|---|---|
| Individual ID | I8952 | 2892-2698 calBCE (4205±20 BP, PSUAMS-10770) |
| | I24093 | 3016-2897 calBCE (4335±25 BP, PSUAMS-10816) |

Grave 3:9 contained two unrelated males. It is stratigraphically the earliest grave in Kurgan 3. They were buried at the same time, but their radiocarbon dates do not quite overlap, missing by five years. The combination of the radiocarbon ages dates the burial to 2907-2877 calBCE (X-Test fails at 5% - X2-Test: df=1 T=16.545(5% 3.8).

The grave pit had a rectangular shape. The bottom of the grave was covered by a layer of brownish-black organic decay, reaching its highest thickness in the center and thinning along the grave walls. In some places, traces of dark red ochre were recorded.

The grave contained two males positioned head to foot. Spots of dark red ochre are recorded around the legs of both of them.

*Individual 1 (ID I8952)* was a 25-35-year-old male, buried next to the northern wall of the grave, on his back, oriented to the southwest and with his face turned to the south. The spine was straight, and the arms were slightly bent at the elbows. The right hand was located on the femur of the right leg, the left hand was missing. The bones of the pelvic girdle lay flat, the legs were bent at an acute angle to the south, and the tibia and femur bones lay parallel to each other. The knees overlapped the right arm of individual 2.

This male was R1b (R L-754) and U5a1a1.

A flint arrowhead 2.0x1.2x0.3 cm, with a deep notch at the base, was found at 4 cm to the west of the left wing of the pelvis of Individual 1. The surface is treated with fine retouching. The flint is translucent and gray.

*Individual 2 (ID I24093)* was a 35-45-years old male, located at the southern wall of the grave pit. The position of the deceased was identical to individual 1, but he was oriented to the opposite direction. The spine was straight, the arms were slightly bent at the elbows, the right hand was on the right femur, the left hand was missing. The bones of the pelvic girdle lay flat, the legs were bent at the knees and piled to the north, the tibia and femur bones lie parallel to each other. The knee joint of the right leg overlapped the elbow of the right arm of individual 1. The feet were in a free position, lying parallel to each other.

Like the other male, this one was R1b (R L-754), but his mt-haplogroup was U4a1e.

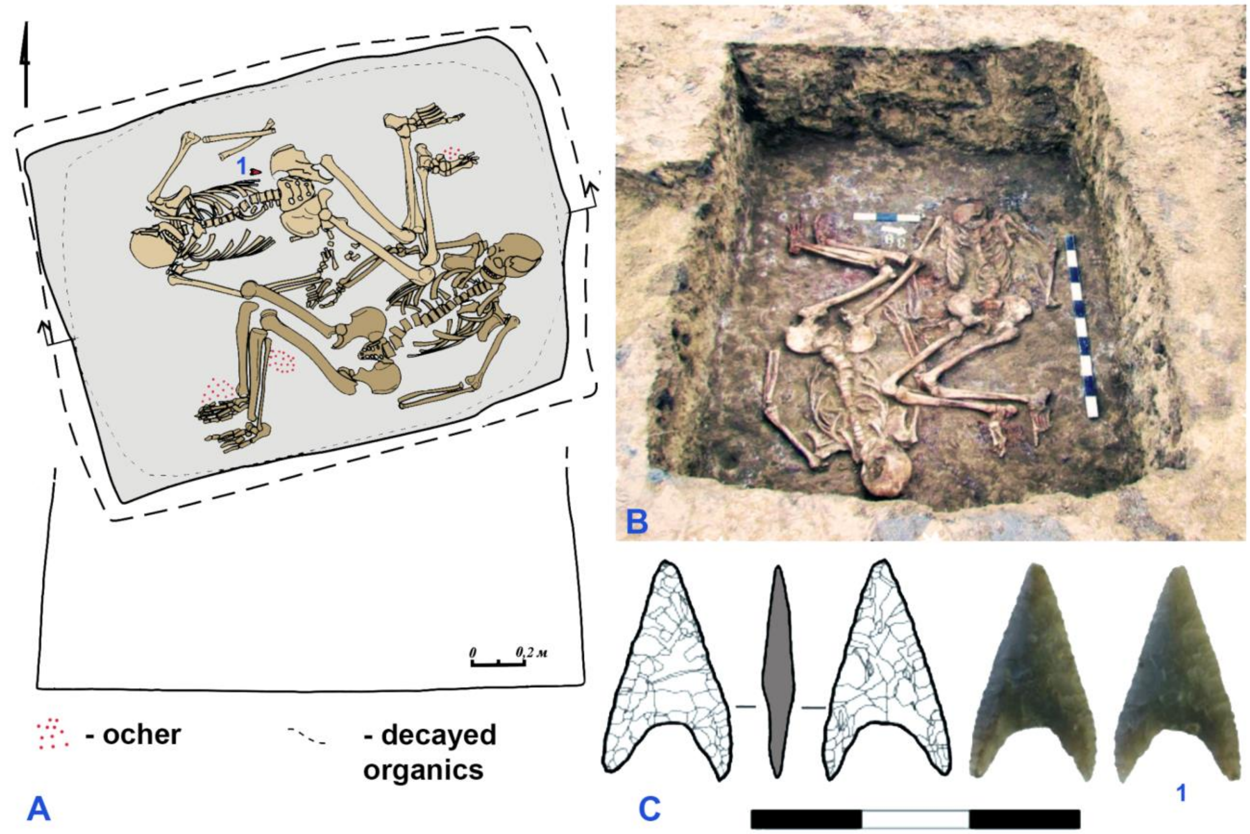

**Fig. 6.18. Mokro-Chaltyrsky-1 site, kurgan 3, burial 9.** A - plan of the burial, B - photo of the burial, C - grave goods: 1 - flint arrowhead (image by Anatoly Faifert).

#### 6.7. Vesely-1 site

*Summary by A. Kiyashko & D. Anthony*

Kurgan Vesely-1 is located in the eastern part of the kurgan group called "Tuzlov," consisting of 6 mounds stretched along the edge of the floodplain terrace of the left bank of the Tuzlov River. The Tuzlov flows eastward for 50km before joining the westward-flowing lower Don, and the triangular watershed between them is a high rolling steppe cut by a dense network of ravines and gullies, often filled by ribbons of forest.

##### *6.7.1 Vesely, kurgan 1 general description*

Kurgan 1 was a circular mound approximately 60 m in diameter and 4.0 m high, overgrown with trees and shrubs. In total, 49 burials belonging to various periods, from the Eneolithic to the Late Middle Ages, were discovered in the kurgan, along with 12 Early Medieval sacrificial pits and two residential structures (semi-subterranean dwellings) dating from the late 19th to early 20th centuries.

The primary burial under the kurgan was Grave 18 in a pit surrounded by a cromlech made of blocks of hard reddish-brown sandstone. Grave 18 was assigned to the Eneolithic-Early Bronze Age but was not sampled for aDNA. Initially, there was no mound above this cromlech. In the Early Bronze Age, a Yamnaya burial (1:17) was dug into the center of the cromlech. The adult in Vesely 1:17 probably was associated with the appearance a contemporaneous Early Yamnaya cemetery located mostly to the west of the cromlech’s center where children and adolescents were buried. These include graves 5, 10, 15, 20, 21, 22, 28, 30, 31, 38, 41, 43, accompanied by various megalithic structures including cromlechs, pavements, and other enclosures. After these burials, the first kurgan mound was constructed over them (mound 1).

Next in chronological order were Graves 11, 44, and probably 29 from the Late Yamnaya culture, inserted into the center of the kurgan. Mound 2 was constructed over them. Burial 29 was of a flint knapper and arrowhead maker.

The first graves of the MBA Catacomb culture were 26 (sampled) and 42 (sampled). Grave 26, inserted into the edge of the late Yamnaya mound 2, was covered with mound 3. Grave 26 belonged to the earliest phase of the Catacomb culture in the region: the Pre-Donetsk stage. Grave 42 was inserted into mound 3 and belonged to the subsequent, Early Donetsk stage. Catacombs 6, 8, 12, 13, 33, 45, 48 are also likely from this period. Mound 4 was constructed over all of them.

Burials 7, 9, 24, 25, 35, 36 belong to the developed and late phases of the Catacomb culture. It is probable that they are associated with the construction of mounds 5 and 6 of the kurgan. By the end of the 3rd millennium BCE, the kurgan had reached its modern size. Subsequently, its mound did not increase.

Burials of the Babino culture from the Middle Bronze Age, 4 and 32, were inserted into it. Then came the burials from the Late Bronze Age: 14, 16, 23, 34. Burial 19 likely belongs to the transitional period from the Bronze Age to the Iron Age. Burial 40, associated with the Early Iron Age Late Sarmatian culture, was surrounded by moat 1. The complex of ritual pits located in the southeastern part of the kurgan mound dates to the Early Middle Ages. Burials 2, 3, 47, and possibly 1 belong to the Late Middle Ages.

Some burials cannot be attributed to any specific period due to the lack of clear ritual features and inventory.

Finally, the latest stage of the kurgan's history is represented by Structures 1 and 2, which are agricultural buildings from the 19th-20th centuries.

The three Kurgan 1 individuals sampled from early MBA graves 26 and 42 were genetic brothers. In these two graves, one brother in 1:26 was an adult, one in 1:26 an adolescent, one in 1:42 a child, and the untested individual in Grave 42 was an infant. All three were I2a (I -L699) and U5a2.

However, according to the stratigraphy described above, the two older brothers were buried first, in Grave 1:26. The mound was then enlarged over their grave before the younger brother(s)? was buried in Grave 1:42. In the few years between their deaths, during the time span of one generation, the Pre-Donetsk early MBA artifact types of Grave 26 were replaced by the Early Donetsk early MBA types deposited in Grave 42. Typological changes that are thought to represent distinct chronological phases happened in this case during one generation, so that the older brother and his younger brother exhibited different chrono-types.

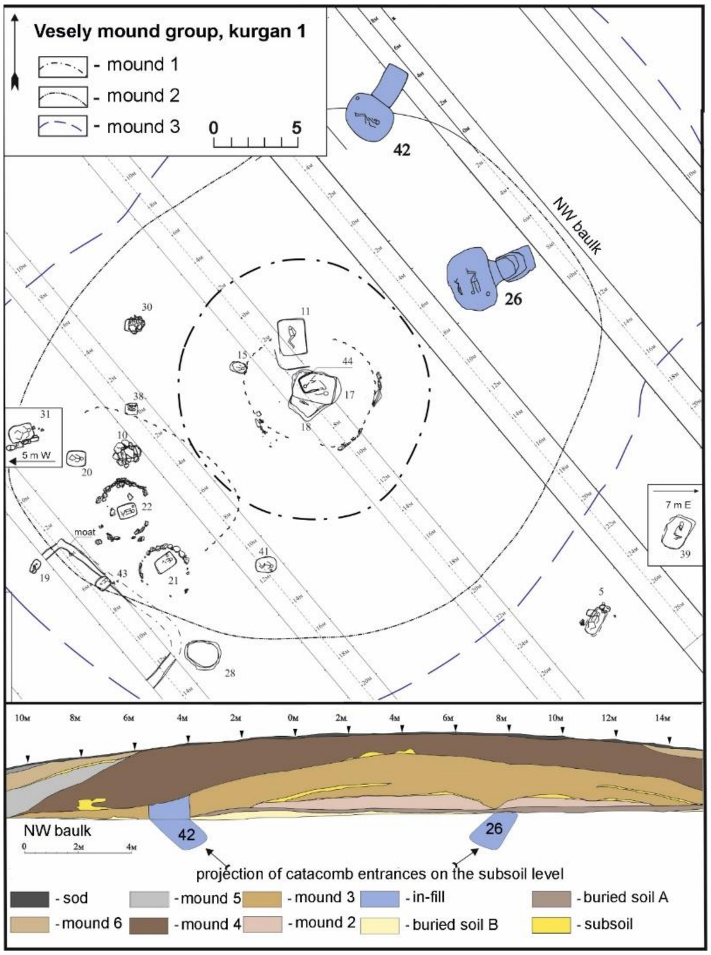

**Fig. 6.19. Vesely, kurgan 1. General plan with burials 26 and 42 highlighted in blue** (image by Aleksey Kiyashko).

*6.7.1.1. Grave 1: 26 (two brothers)*

*Iindividual ID I23654 2911-2881 calBCE (4270±20 BP, PSUAMS-14008)*

*I23655 2894-2701 calBCE (4210±20 BP, PSUAMS-14009)*

Grave 1:26 was discovered in the northeast sector of the kurgan, at a distance of 16.65 m from its center (FIG). It was a Catacomb grave. The entrance shaft was filled with dark-brown loam. It had a trapezoidal shape with rounded corners The top of the shaft measured 2.05 x 1.5 m. The walls were straight, gradually widening towards the bottom.

At a depth of 5.34 m from the top of the kurgan, small steps and a recess were arranged in the north side. Below, at a depth of -6.47 m, shoulder-like protrusions up to 0.37 m wide were observed on the N, E, and W walls. The bottom then sharply declined towards the entrance to the chamber, forming a steep step 0.5 m high. The depth of the entrance shaft reached 2.65 m from the original surface. In the fill of the shaft, several flint flakes, a fragment of pottery, and pieces of charcoal were found. To the right of the entrance of the side chamber, a cluster of charcoal was found. Individual fragments of charcoal were recorded almost throughout the entire floor. All walls, including the steps and protrusions, had vertical traces made by a narrow-bladed chisel-like tool.

The entrance to the side chamber had an oval shape. The dimensions of the entrance were 1.1 x 0.5 m, with a length of 0.4 m. The chamber was ovoid and elongated along the S-N line. The chamber's vault had collapsed, and its height and shape could not be reconstructed. The floor dimensions were 3.35 x 2.5 m.

Individual 1 (ID I23654). A well-preserved adult was positioned near the entrance. The individual was buried on their back with the legs bent to the right, and the skull facing the SW. The skull was shifted towards the entrance, and the lower jaw was found at the right elbow joint. The bones of the right arm were stretched along the torso, with the hand lying next to the right iliac wing. The bones of the left arm were stretched along the torso, with the radius bones lying under the left iliac wing and the hand located under the femoral joint. The legs were bent at the knee joint, to the right. The skeleton was placed on a complex bedding consisting of white decayed matter covered with brown decayed matter, overlaid with ochre.

Individual 2 (ID I23655). A well-preserved adolescent was found closer to the W wall of the chamber. The individual was buried on their right side with the skull facing the SW. The skull was separated from the rest of the skeleton and placed under the left shoulder joint of skeleton 1. The mandible was located between the skeletons. The arms of both skeletons were stretched in front, with the hands between the femurs. The legs were slightly bent at the knee joints, with the knees facing SW, and the left leg bones overlapping the right ones. The body was placed on a bedding of dark organic matter. Under the organic bedding a stone slab was found under individual 2. The displaced position of the adolescent's skull, placed under the adult's shoulder bone, indicates that the adult was placed in the grave after the adolescent. Both skulls were moved a significant distance from their bodies.

At the feet of Individual 2, a skull and long bones of sheep or goat were found.

Inventory:

1. Next to the shoulder bone of skeleton 2 – a flint scraper;
2. A bronze knife was found in the same area;
3. A bronze awl was also found nearby;
4. Near the cervical vertebrae of skeleton 2, a silver ring was found;

5. Next to the shoulder joint of skeleton 2 – an astragalus bone;
6. A second astragalus was found near the lower jaw of skeleton 2;
7. A bone piercing tool was located next to the shoulder bone of skeleton 2;
8. A second silver ring was found during the disassembly of skeleton 2 under the cervical vertebrae.

The items were laid out compactly, suggesting they were packaged in a pouch. All tools were oriented southward. Between the skull of skeleton 2 and the flint scraper was a flint flake. At the northern wall corner, on a burnt board, an ornate clay vessel was placed.

The burial dates to the early phase of the Catacomb Culture of the MBA.

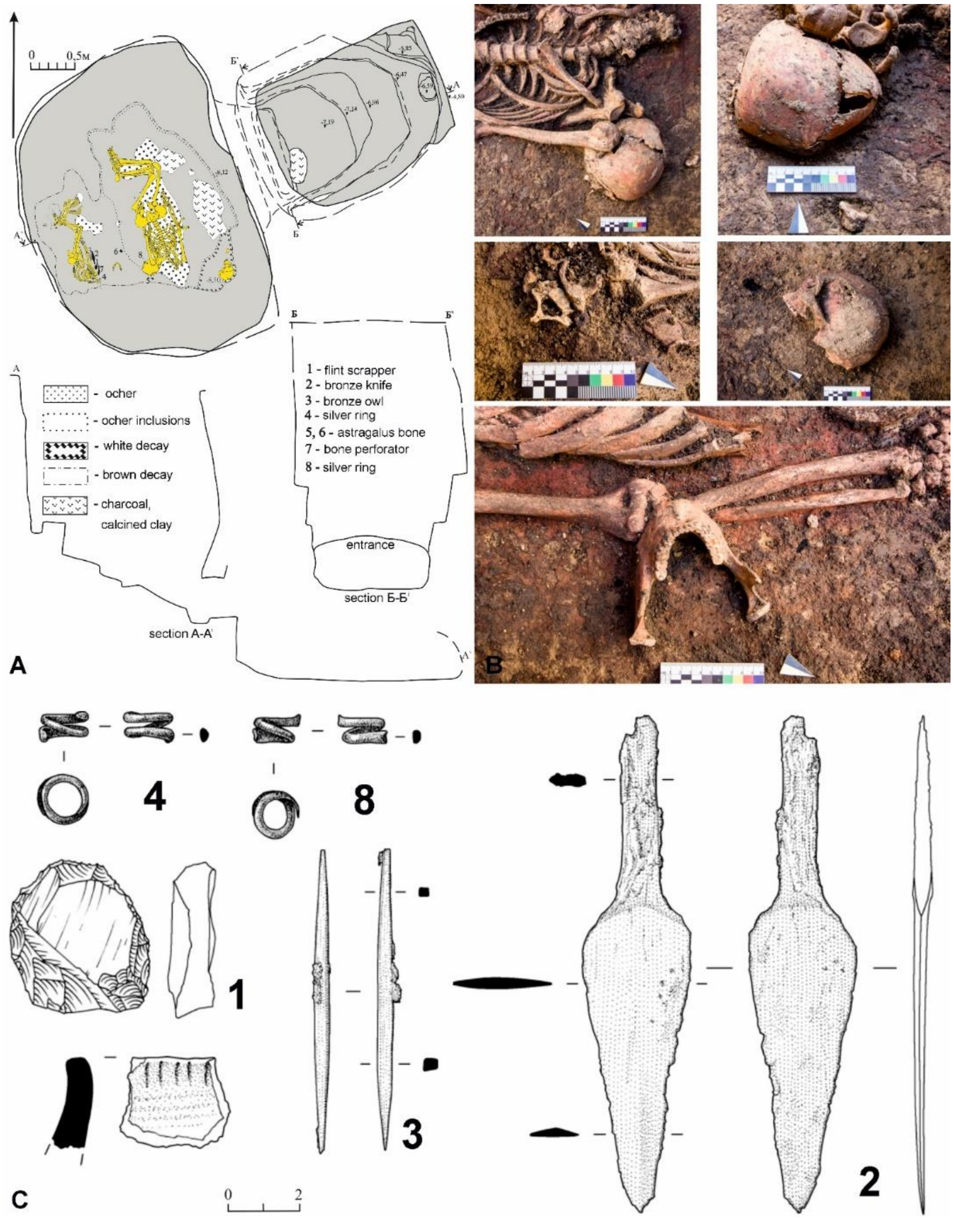

**Fig. 6.20. Burial 26. A – general plan, B – details, C – grave goods (see A for annotaitions)** (image by Aleksey Kiyashko).

*6.7.1.2. Grave 1:42 (individual ID I23656)*

Grave 1:42 was inserted in the northeast sector of the barrow at a distance of 17.25 m from the center of the kurgan (FIG). The dimensions of the main shaft or pit were 2.17 x 1.29 m at its entrance, and depth beneath the top of the kurgan to the floor of the shaft was 7.35 m. There was a step measuring .77 X 1.06m that led down from the floor to the side chamber at a depth of 7.63 m beneath the top of the kurgan. In the SE and NE corners of the shaft, three ledges were traced, serving as supports for descent/ascent into the burial. At the lower level of the walls on the SE and NE sides, there were protruding niches widening towards the entrance to the burial chamber. The entrance to the burial chamber was in the western wall of the shaft. The lower parts of the niches formed smaller steps. A gentle slope from the eastern side to the western side formed a passage to the chamber. The burial chamber had an oval shape, elongated along the SW-NE line. A catacomb was dug out with the following dimensions: burial chamber 2.31 x 2.71 m, bottom level – 7.78 m from the top of the kurgan.

In the central part of the chamber, two skeletons were placed – an adult identified osteologically as a female, but genetically as a male; and an infant (possibly a newborn), both aligned along the SW-NE line, with their skulls to the SW. Only the adult was sampled for aDNA. As was discussed above, this male was a brother of the two males in Grave 1: 26; all had Y-haplogroup I2a.

<u>Individual 1 (infant)</u> was closer to the entrance, nestled against the chest of skeleton 2. It lay on its right side, with the skull separated and placed at chest level. The arms were straight, alongside the torso. The left arm was on the ribs towards the pelvis. The legs were together, slightly bent at the knees, with the knees to the SW.

<u>Individual 2 (individual ID I23656)</u> This individual lay on the right side, with the skull separated and resting on the right shoulder, with the eye sockets facing downwards. The arms were straight, alongside the torso. The left arm was slightly bent at the elbow, with the hand on the right iliac bone. The legs were slightly bent at the knees, with the knees to the SW, and together. The skeletons lay on a rectangular plant bedding of dark brown color with abundant ochre inclusions. At the entrance to the chamber, on the eastern side, traces of a burnt rectangular board were found.

Inventory:

Near Individual 2, to the E, in a row aligned W-E, three items were found: a white elongated flint nodule, a bronze chisel, and a bronze axe. Between Individual 1 and the entrance were a bronze leaf-shaped knife, a bronze awl, three bone piercers, and four flint flakes. The items were laid out compactly, suggesting they were packaged in a pouch. All tools pointed southward. Between the skull of skeleton 2 and the flint nodule was a flint flake. At the northern wall corner, on a burnt board, an ornate clay vessel was placed.

The burial dates typologically to the early stage of the Catacomb Culture of the MBA.

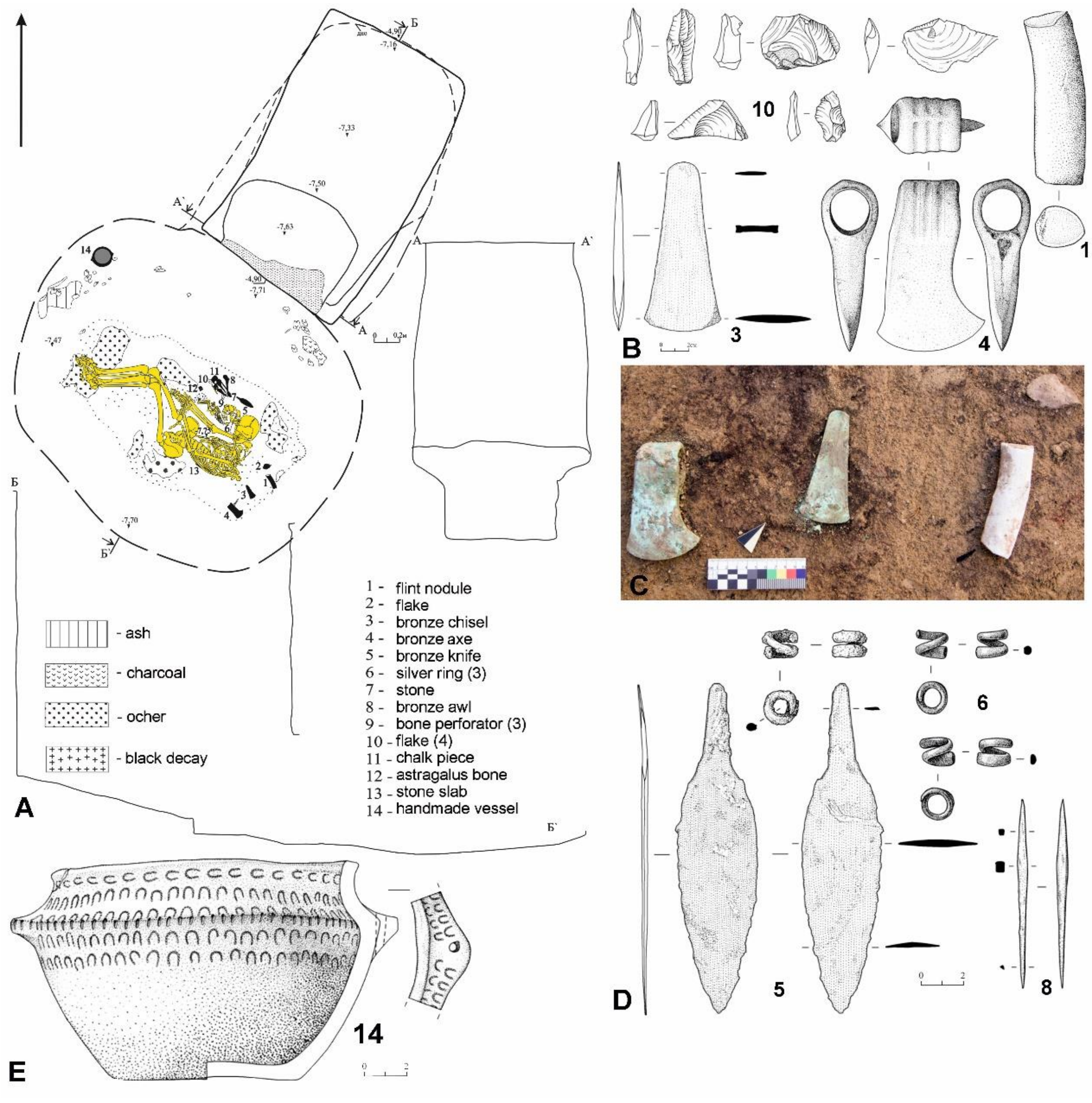

**Fig. 6.21. Burial 42. A – general plan, C – a detail of the burial assemblage, other – grave goods (see A for annotaitions)** (image by Aleksey Kiyashko).

<u>*Bibliography to section 6:*</u>

*Palalidis, S. (2023). A GIS-Based approach to the study of the Yamnaya cultural horizon (ca. 3300-2400 BC): Preliminary results. In InFieri Incontri di Archeologia Sapienza Miscellanea degli atti II (2018-2019) e III (2020), pp. 371-384.*

### 7. THE BALKANS, LOWER DANUBE, PRUT, AND DNIESTER BASINS

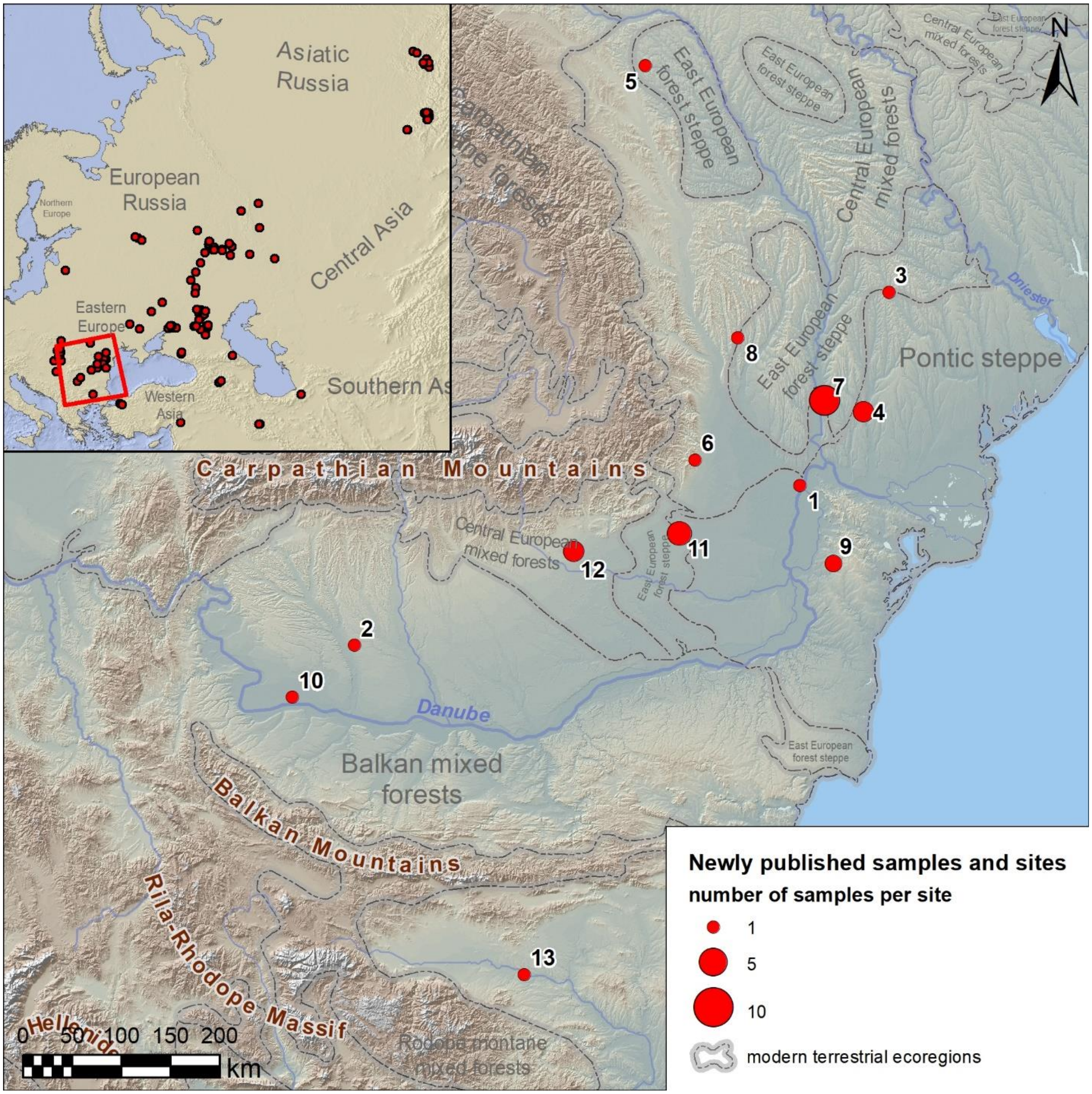

**Fig. 7.1. Newly published samples from the Danube area.** 1 - Brăiliţa (Romania, Brăila County, Brăila); 2 - Cârcea (Romania, Dolj County, Cârcea); 3 - Cimișlia (Moldova, Cimișlia District); 4 - Ciumai (Taraclia District); 5 - Corlăteni (Botoșani County, Corlăteni); 6 - Coțatcu (Buzău County, Podgoria); 7 - Crihana-Veche (Cahul District); 8 - Glăvăneşti (Romania, Bacău County, Glăvăneşti); 9 - Rahman (Romania, Tulcea County, Casimcea); 10 - Rast-Măgura-Barburlui (Romania, Dolj County, Negoi); 11 - Smeeni-Movila Mare (Romania, Buzău County, Smeeni); 12 - Târgșoru-Vechi (Romania, Prahova County, Târgșoru Vechi); 13 - Yabalkovo (Bulgaria, province Haskovo, municipality Dimitrovgrad).

#### 7.1. Smeeni

*Summary by A. Frînculeasa & A. Simalcsik*

##### *7.1.1. Smeeni-Movila Mare*

The mound of Smeeni-Movila Mare (Buzău County) was researched by archaeologist Victor Teodorescu in 1959. Partly destroyed by recent excavations, the mound was 4 m high and 55×49 m in diameter. Thirty-four graves were distributed over six chronological phases. According to the burial ritual, C14-AMS dates and stratigraphy, the first two phases were Yamnaya graves (3050-2650/2600 cal BC), the third– Catacomb-culture graves (2650/2600-2400 cal BC) and the fourth – Middle Bronze Age graves (2150-1750 cal BC); the last two phases were assigned to the Sarmatians (the 2nd-3rd century AD) and to the Late Medieval period (the 18th century).

Yamnaya graves were the most numerous. Their specific characteristics were a rectangular pit with a wooden cover, supine with raised knees body position, west-east orientation, and ochre deposition. Very few had pottery. In one grave three bone pendants were discovered. The askos pot from Grave 13 is an important chronological marker and shows interaction between local and foreign people.

As for the Catacomb graves, the four dates from Smeeni are up until now the only ones available. Catacomb-attributed burials are rather discreet occurrences at the Lower Danube. At the same time, absolute dates for all these graves show in the Lower Danube region a stage of coexistence with the Yamnaya burials.

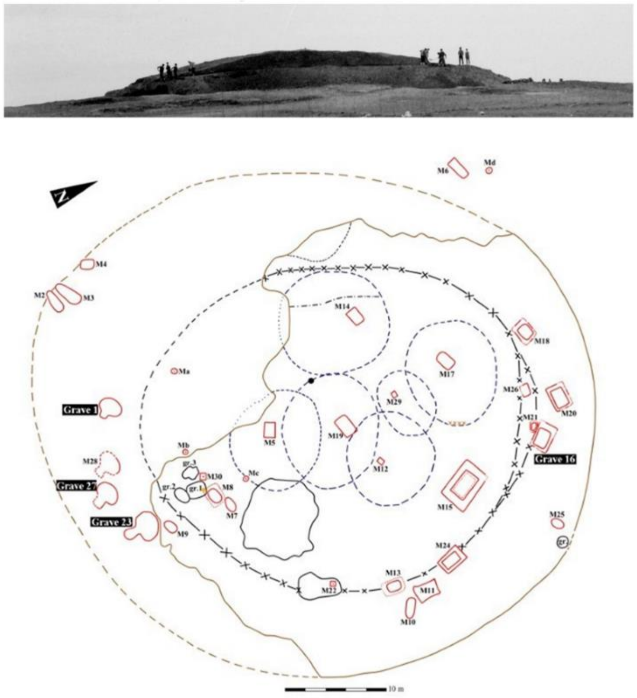

**Fig. 7.2. Smeeni: image of the Movila Mare during archaeological research and the layout of graves on the general plan (image by Alin Frînculeasa).**

*7.1.1.1 Grave 1 (individual ID I12823)*

The deceased was laid in a hole dug deep into the natural ground, consisting of two distinct parts, the access shaft and the arched room (the catacomb proper), both partially destroyed; the first segment was preserved only to a depth of 0.20 m, while the catacomb, almost 1 m. Viewed from the side, Gr. 1 hole has two steps, the first (upper one) represented by the shaft and the second (lower one) marking the catacomb. The maximum depth of the shaft to the current preserved level, which is lower than the ancient level where the funeral hole was dug, was 1.35 m, while that of the catacomb in relation to the same reference point was 1.91 m. The shaft was 1.09 m in diameter, while the catacomb was 2.08 m long and 1.33 m wide. The base of the shaft was +0.56 m higher than the lower part of the catacomb. The arched room plan was bean-shaped. The walls were leaning towards the inside, except for a small portion next to the shaft which was inclined towards the outside. Its bottom was slightly inclined towards the access shaft.

It was in this room, closer to the northern side, that the body was placed, lying on the back, head to W-SW 240°, lower limbs E-NE 60°. The left arm was stretched along the body, the right arm, bent at the elbow, was resting on the hip. Under and around the skeleton traces of brownish-red vegetal bedding 6-9 mm thick were identified.

Anthropological determinations: female, 55-60 years old, but genetically determined to be a male; gracile skeleton; proto-European phenotypic traits; skeletal stature of approx. 165 cm; dental pathologies; torus auricularis; musculoskeletal stress markers; periosteal bone reactions; double fracture on the left ulna produced antemortem; unintentional reshaping of the cranial vault (flattening of the occipital). C14-AMS date: (DeA-5392) 4008 ± 42 BP/2834 - 2457 cal BC, 95.4% probability. The burial is attributed to the Catacomb culture. His Y-haplogroup was R1b (R-L51), mt-haplogroup K1b2b.

*7.1.1.2 Grave 16 (M. 16) (ID I10891)          2875-2585 calBCE (4142±30 BP, DeA-7737)*

Grave 16 was assigned to Yamnaya phase II. The grave had been covered with a small mound, much flattened. The grave pit was wider at the top, the upper chamber being 2.60×1.80 m in size. Below a step the lower chamber was 2.03×1.38 m in size, rectangular with slightly rounded corners.

The deceased was laid supine, with lower limbs flexed, knees raised and fallen leftwise; the head was oriented to west – north-west, while the lower limbs to east – south-east. To the right of the skull, above the left shoulder, a red ochre lump was found. The deceased had been laid on a vegetal bedding. The individual in Grave 16 was distinguished by intentional cranial deformation.

The grave inventory included a flint blade.

Anthropological determinations: male, 35-40 years old; robust skeleton; proto-European phenotypic traits; skeletal stature of approx. 176 cm; very good dental health; metopism; supplimentary sutural ossicles; porotic hyperostosis; periosteal bone reactions; musculoskeletal stress markers; artificial/intentional circular (parieto-occipital) cranial deformation of medium intensity achieved with a cephalic device. His Y-haplogroup was R1b (R-M12149), mt-haplogroup U4c1.

*7.1.1.2 Grave 23 (individual ID I12824)*

This is the best preserved feature; it can be well determined by excavation, both in plan and in section. The access shaft, oval in plan, sized 1.40x1.05 m, preserved only 1.06 m of its depth. On the wall opposing the arched room, there was a small step approximately 0.70 m above the base.

On the same side, a slight panning on the bottom of the shaft indicates the existence of a pole which could probably support the cap sealing the mouth of the catacomb. The access shaft and the arched room were connected through a short “neck” (0.18-0.20 m), which in transverse section is quasi-ogival in shape, with the base slightly curved on the outside (H: 0.98 m, base: 0.87 m). In the plane, the catacomb is kidney-shaped, while in longitudinal section it appears as a triangle with much rounded corners. The arch is irregular (probably due to caving-in as well) and is centred only longitudinally, whereas transversally the “capstone” is closer to the end towards the mouth and the shaft. The irregular catacomb bottom was inclined from the shaft towards the catacomb. Length of catacomb: 2.56 m, width: 1.87 m, maximum height: 1.39 m. The filling of the shaft differs from that of the funerary feature, which may have been related to the existence of a possible cap sealing the entrance to the burial chamber.

The deceased was lying in the catacomb, closer to the wall opposing the entrance, head towards W-NW 281° - legs to E-SE 101°, lying on the back, face oriented to the entrance/south and arms stretched along the body, lower limbs in the runner position to the right. The remains of a vegetal bedding 2-5 mm thick could be identified under the skeleton. A mammalian bone, probably the remnant of an offering deposed about 40 cm in front of the dead, was found in the grave.

Anthropological determinations: probably female, 30-35 years old, confirmed female genetically; moderately robust skeleton; skeletal stature of approx. 161 cm; proto-European and Asian/Mongoloid phenotypic traits; very good dental health; a rib fracture produced antemortem; musculoskeletal stress markers. C14-AMS date: (DeA-5864) 4027 ± 31 BP/2623 - 2472 cal BC, 95.4% probability. The burial is attributed to the Catacomb culture, mt-haplogroup U4c2.

*7.1.1.2 Grave 27 (individual ID I12825)*

The grave was found in the same southern part of the mound, -1.40 m deep from the current preserved level. Modern clay mining strongly affected it, completely destroying the shaft, while in the arched chamber only the base was preserved approximately 10-30 cm deep. The arched chamber, or whatever was kept of it, in the centre of which the deceased was deposed, is bean-shaped (2.16x1.51 m), with the southern side more curved than the northern one.

The dead was placed on his back, head to the W 270° and lower limbs slightly bent at the knees to the left to the E 90°, face oriented to the NE. Arms were stretched along the body, palms resting on the hip, right arm somewhat bent at the elbow. The entire body was slightly oriented towards the north. Under the skeleton, the remnants of a vegetal bedding 2-3 mm thick were identified. Nearby the right knee a bone from a large mammal seems to mark the presence of an offering.

Anthropological determinations: male, 50-55 years old, confirmed genetically; very robust skeleton; skeletal stature of approx. 171 cm; proto-European phenotypic traits; without dental pathologies; torus mandibularis; musculoskeletal stress markers; pronounced osteoarthritis; unintentional reshaping of the cranial vault (flattening of the upper region of the occipital). C14-AMS date: (DeA-5393) 4039 ± 42 BP/2848 - 2468 cal BC, 95.4% probability. The burial is attributed to the Catacomb culture. Y-haplogroup R1b (M-269), mt-haplogroup T1a1.

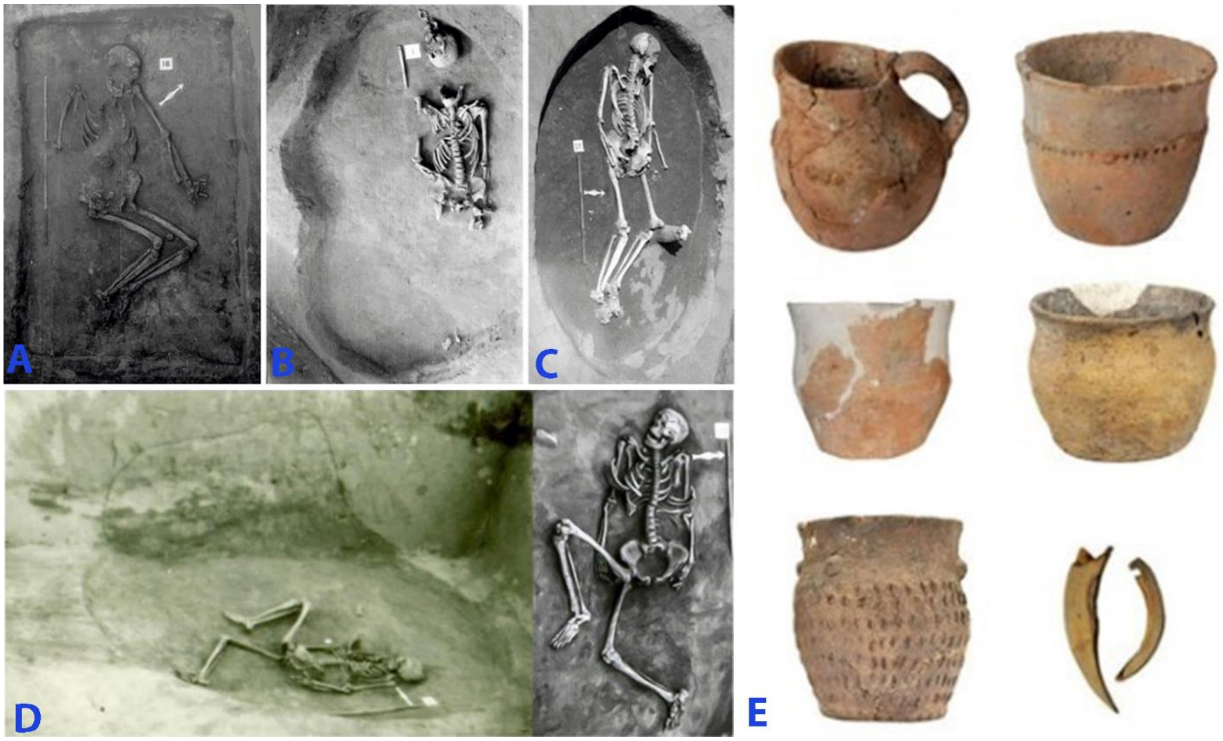

**Fig. 7.3. Smeeni-Movila Mare: A - grave 16; B - grave 1; C - grave 27; D - grave 23; E - goods from Yamnaya graves (image by Alin Frînculeasa).**

#### 7.2. Târgșoru Vechi

*Summry by Alin Frînculeasa, Octav Negrea, Mădălina Nicoleta Frînculeasa*

Târgșoru Vechi is a village located in the southern Romanian plain, in Prahova County, in the metropolitan area of Ploiesti. 'The Mound of the Beizadele Forest', 32 m in diameter and 1.10 m high, was investigated in 2016. Twelve graves contained 16 individuals. Three graves were assigned to the Yamnaya culture, two being double graves (Grave 2 and Grave 10) and one individual (Grave 9). A cremation grave (Grave 12) most likely dates from Iron Age and Grave 8 from the Medieval period (sec. VIII-IX AD). Also, 6 archaeological features dating from the end of the first millennium, assigned to Dridu culture, were discovered.

In addition to Graves 2 and 9, we mention here Grave 10A, not sampled for aDNA, which belongs to a subadult individual, in supine position with flexed legs. A necklace consisting of 7 perforated dog/fox canines had been put around the neck of the deceased. C14-AMS data (DeA-10668) 4140±33 BP/2875-2620 cal BC, 95,4% probability.

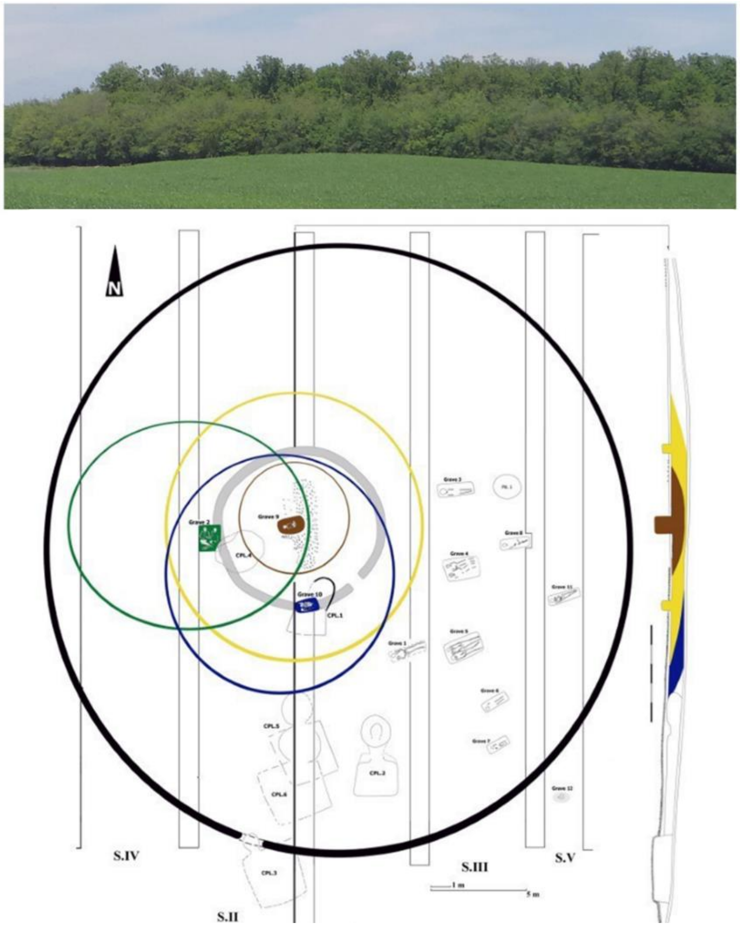

**Fig. 7.4. Târgșoru Vechi: mound before excavation, general plan and eastern stratigraphic profile of trench 2 (image by Alin Frînculeasa).**

*7.2.1 Grave 2, individuals A (ID I18818 ) and B (ID I18819)*

Grave 2 was a secondary grave with a rectangular pit (1.47×1.20 m). In the pit, two individuals were lying in supine position with flexed legs, fallen to the left. They were N-S oriented. Both individuals were adult males (aged 43-45 and no more than 30, respectively), with no grave goods, except an ochre lump placed near the skull of individual B. C14-AMS data/Individual A: (DeA-10666) 4176±32 BP/2886-2636 cal BC, 95,4% probability. The grave was partially destroyed by a medieval feature.

Individual A had the unusual Y-haplogroup P1 (P-P337), mt-haplogroup K1c1. Individual B also had P1 and K1c1. They were 1st-degree relatives, probably brothers.

*7.2.2 Grave 9 (individual ID I18820)*

Grave 9 was primary, located under the center of the mound. It had a rectangular pit with rounded corners, wider at the top and narrowing below a distinct step inward part way down the shaft. The pit was 0.90-1 m deep, 1.33×0.82 m in the upper part and 1.21×0.65 m in the lower part. A subadult was in supine position, with her arms stretched along her body and flexed legs initially raised then fallen to the left. An ochre lump was placed south of the skull. Due to the ochre, the skullcap was red; sprinkled ochre were also present on other bones.

This grave was surrounded by a circular ditch with a diameter of about 9.2 m, width of 0.40-0.45 m and depth of approximately 0.40 m. It had an entrance-break with an opening of 0.65 m in the south-south-east.

Anthropological determination: indeterminable, estimated age of 6 (genetically female H2a1). C14-AMS date: (DeA-10667) 4123±33 BP/2871-2579 cal BC, 95,4% probability. This female child was a 3rd-degree relative of I18819 in grave 2.

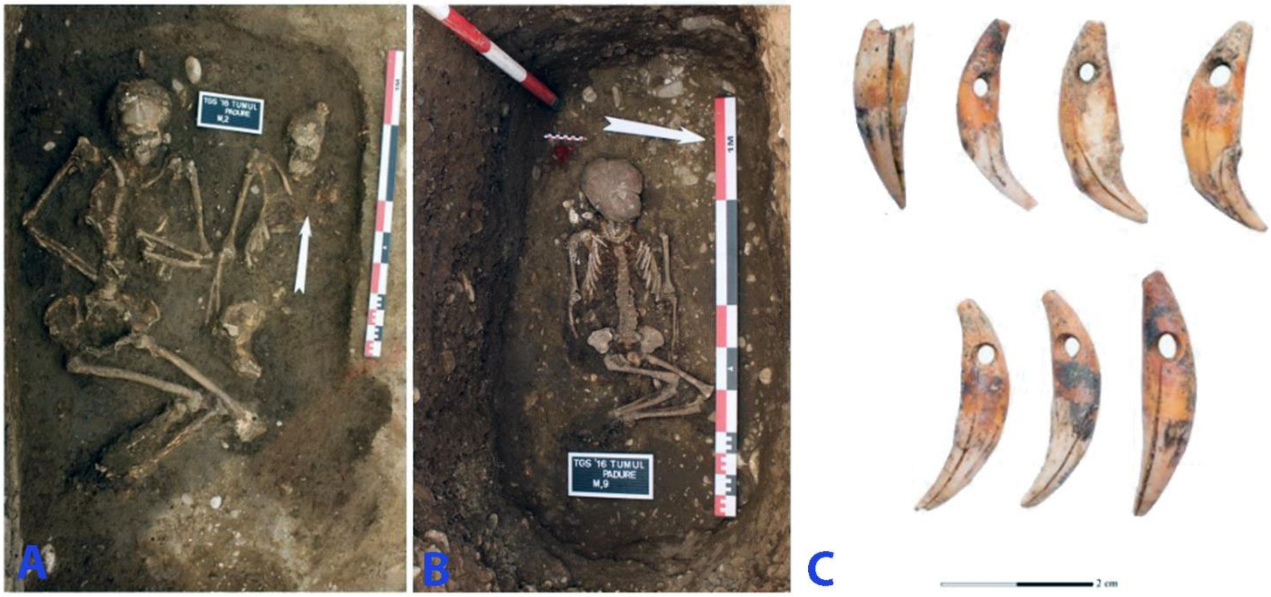

**Fig. 7.5. Târgșoru Vechi, Mound of the Beizadele Forest. A - grave 2, B - grave 9, C - Canine necklace found in grave 10 from Târgșoru Vechi (image by Alin Frînculeasa, Octav Negrea, and Mădălina Nicoleta Frînculeasa).**

#### 7.3 Rahman

*Summary by Sorin-Cristian Ailincăi & Mihai Constantinescu*

The village of Rahman is located in the north-western part of Dobrogea, in a hilly area near the Danube. The area of Casimcea - Topolog is characterized by a particular density of the tumuli. During field research over the last decade, over 1300 mounds have been identified, unfortunately heavily damaged by agricultural works. Although there is little research, it seems that the oldest mounds date back to the Eneolithic period, as indicated by the finds from Casimcea (Popescu 1941).

In 2011 and 2013 two tumuli were investigated west of Rahman (Ailincăi et al 2014; Ailincăi et al 2016). These were part of an alignment oriented along the NNE-SSV direction.

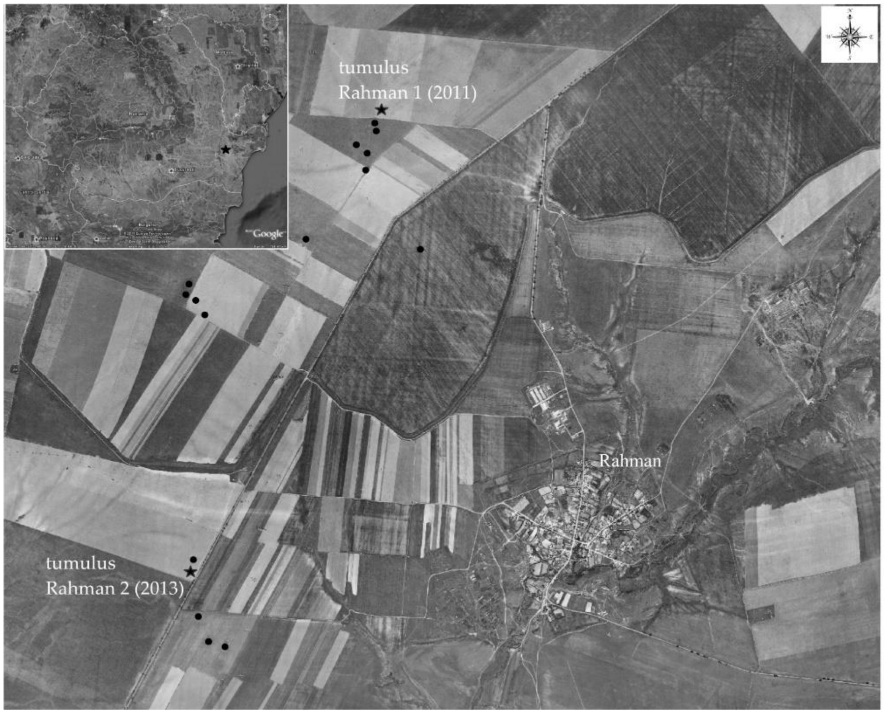

**Fig. 7.6. Burial mounds at Rahman (image by Sorin-Cristian Ailincăi and Mihai Constantinescu).**

##### *7.3.1 Rahman, kurgan 2*

Mound 2 at Rahman after reduction by agricultural work was 40 m in diameter and 1 m high. It contained five graves, with six individuals, grave 5 being double. Of these graves graves 1 and 2 can be considered secondary, and graves 3-5 can be considered primary as their pits were dug in the ancient level.

Graves 2-5 show several elements typical of the Yamnaya package, such as rectangular pits, sometimes with a step, traces of textile cover on the bottom of the pit, the use of ochre and wooden beams. The way in which the deceased were buried (position and orientation) is also specific to this phenomenon. It is possible that Grave 1 dates from a later period, probably from the 2nd millennium BC.

Unfortunately, the only grave dated with 14C is grave 5 (double burial). A first date was obtained from a sample taken from individual no. 5 (Poz-65968: 3950±35 BP; 2570-2330 calBC). A second dating obtained from a sample of wood from the same grave (RoAMS 1370.46: 4196±32 BP; 2894-2672 cal BC) shows a much earlier range (Ailincăi et al 2021). Only in grave no. 3 were identified two loop rings made of copper and encased in silver sheet.

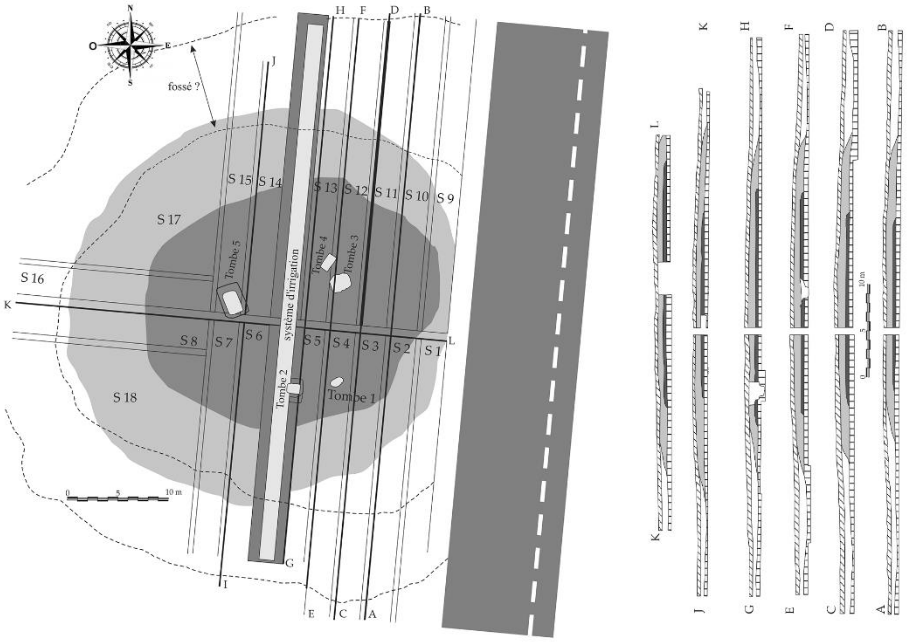

**Fig. 7.7. Kurgan 2 at Rahman, general plan and stratigraphic profiles (image by Sorin-Cristian Ailincăi and Mihai Constantinescu).**

##### *7.3.1.1 burial 4 (individual ID I10498)*

Grave 4 (individual 4) was anthropologically determined as female, 43-58 years old, while grave 5 (individual 6) is a 24-30 years old male (Ailincăi et al 2016).

Grave pit of burial 4 was rectangular in shape and had dimensions of 1.67 × 1.05 m, approximately 1 m depth. The age of the genetically confirmed female individual (individual 4) was estimated at around 43–58 years, and her height was 152.23±3.4 cm. The skeleton was placed at the base of the grave in a contracted position, on her right side, and oriented along south-southeast-north-northwest axis. No traces of pigment were observed on the body. Her mt-haplogroup was U4a.

In the infill of the grave, several animal bone fragments (probably birds and rodents) and a flake without traced of processing (L = 1.8 cm; w = 2.6 cm; thickness = 0.9 cm) were unearthed. The flake was made of a white, opaque flint of fine grain size.

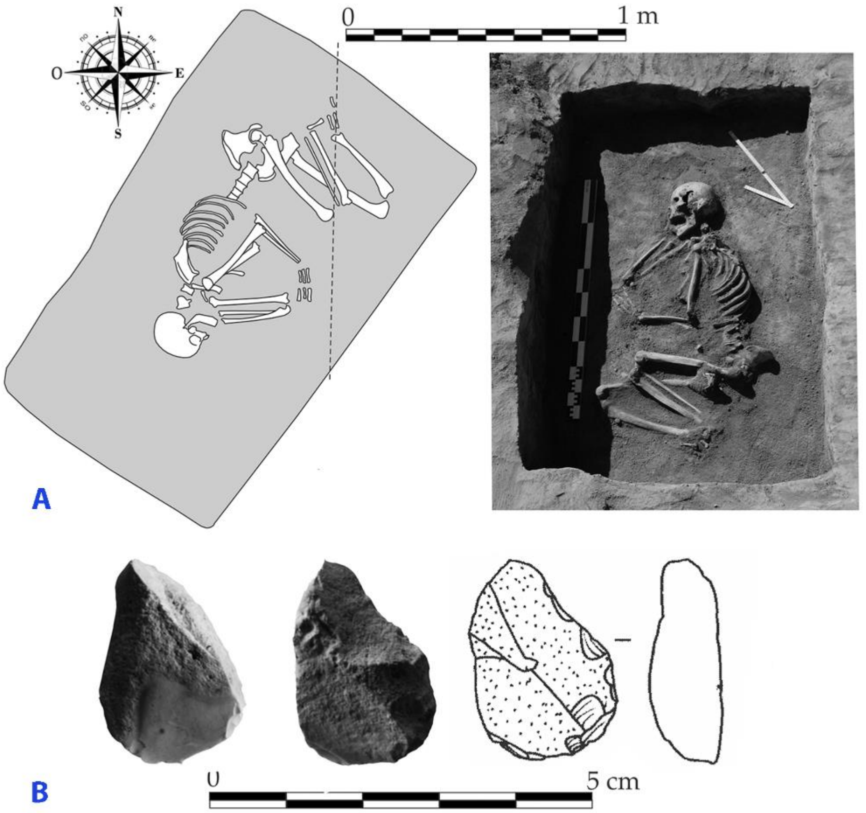

**Fig. 7.8. Rahman, kurgan 2, burial 4. A – a plan and a photo of the burial4 B – flake from infill of the grave (image by Sorin-Cristian Ailincăi and Mihai Constantinescu).**

##### *7.3.1.1 burial 5 (individual ID I10499): 2896-2677 calBCE*

Grave 5 was excavated in two phases. Initially, a large pit measuring 3.73 × 2.88 m was uncovered. Subsequently, a smaller rectangular pit (2.47 × 1.80 m) intersecting the first structure was revealed. Within the latter, the remains of two individuals (designated as individuals no. 5 and no. 6) were discovered. The pit was lined with wooden stakes, and at its base, remnants of a mat stained with red ocher were found. Both bodies were positioned in a contracted posture on the right side, aligned along a northwest to southeast axis. Although the bones were robust, their preservation was poor, with sporadic red pigment observed on some of them. Individual 5 was a male, Y-haplogroup R1b (R-Z2103) and mt-haplogroup T2c1a.

Individual 6, not sampled, displayed intense red pigmentation on the skull, with other bones also exhibiting ocher residue, particularly on the hands and feet. Traces of animal teeth were found on the distal third of the right humerus shaft.

The tomb yielded three bone fragments of animal origin and a fragmentary tooth. Furthermore, a rib fragment (likely left) of individual no. 6 near the sternal end showed a bulge and signs of infection, indicating healing. The grave goods are represented by a flake (L = 3.4 cm; w = 36 mm; thickness = 0.4 cm) made from brown raw material with white inclusions.

The burial is dated to 2896-2677 calBCE (4205±25 BP, PSUAMS-7907).

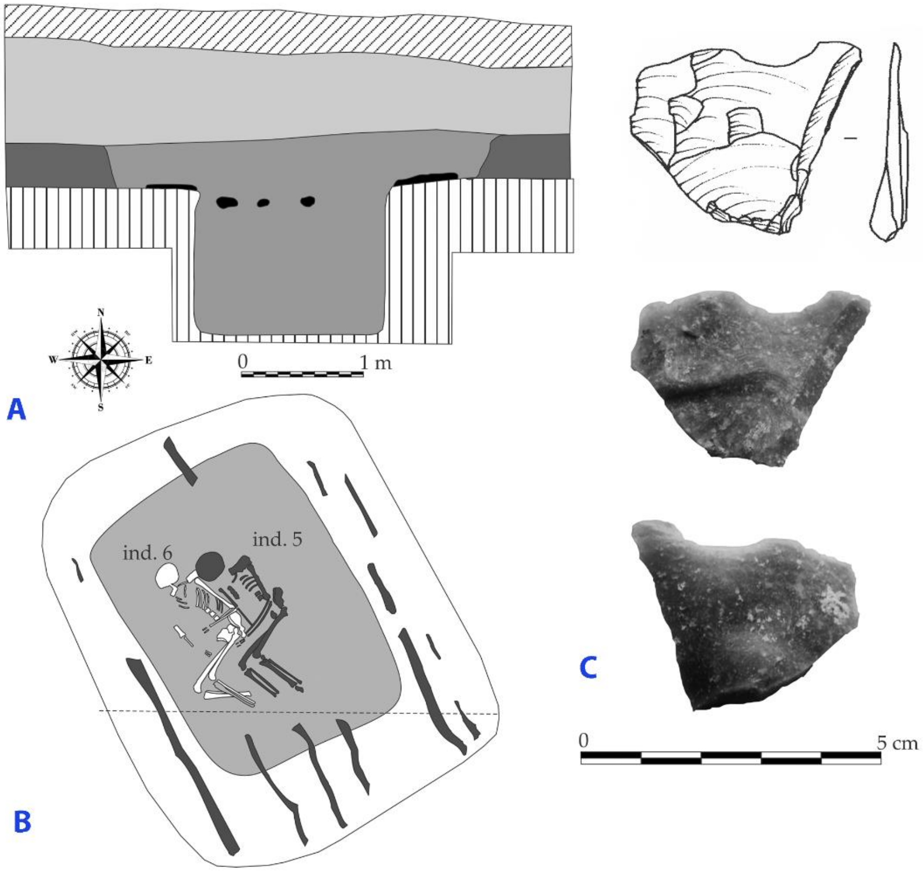

**Fig. 7.9. Rahman, kurgan 2, burial 4. A – stratigraphic profile; B – plan of the burial; C – flake from the grave (image by Sorin-Cristian Ailincăi and Mihai Constantinescu).**

#### 7.4. Crihana Veche

*Summary by A. Simalcsik & I. Ciobanu*

Thousands of burial mounds dot the Lower Prut Plain, but only a few have been investigated using archaeological excavation. Most of the investigated mounds are located in the central and southern regions of this area. The locality that stands out in this respect is Crihana Veche (Cahul District), where 17 mounds have been researched so far. Some of these mounds were built by the Eneolithic or Early Bronze Age populations, while others by Scythian migrants.

##### *7.4.1 Crihana Veche, tumulus 5*

Tumulus 5 from Crihana Veche is located at a point called La Pietricei. The mound, situated approximately 1.95 km north of Cahul International Airport, was investigated in 2016 by archaeologists from the National Archaeological Agency of the Republic of Moldova. It was approximately 0.70 m high and had a diameter of 17 m. Over time, the mantle of this mound merged with that of a neighboring one, resulting in a figure-eight-shaped elevation. Five inhumations were uncovered. The first mantle of the tumulus is directly related to grave no.4/M.4 (Yamnaya). Subsequently, grave no.3/M.3 (Yamnaya) was arranged in the same mantle. The second mantle was erected over grave no.2/M.2 (Scythian). Later, in the third mantle, graves no.1/M.1 and no.5/M.5, both Scythian (4th century BC), were arranged.

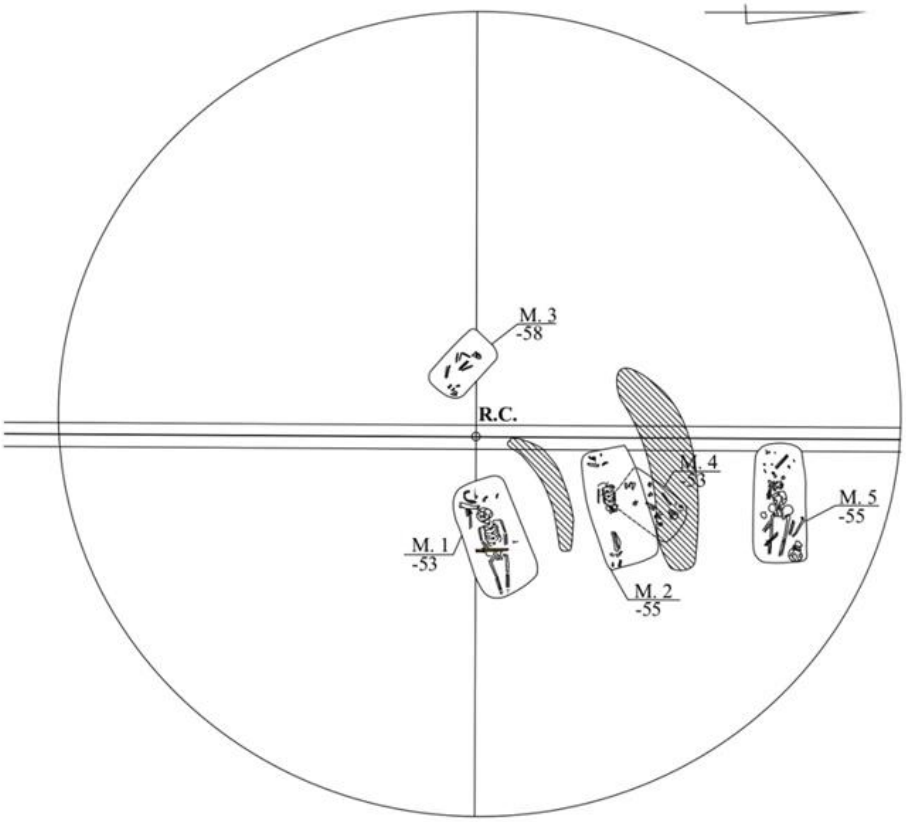

**Fig. 7.10. The general plan of the tumulus 5 from Crihana Veche**

##### *7.4.1.1 Grave 3 (individual ID I10414): 2876-2623 calBCE*

The grave was situated in the western sector of the tumulus, 0.8 m away from the central landmark (R.C.), at a depth of 0.58 m. The grave pit had a rectangular shape with rounded corners and was oriented northwest-southeast. It was identified at a depth of 0.55 m and measured 1.5 × 0.9 m. The skeleton had been partially disturbed by agricultural activities.

The deceased was oriented with the head to the northwest, laid in a crouched position on the right side on the bottom of the pit. The skull was tilted to the right, facing south. The right upper limb was strongly bent at the elbow, while the left one had fallen away from the bones of the ribcage at an angle of about 90º. Only the right tibia from the lower limbs was preserved. The skeleton exhibited red ochre pigmentation, more intense on the parietals and the inner face of the mandible and less distinct on the postcranial bones. Brown traces from the plant layer covering the pit bottom were found near and beneath the skeleton. No grave goods were present.

The human bone remains showed partial anatomical connection. The skeleton is incomplete, poorly preserved, and gracile, and displayed barely visible muscle insertions. Anthropological sex: female. Molecular sex: female with mt-haplogroup H6a1a. Biological age at death: about 20-30 years (young adult). Skeletal stature: middle category. Epigenetic traits: congenital absence of the third molar; torus mandibularis. The burial is attributed to Yamnaya culture and dated to 2876-2623 calBCE (4145±30 BP, PSUAMS-7832).

##### *7.7.1.2 Grave 4, individual A (ID 10415): 2867-2573 calBCE*

The grave was situated in the northeastern sector of the mound, 3.2 m away from the central landmark (R.C.), at a depth of 0.53 m. The grave pit had a roughly rectangular shape with rounded corners and was oriented to the north-northeast – south-southwest. Its dimensions measured 1.65 × 0.95 m, with a depth of 0.95 m from the identification level. Fragments of decayed wood from the burial pit cover and human bone fragments were identified in the fill of the feature. The grave had been disturbed in ancient times.

Both sub-adult individuals were interred at the bottom of the pit, with their heads facing northeast and skulls placed face up. The skeletons exhibited crimson ochre pigmentation, with a higher concentration on the skulls, particularly the frontals. Grave goods: two canines of *Canis familiaris*, a mature individual, were found on the bottom of the grave pit, in the northern corner, near the skull of <u>skeleton A (individual ID 10415)</u>. His Y-haplogroup was R1b (R-M269).

The remains of individual A were partially anatomically connected, with the skeleton being incomplete, partially represented, and poorly preserved. Anthropological sex: indeterminate. Molecular sex: male. Biological age at death: about 9 years ± 24 months (child). Pathologies: active porotic hyperostosis (*cribra cranii, cribra orbitalia*). The burial is attributed to Yamnaya culture and dated to 2867-2573 calBCE (4110±30 BP, PSUAMS-7833)

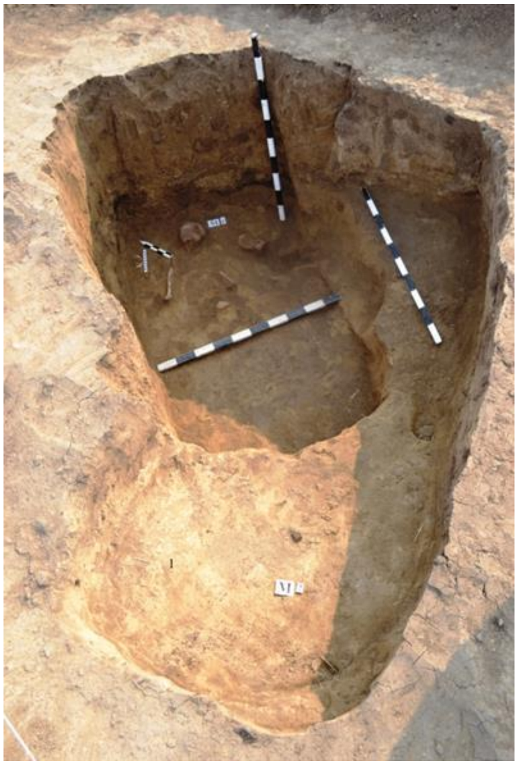

**Fig. 7.11. Crihana Veche, tumulus 5, grave 4.**

##### *7.4.2 Crihana Veche, tumulus 9*

Tumulus 9 from Crihana Veche was situated on the first terrace of the Prut River. It was investigated in 2015 by archaeologists of the National Archaeological Agency of the Republic of Moldova. At the beginning of the research, the mound measured approximately 24 × 27 m with a height of about 1.25 m. Twenty inhumation graves and a cult pit were discovered. In the central part of the mound, at a depth of 1.20 m from the central landmark (R.C.), the prehistoric surface level was identified. The construction of the tumulus revealed three cultural-chronological layers: the Late Eneolithic phase, the Hadžider-Cernavodă-I culture (M.10); Early Bronze Age (Yamnaya), comprising graves nos. 4, 6, 17, 20 from the early Yamnaya stage and nos. 3, 5, 8, 9, 19, and 21 from the late Yamnaya stage; Late Bronze Age phase (Multi-Cordoned Ware and Sabatinovka), including graves nos. 2, 11, 12, 13, 14, 15, 16, and 18. To these three phases is added a grave likely attributed to the Sarmatians from the first centuries of the new era.

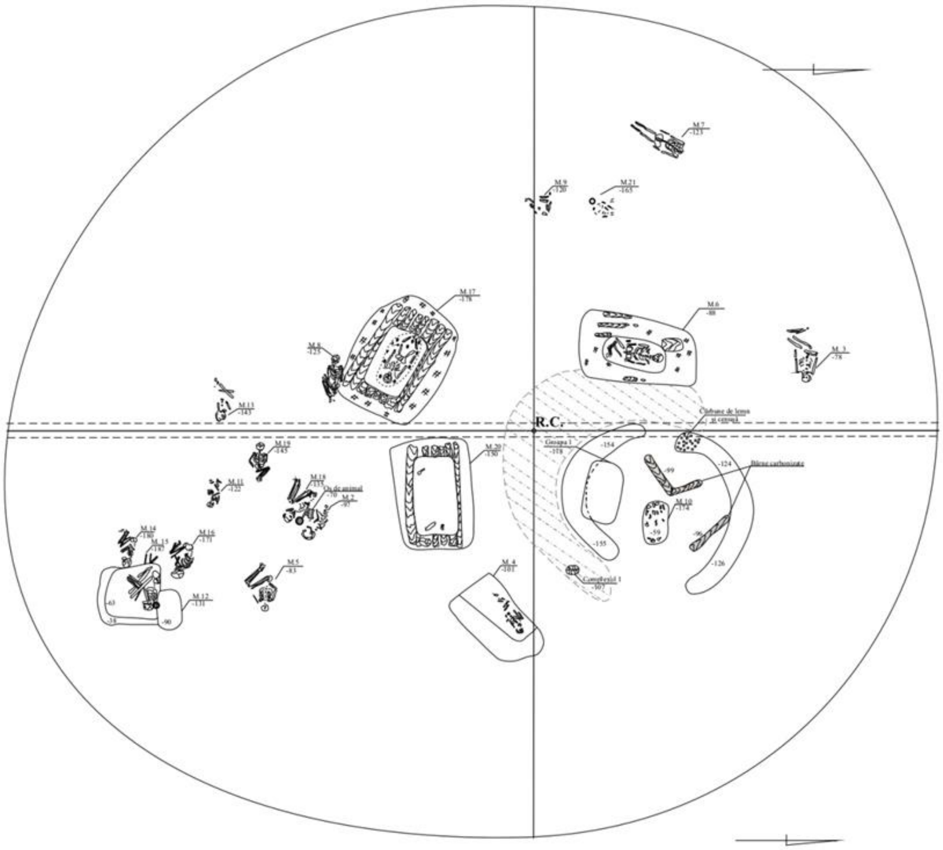

**Fig. 7.12. The general plan of the tumulus 9 from Crihana Veche**

###### *7.4.2.1 Grave 9:9 (individual ID I10174): 2848-2494calBCE*

The grave was situated in the western sector of the tumulus, approximately 6.35 m away from the central landmark (R.C.), at a depth of 1.20 m. The grave pit was not identified. The deceased was laid in a crouched position on the left side, with the head oriented towards the south-southwest, the right arm bent at the elbow, and the hand resting on the femur of the right leg. However, the left hand was missing. The legs were strongly contracted on the left side. Traces of red ochre pigments were found on the skeletal elements of the chest and upper limbs.

The skeleton, which was poorly represented, extremely fragmentary, and poorly preserved, exhibited gracile features with attenuated muscle insertions. Anthropological sex: female. Molecular sex: male. His Y-haplogroup was R1b (R-M269). Biological age at death: about 30-40 years (young to middle-aged adult). The burial is attributed to Yamnaya culture and dated to 2848-2494calBCE (4075±25 BP, PSUAMS-11047).

##### *7.4.3 Crihana Veche, tumulus 12*

Tumulus 12 from Crihana Veche, known locally as Movila Gologan, was investigated in 2016 by the National Archaeological Agency of the Republic of Moldova. Situated on the second terrace of the Prut River, the mound measured approximately 1.25 m in height and had a diameter of about 28 m. During the excavations, 23 graves and three ritual features were discovered. At a depth of 1.40 m from the central landmark, the prehistoric surface level was identified.

Mantle I was raised above grave no.10/M.10 (Late Eneolithic, Cernavodă-I). Mantle II is associated with grave no.1/M.1 (early Yamnaya). Additionally, graves no.5/M.5, no.12/M.12, and no.9/M.9 also belong to this secondary mantle. The two ritual features containing dispersed ceramic fragments are likely connected to this horizon. Mantle III was raised above grave no.16/M.16 (late Yamnaya). Graves no.3/M.3, no.4/M.4, no.6/M.6, no.8/M.8, no.15/M.15, and no.21/M.21 were also arranged within mantle III. Mantle IV was erected above grave no.13/M.13 (Multi-Cordoned Ware), with grave no.14/M.14 being subsequently dug into this mantle. Furthermore, within this fourth addition to the mound, Late Bronze Age graves no.2/M.2, no.18/M.18, no.19/M.19, and no.20/M.20 (Sabatinovka) were arranged. The Antique chronological horizon is represented by two graves, no.17/M.17 and no.22/M.22, both Sarmatians (1st-2nd centuries CE). The final chronological phase in the IVth mantle of the tumulus is represented by grave no.7/M.7, which belongs to medieval migrants.

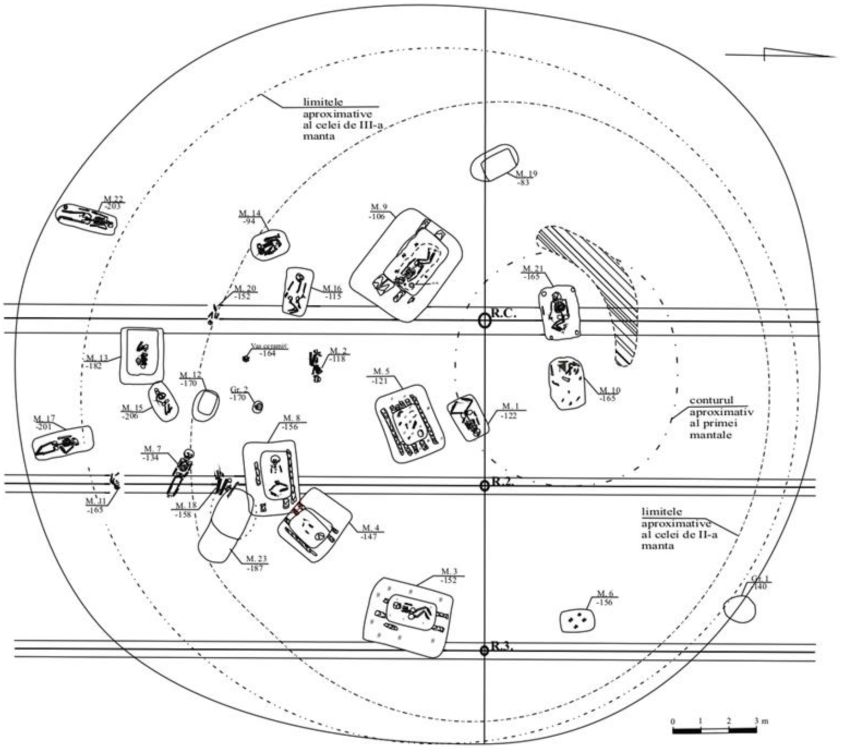

**Fig. 7.13. The general plan of the tumulus 12 from Crihana Veche**

##### *7.4.3.1 Grave 12:5 (individual ID I10424)*

The grave was situated in the central-eastern sector of the mound, 3.30 m south-east of the central landmark, at a depth of 1.21 m. The burial chamber was provided with a step oriented north–east/south–west, positioned at a depth of 0.50 m from the level of the burial pit, with dimensions of 1.95 × 2.50 m. Six oak beams, varying in thickness from 0.10 to 0.15 m and reaching a maximum length of 2.2 m, longitudinally covered the lower level of the step. Beneath the beams, the burial chamber's step was covered with a decayed grey-white mat. The chamber itself had a rectangular shape with rounded corners, oriented to the northeast-southwest. The dimensions of the grave pit were 1.30 × 0.90 m, and the depth of the burial pit from the level of the step was 0.78 m. The grave had been disturbed in ancient times.

Fragments of wood from the beams covering the mortuary chamber were found mixed with human bones arranged in various regions and depths in the filling of the pit. Fragments of the skull were discovered in the eastern corner of the burial chamber. Brown traces from the plant layer covering the pit bottom and scattered human bones were observed at the bottom of the pit. The skeleton, lacking anatomical connection, is nearly complete but poorly preserved and extremely fragmentary. All skeletal elements are intensely pigmented with red ochre. Anthropological sex: probably male. Molecular sex: female, mt-haplogroup U5a1a1+16362. Biological age at death: about 7-8 years (child). Pathologies: supragingival calculus; active porotic hyperostosis (*cribra cranii*). The burial is attributed to Yamnaya culture.

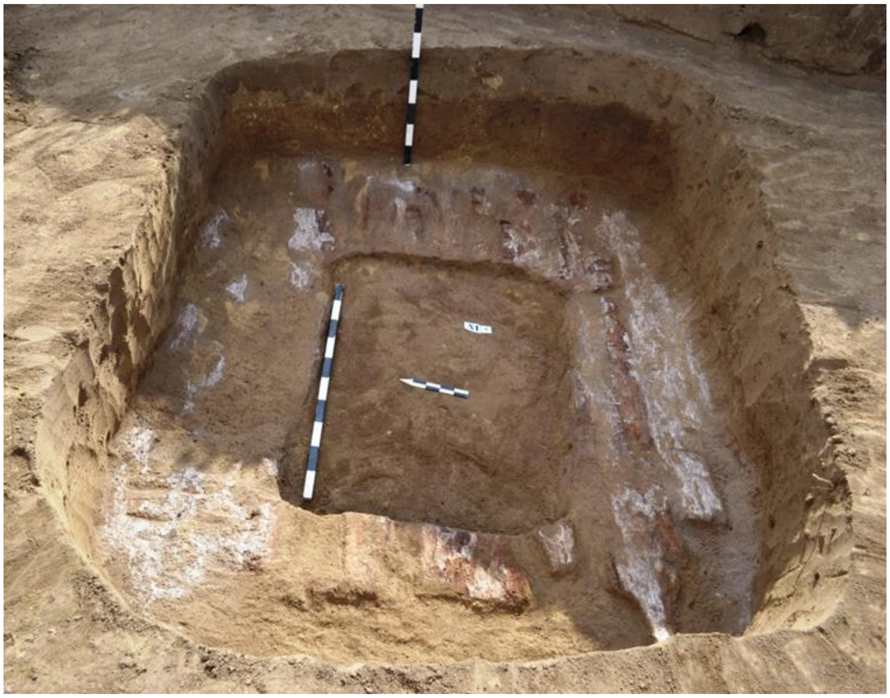

**Fig. 7.14. Crihana Veche, tumulus 12, grave 5**

##### *7.4.3.2 Grave 12:10 (individual ID I10206)*

The grave was situated in the northeastern sector of the mound, 2.80 m away from the central landmark (R.C.), at a depth of 1.65 m. The burial chamber, rectangular with rounded corners and slightly arched sides, was oriented east-west, measuring 1.25 × 1.80 m. Based on the characteristics of the filling of the pit, the arrangement of skeletal remains, and the location of the grave goods, it can be inferred that grave no.10/M.10 was disturbed in ancient times. The skeletal remains were found scattered in different areas of the pit and at different depths.

It was deduced that the deceased was originally placed at the bottom of the pit, with the head oriented to the east. Traces from the plant layer covering the pit bottom were found near and beneath the skeleton. Grave goods: a flint arrowhead of elongated triangular shape with slightly convex sides and a concave base. The artefacts' length is 2 cm, width 1.4 cm, and thickness 0.5 cm.

The skeleton, lacking anatomical connection, is incomplete and poorly preserved. Traces of bright red ochre were identified on all its elements, particularly intense on the skull. Other taphonomic changes observed include cracking, exfoliation (aerial weathering), and marks left by rodent teeth. Anthropological sex: male. Molecular sex: male, with an unusual Y-haplogroup J2b2a1 (J-L283). Biological age at death: about 30 years (young adult). Pathologies: supragingival calculus; active porotic hyperostosis (*cribra cranii*). The skeleton is very robust, with extremely pronounced muscle insertions, with extensive enthesopathic changes on the humerus and femurs. The individual exhibits a very large skeletal stature, at least 181 cm. Traumas: a perimortem fracture in the middle third of the right clavicle. The burial must be attributed to Cernavodă-I or Yamnaya cultures.

*7.4.3.3 Grave 12:19 (individual ID I10208)*

The grave was situated in the central-western sector of the mound, approximately 5.10 m west of the central landmark, at a depth of 0.83 m. The burial chamber was oval-shaped, oriented southeast-northwest, and displayed a step on the northwest, southwest and southeast sides. The step was positioned at a depth of 0.35 m from the level of the burial pit, with a 5 cm deepening towards the edge of the burial pit. The burial chamber was rectangular in shape with rounded corners, measuring 1.05 × 0.65 m. At the bottom level, the pit was displaced about 0.1 m under the northeast wall, thus forming a side niche.

Several small bone fragments intensely pigmented with red ochre were discovered in the filling of the pit. The skeletal remains, extremely fragile (fragments of the parietals, temporals and diaphyses of the long bones of the limbs), were attributed to a sub-adult. Age at death: about 12-14 months (infant). Anthropological sex: indeterminate. Molecular sex: male, Y-haplogroup R1b (R-L51). The burial is attributed to Yamnaya culture.

#### 7.5. Ciumai

*Summary by A. Simalcsik & I. Ciobanu*

##### *7.5.1 Ciumai, tumulus 1*

Tumulus 1 from Ciumai, investigated in 2015 by specialists from the National Archaeological Agency of the Republic of Moldova, is part of a larger burial mound necropolis in the Lower Prut Plain. It is situated on the first terrace, on the left side of the Salcia stream, which flows into the Ialpug river. The mound was partially affected by the works carried out for the development of a channel between the Ialpug and Taraclia lakes, and later by the reconstruction of the national road.

At the beginning of the research, the mound had a maximum height of 0.4 m and a diameter of about 38 m. At least 40 features were discovered, as follows: 14 graves dating from the Late Eneolithic to the Roman era; 21 utility pits (Gr.); four Late Medieval fire installations; a ditch from the Modern era.

The mound consisted of three mantles erected in different historical periods. Mantle I was raised above grave no.7/M.7 (Late Eneolithic, post-Mariupol). Mantle II is attributed to grave no.6/M.6 (Yamnaya). Also in this mantle, grave no.2/M.2 and probably grave no.4/M.4 were arranged. Mantle III was raised above grave no.5/M.5, which can be dated to the middle Yamnaya. Following the construction of the third mantle, burials no.1/M.1, 3/M.3, 8/M.8, 13/M.13, and 14M/14 were arranged within it.

Among the artefacts discovered in the funerary features from the Early Bronze Age, the askos from grave no.11/M.11 stands out, a type of vessel found extremely rarely in Yamnaya features, but which finds more numerous analogues in the ceramics of sedentary populations of the Eneolithic and the Early Bronze Age from the south-Danube regions.

The Classical chronological horizon is represented by grave 12/M.12 (Sarmatian, 1st-2nd centuries AD), situated in the southwestern sector of the mound. The last chronological phase is represented by the 21 pits (Gr.) and four open-air hearths, attributed to Nogai Tatars from the 18th century AD, as well as a ditch from the late Middle Ages or the Modern era.

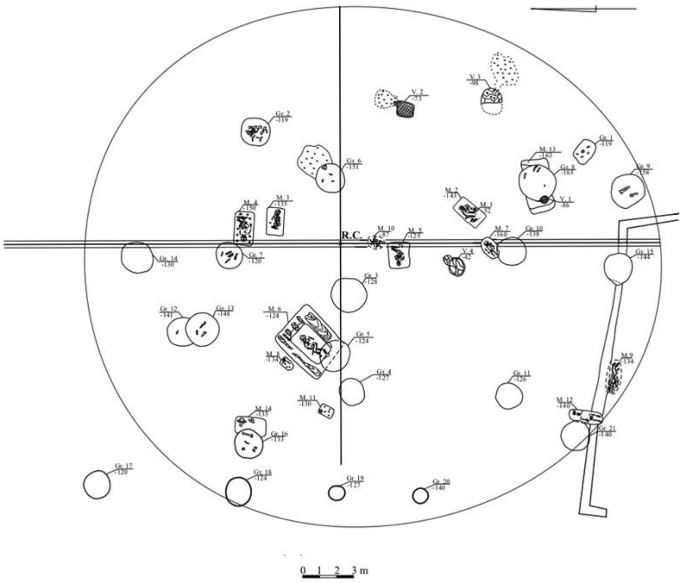

**Fig. 7.15. The general plan of the tumulus 1 from Ciumai**

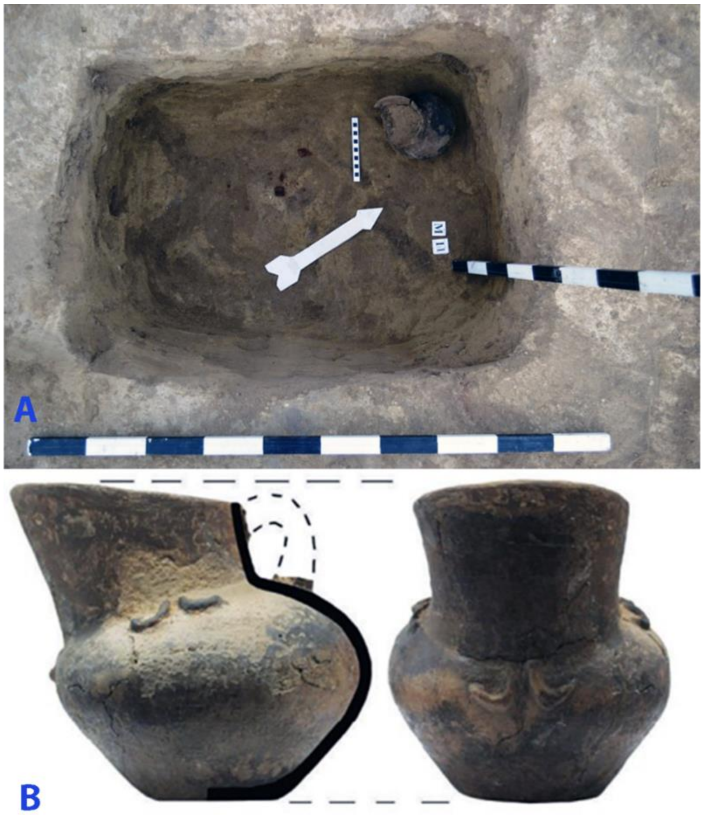

**Fig. 7.16. Ciumai, tumulus 1, grave 11. A – *askos in situ*; B – *askos* after restoration.**

*7.5.1.1 Grave k1:2 (individual ID I7847): 3017-2906 calBCE*

The grave was situated in the southeastern sector of the mound, with a northeast-southwest orientation and a depth of 1.43 m. The pit exhibited an approximately rectangular shape, measuring 1.0 × 1.4 m, with a depth of about 1 m from the identification level. The deceased was laid supine, with the arms extended along the body and the lower limbs initially raised from the knees, later falling to the right side. Bright red ochre covered the deceased, with a higher concentration attested on the skull, the bones of the pelvis, and those of the foot.

An oval-irregular shaped river stone, measuring 3 × 4 × 8 cm was found in the region of the right arm. Brown traces from the plant layer covering the pit bottom were found beneath the skeleton. The skeleton, in anatomical connection, is well represented but precariously preserved.

The skeleton is extremely robust, with developed muscle insertions, and a large skeletal stature. Anthropological sex: male. Molecular sex: male R1b (R-M12149); mt-haplogroup U5a1b1. The burial is dated to 3017-2906 calBCE (4350±20 BP, PSUAMS-4406).

Biological age at death: about 45-50 years (middle-aged adult). Pathologies: porotic hyperostosis (*cribra cranii*); degenerative osteoarthritis; muscle stress markers. Traumas: a perimortem trauma on the left parietal bone caused death; the cracks caused by the blow extend to the frontal and the right parietal. The burial is attributed to Yamnaya culture.

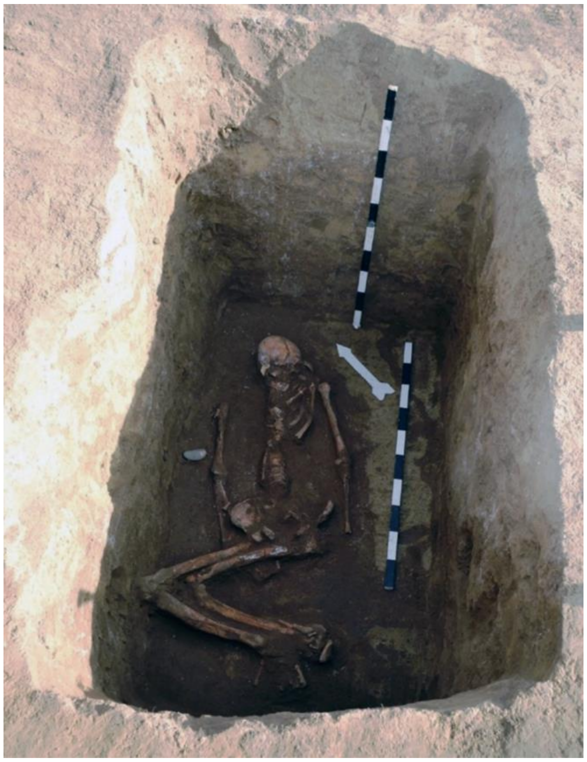

**Fig. 7.17. Ciumai, tumulus 1, grave 2.**

*7.5.1.2 Grave 3 (individual ID I7848): 2905-2706 calBCE*

The grave was situated in the north-eastern sector of the mound, facing west-east. The burial chamber had a rectangular shape, measuring 1.0 × 1.5 m, with a depth of 0.2 m from the identification level. The deceased was laid supine, with raised knees, head oriented to the west, arms bent at the elbows, palms placed on the pelvis, and legs strongly contracted to the right side. The bones of the skeleton were heavily reddened from the red ochre used to pigment the deceased, with a larger amount of ochre present on the skull. Brownish-black traces from the plant layer covering the pit bottom were found.

The skeleton, in anatomical connection, was almost complete and well preserved. It exhibited robust bone structure with very well-marked muscle insertions. The skeletal stature is large, estimated to be at least 170 cm. Anthropological sex: male. Molecular sex: female, mt-haplogroup U4c2a.

Biological age at death: 50-60 years (old adult). Pathologies: antemortem tooth loss; supragingival calculus; degenerative osteoarthritis; muscle stress markers; advanced osteoporosis. The burial is attributed to Yamnaya culture and dated to 2905-2706 calBCE (4235±20 BP, PSUAMS-4241).

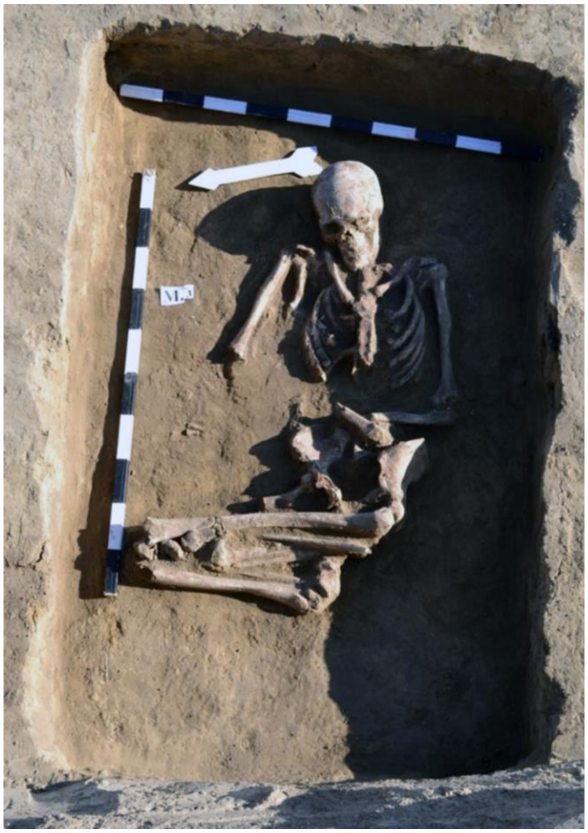

**Fig. 7.18. Ciumai, tumulus 1, grave 3.**

##### *7.5.1.3 Grave 4 (individual ID I10398)*

The grave was situated in the northeastern sector of the mound, 5.2 m from the central landmark, at a depth of 1.7 m. The burial chamber was rectangular, measuring 1.3 × 1.9 m, with a west-east orientation.

The deceased was laid supine, with the head oriented to the west, with raised knees, subsequently fallen to the left side and upper limbs extended along the body. The skeleton exhibited intense pigmentation with dark red ochre, with a larger amount found on the femurs. An oval-irregular-shaped sandstone measuring 4.5 × 7.0 × 16.0 cm was found in the lower abdomen area.

The skeleton, in anatomical connection, was poorly represented and preserved. The bones are gracile. Anthropological sex: female. Molecular sex: female, mt-haplogroup I4a.. Biological age at death: over 60 years (old adult). Pathologies: degenerative osteoarthritis; musculoskeletal stress markers; bone remodelling in the distal third of the diaphyses of the femurs, on the internal side, which could have occupational causes. The burial is attributed to Yamnaya culture.

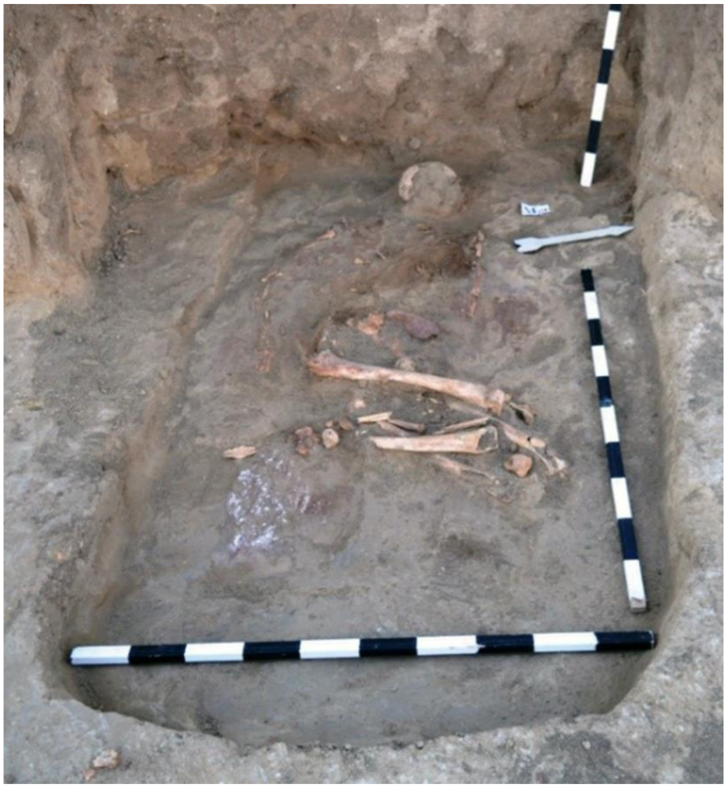

**Fig. 7.19. Ciumai, tumulus 1, grave 4.**

#### 7.6. Cimișlia

*Summary by A. Simalcsik & S. Popovici*

##### *7.6.1 Cimișlia, tumulus 8*

Tumulus 8 from Cimișlia was investigated in 2015 by specialists from the National Archaeological Agency of the Republic of Moldova. The mound is part of a necropolis consisting of approximately six tumuli located on the border between the Central Moldavian Plateau and the Southern Moldavian Plain, on the first terrace of the Cogâlnic River.

The mound was affected by agricultural activities. At the beginning of the research, it measured 36 m in diameter and 0.8 m in height. Ten burials, two agglomerations of bones and ceramics, a cult complex, a circular ditch, and a circular stone platform were identified. The first mantle, stone platform, ditch, and cult complex were constructed together with grave 3, dated to the Late Eneolithic (first half of the 4th millennium BC), attributed to Cernavodă-I communities. Grave 4 also belonged to this layer.

The next cultural-chronological horizon, dating back to the end of the 4th millennium BC, is represented by two Zhivotilovka-type graves from the Early Bronze Age, grave 2 and 8. These are characterized by a crouched position on the right or left sides and the presence of import grave goods from the late Trypillia and Maikop-Novosvobodnaya cultural environments.

Later, grave 6 attributed to the Yamnaya communities was arranged in the first mantle of the mound, which led to the raising of an additional mantle over the entire surface of the mound. Another Yamnaya feature, grave 1, was excavated in the second mantle. Both features can be dated to the middle stage of the evolution of the Yamnaya culture in the northwest Black Sea region. In burial mound 8 from Cimișlia, the graves of the Yamnaya culture were followed by graves belonging to the Catacomb culture communities (grave 9). The subsequent horizon includes grave 7 and grave 10, both of which are attributed to the Sarmatians. The latest burial of the tumulus is grave 5 attributed to late Turanian migrants.

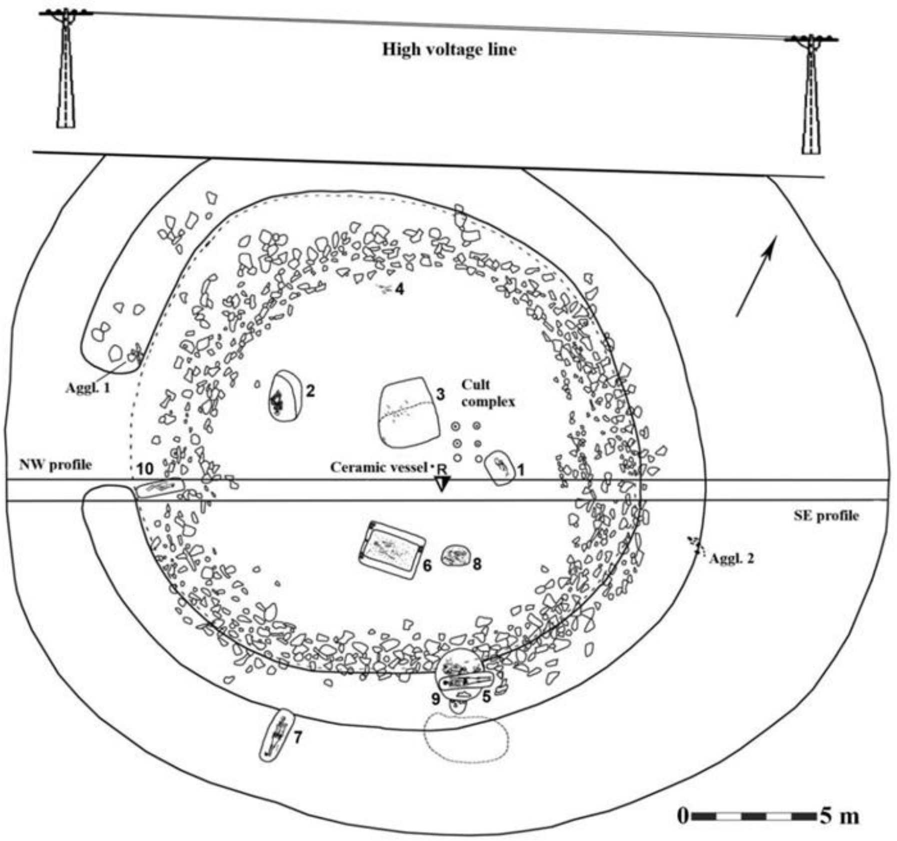

**Fig. 7.20. The general plan of the tumulus 8 from Cimișlia.**

*7.6.1.1 Grave 4 (individual ID I7857): 3903-3647 calBCE*

The grave was situated in the western sector of the mound, at a depth of 0.46 m and a distance of 8.5 m northwest of the central landmark. The graves had been destroyed by anthropogenic factors, particularly agricultural activities. The skeleton is extremely poorly represented and preserved. The bone remains are gracile, with weak muscle insertions. Anthropological sex: female. Molecular sex: female. Biological age at death: 20-35 years (young adult). The burial must be attributed to Eneolithic period or Cernavodă-I culture. Mt-haplogroup U5b1. The bi8urial is dated to 3903-3647 calBCE (4965±35 BP, Poz-121047).

*Bibliography to section 7:*

*Ailincăi, S.C., Constantinescu, M., Dima, A., Sava, G., Mihail, F., Micu, C., Mocanu, M., Stănică, A.D. 2021, Bronze Age graves at Jijila (Southeastern Romania), in Heyd, V., Kulcsar, G., Preda-Bălănică, B. (eds.), Yamnaya Interactions. Proceedings of the International Workshop held in Helsinki, 25-26 Aprilie 2019, Archaeolingua 44, The Yamnaya Impact on Prehistoric Europe 2, Budapest, 207-242.*

*Ailincăi, S.C., Mihail F., Carozza L., Constatinescu M., Soficaru A., Micu C. 2014, Une découverte funéraire de début de l'Age de Bronze en Dobroudja (Sud-est de Roumanie). Le tumulus de Rahman (Com. Casimcea, Dep. Tulcea), Prilozi Instituta za arheologiju u Zagrebu 31,135-149.*

*Ailincăi, S.C., Mihail, F., Constantinescu, M., Carozza, L., Micu, C., Burens, A. 2016, Découverte d'un tumulus de l'age du Bronze a Rahman sur la commune de Casimcea (Dép. de Tulcea), Studii şi Cercetări de Istorie Veche şi Arheologie 67, 1-2, 29-52.*

*Ciobanu I., Agulnicov S. (2016). Investigațiile arheologice de la Crihana Veche. Rezultatele campaniei 2015. Arheologia Preventivă în Republica Moldova III, 2016, pp. 45-60.*

*Ciobanu I., Simalcsik A., Bejenaru L., Pîrnău R., Agulnicov S., Popovici S. (2019). Considerații preliminare privind cercetările arheologice și interdisciplinare de la Crihana Veche (raionul Cahul). Campaniile 2016 și 2017. Arheologia Preventivă în Republica Moldova IV, 2019, pp. 71-118.*

*Ciobanu I., Vornic V., Agulnicov S., Noroc I. (2016). Tumulul 1 de la Ciumai (com. Vinogradovca, r-nul Taraclia). Date preliminare. Arheologia Preventivă în Republica Moldova III, 2016, pp. 29-44.*

*Frînculeasa A. (2021) Burials mounds in the Lower Danube region – from the international to the local and the other way round, În: V. Heyd, G. Kulcsár and B. Preda-Bălănică (eds.), Yamnaya Interactions. Proceedings of the International Workshop held in Helsinki, 25-26th April 2019, pp. 173-205.*

*Frînculeasa A. (2023) Movila Mare of Smeeni, representative research of Romanian archaeology history (new information on the Middle Bronze Age), În: V. Diaconu, A. Gafincu (eds.) Bronze Age in Eastern Europe. Interdisciplinary researches, Brăila-Piatra Neamț, 2023, pp. 87-106.*

*Frînculeasa A., Simalcsik A., Petruneac M., Focșăneanu M., Sîrbu R., Frînculeasa M.N. (2023) From the Eurasian steppe to the Lower Danube: the tradition of intentional cranial deformation during the Bronze Age, Archaeological and Anthropological Sciences, 15 (125), https://doi.org/10.1007/s12520-023-01826-0.*

*Frînculeasa A., Simalcsik A., Preda B., Garvăn D. (2017) Smeeni-Movila Mare: monografia unui sit regăsit, Târgoviște.*

*Frînculeasa Alin, The Children of the Steppe: descendance as a key to Yamnaya success, Studii de Preistorie, 16, 2019, pp. 129-168.*

*Popescu, D. 1941, La tombe a ocre de Casimcea (Dobrogea), Dacia 7-8, 1937-1940, 85-91.*

*Popovici S., Ciobanu I. (2021). Cercetări arheologice la Cimișlia (Bons Offices, Chișinău). ISBN 978-9975-87-915-6.*

*Popovici S., Ciobanu I., Agulnicov S., Noroc I. (2016). Tumulul 8 de la Cimișlia. Considerații preliminare. Arheologia Preventivă în Republica Moldova III, 2016, pp. 19-28.*

### 8. CARPATHIAN BASIN

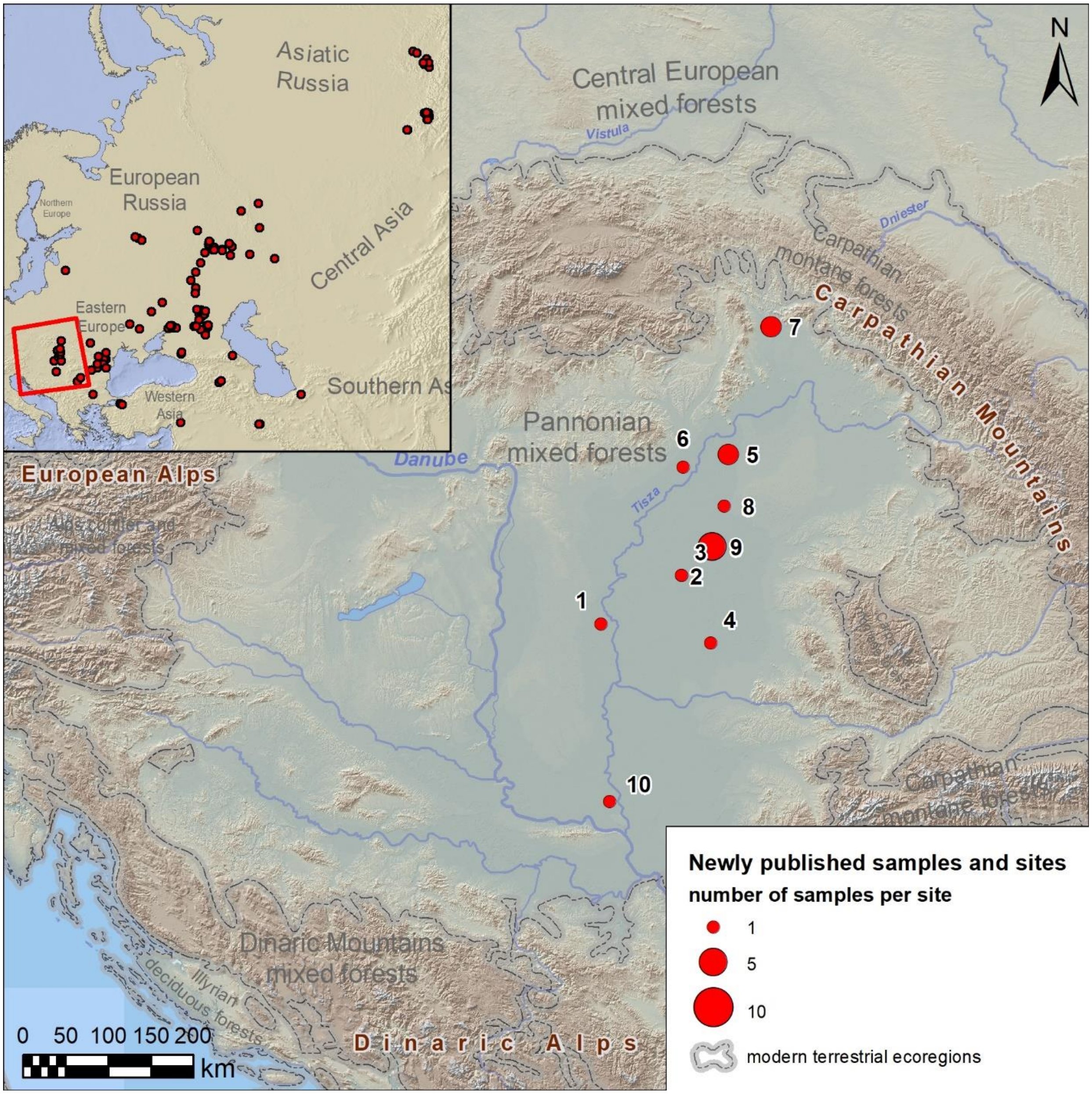

**Fig. 8.1. Newly published samples from the Carpathian Basin.** 1 - Csongrád-Kettőshalom-Bárdostanya (Hungary, Csongrád-Csanád County, Csongrád); 2 - Hungary, Dévaványa-Barcehalom (Békés County, Gyomaendrőd); 3 - Dévaványa-Csordajaras (Hungary, Békés County, Gyomaendrőd); 4 - Kétegyháza-Törökhalom-Kishalom (Hungary, Békés County, Gyula); 5 - Kunhegyes-Nagyállás-halom (Hungary, Hajdú-Bihar County, Hajdúnánás); 6 - Mezőcsát-Hörcsögös (Hungary, Borsod-Abaúj-Zemplén County, Mezőcsát); 7 - Lesné (Slovakia, Košice Region, Michalovce District); 8 - Nagyhegyes-Elep-Mikelapos (Hungary, Hajdú-Bihar County, Hajdúszoboszló); 9 - Sárrétudvari-Őrhalom (Hungary, Hajdú-Bihar County, Püspökladány); 10 - Zabalj-Medisova-humka (Serbia, South Bačka District, Žabalj Municipality, Zabalj).

#### 8.1 Lesné

*Summary by Alena Šefčáková*

Lesné belongs to the sites of the Eneolithic Slovakian Mound culture (East Slovak tumuli) belonging to the northern Carpathian prehistoric cultures (Budinský-Krička, 1967). Seven intact tumuli have been discovered in its vicinity and skeletal findings come from three of them.

The culture of shepherd-peasant builders of eastern Slovak tumuli is characterized by burial mounds on mountain ridges with an altitude of sometimes more than 500 m, which form burial grounds even several kilometres long. Their builders apparently came from Transcarpathia and with their arrival appear in the Carpathian basin first copper ornaments in the shape of a willow leaf. Research of the tumuli began in 1936 and their number is currently estimated at around 2,000.

Burials in mounds were bi-ritual. Unburned individuals usually lay in crouched positions or on their backs with outstretched legs in the shape of a rhomboid (Šefčáková, 1990). Scattered fragments of human bones in some mounds may indicate human sacrifice. Grave goods are rare. These are mostly containers, more often their fragments, or stone tools, exceptionally animal bones. The objects were not located near the skeletons themselves, but outside the grave on the original terrain and in the embankment. The period of the epi-Corded-Ware culture of the group of East Slovak tumuli belongs to the late Eneolithic. It begins in the classical (younger) phase of Corded Ware culture, according to radiocarbon data from Hankovce, perhaps between 2700–2600 BC (Machnik et al., 2008, 236) and disappears at the beginning of the Bronze Age. However, the radiocarbon dates of the human bones from Lesné demonstrate the earlier age of the buried individuals and place the lower chronological boundary of the East Slovakian Mounds to the first centuries of the 3rd millennium BCE.

##### *8.1.1 Lesné, kurgan (tumulus) 1*

###### *8.1.1.1 Undisturbed grave (individual ID I7890)*

Male (obtained osteologically and genetically), ca. 40-55 years (maturus), he was lying on his back with outstretched legs in the shape of a rhomboid. The skeleton was oriented to the W/E. Preserved parts: a cranium without a facial part, only the alveolar part of the maxilla, diaphysis of the right humerus, both femurs and tibias, the remains are of medium build with medium MR, the skull is short and broad, high with a broad forehead. Preserved teeth are without caries, they have partially abraded enamel with exposed dentin (2nd degree), damaged preserved diaphysis of the right humerus has a massive MR, and the tibiae are platycnemic (flattened on the sides).

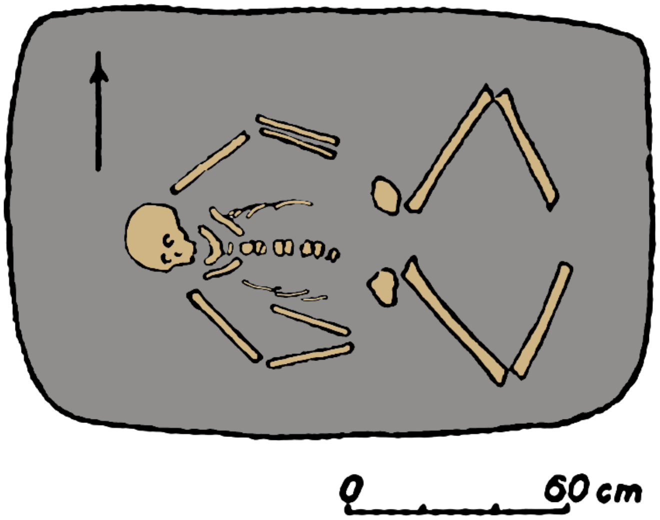

**Fig. 8.2. Lesné. Kurgan 1, undisturbed grave.**

##### *8.1.2 Lesné, kurgan (tumulus) 2*

*8.1.2.1 Grave 2:1 (individual ID I7891).*

Male (obtained osteologically and genetically), ca. 30-40 years (adultus II); 161-165 cm tall, lying on his left side. The skeleton was oriented to the WSW/ENE. Preserved parts: Fragments of the skull and vertebrae, the damaged diaphysis of the right humerus, damaged diaphyses of both femora. The remains are of a more robust structure with a medium MR. There are traces of fire on the bones, abrasion of the teeth corresponds to the 2nd – 3rd degree, damaged preserved diaphysis of the right humerus has a significant exostosis on the proximal part (probably due to some stronger mechanical intravital action), femurs are stenomeric (flattened at the sides).

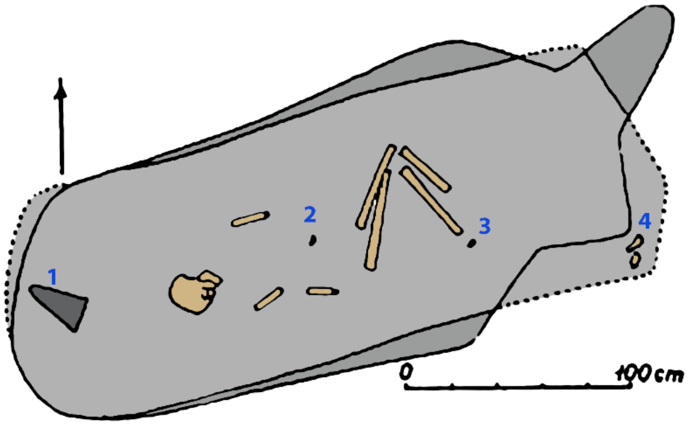

**Fig. 8.3. Lesné. Kurgan 2, burial 2. 1 – stone piece, 2, 3 – flakes, 4 – human and animal bones.**

##### *8.1.3 Lesné, kurgan (tumulus) 3*

*8.1.3.1 Grave 3:3 (individual ID I7889)*

Male (according to DNA), the originally gender indefinite, ca. 30 years (younger adult). Preserved parts: Four fragments from the anterior part of the robust mandible, 15 damaged teeth from the maxilla and mandible, the other remains of the skeleton were not preserved. On the remains, there are traces of fire, tooth enamel erupted and deformed by the glow of the fire. The skeleton was probably oriented to the NNW/SSE.

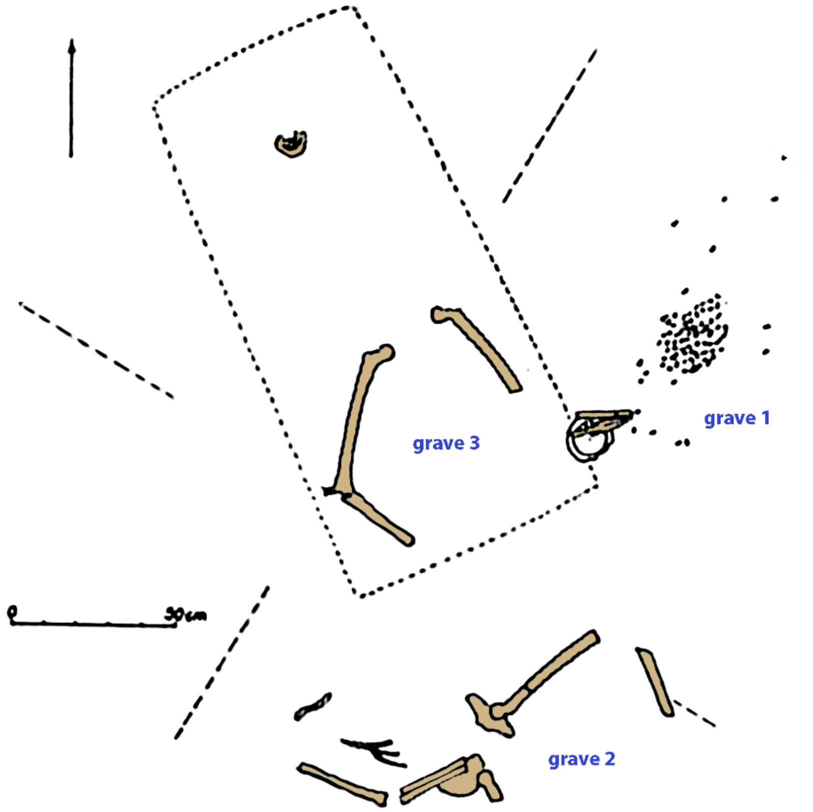

**Fig. 8.4. Lesné. Central part of kurgan 3 with graves 1-3.**

*<u>Bibliography to section 8:</u>*

*Budinský-Krička, V., 1967: Východoslovenské mohyly. Slov. Archeol., 15(2): 277–388.*

*Machnik, J., Mačalová, H., Jarosz, P., Włodarczak, P., 2008: Kurhan Nr 2 kultury ceramiky sznurowej w miejscowości Hankovce okr. Bardejov, stanowisko 1. In: Machnik, J. (ed.): Archaeology and natural background of the Lower Beskid Mountains, Carpathians, Part II, Kurimská brázda. Kraków 2008, 193-240.*

*Šefčáková, A., 1990: Antropologický materiál z východoslovenských eneolitických mohýl v Šapinci a Lesnom. Zbor. Slov. Nár. Múz., Prír. Vedy, 36: 149-163.*

#### Supplementary Information 2

### The proximate and distal origins of the Yamnaya and the origins and dispersals of Indo-Anatolian and Indo-European languages

Iosif Lazaridis, Nick Patterson, and David Reich

#### Introduction

The discovery that the Yamnaya pastoralists of the early Bronze Age (~5kya) had mixed origins[1] related to Eastern European hunter-gatherers and populations of West Asia, led to a search for the populations which contributed to their ancestry and for the geographical location in which the Yamnaya-Afanasievo[2] cluster (which we call here "core Yamnaya") emerged:

- Which populations of the "south" (West Asia and the Caucasus) contributed to steppe populations? The genetic structure of ancient West Asia[3,4] and the Caucasus included broadly speaking "Inland"/"Highland" populations such as Caucasus hunter-gatherers (CHG) and early farmers from the Zagros in Iran[5] as well as "East Mediterranean" or Anatolian-Levantine populations.[5-7] The latter may have contributed to steppe populations either indirectly via the European farmer descendants of the Anatolian farmers (from southeastern Europe, eastward into the steppe)[8] or via the Caucasus and Armenian plateau where Anatolian-Levantine ancestry was also ubiquitous in Neolithic and later times[4,5]
- Which populations of the "north" (the Pontic-Caspian steppe or further north) contributed to steppe populations? Broadly speaking, the steppe was inhabited by populations forming a longitudinal transition zone between Western and Eastern hunter-gatherers towards "Ancient North Eurasians" [5,9] (a WHG-EHG-ANE cline), a part of a longer 7,000-km long cline of boreal hunter-gatherers from Europe to East Asia,[10] with key intermediate populations (between WHG and EHG) in

Ukraine[11] and Siberian-influenced populations such as those of the Progress-2 Eneolithic and Steppe Maikop further east[8].

- When did this admixture take place? Inferences from admixture LD suggest a 5th millennium admixture in the ancestry of the Yamnaya[4,12], but "intermediate" populations between the "south" and "north" already existed in the steppe before the inferred admixture dates[6,8,11]. If admixture in the origin of the Yamnaya was complex, involving two or more episodes, then the admixture date estimates may not correspond to either of these episodes. What these estimates do show is that there was recent admixture in the Yamnaya—within an Eneolithic timeframe—and so their final formation inferred with other methods like $f_3$-statistics[9] cannot be pushed back to the remote past.
- Where did this admixture take place? With the caveat of the possibility of maritime movements, "south" ancestry may have entered the European steppe either from the European farmers to the west of the Dnipro or the West Asian/Caucasus populations via the North Caucasus. However, groups of hunter-gatherers were widely dispersed along the Pontic-Caspian steppe in the Dnipro/Don/Volga regions and in the forest-steppe zone to the north of it, leaving many possibilities on the identity of the population(s) bearing "north" ancestry. Moreover, there were clearly populations of substantial "EHG" ancestry on the North Caucasus piedmont itself[8] and the boundary—or more accurately, transition zone—between "south" and "north" populations remains unknown and may have not been on the steppe itself. This transition may have occurred somewhere along the geographical gap between the Caucasus hunter-gatherers of Georgia (south of the Caucasus) and the hunter-gatherers of the Pontic-Caspian steppe.

In a previous study we identified that Eneolithic steppe populations had CHG ancestry, but Yamnaya also received an Anatolian-Levantine input (from unidentified proximate sources from the North Caucasus through Armenia to Eastern Anatolia and North Mesopotamia).[4] We can use newly available data on hunter-gatherers of Eastern Europe[13] and hunter-gatherers and pastoralists of Eastern Europe (this study) to better understand what the steppe was like genetically before the emergence of the Yamnaya, and how the Yamnaya themselves appeared.

#### Modeling framework and an exploratory Principal Components Analysis

In this section we introduce the populations to be used and show an exploratory PCA analysis of them. We use the following set of Right populations in a qpWave/qpAdm setup[1,14] whose results we will describe in the next section.

**Right:** OldAfrica, Russia_AfontovaGora3, CHG, Iran_GanjDareh_N, Italy_Villabruna, Russia_Sidelkino.SG, Turkey_N

- OldAfrica (N=9) is a set of ancient African individuals (~1-8kya) from Cameroon, Kenya, and Tanzania, represented by in-solution enrichment data.[15-17]
- Russia_AfontovaGora 3[18] (~18kya) "Ancient North Eurasian" from Siberia represented by damage restricted in-solution enrichment data.
- CHG (*N*=2) are Caucasus hunter-gatherers from Kotias and Satsurblia caves (~10-13kya) in Georgia, represented by shotgun sequencing data.[19]
- Italy_Villabruna (*N*=1) (~14kya) "Western hunter-gatherer" is the earliest high-quality individual of the "Villabruna cluster", represented by in-solution enrichment data [18]
- Iran_GanjDareh_N (*N*=8) are early Neolithic farmers from the Central Zagros of Iran[5] (~10kya), represented by in-solution enrichment data.

- Russia_Sidelkino.SG (*N*=1) the oldest (~11kya) individual of "Eastern hunter-gatherer" ancestry, represented by shotgun sequencing data[20]
- Turkey_N (*N*=27) are Anatolian Neolithic farmers from the Marmara (Barcın, Menteşe, and Ilıpınar) region of NW Anatolia, represented by in-solution enrichment data.[6]

Collectively, this set of outgroups includes the main ancestries of boreal Eurasia from mainland Europe through eastern Europe to Siberia (with a representative population of each of the main stops along the WHG-EHG-ANE cline) and of West Asia, including both "Inland" and "East Mediterranean" cluster representatives.[3]

We are interested in modeling the ancestry of the following populations of the Eneolithic/Chalcolithic and Early Bronze Age, all represented by in-solution enrichment data:

**Test:** Armenia_C[5], Azerbaijan_C[21], BPgroup (Berezhnovka, this study and Progress-2 PG2004 [8]), CoreYamnaya (see below how we identified this set and for references), GK1 (this study), GK2 (this study), Igren_o (this study), Kazakhstan_Kumsay_EBA (this study and [22]), Krivyansky (this study), Labazy (this study), Maikop[8], Maximovka (this study), Murzikha (this study), Lebyzahinka_HG[13], PVgroup[8] (Progress-2 PG2001 and Vonjucka-VJ1001), Remontnoye (this study), Russia_Don_EBA_Yamnaya (this study), Russia_Steppe_Maikop[8], TTK[23], the Serednii Stih subsets: SSlo, SSmed, SShi (this study and[11]), the Volga Cline subsets: Klo, Kmed, Khi, KhlopkovBugor, Syezzheye, Ekaterinovka (this study and [6]), and Unakozovskaya (this study and [8]).

We use as potential sources the following populations that include some of the above (as proximate sources), but also earlier Neolithic and hunter-gatherer populations:

**Sources:** Armenia_Aknashen_N[4], Armenia_C[5], Armenia_MasisBlur_N[4], Azerbaijan_N[21,24], Azerbaijan_C[21], BPgroup, CHG[19], GK1 (Golubaya Krinitsa, this study), GK2 (Golubaya Krinitsa, this study), Igren_o (this study), Iran_GanjDareh_N[5], Krivyansky, Labazy (this study), Lebyazhinka_HG, Maikop, Murzikha (this study), PVgroup, Remontnoye, Unakozovskaya (this study and [8]), Russia_Karelia[13], Russia_Steppe_Maikop, Trypillia (this study and [11]), TTK[23], Ukraine_N (this study and [11]), UpperVolga (this study), WSHG[22], and the Serednii Stih and Volga Cline subsets also included in the Test list

Our set of sources is diverse in ancestry, geography and time. We tried to include sources at varying time depths so that we can study the origins of the Test populations in terms of both more "proximal" or more "distal" sources.

We carried out principal components analysis in *smartpca* [25] using the following populations all represented by in-solution enrichment data to form the axes and with parameters newshrink: YES and lsqproject: YES:

OberkasselCluster (set of trans-Alpine WHG individuals identified in[13]), Russia_Firsovo_N, Iran_HajjiFiruz_C[22], Iran_C_SehGabi[5], Iran_C_TepeHissar[26], Israel_C[27], Germany_EN_LBK[1,6,28,29]

This analysis is useful to create a non-model-based depiction of the populations included in the Base, Sources, and Test lists. It includes populations surrounding the steppe but does not include the Test/Sources populations so as to avoid any systematic difference between populations used to compute the PCA and those projected on it. We make the following general observations based on the PCA:

- No pre-Yamnaya populations seem to match exactly the core Yamnaya

- The Yamnaya, broadly speaking, are situated between ancient West Asian and Eastern European populations.
- There are several populations, including BPgroup, PVgroup, Krivyansky, Remontnoye, SShi, and Khi which we call “peri-Yamnaya” that are genetically similar to the Yamnaya in the steppe context, while none of them are in the core Yamnaya cluster.
- The Yamnaya are on one end of a genetic cline that also includes populations of the Serednii Stih archaeological culture of the Dnipro-Don area. This strongly suggests that whatever the locality of Yamnaya formation was, it may have been either within the Serednii Stih culture or nearby in the east, given that the Yamnaya have less of the Ukraine_N-related ancestry that varies along the Serednii Stih cline.

The genetic intermediacy of the Yamnaya cluster, both in terms of distal sources (of Europe and West Asia), and the proximate “peri-Yamnaya” sources, raise several possibilities as to the cluster’s origin. For example, the Yamnaya are different with respect to BPgroup (\), Krivyansky (|), PVgroup (—), or Serednii Stih (/). If any of these peri-Yamnaya populations were ancestral to them, then corresponding “matched” sources must also be included to account for the position of the Yamnaya.

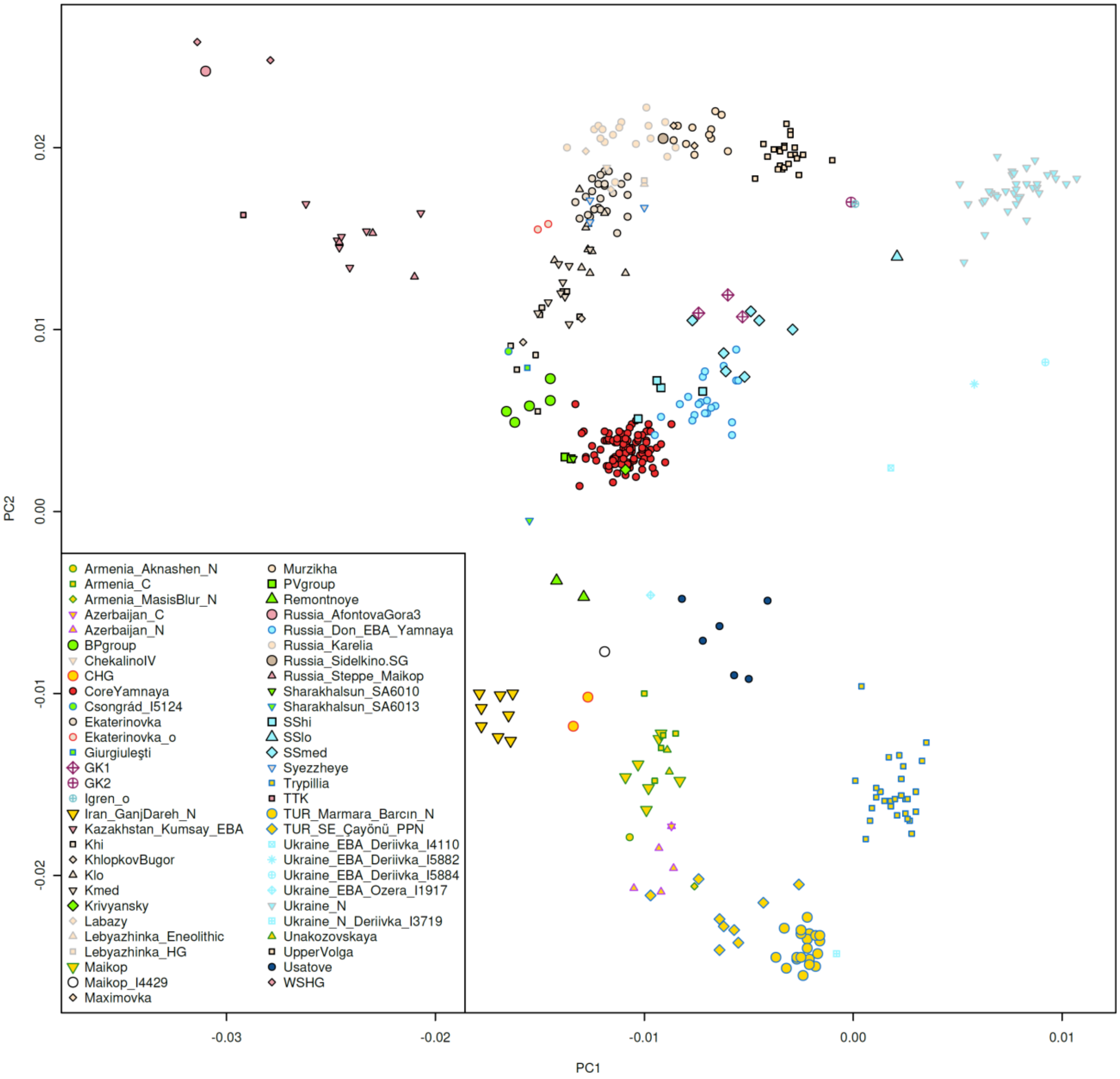

**Fig. S 1** A principal components analysis of steppe and neighboring populations.

### Results of qpAdm/qpWave analysis

#### Grouping core Yamnaya individuals

In Fig. S 1 we identified a set of individuals to represent the "Core Yamnaya" ancestry profile. We identified this set according to the following criteria: (i) labeled as either Yamnaya or Afanasievo based on archaeological considerations, (ii) at least 300,000 1240K autosomal SNPs covered, (iii) p-value of qpWave with Samara Yamnaya as the source of at least 0.2, (iv) date between 6000 and 4000 ky BP, and (v) no related individuals included.

The very stringent criteria may miss some Yamnaya-Afanasievo individuals, for example, with p-values <0.2, or fewer than 300,000 SNPs, or a population label that was not identified archaeologically as Yamnaya or Afanasievo. However, our aim here is to identify a set of high quality and reliably genetically similar individuals, and we were still able to recover 104 Core Yamnaya individuals which represent a high quality subset that is used in our qpAdm and qpWave analyses and represents numerous geographically dispersed populations, establishing the expansiveness of the Yamnaya population under strong criteria (Table S 1).

| Population | Sample size | Reference |
|---|---|---|
| Russia_Samara_EBA_Yamnaya | 24 | This study and ref.[1,6,22,30] |
| Russia_Afanasievo | 18 | This study and ref.[22] |
| Russia_CaspianInland_EBA_Yamnaya | 12 | This study |
| Russia_UpperOb_Eneolithic_Afanasievo | 6 | This study |
| Russia_Chelyabinsk_EBA_Yamnaya | 5 | This study |
| Russia_Remontnoye_EBA_Yamnaya | 5 | This study |
| Russia_Ural_EBA_Yamnaya | 5 | This study |
| Moldova_EBA_Yamnaya | 4 | This study |
| Russia_Volga_EBA_Yamnaya | 4 | This study |
| Ukraine_EBA_Yamnaya | 4 | This study and ref.[11,30] |
| Russia_Volgograd_EBA_Yamnaya | 3 | This study |
| Romania_EBA_Yamnaya | 2 | This study |
| Russia_Don_EBA_Yamnaya | 2 | This study |
| Russia_Kalmykia_EBA_Yamnaya.SG | 2 | ref.[2] |
| China_Xinjiang_G218_BA_Afanasievo_oWestEurasia | 1 | ref.[31] |
| Hungary_EBA_Yamnaya | 1 | This study |
| Kazakhstan_EBA_Yamnaya.SG | 1 | ref.[20] |
| Russia_Afanasievo_Yenisei | 1 | This study |
| Russia_Caucasus_EBA_Yamnaya | 1 | ref.[8] |
| Russia_Ishkinovka_EBA_Yamnaya | 1 | ref.[1,6] |
| Russia_Orlovka_EBA_Yamnaya | 1 | This study |
| Russia_UpperYenisey_Eneolithic_Afanasievo | 1 | This study |

**Table S 1 Subsets of Core Yamnaya individuals**

#### Which populations are simple clades of the Sources?

First, we show which populations can be modeled in qpWave as clades of other Source populations with a cutoff of p>0.05. (Table S 2)

| Test | Source | P-value |
|---|---|---|
| Azerbaijan_C | Azerbaijan_N | 0.478 |
| Ekaterinovka | Labazy | 0.607 |
| GK1 | SSlo | 0.078 |
| GK2 | Igren_o | 0.997 |
| Igren_o | UpperVolga | 0.197 |

| | | |
|---|---|---|
| Kazakhstan_Kumsay_EBA | Russia_Steppe_Maikop | 0.072 |
| Khi | KhlopkovBugor | 0.172 |
| Lebyazhinka_HG | Murzikha | 0.063 |
| Lebyazhinka_HG | Russia_Karelia | 0.212 |
| Maximovka | Lebyazhinka_HG | 0.561 |
| Syezzheye | Ekaterinovka | 0.128 |
| Syezzheye | Labazy | 0.146 |

**Table S 2 Populations that can be modeled as simple clades (*N*=1) with one of the sources**

The Chalcolithic population of Azerbaijan is cladal with its Neolithic one, but none of the other pairs of Chalcolithic and Neolithic populations from the South Caucasus (Armenia and Azerbaijan) are cladal to each other.

The Serednii Stih cline (that includes the Don Yamnaya in the middle and the Core Yamnaya including Afanasievo on one end) is visibly continuous (Fig. S 1) and so its division into subgroups is somewhat subjective, but useful given the clear diversity of ancestry along it. The two main groups, SShi and SSmed are clearly not cladal to each other (p=4.8e-09) and neither is the core Yamnaya with either of them (SShi=1.5e-07 and SSmed=9.7e-44).

The hunter-gatherer from Lebyazhinka in Samara is cladal to the set of hunter-gatherers from Karelia; these two widely dispersed locations were the basis for the original definition of an "Eastern hunter-gatherer" (EHG) population.[13]

On the Volga, we see that Ekaterinovka, Labazy, and Syezzheye are mutually all cladal to each other, paralleling the PCA in which all three populations are in the high-EHG end of the cline. The two Labazy individuals are somewhat separated with one (I6916) clustering with Karelia and another (I6910) with Ekaterinovka. However, we cannot reject that the two individuals form a clade (p=0.12). A clade is seen between Khi (a subset of Khvalynsk) and KhlopkovBugor; both of these populations are on the opposite, low-EHG end, of the Volga cline (Fig. S 1).

The core Yamnaya cannot be modeled with a single source (the highest p-value is for Krivyansky p=0.00009), and there is no good single source for the other peri-Yamnaya populations.

The two groups from Golubaya Krinitsa (GK1 and GK2) are not cladal to each other (p=2.5e-12), but GK2 is cladal to a single outlier individual of the Serednii Stih culture (p=0.997) (Igren_o, individual I27930) according to qpWave, and also clusters with it in PCA. This suggests a long-range connection across the span of the Don-Dnipro area between the Don (Golubaya Krinitsa) and Dnipro (Igren). The closest neighbors of these two individuals are the Upper Volga individuals: Igren_o forms a clade with Upper Volga (p=0.197), however, GK2 does not (p=9.7e-8), possibly reflecting the better data quality of GK2.

Another Serednii Stih outlier (SSlo / I1424 from the Middle Dnipro and the mid-4$^{th}$ millennium BCE) clusters visibly away from both Sshi/Ssmed and Igren_o in the direction of extra relatedness to hunter-gatherers of Ukraine, although it forms a weak clade with GK1 (p=0.078)

We draw the following conclusion: many (15/31) Test populations cannot be modeled with a single source, and those that do are mostly modeled in terms of other proximate sources. Some admixture is needed to understand the origins of the Test populations considered, so we now turn to models that involve admixture.

#### Which Test populations are consistent with being simple 2-way admixtures?

We next examine models with *N*=2 sources for each Test population. We define a feasible model as one in which:

i. the qpWave p-value is >0.05 (so that the Test population and the *N* sources are consistent with being derived from *N* waves of ancestry in relation to the outgroups),
ii. the standard error of the estimated admixture proportions is <10% (so that the admixture proportions are estimated reasonably tightly), and
iii. the admixture proportions are within 2 standard errors of the [0, 1] interval endpoints (so as not to penalize models with slightly <0 or >1 proportions that are consistent, respectively, with 0% or 100% of ancestry derived from one of the N sources, e.g., a model with a $-0.2\% \pm 3\%$ proportion of ancestry will be considered.

30/31 Test populations can be modeled with two sources, with a total of 768 feasible models. The only one that cannot be modeled is TTK which was inferred to be a mixture of Ancient North Eurasians and early Iranian food producers in ref. [13]. Thus, multiple 2-way models are feasible for many Test populations. These are not all independent but tend to involve similar population sources. We can assess these models by examining what is common between them and also by comparing them to each other.

#### A tournament approach for identifying the most plausible admixture models

How can we choose between competing models for the same population? We use the following "tournament" approach in which each fitting model A faces off against each other fitting model B in two matches:

- (A, B): A is fitted, and all source populations used in model B (except those sources that are shared with model A if any) are placed on the Right list. This tests the *resilience* of model A to B. If, for example, model A has a P-value of 0.67 when fitted using Right=Base, but this drops to 0.01 when the sources included in model B are added to the Right, then this raises a red flag about A.
- (B, A): conversely, B is fitted, and all populations of model A are placed on the Right list (except those sources that are shared with model B if any). This tests the *fortitude* of model A, in the sense that if it causes B to fit poorly, then this is problematic for B and an indication that A has some connection with the Test population that is not captured by the model B.

Thus, model A is attractive if it is resilient to other models placed on the Right while, at the same time, casting doubt on these other models when A itself is placed on the Right.

We may assess a model via a score formulated by subtracting its "losses" from its "wins" in the tournament. Whenever A is resilient to B, but B is not resilient to A, we count a "win" for A and a "loss" for B. In the symmetric case (when both models are resilient or not resilient to each other, we call a "draw" which does not help us decide between them). Clearly, losses, wins, and draws depend on the p-value threshold used, with more wins/losses awarded if this is higher and more draws if this is lower; we use a p-value threshold of 0.05 in our tournament.

It is important to note that this procedure provides no guarantee of choosing the right model. But, if a model consistently loses to other models, then it is in some sense questionable, whereas if it rarely loses then it can be provisionally considered further. Our tournament procedure also has the attractive feature that it does not simply favor source populations with less data that are more difficult to falsify which is a pitfall of qpAdm model fitting exercises that rely only on *resilience*; the *fortitude* criterion favors source populations that have more data, as these are the ones that have most power to falsify competing models.

Naturally, if populations are formed from entirely different sources from those sampled, then the tournament approach cannot discover the right model: it is limited to the available sources. Therefore, we consider the tournament approach as a semi-supervised and data-driven procedure of hypothesis filtering, taking hypotheses (fitting models) as input and ranking them in order to output those that seem worth additional exploration.

#### Tournament for fitting the core Yamnaya identifies a Remontnoye+Serednii Stih (SShi) model with high resilience and fortitude

We apply the tournament approach to the core Yamnaya for which a total of four 2-way models are feasible. These are listed in Table S 3. All these models involve SShi and an "eastern" population, either from the Caucasus (Unakozovskaya, Maikop, or Aknashen) or Remontnoye. The Remontnoye+SShi model has the highest p-value.

In the tournament, SShi+Remontnoye or SShi+Maikop models suffer no losses, while the other two models do. The full results of the tournament are shown in Table S 4. We note that the SShi+Remontnoye and SShi+Maikop models "draw" under our definition. However, the SShi+Maikop model fails convincingly (p=0.0002) when Remontnoye is placed on the Right, while the SShi+Remontnoye model does not when Maikop is placed on the Right (p=0.0368). As we will see below, Remontnoye itself can be modeled with Maikop as a source, so the two models are not inconsistent with each other.

| A | B | P-value | A | B | S.E. | Win | Draw | Lose | Score |
|---|---|---|---|---|---|---|---|---|---|
| **Remontnoye** | **SShi** | **0.675** | **26.3%** | **73.7%** | **3.4%** | **2** | **1** | **0** | **2** |
| **Maikop** | **SShi** | **0.068** | **13.1%** | **86.9%** | **2.1%** | **2** | **1** | **0** | **2** |
| Armenia_Aknashen_N | SShi | 0.075 | 11.3% | 88.7% | 1.9% | 0 | 1 | 2 | -2 |
| SShi | Unakozovskaya | 0.072 | 87.3% | 12.7% | 2.1% | 0 | 1 | 2 | -2 |

**Table S 3 The four fitting qpAdm models for core Yamnaya that have no more than two sources**.

| | **Remontnoye+SShi** | **Maikop+SShi** | Armenia_Aknashen_N+SShi | SShi+Unakozovskaya |
|---|---|---|---|---|
| **Remontnoye+SShi** | | 0.0002 | 0.0007 | 0.0012 |
| **Maikop+SShi** | 0.0368 | | 0.0005 | 0.0094 |
| Armenia_Aknashen_N+SShi | 0.6392 | 0.1060 | | 0.1090 |

| SShi+Unakozovskaya | 0.7880 | 0.0879 | 0.0600 |
|---|---|---|---|

**Table S 4 Model tournament for core Yamnaya.** Results of (A, B) matches are shown in A=columns and B=rows of the matrix. The Remontnoye+SShi model is highlighted in bold. Red coloring shows cases where the row model does not have good fortitude, that is, is not able to falsify the column model at the p<0.05 level.

#### The uniqueness of the Remontnoye+Serednii Stih blend in the ancestry of the Yamnaya

We next show the fits of the Remontnoye+SShi model in all considered Test populations (Table S 5). For most populations, this model fails with a low p-value. For GK1 the p-value is >0.05, but the admixture proportion of Remontnoye ancestry is significantly lower than zero. While these results suggest the mathematical possibility that the GK1 individuals were more extreme members of the Remontnoye-Core Yamnaya-SShi cline (beyond the SShi end of it), from the historical point of view they cannot have been formed by admixture of -58% Remontnoye- and 158% SShi-related ancestries but from an unsampled population beyond SShi which contributed some proportion within [0, 100]% of ancestry. Thus none of the Test populations (other than Core Yamnaya) have strong evidence of dual ancestries related to Remontnoye and SShi. Only the Core Yamnaya can be described as a mixture of these two sources.

| Test | P-value | Remontnoye | SShi | S.E. |
|---|---|---|---|---|
| Armenia_C | 3.28E-16 | 207.4% | -107.4% | 15.6% |
| Azerbaijan_C | 7.02E-09 | 226.1% | -126.1% | 16.0% |
| BPgroup | 2.90E-22 | -10.6% | 110.6% | 12.0% |
| CoreYamnaya | 6.75E-01 | 26.3% | 73.7% | 3.4% |
| Ekaterinovka | 8.71E-06 | -130.4% | 230.4% | 12.5% |
| GK1 | 8.61E-02 | -57.9% | 157.9% | 9.9% |
| GK2 | 1.89E-06 | -128.7% | 228.7% | 14.3% |
| Igren_o | 3.23E-03 | -124.9% | 224.9% | 16.5% |
| Kazakhstan_Kumsay_EBA | 4.28E-27 | -135.9% | 235.9% | 20.8% |
| Khi | 6.98E-13 | -49.4% | 149.4% | 9.3% |
| KhlopkovBugor | 2.70E-07 | -51.6% | 151.6% | 10.0% |
| Klo | 6.95E-05 | -96.0% | 196.0% | 10.3% |
| Kmed | 3.19E-08 | -73.1% | 173.1% | 10.1% |

| | | | | |
|---|---|---|---|---|
| Krivyansky | 1.53E-02 | 45.0% | 55.0% | 8.1% |
| Labazy | 3.50E-06 | -137.8% | 237.8% | 14.7% |
| Lebyazhinka_HG | 5.30E-05 | -153.2% | 253.2% | 16.3% |
| Maikop | 4.91E-06 | 188.9% | -88.9% | 11.5% |
| Maximovka | 2.29E-03 | -145.9% | 245.9% | 13.3% |
| Murzikha | 9.63E-04 | -151.2% | 251.2% | 13.9% |
| PVgroup | 2.19E-11 | 38.9% | 61.1% | 9.9% |
| Russia_Don_EBA_Yamnaya | 3.24E-06 | -3.5% | 103.5% | 5.7% |
| Russia_Steppe_Maikop | 2.66E-22 | -112.8% | 212.8% | 17.1% |
| SSlo | 3.20E-05 | -64.6% | 164.6% | 13.8% |
| SSmed | 6.56E-05 | -29.0% | 129.0% | 8.0% |
| Syezzheye | 9.56E-05 | -111.5% | 211.5% | 13.0% |
| TTK | 4.72E-31 | -273.1% | 373.1% | 47.0% |
| Unakozovskaya | 3.01E-05 | 193.1% | -93.1% | 11.9% |
| UpperVolga | 2.88E-03 | -144.5% | 244.5% | 13.2% |
| Usatove | 9.58E-32 | 215.7% | -115.7% | 24.8% |

**Table S 5 The Remontnoye+SShi model applied to all Test populations.** We highlight test populations with p>0.05.

#### The origins of Remontnoye: half Maikop/Aknashen Neolithic and half Lower Volga Eneolithic

Thus, we have arrived at the Remontnoye+SShi model as the only plausible solution, given the available sampled individuals, for the origins of the core Yamnaya population. It is possible that other such models would work but we lack the sources for them in our dataset; for example, models with SSmed+X as sources could be possible but we have not sampled an unknown "X" population. This is of course, a general problem in admixture modeling not unique to our study. However, with these limitations, we know for a fact that core Yamnaya are not a clade with any of the Serednii Stih subsets and Remontnoye is one source of extra-Serednii Stih ancestry that results in a successful model. So, regardless of what other possible combinations may have occurred historically, investigating the origins of Remontnoye is important as its composition will contain forms of ancestry not represented in the sampled Serednii Stih. Thus, we turn to the question of the origins of Remontnoye itself.

We show in Table S 6 the feasible models and the results of the tournament comparing them. The models are all quite similar; they involve one population from the south (either Maikop, from the North Caucasus, or Aknashen Neolithic from Armenia or Azerbaijan Neolithic/Chalcolithic, in the South Caucasus) and one population from the north (BPgroup or PVgroup or Steppe Maikop or WSHG). The best fitting models both in terms of the tournament score and in terms of p-value involve BPgroup and either Aknashen Neolithic or Maikop.

The results of the full tournament are presented in Table S 7. The best fitting models with greatest difference between wins and losses the BPgroup+Aknashen or BPgroup+Azerbaijan ones, which the BPgroup+Maikop model is almost as good (one fewer win and one more draw) and is also geographically and temporally more plausible. Strictly speaking, however, the non-BPgroup-related source for the Remontnoye must ultimately have had genetic roots further south, which is consistent with the observation that the models with Neolithic southern Caucasus sources have the most wins in the tournament. The models with either Steppe Maikop or WSHG as the northern source consistently fail in the competition. When did the Aknashen-related ancestry reach the north Caucasus and via it steppe populations like Remontnoye? All sampled individuals from the north Caucasus are from the 5th millennium BCE or later, while Aknashen is from the beginning of the 6th millennium BCE. We know that the native population of the South Caucasus consisted of Caucasus hunter-gatherers as late as the Kotias individual[19] of the mid-8th millennium BCE. Thus, the genetic Neolithization of the South Caucasus must have occurred sometime between the 8th and 5th millennium BCE, i.e., during the flourishing of the Shulaveri–Shomu culture Neolithic culture. This ancestry could then have reached the north Caucasus by the time of the Unakozovskaya individuals (ref.[8] and this study) of the mid-5th millennium BCE. Thus, the time window of ~6000-4500BCE appears to be a plausible time frame for the spread of Aknashen-related ancestry into the North Caucasus. We note that Aknashen itself differs from the Masis Blur Neolithic of a few centuries later (mid-6th millennium BCE),[4] in which the CHG-related ancestry is greatly reduced. Thus, it seems probable that the Aknashen-related ancestry reached the North Caucasus before the 5633-5532 calBCE date of the Masis Blur individual when the (undiluted) Aknashen-related ancestry still existed in the South Caucasus.

Regardless of the temporality of this process, all fitting models agree on is that Remontnoye was formed as a mixture of people of the Caucasus-Volga north (BPgroup or PVgroup) and Neolithic-Chalcolithic south (from Maikop or Armenia-Azerbaijan).

| A | B | P-value | A | B | S.E. | Win | Draw | Lose | Score |
|---|---|---|---|---|---|---|---|---|---|
| **Armenia_Aknashen_N** | **BPgroup** | **0.661** | **44.6%** | **55.4%** | **2.7%** | **4** | **5** | **0** | **4** |
| **Azerbaijan_N** | **PVgroup** | **0.118** | **33.2%** | **66.8%** | **2.9%** | **4** | **5** | **0** | **4** |
| **Maikop** | **BPgroup** | **0.445** | **51.9%** | **48.1%** | **2.9%** | **3** | **6** | **0** | **3** |
| Azerbaijan_C | BPgroup | 0.143 | 42.8% | 57.2% | 2.6% | 2 | 7 | 0 | 2 |
| Maikop | PVgroup | 0.248 | 45.0% | 55.0% | 3.7% | 2 | 6 | 1 | 1 |
| Armenia_Aknashen_N | PVgroup | 0.290 | 38.0% | 62.0% | 3.4% | 2 | 4 | 3 | -1 |
| Azerbaijan_C | PVgroup | 0.227 | 35.8% | 64.2% | 3.3% | 0 | 8 | 1 | -1 |
| Unakozovskaya | WSHG | 0.102 | 74.7% | 25.3% | 1.7% | 0 | 5 | 4 | -4 |
| Unakozovskaya | Russia_Steppe_Maikop | 0.091 | 65.1% | 34.9% | 2.2% | 0 | 5 | 4 | -4 |
| Armenia_MasisBlur_N | PVgroup | 0.085 | 29.5% | 70.5% | 2.8% | 0 | 5 | 4 | -4 |

**Table S 6 Considered 2-way models for Remontnoye**

| | **Armenia_Aknashen_N+BPgroup** | **Azerbaijan_N+PVgroup** | **BPgroup+Maikop** | Azerbaijan_C+BPgroup | Maikop+PVgroup | Armenia_Aknashen_N+PVgroup | Azerbaijan_C+PVgroup | Unakozovskaya+WSHG | Russia_Steppe_Maikop+Unakozovskaya | Armenia_MasisBlur_N+PVgroup |
|---|---|---|---|---|---|---|---|---|---|---|
| **Armenia_Aknashen_N+BPgroup** | | 3.12E-03 | 4.16E-01 | 1.88E-01 | 1.54E-01 | 3.70E-02 | 1.67E-01 | 2.49E-07 | 1.21E-03 | 3.43E-04 |
| **Azerbaijan_N+PVgroup** | 4.82E-02 | | 4.39E-02 | 6.14E-03 | 3.61E-02 | 2.92E-02 | 8.86E-03 | 1.64E-04 | 1.33E-02 | 2.06E-02 |
| **Maikop+BPgroup** | 7.59E-01 | 6.20E-03 | | 8.76E-02 | 1.64E-01 | 6.02E-02 | 1.87E-01 | 2.24E-08 | 1.43E-04 | 6.03E-04 |
| Azerbaijan_C+BPgroup | 7.12E-01 | 5.58E-03 | 2.12E-01 | | 8.20E-02 | 4.95E-02 | 1.45E-01 | 2.72E-07 | 1.44E-03 | 2.17E-04 |
| Maikop+PVgroup | 3.47E-01 | 1.69E-01 | 2.06E-01 | 7.89E-02 | | 3.48E-01 | 3.18E-01 | 1.04E-04 | 4.64E-03 | 1.33E-01 |
| Armenia_Aknashen_N+PVgroup | 2.72E-01 | 1.32E-01 | 1.85E-01 | 1.81E-01 | 2.47E-01 | | 2.68E-01 | 5.33E-04 | 2.73E-02 | 9.62E-02 |
| Azerbaijan_C+PVgroup | 3.42E-01 | 1.65E-01 | 8.80E-02 | 1.42E-01 | 1.38E-01 | 3.33E-01 | | 4.69E-04 | 3.29E-02 | 6.64E-02 |
| Unakozovskaya+WSHG | 2.48E-01 | 3.44E-02 | 5.81E-01 | 2.35E-03 | 3.26E-01 | 1.33E-01 | 2.29E-02 | | 9.51E-02 | 1.27E-02 |

| | | | | | | | | | | |
|---|---|---|---|---|---|---|---|---|---|---|
| Russia_Steppe_Maikop+Unakozovskaya | 1.83E-01 | 3.33E-02 | 3.26E-01 | 2.06E-03 | 2.21E-01 | 1.09E-01 | 2.11E-02 | 6.27E-02 | | 1.33E-02 |
| Armenia_MasisBlur_N+PVgroup | 3.24E-01 | 1.06E-01 | 1.38E-01 | 1.08E-01 | 1.61E-01 | 3.12E-01 | 1.75E-01 | 1.39E-04 | 1.30E-02 | |

**Table S 7 Model tournament for Remontnoye.** Results of (A, B) matches are shown in A=columns and B=rows of the matrix.

#### The origins of Maikop: Aknashen Neolithic with minor BPgroup influence

The Maikop population is very similar to that of Aknashen and the only feasible models for Maikop involve admixture between Aknashen and populations of eastern Europe.

| A | B | P-value | A | B | S.E. |
|---|---|---|---|---|---|
| **Armenia_Aknashen_N** | **BPgroup** | **0.502** | **86.2%** | **13.8%** | **2.9%** |
| Armenia_Aknashen_N | Ekaterinovka | 0.274 | 91.9% | 8.1% | 2.0% |
| Armenia_Aknashen_N | GK1 | 0.531 | 89.0% | 11.0% | 2.4% |
| Armenia_Aknashen_N | GK2 | 0.351 | 91.4% | 8.6% | 2.0% |
| Armenia_Aknashen_N | Igren_o | 0.323 | 91.5% | 8.5% | 2.0% |
| Armenia_Aknashen_N | Khi | 0.425 | 88.5% | 11.5% | 2.6% |
| Armenia_Aknashen_N | KhlopkovBugor | 0.375 | 89.2% | 10.8% | 2.5% |
| Armenia_Aknashen_N | Klo | 0.343 | 90.8% | 9.2% | 2.0% |
| Armenia_Aknashen_N | Kmed | 0.356 | 89.9% | 10.1% | 2.2% |
| Armenia_Aknashen_N | Krivyansky | 0.761 | 82.1% | 17.9% | 3.4% |
| Armenia_Aknashen_N | Labazy | 0.244 | 92.1% | 7.9% | 1.9% |
| Armenia_Aknashen_N | Lebyazhinka_HG | 0.302 | 92.4% | 7.6% | 1.9% |
| Armenia_Aknashen_N | Murzikha | 0.332 | 92.2% | 7.8% | 1.7% |
| Armenia_Aknashen_N | PVgroup | 0.467 | 84.5% | 15.5% | 3.4% |
| Armenia_Aknashen_N | Remontnoye | 0.451 | 75.2% | 24.8% | 5.0% |
| Armenia_Aknashen_N | Russia_Karelia | 0.234 | 92.9% | 7.1% | 1.7% |
| Armenia_Aknashen_N | Russia_Steppe_Maikop | 0.440 | 90.0% | 10.0% | 2.3% |
| Armenia_Aknashen_N | SSlo | 0.497 | 89.1% | 10.9% | 2.4% |
| Armenia_Aknashen_N | SSmed | 0.475 | 87.6% | 12.4% | 2.7% |
| Armenia_Aknashen_N | SShi | 0.520 | 86.2% | 13.8% | 2.9% |
| Armenia_Aknashen_N | TTK | 0.226 | 92.5% | 7.5% | 1.8% |
| Armenia_Aknashen_N | Ukraine_N | 0.286 | 91.5% | 8.5% | 1.9% |
| Armenia_Aknashen_N | UpperVolga | 0.279 | 92.2% | 7.8% | 1.9% |
| Armenia_Aknashen_N | WSHG | 0.261 | 92.9% | 7.1% | 1.6% |

**Table S 8 Models fitting the Maikop**

Model competition does not help us distinguish between these models, but they all agree in that the Maikop population is very similar to Aknashen but with some additional northern influence. In particular, the model that fits the Remontnoye (BPgroup+Aknashen) also fits the Maikop, but with substantially more Aknashen-related ancestry (86.2±2.9% in Maikop vs. 44.6±2.7% in Remontnoye).

The Maikop group here includes individuals: OSS001, OSS002.B0101, I1720, I6266, I6267, I6268, I6272. We separately analyzed I4429 (a PCA ancestry outlier) which we left out of the main group in our fitting analyses. We show the fits of the Aknashen+BPgroup model for all Maikop individuals (Table S 9), demonstrating the relative homogeneity of the Maikop label and the poor fit and higher BPgroup-related ancestry of the outlier individuals that we did not include in the Maikop label used for the model fitting.

| Individual | P-value | Aknashen | BPgroup | S.E. | Label |
|---|---|---|---|---|---|
| I4429 | 1.47E-03 | 59.8% | 40.2% | 3.9% | Maikop_I4429 |
| I6266 | 1.09E-01 | 84.3% | 15.7% | 4.2% | Maikop |
| I6267 | 2.20E-01 | 84.3% | 15.7% | 4.5% | Maikop |
| I1720_wNonUDG | 2.92E-01 | 85.4% | 14.6% | 6.8% | Maikop |
| OSS001 | 3.63E-01 | 86.6% | 13.4% | 4.2% | Maikop |
| I6268 | 7.06E-01 | 88.4% | 11.6% | 4.2% | Maikop |
| I6272 | 6.92E-01 | 89.9% | 10.1% | 4.2% | Maikop |
| OSS002.B0101 | 9.80E-03 | 95.1% | 4.9% | 6.3% | Maikop |

**Table S 9 Model fits for Maikop individuals**

Individual I4429 (Novosvobodnaya, Tsarskaya; 3700-3000 BCE) does not fit the model well but has more BPgroup-related ancestry than those of the main Maikop cluster, consistent also with his position in PCA space (Fig. S 1). This suggested to us that perhaps his northern ancestry is not well-modeled as BPgroup and some other population from the north is a better fit for them. Indeed, we can model I4429 with Aknashen as one source and different populations as the second source. We list feasible models in Table S 10 which show that the northern ancestry is better modeled as Lebyazhinka_HG or other populations of high hunter-gatherer ancestry.

| A | P-value | Aknashen | A | S.E. |
|---|---|---|---|---|
| Lebyazhinka_HG | 0.391 | 74.6% | 25.4% | 2.3% |
| Murzikha | 0.273 | 75.2% | 24.8% | 2.4% |
| Klo | 0.244 | 70.5% | 29.5% | 2.7% |
| Russia_Karelia | 0.227 | 76.6% | 23.4% | 2.3% |
| SSmed | 0.205 | 61.9% | 38.1% | 3.4% |
| Ekaterinovka | 0.204 | 73.5% | 26.5% | 2.5% |
| GK1 | 0.198 | 66.9% | 33.1% | 3.2% |
| Kmed | 0.176 | 67.8% | 32.2% | 2.9% |
| Labazy | 0.172 | 73.9% | 26.1% | 2.5% |
| SShi | 0.145 | 58.3% | 41.7% | 3.8% |
| Igren_o | 0.117 | 72.5% | 27.5% | 2.9% |
| UpperVolga | 0.101 | 75.2% | 24.8% | 2.3% |

**Table S 10 Modeling Maikop outlier I4429**

The admixture in Remontnoye is geographically plausible: Maikop is modeled as having formed from more ancient populations of the Caucasus (similar to the earliest Neolithic of the southern Caucasus sampled in Aknashen in Armenia), but is in contact and experienced admixture with steppe Eneolithic populations like BPgroup which geographically spanned at least the area from the Lower Volga (where the four Berezhnovka individuals were sampled) and the North Caucasus piedmont (the site of Progress-2). Remontnoye is the result of one such admixture, with roughly half of its ancestry modeled as being of Maikop/Aknashen origin, and the other half derived from the BPgroup.

The Unakozovskaya Pre-Maikop individuals from the North Caucasus predate the Maikop. These include three relatives published in ref. [8] of which the higher quality I2056 is used here, together with individual I1717. The Unakozovskaya population is not a clade with Maikop (p=2e-11), because it shares much more genetic drift with CHG, as evidenced by the statistic $f_4$(Maikop, Unakozovskaya; CHG, OldAfrica) which has a Z-score of -6.0. However, it can be modeled as a mixture of Maikop and CHG (p=0.46) with predominantly Maikop-related ancestry (95.3±6.4%) and conversely Maikop can be modeled as 105.4±7.1% Unakozovskaya with a negative CHG contribution. Thus, Unakozovskaya (the precursors of Maikop) were similar if not quite like them. The Aknashen+BPgroup model does not fit Unakzovskaya (p=6e-9) as it underestimates CHG shared drift as well (Z=-4.8). Therefore, Unakozovskaya was not quite on the Aknashen-BPgroup (Caucasus-Lower Volga) cline but occupied a position similar to the later Maikop, offset by higher CHG affinity.

#### Chalcolithic Armenia: Masis Blur Neolithic plus steppe ancestry

Ever since its publication[5] it has been known that the Chalcolithic of Armenia at Areni-1 cave harbored "northern"/EHG ancestry that was not found in other individuals from Armenia before the Middle Bronze Age.[4] We thus included Armenia_C as a Test population in our analysis to obtain insights into which more proximate sources of steppe ancestry it possessed.

In contrast to the Remontnoye (Table S 6) and Maikop (Table S 8) population just discussed for which only Aknashen Neolithic was feasible as the local (Caucasus) source, for the Areni-1 Chalcolithic only Masis Blur Neolithic and the populations from Azerbaijan result in feasible models. Recall that Masis Blur and Aknashen Neolithic are highly differentiated populations of Neolithic Armenia, with the earlier Aknashen having more CHG and the later Masis Blur more Anatolian-Levantine ancestry, while those from Azerbaijan are intermediate[4] The feasible models for Armenia_C are shown in
Table S 11: they all involve Masis Blur Neolithic with a northern source. (The results of the tournament are not conclusive for determining the northern source and are not shown).

| A | B | P-value | A | B | S.E. | | A=Aknashen |
|---|---|---|---|---|---|---|---|
| Armenia_MasisBlur_N | BPgroup | 0.845 | 73.1% | 26.9% | 2.3% | | 1.83E-10 |
| Armenia_MasisBlur_N | Ekaterinovka | 0.124 | 81.5% | 18.5% | 1.7% | | 1.22E-05 |
| Armenia_MasisBlur_N | GK1 | 0.061 | 77.1% | 22.9% | 2.2% | | 9.20E-06 |
| Armenia_MasisBlur_N | Khi | 0.982 | 75.9% | 24.1% | 2.0% | | 9.52E-09 |
| Armenia_MasisBlur_N | KhlopkovBugor | 0.959 | 76.9% | 23.1% | 2.0% | | 1.22E-08 |
| Armenia_MasisBlur_N | Klo | 0.347 | 79.6% | 20.4% | 1.8% | | 2.74E-06 |
| Armenia_MasisBlur_N | Kmed | 0.750 | 77.9% | 22.1% | 1.9% | | 2.64E-07 |
| Armenia_MasisBlur_N | Krivyansky | 0.286 | 68.1% | 31.9% | 2.8% | | 2.05E-10 |
| Armenia_MasisBlur_N | Labazy | 0.095 | 81.9% | 18.1% | 1.7% | | 2.04E-05 |
| Armenia_MasisBlur_N | PVgroup | 0.581 | 70.8% | 29.2% | 2.5% | | 4.89E-11 |
| Armenia_MasisBlur_N | Remontnoye | 0.119 | 59.5% | 40.5% | 3.4% | | 1.12E-11 |
| Armenia_MasisBlur_N | Russia_Steppe_Maikop | 0.565 | 79.0% | 21.0% | 1.9% | | 1.48E-08 |
| Armenia_MasisBlur_N | SShi | 0.639 | 72.0% | 28.0% | 2.4% | | 1.09E-07 |
| Armenia_MasisBlur_N | TTK | 0.072 | 83.5% | 16.5% | 1.5% | | 2.79E-09 |
| Armenia_MasisBlur_N | WSHG | 0.082 | 84.1% | 15.9% | 1.5% | | 3.40E-07 |
| Azerbaijan_C | Ekaterinovka | 0.066 | 87.6% | 12.4% | 2.1% | | 1.22E-05 |
| Azerbaijan_C | Igren_o | 0.085 | 87.0% | 13.0% | 2.3% | | 8.30E-04 |
| Azerbaijan_C | Labazy | 0.089 | 87.6% | 12.4% | 2.0% | | 2.04E-05 |
| Azerbaijan_C | Lebyazhinka_HG | 0.078 | 88.2% | 11.8% | 1.9% | | 1.09E-04 |
| Azerbaijan_C | Murzikha | 0.072 | 88.2% | 11.8% | 2.0% | | 1.68E-04 |
| Azerbaijan_C | Russia_Karelia | 0.089 | 88.8% | 11.2% | 1.9% | | 9.07E-05 |
| Azerbaijan_C | UpperVolga | 0.102 | 87.8% | 12.2% | 2.0% | | 2.32E-04 |
| Azerbaijan_N | Ekaterinovka | 0.180 | 85.5% | 14.5% | 1.4% | | 1.22E-05 |
| Azerbaijan_N | Khi | 0.057 | 81.5% | 18.5% | 1.7% | | 9.52E-09 |
| Azerbaijan_N | KhlopkovBugor | 0.087 | 82.2% | 17.8% | 1.7% | | 1.22E-08 |
| Azerbaijan_N | Klo | 0.185 | 84.1% | 15.9% | 1.5% | | 2.74E-06 |

| | | | | | | | |
|---|---|---|---|---|---|---|---|
| Azerbaijan_N | Kmed | 0.160 | 82.8% | 17.2% | 1.6% | | 2.64E-07 |
| Azerbaijan_N | Labazy | 0.238 | 85.7% | 14.3% | 1.3% | | 2.04E-05 |
| Azerbaijan_N | Lebyazhinka_HG | 0.085 | 86.6% | 13.4% | 1.4% | | 1.09E-04 |
| Azerbaijan_N | Russia_Karelia | 0.097 | 87.2% | 12.8% | 1.3% | | 9.07E-05 |

**Table S 11 Models fitting the Areni-1 Chalcolithic of Armenia.** In the rightmost column we show the p-values for the same models with Aknashen Neolithic as the source (all of which fail to model the Chalcolithic of Armenia).

These results show that sampled Armenian Chalcolithic people had evidence for admixture with steppe people but that it represents a different mixture from that in the North Caucasus. The Armenian Chalcolithic follows in the footsteps of the Masis Blur Neolithic that preceded it, but with additional steppe influences.

By contrast, in Maikop and Remontnoye the Caucasus ancestry was similar to that in Aknashen Neolithic (Table S 8; the Masis Blur + BPgroup model is rejected for Maikop; p=3e-09). We hypothesize that Aknashen-like ancestry was widely distributed in the Caucasus, hence its presence in the Maikop and via the Maikop in Remontnoye, and via Remontnoye in Yamnaya. By contrast, Armenia was the recipient of later additional Anatolian-Levantine ancestry represented by the Masis Blur individual.[4] Hence, Armenia_C is a poor distal source for the Maikop/Remontnoye/Yamnaya; it is a genetic dead-end, not relevant to what happened later in the Steppe.

#### The origins of the Lower Volga-North Caucasus Eneolithic: more CHG-admixed than the rest of the Volga cline

We have so far shown that core Yamnaya was formed by admixture between SShi- and Remontnoye-related sources, and that Remontnoye was consistent with being part of a cline (together with Maikop) of admixture between people of the Caucasus (most robustly represented by Aknashen Neolithic) and people of the north (most robustly represented by BPgroup). What of the origins of BPgroup?

Several models fit the ancestry of BPgroup (Table S 12). The only model that does not lose any matches is the one that involves Khlopkov Bugor and PVgroup ancestries. All but one models involve ancestry from the north (Khi, Kmed, or Khlopkov Bugor) and southern ancestry (CHG or PVgroup). The one exception is the Krivyansky+TTK model which captures a west (Don/Krivyansky) to east (Central Asia/TTK) distinction. This model loses many matches in the tournament, but this is because TTK is from a Central Asian ancestry source that likely diverged from the TTK-related ancestors of BP-group in Neolithic times at the most recent, and thus underestimates later shared genetic drift between BPgroup and other Eneolithic populations of the Volga.

| A | B | P-value | A | B | S.E. | Win | Draw | Lose | Score |
|---|---|---|---|---|---|---|---|---|---|
| **KhlopkovBugor** | **PVgroup** | **0.217** | **28.8%** | **71.2%** | **6.2%** | **4** | **2** | **0** | **4** |
| **Khi** | **PVgroup** | **0.088** | **31.9%** | **68.1%** | **7.2%** | **5** | **0** | **1** | **4** |
| WSHG | PVgroup | 0.127 | 10.1% | 89.9% | 2.5% | 1 | 4 | 1 | 0 |
| Kmed | PVgroup | 0.055 | 19.9% | 80.1% | 4.8% | 1 | 4 | 1 | 0 |
| Russia_Steppe_Maikop | PVgroup | 0.120 | 19.3% | 80.7% | 4.6% | 1 | 3 | 2 | -1 |
| Khi | CHG | 0.159 | 78.6% | 21.4% | 1.9% | 0 | 4 | 2 | -2 |

| | | | | | | | | | |
|---|---|---|---|---|---|---|---|---|---|
| Krivyansky | TTK | 0.128 | 76.3% | 23.7% | 3.0% | 0 | 1 | 5 | -5 |

**Table S 12 Considered 2-way models for BPgroup**

We examine the models for PVgroup (Table S 13). We note that this population too fits the CHG+Khi model albeit with a higher proportion of CHG ancestry compared to BPgroup. When BPgroup is the source, then additional ancestry from the Caucasus is included in the successful models.

| A | B | P-value | A | B | S.E. | Win | Draw | Lose | Score |
|---|---|---|---|---|---|---|---|---|---|
| Armenia_Aknashen_N | BPgroup | 0.753 | 11.1% | 88.9% | 2.5% | 3 | 8 | 0 | 3 |
| Armenia_C | BPgroup | 0.274 | 10.9% | 89.1% | 2.6% | 3 | 8 | 0 | 3 |
| Armenia_MasisBlur_N | BPgroup | 0.358 | 8.2% | 91.8% | 1.8% | 3 | 8 | 0 | 3 |
| Azerbaijan_C | BPgroup | 0.485 | 10.0% | 90.0% | 2.3% | 3 | 8 | 0 | 3 |
| Azerbaijan_N | BPgroup | 0.376 | 9.2% | 90.8% | 2.1% | 3 | 8 | 0 | 3 |
| CHG | BPgroup | 0.247 | 14.7% | 85.3% | 3.4% | 3 | 8 | 0 | 3 |
| Maikop | BPgroup | 0.655 | 12.7% | 87.3% | 2.7% | 3 | 8 | 0 | 3 |
| Remontnoye | BPgroup | 0.487 | 23.7% | 76.3% | 5.4% | 3 | 8 | 0 | 3 |
| Unakozovskaya | BPgroup | 0.350 | 12.4% | 87.6% | 2.9% | 3 | 8 | 0 | 3 |
| CHG | Khi | 0.257 | 32.7% | 67.3% | 2.6% | 2 | 0 | 9 | -7 |
| CHG | KhlopkovBugor | 0.072 | 36.2% | 63.8% | 2.8% | 0 | 1 | 10 | -10 |
| CHG | Kmed | 0.064 | 42.2% | 57.8% | 2.4% | 0 | 1 | 10 | -10 |

**Table S 13 Considered 2-way models for PVgroup**

To summarize: BPgroup and PVgroup are populations with more CHG ancestry than Khi: PVgroup more than BPgroup. These are then transitional populations between the Caucasus and the Volga cline that share ancestry with each other but also differ in terms of their relative proportions of CHG ancestry. A simple test for cladality shows that they are different more directly (BPgroup and PVgroup p=0.0006). BPgroup and PVgroup can be seen as belonging to a variable Lower Volga-North Caucasus Eneolithic population. The two groups are clearly similar to each other, fitting many of the same models with different proportions. As we have seen, members of this meta-population was mixing with the descendants of Aknashen-related Neolithic farmers in the North Caucasus piedmont to form populations like the Maikop (Table S 8) which was mostly derived from the farmers and Remontnoye (Table S 6) in which the two components were balanced.

#### Modeling the Volga Cline as a whole: admixture between Lower Volga Eneolithic and upriver Eastern hunter-gatherers

We have seen that Khi and Khlopkov Bugor are cladal to each other, and also that Ekaterinovka, Labazy, and Syezzheye are also cladal. We have also seen that BPgroup, at the end of the Volga cline can be modeled as Khi (Khvalynsk subset) with extra CHG ancestry. We now try to study the ancestry of Volga populations as a whole from the following set.

**Volga:** BPgroup, Ekaterinovka, Khi, KhlopkovBugor, Klo, Kmed, Labazy, Lebyazhinka_HG, Maximovka, Murzikha, Syezzheye, UpperVolga

As some of these populations (notably Upper Volga and Murzikha) are upriver compared to the rest and noticeably not clinal in PCA (Fig. S 1) with the rest, we expect them to be of possibly different ancestral origins than the main cline which extends from BPgroup on the Lower Volga through Khvalynsk, Ekaterinovka and the populations similar to them, and finally to EHG.

To find a common model for Volga populations, we take all pairs A, B of the Sources and tabulated how many of them Volga groups fit as A+B mixtures. As there are 12 target populations in total, we expect that—if the two sources are from the Volga itself—then a maximum of 10 (excluding A and B) could possibly be fitted. As there are at least two populations notably off cline in PCA (Upper Volga and Murzikha) we expect that realistically we will be able to model only 8 using this automated procedure.

We find two models (with either Khi or Khlopkov Bugor and Lebyazhinka_HG as the sources) that fit 8 populations and one (with BPgroup and Lebyazhinka_HG as the sources) that fit 7 (Table S 14). None of the models fit Upper Volga (the most off-cline population). Murzikha can be modeled with p-values barely above the 0.05 threshold and with a negative coefficient of Khi/Khlopkov Bugor ancestry and so is not convincingly on the cline. The models with Khi or Khlopkov Bugor as a source cannot model Bpgroup which has a substantial (~1/3-1/2) negative coefficient of Lebyazhinka ancestry, i.e., it has substantially less EHG ancestry than the Khvalynsk/Khlopkov Bugor populations.

The BPgroup+Lebyazhinka_HG model is successful for all Volga populations except the off-cline Upper Volga and Murzikha, and Khlopkov Bugor with p=0.042. Admixture proportions under this model are as one might expect from the PCA: ~3/4 BPgroup ancestry in the Khi subset of Khvalynsk down to <1/4 in Labazy, Syezzheye, and Ekaterinovka. While the marginally poor fit for Khlopkov Bugor might in theory be a statistical fluctuation due to the substantial number of groups tested, we also observed a marginally passing fit for the genetically similar Khi (p=0.061), and we believe this is likely due to real but subtle deviation from a simple two-way cline, a point we return to in what follows.

Overall, the Volga cline can be seen as largely driven by BPgroup-Lebyazhinka admixture tracing the Volga river from its downstream BPgroup end, to groups with variable ancestry in the Middle Volga ranging from ones with high BP relatedness (such as Khi), to predominant EHG ancestry (such as Ekaterinovka).

| Test | A | B | P-value | A | B | S.E. |
|---|---|---|---|---|---|---|
| Ekaterinovka | BPgroup | Lebyazhinka_HG | 3.31E-01 | 17.0% | 83.0% | 3.4% |
| Khi | BPgroup | Lebyazhinka_HG | 6.09E-02 | 74.7% | 25.3% | 2.3% |
| KhlopkovBugor | BPgroup | Lebyazhinka_HG | 4.20E-02 | 68.4% | 31.6% | 3.3% |
| Klo | BPgroup | Lebyazhinka_HG | 5.02E-01 | 35.2% | 64.8% | 2.8% |
| Kmed | BPgroup | Lebyazhinka_HG | 3.76E-01 | 53.6% | 46.4% | 2.6% |
| Labazy | BPgroup | Lebyazhinka_HG | 3.18E-01 | 13.1% | 86.9% | 4.3% |
| Maximovka | BPgroup | Lebyazhinka_HG | 4.66E-01 | 2.1% | 97.9% | 4.5% |
| Murzikha | BPgroup | Lebyazhinka_HG | 4.64E-02 | -3.5% | 103.5% | 4.1% |
| Syezzheye | BPgroup | Lebyazhinka_HG | 2.83E-01 | 24.4% | 75.6% | 4.0% |
| UpperVolga | BPgroup | Lebyazhinka_HG | 1.42E-07 | -8.1% | 108.1% | 4.3% |
| BPgroup | Khi | Lebyazhinka_HG | 6.23E-02 | 134.0% | -34.0% | 4.1% |
| Ekaterinovka | Khi | Lebyazhinka_HG | 4.21E-01 | 23.1% | 76.9% | 4.1% |
| KhlopkovBugor | Khi | Lebyazhinka_HG | 2.91E-01 | 92.8% | 7.2% | 4.3% |
| Klo | Khi | Lebyazhinka_HG | 4.94E-01 | 47.4% | 52.6% | 3.3% |
| Kmed | Khi | Lebyazhinka_HG | 5.96E-01 | 72.4% | 27.6% | 2.9% |
| Labazy | Khi | Lebyazhinka_HG | 3.67E-01 | 18.0% | 82.0% | 5.7% |
| Maximovka | Khi | Lebyazhinka_HG | 4.54E-01 | 2.2% | 97.8% | 6.0% |
| Murzikha | Khi | Lebyazhinka_HG | 5.54E-02 | -6.1% | 106.1% | 5.9% |
| Syezzheye | Khi | Lebyazhinka_HG | 2.75E-01 | 32.6% | 67.4% | 5.0% |
| UpperVolga | Khi | Lebyazhinka_HG | 5.14E-07 | -14.6% | 114.6% | 6.4% |
| BPgroup | KhlopkovBugor | Lebyazhinka_HG | 4.27E-02 | 146.5% | -46.5% | 7.2% |
| Ekaterinovka | KhlopkovBugor | Lebyazhinka_HG | 6.25E-01 | 25.4% | 74.6% | 4.4% |
| Khi | KhlopkovBugor | Lebyazhinka_HG | 2.91E-01 | 108.0% | -8.0% | 5.0% |

| | | | | | | |
|---|---|---|---|---|---|---|
| Klo | KhlopkovBugor | Lebyazhinka_HG | 6.84E-01 | 51.0% | 49.0% | 4.0% |
| Kmed | KhlopkovBugor | Lebyazhinka_HG | 4.51E-01 | 77.3% | 22.7% | 4.0% |
| Labazy | KhlopkovBugor | Lebyazhinka_HG | 4.87E-01 | 20.3% | 79.7% | 6.1% |
| Maximovka | KhlopkovBugor | Lebyazhinka_HG | 4.68E-01 | 3.0% | 97.0% | 6.3% |
| Murzikha | KhlopkovBugor | Lebyazhinka_HG | 5.91E-02 | -7.0% | 107.0% | 6.5% |
| Syezzheye | KhlopkovBugor | Lebyazhinka_HG | 4.96E-01 | 36.2% | 63.8% | 5.7% |
| UpperVolga | KhlopkovBugor | Lebyazhinka_HG | 6.83E-07 | -17.0% | 117.0% | 7.7% |

**Table S 14 Modeling the Volga cline.** We show the 30 models that involve Lebyazhinka_HG as one source and Khi, KhlopkovBugor, or BPgroup as the other as these are feasible for most (7-8) populations of the Volga Cline. Other models are feasible for 6 or fewer populations and are not shown for brevity, as they are either feasible for fewer populations or describe shorter subsets of the cline (e.g., Kmed+Lebyazhinka_HG is infeasible for the less EHG-admixed Khi subset with an inferred -38.3±5.5% Lebyazhinka_HG contribution).

We were intrigued by the marginal p-values of the fits for Khi and Khlopkov Bugor when BPgroup was used as a source. To better understand how these three populations at the low-EHG end of the Volga Cline came about, we examined the fits of 3-way models for the Volga cline populations. A single model (Table S 15) had the property of fitting BPgroup and most other Volga populations (except Upper Volga, Murzikha, and Maximovka).

The notable feature of this model is that it does not only include CHG/EHG (represented by Lebyazhinka_HG) ancestry in the Volga Cline, i.e., the proximate hunter-gatherer sources of south and north / Caucasus-steppe area, but also includes ancestry from the Central Asian TTK-related source. The presence of this ancestry was hinted already in our modeling of BPgroup in which its presence in the Lower Volga was inferred in contrast to the Lower Don individual from Krivyansky (Table S 12). In Table S 15 we see its presence more directly in terms of the distal 3-way model.

| | | Proportions | | | Std. errors | | |
|---|---|---|---|---|---|---|---|
| **Test** | **P-value** | **A=CHG** | **B=Lebyazhinka_HG** | **C=TTK** | **A** | **B** | **C** |
| BPgroup | 0.228 | 47.6% | 36.7% | 15.7% | 1.6% | 2.9% | 3.2% |
| Ekaterinovka | 0.268 | 7.4% | 89.6% | 3.0% | 2.2% | 4.0% | 4.6% |
| Khi | 0.103 | 33.8% | 50.3% | 15.9% | 1.6% | 3.0% | 3.4% |
| KhlopkovBugor | 0.374 | 30.4% | 55.8% | 13.8% | 2.2% | 4.0% | 4.4% |
| Klo | 0.470 | 17.1% | 80.3% | 2.6% | 2.2% | 3.9% | 4.4% |
| Kmed | 0.444 | 24.4% | 64.6% | 11.0% | 1.8% | 3.4% | 3.7% |
| Labazy | 0.260 | 5.4% | 92.8% | 1.8% | 2.7% | 5.1% | 5.6% |
| Maximovka | 0.642 | 2.7% | 104.5% | -7.2% | 2.9% | 5.1% | 5.7% |
| Murzikha | 0.309 | 2.9% | 111.8% | -14.7% | 2.8% | 5.0% | 5.6% |
| Syezzheye | 0.310 | 11.7% | 87.5% | 0.8% | 2.7% | 5.0% | 5.8% |
| UpperVolga | 0.354 | 7.4% | 124.1% | -31.4% | 3.2% | 5.4% | 6.2% |

**Table S 15 A 3-way model of the Volga cline.**

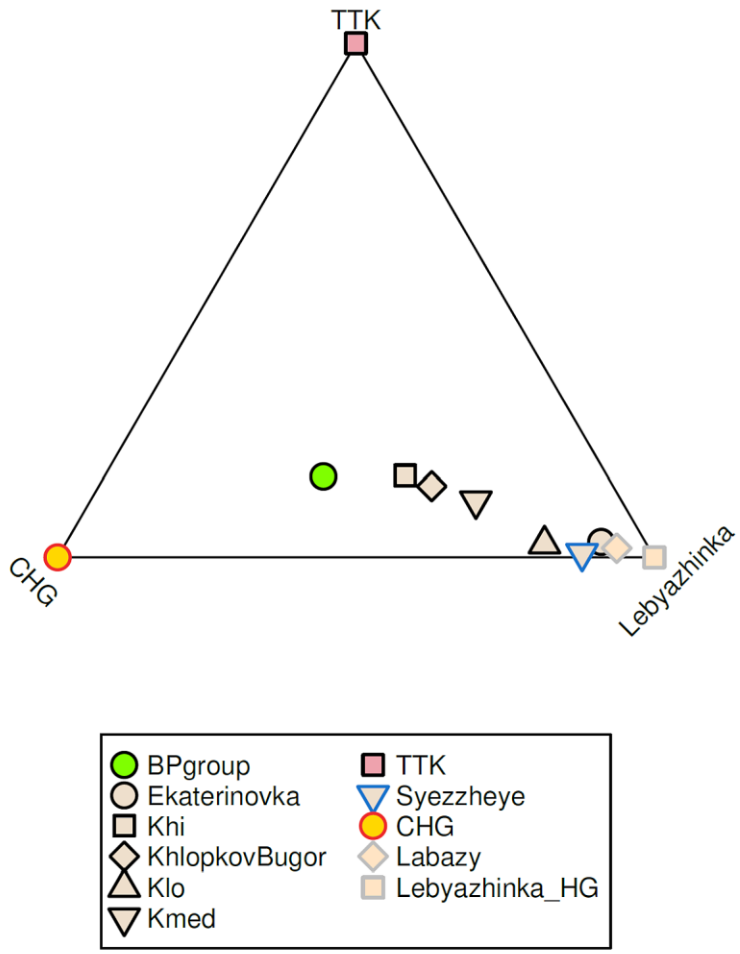

**Fig. S 2 A visualization of the 3-way model of the Volga cline**

We visualize the proportions of Table S 15 in Fig. S 2 which highlights their key feature of the presence of Central Asian-related ancestry in the Lower Volga (BPgroup), and low-medium EHG ancestry groups (Khi, Kmed, and Khlopkov Bugor), in contrast to the high EHG ancestry groups (Klo, Syezzheye, Ekaterinovka, and Labazy) all of which have a low and non-significant inferred proportion of TTK ancestry.

Thus, the Volga cline can indeed be seen as having been formed by an admixture of a population like BPgroup representing the Lower Volga and an EHG population like Lebyazhinka representing the older occupants of eastern Europe. However, this 2-way mix occurred among populations that were also impacted by migrants related to Central Asians and thus presumably coming from the east. The impact of TTK-related ancestry appears to have been variable along the Volga Cline, which provides a likely explanation for why the simple two-way models do not fit for groups like Khi. For example, the proportion of TTK ancestry is inferred to be approximately equal in Khi and in BPgroup (~16%) and is not diminished in Khi by ~1/4 as one might expect from the 2-way model for Khi as BPgroup+Lebyazhinka.

#### The Golubaya Krinitsa individuals from the Middle Don: Ukraine and Eastern hunter-gatherer and ancestry; migrations from the Volga and to the Dnipro

We discuss the modeling results for the Golubaya Krinitsa groups GK1 and GK2. Excluding models that involve Igren_o as a source (with which GK2 is cladal Table S 2), GK2 can be modeled itself as a mixture of people of the Volga (with high EHG ancestry) and hunter-gatherers from Ukraine. Its geographical position on the Don, and thus between the Dnipro-sampled "Ukraine_N" and Volga-sampled individuals, corresponds precisely to its genetic composition.

| A | B | P-value | A | B | S.E. | Win | Draw | Lose | Score |
|---|---|---|---|---|---|---|---|---|---|
| *1-way models* | | | | | | | | | |
| Igren_o | | 0.997 | | | | 0 | 21 | 0 | 0 |
| *2-way models* | | | | | | | | | |
| Murzikha | Ukraine_N | 0.655 | 44.8% | 55.2% | 6.0% | 3 | 18 | 0 | 3 |
| Russia_Karelia | Ukraine_N | 0.539 | 33.4% | 66.6% | 4.7% | 3 | 18 | 0 | 3 |
| Ekaterinovka | Ukraine_N | 0.969 | 36.6% | 63.4% | 4.8% | 2 | 19 | 0 | 2 |
| UpperVolga | Ukraine_N | 0.478 | 53.4% | 46.6% | 7.4% | 3 | 17 | 1 | 2 |
| Labazy | Ukraine_N | 0.990 | 36.3% | 63.7% | 4.9% | 1 | 20 | 0 | 1 |
| Lebyazhinka_HG | Ukraine_N | 0.844 | 38.0% | 62.0% | 5.7% | 1 | 20 | 0 | 1 |
| Klo | Ukraine_N | 0.349 | 35.4% | 64.6% | 5.0% | 2 | 15 | 4 | -2 |
| WSHG | Ukraine_N | 0.345 | 20.7% | 79.3% | 2.8% | 0 | 15 | 6 | -6 |
| TTK | Ukraine_N | 0.164 | 17.9% | 82.1% | 2.6% | 0 | 13 | 8 | -8 |

**Table S 16 1- and 2-way models for GK2.** Only a single 1-way model fits (with Igren_o as a clade). We also list all fitting 2-way models that do not include Igren_o, all of which have Ukraine_N as one source. The models that win in the model tournament have high EHG-related ancestry.

| A | B | P-value | A | B | S.E. | Win | Draw | Lose | Score |
|---|---|---|---|---|---|---|---|---|---|
| *1-way models* | | | | | | | | | |
| SSlo | | 0.078 | | | | 0 | 26 | 19 | -19 |
| *2-way models* | | | | | | | | | |
| GK2 | PVgroup | 0.915 | 62.3% | 37.7% | 4.1% | 20 | 25 | 0 | 20 |
| GK2 | SShi | 0.860 | 46.0% | 54.0% | 5.6% | 19 | 26 | 0 | 19 |
| **GK2** | **BPgroup** | **0.846** | **59.7%** | **40.3%** | **4.3%** | **18** | **27** | **0** | **18** |
| GK2 | CHG | 0.421 | 78.5% | 21.5% | 2.6% | 18 | 27 | 0 | 18 |
| GK2 | Krivyansky | 0.699 | 61.1% | 38.9% | 4.2% | 17 | 28 | 0 | 17 |
| GK2 | SSmed | 0.090 | 27.7% | 72.3% | 8.2% | 16 | 29 | 0 | 16 |
| Ekaterinovka | SSlo | 0.366 | 24.9% | 75.1% | 9.6% | 7 | 38 | 0 | 7 |
| GK2 | Iran_GanjDareh_N | 0.171 | 79.7% | 20.3% | 2.5% | 15 | 22 | 8 | 7 |
| Kmed | SSlo | 1.000 | 38.6% | 61.4% | 8.7% | 7 | 38 | 0 | 7 |
| Labazy | SSlo | 0.308 | 22.9% | 77.1% | 9.3% | 6 | 38 | 1 | 5 |
| SSlo | WSHG | 0.615 | 83.2% | 16.8% | 5.1% | 5 | 40 | 0 | 5 |
| Russia_Steppe_Maikop | SSlo | 0.861 | 26.1% | 73.9% | 7.1% | 4 | 41 | 0 | 4 |
| Khi | Ukraine_N | 0.818 | 56.8% | 43.2% | 3.9% | 10 | 29 | 6 | 4 |
| Russia_Karelia | SSlo | 0.142 | 14.2% | 85.8% | 7.7% | 7 | 35 | 3 | 4 |
| KhlopkovBugor | SSlo | 0.931 | 33.7% | 66.3% | 8.9% | 3 | 42 | 0 | 3 |
| BPgroup | Igren_o | 0.828 | 39.5% | 60.5% | 5.8% | 3 | 42 | 0 | 3 |
| Igren_o | SShi | 0.886 | 46.9% | 53.1% | 6.9% | 3 | 42 | 0 | 3 |
| Igren_o | PVgroup | 0.939 | 62.8% | 37.2% | 5.6% | 2 | 43 | 0 | 2 |
| Khi | SSlo | 0.871 | 32.9% | 67.1% | 8.7% | 2 | 43 | 0 | 2 |
| KhlopkovBugor | Ukraine_N | 0.473 | 57.1% | 42.9% | 4.0% | 10 | 27 | 8 | 2 |
| SSlo | TTK | 0.895 | 82.9% | 17.1% | 5.0% | 1 | 44 | 0 | 1 |
| BPgroup | Ukraine_N | 0.221 | 49.5% | 50.5% | 3.2% | 7 | 31 | 7 | 0 |
| Ekaterinovka | SSmed | 0.270 | 24.9% | 75.1% | 5.7% | 7 | 31 | 7 | 0 |

| | | | | | | | | | |
|---|---|---|---|---|---|---|---|---|---|
| Igren_o | Krivyansky | 0.937 | 61.1% | 38.9% | 5.7% | 0 | 45 | 0 | 0 |
| Igren_o | Remontnoye | 0.385 | 71.0% | 29.0% | 4.8% | 0 | 45 | 0 | 0 |
| CHG | Igren_o | 0.861 | 22.2% | 77.8% | 3.9% | 0 | 44 | 1 | -1 |
| Murzikha | SSmed | 0.518 | 24.1% | 75.9% | 5.3% | 6 | 32 | 7 | -1 |
| Igren_o | Iran_GanjDareh_N | 0.779 | 79.4% | 20.6% | 3.9% | 0 | 44 | 1 | -1 |
| Russia_Karelia | SSmed | 0.302 | 18.7% | 81.3% | 4.5% | 6 | 32 | 7 | -1 |
| Labazy | SSmed | 0.227 | 23.1% | 76.9% | 5.6% | 5 | 33 | 7 | -2 |
| Lebyazhinka_HG | SSlo | 0.148 | 17.0% | 83.0% | 9.5% | 1 | 41 | 3 | -2 |
| Klo | SSmed | 0.187 | 31.4% | 68.6% | 7.4% | 6 | 30 | 9 | -3 |
| Krivyansky | UpperVolga | 0.504 | 45.3% | 54.7% | 3.5% | 4 | 33 | 8 | -4 |
| Igren_o | SSmed | 0.218 | 31.5% | 68.5% | 10.0% | 1 | 39 | 5 | -4 |
| GK2 | Khi | 0.056 | 53.5% | 46.5% | 5.2% | 6 | 28 | 11 | -5 |
| SSmed | UpperVolga | 0.362 | 75.5% | 24.5% | 5.5% | 2 | 36 | 7 | -5 |
| SShi | UpperVolga | 0.145 | 60.9% | 39.1% | 4.7% | 3 | 34 | 8 | -5 |
| Igren_o | Khi | 0.118 | 56.7% | 43.3% | 7.0% | 2 | 34 | 9 | -7 |
| Lebyazhinka_HG | SSmed | 0.347 | 21.8% | 78.2% | 5.2% | 1 | 36 | 8 | -7 |
| Igren_o | Unakozovskaya | 0.136 | 79.6% | 20.4% | 4.0% | 1 | 33 | 11 | -10 |
| BPgroup | SSlo | 0.279 | 20.6% | 79.4% | 8.4% | 1 | 33 | 11 | -10 |
| Krivyansky | Murzikha | 0.084 | 46.9% | 53.1% | 3.6% | 4 | 26 | 15 | -11 |
| PVgroup | SSlo | 0.169 | 16.4% | 83.6% | 8.1% | 1 | 25 | 19 | -18 |
| CHG | UpperVolga | 0.068 | 27.4% | 72.6% | 2.2% | 1 | 19 | 25 | -24 |
| Remontnoye | SSlo | 0.054 | 4.6% | 95.4% | 7.0% | 0 | 18 | 27 | -27 |

**Table S 17 Considered 1- and 2-way models for GK1**

Thel GK1 population is visibly more CHG-shifted in PCA (Fig. S 1) and many 2-way models fit it (Table S 17). None of the models involve only populations of high steppe ancestry. A particularly good model with a high score, no losses, and a high p-value involves 40.3±4.3% BPgroup ancestry and 59.7±4.3% GK2 ancestry, and all the models that do well in the tournament involve GK2. This suggests that populations from the Volga and/or the Dnipro clines were admixing with populations of the Don (like GK2).

Individuals from Golubaya Krinitsa of a similar age as GK1/GK2 were reported by Allentoft et al.[32] where it is observed that they already had CHG-related ancestry. These individuals thus appear to be similar to our GK1 individuals rather than the GK2 one that appears to have simpler ancestry (Ukraine_N/EHG hunter-gatherer ancestry). They further suggested in that study that Yamnaya steppe pastoralists can be modeled as ~2/3 Golubaya Krinitsa Middle Don people and ~1/3 Caucasus hunter-gatherers.

We obtain a similar estimate when we model the core Yamnaya using the CHG+GK1 model as in ref.[32] of 31.5±2.2% CHG ancestry and 68.5±2.2% GK1 ancestry. However, this model fails (p=1.3e-7) as it underestimates shared genetic drift with both Afontova Gora 3 (Z=-2.6) and with Turkey_N (Z=-3.0). The model proposed in our note (Remontnoye + SShi) fits well because it can model both the Anatolian Neolithic affinity of the Core Yamnaya (via the Aknashen/Maikop component of Remontnoye) and their Siberian affinity (via the BPgroup component of Remontnoye which itself has a Siberian/Central Asian affinity).

After the publication of the data of ref.[32] we modeled each Golubaya Krinitsa individual from that study using the very same sources as in the current study (Table S 18); all of them can be modeled as cladal with GK1. Overall, this indicates that the ref.[32] individuals represent a similar population to the GK1 subset.

| Individual | Source | P-value |
|---|---|---|
| NEO113.SG | GK1 | 0.710 |
| NEO204.SG | GK1 | 0.624 |
| NEO207.SG | GK1 | 0.907 |
| NEO209.SG | GK1 | 0.759 |
| NEO210.SG | GK1 | 0.757 |
| NEO212.SG | GK1 | 0.176 |
| NEO207.SG | Igren_o | 0.170 |
| NEO207.SG | SSlo | 0.393 |
| NEO209.SG | SSlo | 0.220 |
| NEO210.SG | SSlo | 0.075 |
| NEO204.SG | SSmed | 0.061 |
| NEO209.SG | SSmed | 0.123 |

**Table S 18 Golubaya Krinitsa individuals from ref.[32] can be modeled as a clade with our GK1 population from the same site.**

Furthermore, we model these individuals using the same ~ 60:40 GK2:BPgroup model of Table S 17 and show the results in Table S 19 which confirm the similarity with the GK1 subset.

| Individual | P-value | BPgroup | GK2 | S.E. |
|---|---|---|---|---|
| NEO113.SG | 0.536 | 40.7% | 59.3% | 5.9% |
| NEO204.SG | 0.401 | 50.6% | 49.4% | 6.5% |
| NEO207.SG | 0.931 | 30.6% | 69.4% | 7.5% |
| NEO209.SG | 0.686 | 47.0% | 53.0% | 7.0% |
| NEO210.SG | 0.942 | 44.4% | 55.6% | 7.5% |
| NEO212.SG | 0.542 | 55.2% | 44.8% | 4.5% |

**Table S 19 Golubaya Krinitsa individuals from ref.[32] using the BPgroup+GK2 mode**

The Golubaya Krinitsa individuals present an important data point for the early presence of populations of mixed Caucasus and steppe origins in the Middle Don, well to the north of the Caucasus; but they cannot be seen as having formed the Yamnaya in the kind of admixture proposed in ref.[32] Rather, the Yamnaya have ancestry related to populations of the North Caucasus-Lower Volga (BPgroup) and to the Caucasus Neolithic and Bronze Age (Aknashen/Maikop) and were formed by the admixture of such populations via an intermediate stage like Remontnoye with people of the Don-Dnipro area.

#### Modeling the Volga Cline and Golubaya Krinitsa

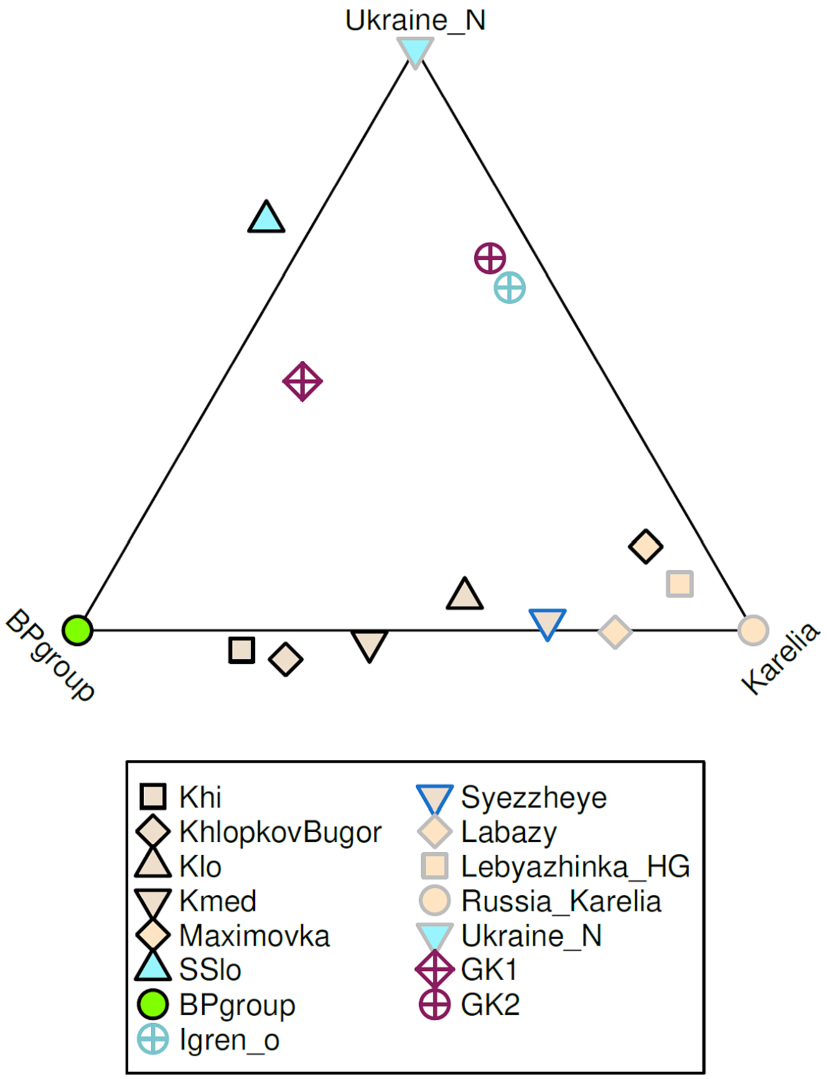

**Fig. S 3 Modeling the Don-Volga region jointly.** All populations that fit the specified model are plotted.

The model of Fig. S 2 includes the Volga Cline as well as the "eastern" Central Asian influence (TTK) that is needed for some of its populations. We have also seen that the Golubaya Krinitsa individuals have ancestry from the Volga cline and we show in Fig. S 3 a model which includes populations of the Volga and "western" populations like GK1/GK2 that fit the model. We plot all populations that fit the model in Fig. S 3. Notable in their absence are Krivyansky and other Serednii Stih populations other than the SSlo subset (which has the most Ukraine_N-related ancestry along the Dnipro cline, meaning that there is little power to reject models for the more CHG-related part of their ancestry). The absent Serednii Stih populations do have Ukraine_N-related ancestry as is apparent from the PCA, but as we will next see, this ancestry is combined with additional sources not present in the Volga cline and the mid-6$^{th}$ millennium BCE Don at Golubaya Krinitsa, explaining why they do not fit in Fig. S 3.

#### Modeling the Serednii Stih-Yamnaya (Dnipro) cline

Our exploration of Yamnaya origins has revealed that they can be well-modeled formed as a mixture of Remontnoye + SShi ancestry (Table S 3). The Remontnoye in turn were consistent with being formed by admixture between groups from the Caucasus (Aknashen or Maikop) plus BPgroup (Table S 12). The BP-group was one of the sources of the Volga cline with the other being an EHG-related group (Table S 14).

We turn now to the second component of the Yamnaya, the SShi subset of the Serednii Stih culture. This consists in this analysis of the following four individuals: I1430, I1924, I2108, and I6559 from Igren, Olexandria, and Vinogradnoe.

We assess the following potential members of the proposed Serednii Stih-Yamnaya cline: CoreYamnaya, GK1, GK2, Russia_Don_EBA_Yamnaya, SShi, SSlo, SSmed, Ukraine_N. Some of these are visibly "off-cline" in PCA, such as some Serednii Stih individuals, or the termini of the cline at either Ukraine_N (composed of many individuals) or GK2 (a single individual). Taking pairs of these populations as sources, we list the feasible models for them (Table S 20).

| Test | A | B | P-value | A | B | S.E. |
|---|---|---|---|---|---|---|
| SShi | CoreYamnaya | GK1 | 0.419 | 69.3% | 30.7% | 5.8% |
| SSmed | CoreYamnaya | GK1 | 0.054 | 29.4% | 70.6% | 7.0% |
| GK1 | CoreYamnaya | GK2 | 0.382 | 44.9% | 55.1% | 4.8% |
| SShi | CoreYamnaya | GK2 | 0.079 | 83.9% | 16.1% | 2.9% |
| SSlo | CoreYamnaya | GK2 | 0.075 | 33.0% | 67.0% | 7.5% |
| SSmed | CoreYamnaya | GK2 | 0.435 | 60.1% | 39.9% | 3.0% |
| Russia_Don_EBA_Yamnaya | CoreYamnaya | SSlo | 0.361 | 65.3% | 34.7% | 3.7% |
| SSmed | CoreYamnaya | SSlo | 0.599 | 42.0% | 58.0% | 6.6% |
| Russia_Don_EBA_Yamnaya | CoreYamnaya | SSmed | 0.237 | 40.0% | 60.0% | 4.7% |
| Russia_Don_EBA_Yamnaya | CoreYamnaya | Ukraine_N | 0.079 | 79.4% | 20.6% | 1.1% |
| SSlo | CoreYamnaya | Ukraine_N | 0.850 | 39.1% | 60.9% | 5.7% |
| SSmed | CoreYamnaya | Ukraine_N | 0.272 | 65.2% | 34.8% | 2.2% |
| SSmed | GK1 | Russia_Don_EBA_Yamnaya | 0.410 | 45.9% | 54.1% | 9.1% |
| GK1 | GK2 | Russia_Don_EBA_Yamnaya | 0.093 | 44.1% | 55.9% | 6.0% |
| SSlo | GK2 | Russia_Don_EBA_Yamnaya | 0.135 | 56.0% | 44.0% | 9.5% |
| SSmed | GK2 | Russia_Don_EBA_Yamnaya | 0.868 | 22.1% | 77.9% | 3.6% |
| GK1 | GK2 | SShi | 0.860 | 46.1% | 53.9% | 5.5% |
| SSlo | GK2 | SShi | 0.062 | 61.7% | 38.3% | 8.8% |
| SSmed | GK2 | SShi | 0.100 | 29.4% | 70.6% | 4.5% |
| GK1 | GK2 | SSmed | 0.090 | 27.8% | 72.2% | 7.9% |
| SSmed | Russia_Don_EBA_Yamnaya | SSlo | 0.552 | 62.7% | 37.3% | 8.7% |
| SSlo | Russia_Don_EBA_Yamnaya | Ukraine_N | 0.782 | 48.8% | 51.2% | 7.5% |
| SSmed | Russia_Don_EBA_Yamnaya | Ukraine_N | 0.289 | 82.1% | 17.9% | 3.0% |
| SSmed | SShi | SSlo | 0.501 | 51.8% | 48.2% | 8.5% |
| SSlo | SShi | Ukraine_N | 0.931 | 44.6% | 55.4% | 6.5% |
| SSmed | SShi | Ukraine_N | 0.530 | 74.1% | 25.9% | 3.4% |
| SSlo | SSmed | Ukraine_N | 0.805 | 59.4% | 40.6% | 9.3% |

**Table S 20 Modeling the Serednii Stih-Yamnaya cline.**

These results point to the Serednii Stih-Yamnaya cline being formed by the admixture of high hunter-gatherer ancestry populations from the Dnipro and Don (such as Ukraine_N, GK2, and SSlo) with those of the opposing end of the cline (SShi, Don/Core Yamnaya). Notably, the Core Yamnaya cannot be formed in terms of mixtures of members of the cline itself, as one might expect given its position as the terminus of the "low hunter-gatherer" end of the Cline. As we have seen, however, the Core Yamnaya can be modeled when populations beyond the Cline are included as sources, such as Remontnoye and SShi (Table

S 3). By contrast, the Don Yamnaya are modeled as a mixture of Core Yamnaya and high hunter-gatherer ancestry groups (SSlo, SSmed, and Ukraine_N) and as we will see below, they has evidence for having been formed by such a mixture a few centuries before the sampling of the Don Yamnaya individuals.

The hunter-gatherer end of the cline could be modeled alternatively as either Ukraine_N or GK2, and as we have seen (Table S 16) GK2 is itself intermediate between the Ukraine_N and EHG groups (matching its intermediate geography on the Don between the Dnipro and the Volga). We thus allowed for a 3rd source (adding Lebyazhinka_HG to the set of possible sources) (Table S 21).

| | | Proportions | | | Std. errors | | | |
|---|---|---|---|---|---|---|---|---|
| Test | P-value | A=Lebyazhinka_HG | B=Ukraine_N | C=CoreYamnaya | A | B | C | A/(A+B) |
| GK1 | 0.334 | 24.8% | 27.8% | 47.4% | 5.1% | 5.1% | 4.2% | 47.1% |
| GK2 | 0.968 | 34.9% | 59.4% | 5.7% | 6.0% | 5.9% | 4.4% | 37.0% |
| Russia_Don_EBA_Yamnaya | *0.050* | 0.3% | 20.5% | 79.2% | 1.5% | 1.5% | 1.2% | 1.4% |
| SShi | 0.082 | 10.2% | 5.4% | 84.4% | 3.4% | 3.3% | 2.8% | 65.4% |
| SSlo | 0.735 | 0.5% | 60.5% | 39.0% | 8.3% | 8.3% | 6.2% | 0.8% |
| SSmed | 0.751 | 6.9% | 29.8% | 63.3% | 3.2% | 3.3% | 2.5% | 18.8% |

**Table S 21 A 3-way model for the Serednii Stih-Yamnaya cline**. The unrounded p-value for Don Yamnaya is 0.0496.

It seems that relative to Core Yamnaya (the low-hunter gatherer end of the cline), SSlo, SSmed, and Don Yamnaya (the bulk of the cline) draw their hunter-gatherer ancestry primarily from Ukraine_N; the Golubaya Krinitsa individuals previously discussed from both EHG and Ukraine_N sources. The proportions for the SShi are low (as this population has the least hunter-gatherer ancestry of the Serednii Stih subsets), but seem to slightly favor the EHG source.

#### The origins of Krivyansky on the Lower Don: a Seredniih Stih population with excess Caucasus-related ancestry

The Krivyansky individual does not fit the Remontnoye+SShi model of Table S 5 well (p=0.015) and it also does not fit the model of Table S 21 well (p=0.025). This individual is also unusual as it belonged to Y-haplogroup J2 suggesting Caucasus/West Asian connections.

The results of the model tournament (Table S 22) show that relative to populations of the Dnipro-Don Serednii Stih and Golubaya Krinitsa, Krivyansky has an excess of CHG-related ancestry. The presence of this ancestry in the lower Don at a higher level than in the middle Don (at the GK1 subset of Golubaya Krinitsa parallels the situation of the Volga where CHG-related ancestry was higher in the lower Volga (at Berezhnovka) than in the middle Volga (Fig. S 2) and of the Caucasus where CHG-related ancestry in Maikop and (in slight excess Unakozovskaya) was higher in the North Caucasus than in the South Caucasus where the Aknashen earliest known Neolithic had seen its CHG-related ancestry diluted twice in Masis Blur and Areni-1 Neolithic and Chalcolithic populations. While CHG-related ancestry refers to the pre-Neolithic inhabitants of Georgia[33] in the South Caucasus, it seems that it is in the triangle formed by the lower Don (at Krivyansky), the lower Volga (at Berezhnovka), and north Caucasus (at Unakozovskaya) that it remained a strong source of ancestry in the Eneolithic while CHG-related ancestry diminished upriver along both Don and Volga to the north, and across the Caucasus to the south.

| A | B | P-value | A | B | S.E. | Win | Draw | Lose | Score |
|---|---|---|---|---|---|---|---|---|---|
| CHG | SSmed | 0.578 | 38.0% | 62.0% | 3.5% | 4 | 4 | 0 | 4 |

| | | | | | | | | | |
|---|---|---|---|---|---|---|---|---|---|
| CHG | SShi | 0.504 | 27.5% | 72.5% | 4.3% | 2 | 6 | 0 | 2 |
| Iran_GanjDareh_N | SShi | 0.132 | 24.9% | 75.1% | 4.1% | 2 | 6 | 0 | 2 |
| CHG | GK1 | 0.645 | 44.9% | 55.1% | 3.3% | 0 | 8 | 0 | 0 |
| CHG | Igren_o | 0.643 | 57.0% | 43.0% | 3.0% | 0 | 8 | 0 | 0 |
| CHG | GK2 | 0.369 | 56.7% | 43.3% | 2.6% | 0 | 7 | 1 | -1 |
| CHG | SSlo | 0.087 | 47.2% | 52.8% | 3.8% | 0 | 7 | 1 | -1 |
| CHG | Murzikha | 0.061 | 60.2% | 39.8% | 2.3% | 0 | 5 | 3 | -3 |
| CHG | UpperVolga | 0.110 | 59.9% | 40.1% | 2.3% | 0 | 5 | 3 | -3 |

**Table S 22 Considered 2-way models for Krivyansky.**

We also explored 3-way models for Krivyansky after fixing CHG as one source (to account for this population's excess CHG ancestry) and Ukraine_N as another (as a stand-in for the source that generates the Serednii Stih cline). All fitting models (Table S 23) involve some "eastern" ancestry (in addition to the CHG and Ukraine_N fixed sources; Table S 23). Three models have no losses in the tournament (Table S 23) and these involve Lower Volga-North Caucasus Eneolithic ancestry (BPgroup or PVgroup) or the SShi subset of Serednii Stih. Thus, Krivyansky appears like a population of "western" affinity (due to its Ukraine_N-related ancestry) but also "eastern" affinity (due to its Lower Volga-North Caucasus Eneolithic-related ancestry) and also "southern" affinity (due to its CHG-related ancestry). As we will see further on, the people of the Serednii Stih and the Yamnaya share two of these ancestries but their southern source is related to Neolithic and post-Neolithic people of the Caucasus, not the much earlier CHG as is the case for Krivyansky.

| | | Proportions | | | Std. errors | | | | | | |
|---|---|---|---|---|---|---|---|---|---|---|---|
| B | P-value | A=CHG | B | C=Ukraine_N | A | B | C | Win | Draw | Lose | Score |
| **BPgroup** | **0.387** | **38.8%** | **37.2%** | **8.1%** | **24.0%** | **5.2%** | **4.2%** | **11** | **4** | **0** | **11** |
| **PVgroup** | **0.403** | **32.5%** | **43.9%** | **23.6%** | **6.7%** | **9.8%** | **4.3%** | **13** | **2** | **0** | **13** |
| Khi | 0.311 | 46.6% | 29.7% | 23.7% | 3.8% | 6.6% | 4.2% | 9 | 5 | 1 | 8 |
| **SShi** | **0.518** | **32.6%** | **59.6%** | **7.8%** | **6.5%** | **13.4%** | **7.7%** | **6** | **9** | **0** | **6** |
| Kmed | 0.434 | 50.3% | 27.6% | 22.1% | 3.2% | 6.1% | 4.6% | 7 | 6 | 2 | 5 |
| Russia_Steppe_Maikop | 0.417 | 50.8% | 21.1% | 28.1% | 3.1% | 4.6% | 3.4% | 5 | 5 | 5 | 0 |
| KhlopkovBugor | 0.281 | 48.1% | 28.5% | 23.5% | 3.5% | 6.4% | 4.5% | 1 | 13 | 1 | 0 |
| Klo | 0.304 | 53.3% | 26.8% | 19.9% | 2.7% | 5.7% | 4.8% | 1 | 12 | 2 | -1 |
| Ekaterinovka | 0.261 | 56.0% | 23.2% | 20.8% | 2.5% | 5.2% | 4.9% | 1 | 10 | 4 | -3 |
| Labazy | 0.311 | 56.5% | 22.7% | 20.8% | 2.5% | 5.1% | 4.8% | 0 | 12 | 3 | -3 |
| WSHG | 0.494 | 56.2% | 14.0% | 29.7% | 3.2% | 2.5% | 3.0% | 1 | 9 | 5 | -4 |
| Lebyazhinka_HG | 0.440 | 58.2% | 23.4% | 18.4% | 2.4% | 5.1% | 5.2% | 0 | 10 | 5 | -5 |
| Murzikha | 0.257 | 58.7% | 27.2% | 14.1% | 2.3% | 6.1% | 6.2% | 0 | 9 | 6 | -6 |
| Russia_Karelia | 0.298 | 58.9% | 20.5% | 20.6% | 2.4% | 4.5% | 4.8% | 0 | 9 | 6 | -6 |
| UpperVolga | 0.121 | 59.0% | 31.0% | 10.0% | 7.7% | 2.4% | 7.7% | 0 | 8 | 7 | -7 |
| TTK | 0.268 | 54.3% | 13.0% | 32.8% | 2.7% | 2.9% | 2.9% | 0 | 7 | 8 | -8 |

**Table S 23 Considered 3-way models for Krivyansky.**

We show in Fig. S 4 the populations that fit the BPgroup-Ukraine_N-CHG model, showing how Krivyansky fits, but most other Serednii-Stih and Yamnaya groups do not fit.

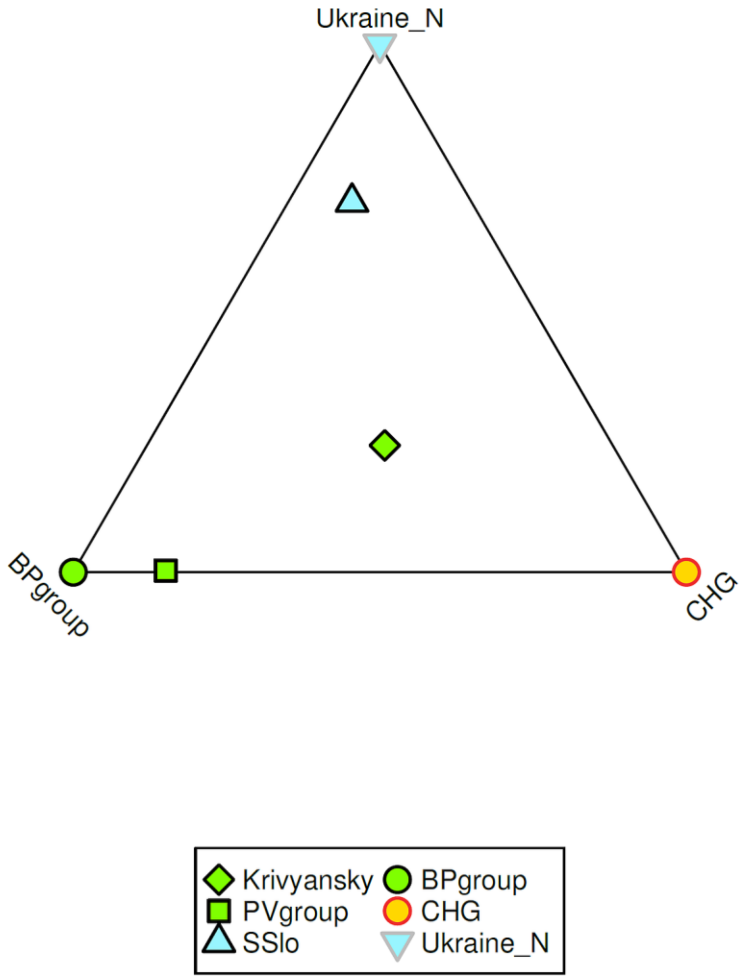

**Fig. S 4 Populations that can fit the model for Krivyansky.**

#### Variable Central Asian ancestry in the Eneolithic steppe and the origins of the Steppe Maikop

The Krivyansky population which we just discussed can be modeled without any TTK-related ancestry. As such, it contrasts with the Eneolithic populations of the North Caucasus piedmont-Lower Volga (PVgroup and BPgroup) that have evidence of such ancestry (Table S 12and Table S 13). People at Khvalynsk and Khlopkov Bugor also had such extra ancestry, beyond that expected based on their inferred proportion of mixture from people of the North Caucasus piedmont-Lower Volga, but people at Ekaterinovka and Syezzheye did not have any evidence of excess (Table S 15, Fig. S 2). Thus, the Central Asian influence appears to be variable: found on the Lower Volga and upriver to Khvalynsk, but not on the eastern edges of the Serednii Stih culture as represented by Krivyansky or indeed the other Serednii Stih populations that can also be modeled without any extra such ancestry (Table S 20).

To this evidence of variability, we add the modeling results for the Steppe Maikop, a population whose eastern “Siberian” affinities were noticed when it was first published.[8] We can model it with proximate sources here (

| | | | | | | | | | |
|---|---|---|---|---|---|---|---|---|---|
| Krivyansky | WSHG | 0.068 | 42.2% | 57.8% | 2.6% | 0 | 1 | 2 | -2 |

Table S 24) as a mixture of approximately half Serednii Stih and half TTK/WSHG ancestry.

| A | B | P-value | A | B | S.E. | Win | Draw | Lose | Score |
|---|---|---|---|---|---|---|---|---|---|
| BPgroup | WSHG | 0.236 | 53.0% | 47.0% | 2.7% | 1 | 2 | 0 | 1 |
| PVgroup | WSHG | 0.237 | 47.7% | 52.3% | 2.6% | 1 | 2 | 0 | 1 |
| SShi | TTK | 0.179 | 46.5% | 53.5% | 3.5% | 0 | 3 | 0 | 0 |
| Krivyansky | WSHG | 0.068 | 42.2% | 57.8% | 2.6% | 0 | 1 | 2 | -2 |

**Table S 24 Consider models for Russia_Steppe_Maikop.**

The sources of the TTK/WSHG-related ancestry are unclear, and more sampling will be required in western Siberia and central Asia to identify proximate sources for it. But, the results of this analysis suggest its variable, and probably late, impact in the Don-Volga-North Caucasus region. Some of this ancestry entered the Yamnaya via BPgroup ancestors in our reconstruction of Yamnaya origins. However, the presence of it in the Steppe Maikop and in the Volga cline (above and beyond what was contributed by BPgroup) suggest a more complex history of contact between the Eneolithic people of eastern Europe and their Central Asian and Siberian neighbors.

#### When was the Yamnaya formed and when did it expand?

The modeling of the Serednii-Stih with the Yamnaya as the proxy for its “eastern” source (Table S 20) suggests that the population ancestral to the Yamnaya may have been formed during the 5$^{th}$ millennium BCE. This “pre-Yamnaya” population can be seen as a population of similar autosomal genetic ancestry. The archaeologically defined Yamnaya of the late 4$^{th}$ millennium BCE would then be descendants of the pre-Yamnaya of a thousand years earlier, defined not only by their autosomal genetic composition, but also (archaeologically) from their distinctive Pit Grave culture, and (genetically) their possession of a specific Y-chromosome lineage (R-Z2103/R-M12149).

The admixture in the ancestry of the Yamnaya has been dated using DATES[12] to ~4100BCE[12] and ~4555BCE[4]. Using the core Yamnaya set (*n*=104) of which a large number (*n*=61) have radiocarbon dates with a mean of 2877BCE, we repeat this computation and estimate that West Asian (*n*=125)-European hunter-gatherer (*n*=215) admixture took place 41.5±1.7 generations before that time. Assuming a generation time of 28 years[34] yields an estimate of 4038±48 years BCE. Thus, admixture LD points to an admixture date between the 5$^{th}$ and 4$^{th}$ millennium BCE. As this admixture may not have been instantaneous it may have stretched from the 5$^{th}$ millennium BCE to the dawn of the Bronze Age in the 4$^{th}$ millennium BCE.

Our reconstruction reveals population admixture at different time scales in the ancestry of the Yamnaya. What is significant about the obtained admixture LD date is it shows that there was major admixture of “West Asian” and “steppe” related populations in the ancestry of the Yamnaya at least as recent into time of the Serednii Stih culture. Thus, the hypothesis that the Yamnaya represent a stable population at the terminus of the Dnipro Cline and that they themselves did not experience major admixture in the 5$^{th}$ millennium, is not feasible.

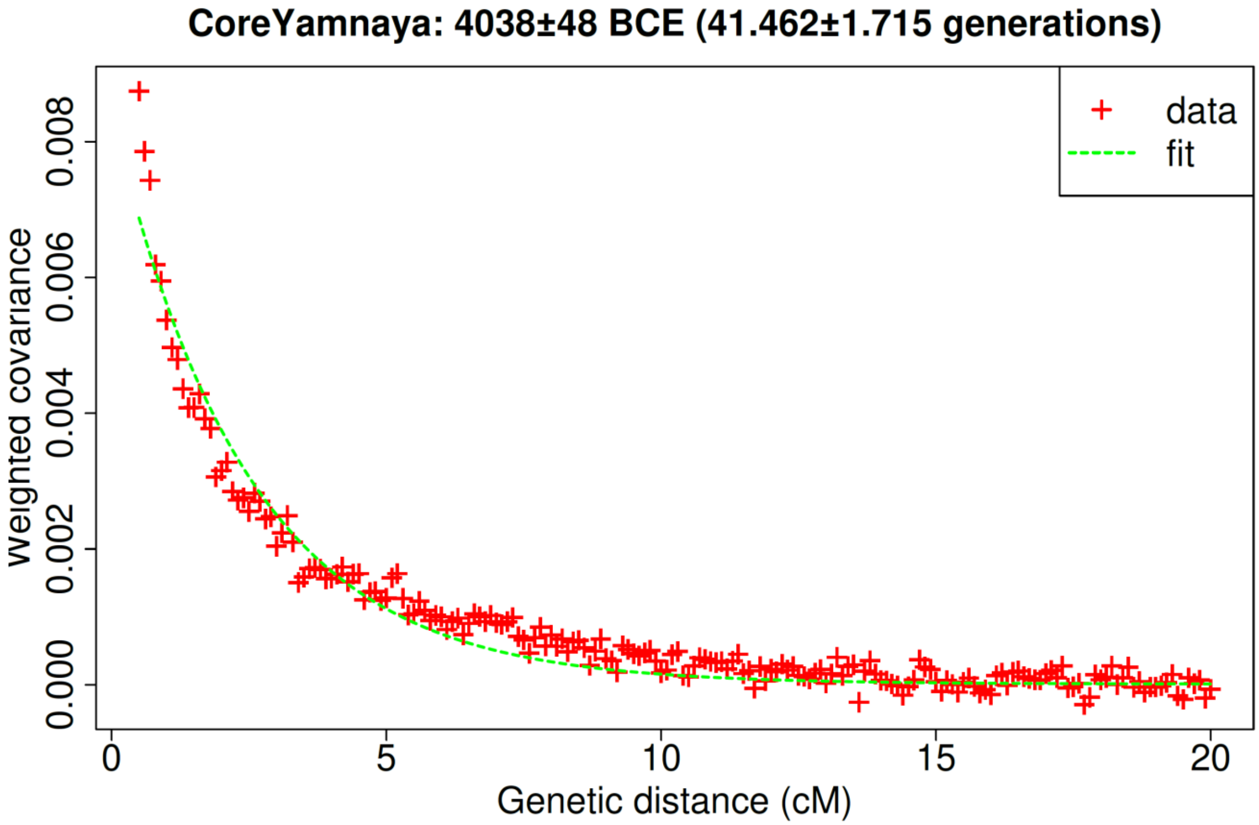

**Fig. S 5 Admixture LD estimation of Core Yamnaya formation**

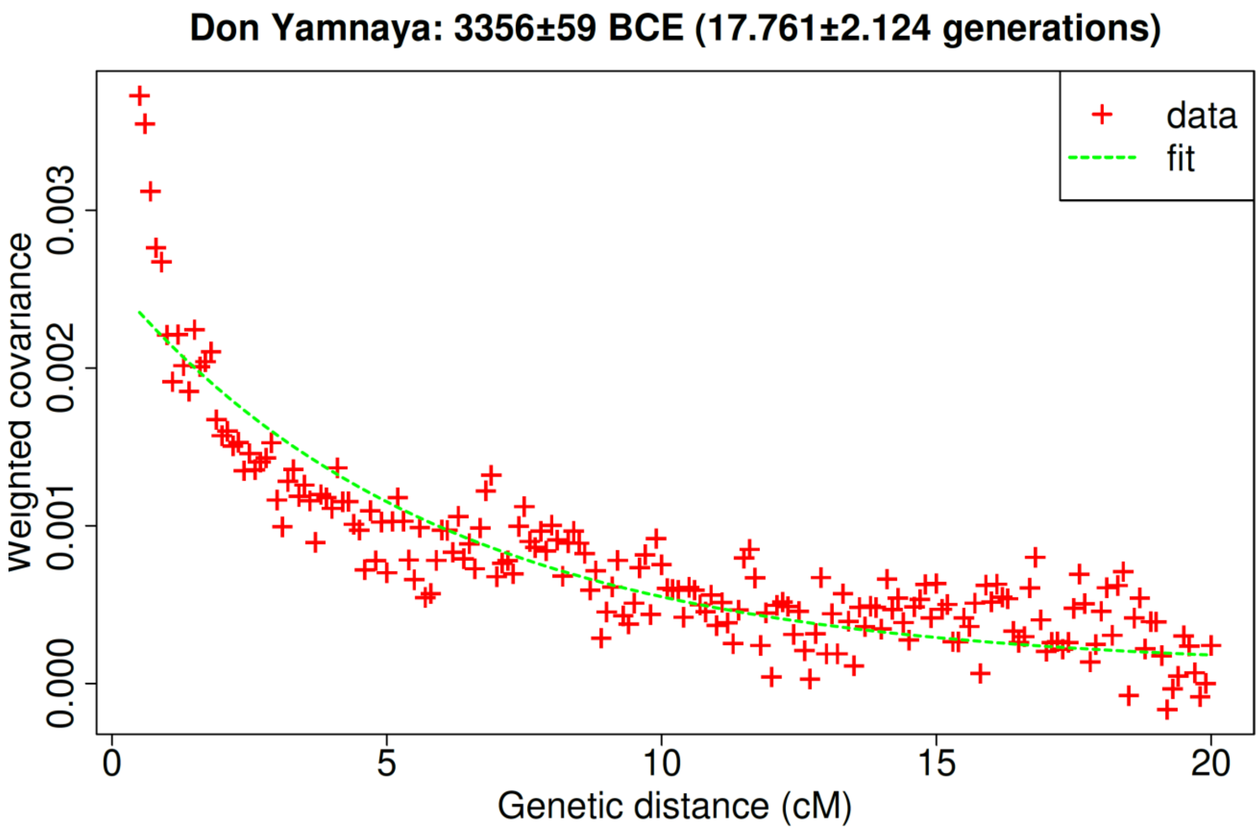

**Fig. S 6 Admixture LD estimation of Don Yamnaya formation**

We can also date the genetic formation of the Don Yamnaya (*n*=21) by using Core Yamnaya as one source and Ukraine_N as the other (Fig. S 6). This is 17.8±2.1 generations. As the Don Yamnaya have an average calibrated date of 2858BCE, using a generation of 28 years yields an admixture time of 3356BCE, slightly preceding or coinciding with the emergence of the Yamnaya archaeological culture itself, and around 700 years after the mean date of formation of the core Yamnaya.

We can also date the admixture in Yamnaya that went westwards into Southeastern Europe. Grouping Yamnaya individuals of mean date 2796BCE from Bulgaria-Moldova-Romania-Serbia and Neolithic/Chalcolithic farmers from Fig. 4b we obtain an estimate of 19.7±3.1 generations which corresponds to 3349BCE (Fig. S 7), a date that is similar to that of the Don Yamnaya at the beginning of the Yamnaya expansion. These two dates suggest that soon after the core Yamnaya appeared—and we will argue that this happened in the Dnipro-Don region in the next section—it admixed with its geographical neighbors: with at least one population of high Ukraine_N ancestry to form the Don Yamnaya, and with at least one population of high European Neolithic ancestry to form the BMRS Yamnaya.

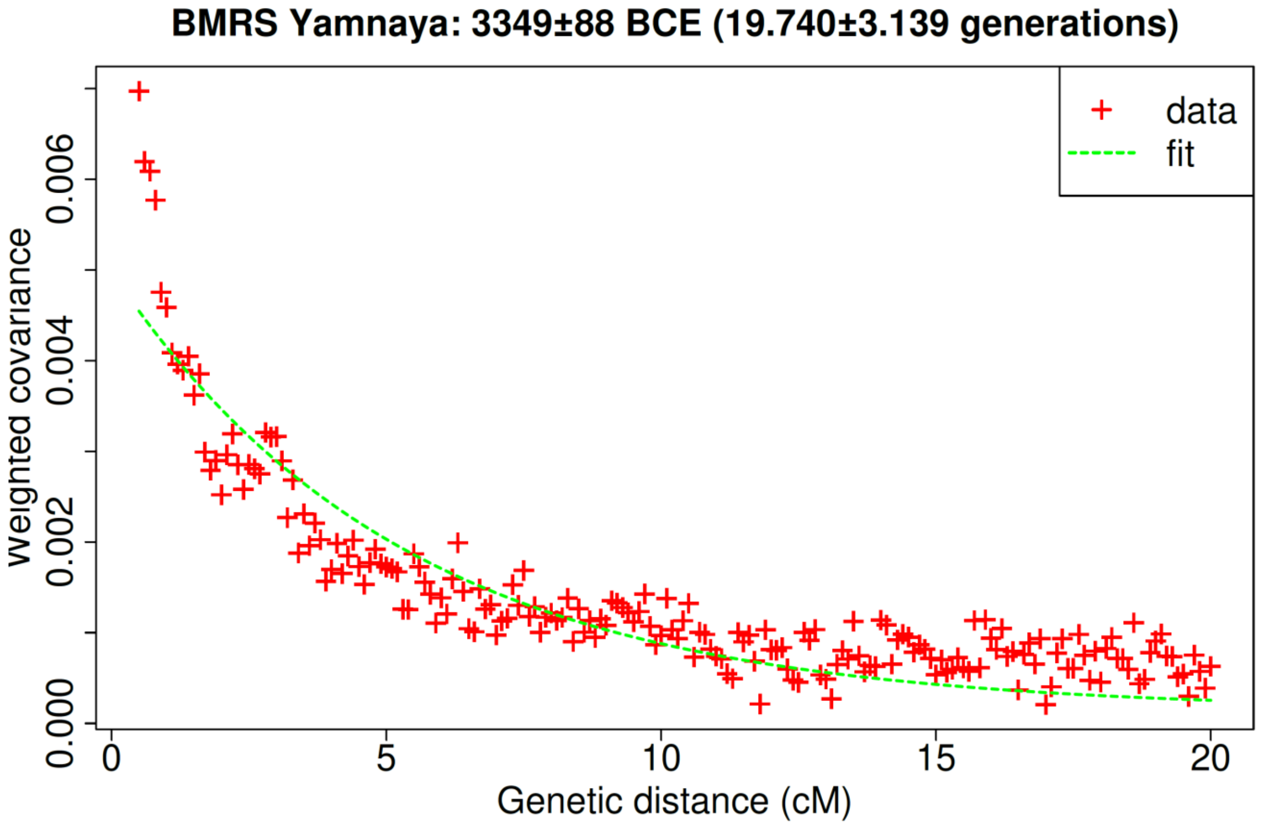

**Fig. S 7 Admixture LD estimation of BMRS Yamnaya formation**

Finally, we also dated the admixture in the Corded Ware complex, a population estimated to have ~3/4 of Yamnaya-related ancestry.[1] It has recently been discovered that this population shares a high rate of IBD segments with the people of the Globular Amphora farming culture[35] who may thus have been responsible for the remaining ~1/4 of (non-steppe) ancestry. It has been estimated that admixture in diverse Corded Ware populations occurred in a narrow date of ~3000-2900BCE.[12] We combine 86 Corded Ware individuals from the literature[2,6,11,22,30,36-43] and date them using the Core Yamnaya as one source and 27 Globular Amphora individuals[11,30,42,44,45] as the other. The obtained estimate is 12.8±0.6 generations corresponding to a date of 2933±17 BCE (Fig. S 8). This date is younger than those of either the Don or BMRS Yamnaya and the admixture in the Corded Ware occurred after the Yamnaya expansion had been ongoing for centuries. The Corded Ware also shares IBD segments with the Yamnaya[35] and thus its own expansion likely followed an admixture event during the 2900s BCE of a Yamnaya population with a Globular Amphora-related one.

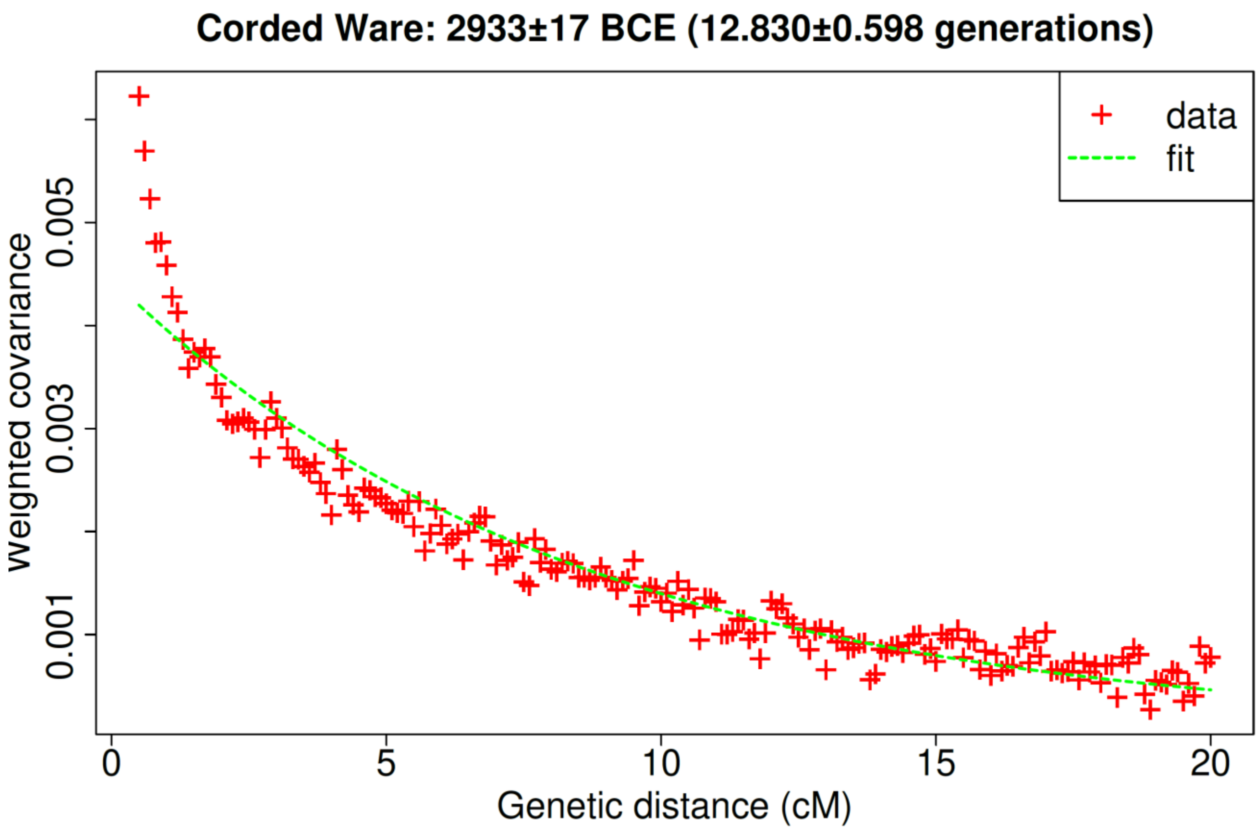

**Fig. S 8 Admixture LD estimation of Corded Ware formation**

#### Where and how was the Yamnaya formed?

Multiple observations build on each other to support the hypothesis that the Yamnaya were formed by movement of people like Remontnoye—groups of mixed Maikop/Aknashen "Caucasus" and BPgroup "Lower Volga" ancestry from the middle of the CLV cline—into the territory of the Serednii Stih or an adjacent area.

*First*, the Yamnaya are inferred (Table S 3) to derive from a mixture of SShi (~74%) Remontnoye (~26%) ancestry.

*Second*, the Serednii Stih people on the Lower Don (represented by Krivyansky) had about the same (~73%) SShi ancestry as the Yamnaya (Table S 22), albeit with other southern components of ancestry, consistent with multiple movements of different groups from the east or south into Serednii Stih territory. Conversely, east or south of the Serednii Stih, we see that Remontnoye, Maikop, and the Volga Cline can all be well-modeled without any Serednii Stih ancestry whatsoever. Thus, there is little evidence of expansion of Serednii Stih groups east and south, again providing evidence that the admixture was driven by movement from these regions into Serednii Stih territory. A possible exception is Steppe Maikop who we can model as having Serednii Stih ancestry, but Table S 21 shows that models without SShi fit even better in the tournament (the Steppe Maikop are also an outlier with respect to their extremely high Siberian-related ancestry, suggesting that their formation history is unusual).

*Third*, the Core Yamnaya are consistent with being part of the cline to which most Serednii Stih individuals also belong. While we do not have a sampled Serednii Stih population that is a perfect clade with the Core Yamnaya, we do have subsets of variable position along the Cline, including some (SShi) that approach the Core Yamnaya. Therefore it is parsimonious to assume that the Core Yamnaya are descended from an unsampled Serednii Stih population (of even lower hunter-gatherer ancestry), rather than the alternative that a different admixture proportion led to their formation and coincidentally resulted in a mixed population at precisely the edge of the Serednii Stih cline.

*Fourth*, the Serednii Stih-Yamnaya cline as a whole can be well-modeled with Core Yamnaya as one source. Naturally, the Core Yamnaya cannot have been the actual source (as it is later than the Eneolithic Serednii Stih culture). But, this fact argues strongly for the existence of some population (from which the Core Yamnaya itself emerged) that was driving differentiation across the Don-Dnipro area. This ancestral population of the Core Yamnaya (which we have called the "Pre-Yamnaya") must have been in close proximity to the Don-Dnipro area (if not in the area itself).

*Fifth*, either in the late Pre-Yamnaya period before the archaeological emergence of the Yamnaya horizon, or early in Yamnaya period, the "ancestral blend" characteristic of the Yamnaya contributed to both the Don Yamnaya and by the 3rd millennium BCE, the steppe ancestry in people of the Corded Ware culture[1,4,39,46-48] The date of shared ancestry between Yamnaya ancestors and people of the Corded Ware is definitively around the dawn of the Yamnaya culture in the second half of the 4th millennium BCE—not in the 5th millennium BCE or the beginning of the 4th millennium BCE as was recently hypothesized[42]—based on the finding of sharing of many large segments of DNA identical-by-descent between people of these two groups dating to the second half of the 4th millennium BCE[35], most plausibly to the core Yamnaya founder event that we date in this paper to ~3800-3400 BCE. While the location of emergence of the people of the Corded Ware is, itself, an open question given the expansive history of that culture after its emergence, it must have certainly been to the west of the Core Yamnaya, and at the same time the geographic neighbor of the Yamnaya.

*Sixth*, the Bell Beaker culture of Central/Western Europe[41] and early Corded Ware of Bohemia[46] was dominated by Y-haplogroup R-L151, a relative of the R-Z2103 Y-haplogroup of the core Yamnaya within haplogroup R-L23 (https://www.yfull.com/tree/R-L23//; v11.04.00) formed in 4450BCE. This argues for a relatively western origin of the core Yamnaya similarly to the Fifth point above.

We thus propose the following scenario:

Between the North Caucasus piedmont and Lower Volga population, mixtures were taking place during the 5th and 4th millennia BCE between populations with ancestry characteristic of the Neolithic and Eneolithic Caucasus (represented by Aknashen and Maikop) and Lower Volga-North Caucasus Eneolithic populations (represented by BPgroup), both of which had substantial proportions of CHG ancestry from the earlier hunter-gatherer periods. Steppe ancestry was present in the later Maikop (Table S 9) and Remontnoye (Table S 6, Table S 7) individuals from this area. It was also present further south in Chalcolithic Armenia at Areni-1 cave, but there it was added to a different, Masis Blur Neolithic population (Table S 11).[5] Thus, the Caucasus area was seeing admixture in both south-north and north-south directions: Remontnoye had Aknashen Neolithic/Maikop ancestry and Maikop and Armenian Chalcolithic had steppe ancestry.

People out of this Caucasus-Lower Volga admixture zone people flowed outwards: along the Volga where all the "southern" ancestry can be well explained as of BPgroup origin alone (Table S 14) and into the Don-Dnipro area where it interacted with the descendants of hunter-gatherers of the Dnipro-Don forming the Serednii-Stih cline. A "Pre-Yamnaya" population quite like the Core Yamnaya was at the other end of the cline (Table S 20) and Serednii Stih people had therefore not only BPgroup-related ancestry from the south (as people on the Volga did), but also some Aknashen/Maikop-related ancestry.

The Yamnaya themselves were thus the product of admixture between the Caucasus-Lower Volga easterners (for which Remontnoye is a genetic stand-in) with the Ukraine_N-admixed Serednii Stih westerners (for which SShi is a genetic stand-in). The Yamnaya must have been a subset of the wider "Pre-Yamnaya" population experiencing this admixture, although we can find only scant evidence for others like them, except perhaps the Vin1 sample from Vinogradnoe, one of the individuals of SShi most similar to the Yamnaya.

With our available sampling we cannot determine the geographical extent of the "Pre-Yamnaya", although it is plausible that it is within the Serednii Stih area itself or close-by, as these groups and the core Yamnaya all had Ukraine Neolithic Hunter-Gatherer ancestry. We also cannot determine the extent of admixture between the Pre-Yamnaya and their neighbors in the Serednii Stih culture leading to groups like the Don Yamnaya (Table S 20), the Core Yamnaya, and the Serednii Stih culture Krivyansky individual who had ancestry distinctive from the others we samples (Table S 22, Table S 23). Whatever the origins of the "Pre-Yamnaya" they gave rise to a very distinctive population of remarkable genetic homogeneity: many Yamnaya-Afanasievo sub-populations fit exactly the Remontnoye+SShi model (Table S 25).

| Test | P-value | Remontnoye | SShi | S.E. |
|---|---|---|---|---|
| Russia_Don_EMBA_Yamnaya_Catacomb_transitional | 0.188 | 2.4% | 97.6% | 7.0% |
| Russia_Kalmykia_EBA_Yamnaya | 0.071 | 8.6% | 91.4% | 8.8% |
| Ukraine_EBA_Yamnaya_o.SG | 0.129 | 13.0% | 87.0% | 11.8% |
| Russia_Afanasievo_Yenisei | 0.875 | 15.2% | 84.8% | 8.7% |
| Russia_Orlovka_EBA_Yamnaya | 0.402 | 20.9% | 79.1% | 7.5% |
| Russia_Ishkinovka_EBA_Yamnaya | 0.613 | 23.2% | 76.8% | 7.8% |
| Russia_Volga_EBA_Yamnaya | 0.445 | 23.9% | 76.1% | 4.9% |
| Russia_LowerVolga_EBA_Yamnaya | 0.576 | 24.0% | 76.0% | 22.3% |
| Russia_UpperOb_Eneolithic_Afanasievo | 0.139 | 24.2% | 75.8% | 4.7% |
| Russia_Samara_EBA_Yamnaya | 0.344 | 24.3% | 75.7% | 3.8% |
| Russia_Ural_EBA_Yamnaya | 0.269 | 24.5% | 75.5% | 4.6% |
| Russia_Afanasievo.SG | 0.337 | 24.6% | 75.4% | 6.8% |
| Russia_Chelyabinsk_EBA_Yamnaya | 0.399 | 24.6% | 75.4% | 4.9% |
| Russia_Kalmykia_EBA_Yamnaya.SG | 0.209 | 26.2% | 73.8% | 4.9% |
| Usatove_Yamnaya | 0.197 | 27.3% | 72.7% | 7.9% |
| Russia_Volgograd_EBA_Yamnaya | 0.633 | 27.6% | 72.4% | 4.3% |
| China_Xinjiang_G218_BA_Afanasievo_oWestEurasia | 0.858 | 28.9% | 71.1% | 7.5% |
| Kazakhstan_EBA_Yamnaya.SG | 0.317 | 29.2% | 70.8% | 8.0% |
| Russia_UpperYenisey_Eneolithic_Afanasievo | 0.101 | 29.7% | 70.3% | 5.3% |
| Mongolia_Chalcolithic_Afanasievo_1 | 0.116 | 30.1% | 69.9% | 7.7% |
| Russia_Afanasievo | 0.234 | 30.3% | 69.7% | 3.8% |
| Russia_CaspianInland_EBA_Yamnaya | 0.766 | 30.5% | 69.5% | 3.7% |
| Russia_LowerDon_EBA_Yamnaya | 0.513 | 31.9% | 68.1% | 7.3% |
| Russia_Remontnoye_EBA_Yamnaya | 0.908 | 32.2% | 67.8% | 4.3% |
| Ukraine_EBA_Yamnaya | 0.571 | 32.9% | 67.1% | 4.4% |
| Moldova_EBA_Yamnaya | 0.183 | 34.2% | 65.8% | 3.9% |
| Romania_EBA_Yamnaya | 0.678 | 35.5% | 64.5% | 4.3% |
| Russia_Caucasus_EBA_Yamnaya | 0.143 | 36.3% | 63.7% | 5.5% |

| | | | | |
|---|---|---|---|---|
| Romania_Brailita_EBA_Yamnaya | 0.099 | 37.7% | 62.3% | 8.2% |
| Russia_Volgograd_EBA_Yamnaya_o | 0.512 | 39.1% | 60.9% | 15.9% |
| Russia_StavropolKrai_EBA_Yamnaya | 0.105 | 41.7% | 58.3% | 7.3% |

**Table S 25 The Remontnoye+SShi model fits multiple Yamnaya-Afanasievo related populations**

#### A combined model of the ancestry of the Yamnaya and their Eneolithic neighbors

We have thus argued that Yamnaya was formed when people of "eastern" Aknashen-Maikop/BPgroup origins (of proximate Remontnoye-related origins) moved westward and admixed with people of the Serednii Stih culture. Motivated by this, we tested whether we could use this framework to jointly model all these populations involved in our reconstruction:

**Combined:** CoreYamnaya, Russia_Don_EBA_Yamnaya, SSlo, SSmed, SShi, Remontnoye, Maikop

A single 3-way model fits all these populations and three fit all of them but one. We show all these models as well as the Aknashen+BPgroup+Ukraine_N model in Table S 26. The difference between the models is in the 3rd ancestral source. The model that fits all seven populations well is Aknashen+BPgroup+GK2. The model with GK1 (instead of GK2) as a source predicts a negative admixture coefficient for SSlo (which is the one population that fails it). The model with Igren_o as a source has a high standard error for the estimation of the ancestry of SSlo; this model is qualitatively similar to the one with the GK2 source, since as we have seen Igren_o forms a clade with GK2, but this model has higher standard errors due to lower data quality of Igren_o. The model with SSlo as the source explains all populations (except itself, as SSlo is included as a source). While the model with Ukraine_N as the source fails for three of the seven populations, in fact these three population almost fit (p=0.02-0.05), and it is plausibly that this model is in fact as good as the others and it is producing more evidence of imperfect fits simply because the much larger sample size of Ukraine_N than GK1, GK2, SSlo, or Igren_o provides more power to falsify models.

| | | | Proportions | | | Std. errors | | |
|---|---|---|---|---|---|---|---|---|
| **Test** | **C** | **P-value** | **A=Aknashen** | **B=BPgroup** | **C** | **A=Aknashen** | **B=BPgroup** | **C** |
| CoreYamnaya | GK1 | 0.801 | 19.7% | 43.3% | 37.0% | 1.5% | 3.7% | 3.2% |
| Maikop | GK1 | 0.474 | 87.2% | 6.7% | 6.1% | 3.2% | 8.6% | 6.9% |
| Remontnoye | GK1 | 0.524 | 44.4% | 56.7% | -1.1% | 2.8% | 7.2% | 5.8% |
| Russia_Don_EBA_Yamnaya | GK1 | 0.829 | 18.5% | 10.0% | 71.4% | 2.4% | 6.1% | 5.2% |
| SShi | GK1 | 0.627 | 11.4% | 36.6% | 52.0% | 2.6% | 6.4% | 5.3% |
| SSlo | GK1 | 0.519 | 10.7% | -41.9% | 131.2% | 6.3% | 16.9% | 14.0% |
| SSmed | GK1 | 1.000 | 13.9% | -6.2% | 92.4% | 3.0% | 8.2% | 6.9% |
| **CoreYamnaya** | **GK2** | **0.934** | **20.7%** | **56.8%** | **22.5%** | **1.3%** | **2.5%** | **1.8%** |
| **Maikop** | **GK2** | **0.458** | **87.4%** | **9.1%** | **3.5%** | **3.3%** | **6.5%** | **4.2%** |
| **Remontnoye** | **GK2** | **0.520** | **44.4%** | **56.1%** | **-0.5%** | **2.9%** | **5.4%** | **3.5%** |
| **Russia_Don_EBA_Yamnaya** | **GK2** | **0.708** | **20.6%** | **36.3%** | **43.2%** | **1.8%** | **3.2%** | **2.2%** |
| **SShi** | **GK2** | **0.305** | **13.1%** | **55.6%** | **31.3%** | **2.3%** | **4.3%** | **3.1%** |
| **SSlo** | **GK2** | **0.102** | **13.6%** | **7.7%** | **78.7%** | **5.3%** | **10.0%** | **7.1%** |
| **SSmed** | **GK2** | **0.851** | **16.5%** | **27.4%** | **56.1%** | **2.7%** | **5.1%** | **3.5%** |
| CoreYamnaya | Igren_o | 0.999 | 21.3% | 55.8% | 22.9% | 1.8% | 3.4% | 2.4% |
| Maikop | Igren_o | 0.427 | 87.3% | 9.7% | 3.0% | 3.3% | 6.9% | 4.5% |
| Remontnoye | Igren_o | 0.524 | 44.3% | 56.2% | -0.6% | 2.9% | 5.7% | 3.8% |
| Russia_Don_EBA_Yamnaya | Igren_o | 0.633 | 21.4% | 34.7% | 44.0% | 2.9% | 5.4% | 3.8% |
| SShi | Igren_o | 0.666 | 13.4% | 54.7% | 32.0% | 2.9% | 5.3% | 3.5% |
| SSlo | Igren_o | 0.190 | 15.2% | 3.9% | 80.9% | 7.4% | 13.8% | 9.5% |
| SSmed | Igren_o | 0.875 | 17.7% | 25.8% | 56.5% | 3.9% | 6.7% | 4.6% |
| CoreYamnaya | SSlo | 0.104 | 16.7% | 55.6% | 27.7% | 1.6% | 3.7% | 3.2% |
| Maikop | SSlo | 0.550 | 86.8% | 7.5% | 5.7% | 3.0% | 6.6% | 5.4% |
| Remontnoye | SSlo | 0.522 | 44.5% | 56.1% | -0.6% | 2.8% | 5.7% | 4.5% |
| Russia_Don_EBA_Yamnaya | SSlo | 0.269 | 13.2% | 32.5% | 54.4% | 2.6% | 5.5% | 4.7% |
| SShi | SSlo | 0.283 | 7.2% | 54.4% | 38.4% | 2.7% | 5.6% | 4.7% |
| SSmed | SSlo | 0.328 | 6.6% | 23.5% | 69.9% | 3.7% | 7.3% | 6.1% |

| | | | | | | | | |
|---|---|---|---|---|---|---|---|---|
| CoreYamnaya | Ukraine_N | 0.038 | 18.0% | 64.2% | 17.7% | 1.1% | 1.8% | 1.3% |
| Maikop | Ukraine_N | 0.530 | 87.0% | 9.3% | 3.7% | 3.1% | 5.4% | 3.6% |
| Remontnoye | Ukraine_N | 0.516 | 44.5% | 55.9% | -0.3% | 2.7% | 4.4% | 2.9% |
| Russia_Don_EBA_Yamnaya | Ukraine_N | 0.020 | 15.3% | 49.8% | 34.9% | 1.2% | 1.8% | 1.3% |
| SShi | Ukraine_N | 0.064 | 8.8% | 66.4% | 24.8% | 2.1% | 3.3% | 2.3% |
| SSlo | Ukraine_N | 0.878 | 4.7% | 29.2% | 66.1% | 4.4% | 7.2% | 5.2% |
| SSmed | Ukraine_N | 0.046 | 9.8% | 44.3% | 45.9% | 1.9% | 3.2% | 2.2% |

**Table S 26 Joint models of populations involved in our scenario of Yamnaya origins.** We highlight in bold the model shown in Fig. S 9

Based on this analysis, the hunter-gatherer source of the Serednii Stih-Yamnaya cline is ambiguous and is drawn from the continuum of EHG-Ukraine_N (along which Ukraine_N and GK2 are both points). We show a ternary plot of the Aknashen+BPgroup+GK2 model (Fig. S 9), as a contrast to the also-plausible models with Ukraine_N in place of GK2 that we discuss elsewhere in this note and in the text. If we assume that the Serednii Stih admixing population is like SShi and that the eastern population X was an Aknashen/Bpgroup admixture, we may estimate the proportions of that population as follows.

Sshi = (13.1, 55.6, 31.3)
CoreYamnaya = (20.7, 56.8, 22.5)
X = (x, y, 0)
CoreYamnaya = αX+(1-α)Sshi
Which yields the following system of equations:

$$20.7 = ax + 13.1 - 13.1a$$
$$56.8 = ay + 55.6 - 55.6a$$
$$22.5 = 31.3 - 31.3a$$

Solving implies that Yamnaya had α=28.1% eastern ancestry which itself was composed of 40.1% Aknashen and 59.9% Bpgroup ancestry, quite similar to the balance of the two components in the sampled Remontnoye. In Fig. S 9 we also show the two Remontnoye individuals separately which fall on slightly different positions on the Aknashen-Bpgroup cline.

When we analyze the two Remontnoye individuals separately, we do indeed see that they vary significantly in their relative proportions of Aknashen and BPgroup ancestry and that the earlier individual (I28683; ~4000BCE) has more BPgroup ancestry than the later one (I28682; ~3700BCE), a difference of 61.6±3.6% vs. 47.8±3.6%. The difference between the two individuals is also visible in the 3-way model of Fig. S 9 and is direct evidence for the variable cline between Caucasus and steppe genetic ancestry on which our population X resides, with the combined Remontnoye population virtually identical to it.

Observe also the previously mentioned contrast between Eneolithic populations of the Don-Volga who did not require Aknashen/Maikop ancestry but can be modeled with BPgroup/PVgroup ancestry alone (Fig. S 2, Fig. S 3) with those of the Serednii Stih culture of the Don-Dnipro area that also had Aknashen-related ancestry (Fig. S 9).

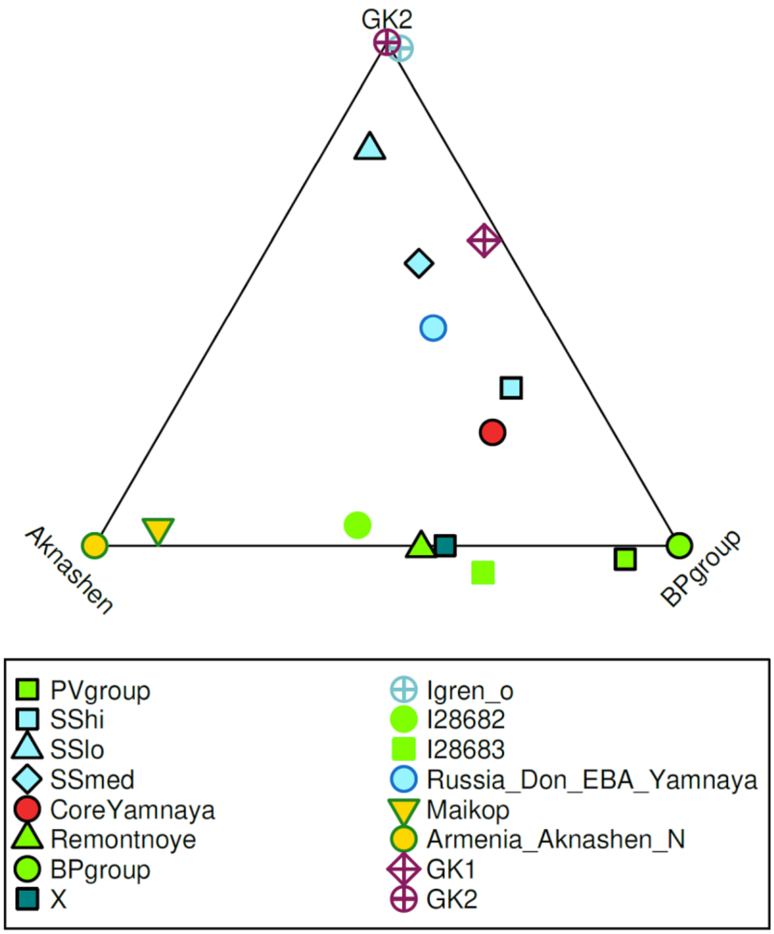

**Fig. S 9 A combined model of Yamnaya origins.** Population X represents the point on the Aknashen-BPgroup cline if SShi is fixed as one of the two sources of the core Yamnaya. The two Remontnoye individuals I28683 and I28682 are also shown separately.

##### Full exploration of 3-way models for the core Yamnaya

The models of Table S 1 are derived by fixing two of the sources (Aknashen and BPgroup) since these two sources can be used effectively to model Maikop and the Remontnoye individuals which, according to our reconstruction, may be proximate sources for the core Yamnaya.

However, other 3-way models are also feasible for the Yamnaya if we remove this constraint; a total of 274 3-way models are possible, up from the 4 2-way models of Table S 3. We can add a further constraint by requiring the models to fit not only the Yamnaya but many of the 7 populations of the Combined set. A total of 22 3-way models can be used to model at least 6 of these populations and we list them in Table S 27. These models invariably include a Lower Volga-North Caucasus Eneolithic population (BPgroup or PVgroup), a Dnipro-Don population, and a Caucasus Neolithic or Chalcolithic population.

| A | B | C | P-value | A | B | C | S.E. A | S.E. B | S.E. C | Win | Draw | Lose | Score |
|---|---|---|---|---|---|---|---|---|---|---|---|---|---|
| **Armenia_Aknashen_N** | **BPgroup** | **Igren_o** | **0.999** | **21.3%** | **55.8%** | **22.9%** | **1.8%** | **3.4%** | **2.4%** | **12** | **9** | **0** | **12** |
| **Azerbaijan_C** | **BPgroup** | **Igren_o** | **0.553** | **19.5%** | **60.2%** | **20.4%** | **1.6%** | **3.0%** | **2.2%** | **12** | **9** | **0** | **12** |
| Armenia_Aknashen_N | BPgroup | GK1 | 0.801 | 19.7% | 43.3% | 37.0% | 1.5% | 3.7% | 3.2% | 9 | 11 | 1 | 8 |
| **Armenia_Aknashen_N** | **Igren_o** | **PVgroup** | **0.874** | **15.5%** | **25.2%** | **59.3%** | **2.5%** | **2.8%** | **4.3%** | **4** | **17** | **0** | **4** |
| **Azerbaijan_C** | **Igren_o** | **PVgroup** | **0.667** | **13.6%** | **23.0%** | **63.4%** | **2.2%** | **2.7%** | **4.0%** | **4** | **17** | **0** | **4** |
| Azerbaijan_N | Igren_o | PVgroup | 0.770 | 12.6% | 22.1% | 65.3% | 2.0% | 2.5% | 3.5% | 4 | 16 | 1 | 3 |
| Armenia_Aknashen_N | BPgroup | SSlo | 0.104 | 16.7% | 55.6% | 27.7% | 1.6% | 3.7% | 3.2% | 6 | 10 | 5 | 1 |
| GK2 | Maikop | PVgroup | 0.459 | 23.4% | 16.6% | 60.0% | 2.1% | 2.2% | 3.4% | 3 | 16 | 2 | 1 |
| BPgroup | GK2 | Maikop | 0.418 | 54.2% | 21.9% | 23.9% | 2.2% | 1.5% | 1.3% | 4 | 14 | 3 | 1 |
| Armenia_Aknashen_N | GK2 | PVgroup | 0.634 | 14.4% | 24.1% | 61.5% | 2.0% | 2.2% | 3.5% | 3 | 15 | 3 | 0 |
| Armenia_Aknashen_N | BPgroup | GK2 | 0.934 | 20.7% | 56.8% | 22.5% | 1.3% | 2.5% | 1.8% | 2 | 16 | 3 | -1 |
| Armenia_MasisBlur_N | Igren_o | PVgroup | 0.782 | 11.1% | 21.6% | 67.3% | 1.8% | 2.6% | 3.4% | 1 | 17 | 3 | -2 |
| Armenia_MasisBlur_N | PVgroup | Ukraine_N | 0.321 | 8.2% | 74.5% | 17.2% | 1.5% | 2.3% | 1.7% | 1 | 17 | 3 | -2 |
| Azerbaijan_C | GK2 | PVgroup | 0.358 | 12.6% | 22.1% | 65.3% | 1.9% | 2.1% | 3.1% | 2 | 15 | 4 | -2 |
| Azerbaijan_C | PVgroup | Ukraine_N | 0.102 | 9.5% | 72.8% | 17.7% | 1.8% | 2.6% | 1.7% | 1 | 17 | 3 | -2 |
| Azerbaijan_N | GK2 | PVgroup | 0.505 | 11.8% | 21.4% | 66.8% | 1.6% | 2.2% | 2.9% | 2 | 15 | 4 | -2 |
| Armenia_Aknashen_N | PVgroup | Ukraine_N | 0.113 | 10.3% | 70.9% | 18.8% | 1.9% | 2.6% | 1.6% | 1 | 17 | 3 | -2 |
| Armenia_Aknashen_N | GK1 | PVgroup | 0.614 | 14.9% | 39.3% | 45.8% | 2.1% | 3.7% | 4.8% | 2 | 15 | 4 | -2 |
| Azerbaijan_N | PVgroup | Ukraine_N | 0.162 | 9.0% | 73.7% | 17.3% | 1.7% | 2.4% | 1.7% | 1 | 17 | 3 | -2 |
| Armenia_Aknashen_N | PVgroup | SSlo | 0.153 | 10.0% | 61.2% | 28.8% | 2.2% | 4.6% | 3.6% | 0 | 18 | 3 | -3 |
| Armenia_Aknashen_N | GK1 | Khi | 0.105 | 26.0% | 26.9% | 47.1% | 1.2% | 4.2% | 4.4% | 0 | 13 | 8 | -8 |
| Armenia_Aknashen_N | KhlopkovBugor | Ukraine_N | 0.160 | 29.0% | 60.5% | 10.5% | 1.5% | 2.8% | 2.1% | 0 | 3 | 18 | -18 |

**Table S 27 Full exploration of 3-way models.** Models shown fit the Core Yamnaya and at least 5 other populations of the Combined Set. Models without any losses in the tournament are highlighted in bold.

In Table S 27 that only models involving Aknashen_B/Azerbaijan_C and BPgroup/PVgroup and Igren_o ancestries have no losses in the tournament, the full results of which are shown in Table S 28. While no unique solution emerges out of this tournament, it is useful to weigh our confidence in the different models. With the knowledge that we have only partially sampled the genetic variation of the Caucasus, Lower Volga, and Dnipro-Don areas, it is nonetheless interesting that the 3-way models directly recapitulate the conclusions we reached by exploring 2-way models: that all three of these areas contributed to the formation of the Yamnaya.

What was the order in which the three components admixed? The existence of the Serednii Stih-Yamnaya cline is itself a powerful argument in favor of Dnipro-Don ancestry admixing with Caucasus-Volga ancestry, a conclusion which maps well to the Remontnoye+SShi proximal model. Thus, we can tentatively conclude that admixture between people from the Caucasus-Lower Volga cline (like Remontnoye if not necessarily identical to it), admixed with people from the Dnipro-Don area (like SShi, if not necessarily identical to it) to form the Yamnaya.

| | Armenia_MasisBlur_N+Igren_o+PVgroup | Armenia_Aknashen_N+BPgroup+GK1 | Armenia_Aknashen_N+BPgroup+GK2 | Armenia_Aknashen_N+BPgroup+Igren_o | Armenia_Aknashen_N+BPgroup+SSlo | Armenia_MasisBlur_N+PVgroup+Ukraine_N | Azerbaijan_C+BPgroup+Igren_o | Azerbaijan_C+GK2+PVgroup | Azerbaijan_C+Igren_o+PVgroup | Azerbaijan_C+PVgroup+Ukraine_N | Azerbaijan_N+GK2+PVgroup | Azerbaijan_N+Igren_o+PVgroup | Armenia_Aknashen_N+GK1+Khi | Armenia_Aknashen_N+GK1+PVgroup | Azerbaijan_N+PVgroup+Ukraine_N | Armenia_Aknashen_N+GK2+PVgroup | BPgroup+GK2+Maikop | Armenia_Aknashen_N+Igren_o+PVgroup | Armenia_Aknashen_N+KhlopkovBugor+Ukraine_N | GK2+Maikop+PVgroup | Armenia_Aknashen_N+PVgroup+SSlo | Armenia_Aknashen_N+PVgroup+Ukraine_N |
|---|---|---|---|---|---|---|---|---|---|---|---|---|---|---|---|---|---|---|---|---|---|---|
| Armenia_MasisBlur_N+Igren_o+PVgroup | | 5.9E-01 | 1.7E-02 | 4.2E-01 | 5.5E-02 | 4.4E-01 | 9.4E-02 | 4.7E-01 | 3.2E-01 | 2.1E-01 | 4.9E-01 | 4.6E-01 | 1.7E-01 | 7.9E-01 | 2.9E-01 | 5.8E-01 | 2.1E-04 | 5.2E-01 | 2.1E-02 | 3.1E-01 | 3.0E-01 | 2.2E-01 |
| Armenia_Aknashen_N+BPgroup+GK1 | 5.3E-02 | | 1.9E-01 | 1.0E+00 | 1.4E-01 | 4.3E-06 | 7.4E-01 | 1.0E-02 | 3.9E-01 | 1.5E-04 | 2.2E-04 | 1.2E-01 | 1.1E-02 | 4.8E-01 | 4.4E-06 | 4.4E-03 | 8.4E-02 | 5.2E-01 | 3.5E-02 | 6.7E-03 | 1.6E-02 | 4.4E-05 |
| Armenia_Aknashen_N+BPgroup+GK2 | 3.8E-02 | 5.1E-02 | | 9.2E-01 | 1.7E-01 | 3.7E-06 | 4.3E-01 | 1.1E-02 | 2.0E-01 | 1.6E-04 | 3.0E-04 | 7.6E-02 | 1.0E-02 | 1.7E-02 | 4.3E-06 | 2.0E-02 | 3.9E-01 | 3.4E-01 | 3.9E-02 | 2.2E-02 | 2.0E-02 | 4.2E-05 |
| Armenia_Aknashen_N+BPgroup+Igren_o | 3.0E-02 | 7.6E-01 | 4.4E-01 | | 9.7E-02 | 4.7E-06 | 6.8E-01 | 2.1E-02 | 3.0E-01 | 1.8E-04 | 4.3E-04 | 7.4E-02 | 2.2E-02 | 5.6E-01 | 4.2E-06 | 2.2E-02 | 6.2E-02 | 3.9E-01 | 4.5E-02 | 1.9E-02 | 1.7E-02 | 4.0E-05 |
| Armenia_Aknashen_N+BPgroup+SSlo | 4.8E-02 | 5.9E-01 | 8.3E-01 | 9.4E-01 | | 4.3E-06 | 7.2E-01 | 2.0E-02 | 4.1E-01 | 1.9E-04 | 7.0E-04 | 1.2E-01 | 1.4E-02 | 4.9E-01 | 4.7E-06 | 3.7E-02 | 4.9E-01 | 4.9E-01 | 2.5E-02 | 3.9E-02 | 1.1E-02 | 4.2E-05 |
| Armenia_MasisBlur_N+PVgroup+Ukraine_N | 4.0E-01 | 4.9E-01 | 3.0E-02 | 3.1E-01 | 2.6E-02 | | 1.1E-01 | 3.3E-01 | 2.7E-01 | 1.3E-01 | 4.7E-01 | 3.6E-01 | 1.6E-01 | 5.1E-01 | 2.3E-01 | 4.0E-01 | 7.4E-04 | 3.0E-01 | 1.4E-02 | 2.2E-01 | 1.3E-01 | 1.7E-01 |
| Azerbaijan_C+BPgroup+Igren_o | 4.8E-03 | 8.4E-01 | 4.2E-01 | 6.8E-01 | 1.1E-01 | 4.5E-06 | | 1.7E-02 | 2.4E-01 | 1.2E-04 | 4.8E-04 | 2.8E-02 | 3.5E-02 | 6.6E-01 | 6.5E-06 | 2.5E-02 | 9.6E-02 | 1.9E-01 | 7.3E-02 | 3.2E-02 | 2.1E-02 | 8.0E-05 |
| Azerbaijan_C+GK2+PVgroup | 2.3E-01 | 6.0E-02 | 2.3E-02 | 3.7E-01 | 6.4E-02 | 4.0E-01 | 1.3E-01 | | 2.6E-01 | 8.1E-02 | 5.3E-01 | 3.0E-01 | 7.5E-02 | 2.7E-02 | 2.4E-01 | 6.4E-01 | 5.2E-03 | 3.2E-01 | 1.6E-02 | 5.7E-01 | 2.0E-01 | 1.9E-01 |
| Azerbaijan_C+Igren_o+PVgroup | 3.9E-01 | 5.7E-01 | 1.5E-02 | 3.9E-01 | 3.9E-02 | 5.1E-01 | 2.2E-01 | 4.8E-01 | | 1.7E-01 | 4.7E-01 | 5.4E-01 | 1.6E-01 | 7.8E-01 | 3.1E-01 | 5.3E-01 | 1.1E-03 | 5.2E-01 | 2.1E-02 | 4.1E-01 | 2.3E-01 | 2.3E-01 |
| Azerbaijan_C+PVgroup+Ukraine_N | 2.4E-01 | 4.7E-01 | 2.6E-02 | 2.6E-01 | 1.8E-02 | 4.0E-01 | 2.1E-01 | 3.1E-01 | 4.6E-01 | | 4.3E-01 | 4.3E-01 | 1.5E-01 | 5.1E-01 | 2.5E-01 | 3.6E-01 | 3.6E-03 | 2.8E-01 | 1.4E-02 | 2.8E-01 | 9.1E-02 | 1.8E-01 |
| Azerbaijan_N+GK2+PVgroup | 5.2E-01 | 5.5E-02 | 3.4E-02 | 6.2E-01 | 7.0E-02 | 4.4E-01 | 1.2E-01 | 3.2E-01 | 2.5E-01 | 1.2E-01 | | 4.1E-01 | 7.7E-02 | 2.2E-02 | 1.6E-01 | 6.5E-01 | 5.0E-03 | 5.7E-01 | 1.1E-02 | 5.2E-01 | 2.7E-01 | 1.9E-01 |
| Azerbaijan_N+Igren_o+PVgroup | 8.0E-01 | 5.7E-01 | 2.0E-02 | 6.7E-01 | 4.2E-02 | 5.6E-01 | 1.9E-01 | 4.2E-01 | 5.7E-01 | 2.3E-01 | 4.4E-01 | | 1.6E-01 | 7.4E-01 | 2.2E-01 | 5.3E-01 | 1.1E-03 | 8.5E-01 | 1.6E-02 | 3.5E-01 | 2.9E-01 | 2.3E-01 |
| Armenia_Aknashen_N+GK1+Khi | 8.7E-01 | 8.7E-01 | 2.8E-01 | 1.0E+00 | 1.3E-01 | 2.4E-01 | 8.2E-01 | 2.1E-01 | 9.0E-01 | 1.9E-01 | 1.6E-01 | 9.2E-01 | | 7.4E-01 | 1.9E-01 | 8.9E-02 | 1.3E-01 | 9.4E-01 | 1.5E-01 | 8.4E-02 | 1.7E-01 | 1.1E-01 |
| Armenia_Aknashen_N+GK1+PVgroup | 8.3E-01 | 4.4E-01 | 6.4E-03 | 7.6E-01 | 4.6E-02 | 3.7E-01 | 3.3E-01 | 1.4E-01 | 8.4E-01 | 1.9E-01 | 1.1E-01 | 8.6E-01 | 6.4E-02 | | 2.3E-01 | 5.2E-02 | 7.1E-04 | 9.0E-01 | 9.5E-03 | 5.4E-02 | 1.3E-01 | 1.2E-01 |
| Azerbaijan_N+PVgroup+Ukraine_N | 4.4E-01 | 4.7E-01 | 3.6E-02 | 4.4E-01 | 2.2E-02 | 4.5E-01 | 1.7E-01 | 2.8E-01 | 3.7E-01 | 1.5E-01 | 4.2E-01 | 5.3E-01 | 1.5E-01 | 4.7E-01 | | 3.8E-01 | 3.5E-03 | 4.3E-01 | 1.0E-02 | 2.6E-01 | 1.3E-01 | 1.8E-01 |
| Armenia_Aknashen_N+GK2+PVgroup | 5.4E-01 | 3.5E-02 | 2.3E-02 | 6.0E-01 | 5.4E-02 | 3.9E-01 | 1.8E-01 | 4.2E-01 | 3.4E-01 | 1.1E-01 | 5.6E-01 | 5.0E-01 | 5.0E-02 | 1.4E-02 | 2.1E-01 | | 1.8E-03 | 5.4E-01 | 8.5E-03 | 4.5E-01 | 1.9E-01 | 1.3E-01 |
| BPgroup+GK2+Maikop | 4.6E-02 | 2.0E-02 | 5.3E-01 | 9.3E-01 | 3.1E-03 | 5.1E-06 | 2.4E-01 | 1.1E-02 | 2.3E-01 | 1.4E-04 | 5.0E-04 | 8.8E-02 | 9.5E-03 | 2.9E-02 | 6.7E-06 | 3.3E-02 | | 4.5E-01 | 6.1E-03 | 2.4E-02 | 5.0E-03 | 6.9E-05 |
| Armenia_Aknashen_N+Igren_o+PVgroup | 8.4E-01 | 4.7E-01 | 1.5E-02 | 6.5E-01 | 3.2E-02 | 5.1E-01 | 3.0E-01 | 5.3E-01 | 7.5E-01 | 2.3E-01 | 5.1E-01 | 8.3E-01 | 1.1E-01 | 6.9E-01 | 2.8E-01 | 5.3E-01 | 4.0E-04 | | 1.2E-02 | 3.4E-01 | 2.0E-01 | 1.6E-01 |
| Armenia_Aknashen_N+KhlopkovBugor+Ukraine_N | 6.1E-01 | 7.0E-01 | 7.7E-01 | 7.9E-01 | 7.7E-02 | 5.2E-01 | 7.2E-01 | 3.0E-01 | 5.6E-01 | 1.9E-01 | 4.8E-01 | 6.9E-01 | 2.4E-01 | 2.1E-01 | 3.0E-01 | 3.0E-01 | 2.7E-01 | 4.9E-01 | | 1.3E-01 | 3.8E-02 | 1.7E-01 |
| GK2+Maikop+PVgroup | 4.8E-01 | 1.1E-02 | 6.0E-03 | 6.5E-01 | 5.6E-04 | 4.3E-01 | 9.0E-02 | 5.0E-01 | 3.6E-01 | 1.3E-01 | 6.0E-01 | 4.2E-01 | 3.3E-02 | 2.7E-02 | 2.3E-01 | 7.5E-01 | 2.5E-03 | 6.0E-01 | 1.8E-03 | | 1.7E-01 | 1.9E-01 |
| Armenia_Aknashen_N+PVgroup+SSlo | 8.4E-01 | 3.5E-01 | 3.9E-02 | 6.8E-01 | 3.0E-02 | 4.4E-01 | 3.8E-01 | 5.4E-01 | 8.2E-01 | 2.2E-01 | 6.6E-01 | 8.6E-01 | 8.7E-02 | 5.9E-01 | 2.8E-01 | 7.1E-01 | 3.9E-03 | 8.9E-01 | 5.5E-03 | 5.7E-01 | | 1.4E-01 |
| Armenia_Aknashen_N+PVgroup+Ukraine_N | 5.0E-01 | 3.7E-01 | 2.6E-02 | 4.3E-01 | 1.4E-02 | 4.0E-01 | 2.9E-01 | 3.8E-01 | 5.7E-01 | 1.5E-01 | 4.9E-01 | 6.3E-01 | 1.0E-01 | 3.9E-01 | 2.2E-01 | 3.4E-01 | 1.6E-03 | 4.3E-01 | 7.2E-03 | 2.4E-01 | 7.6E-02 | |

**Table S 28 Model tournament between models of Table S 27.** Results of (A, B) matches are shown in A=columns and B=rows of the matrix.

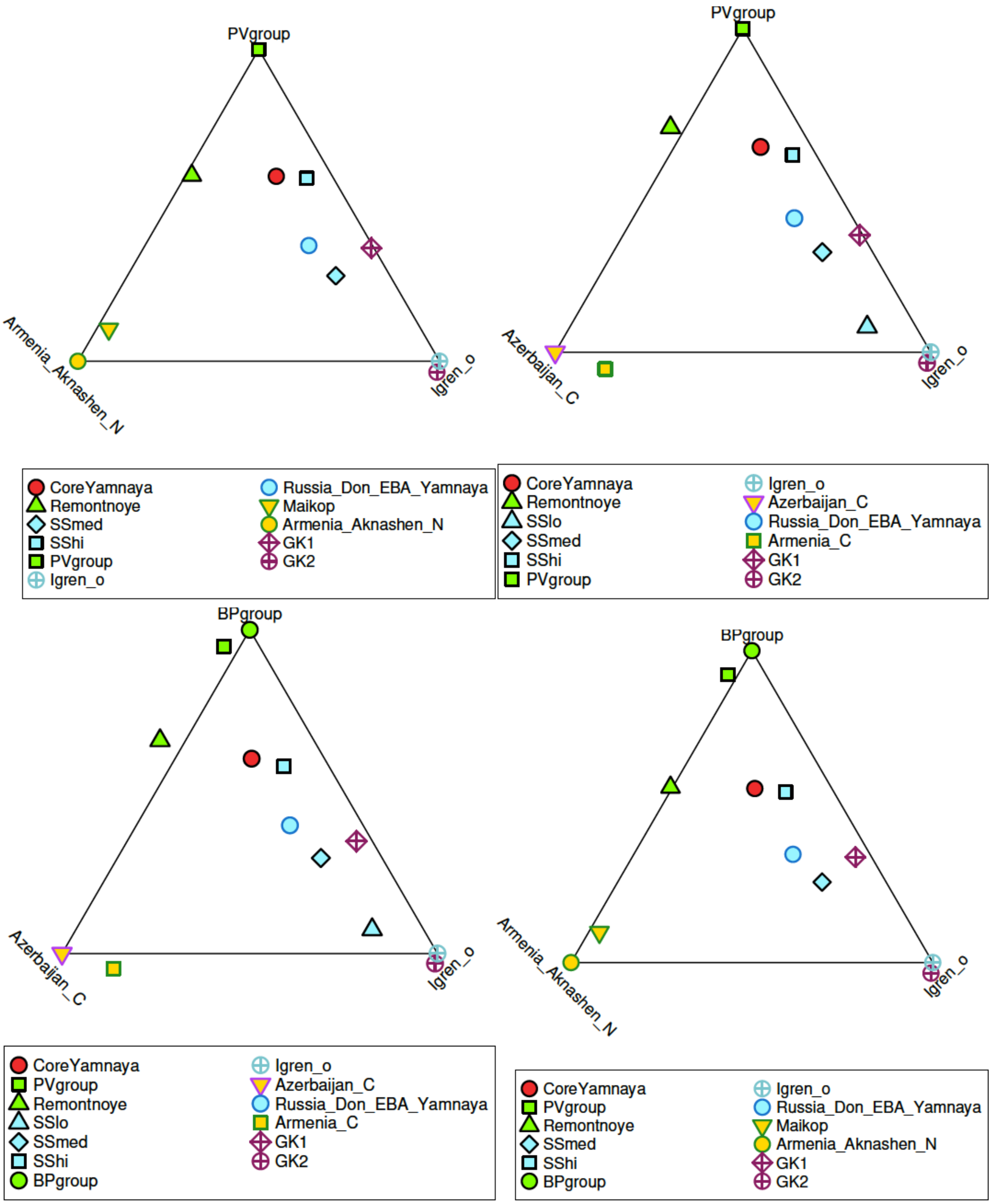

**Fig. S 10 Unconstrained 3-way models that have no losses in tournament of Table S 27.** All Test populations that fit each of these models is shown.

The four 3-way models with no losses are shown in Fig. S 10

**A combined model for the entire Dnipro-Don-Volga-Caucasus region**

Finally, we observe that since we can model the Don-Volga populations as a 3-way mixture of BPgroup, Aknashen, and Ukraine_N/GK2/Igren_o and we can also model the Volga cline populations as a mixture of BPgroup and an Eastern hunter-gatherer related source, we might be able to model all populations of interest using a 4-way model. We show in Table S 29 a model with BPgroup, Aknashen, Ukraine_N, and Lebyazhinka_HG sources which summarizes the ancestry of all populations of interest across the Dnipro-Don-Volga-Caucasus region of interest. And fits all of them with relatively small standard errors.

| Test | P-value | A= Lebyazhinka_HG | B= Armenia_Aknashen_N | C=BPgroup | D= Ukraine_N | S.E. A | S.E. B | S.E. C | S.E. D |
|---|---|---|---|---|---|---|---|---|---|
| CoreYamnaya | 0.631 | 9.9% | 22.0% | 55.8% | 12.4% | 3.6% | 1.9% | 3.6% | 2.3% |
| Ekaterinovka | 0.223 | 89.9% | 0.8% | 15.9% | -6.7% | 8.8% | 4.2% | 8.1% | 6.0% |
| GK1 | 0.796 | 16.2% | 2.1% | 41.7% | 40.1% | 8.2% | 4.4% | 8.0% | 5.9% |
| GK2 | 0.871 | 38.7% | 3.7% | -0.1% | 57.6% | 9.8% | 5.0% | 9.0% | 7.0% |
| Khi | 0.208 | 30.7% | -0.6% | 75.8% | -6.0% | 5.1% | 2.7% | 5.0% | 3.3% |
| KhlopkovBugor | 0.074 | 32.0% | -4.6% | 77.1% | -4.4% | 8.9% | 4.7% | 8.6% | 6.1% |
| Klo | 0.205 | 65.3% | 0.2% | 34.9% | -0.4% | 7.2% | 3.6% | 6.7% | 4.9% |
| Kmed | 0.674 | 54.9% | 2.2% | 50.6% | -7.7% | 6.4% | 3.3% | 6.1% | 4.4% |
| Krivyansky | 0.187 | -27.0% | 11.0% | 89.3% | 26.6% | 11.1% | 5.8% | 11.0% | 7.0% |
| Labazy | 0.237 | 97.3% | 3.2% | 9.1% | -9.6% | 11.7% | 5.9% | 11.1% | 7.8% |
| Maikop | 0.553 | -10.2% | 81.9% | 19.1% | 9.2% | 8.8% | 5.3% | 10.0% | 5.6% |
| Maximovka | 0.609 | 80.3% | -6.3% | 13.0% | 12.9% | 9.8% | 5.1% | 9.2% | 6.7% |
| Murzikha | 0.665 | 80.2% | -3.8% | 3.5% | 20.2% | 8.1% | 4.2% | 7.6% | 5.6% |
| PVgroup | 0.553 | -0.8% | 10.5% | 91.7% | -1.4% | 7.0% | 3.6% | 7.2% | 4.2% |
| Remontnoye | 0.324 | 1.9% | 45.0% | 54.3% | -1.2% | 8.0% | 4.6% | 8.5% | 5.2% |
| Russia_Don_EBA_Yamnaya | 0.796 | 11.0% | 19.6% | 40.3% | 29.0% | 3.6% | 2.0% | 3.8% | 2.3% |
| Russia_Karelia | 0.588 | 118.5% | 1.0% | -9.5% | -10.0% | 11.9% | 6.0% | 11.1% | 8.0% |
| SShi | 0.103 | 9.9% | 12.9% | 57.7% | 19.5% | 6.4% | 3.5% | 6.4% | 4.3% |
| SSlo | 0.783 | -6.8% | 2.2% | 35.2% | 69.4% | 13.6% | 6.8% | 13.3% | 9.1% |
| SSmed | 0.691 | 16.9% | 16.4% | 30.9% | 35.8% | 6.2% | 3.1% | 5.9% | 4.2% |
| Syezzheye | 0.189 | 86.4% | 4.7% | 17.4% | -8.5% | 11.2% | 5.7% | 10.6% | 7.5% |

**Table S 29 A 4-way model for the entire Dnipro-Don-Volga-Caucasus region.** Proportions are feasible except for Krivyansky which has a negative proportion -27.0±11.1% of Lebyazhinka_HG ancestry. As Krivyansky has more CHG-related ancestry than BPgroup (Table S 23), the 4-way model adjusts the BPgroup-related ancestry upwards to 89.3±11.0%: this accounts for the CHG-related ancestry in Krivyansky but also brings in more EHG-related ancestry than exists in Krivyansky, hence the negative Lebyazhinka_HG ancestry coefficient.

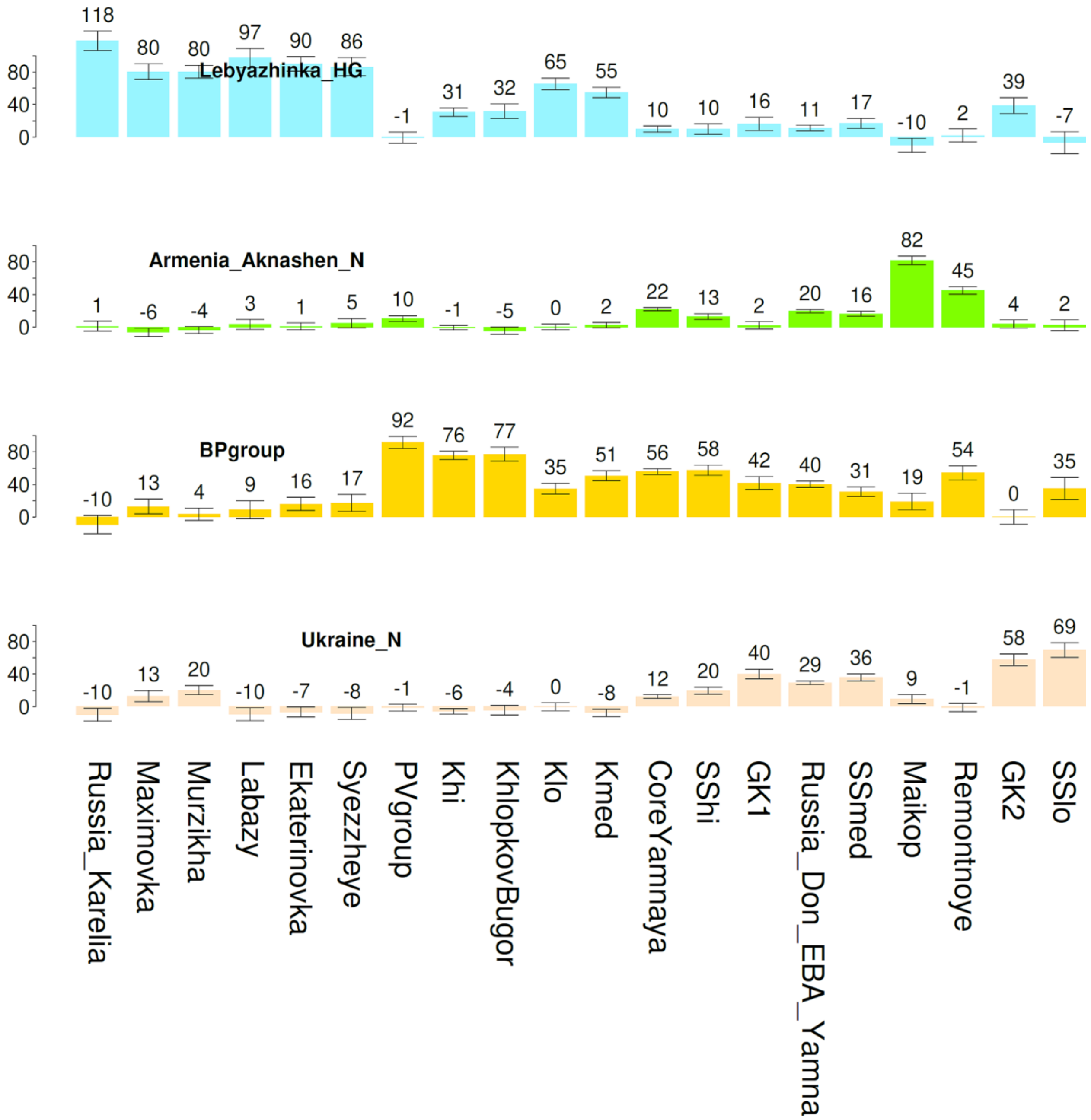

**Fig. S 11 A 4-way model for the entire Dnipro-Don-Volga-Caucasus region.** Proportions shown are those of Table S 29. Error bars show ± 1 standard error.

Finally, we plot in Fig. S 11 the proportions of Table S 29 which visually demonstrates all our inferences. The contrast between the Don-Dnipro and Volga in terms of Ukraine_N vs. EHG ancestry; the presence of Aknashen ancestry in Maikop, Remontnoye, the Serednii Stih, and the Yamnaya, but not in Golubaya Krinitsa or the Volga; and the contribution of a population like BPgroup of ancestry in populations across the entire region except the far north of eastern Europe (Karelia) and starting from at least the mid-6$^{th}$ millennium date of the Golubaya Krinitsa individuals, one of which has it (GK1) while the other one does not (GK2).

**Evidence from the 4th millennium BCE**

The scenario for the formation of the core Yamnaya developed in this note predicts that a "pre-Yamnaya" population was formed ~4000BCE and gave rise to the sampled Yamnaya of the late 4th millennium BCE when the latter acquired the high mobility characteristic of that culture and appeared on the archaeological record as the Yamnaya archaeological horizon.

Here we examine 4th millennium BCE individuals understand see if any of them are plausible early representatives of that nascent Yamnaya population. We show their distribution in Fig. S 12 and a PCA (using the same populations to form the axes as in Fig. S 1) in Fig. S 13. As can be seen, none of these individuals correspond to the core Yamnaya population, a conclusion that we test specifically using qpWave (Table S 30). It is clear that during the 4th millennium BCE there is evidence for the presence of populations distinct from the core Yamnaya in the Middle Volga (at Maximovka and Chekalino IV), the Upper Volga, in the North Caucasus and steppes north of it, and in different sites of the Dnipro area (or west of it).

We cannot, of course, be certain that the sampled individuals in each location did not live side by side with the elusive Pre-Yamnaya population. The fact, however, that these individuals do *not* appear to be genetically related to the core Yamnaya should reduce any prior belief in the emergence of the Yamnaya in the locations of Fig. S 12. Future research must, as a priority, investigate the gaps of the map of Fig. S 12, and especially additional sites of the Serednii Stih archaeological culture. It is not guaranteed that the location of Yamnaya emergence will be recoverable, especially as this may be geographically constrained and archaeologically unremarkable: the corollary of the high mobility of the Yamnaya is that they quickly dispersed from the cradle of their culture. However, even if the geographical locus of their formation is never found positively, comprehensive sampling of both the Serednii Stih territory and the steppe as a whole may still continue to do add—through exclusion of alternatives—to theories of Yamnaya emergence.

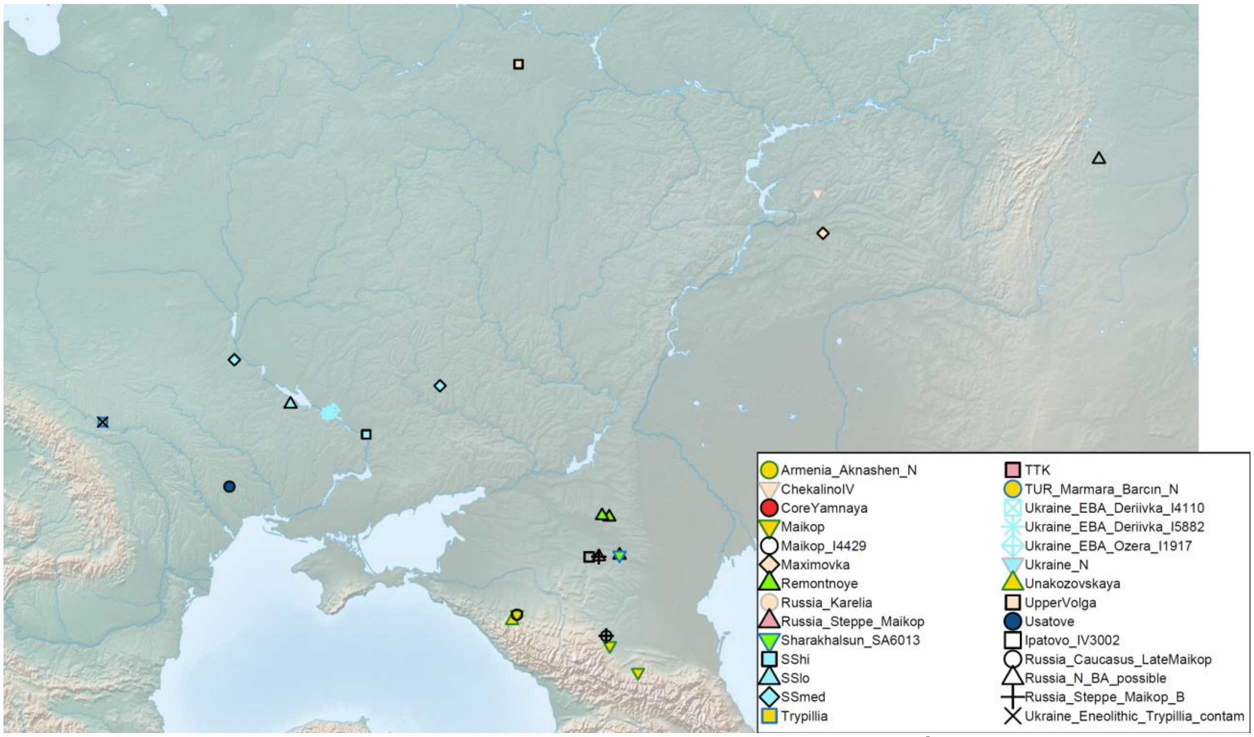

**Fig. S 12 Distribution of non-Yamnaya/Afanasievo individuals from the 4th millennium BCE**

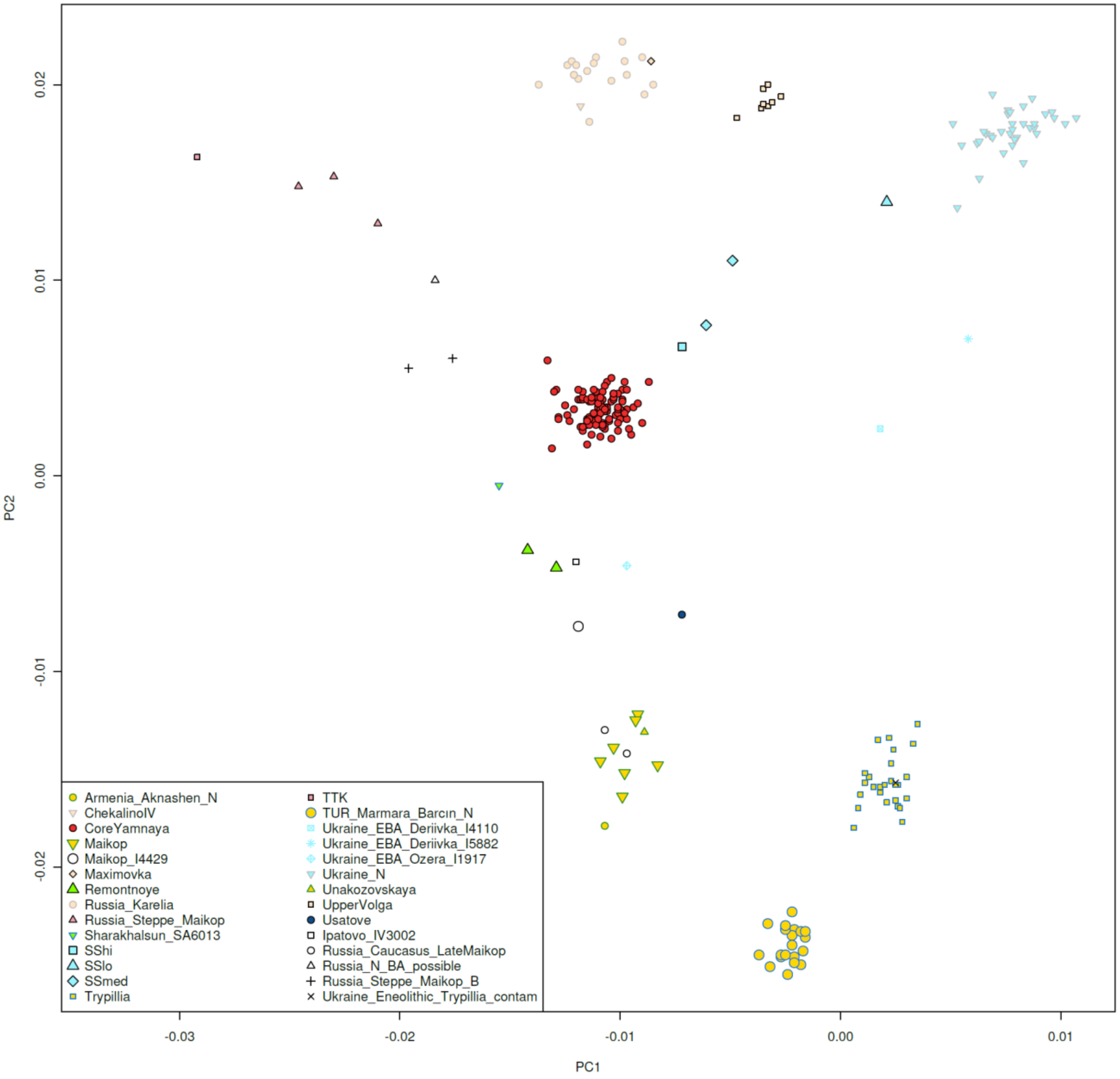

**Fig. S 13 PCA of non-Yamnaya/Afanasievo 4th millennium BCE individuals.** Individuals were selected to be from 4000-3000BCE, from Russia or Ukraine, not to be labeled as Yamnaya or Afanasievo, and to be west of 70E longitude (to avoid plotting Siberian individuals from the easternmost parts of the Russian Federation that are not relevant to our question).

| | | | | | | | |
|---|---|---|---|---|---|---|---|
| I6303 | F | ChekalinoIV | Russia | 53.9 | 50.9 | 5528 | 1.02E-19 |
| IV3002 | M | Ipatovo_IV3002 | Russia | 45.7 | 42.9 | 5201 | 8.60E-69 |
| I4429 | M | Maikop_I4429 | Russia | 44.4 | 40.4 | 5300 | 1.95E-186 |
| OSS001 | F | Maikop | Russia | 43.1 | 44.6 | 5582 | 2.89E-125 |
| OSS002.B0101 | M | Maikop | Russia | 43.1 | 44.6 | 5711 | 7.80E-29 |
| I1720_wNonUDG | M | Maikop | Russia | 43.7 | 43.6 | 5300 | 1.68E-118 |
| I6272 | M | Maikop | Russia | 44.4 | 40.4 | 5040 | 1.32E-24 |
| I6266 | M | Maikop | Russia | 44.4 | 40.4 | 5385 | 0.00E+00 |
| I6267 | F | Maikop | Russia | 44.4 | 40.4 | 5410 | 6.67E-21 |
| I6268 | M | Maikop | Russia | 44.4 | 40.4 | 5550 | 1.81E-131 |
| I8446 | M | Maximovka | Russia | 53.0 | 51.1 | 5792 | 9.30E-28 |
| I28682 | F | Remontnoye | Russia | 46.6 | 43.6 | 5632 | 1.96E-258 |
| I28683 | M | Remontnoye | Russia | 46.6 | 43.4 | 5913 | 3.78E-41 |
| MK5001 | M | Russia_Caucasus_LateMaikop_rel.MK5004 | Russia | 43.9 | 43.5 | 5150 | 2.27E-134 |
| MK5004 | M | Russia_Caucasus_LateMaikop | Russia | 43.9 | 43.5 | 5150 | **9.58E-02** |
| MK5008 | M | Russia_Caucasus_LateMaikop | Russia | 43.9 | 43.5 | 5187 | 5.76E-138 |
| I8742 | M | Russia_Eneolithic_1d.rel.I6904 | Russia | 53.0 | 51.1 | 5617 | 1.40E-223 |
| I8447 | F | Russia_Eneolithic_1d.rel.I8446 | Russia | 53.0 | 51.1 | 5820 | 2.70E-04 |
| SIJ003 | F | Russia_LateMaikop_1d.rel.I11131.I11133_dup.SIJ003 | Russia | 45.1 | 39.9 | 5164 | 2.93E-272 |
| SIJ002 | M | Russia_LateMaikop_1d.rel.I11132.I11133_dup.SIJ002 | Russia | 45.1 | 39.9 | 5164 | 1.32E-133 |
| SA6002 | F | Russia_LateMaikop_dup.SA6002 | Russia | 45.1 | 39.9 | 5126 | 1.08E-55 |
| I10632_d | M | Russia_N_BA_possible | Russia | 54.6 | 60.8 | 5350 | 1.15E-165 |
| MK5005.C0101 | F | Russia_Steppe_Maikop_B | Russia | 43.9 | 43.5 | 5450 | 9.76E-109 |
| AY2001 | F | Russia_Steppe_Maikop_B | Russia | 45.7 | 43.3 | 5397 | 5.94E-63 |
| AY2003 | F | Russia_Steppe_Maikop | Russia | 45.7 | 43.3 | 5452 | 4.78E-93 |
| SA6004 | M | Russia_Steppe_Maikop | Russia | 45.7 | 44.0 | 5159 | 4.80E-41 |
| SA6001 | F | Russia_Steppe_Maikop | Russia | 45.7 | 44.0 | 5397 | 1.54E-52 |
| SA6013 | M | Sharakhalsun_SA6013 | Russia | 45.7 | 44.0 | 5167 | 8.26E-109 |
| I1924 | F | SShi | Ukraine | 48.4 | 35.1 | 5828 | 6.56E-11 |
| I1424 | M | SSlo | Ukraine | 49.1 | 32.5 | 5658 | 5.77E-289 |
| I6558 | M | SSmed | Ukraine | 49.5 | 37.7 | 5456 | 5.71E-109 |
| I7585_enhanced | F | SSmed | Ukraine | 50.1 | 30.5 | 5935 | 5.85E-278 |
| I3151_enhanced | M | Trypillia | Ukraine | 48.7 | 25.9 | 5528 | 1.04E-100 |
| I13064 | M | Trypillia | Ukraine | 48.7 | 25.9 | 5533 | 3.08E-128 |
| VERT015_wNonUDG.SG | F | Trypillia | Ukraine | 48.7 | 25.9 | 5589 | 1.09E-40 |
| VERT103B_wNonUDG.SG | M | Trypillia | Ukraine | 48.7 | 25.9 | 5600 | 1.39E-51 |
| VERT118_wNonUDG.SG | F | Trypillia | Ukraine | 48.7 | 25.9 | 5600 | 1.44E-282 |
| VERT111_wNonUDG.SG | F | Trypillia | Ukraine | 48.7 | 25.9 | 5600 | 2.06E-293 |
| VERT115_wNonUDG.SG | F | Trypillia | Ukraine | 48.7 | 25.9 | 5600 | 2.13E-213 |
| VERT105B_wNonUDG.SG | M | Trypillia | Ukraine | 48.7 | 25.9 | 5600 | 2.96E-100 |
| VERT033_wNonUDG.SG | M | Trypillia | Ukraine | 48.7 | 25.9 | 5600 | 3.45E-260 |
| VERT117_wNonUDG.SG | M | Trypillia | Ukraine | 48.7 | 25.9 | 5600 | 6.23E-237 |
| VERT030_wNonUDG.SG | M | Trypillia | Ukraine | 48.7 | 25.9 | 5600 | 6.83E-119 |
| VERT107_wNonUDG.SG | M | Trypillia | Ukraine | 48.7 | 25.9 | 5600 | 7.75E-22 |
| VERT028_wNonUDG.SG | M | Trypillia | Ukraine | 48.7 | 25.9 | 5600 | 8.64E-13 |
| VERT029_wNonUDG.SG | M | Trypillia | Ukraine | 48.7 | 25.9 | 5600 | 9.53E-109 |
| I7584 | F | Trypillia | Ukraine | 48.7 | 25.9 | 5607 | 5.36E-143 |
| I2111_enhanced | M | Trypillia | Ukraine | 48.7 | 25.9 | 5622 | 9.37E-166 |
| I1929 | F | Trypillia | Ukraine | 48.7 | 25.9 | 5628 | 8.51E-264 |
| I7586 | F | Trypillia | Ukraine | 48.7 | 25.9 | 5639 | 6.69E-04 |
| VERT104B_wNonUDG.SG | M | Trypillia | Ukraine | 48.7 | 25.9 | 5642 | 5.50E-43 |
| VERT031_wNonUDG.SG | M | Trypillia | Ukraine | 48.7 | 25.9 | 5646 | 2.04E-303 |
| VERT100B_wNonUDG.SG | M | Trypillia | Ukraine | 48.7 | 25.9 | 5646 | 3.57E-114 |
| I1926_enhanced | M | Trypillia | Ukraine | 48.7 | 25.9 | 5649 | 9.68E-69 |
| VERT106C_wNonUDG.SG | M | Trypillia | Ukraine | 48.7 | 25.9 | 5669 | 7.76E-288 |
| I7923 | M | Trypillia | Ukraine | 48.7 | 25.9 | 5670 | 1.50E-128 |
| VERT035_wNonUDG.SG | F | Trypillia | Ukraine | 48.7 | 25.9 | 5671 | 1.06e-314 |
| I2110 | M | Trypillia | Ukraine | 48.7 | 25.9 | 5697 | 1.97E-228 |
| I4110 | F | Ukraine_EBA_Deriivka_I4110 | Ukraine | 48.9 | 33.8 | 5394 | 3.75E-87 |
| I5882_enhanced | F | Ukraine_EBA_Deriivka_I5882 | Ukraine | 48.9 | 33.8 | 5005 | 2.75E-234 |
| I1917 | F | Ukraine_EBA_Ozera_I1917 | Ukraine | 49.0 | 34.0 | 4955 | 3.15E-110 |
| I2109 | M | Ukraine_Eneolithic_Trypillia_contam | Ukraine | 48.7 | 25.9 | 5589 | 5.39E-120 |
| I1717 | F | Unakozovskaya | Russia | 44.3 | 40.2 | 5900 | 4.14E-288 |
| I8437 | F | UpperVolga | Russia | 56.8 | 40.4 | 5192 | 8.70E-44 |
| I12494 | M | UpperVolga | Russia | 56.8 | 40.4 | 5409 | 1.80E-118 |
| I12964 | M | UpperVolga | Russia | 56.8 | 40.4 | 5192 | 1.66E-70 |
| I12962 | F | UpperVolga | Russia | 56.8 | 40.4 | 5648 | 2.45E-07 |
| I8408 | M | UpperVolga | Russia | 56.8 | 40.4 | 5650 | 1.72E-296 |
| I8404 | M | UpperVolga | Russia | 56.8 | 40.4 | 5820 | 7.52E-11 |
| I8419 | F | UpperVolga | Russia | 56.8 | 40.4 | 5950 | 5.48E-53 |
| I12500 | F | UpperVolga | Russia | 56.8 | 40.4 | 5950 | 6.07E-94 |
| I7929 | M | Usatove | Ukraine | 47.3 | 30.3 | 5628 | 2.35E-20 |

**Table S 30 qpWave modeling of non-Yamnaya/Afanasievo 4th millennium BCE individuals.**

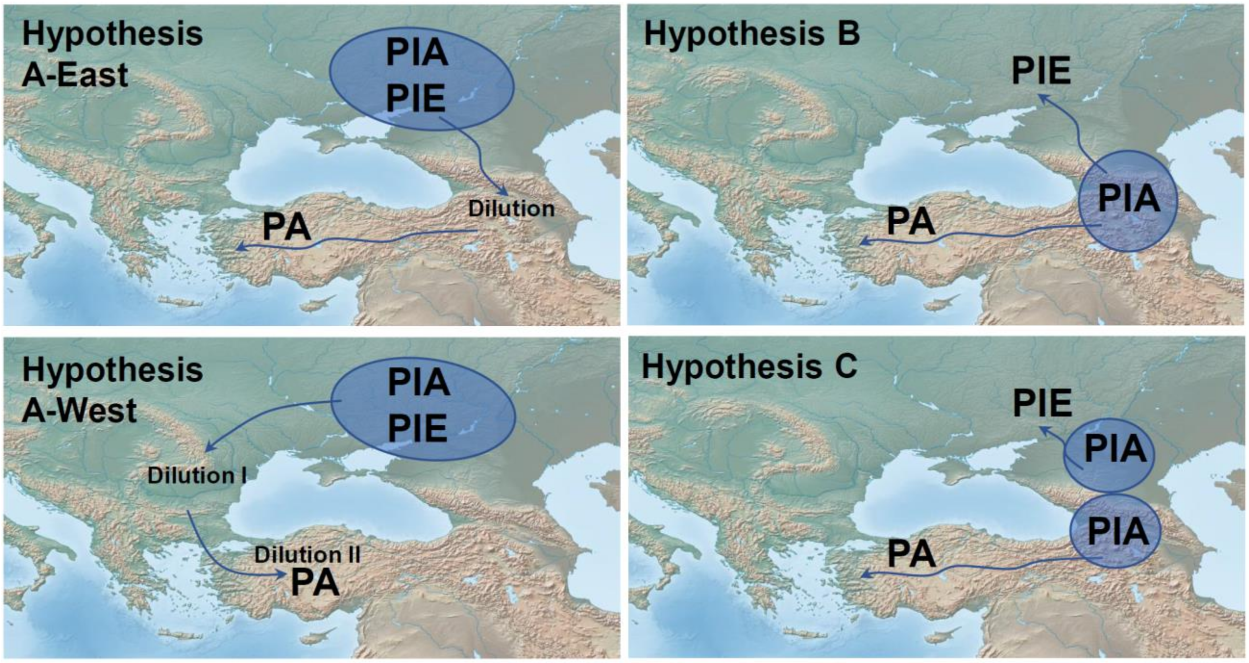

**Fig. S 14 Competing hypotheses of Indo-Anatolian and Indo-European origins.** Hypothesis A: Proto-Indo-Anatolian was a steppe language; variant A-East: Anatolian languages spread from the steppe via the Caucasus; variant A-West: Anatolian languages spread from the steppe via the Balkans. Hypothesis B: Proto-Indo-Anatolian was a language of the Caucasus-West Asian highlands. Hypothesis C: Proto-Indo-Anatolian had been a common language understood by steppe and Caucasus neighbors, regardless of its deeper origins; whatever admixtures were taking place between south and north were incidental and not instrumental to the spread of the language as the Proto-Indo-Anatolian language was used by people of both "steppe" and "West Asian" genetic background.

#### Links between Anatolian and Indo-European speakers

We have arrived at a synthesis of Yamnaya origins which ties their ancestry to both the "north" (via SShi and BPgroup, themselves mixtures of earlier "north"-"south" contact) and the "south" (via Maikop/Aknashen in the Caucasus). In this section we try to link the Yamnaya, who presumably spoke an Indo-European language and may have spoken the Proto-Indo-European ancestral language, with ancient Anatolians where languages of the Anatolian branch (such as Hittite and Luwian) of the broader Indo-Anatolian family were spoken.

##### Leading hypotheses of Proto-Indo-Anatolian origins

In a recent paper[4] we argued that two hypotheses remain for the origin of Proto-Indo-Anatolian languages (the proto-language of which Anatolian languages and Indo-European languages were twin daughters) (Fig. S 14). We identified the Yamnaya as speakers of Proto-Indo-European and argued (in Hypothesis B) that the Proto-Indo-Anatolians were a population of the highlands of West Asia and the Caucasus as:

1. Migrations from this area transformed Anatolia during the Chalcolithic and Bronze Age period
2. Migrations from this area contributed at least two pulses of ancestry to the Yamnaya
3. Anatolian populations lacked Eastern hunter-gatherer ancestry that would tie them to steppe populations

The competing hypothesis (Hypothesis A) proposes that Proto-Indo-Anatolians were a steppe population. However, making a connection between from the steppe into Anatolia has proven elusive[4,20,21,49,50]

One possibility ("Hypothesis A-West") is that steppe migrants came from Southeastern Europe into Anatolia.[4] If they had shed their steppe ancestry by admixing with local Southeastern European farmers then perhaps their entry into Anatolia would not be detectible. However, we know that the farmers of Southeastern Europe were of largely Anatolian Neolithic ancestry with local Balkan hunter-gatherer admixture.[11] Admixture from that area would then presumably introduce back to Anatolia some of this Anatolian Neolithic ancestry as well as some Balkan hunter-gatherer ancestry. Yet, in Anatolia during the Chalcolithic and Bronze Age we see a *decrease* of Anatolian Neolithic ancestry and a lack of Balkan hunter-gatherer ancestry.[4]

Another possibility ("Hypothesis A-East") is that steppe migrants came via the Caucasus into Anatolia. This hypothesis also requires some shedding of steppe ancestry to have occurred (to account for its lack or paucity in Chalcolithic and Bronze age Anatolia). However, it has the advantage (compared to "Hypothesis A-West") of going "with the grain" of the transformation of Chalcolithic and Bronze Age Anatolia. Any Proto-Anatolian-speaking migrants from the east would indeed bring some ancestry from the eastern highlands of West Asia into the west of Anatolia even if—under this hypothesis— they had shed most of their steppe ancestry while adopting the language of their steppe (linguistic) forebears.

All three hypotheses remain viable.

Hypothesis B harmonizes with all known facts and the results of our reconstruction of Yamnaya origins strengthen it, as the Yamnaya do indeed have ancestry from the south: both early ones via their BPgroup ancestors which experienced gene flow from the Caucasus and contributing to the Serednii Stih and Volga clines; but, also later ones via the migration of Remontnoye-related people (who also had Maikop/Aknashen ancestry). What was only indistinct before (the CHG ancestry in the Eneolithic steppe and the extra Anatolian-Levantine ancestry in the Yamnaya[4]) has now come into better focus.

Hypothesis A-West is not the focus of this study and has the difficulties we outlined above. However, we think that it must continue to be explored – provided that a mechanism can be found to explain how language spread may have been effected despite twin dilutions *en route* from the steppe to Anatolia: first in the Balkans—to explain the paucity of EHG ancestry in Anatolia—and second in Anatolia itself, to explain the decrease of Anatolian ancestry there during the crucial Chalcolithic and Bronze Age periods.

Hypothesis A-East has fewer difficulties from the genetic point of view, as it requires only one dilution of steppe ancestry: between the steppe and the highlands of West Asia. But, such a dilution[5] did indeed take place as we see the Maikop and Armenian Chalcolithic had limited steppe ancestry (Table S 8; Table S 11). Thus, Hypothesis A-East must only explain why despite the numerical disadvantage of their ancestry the language of the steppe migrants prevailed – a difficulty not faced by Hypothesis B.

The strength of Hypothesis A-West is that the historical distribution of Anatolian languages is western.[51] Thus, both Hypothesis A-East and Hypothesis B must explain why the Proto-Indo-Anatolian languages of the eastern highlands—whether they were native there or the result of linguistic conversion of natives by steppe migrants—disappeared. This is a real difficulty with the eastern hypotheses, but it can be argued that movements as early as the halving of CHG ancestry between the Aknashen and Masis Blur Neolithic[4] in the 6$^{th}$ millennium BCE, the expansion of the Kura-Araxes culture of the Early Bronze Age[4] that represented a disruption of the genetic continuity compared to the earlier Chalcolithic, or even the Levantine influence in the ancestry of the Urartians of the eastern highlands[52] may all have combined to remove traces of the Proto-Indo-Anatolian presence there.

After all, the descendants of the Yamnaya themselves, to whose origins we have devoted the previous sections, were themselves largely displaced during the 3rd millennium BCE from the steppe only to be replaced by the descendants of their Corded Ware relatives moving eastwards,[4,22,48] the first of multiple genetic and presumably linguistic turnovers on the steppe in the ensuing millennia. Similarly, there is no necessity that descendants of a proto-language including the speakers of those languages (be it Proto-Indo-European on the steppe or Proto-Indo-Anatolian in the Caucasus and West Asian highlands) should persist in their homeland. To summarize: the existence of non-Indo-Anatolian languages in the east of Anatolia and the Caucasus does not disprove either Hypothesis B or Hypothesis A-East, although it must be reckoned as one of the strengths of Hypothesis A-West.

If either variant of Hypothesis A is right, then the Proto-Indo-Anatolian homeland could either be in the Don-Dnipro area or in the Don-Volga interfluvial (which furnishes one component in the ancestry of BPgroup) or even in Siberia/Central Asia (as BPgroup has ~25% ancestry from such a source; Table S 12) The fact that the core Yamnaya have substantial SShi-related ancestry may argue in favor of Don-Dnipro; yet the ancestry of the Don-Dnipro Serednii Stih itself was that of local hunter-gatherers admixing with CLV cline easterners and forming the Don-Dnipro cline. The Don-Volga interfluvial would thus be an alternative option that would also be geographically more proximate to the south where Don-Volga ancestry is found in the Armenian Chalcolithic (
Table S 11) and as sporadic Y-chromosomes of the R-V1636 clade in West Asia.[4,5,21] Thus, under Hypothesis A, Proto-Indo-Anatolian was the language spoken in the North Caucasus piedmont and Lower Volga by people of mixed CHG-EHG-Central Asian/Siberian ancestry and Proto-Indo-European was the language spoken by people of Serednii Stih descendants mixing with migrants from this area. Other migrants headed south (Hypothesis A-East), admixing into the Maikop and ancient Armenia; admixture with locals shed the traces of their steppe origin before they reached Anatolia. Alternatively, under Hypothesis A-West, Proto-Indo-Anatolian may have been spoken either in the Don-Volga or Don-Dnipro areas. Migrants from the steppe reached the Balkans where they admixed with locals in a first dilution event, and then reached Anatolia where they admixed with locals in a second dilution event.

If, on the other hand, Hypothesis B is right, then the Proto-Indo-Anatolian homeland could either be the language of early pre-agricultural people of the Caucasus that admixed with hunter-gatherers to form the Don-Dnipro and Volga clines or, alternatively, of the agricultural Aknashen/Maikop people of the Caucasus who contributed ancestry to the steppe via transitional populations like Remontnoye. The second possibility seems more in line with the evidence for a relatively shallow time of separation of Anatolian and Indo-European languages.[53] However, it has been argued that the paucity of shared agricultural vocabulary in Anatolian and core Indo-European languages speaks in favor of the scenario that the Proto-Indo-Anatolians were pastoralists that did not practice agriculture (thus unlike the people of the Maikop and Aknashen).[54] This evidence would argue in favor of either Hypothesis A or the version of Hypothesis B in which it is early (pre-agricultural) gene flow from the south bringing Indo-Anatolian languages to the steppe. However, the paucity of shared agricultural vocabulary could also be explained by the hiatus in the practice of agriculture in the steppe between the arrival of Indo-Anatolian languages and the dispersal of Indo-European languages with the Yamnaya.

To conclude: good arguments have been brought forth for all hypotheses of the deep origins of Indo-Anatolian and Indo-European languages. To these we may also add another “Hypothesis C” (Fig. S 14): that the existence of a common genetic component mediating the spread of language into both the steppe and Anatolia—however diluted or substantial it may be—is uncoupled from the spread of the speakers of Proto-Indo-Anatolian.

Under this scenario, at least some of the people of the steppe in the Lower Don-Lower Volga interfluvial and some of their southern neighbors in the North Caucasus shared a language for communicating with each other even though they belonged to different genetic backgrounds. This language may have been originally spoken in either the north or the south, but, early on, it came to be known to people of both north and south, perhaps long before its breakdown into Anatolian and Indo-European branches. Genetic admixture between the Caucasus and the steppe—at different time scales and from different sources—did occur, and is a direct evidence of contact, supplementing that of archaeology.

But, under Hypothesis C, migration followed by admixture was incidental and not the main medium of language spread: the language was already in use by people of the North Caucasus-steppe transition zone regardless of genetic background. Migrations from the Don-Volga interfluvial, associated with "steppe" genetics, then brought a variant of this language to the Don-Dnipro area, setting the stage for the emergence and expansion of Proto-Indo-Europeans; and migrations from the Caucasus and West Asian highlands, associated with "Caucasus" genetics, brought another variant of this language into Anatolia. It may turn out that genetics, which has made a convincing case for the transmission of language by migration in Europe, West Asia, Central and South Asia,[2,4,9,20,22,49] may have little to say about the earliest history of Proto-Indo-Anatolian. At the very least, it has helped track the speakers of Indo-Anatolian languages to the slopes of the Caucasus and the rolling steppes beyond them: whether the language was spoken by one, or many, of the diverse people living there may be a question for other disciplines to answer.

#### Modeling Anatolian Chalcolithic and Bronze Age populations

We re-assessed the Anatolian archaeogenetic record[4,20,21,49] of the Chalcolithic and Bronze Age periods by studying the ancestry of the following Test populations:

**TestA:** Armenia_Aknashen_N[4], Armenia_C[5], Azerbaijan_N[21,24], Azerbaijan_C[21], Armenia_KuraAraxes_EBA[4,8], Armenia_MasisBlur_N[4], Armenia_MLBIA[4,5], Israel_C[27], Maikop[8], TUR_Aegean_BA[4], TUR_BlackSea_BA[4], TUR_BlackSea_ChL[21], TUR_C_BA[20], TUR_C_ChL[21], TUR_E_BA[21], TUR_E_ChL[21], TUR_Hatay_BA[21], TUR_Hatay_ChL[21], TUR_Marmara_ChL[4,5], TUR_Med_BA[49], TUR_SE_BA[4,21], TUR_SE_ChL[4], Unakozovskaya (ref.[8] and this study)

We grouped Anatolian samples by region (C: Central, E: East, Med: Mediterranean (SW Anatolia), Aegean: Aegean, Black Sea: Black Sea (NE Anatolia), Marmara: Marmara (NW Anatolia), SE: Southeast, Hatay: Hatay Province) and by Chalcolithic (ChL) or Bronze Age (BA) period, and also included comparative samples from the rest of West Asia.

We use the following populations as candidate sources, which includes sources of the steppe, southeastern Europe, and West Asia.

**SourcesA:** Armenia_Aknashen_N, Armenia_C, Armenia_KuraAraxes_EBA, Armenia_MasisBlur_N, Azerbaijan_C, Azerbaijan_N, Bulgaria_C[11], BPgroup, CoreYamnaya, Iran_GanjDareh_N, Maikop, PVgroup, Remontnoye, Russia_Caucasus_Eneolithic, SShi, SSo, SSmed, Trypillia, Turkey_N, TUR_C_Boncuklu_PPN[7], TUR_C_AşıklıHöyük_PPN[55], TUR_C_Çatalhöyük_N[55], TUR_SE_Çayönü_PPN[56], TUR_C_Musular_PPN[57], Israel_C, Levant_N[5], Armenia_KuraAraxes_EBA,

We used the following set of Right outgroup populations:

**BaseA:** OldAfrica, CHG, Iran_GanjDareh_N, Italy_Villabruna, Russia_AfontovaGora3, Russia_Sidelkino.SG, TUR_Marmara_Barcın_N, TUR_C_Boncuklu_PPN, TUR_C_Çatalhöyük_N, Natufian

This set includes the same populations as the Base set used in the analysis of steppe populations, but since our Test set consists of West Asian populations, we have added Levantine and Anatolian outgroups to better distinguish ancestry within this region. Moreover, we added different Anatolian Neolithic populations (NW Anatolian Neolithic was included in Base, but here we also include TUR_C_Boncuklu_PPN, TUR_C_Çatalhöyük_N in BaseA), so as to treat them symmetrically, i.e., not assume that any of them represent the Neolithic ancestry prior to the Chalcolithic and Bronze Age transformation of Anatolia.

#### Which populations are simple clades of the SourcesA?

First, we show populations that can be modeled as simple clades of one of the sources (Table S 31).

| Test | Source | P-value |
|---|---|---|
| Armenia_KuraAraxes_EBA | Maikop | 0.058 |
| Azerbaijan_N | Azerbaijan_C | 0.670 |
| Azerbaijan_C | Azerbaijan_N | 0.670 |
| Maikop | Armenia_KuraAraxes_EBA | 0.058 |
| TUR_BlackSea_ChL | Armenia_MasisBlur_N | 0.517 |
| TUR_E_BA | Armenia_MasisBlur_N | 0.423 |
| TUR_E_ChL | Armenia_MasisBlur_N | 0.311 |

**Table S 31 Populations that can be modeled as simple clades (*N*=1) with one of the sources**

The Kura Araxes population from Armenia is a clade with Maikop with a low p-value (p=0.058). The previously observed (Table S 2) clade between Chalcolithic and Neolithic Azerbaijan is observed also with the BaseA set of outgroup populations. The most interesting observation is that the Bronze Age and Chalcolithic population from Eastern Turkey (which is adjacent to Armenia) is a clade with the Neolithic population of Masis Blur.

It is unclear whether the "MasisBlur"-like population of Eastern Turkey represents a Neolithic stratum there, as we lack Neolithic-era samples from the region. What we can say is that by the Late Chalcolithic (or the early part of the 4$^{th}$ millennium BCE) it stretched across Eastern Anatolia where it continued to the Bronze Age. Therefore, our Armenian Masis Blur source should be understood as encompassing a source of ancestry not localized to present-day Armenia but encompassing eastern parts of Turkey as well.

We can model Masis Blur in the framework used here as a mixture of Aknashen Neolithic and 30.6±4.8% TUR_Marmara_Barcın_N (p=0.36) or 37.2±4.7% TUR_C_Çatalhöyük_N (p=0.42). However, we can also model Aknashen Neolithic as 69.1±4.5% Masis Blur Neolithic and 30.9±4.5% CHG ancestry. The only feasible for both Aknashen and Masis Blur Neolithic populations that does not involve them mutually modeling each other includes TUR_SE_Çayönü_PPN and CHG ancestry. Masis Blur has 13.7±3.8% CHG (p=0.13) and Aknashen has 42.0±3.7% CHG (p=0.09) ancestry according to this model. It is unclear[4] if this represents population change during the 6$^{th}$ millennium BCE in Armenia or pre-existing population structure. That Masis Blur and Aknashen share the majority of their ancestry could be consistent with being part of a genetically variable population that shared a language, although their genetic differentiation could also be consistent with language shift in the area.

If Proto-Indo-Anatolian was spoken in the highlands of West Asia (as Hypothesis B postulates), then both Anatolian Neolithic populations may have spoken either the ancestral language of Proto-Indo-Anatolian or another of the many languages spoken in the region. Alternatively (as Hypothesis A-East postulates) people of ancestry from Armenia-Eastern Turkey may have received their language—but not much genetic ancestry—from the north and may have spread it westward into central/western Anatolia where it was historically recorded.

##### Which populations are 2-way admixtures?

Since only 7 populations can be modeled as simple clades of the sources, and none of these are from central and western regions of Anatolia where ancient Anatolian speakers must have been present, we next looked at 2-way admixture models. We discuss these below.

First, we look at Chalcolithic NW Anatolia (TUR_Marmara_ChL) (Table S 32). All feasible models include ancestry from the east. Of the models that have NW Anatolian as a source (the NW Anatolian Neolithic population that preceded the Chalcolithic in the Marmara region), the amount of eastern ancestry is substantial (~1/2 or more). This includes a model with Maikop as a source. This is likely ahistorical given that this is an Early Bronze Age source and predicts a large component of ancestry from the North Caucasus and faraway NW Anatolia. However, it does underscore that models in which the source has a heavily diluted component of steppe ancestry may be feasible (as Maikop has a small proportion of Steppe ancestry).

| A | B | P-value | A | B | S.E |
|---|---|---|---|---|---|
| Armenia_Aknashen_N | Bulgaria_C | 0.231 | 49.4% | 50.6% | 2.6% |
| Armenia_Aknashen_N | TUR_C_Boncuklu_PPN | 0.944 | 52.8% | 47.2% | 2.5% |
| Armenia_Aknashen_N | TUR_C_Çatalhöyük_N | 0.065 | 46.5% | 53.5% | 3.0% |
| Armenia_Aknashen_N | TUR_C_Musular_PPN | 0.394 | 41.6% | 58.4% | 3.4% |
| Armenia_Aknashen_N | TUR_Marmara_Barcın_N | 0.476 | 53.0% | 47.0% | 2.8% |
| Armenia_C | TUR_C_AşıklıHöyük_PPN | 0.061 | 53.8% | 46.2% | 4.5% |
| Armenia_KuraAraxes_EBA | TUR_C_Musular_PPN | 0.410 | 38.1% | 61.9% | 2.9% |
| Armenia_KuraAraxes_EBA | TUR_Marmara_Barcın_N | 0.090 | 49.6% | 50.4% | 1.9% |
| Armenia_MasisBlur_N | Bulgaria_C | 0.082 | 75.1% | 24.9% | 4.9% |
| Armenia_MasisBlur_N | Trypillia | 0.132 | 73.9% | 26.1% | 4.9% |
| Armenia_MasisBlur_N | TUR_C_AşıklıHöyük_PPN | 0.112 | 76.2% | 23.8% | 7.4% |
| Armenia_MasisBlur_N | TUR_C_Boncuklu_PPN | 0.242 | 76.3% | 23.7% | 4.3% |
| Armenia_MasisBlur_N | TUR_C_Musular_PPN | 0.057 | 69.7% | 30.3% | 6.3% |
| Azerbaijan_C | TUR_C_AşıklıHöyük_PPN | 0.207 | 56.0% | 44.0% | 4.8% |
| Azerbaijan_C | TUR_C_Boncuklu_PPN | 0.202 | 58.7% | 41.3% | 3.2% |
| Azerbaijan_C | TUR_C_Musular_PPN | 0.107 | 46.2% | 53.8% | 4.1% |
| Azerbaijan_N | TUR_C_AşıklıHöyük_PPN | 0.332 | 61.9% | 38.1% | 4.6% |
| Maikop | TUR_C_AşıklıHöyük_PPN | 0.071 | 45.9% | 54.1% | 4.2% |
| Maikop | TUR_C_Çatalhöyük_N | 0.156 | 41.8% | 58.2% | 2.2% |
| Maikop | TUR_C_Musular_PPN | 0.603 | 37.0% | 63.0% | 2.9% |
| Maikop | TUR_Marmara_Barcın_N | 0.081 | 48.0% | 52.0% | 1.9% |

| | | | | | |
|---|---|---|---|---|---|
| TUR_C_Musular_PPN | Unakozovskaya | 0.187 | 69.1% | 30.9% | 2.5% |

**Table S 32 Feasible models for TUR_Marmara_ChL**

Next, we look at Chalcolithic Central Anatolia. The amount of eastern ancestry is even higher here (~70-80%) compared to the Marmara region, suggesting major discontinuity between the Neolithic and Chalcolithic periods mediated by migrants from the east.

| A | B | P-value | A | B | S.E |
|---|---|---|---|---|---|
| Armenia_Aknashen_N | TUR_C_Çatalhöyük_N | 0.170 | 49.9% | 50.1% | 2.8% |
| Armenia_KuraAraxes_EBA | TUR_C_Çatalhöyük_N | 0.172 | 46.0% | 54.0% | 1.8% |
| Armenia_MasisBlur_N | Bulgaria_C | 0.650 | 80.2% | 19.8% | 4.6% |
| Armenia_MasisBlur_N | Israel_C | 0.050 | 85.2% | 14.8% | 8.1% |
| Armenia_MasisBlur_N | Levant_N | 0.052 | 89.8% | 10.2% | 5.4% |
| Armenia_MasisBlur_N | Trypillia | 0.606 | 80.2% | 19.8% | 4.6% |
| Armenia_MasisBlur_N | TUR_C_AşıklıHöyük_PPN | 0.128 | 87.5% | 12.5% | 7.2% |
| Armenia_MasisBlur_N | TUR_C_Boncuklu_PPN | 0.609 | 82.3% | 17.7% | 4.3% |
| Armenia_MasisBlur_N | TUR_C_Çatalhöyük_N | 0.517 | 80.2% | 19.8% | 5.7% |
| Armenia_MasisBlur_N | TUR_C_Musular_PPN | 0.368 | 77.6% | 22.4% | 6.3% |
| Armenia_MasisBlur_N | TUR_Marmara_Barcın_N | 0.273 | 84.5% | 15.5% | 5.2% |
| Azerbaijan_C | TUR_C_AşıklıHöyük_PPN | 0.096 | 62.9% | 37.1% | 4.8% |
| Azerbaijan_C | TUR_C_Boncuklu_PPN | 0.372 | 63.4% | 36.6% | 2.9% |
| Azerbaijan_C | TUR_C_Çatalhöyük_N | 0.412 | 56.2% | 43.8% | 3.3% |
| Azerbaijan_C | TUR_C_Musular_PPN | 0.416 | 51.9% | 48.1% | 3.8% |
| Azerbaijan_C | TUR_Marmara_Barcın_N | 0.156 | 63.1% | 36.9% | 3.3% |
| Azerbaijan_N | TUR_C_AşıklıHöyük_PPN | 0.219 | 68.7% | 31.3% | 4.0% |
| Azerbaijan_N | TUR_C_Boncuklu_PPN | 0.284 | 70.8% | 29.2% | 2.5% |
| Azerbaijan_N | TUR_C_Çatalhöyük_N | 0.311 | 62.1% | 37.9% | 3.0% |
| Azerbaijan_N | TUR_C_Musular_PPN | 0.393 | 58.3% | 41.7% | 3.4% |
| Azerbaijan_N | TUR_Marmara_Barcın_N | 0.056 | 67.5% | 32.5% | 2.7% |
| Maikop | TUR_C_Çatalhöyük_N | 0.067 | 44.6% | 55.4% | 1.9% |

**Table S 33 Feasible models for TUR_C_ChL**

Next we look at Bronze Age Central Anatolia (Table S 34). A different mix is inferred here, of an eastern farmer population (the Pre-Pottery Neolithic from Çayönü[56]) but with a small contribution of several populations from the steppe.

The only model that does not involve a steppe contribution has Masis Blur + Ganj Dareh ancestry, however note that this model is evaluated with a Right set that does not include Ganj Dareh (which is moved to the Left set of sources) whereas the models with steppe ancestry have Ganj Dareh on the Right and are thus resilient to ancestry from Iran. When we add a steppe source (BPgroup) as a 3$^{rd}$ source to the Masis Blur + Ganj Dareh model, the estimated proportion of steppe ancestry is 9.0±3.7% (p=0.89), similar to the (simpler) 2-way models with steppe ancestry.

| A | B | P-value | A | B | S.E |
|---|---|---|---|---|---|
| Armenia_C | TUR_SE_Çayönü_PPN | 0.104 | 33.5% | 66.5% | 4.8% |
| Armenia_MasisBlur_N | Iran_GanjDareh_N | 0.302 | 94.3% | 5.7% | 3.3% |
| BPgroup | TUR_SE_Çayönü_PPN | 0.137 | 10.8% | 89.2% | 1.7% |

| | | | | | |
|---|---|---|---|---|---|
| CoreYamnaya | TUR_SE_Çayönü_PPN | 0.101 | 12.2% | 87.8% | 2.0% |
| PVgroup | TUR_SE_Çayönü_PPN | 0.115 | 11.6% | 88.4% | 2.0% |
| Remontnoye | TUR_SE_Çayönü_PPN | 0.190 | 16.3% | 83.7% | 2.4% |
| SShi | TUR_SE_Çayönü_PPN | 0.076 | 11.0% | 89.0% | 1.8% |

**Table S 34 Feasible models for TUR_C_BA**

| | BPgroup+TUR_SE_Çayönü_PPN | CoreYamnaya+TUR_SE_Çayönü_PPN | PVgroup+TUR_SE_Çayönü_PPN | Remontnoye+TUR_SE_Çayönü_PPN | SShi+TUR_SE_Çayönü_PPN | Armenia_C+TUR_SE_Çayönü_PPN | Armenia_MasisBlur_N+Iran_GanjDareh_N |
|---|---|---|---|---|---|---|---|
| BPgroup+TUR_SE_Çayönü_PPN | | 0.119 | 0.068 | 0.171 | 0.099 | 0.100 | 0.114 |
| CoreYamnaya+TUR_SE_Çayönü_PPN | 0.101 | | 0.090 | 0.130 | 0.080 | 0.066 | 0.042 |
| PVgroup+TUR_SE_Çayönü_PPN | 0.159 | 0.147 | | 0.250 | 0.115 | 0.128 | 0.096 |
| Remontnoye+TUR_SE_Çayönü_PPN | 0.169 | 0.119 | 0.142 | | 0.073 | 0.153 | 0.035 |
| SShi+TUR_SE_Çayönü_PPN | 0.172 | 0.137 | 0.148 | 0.249 | | 0.131 | 0.228 |
| Armenia_C+TUR_SE_Çayönü_PPN | 0.151 | 0.115 | 0.135 | 0.249 | 0.074 | | 0.461 |
| Armenia_MasisBlur_N+Iran_GanjDareh_N | 0.132 | 0.098 | 0.109 | 0.146 | 0.080 | 0.141 | |

**Table S 35 Model tournament for TUR_C_BA.** Results of (A, B) matches are shown in A=columns and B=rows of the matrix.

The results of the tournament between these models are inconclusive (Table S 35), but the only model that does not involve any steppe ancestry (Masis Blur + Ganj Dareh) loses out to the models that include either Remontnoye or Core Yamnaya ancestry, albeit weakly (p=0.04).

Next, we look at Bronze Age SW Anatolia in the Mediterranean region (Table S 36). We do not have Neolithic samples from this area, but the Neolithic was presumably similar to that of the early farmers of NW and Central Anatolia. In comparison to them, there is eastern ancestry here.[49] Proportions are virtually identical to those from the Chalcolithic of NW Anatolia discussed above. Thus, during the Chalcolithic and Bronze Age the populations of Western Anatolia were quite similar to each other, composed of a blend of eastern migrants with some survival of the pre-Neolithic inhabitants.

| A | B | P-value | A | B | S.E |
|---|---|---|---|---|---|
| Armenia_Aknashen_N | Bulgaria_C | 0.298 | 48.6% | 51.4% | 3.0% |
| Armenia_Aknashen_N | TUR_C_Çatalhöyük_N | 0.185 | 47.5% | 52.5% | 3.2% |

| | | | | | |
|---|---|---|---|---|---|
| Armenia_KuraAraxes_EBA | TUR_C_Çatalhöyük_N | 0.199 | 43.0% | 57.0% | 2.6% |
| Armenia_KuraAraxes_EBA | TUR_Marmara_Barcın_N | 0.256 | 52.8% | 47.2% | 2.3% |
| Armenia_MasisBlur_N | Bulgaria_C | 0.379 | 73.0% | 27.0% | 5.4% |
| Armenia_MasisBlur_N | Trypillia | 0.077 | 77.7% | 22.3% | 5.7% |
| Armenia_MasisBlur_N | TUR_C_Boncuklu_PPN | 0.127 | 78.8% | 21.2% | 5.1% |
| Armenia_MasisBlur_N | TUR_C_Çatalhöyük_N | 0.196 | 76.7% | 23.3% | 6.5% |
| Armenia_MasisBlur_N | TUR_Marmara_Barcın_N | 0.052 | 82.2% | 17.8% | 6.1% |
| Azerbaijan_C | Bulgaria_C | 0.074 | 56.3% | 43.7% | 3.8% |
| Azerbaijan_C | TUR_C_AşıklıHöyük_PPN | 0.064 | 58.6% | 41.4% | 5.8% |
| Azerbaijan_C | TUR_C_Boncuklu_PPN | 0.534 | 60.8% | 39.2% | 3.3% |
| Azerbaijan_C | TUR_C_Çatalhöyük_N | 0.313 | 53.1% | 46.9% | 4.0% |
| Azerbaijan_C | TUR_C_Musular_PPN | 0.189 | 50.2% | 49.8% | 4.4% |
| Azerbaijan_C | TUR_Marmara_Barcın_N | 0.359 | 61.2% | 38.8% | 3.7% |
| Azerbaijan_N | TUR_C_AşıklıHöyük_PPN | 0.068 | 63.6% | 36.4% | 5.4% |
| Azerbaijan_N | TUR_C_Boncuklu_PPN | 0.099 | 67.1% | 32.9% | 3.2% |
| Azerbaijan_N | TUR_C_Çatalhöyük_N | 0.109 | 58.0% | 42.0% | 3.7% |
| Maikop | TUR_C_Çatalhöyük_N | 0.454 | 42.3% | 57.7% | 2.5% |
| Maikop | TUR_C_Musular_PPN | 0.066 | 38.9% | 61.1% | 3.0% |
| Maikop | TUR_Marmara_Barcın_N | 0.466 | 50.9% | 49.1% | 2.2% |

**Table S 36 Feasible models for TUR_Med_BA**

Next, we looked at the Bronze Age population of the Aegean region from Yassıtepe[4] and Ulucak[57] (Table S 37). This can also be modeled in terms of the Neolithic people of western Anatolia with eastern ancestry.

| A | B | P-value | A | B | S.E |
|---|---|---|---|---|---|
| Armenia_MasisBlur_N | Bulgaria_C | 0.075 | 64.7% | 35.3% | 4.7% |
| Azerbaijan_C | Bulgaria_C | 0.697 | 50.6% | 49.4% | 3.4% |
| Azerbaijan_C | TUR_C_Boncuklu_PPN | 0.484 | 55.7% | 44.3% | 3.2% |
| Azerbaijan_C | TUR_Marmara_Barcın_N | 0.094 | 62.0% | 38.0% | 3.9% |
| Azerbaijan_N | Bulgaria_C | 0.411 | 56.1% | 43.9% | 3.0% |
| Azerbaijan_N | TUR_C_Boncuklu_PPN | 0.053 | 62.1% | 37.9% | 2.9% |

**Table S 37 Feasible models for TUR_Aegean_BA**

#### What was the source of steppe ancestry in Central Anatolian Bronze Age?

The models of Table S 34 include multiple possible steppe sources of the Central Anatolian Bronze Age population and the results of the model tournament were not conclusive (Table S 35). We tried a different approach to better understand the source of the steppe ancestry, by investigating where, along each of the three steppe clines the admixing population could be derived. To this end, we fixed Çayönü as one source and considered the endpoints of the steppe clines as additional sources: BPgroup, Russia_Karelia, Armenia_Aknashen_N, Ukraine_N, GK2, CoreYamnaya. We fit models with 3 sources in total which allowed us to place the steppe component in TUR_C_BA along both the clines inferred by analysis (e.g., Volga=BPgroup+Russia_Karelia), as well as others non-attested with any genetic data (e.g., Aknashen+Ukraine_N) and unlikely on geographical grounds.

The results of this analysis can be seen in Table S 38. We observe (in the order of rows of the table) that:

1. Along the Volga Cline (BPgroup-Karelia) no evidence that the steppe ancestry is from an upriver population, as the proportion of Karelia ancestry is -3.4±2.6%

2. The steppe ancestry could be from the CLV cline (8.8±2.7% BPgroup and 5.3%±8.0% Aknashen) with a significant amount of lower Volga ancestry (Z-score for BPgroup ancestry is 3.3). The standard error of the Aknashen-related ancestry is high, so we can infer that the admixing population definitely had lower Volga ancestry and more likely than not (but not certainly) Aknashen-related ancestry. Nominally, the fraction of BPgroup ancestry of the admixing population was $\frac{8.8}{8.8+5.3} \approx 62\%$ which is similar to the $\frac{56.8}{56.8+20.7} \approx 73\%$ ratio inferred for the Yamnaya (Table S 26). Given the large uncertainty for the fraction of Aknashen-related ancestry in the TUR_C_BA population we cannot be certain that the steppe population admixing into the Dnipro-Don region and that admixing into the ancestors of the Anatolian Bronze Age was from the exact same point in the CLV cline, although both were derived from the CLV cline and had significant amounts of lower Volga (BPgroup-related) ancestry.
3. Models of the form BPgroup+(Ukraine_N or GK2) correspond to the Dnipro cline. In these models the Ukraine_N and GK2 ancestry is nominally negative and non-significant. Thus, there is no compelling evidence for the steppe ancestors of the Central Anatolian Bronze Age having any contribution from the Dnipro-Don area.
4. The model involving Aknashen+Karelia fits (p=0.062) but is unlikely geographically as it involves a hypothetical mixture of Caucasus Neolithic with EHG and there are no known populations of such intermediate ancestry.
5. The model involving Core Yamnaya+Karelia is also feasible according to our criteria but with a negative Karelia contribution of -6.0±3.1% which would point to a source with lower hunter-gatherer ancestry than the Yamnaya. It is unlikely on chronological grounds (given the lateness of the Yamnaya)
6. The models involving Aknashen+(Ukraine_N or GK2) are also feasible but also geographically unlikely for the same reasons as the Aknashen+Karelia one.
7. The models involving Core Yamnaya and Aknashen or Ukraine_N or GK2 ancestry are also rejected on chronological grounds but point to the steppe population being "less Dnipro-Don" and "more Aknashen" than the Yamnaya. Given that the Yamnaya were formed as a mixture of CLV cline people with Dnipro-Don people, these results point indeed to the CLV cline (subtracting the Dnipro-Don component and adding extra Aknashen-related ancestry.

| | | | Proportions | | | Std. errors | | |
|---|---|---|---|---|---|---|---|---|
| **B** | **C** | **P-value** | **Çayönü** | **B** | **C** | **Çayönü** | **B** | **C** |
| BPgroup | Russia_Karelia | 0.149 | 88.8% | 14.6% | -3.4% | 2.0% | 4.0% | 2.6% |
| **BPgroup** | **Armenia_Aknashen_N** | **0.129** | **85.9%** | **8.8%** | **5.3%** | **6.1%** | **2.7%** | **8.0%** |
| BPgroup | Ukraine_N | 0.105 | 90.0% | 10.6% | -0.6% | 1.7% | 3.0% | 2.3% |
| BPgroup | GK2 | 0.155 | 89.8% | 11.6% | -1.4% | 1.8% | 3.4% | 2.6% |
| Russia_Karelia | Armenia_Aknashen_N | 0.062 | 78.8% | 4.0% | 17.2% | 5.3% | 1.2% | 5.6% |
| Russia_Karelia | CoreYamnaya | 0.183 | 85.1% | -6.0% | 20.9% | 2.8% | 3.1% | 5.5% |
| Armenia_Aknashen_N | Ukraine_N | 0.121 | 76.2% | 19.0% | 4.8% | 4.6% | 4.8% | 1.3% |
| Armenia_Aknashen_N | GK2 | 0.181 | 77.2% | 17.8% | 5.0% | 5.0% | 5.3% | 1.4% |
| Armenia_Aknashen_N | CoreYamnaya | 0.123 | 82.6% | 8.2% | 9.2% | 5.6% | 7.1% | 2.7% |
| Ukraine_N | CoreYamnaya | 0.084 | 88.2% | -2.3% | 14.1% | 2.2% | 2.7% | 4.1% |
| GK2 | CoreYamnaya | 0.143 | 87.4% | -3.9% | 16.5% | 2.4% | 3.5% | 5.1% |

**Table S 38 Origin of the steppe ancestry in Central Anatolian Bronze Age.** Feasible models shown and the most plausible model is highlighted.

In conclusion, the models of Table S 38 show no evidence of the steppe population admixing into the Central Anatolian Bronze Age being on either the Volga or Dnipro clines but are consistent with it being on the CLV cline and having lower Volga (BPgroup-related) ancestry.

#### How did steppe ancestry reach Central Anatolia during the Bronze Age?

We were curious about the fact that steppe ancestry is combined with Mesopotamian ancestry from Çayönü[9] when modeling the Central Anatolian Bronze Age, so we considered an alternative Mesopotamian source from Boncuklu Tarla in Mardin[3]. However, the BPgroup+TUR_SE_Mardin_PPN model does not fit for the TUR_C_BA population (p=1e-10), as it overestimates shared genetic drift with Ganj Dareh (Z=3.6) and underestimates it with TUR_Marmara_Barcın_N (Z=-2.5), suggesting that Mardin is more to the "east" along the Anatolian-Iranian or "East Mediterranean-Inland"[3] genetic gradient. We estimate that indeed the Çayönü PPN individuals, sampled 200km west of the Mardin individual (and thus geographically closer to Central Anatolia) are a mixture of 49.2±4.2% Mardin and 50.8±4.2% TUR_C_Çatalhöyük_N ancestry (p=0.373), and thus intermediate between the Neolithic population of Central Anatolia and that further east on the Tigris at Mardin in north Mesopotamia.

Where exactly the admixture took place is uncertain, but we may plausibly place it in Southeastern Anatolia from which it would then reach Central Anatolia. We wanted to identify proximate sources of the "Mesopotamian" ancestry, and so we analyzed separately Chalcolithic and Bronze Age people from eastern regions of Anatolia (Southeast, East, and Black Sea) as well as Chalcolithic people from central Anatolia. First, we examined if any such populations were a clade with the Çayönü population (Table S 39).

| Test | P-value |
|---|---|
| TUR_SE_Titriş_Höyük_EBA | 7.31E-02 |
| TUR_C_Büyükkaya_ChL | 5.91E-02 |
| TUR_SE_Gaziantep_BA | 4.05E-02 |
| TUR_C_ÇamlıbelTarlası_ChL | 5.64E-04 |
| TUR_SE_Şırnak_ChL_B | 1.21E-05 |
| TUR_BlackSea_Amasya_EBA | 2.51E-06 |
| TUR_BlackSea_İkiztepe_ChL | 2.18E-06 |
| TUR_SE_Kilis_EBA_A | 9.23E-09 |
| TUR_E_Arslantepe_ChL | 1.93E-09 |
| TUR_SE_Batman_ChL | 4.78E-10 |
| TUR_E_Arslantepe_EBA | 7.90E-11 |
| TUR_SE_Şırnak_ChL_C | 3.24E-11 |
| TUR_SE_Kilis_MBA | 3.49E-14 |
| TUR_Hatay_Alalakh_MLBA | 1.21E-16 |
| TUR_Hatay_TellKurdu_ChL | 3.51E-24 |
| TUR_SE_Şırnak_BA | 2.45E-28 |
| TUR_SE_Şırnak_ChL_A | 7.49E-61 |

**Table S 39 Testing Central and eastern Anatolian Chalcolithic/Bronze Age sub-populations for being a clade with TUR_SE_Çayönü_PPN**

Some populations in both Southeastern Anatolia and Central Anatolia are consistent with being a clade with TUR_SE_Çayönü_PPN, and thus it is plausible that the Pre-Pottery Neolithic population represented by Çayönü may have existed until a Chalcolithic and Bronze age time frame so that it could have combined with steppe ancestry to form the Central Anatolian Bronze Age population. Our second analysis is to include all subset populations of Table S 39, together with all steppe populations of Table S 34 to see if feasible models exist for TUR_C_BA. None of the inferred models are feasible, but they agree in deriving TUR_C_BA from a mixture of a steppe source with a Çayönü-like population, like Bronze Age Gaziantep or Early Bronze Age Titriş Höyük (Table S 40).

| A | B | P-value | A | B | S.E. |
|---|---|---|---|---|---|
| TUR_SE_Gaziantep_BA | TUR_SE_Şırnak_ChL_A | 0.023 | 117.8% | -17.8% | 8.9% |
| TUR_SE_Gaziantep_BA | BPgroup | 0.020 | 93.1% | 6.9% | 2.9% |
| TUR_SE_Gaziantep_BA | CoreYamnaya | 0.022 | 91.7% | 8.3% | 3.4% |
| TUR_SE_Gaziantep_BA | PVgroup | 0.020 | 92.6% | 7.4% | 3.1% |
| TUR_SE_Gaziantep_BA | Remontnoye | 0.014 | 90.8% | 9.2% | 4.5% |
| TUR_SE_Gaziantep_BA | SShi | 0.024 | 92.3% | 7.7% | 3.2% |
| TUR_SE_Titriş_Höyük_EBA | BPgroup | 0.011 | 95.7% | 4.3% | 4.2% |
| TUR_SE_Titriş_Höyük_EBA | CoreYamnaya | 0.012 | 94.1% | 5.9% | 4.8% |
| TUR_SE_Titriş_Höyük_EBA | PVgroup | 0.007 | 96.2% | 3.8% | 4.5% |
| TUR_SE_Titriş_Höyük_EBA | Remontnoye | 0.006 | 96.1% | 3.9% | 6.6% |
| TUR_SE_Titriş_Höyük_EBA | SShi | 0.023 | 93.7% | 6.3% | 4.1% |

**Table S 40 Modeling Central Anatolian Bronze Age with proximate Chalcolithic/Bronze Age sources.** We show feasible models with p>0.001 as no feasible models with p>0.05 exist.

Next, we investigated how the different proximate Chalcolithic/Bronze Age sources differ from each other in terms of their ancestry, reasoning that since Çayönü itself is halfway along the Çatalhöyük-Mardin PPN gradient, the Chalcolithic/Bronze Age sources might represent different points along this gradient as well. We do indeed observe that most of these populations can be well-modeled with these two sources (Table S 41). We observe that the two genetically closest populations of Çayönü along this gradient are from Chalcolithic Central Anatolia at Büyükkaya and Çamlıbel Tarlası.[21]

| Test | P-value | TUR_SE_Mardin_PPN | TUR_C_Çatalhöyük_N | S.E. |
|---|---|---|---|---|
| TUR_Hatay_TellKurdu_ChL | 6.16E-07 | 11.7% | 88.3% | 4.5% |
| TUR_C_Büyükkaya_ChL | 3.46E-01 | 37.2% | 62.8% | 5.4% |
| TUR_SE_Çayönü_PPN | 3.73E-01 | 49.2% | 50.8% | 4.2% |
| TUR_C_ÇamlıbelTarlası_ChL | 1.50E-01 | 55.9% | 44.1% | 3.8% |
| TUR_BlackSea_Amasya_EBA | 3.71E-01 | 57.2% | 42.8% | 3.9% |
| TUR_SE_Gaziantep_BA | 5.10E-01 | 65.0% | 35.0% | 6.0% |
| TUR_BlackSea_İkiztepe_ChL | 6.54E-02 | 65.1% | 34.9% | 4.3% |
| TUR_E_Arslantepe_ChL | 2.38E-01 | 66.4% | 33.6% | 3.8% |
| TUR_SE_Titriş_Höyük_EBA | 6.73E-01 | 67.3% | 32.7% | 7.7% |
| TUR_SE_Kilis_EBA_A | 3.56E-01 | 70.7% | 29.3% | 4.5% |
| TUR_SE_Şırnak_ChL_B | 6.01E-03 | 72.3% | 27.7% | 7.1% |
| TUR_E_Arslantepe_EBA | 1.55E-01 | 72.6% | 27.4% | 4.7% |
| TUR_Hatay_Alalakh_MLBA | 2.87E-01 | 73.9% | 26.1% | 4.3% |
| TUR_SE_Kilis_MBA | 2.67E-01 | 75.4% | 24.6% | 4.6% |
| TUR_SE_Batman_ChL | 9.88E-01 | 84.6% | 15.4% | 6.0% |
| TUR_SE_Şırnak_ChL_C | 3.33E-01 | 86.9% | 13.1% | 6.7% |
| TUR_SE_Şırnak_BA | 5.61E-01 | 90.2% | 9.8% | 5.4% |
| TUR_SE_Şırnak_ChL_A | 4.00E-01 | 134.1% | -34.1% | 8.4% |

**Table S 41 Chalcolithic/Bronze Age populations on Çatalhöyük-Mardin PPN gradient**

In conclusion it is possible that the Çayönü-related ancestry in TUR_C_BA could reflect populations on the path from southeastern Anatolia to central Anatolia and future studies may clarify if the admixture occurred in the east of the Hittite area followed by migration of the admixed population, or in Central Anatolia itself.

TUR_C_BA that includes Hittite era samples[20] is the only population for which there is evidence of steppe ancestry. We further analyzed the Central Anatolian Bronze Age samples by period (Table S 42), following their assignment in ref[20] into Early Bronze Age (3000-2500BCE), Assyrian Colony (~2000-1750 BCE), and Old Hittite periods (~1750-1200 BCE). This shows that the inferred steppe ancestry, either via a population in which it had been diluted (such as Chalcolithic Armenia) or from populations of the steppe itself, is significant and present in all three subsets of this population.

| Subset | A | B | P-value | A | B | S.E. | Z-score Steppe |
|---|---|---|---|---|---|---|---|
| Turkey_AssyrianColonyPeriod | Armenia_C | TUR_SE_Çayönü_PPN | 0.876 | 31.9% | 68.1% | 6.4% | 5.0 |
| Turkey_EBA_II | Armenia_C | TUR_SE_Çayönü_PPN | 0.084 | 28.5% | 71.5% | 6.4% | 4.5 |
| Turkey_OldHittitePeriod | Armenia_C | TUR_SE_Çayönü_PPN | 0.204 | 38.5% | 61.5% | 7.3% | 5.3 |
| Turkey_AssyrianColonyPeriod | Armenia_MasisBlur_N | Iran_GanjDareh_N | 0.535 | 98.6% | 1.4% | 4.2% | |
| Turkey_EBA_II | Armenia_MasisBlur_N | Iran_GanjDareh_N | 0.812 | 92.5% | 7.5% | 3.7% | |
| Turkey_OldHittitePeriod | Armenia_MasisBlur_N | Iran_GanjDareh_N | 0.117 | 94.0% | 6.0% | 3.9% | |
| Turkey_AssyrianColonyPeriod | BPgroup | TUR_SE_Çayönü_PPN | 0.736 | 9.6% | 90.4% | 2.3% | 4.2 |
| Turkey_EBA_II | BPgroup | TUR_SE_Çayönü_PPN | 0.079 | 8.8% | 91.2% | 2.0% | 4.4 |
| Turkey_OldHittitePeriod | BPgroup | TUR_SE_Çayönü_PPN | 0.217 | 12.1% | 87.9% | 2.3% | 5.3 |
| Turkey_AssyrianColonyPeriod | CoreYamnaya | TUR_SE_Çayönü_PPN | 0.598 | 10.7% | 89.3% | 2.6% | 4.1 |
| Turkey_EBA_II | CoreYamnaya | TUR_SE_Çayönü_PPN | 0.055 | 9.6% | 90.4% | 2.3% | 4.2 |
| Turkey_OldHittitePeriod | CoreYamnaya | TUR_SE_Çayönü_PPN | 0.178 | 13.8% | 86.2% | 2.6% | 5.3 |
| Turkey_AssyrianColonyPeriod | PVgroup | TUR_SE_Çayönü_PPN | 0.764 | 10.5% | 89.5% | 2.4% | 4.4 |
| Turkey_EBA_II | PVgroup | TUR_SE_Çayönü_PPN | 0.058 | 9.4% | 90.6% | 2.3% | 4.1 |
| Turkey_OldHittitePeriod | PVgroup | TUR_SE_Çayönü_PPN | 0.154 | 13.0% | 87.0% | 2.5% | 5.2 |
| Turkey_AssyrianColonyPeriod | Remontnoye | TUR_SE_Çayönü_PPN | 0.847 | 14.8% | 85.2% | 3.3% | 4.5 |
| Turkey_EBA_II | Remontnoye | TUR_SE_Çayönü_PPN | 0.130 | 14.0% | 86.0% | 3.0% | 4.7 |
| Turkey_OldHittitePeriod | Remontnoye | TUR_SE_Çayönü_PPN | 0.180 | 18.0% | 82.0% | 3.4% | 5.3 |
| Turkey_AssyrianColonyPeriod | SShi | TUR_SE_Çayönü_PPN | 0.487 | 9.4% | 90.6% | 2.4% | 3.9 |
| Turkey_EBA_II | SShi | TUR_SE_Çayönü_PPN | *0.039* | 8.7% | 91.3% | 2.2% | 4.0 |
| Turkey_OldHittitePeriod | SShi | TUR_SE_Çayönü_PPN | 0.122 | 12.6% | 87.4% | 2.4% | 5.3 |

**Table S 42 Models of Table S 34 for subsets of Bronze Age Central Anatolia**

The Maikop culture was previously suggested to be the archaeological culture of either the Proto-Anatolians or Proto-Indo-Anatolians as a whole.[58] The model in which the steppe ancestry is mediated by Maikop rather than Armenia_C fails weakly for TUR_C_BA with p=0.022 and predicts 23.2±3.8% Maikop ancestry. However, that does not mean that Maikop is not involved in the ancestry of TUR_C_BA, since, as we have seen it is a source for Caucasus-Lower Volga populations like Remontnoye (Table S 6) which could be the source for both TUR_C_BA (Table S 34) and Armenia_C (Table S 11). The Maikop (archaeological culture) was an Early Bronze Age phenomenon and the sampled Maikop individuals span the 4th millennium BCE, with the Remontnoye individuals proving that the steppe-Caucasus mixtures had started by the late 5th-early 4th millennium BCE. This is also the time frame of the Areni-1 Chalcolithic population in Armenia.[5] This would allow for, perhaps, 1,000 years of linguistic differentiation to occur between the "high-steppe" groups derived from Remontnoye-like speakers of Proto-Indo-Anatolian and the "low-steppe" groups from which the Maikop/Areni-1 groups would emerge, and coincide well with inferred dates for Proto-Indo-Anatolian of 4000BCE[59], 4200BCE[60], 4139–3450 BCE[53], and 4300-4200BCE.[51] With all due caution as to its meaning, we also note the curious coincidence of the dates of Proto-Indo-Anatolian with the ~4000BCE date of admixture in the ancestry of the Yamnaya (Fig. S 5).

#### Unified models of Anatolian and Caucasus populations

The fact that TUR_C_BA can be modeled quite distinctly from the other central/western Anatolian populations raises the question whether the ancestry we detect there could also be discerned in other Chalcolithic/Bronze Age Anatolians that can be, strictly speaking, modeled without any such ancestry. As the 2-way model for TUR_C_BA does not fit the other Chalcolithic and Bronze Age populations, we identified 3-way models that (i) include one of the steppe sources, (ii) fit TUR_C_BA, and (iii) fit as many populations of the TestA set as possible.

Using this procedure, we identify four models (that differ in terms of the steppe source) that fit a total of 9 populations. The two other sources of these models are (i) Masis Blur (which we have seen represents a population that is the substratum of Armenian Chalcolithic (Table S 11) and of the Eastern Turkey Chalcolithic and Bronze Age (Table S 31), and (ii) Zagros Neolithic from Ganj Dareh[5] which makes a positive contribution to populations from Azerbaijan whose Iranian Neolithic affinity was previously noted.[3]

These models agree that Armenia_C is distinctive from the populations of Anatolia having a much greater steppe affinity and systematically show that TUR_C_BA also has some such affinity. But, what is most notable is that the populations of Eastern Turkey and Azerbaijan cluster closely together and with Masis Blur. So, the conclusion is that, overall, this is a region in which the old Neolithic ancestry persists with only slight variations either towards Iran (as in the case of Azerbaijan) or the steppe (as is the case for TUR_C_BA). For the Indo-Anatolian question, the data is consistent with either Hypothesis B (given the preponderance of the local Neolithic ancestry) or with Hypothesis A (given the evidence for different levels of diluted ancestry in Armenia_C and TUR_C_BA).

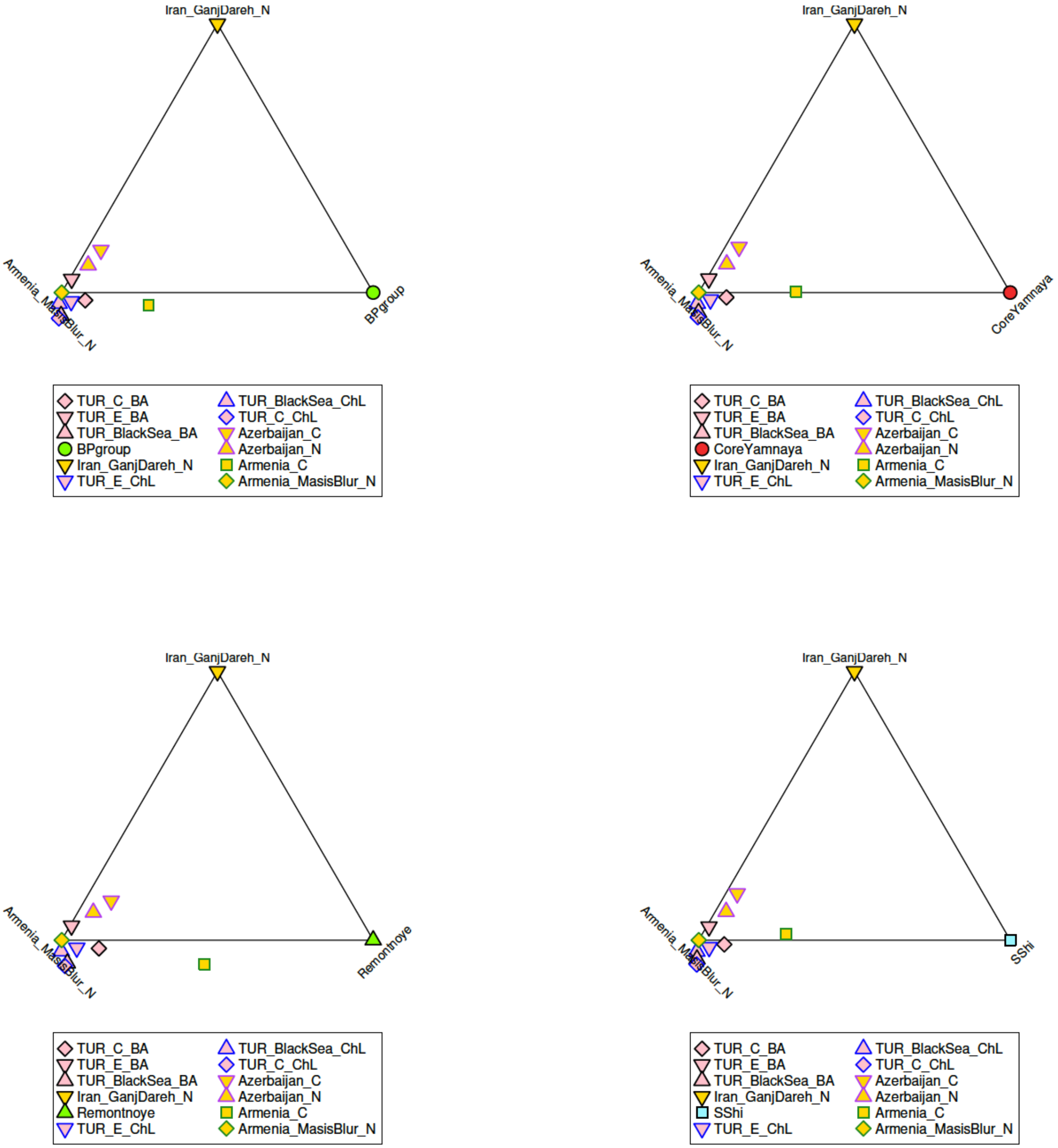

**Fig. S 15 Unified 3-way models for Anatolia and the Caucasus**

The models of Fig. S 15 have Masis Blur as the West Asian source, while those of Table S 34 have Çayönü PPN as the source. It was previously observed that Masis Blur could be modeled as a mixture of the earlier Neolithic from Aknashen with Anatolian-Levantine ancestry presumably from Mesopotamia.[3,4] We can indeed model Masis Blur as 33.9±8.6% Aknashen and 66.1±8.6% Çayönü PPN (p=0.466). Given that the steppe ancestry also included Aknashen ancestry via the Caucasus-Lower Volga populations, it is thus possible to fit models in which the West Asian ancestry is Çayönü (in which the Aknashen ancestry could be derived from the north, from the Caucasus-Lower Volga admixed populations) or in which it is Masis Blur (in which the Aknashen ancestry is derived from the south, from Mesopotamian-South Caucasus admixed populations), or indeed a combination of both. Regardless of the history, the two classes of models agree in the derivation of part of the ancestry of TUR_C_BA from West Asia with a smaller contribution from the steppe.

#### A Western route via Southeastern Europe into Anatolia?

We now consider the possibility of a western route of steppe people into Anatolia (Hypothesis "A-West"). The strongest arguments for this hypothesis is the western distribution of Anatolian languages within Anatolia[51] and the archaeological and genetic evidence for pre-Yamnaya expansions from the steppe into southeastern Europe.[11,60,61] The main counter-arguments have been (i) the lack of steppe ancestry in Anatolia[4,20,21,49,50], and (ii) the fact that the trajectory of ancestry change in Chalcolithic/Bronze Age Anatolia was strongly in the direction of an increase of "eastern" (Caucasus-Mesopotamian) ancestry[4], whereas migrations from the Balkans would have introduced a mixture of Anatolian Neolithic-WHG ancestry.

In this section we explore the western hypothesis directly, by making use of the data recently published by Penske et al.[61] as well as data from our study and Anatolian populations to see if we can model the Central Anatolian Bronze Age population as well with a western source as we did with a combination of an eastern Mesopotamian and a steppe source (Table S 34). We use the following set of sources (we use the population names of ref.[61] and include the totality of their populations, that includes pre-steppe migration, early steppe migrants, and Yamnaya-related migrants for the sake of completeness):

**SourcesW**: BOY_EBA, KTL_A, KTL_B, MAJ, MAJ_EBA, PIE039, PIE060, PIE078, PIE_CA, PTK_CA, TUR_C_AşıklıHöyük_PPN, TUR_C_Boncuklu_PPN, TUR_C_Çatalhöyük_N, TUR_C_Musular_PPN, TUR_Marmara_Barcın_N, TUR_SE_Çayönü_PPN, Usatove, USV, VAR_CA, YUN041, YUN_CA, YUN_EBA

| A | B | P-value | A | B | S.E. |
|---|---|---|---|---|---|
| BOY_EBA | TUR_SE_Çayönü_PPN | 8.89E-02 | 12.5% | 87.5% | 2.2% |
| MAJ_EBA | TUR_SE_Çayönü_PPN | 8.17E-02 | 12.4% | 87.6% | 2.2% |
| Usatove | TUR_SE_Çayönü_PPN | 5.08E-02 | 22.2% | 77.8% | 3.9% |
| USV | TUR_SE_Çayönü_PPN | 3.92E-02 | 20.7% | 79.3% | 3.6% |
| KTL_A | TUR_SE_Çayönü_PPN | 3.89E-02 | 16.6% | 83.4% | 3.0% |
| MAJ | TUR_SE_Çayönü_PPN | 1.92E-02 | 20.9% | 79.1% | 3.7% |
| YUN041 | TUR_SE_Çayönü_PPN | 1.13E-02 | 20.4% | 79.6% | 4.2% |
| KTL_B | TUR_SE_Çayönü_PPN | 3.66E-06 | 13.1% | 86.9% | 9.5% |
| PIE060 | TUR_SE_Çayönü_PPN | 1.78E-06 | 3.5% | 96.5% | 3.5% |
| YUN_EBA | TUR_SE_Çayönü_PPN | 1.36E-06 | -5.5% | 105.5% | 6.9% |
| PIE078 | TUR_SE_Çayönü_PPN | 1.18E-06 | -1.9% | 101.9% | 7.0% |

**Table S 43 Modeling Central Anatolian Bronze Age with western sources**

In Table S 43 we show feasible models as well as non-feasible ones p>1e-06. What all these models have in common is that they too (like the models of Table S 34) involve the pairing of Mesopotamian ancestry with steppe-admixed sources. The top two sources (BOY_EBA and MAJ_EBA) are Yamnaya-related and thus correspond to the model with Core Yamnaya already examined (Table S 34). These models point to our inability to precisely determine the steppe-related source of ancestry of the Anatolian Bronze Age but seem implausible on chronological grounds as the Bronze Age individuals from Central Anatolia date to as early as the Early Bronze Age (archaeologically dated to 2750-2500BCE), soon after the Yamnaya expansion, yet their Yamnaya ancestry is greatly diluted. Moreover, if Yamnaya-related ancestors brought steppe ancestry into Anatolia then this would make the linguistic inference that Anatolian languages are a sister group to Indo-European languages problematic, and predict that the whole of the Indo-Anatolian language family has a common ancestor ~5kya, which is outside of mainstream opinion.

Pre-Yamnaya populations which appear as sources in Table S 34 include Usatove/USV and Kartal cluster A (KTL_A) and Mayaki (MAJ) that either narrowly pass or miss the p=0.05 threshold. However, these models appear contrived and implausible as they predict the almost choreographed arrival of people from the Balkans and Mesopotamia, bypassing from both directions the people that lived on the path to their Central Anatolian destination, and their admixture there to form the Central Anatolian Bronze Age.

An eastern route into Anatolia seems more plausible as steppe ancestors admixed with an Aknashen-related population to form the ancestors of the Maikop (Table S 8) in the North Caucasus, and with Masis Blur-related ancestors to form the ancestors of the Armenian Chalcolithic (Table S 11), and finally with Mesopotamian-related ancestors to form the ancestors of the Central Anatolian Bronze Age (Table S 34), i.e., moving from the steppe via the North Caucasus, to the South Caucasus, to Mesopotamia and admixing with the people that lived there along the entire Caucasus-Mesopotamia genetic cline.

### A reconstruction of the origin and dispersal of Proto-Indo-Anatolian and Proto-Indo-European languages

We present below (Table S 44) a unified scenario of Indo-Anatolian and Indo-European origins that seems consistent with the genetic transformations we observe.

| Period | Mainland Europe | Don-Dnipro | Volga | Caucasus | Anatolia |
|---|---|---|---|---|---|
| >6000 years BCE | WHG with some EHG ancestry in Scandinavia, the Baltics, and Southeastern Europe.[11,39,62-67] | Ukrainian hunter-gatherers: within WHG-EHG cline[11] | Largely EHG | CHG[19] (South)<br>? (North) | “Anatolian pottery Neolithic” in Central-Western Anatolia[3,6,7,55,68] descended from Epipaleolithic Anatolians[7] but with Mesopotamian[3,56]/Levantine ancestry.[3] |
| | **Transformations by movements of people from Anatolia and the Caucasus** | | | | |
| 6000-4000 years BCE | Early European farmers descended from Neolithic Anatolians across the continent.[6,28,62,64,69,70] | Gene flow from the Volga at Golubaya Krinitsa (GK1) and thence to the Dnipro (Igren_o / GK2);<br><br>Formation of the Serednii Stih Cline: a mixture of “Pre-Yamnaya”+Ukraine_N/GK2<br><br>Haplogroup J2 man on the Lower Don at Krivyansky: Serednii Stih + CHG ancestry.<br><br>Pre-Yamnaya are formed by 4000BCE: Remontnoye + Serednii Stih sources (Proto-Indo-European speakers) | Admixture of Caucasus (CHG) and Central Asian (TTK) ancestry;<br><br>Formation of the Lower Volga+North Caucasus Eneolithic (BPgroup+PVgroup) (Proto-Indo-Anatolian speakers? Hypothesis A)<br><br>Formation of the Volga cline: BPgroup+EHG ancestry | Anatolian-Levantine ancestry reaches the Caucasus with the Neolithic expansion: Aknashen (less), Masis Blur (more)[4];<br><br>Pre-Maikop in the North Caucasus: Majority Aknashen with some BPgroup ancestry (Proto-Indo-Anatolian speakers? Hypothesis B)<br><br>Armenia Chalcolithic[4]: Majority Masis Blur + some steppe<br><br>Remontnoye: Volga cline + Maikop/Aknashen | Eastern migrants from Caucasus / Eastern Anatolia into central/western Anatolia;<br>Drastic reduction of “Anatolian Neolithic ancestry”[4];<br><br>(Proto-Anatolians from the east?) |
| 4000-3000 years BCE | | Yamnaya Y-chromosome R-Z2103/R-M12149 patrilineal clan emerges out of the Pre-Yamnaya: location unknown;<br><br>Horse domestication[71], wheeled vehicles, interaction with Maikop, | Continuation of Volga cline populations until their replacement by the Yamnaya | Maikop culture emerges in the NW Caucasus, interaction with Pre-Yamnaya and with Steppe Maikop | |

| | | | | | |
|---|---|---|---|---|---|
| | | (Likely Proto-Indo-European speakers) | | At the end of this period: Kura Araxes expansion[4,8] | |
| | **The era of steppe migrations** | | | | |
| 3000-2000 years BCE | Steppe-derived cultures: Corded Ware[1,2,46] and Bell Beakers[41] in mainland Europe. Admixture with European farmers.<br>Yamnaya in Southeastern Europe beyond the steppe.[4,11] | The Yamnaya-Afanasievo dominate the Eurasian steppe from Mongolia to the Pannonian Basin.<br><br>Successor cultures: Poltavka and Catacomb on the steppe.<br><br>The end of the Yamnaya and their replacement by the back-migration of Fatyanovo[48] / Sintashta[72] culture Corded Ware R-M17 patrilineal clans.[1,2,4,22] | | Kura Araxes dominance until Middle Bronze Age;<br><br>Mid-3rd millennium BCE: Yamnaya descendants re-introduce steppe ancestry to Armenia[4] | |
| | **The height of the Bronze Age** | | | | |
| 2000-1000BCE | Mycenaean Greece[21,49,52]: ~10% Yamnaya ancestry. ~30% in Southeastern Europe.[4]<br>Corded Ware/Bell Beaker-Farmer descendants across Europe. | Sintashta-Srubnaya-Andronovo (Corded Ware descendants) have replaced the Yamnaya on the steppe.[6,22,73] | | Middle-Late Bronze Age Armenians: an enclave of ~1/4 Yamnaya ancestry[4] and ~30% Yamnaya Y-chromosomes in West Asia | Diluted CLV ancestry Bronze Age Hittite-era Central Anatolia[20] |

**Table S 44 A unified model of Indo-Anatolian and Indo-European origins**

#### Summary of Findings, Conclusions, and Future directions

In this note we first modeled the ancestry of the core Yamnaya and pre-Yamnaya Eneolithic populations of the Pontic-Caspian steppe. We find a proximate 2-source model for the core Yamnaya involving ancestry from the North Caucasus piedmont (represented by Remontnoye) and the SShi subset of the Serednii Stih culture of the Don-Dnipro region. The Yamnaya are more distally derived from hunter-gatherers of eastern Europe (both of Ukraine, via their Serednii Stih ancestry, and of Russia, via their Remontnoye ancestry), and from populations of the Caucasus that admixed with them to form the Serednii-Stih cline in the Don-Dnipro region and the Volga cline that reached the North Caucasus piedmont.

The Yamnaya also had extra ancestry related to the North Caucasus Maikop culture and more generally the Neolithic of Armenia; people of this ancestry admixed with people of the Lower Volga (Berezhnovka)-North Caucasus piedmont (Progress-2) to form intermediate populations like Remontnoye. The migration of people like those of Remontnoye into the Serednii Stiih territory created the precursors of the Yamnaya which then emerged as a homogeneous population at one extreme of Serednii Stih culture variation before their late 4th millennium BCE expansion across Eurasia.

We explored alternative hypotheses about the origins of Indo-Anatolian languages and we summarize the relative findings below:

- Hypothesis A: a steppe origin of Indo-Anatolian languages, places the homeland in the North Caucasus-Lower Volga region and identifies the Proto-Indo-Anatolians as people of steppe ancestry. Expansions out of this region to the Caucasus and thence to Anatolia may have greatly diluted the steppe component (Hypothesis A-East) and given rise to Anatolian languages. The Maikop culture of the North Caucasus could be identified as the harbinger of Proto-Indo-Anatolian languages (which would eventually become Proto-Anatolian) to the south. Expansions via southeastern Europe and thence to Anatolia (Hypothesis A-West) would also require a great dilution of the steppe component as steppe ancestry is not seen in western Anatolia and the Anatolian Neolithic/SE European Neolithic ancestry diminishes in Chalcolithic and Bronze Age Anatolians. However, we do in fact see dilution of this type, so this scenario is more plausible.
- Hypothesis B: a Caucasus-West Asian origin of Indo-Anatolian origins is strengthened by the finding of early migrations from the Caucasus into the Volga/Don-Dnipro Eneolithic populations followed by later Maikop/Armenian Neolithic ancestry into the ancestors of the Yamnaya. This

hypothesis also maps to the transformation of Chalcolithic and Bronze Age central/western Anatolia which saw half to all its Neolithic population replaced.

- Hypothesis C: the Indo-Anatolian languages were spoken by people of diverse genetic background in the North Caucasus-steppe interaction zone and spread from there by largely disjoint ancestors of the Yamnaya in the north and Caucasus-West Asian populations in the south.

In the future it is important to study the Pontic-Caspian steppe in even finer spatio-temporal detail to identify the pre-Yamnaya population in the Eneolithic mix of Don-Volga with Serednii Stih populations out of which we think that the Yamnaya emerged. Where did the "core Yamnaya" patrilineal clan (R-Z2103/R-M12149 bearers) live and why did they become so successful? Did the Corded Ware descend from the core Yamnaya directly (via a shift in Y-chromosomes) or from a late pre-Yamnaya group that was genetically similar but did not belong to the "core Yamnaya" clan? It is also crucial to study interactions between populations of the North Caucasus such as those of the Maikop and preceding cultures with the inhabitants of the steppe as we think that here was the setting of the earliest history of Indo-Anatolian languages. At the same time, it is necessary to continue work in Anatolia, in order to better understand the proximate origins of the Chalcolithic and Bronze Age newcomers in central/western Anatolia and their more distal sources in eastern Anatolia, Armenia, and the Caucasus.
